# A telomere-to-telomere genome of Holstein cattle resolves chromosome-end architecture and the genetic basis of telomere length

**DOI:** 10.64898/2026.10.10.758172

**Authors:** Fangyuan Liu, Tao Wang, Pengbo Cao, Wei Wang, Fatima, Zhuangbiao Zhang, Fanxin Meng, Pengfei Lin, Zhenyu Wei, Jie Liu, Rihan Wu, Lixia Wang, Yunxia Li, Gaoping Zhao, Xiangnan Bao, Wei Sun, Yu Jiang, Yu Wang, Xihe Li

**Author notes:** Correspondence (W.S.), (Y.J.), (Y.W.), (X.L.). These authors contributed equally (Fangyuan Liu, Tao Wang, Pengbo Cao, Wei Wang, Fatima).

## Abstract

Telomeres safeguard chromosome-end integrity, yet population-scale dissection of telomere-length genetics has been largely restricted to humans. Here, we report a 3.01-Gb telomere-to-telomere (T2T) genome of Holstein cattle that resolves repeat-rich chromosome ends and centromeres and enables telomere-length quantification at single-chromosome-end resolution. Holstein chromosome ends show marked telomere-length asymmetry and end-specific variation in telomere variant repeats, subtelomeric tandem repeats, GC/CpG composition, and DNA methylation. Using sequencing-depth-corrected telomere-length phenotypes from 9,591 cows, teloGWAS identifies major loci encompassing *RTEL1*, *CTC1*, and *TERT*, while multi-tissue eQTL, sQTL, TWAS, and co-expression analyses prioritize *CTC1* and implicate DNA replication, DNA repair, and stress-response networks. Together, these findings extend population-scale telomere genetics beyond humans and establish a sequence-resolved framework for chromosome-end biology and telomere-informed livestock genomics.

**In brief:** Liu et al. generate a telomere-to-telomere Holstein cattle genome and resolve chromosome-end repeat architecture. Integrating end-resolved telomere annotation with population-scale GWAS and transcriptomic analyses, they identify conserved telomere-maintenance loci and nominate *CTC1* as a candidate effector gene for bovine telomere length variation.

**Graphical abstract:** 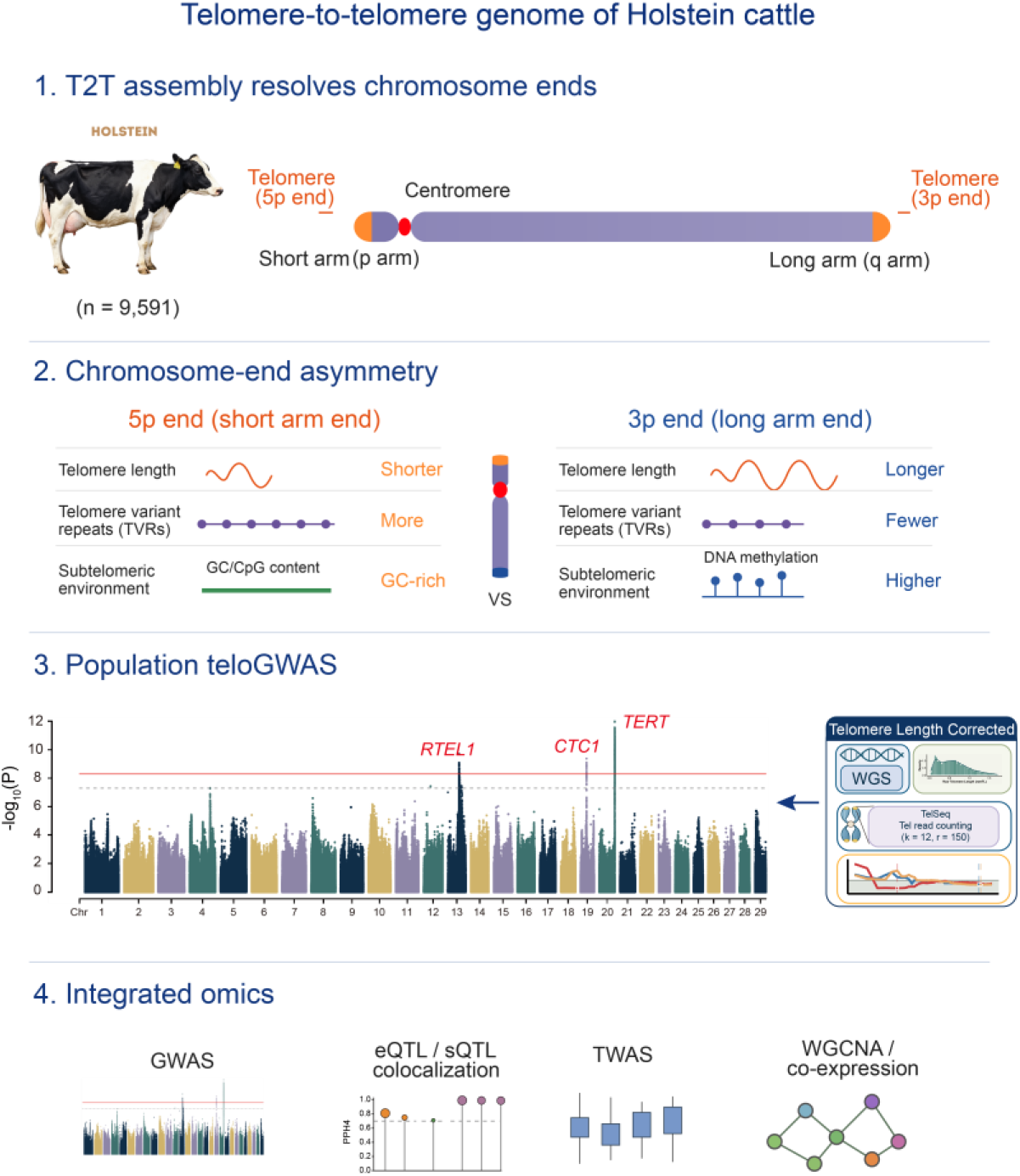

**Highlights:**

Holstein_T2T resolves bovine chromosome termini and centromeres
Acrocentric chromosome architecture is linked to telomere length asymmetry
TVR and subtelomeric GC/CpG landscapes distinguish chromosome ends
The teloGWAS identifies regulatory loci near RTEL1, CTC1 and TERT

## Introduction

Telomeres cap the ends of linear chromosomes and prevent chromosome termini from being recognized as DNA double-strand breaks. In vertebrates, this protective function is mediated by tandem arrays of TTAGGG repeats together with associated shelterin proteins, which safeguard chromosome-end integrity^1,2^. As a hallmark of aging, telomere attrition is intimately linked to cellular senescence, genome instability and organismal aging ^3,4^. Work at single-telomere resolution has further shown that, in telomerase-negative cells, genome instability and replicative senescence are triggered by the shortest telomeres rather than by the average telomere length of a cell ^5^, underscoring the biological significance of telomere heterogeneity among individual chromosome ends. In livestock, such heterogeneity may also influence fertility, health, stress resilience, and productive lifespan^6^. The repeat-rich nature of chromosome ends has long prevented their complete assembly in livestock reference genomes; as a consequence, telomere length has usually been reduced to a single genome-wide, sample-averaged phenotype, leaving population-scale variation at individual chromosome termini, its genetic basis, and its relationship to complex traits largely unresolved.

Telomere-to-telomere (T2T) assembly brings chromosome termini, subtelomeres, centromeres, and large satellite repeat arrays into sequence-resolved view, closing gaps that remained in previous reference genomes and enabling direct comparative analysis of regions that had been inaccessible to conventional assemblies^7,8^. Complete assemblies from humans and other vertebrates have shown that these regions are far from uniform structural entities: centromeric arrays vary among haplotypes and individuals in length, sequence composition, and higher-order repeat (HOR) organization, whereas subtelomeric regions are frequently enriched for structural variation and repeat-mediated diversification^7,9,10^. This layer of variation remains largely invisible to analyses anchored to conventional references, which primarily capture SNPs, short indels, and copy-number variants. In domestic animals, population history, introgression, and artificial selection have jointly shaped genomic diversity among breeds ^11-13^, but the extent to which this diversity extends into telomeric, subtelomeric, and centromeric repeat domains is still poorly understood. Bovine autosomes are predominantly acrocentric, with the centromere positioned near one chromosome end, creating pronounced structural polarity between the two chromosome termini and providing a natural system for dissecting chromosome-end organization^14^. Systematic resolution of chromosome-end sequence architecture, including canonical telomeric repeats, telomere variant repeats, and subtelomeric sequences, is therefore essential for understanding telomeric and centromeric variation across species, breeds, and individuals.

Telomeres and subtelomeres form a continuous sequence transition at chromosome ends, making their boundaries difficult to define precisely. Telomere variant repeats (TVRs), which occur within terminal repeat arrays or near telomere–subtelomere junctions, punctuate otherwise continuous tracts of canonical TTAGGG repeats and can alter telomere function^15^. Although these variants provide informative sequence features for anchoring long reads to specific chromosome ends or alleles, they also make accurate telomere-length determination more challenging^15,16^. The adjoining subtelomeres add another layer of heterogeneity: mammalian subtelomeric regions are enriched for segmental duplications, interspersed repeats, CpG-rich sequences, and structural variants, and their composition varies among chromosome arms, individuals, and lineages ^17,18^. Such features can shape local chromatin state, recombination activity, and gene regulation, while also influencing how telomere length is measured and interpreted.

Telomere length has emerged as a tractable quantitative trait for population-scale genetic analysis. Genome-wide association studies (GWAS) have identified numerous telomere-length-associated loci, implicating pathways involved in telomere maintenance, DNA repair, cell-cycle regulation and aging-related processes in humans^19-21^. Integrative analyses that combine association signals with transcriptomic data have further prioritized candidate causal genes and regulatory mechanisms ^20^, while the incorporation of qPCR- and sequencing-derived telomere measurements in large WGS cohorts has increased the power to detect genetic effects on telomere length ^21^. In livestock, studies of telomere-length genetics are only beginning to emerge: bovine leukocyte telomere length has been shown to be moderately heritable, and early GWAS have begun to map loci and candidate genes underlying its variation ^22,23^. However, the reliability of telomere-length GWAS depends critically on how telomere length is defined, measured and anchored to a reference genome. Short-read-based estimators generally infer the average telomeric-repeat content of a sample rather than resolving individual chromosome ends ^24,25^. Even in T2T assemblies, terminal telomeric arrays, TVRs and adjacent subtelomeric sequences can remain compressed, incompletely recovered or misclassified during assembly and annotation ^7,15,16^. Such uncertainties can propagate into population-scale telomere phenotyping, distort association signals and complicate the interpretation of GWAS loci. Accurate assembly and annotation of chromosome termini are therefore essential for building robust telomere phenotypes and for interpreting telomere-length associations at chromosome-end resolution ^15,26,27^.

In this study, we generate a T2T reference genome for Holstein cattle (Holstein_T2T) and use it to systematically resolve complex chromosome-end regions that remain incomplete in conventional cattle references. This assembly enables chromosome-arm-specific characterization of telomere architecture. We further combine telomere length GWAS and TWAS in 9,591 Holstein cattle with large-scale blood and rumen transcriptomic data to identify candidate loci and genes associated with telomere length variation. Together, our study establishes, to our knowledge, the first chromosome-end-resolved framework that couples a T2T reference genome with population-scale telomere-length association analysis, providing a general paradigm for dissecting telomere variation and its genetic basis in livestock.

## Results

### High-contiguity Holstein_T2T resolves complex chromosome regions

To construct a high-quality T2T assembly, we generated PacBio HiFi (∼71×), Oxford Nanopore Technologies (ONT; ∼62×), and Hi-C (∼135×) sequencing data from peripheral blood of a seven-year-old Holstein cow and assembled the genome using an integrated multi-platform strategy (Figure S1 and Supplementary Table 1). The final assembly spans 3.01 Gb, with a contig N50 of 111.23 Mb, a Merqury consensus quality value (QV) of 72.54, and 99.92% BUSCO completeness (Figure 1A and Supplementary Table 2). Chromosome-wide Hi-C contact maps showed clear intrachromosomal interaction patterns without evidence of large-scale interchromosomal misjoins, supporting the structural accuracy of the assembly (Figure S2). Read-mapping analyses further confirmed continuous and accurate sequence representation across the Holstein genome (Figure S3). Compared with the previous cattle reference genome ARS-UCD2.0, the Holstein_T2T assembly resolved complex regions that were missing or incompletely represented, with newly resolved sequences concentrated at chromosome termini, subtelomeres, centromeres, and other repeat-rich domains. Annotation of these newly resolved intervals showed that they were dominated by repetitive sequences. Centromeric sequences represented the largest component, accounting for 71.76% of the newly resolved sequence space (∼243.37 Mb), followed by LINEs (24.87 Mb), LTR retrotransposons (24.86 Mb), satellite repeats (20.80 Mb), SINEs, and other repeat classes (Figures 1B and 1C).

**Figure 1.**
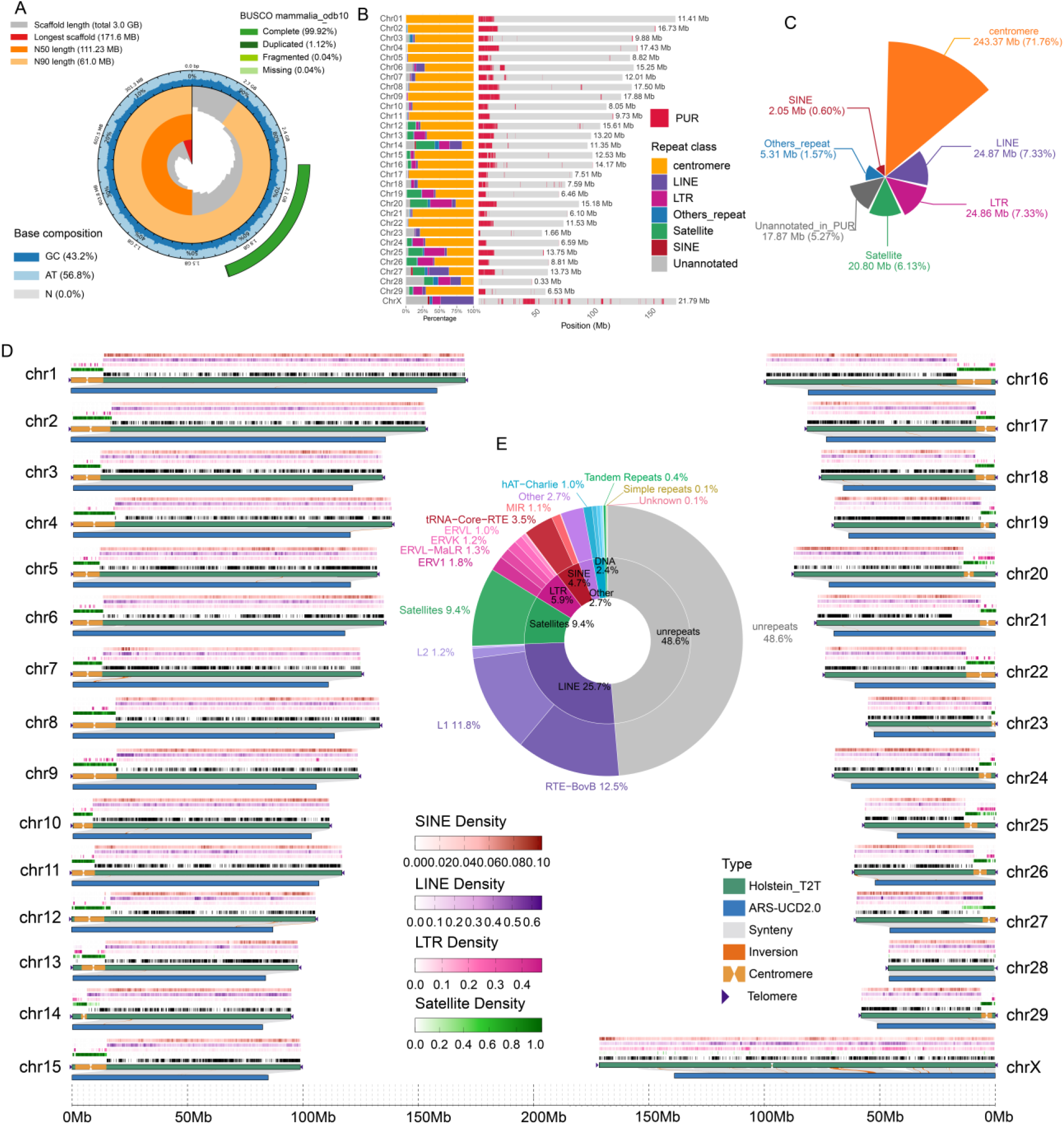
Holstein_T2T assembly and genome-wide repeat landscape. (A) Overview of assembly quality, including genome size, contig N50, Merqury quality value, and BUSCO completeness. (B) Chromosomal distribution of sequences newly resolved in the Holstein_T2T assembly relative to ARS-UCD2.0. (C) Repeat composition of newly resolved sequences, including centromeric sequences, LINEs, LTR retrotransposons, satellite repeats, SINEs, and other repeat classes. (D) Genome-wide distribution of repeat density along chromosomes, highlighting enrichment patterns in centromeric and subtelomeric regions. (E) Composition of repetitive elements across the whole genome, including LINEs and the RTE-BovB and L1 subfamilies, satellite repeats, LTR retrotransposons, and other repeat classes.

Genome-wide annotation identified approximately 1.55 Gb of repetitive sequence, representing 51.36% of the genome. Transposable elements accounted for 41.44%, with LINEs forming the predominant class (25.75%), followed by LTR retrotransposons (5.88%) and SINEs (4.69%). Within the LINE fraction, RTE-BovB and L1 were the two major subfamilies, accounting for 12.46% and 11.83% of the genome, respectively. Satellite repeats contributed a further 9.42% of the genome, whereas tandem repeats and other repeat classes constituted smaller fractions. Repeat-density profiling revealed pronounced domain-specific enrichment in centromeric and subtelomeric regions, highlighting the heterogeneous organization of repetitive sequence across bovine chromosomes (Figures 1D and 1E).

Integrated gene annotation combining transcriptome-based, homology-based, and ab initio predictions identified 21,200 protein-coding genes, with a mean gene length of 41.0 kb, a mean coding-sequence length of 1.60 kb, and an average of 9.07 exons per gene. BUSCO analysis of the predicted gene set recovered 98.28% of mammalian conserved orthologs as complete, supporting the high completeness of the annotation. Functional annotation assigned at least one annotation to 20,358 genes (96.03%) across NR, Swiss-Prot, KEGG, KOG, and GO databases, and 19,841 genes (93.59%) were supported by transcriptomic expression evidence. Non-coding RNA annotation additionally identified rRNAs, small RNAs, regulatory RNAs, and tRNAs, further expanding the functional annotation of the Holstein_T2T. Together, the Holstein_T2T assembly closes major gaps in previous cattle references while providing a highly complete annotation of both repetitive and gene-containing regions. The resulting repeat-resolved and gene-annotated genome provides a sequence framework for subsequent analyses of bovine centromere organization, chromosome-end architecture, and telomere biology.

### Structural Diversity of Acrocentric chromosomes in Cattle Against a Conserved Karyotypic Background

The complete Holstein_T2T assembly enabled fine-scale dissection of centromeric and telomeric organization. Using CENPA CUT&Tag, we annotated centromeric regions across the Holstein genome and confirmed that Holstein autosomes are predominantly acrocentric, with centromeres positioned close to one chromosome end (Figure S4 and Supplementary Table 3). This organization provides a natural basis for examining how centromeres and chromosome termini are structurally related. To evaluate conservation of acrocentric centromere architecture within cattle, homologous centromeric intervals were extracted from the Holstein_T2T and the Wagyu genome UOA_Wagyu_1^28^. Despite strong chromosome-scale collinearity, homologous centromeres showed marked local sequence divergence (Figure S5 and Figure S6). Although broad sequence similarity was retained between corresponding centromeric regions, repeat arrays within homologous intervals frequently displayed expansion, contraction, or fragmentation, reflecting insertions, deletions, and altered repeat organization within otherwise conserved centromeric regions (Figure S7A and S7B). Indel profiling further revealed abundant insertions and deletions across homologous centromeres, with repeat-rich intervals often dominated by either insertion- or deletion-biased pattern (Figure S7C).

Analysis of satellite repeat-unit organization further showed that higher-order repeat units in Holstein and Wagyu centromeres are relatively conserved in sequence composition but differ substantially in copy abundance and higher-order arrangement (Figure S7D–S7F). These findings indicate that considerable centromeric sequence diversity lies beneath the conserved cattle karyotype, with dynamic repeat-array turnover and local structural rearrangement emerging as major contributors to intraspecific variation in bovine centromere architecture. Because this comparison relies on single representative genomes, the full extent of centromere diversity within cattle will require additional individuals and complementary experimental approaches.

### Chromosome-end-specific repetitive sequence architecture is associated with telomere length asymmetry

The acrocentric organization of cattle chromosomes provides a natural model for examining how chromosome-end architecture relates to telomere-length variation. Because long terminal TTAGGG arrays are difficult to quantify accurately from assembled sequences alone, we applied Telogator2 to raw PacBio HiFi and ONT long reads to independently evaluate and calibrate telomeric repeat tracts and refine chromosome-end boundaries^15^. After calibration, 42 chromosome ends reached base-level (±1 bp) agreement with read-supported telomere lengths, whereas larger residual deviations remained only at the chromosome 13 3p end (1,083 bp) and the chromosome 19 5p end (266 bp). This calibrated framework enabled genome-wide telomere-length (TL) analysis at single-chromosome-end resolution (Supplementary Table 4).

Calibrated TL varied substantially among chromosome ends, ranging from 1.08 to 65.76 kb at 5p termini and from 0.94 to 34.77 kb at 3p termini (Figure 2A and Supplementary Table 5). Paired comparison of the two ends of each chromosome showed that 21 of 30 chromosomes carried longer telomeres at the 3p end, with a median absolute inter-end difference of 7.68 kb (Figure 2B). Although this paired comparison did not reach statistical significance (paired Wilcoxon signed-rank test, P = 0.074), the frequency and magnitude of inter-end differences point to strong chromosome-end dependence of telomere length. This asymmetry was accompanied by variation in repeat architecture: TVR profiling showed that non-canonical repeats are unevenly distributed among chromosome ends (Figure 2C), with many short-arm 5p termini displaying broader and stronger enrichment of non-canonical TVRs and other non-canonical repeat sequences (Figure S8A and S8C), whereas long-arm 3p termini were generally dominated by extended canonical-repeat tracts with lower non-canonical content (Figure S8B and S8D). Quantitatively, TVR abundance was higher at 5p than at 3p termini for 22 of 30 chromosomes (paired Wilcoxon test, P = 6.0 × 10−5) and was negatively correlated with mean TL across chromosome ends (Spearman’s ρ = −0.405, P = 0.026; Figure 2D). Stratified analysis showed that this negative association was stronger at 3p termini than at 5p termini, indicating that the relationship between TVR abundance and TL is chromosome-end specific (Figure 2E).

**Figure 2.**
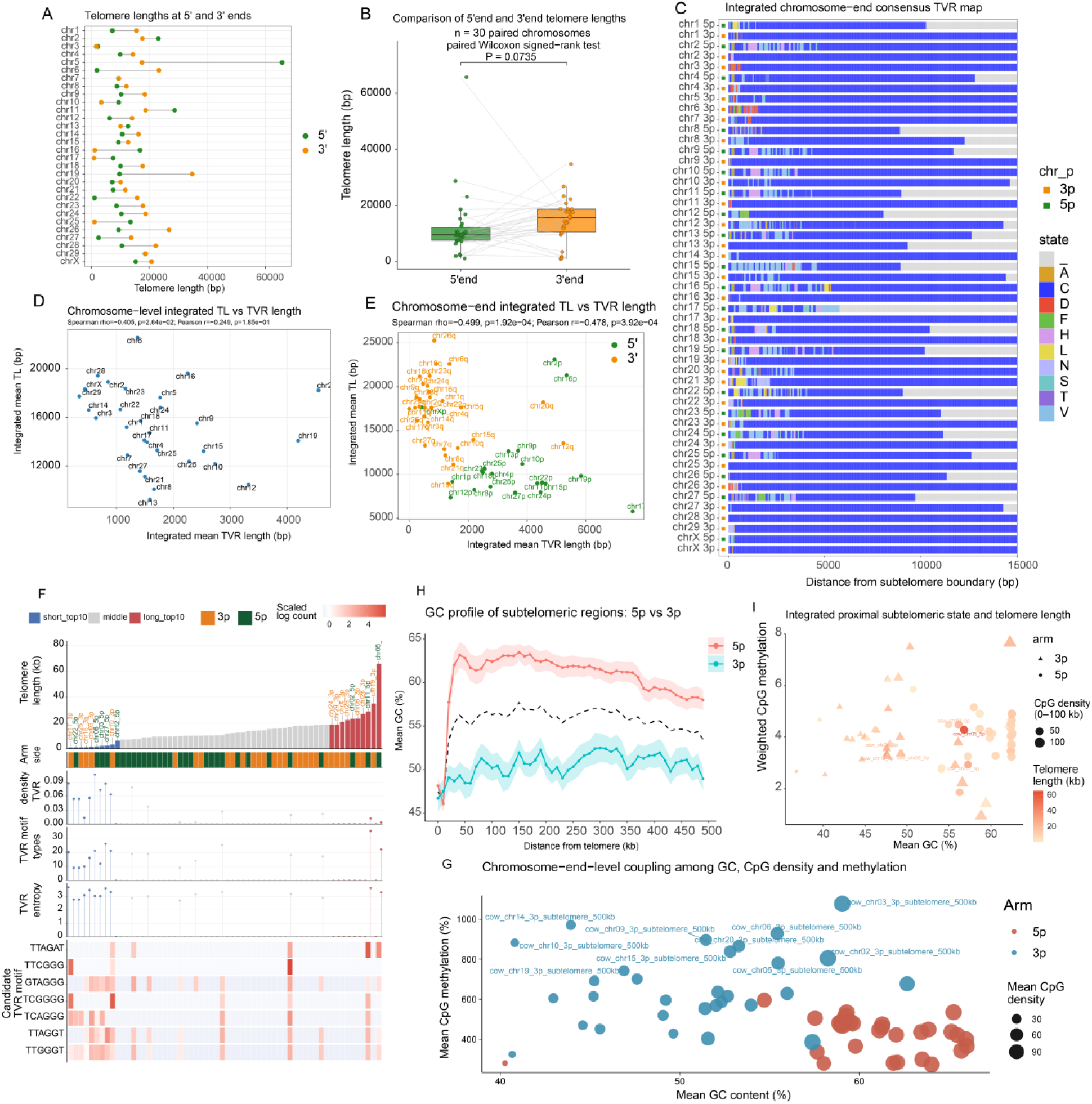
Chromosome-end-resolved telomere length, TVRs, and subtelomeric sequence environment. (A) Distribution of telomere length at 5p and 3p chromosome ends. (B) Paired comparison of telomere length between the two ends of each chromosome. (C) Genome-wide map of telomere variant repeat (TVR) distribution across chromosome ends. (D) Correlation between TVR abundance and mean telomere length across chromosome ends (Spearman’s ρ = −0.405, *P* = 0.026). (E) Chromosome-end-specific comparison of the relationship between TVR abundance and telomere length at 5p and 3p termini. (F) Motif-level divergence of TVRs across the telomere–subtelomere transition zone. (G) Comparison of subtelomeric GC content between 5p and 3p termini (60.65% versus 50.64%, *P* = 1.2 × 10^−8^). (H) Comparison of CpG density between 5p and 3p subtelomeric regions (106.4 versus 56.2 CpG sites per kb, *P* = 4.8 × 10^−7^). (I) Quantitative associations between subtelomeric GC/CpG composition and telomere length.

Motif-level analysis of TVR-containing regions showed that sequence divergence extends beyond terminal TVR tracts into the telomere–subtelomere transition zone and adjacent tandem-repeat regions (Figure 2F). Repeat-unit distributions at 5p and 3p termini indicated that the two chromosome ends differ not only in TVR abundance but also in the abundance and composition of neighboring tandem repeats (Figure S9A and Supplementary Table 5): 25 of the 30 chromosomes harbored a higher proportion of tandem repeats at the 3p end, whereas only five showed 5p-biased enrichment (Figure S9B). Telomere-like motif density was likewise heterogeneous among termini, with marked enrichment of telomere-like motifs in the subtelomeric regions of a subset of chromosome ends (Figure S9C). Correlation analyses suggested that TL variation is associated not only with TVR quantity but also with the compositional complexity of TVR-containing regions. This negative association was driven mainly by 3p termini and attenuated at 5p termini (Figure S9D–S9F). Motif-abundance clustering separated 3p and 5p subtelomeric regions and identified telomere-like and non-telomere-like motifs enriched in a chromosome-arm-specific manner (Figure S9G). Principal-component analysis (PCA) of motif density similarly distinguished the two classes of termini (PERMANOVA, *R*² = 0.178, *P* = 0.001; Figure S9H), while differential motif analysis identified repeat motifs preferentially enriched in either 5p or 3p subtelomeric regions (Figure S9I).

These results indicate that TVR abundance, motif composition, and the organization of the telomere– subtelomere transition zone are associated with TL variation in Holstein cattle. Telomere-length asymmetry therefore reflects a broader chromosome-end-specific repeat architecture shaped by terminal TVR organization, neighboring motif composition, local repeat density, and chromosome-end identity.

### Subtelomeric GC/CpG Composition and DNA Methylation Are Associated with Telomere Length

To assess how sequence composition and epigenetic features contribute to telomere-length variation and chromosome-end asymmetry, we analyzed the 500-kb subtelomeric intervals immediately internal to terminal telomeric arrays. Subtelomeric regions differed markedly between 5p and 3p termini in GC content, CpG density, and DNA methylation (Figures 2G and 2H; Figure S10A–S10E). Mean GC content was significantly higher at 5p than at 3p termini (60.65% versus 50.64%; paired Wilcoxon test, P = 1.99 × 10^−10^; Figure 2G and Figure S10A), and CpG density was likewise elevated at 5p termini (106.4 versus 56.2 CpG sites per kb; P = 4.8 × 10^−7^; Figure 2H). Distance-resolved analysis further showed consistently higher CpG density at 5p than at 3p chromosome ends across the adjacent subtelomeric region (Figure S10B). By contrast, DNA methylation was enriched at 3p termini relative to 5p termini (P = 5.6 × 10^−7^). Chromosome-end-resolved heatmaps further revealed substantial heterogeneity in both CpG density and DNA methylation across the 500-kb subtelomeric intervals, with 5p termini generally displaying higher CpG density and 3p termini showing stronger methylation enrichment (Figures S10C and S10D). This arm-specific methylation difference was also evident in distance-resolved profiles across the subtelomeric region (Figure S10E).

Subtelomeric sequence composition was also quantitatively associated with TL (Figure 2I). Across chromosome ends, GC content was negatively associated with TL in both the 100–250 kb subtelomeric interval and the full 0–500 kb region (P = 4.6 × 10^−4^, FDR = 0.021 and P = 0.0016, FDR = 0.024, respectively). CpG density in the distal 250–500 kb interval was similarly negatively associated with TL (P = 0.0021, FDR = 0.024). Relative to sequence composition, DNA methylation showed a more localized relationship with TL: distance-resolved window analysis showed that methylation at the 5p chromosome-end boundary was positively associated with TL (P = 0.0050), whereas methylation 10 kb upstream of the terminus was negatively associated with TL (P = 0.0078).

GC content was strongly coupled with CpG density across subtelomeric windows, particularly at 3p chromosome ends (Figure S10F). By contrast, the relationship between GC content and DNA methylation was weak, with only modest arm-specific correlations (5p, Spearman’s ρ = 0.08; 3p, Spearman’s ρ = 0.06; Figure S10G). To evaluate the independent contributions of subtelomeric features to TL variation, we fitted multivariable linear models using log10-transformed TL as the response variable and chromosome arm as a covariate. The standardized effects of GC content, CpG density, and methylation on TL were interval dependent: GC content in the 100–250 kb interval and CpG density in the distal 250–500 kb interval showed the strongest negative effects, whereas the independent contribution of DNA methylation was comparatively limited. These sequence and epigenetic analyses indicate that TL variation is associated with chromosome-end-specific subtelomeric environments, with GC/CpG-rich subtelomeric domains generally corresponding to shorter telomeres. In conjunction with TVR analyses, these findings suggest that the distinctive chromosome-end telomere phenotype of Holstein cattle is shaped jointly by terminal repeat architecture and the surrounding subtelomeric genomic environment, providing a structural basis for population-scale analysis of telomere-length variation and its molecular determinants.

### Corrected telomere length estimates enable robust identification of telomere maintenance loci in cattle

Using whole-genome resequencing data from 9,591 Holstein cows, we performed a genome-wide screen for loci associated with telomere length (Supplementary Table 6). Because sequencing-based telomere-length estimates can be confounded by depth, batch effects, and other technical sources of coverage variation, we applied next-generation sequencing principal component analysis (NGS-PCA) to a genome-wide 1-kb-window coverage matrix. The resulting coverage-derived principal components, hereafter referred to as bPCs, were used to model systematic technical variation, and bPC-corrected telomere-length phenotypes were carried forward into GWAS (Figure 3A). Raw TelSeq estimates were strongly right-skewed, whereas bPC-corrected values were approximately normally distributed (Figure S11A and S11B). To cross-validate WGS-based telomere-length estimation, 57 randomly selected samples were additionally analyzed with Computel; TelSeq- and Computel-derived estimates were highly correlated (r = 0.965, P < 0.001; Figure S11C).

**Figure 3.**
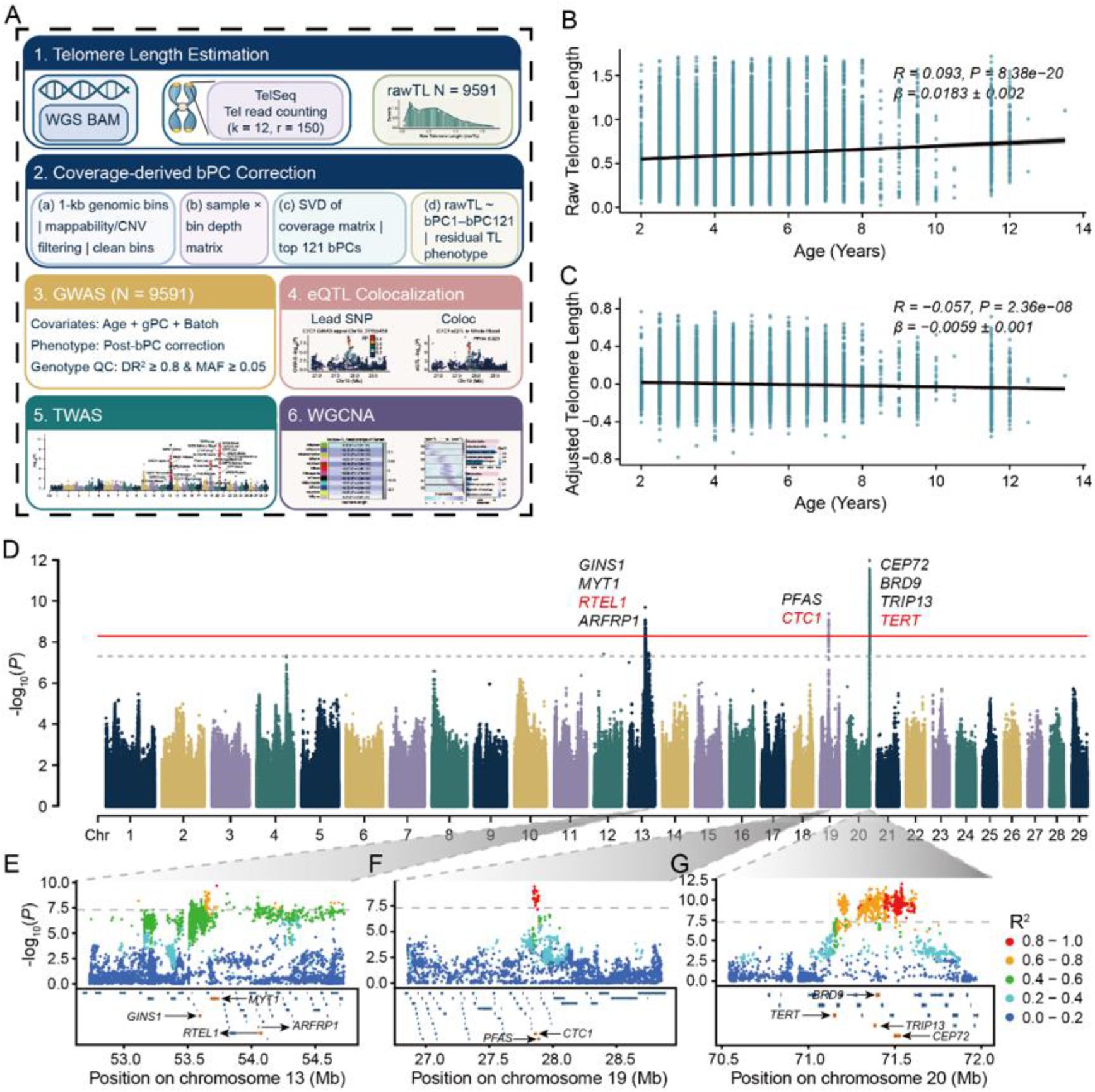
Genome-wide association analysis of telomere length (teloGWAS). (A) Workflow for correction of the telomere length phenotype based on genome-wide sequencing coverage profiles. (B) Relationship between raw telomere length estimates and age before correction (*r* = 0.093, *P* = 8.38 × 10^−20^). (C) Relationship between corrected telomere length estimates and age after correction (*r* = −0.049, *P* = 2.36 × 10^−8^). (D) Manhattan plot of teloGWAS showing major association signals on chromosomes 13, 19, and 20. The red solid line indicates the Bonferroni-corrected genome-wide significance threshold (*P* = 5.037 × 10^−9^), and the gray dashed line indicates the reference threshold (*P* = 5 × 10^−8^). Representative candidate genes within associated regions are annotated, with *RTEL1*, *CTC1*, and *TERT* highlighted in red. (E) Regional association plot for the *RTEL1* locus on chromosome 13 (approximately 53.0–54.5 Mb). (F) Regional association plot for the *CTC1* locus on chromosome 19 (approximately 27.0–28.5 Mb). (G) Regional association plot for the *TERT* locus on chromosome 20 (approximately 70.5–72.0 Mb). Colors indicate the degree of linkage disequilibrium (LD; *R*^2^) between each SNP and the corresponding lead SNP.

Raw telomere length showed a weak negative correlation with genome-wide mean sequencing depth (r = −0.059, P = 8.25 × 10−9), whereas this association was largely eliminated after bPC correction (r = −0.008, P = 0.458; Figures S11D and S11E). Raw and corrected estimates remained moderately correlated (r = 0.523, P < 0.001; Figure S11F), indicating that the correction reduced technical noise while preserving a substantial component of inter-individual variation. Batch-associated differences that were apparent in raw telomere length were also markedly reduced after correction, supporting the effectiveness of bPC adjustment in mitigating systematic variation related to sequencing procedure and batch structure (Figure S11G and S11H).

To evaluate the influence of low sequencing depth on TelSeq estimates, we performed downsampling analysis in ten samples across a range of depths. Telomere-length estimates stabilized rapidly as depth increased, and most samples showed reduced depth sensitivity once coverage reached approximately 1× (Figure S11I). These quality-control and sensitivity analyses indicate that low-depth WGS data can capture relative inter-individual telomere-length variation, provided that technical variation associated with depth and batch is appropriately corrected. The 9,591 samples were then analyzed as a full cohort, a high-depth subgroup (≥4.5×; n = 1,498), and a low-depth subgroup (<4.5×; n = 8,093), using 121, 5, and 131 bPCs for correction, respectively (Supplementary Table 7). Age served as a biological benchmark for evaluating correction performance. Before correction, raw telomere length showed a weak but highly significant positive correlation with age, potentially reflecting residual technical bias or population structure (r = 0.093, P = 8.38 × 10−20; Figure 3B). After bPC correction, the association shifted to a negative correlation (r = −0.049, P = 2.36 × 10−8; Figure 3C), consistent with expected telomere attrition; similar improvements were observed in both high- and low-depth subgroups (Figure S11J).

We next performed two-step LDAK-KVIK GWAS using the bPC-corrected TL phenotype, with age, sequencing batch, and the first 20 genotype principal components included as covariates. This analysis identified strong association signals on chromosomes 13, 19, and 20 (Figure 3D and Supplementary Table 8). On chromosome 13, the association interval (∼53.0–54.5 Mb) harbored *RTEL1*, a helicase involved in telomeric DNA replication (Figure 3E); on chromosome 19, the peak at ∼27.0–28.5 Mb encompassed *CTC1*, which encodes a core component of the CST telomere-protection complex (Figure 3F); and the strongest signal, on chromosome 20 (∼70.5– 72.0 Mb), covered several genes, including *TERT*, which encodes the catalytic subunit of telomerase (Figure 3G). The convergence of telomere-replication, chromosome-end-protection, and telomerase genes within these loci indicates that telomere-length variation in cattle is governed by evolutionarily conserved telomere-maintenance pathways.

To assess the sensitivity of these signals to sequencing depth, we stratified the cohort by genome-wide mean depth into low-depth (<4.5×; n = 8,093) and high-depth (≥4.5×; n = 1,498) subgroups and repeated GWAS separately in each. As expected from the difference in sample size, statistical power differed substantially between the two analyses: the major association regions were largely retained in the larger low-depth subgroup, whereas association significance was attenuated in the smaller high-depth subgroup (Figure S12A and S12B). Conditional GWAS was then performed with the three lead variants, chr13_53727328, chr19_27855458, and chr20_71539969, included as covariates (Supplementary Table 9). After conditioning on the corresponding lead SNP, the original association peak at each locus was markedly attenuated, and no residual signal reached the predefined significance threshold (Figure S12C–S12E). The maximum residual −log10(P) values at the three conditioned loci decreased to 2.9, 1.7, and 1.7, respectively, while the other two unconditioned loci retained strong signals in each analysis (Figure S12F), indicating that each major locus is driven predominantly by a single detectable association signal rather than by multiple independent lead variants. Finally, tissue-expression profiling showed that *RTEL1* and *CTC1* are broadly expressed across cattle tissues, whereas *TERT* displays lower overall expression and greater tissue specificity (Figure S12G).

### Integrated eQTL, sQTL and TWAS analyses identify candidate effector genes for telomere length variation

Having mapped the major loci, we integrated multi-tissue expression quantitative trait locus (eQTL), splicing QTL (sQTL), and transcriptome-wide association (TWAS) data to prioritize candidate effector genes within them. At the chromosome 19 locus, the lead variant chr19_27855458 defined a pronounced association peak adjacent to *CTC1* (Figure 4A). The *CTC1* eQTL signal colocalized with the telomere-length GWAS signal, quantified as the posterior probability of a shared causal variant (PPH4), in whole blood (PPH4 = 0.923), liver (0.985), testis (0.952), muscle (0.965), and adipose tissue (0.981) (Figure 4B–4F and Supplementary Table 10), consistent with a shared putative causal variant underlying both association signals. Additional tissues reinforced *CTC1* as the candidate effector gene at the chromosome 19 locus. Colocalization between the *CTC1* eQTL and telomere-length GWAS signals was also observed in jejunum (PPH4 = 0.786), rumen (PPH4 = 0.980), visceral fat (PPH4 = 0.939), and subcutaneous adipose tissue (PPH4 = 0.831) (Figure S13A–S13D). The lead variant itself tracked with both phenotypes: bPC-corrected telomere length differed significantly among chr19_27855458 genotype classes (Kruskal–Wallis test, *P* = 8.75 × 10−12; Figure S13E), and genotype-stratified analysis of CattleGTEx individuals revealed significant genotype-dependent *CTC1* expression in testis, liver, adipose tissue, muscle, subcutaneous adipose tissue, rumen, and jejunum, with weaker effects in whole blood and visceral fat (Figure S13F).

**Figure 4.**
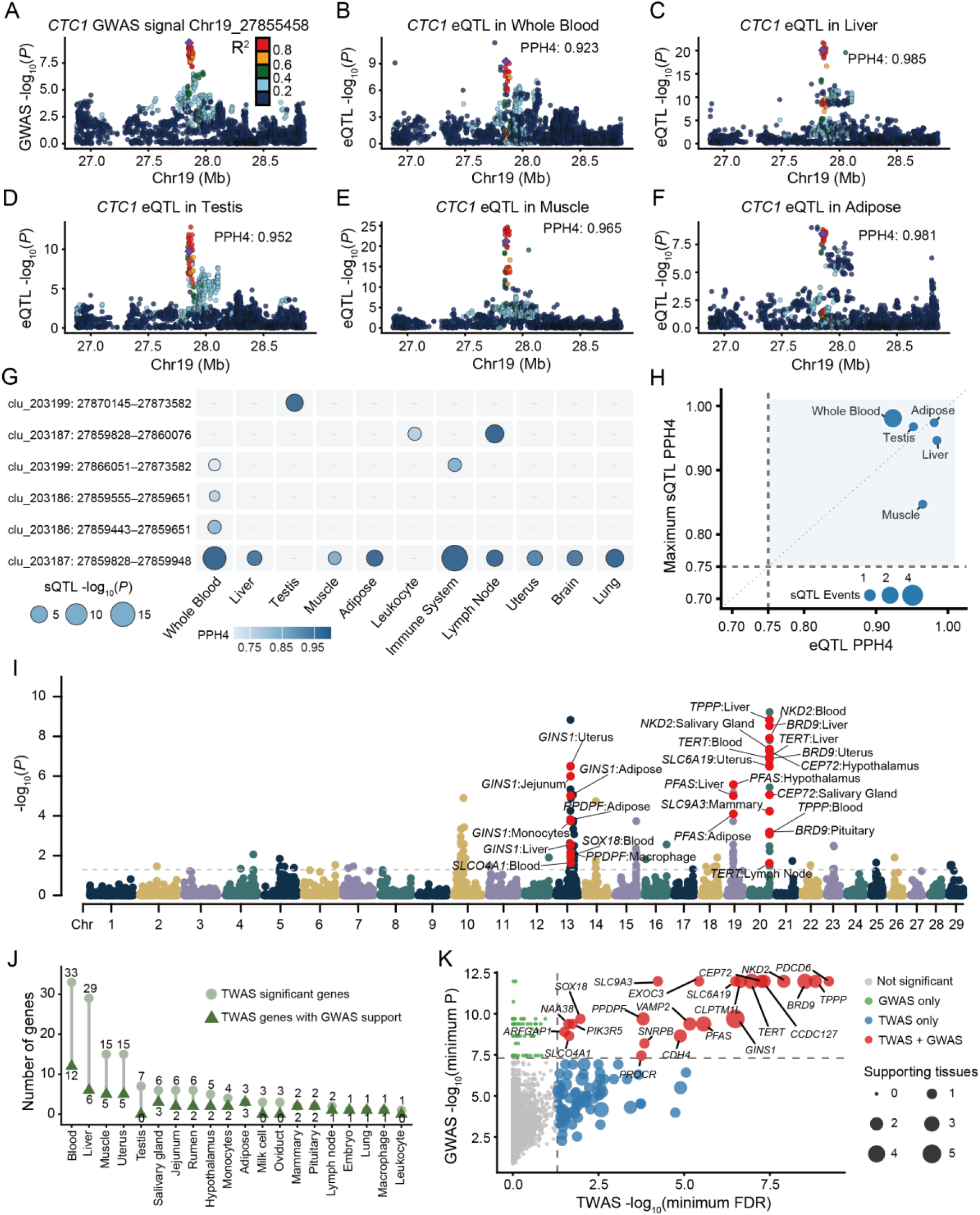
Multi-omic regulatory evidence prioritizes *CTC1* and candidate effector genes for telomere length variation. (A) Regional association signal for telomere length at the chromosome 19 *CTC1* locus. (B–F) Colocalization of *CTC1* eQTL signals with teloGWAS signals in whole blood, liver, testis, muscle, and adipose tissue. PPH4 values are shown for each tissue. (G) Tissue-resolved colocalization between *CTC1*-associated sQTL events and the teloGWAS signal. Rows represent *CTC1* splicing events and columns represent tissues; point color indicates GWAS–sQTL colocalization probability (PPH4), and point size denotes the association strength of the lead SNP for each sQTL event. (H) Cross-tissue comparison of *CTC1* eQTL and sQTL colocalization evidence. The x axis shows the PPH4 for *CTC1* eQTL– GWAS colocalization, the y axis shows the maximum PPH4 among *CTC1* sQTL events in each tissue, and point size indicates the number of sQTL events analyzed. (I) Genome-wide distribution of multi-tissue telomere length TWAS signals. Each point represents a gene–tissue association, with significant TWAS associations highlighted in red (FDR < 0.05). The gray dashed line denotes the FDR = 0.05 threshold. Candidate genes supported by nearby GWAS signals are labeled as gene:tissue pairs. (J) Number of TWAS-significant genes across tissues and the subset supported by nearby GWAS signals. Light-green circles indicate genes with TWAS FDR < 0.05, whereas dark-green triangles indicate genes with at least one GWAS variant reaching *P* < 5 × 10^−8^ within ±500 kb. (K) Gene-level integration of multi-tissue TWAS and local GWAS evidence. The x axis shows −log10 of the minimum TWAS FDR for each gene across available tissues, and the y axis shows −log10 of the minimum GWAS P value within ±500 kb of the gene. Point color indicates the category of genetic support, and point size indicates the number of supporting tissues.

The chromosome 19 locus also showed evidence of splicing-level regulation. Multiple *CTC1*-associated splicing events colocalized with the GWAS signal across tissues (Figure 4G and Supplementary Table 11), and in whole blood, adipose tissue, testis, liver, and muscle, both the *CTC1* eQTL PPH4 and the maximum sQTL PPH4 exceeded 0.75 (Figure 4H). Expression- and splicing-level regulation at *CTC1* therefore converge on the same association region, strengthening the case for *CTC1* as the leading candidate effector gene at this locus.

To search for additional effector genes genome-wide, we performed multi-tissue TWAS. Significant gene-tissue associations with telomere length (FDR < 0.05) were detected across multiple tissues, most frequently in whole blood and liver (Figure 4I, 4J and Supplementary Table 12). Intersecting TWAS results with local GWAS evidence, defined as TWAS FDR < 0.05 in at least one tissue and at least one GWAS variant with P < 5 × 10−8 within ±500 kb of the gene, yielded a smaller set of jointly supported genes that was again enriched in whole blood and liver (Figure 4J and Supplementary Table 13). These signals mapped mainly to the three major GWAS regions, including *GINS1* and neighboring genes on chromosome 13, *PFAS* on chromosome 19, and *TERT*, *BRD9*, *CEP72*, *NKD2*, and *TPPP* on chromosome 20 (Figure 4I; Figure S14A).

A gene-level integration combining each gene’s minimum TWAS FDR across tissues with the minimum GWAS P value within ±500 kb confirmed that jointly supported candidates were concentrated in the three major association regions (Figure 4K). Several candidates were TWAS significant in multiple tissues, suggesting that the same loci may influence telomere length through tissue-dependent expression effects. For three selected candidates, *BRD9*, *GINS1*, and *PFAS*, eQTL–GWAS colocalization returned high PPH4 in specific tissues (Figure S14B and Supplementary Table 14), and each displayed a distinct tissue-expression pattern (Figure S14C), consistent with expression regulation that shares putative causal variation with the GWAS signals.

Collectively, the multi-tissue regulatory evidence nominates *CTC1* as the leading candidate effector gene at the chromosome 19 locus, while supporting *GINS1*, *PFAS*, *TERT*, *BRD9*, and *CEP72* at other major regions. Holstein telomere-length variation therefore maps near conserved telomere-maintenance genes and may act, at least in part, through modulation of gene expression or transcript structure.

### Tissue-specific co-expression networks underlie telomere length variation in Holstein cattle

To examine how telomere length relates to tissue-specific transcriptional states, we analyzed 317 rumen and 289 whole-blood samples with matched TL and transcriptome measurements at three levels: single-gene expression associations, co-expression modules, and TL-stratified network connectivity. Principal-component and inter-sample correlation analyses revealed no outlier samples or unexpected clustering in either tissue (Figure S15A and S15B). We first evaluated telomere-maintenance genes catalogued**^20^** and found that several were significantly associated with TL, with some genes showing opposite directions of association in rumen and whole blood, indicating tissue-dependent regulation of telomere-related transcription (Figure S15C and Supplementary Table 15). Among GWAS candidates, *SIRPA*, *ABHD16B*, *GINS1*, *DIDO1*, *SOX18*, *TCEA2*, *ARFRP1*, *PRPF6*, *ZGPAT*, and *SLCO4A1* correlated positively with TL in whole blood, whereas *GMEB2*, *PRPF6*, and *C13H20orf204* correlated negatively in rumen (Figure 5A; Figure S15D and Supplementary Table 16), with *PRPF6* changing direction between the two tissues.

**Figure 5.**
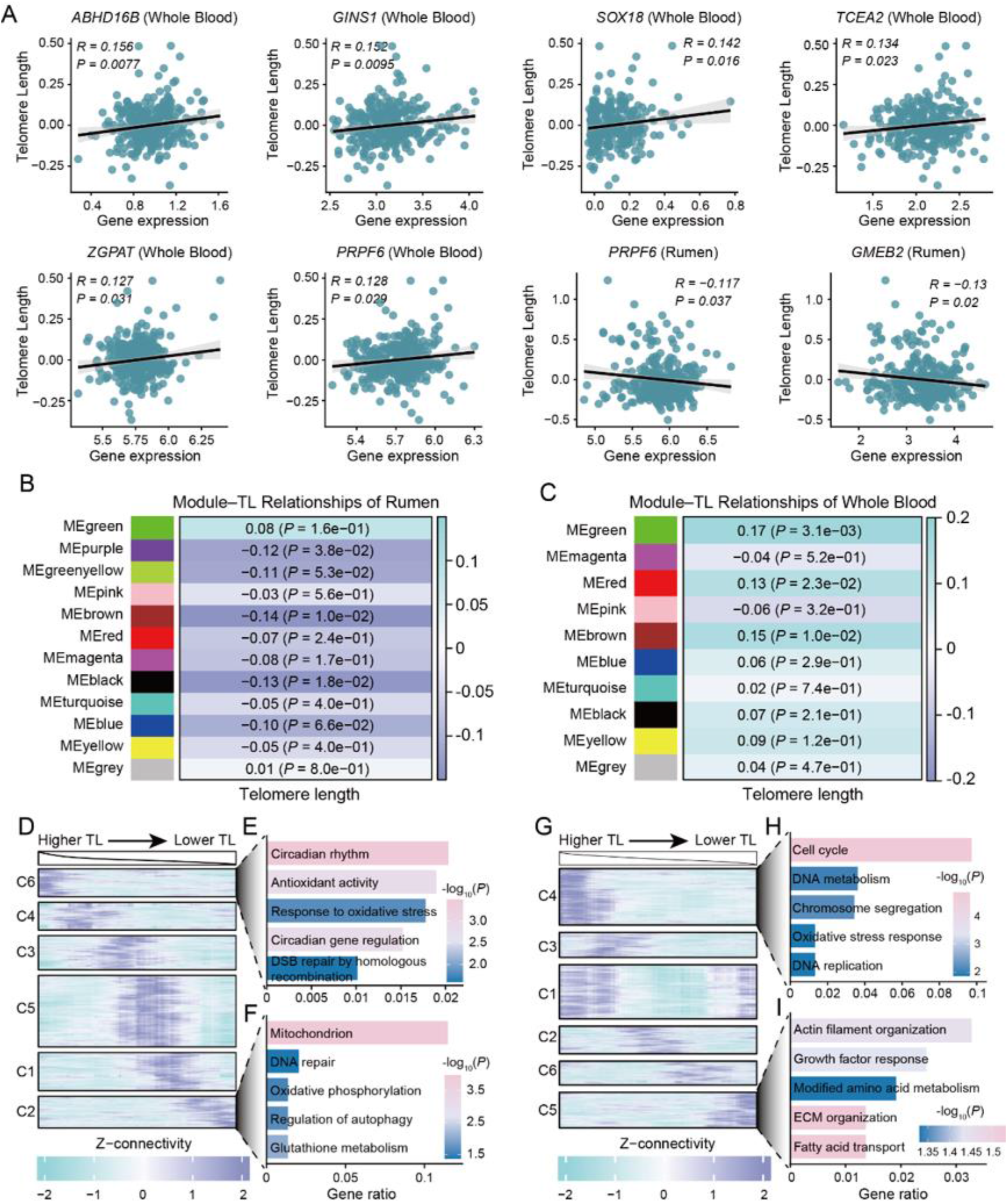
Tissue-specific transcriptional networks and connectivity dynamics associated with telomere length. (A) Correlations between expression levels of GWAS candidate genes and telomere length (TL) in rumen and whole blood. (B and C) WGCNA module–TL associations in rumen (B) and whole blood (C). Heatmaps show Pearson correlations between module eigengenes (MEs) and corrected TL, with corresponding *P* values. (D) TL-stratified sliding-window analysis of gene network connectivity in rumen. Samples were ordered from longer to shorter TL, and connectivity was calculated for highly variable genes in consecutive windows. Rows represent genes, columns represent TL-ordered windows, and colors indicate Z-scored connectivity across windows. Genes were grouped into six K-means clusters (C1–C6) according to their connectivity trajectories. (E and F) GO enrichment of rumen connectivity clusters showing higher connectivity in long-TL windows (Cluster 6; E) or short-TL windows (Cluster 2; F). Abbreviated GO term labels correspond to the following full terms: Glutathione metabolism, glutathione metabolic process; Circadian gene regulation, circadian regulation of gene expression; and DSB repair by homologous recombination, double-strand break repair via homologous recombination. (G) TL-stratified sliding-window analysis of gene network connectivity in whole blood, performed as in (D). (H and I) GO enrichment of whole-blood connectivity clusters showing higher connectivity in long-TL samples (Cluster 4; H) or short-TL samples (Cluster 5; I). Abbreviated GO term labels correspond to the following full terms: DNA metabolism, DNA metabolic process; Oxidative stress response, response to oxidative stress; Growth factor response, response to growth factor; Modified amino acid metabolism, modified amino acid metabolic process; and ECM organization, extracellular matrix organization.

Weighted gene co-expression networks were then constructed separately for each tissue to identify TL-associated transcriptional programs. Hierarchical clustering again showed no clear outliers (Figure S16A and S16E), and soft-thresholding powers were selected according to scale-free topology fit and network connectivity (Figure S16B and S16F). The rumen network resolved 12 modules (Figure S16C), three of which correlated negatively with TL: MEpurple (R = −0.12, P = 0.038), MEbrown (R = −0.14, P = 0.010), and MEblack (R = −0.13, P = 0.018) (Figure 5B and Supplementary Table 17). MEblack was enriched for DNA-templated DNA replication, replication-fork processing, DNA-damage sensing, mitotic checkpoint signaling, and homologous recombination (Figure S16D and Supplementary Table 18). In whole blood, 10 modules were resolved (Figure S16G); in contrast to rumen, the TL-associated modules, including MEgreen (R = 0.17, P = 3.1 × 10−3), MEred (R = 0.13, P = 0.023), and MEbrown (R = 0.15, P = 0.010), correlated positively with TL (Figure 5C). MEgreen was enriched for DNA-replication initiation, replication-fork processing, DNA-damage response, double-strand-break repair via homologous recombination, and cell-cycle checkpoint signaling (Figure S16H and Supplementary Table 19). Although the direction of module-trait association differed between tissues, the enriched biology converged on a DNA replication-replication stress-DNA repair axis, implicating replication homeostasis as a shared molecular correlate of telomere-length variation.

Beyond static module-trait relationships, we asked whether network connectivity itself changes along the TL gradient. In rumen, samples were ordered by bPC-corrected TL residuals, and a 79-sample window, corresponding to approximately 25% of the cohort, was moved along this gradient with a step size of one sample. Within each window, the connectivity of highly variable genes was computed using WGCNA (β = 5), after which connectivity trajectories were Z-score standardized and clustered by K-means. Six connectivity patterns were identified (Figure 5D and Supplementary Table 20). Cluster 6, whose connectivity was highest in TL-rich windows and decreased as TL shortened, was enriched for circadian rhythm, antioxidant activity, response to oxidative stress, and homologous recombination repair (Figure 5E). Cluster 2 showed the opposite pattern, with highest connectivity in low-TL windows, and was enriched for mitochondrial function, oxidative phosphorylation, glutathione metabolism, autophagy, and DNA repair (Figure 5F and Supplementary Table 21). Rumen network topology therefore tracks TL status: longer telomeres are associated with stress homeostasis and genome maintenance, whereas shorter telomeres are associated with mitochondrial metabolism and cellular stress responses.

Applying the same strategy to whole blood, with TL residuals corrected for bPCs, age, and mean sequencing depth and connectivity calculated using β = 3, again yielded six connectivity clusters (Figure 5G). Cluster 4, which was most connected in high-TL samples, was enriched for cell cycle, chromosome segregation, DNA replication, DNA metabolic processes, and oxidative-stress response (Figure 5H), consistent with a program linked to genome stability and telomere integrity. Conversely, cluster 5 dominated low-TL samples and was enriched for actin-filament organization, fatty-acid transport, growth-factor response, modified amino-acid metabolism, and extracellular-matrix organization (Figure 5I and Supplementary Table 22). TL variation therefore extends beyond replication- and genome-stability-related networks, encompassing metabolic, cytoskeletal, and extracellular-matrix programs as well.

## Discussion

T2T assembly has opened previously inaccessible genomic territories, including telomeres, subtelomeres, centromeres, and other repeat-rich domains, to direct analysis of natural variation^7,8,26^. With recent T2T assemblies reported for goat, sheep, and horse, livestock genomics is entering the complete-assembly era^29-31^. Most of these efforts, however, have focused on generating and characterizing single-individual assemblies; how chromosome-end variation can be translated into population phenotyping and genetic association analysis has remained largely unexplored. The Holstein_T2T presented here addresses both aspects of this gap by completing the reference coordinate framework for telomeric and centromeric regions and linking that framework to population-scale telomere phenotyping and association analysis, thereby laying groundwork for chromosome-end genetics in livestock.

The acrocentric configuration is a common mammalian karyotypic architecture in which each autosome carries a short arm adjacent to the centromere and a long arm extending away from it, so that a single chromosome presents two structurally distinct ends^32^. In bovids, this architecture has been shaped in part by Robertsonian fusions, which segregate in cattle populations and can impair fertility^33,34^. Notably, the chromosome-wide Hi-C interaction map of the donor Holstein cow showed no evidence of large-scale interchromosomal fusion, consistent with the canonical bovine karyotype of 2n = 60 and indicating that the chromosome-end patterns described here are unlikely to reflect a major Robertsonian rearrangement in this individual. Recent T2T studies add further nuance: a natural neocentromere on the cattle X chromosome shows that centromere position can shift outside pre-existing satellite contexts ^28^, while goat and sheep assemblies document rapid repeat turnover in acrocentric and sex-chromosome regions^29,35^. These observations indicate that centromere function is maintained on a highly plastic sequence substrate, such that chromosome-scale collinearity alone is insufficient for centromere comparison and repeat-array copy number and organization must be treated as core features of centromere diversity^9,10^. Against this evolutionary backdrop, the 5p/3p telomere-length asymmetry observed in acrocentric cattle chromosomes echoes the chromosome-specific, inter-individually conserved telomere-length patterns reported in humans^27^. One interpretation is that the two chromosome ends experience distinct sequence and chromatin environments: the centromere-proximal short-arm end resides within a heterochromatic neighborhood that may constrain telomere behavior and dampen the effect of TVRs, whereas the centromere-distal long-arm end, which is more often dominated by continuous canonical-repeat tracts, permits a more direct inverse relationship between TVR abundance and TL. Mechanistically, non-canonical repeats have been proposed to weaken shelterin binding and telomerase substrate efficiency and may also mark recombination- or ALT-like telomere-maintenance activity^36,37^. Subtelomeric GC/CpG enrichment could further modulate recruitment of telomere-associated complexes through effects on nucleosome occupancy and chromatin accessibility, extending a broader form of telomere position effect^38^. The chromosome-end telomere phenotype of Holstein cattle can therefore be decomposed into three interconnected layers: terminal repeat architecture, subtelomeric sequence context, and local epigenetic state.

Beyond the single reference genome, this study provides a population-scale strategy for telomere phenotyping in dairy cattle. WGS-based estimators such as TelSeq and Computel are widely used in large human cohorts, yet the technical variation they carry, driven by sequencing depth and batch effects, is particularly pronounced in low-depth livestock datasets^39^. Correcting telomere length with genome-wide coverage-derived principal components reduced this systematic bias and recovered the expected age-dependent pattern of telomere shortening. The three major loci identified using the corrected phenotype are notable because they harbor genes that rank among the most robust signals in human telomere GWAS^19^, arguing that these regions represent conserved components of telomere-length regulation rather than breed- or population-specific coincidences. The implicated genes accord with established telomere biology: *RTEL1* supports telomeric replication and replication stability, *CTC1* encodes a core CST component responsible for C-strand fill-in during chromosome-end replication, and *TERT* encodes the catalytic subunit of telomerase, whose expression is silenced in most adult somatic tissues^40-42^. Multi-tissue regulatory evidence points to distinct routes for these loci. At chromosome 19, convergence of expression- and splicing-level colocalization across tissues identifies *CTC1* as the most plausible effector. The chromosome 20 region, despite carrying the strongest association, contains *TERT*, a gene expressed at very low levels in adult tissues, and may therefore act through non-coding regulatory variation or developmental windows of expression. TWAS adds an additional layer by nominating *GINS1*, a replication-fork component, and *PFAS*, an enzyme involved in de novo purine biosynthesis, linking telomere-length variation to replication capacity and nucleotide supply.

Transcriptomic evidence reinforces the same mechanistic axis. Because *GINS1* is a core component of the CMG replicative helicase and replication forks are intrinsically challenged at telomeric repeats^43^, the *GINS1* association links telomere-length variation to replication efficiency. The dual eQTL and sQTL colocalization at *CTC1* suggests that this gene may modulate telomere homeostasis through both expression-level and transcript-structure regulation. More broadly, co-expression and connectivity analyses in rumen and whole blood converged on DNA replication, replication stress, and DNA repair, consistent with the view that telomeres are difficult-to-replicate regions that behave as fragile sites when replication stress is unresolved ^44,45^. The fact that some TL-associated modules changed direction between tissues emphasizes that the relationship between telomere length and the transcriptome is strongly dependent on cellular context.

For dairy breeding, the implications are twofold. Telomere length has been proposed as an indicator of health and productive lifespan^22^, and the scalable, low-bias phenotyping framework developed here offers a practical route for bringing telomere-related traits into genomic evaluation. In parallel, the extensive structural variation documented in acrocentric and centromeric regions argues for karyotype-level screening in breeding programs, particularly in populations where Robertsonian fusions segregate^33,46^.

### Limitations of the study

Several limitations should be considered. The T2T assembly derives from a single Holstein individual, and whether the chromosome-end architectures described here generalize across the breed or to other cattle populations will require additional T2T genomes. Among the chromosome ends amenable to stringent long-read-based calibration, 42 achieved single-base agreement, whereas other termini could not be evaluated with the same level of confidence, which may limit some chromosome-arm comparisons. The GWAS, colocalization, and TWAS evidence supporting *CTC1* and other candidate genes is statistical in nature, and the causal roles of the lead variants and their regulatory mechanisms will require experimental validation. Telomere length phenotypes were estimated from whole-genome sequencing of peripheral blood; because telomere length can vary among tissues, the extent to which the genetic associations identified here reflect telomere-length regulation in other tissues remains to be determined.

By moving from assembling chromosome ends to interpreting them, this study turns Holstein cattle termini from passive assembly targets into measurable, associable, and comparable genetic features. The combination of a T2T reference, chromosome-end-resolved telomere phenotyping, and population genetic evidence provides a general framework for livestock telomere biology. As Bos taurus pangenome resources and functional validation datasets accumulate, telomeric and centromeric regions may become increasingly tractable substrates for cattle breeding and biomedical research.

## RESOURCE AVAILABILITY

### Lead contact

Further information and requests for resources and data should be directed to and will be fulfilled by the lead contact Xihe Li.

### Materials availability

This study did not generate new unique reagents or materials. Requests for access to non-public datasets or derived resources should be directed to the lead contact and will be subject to institutional and data-use restrictions.

### Data and code availability

Raw sequencing data generated in this study have been deposited under BioProject PRJNA1413493, and CENP-A CUT&Tag sequencing data are available under accession numbers SRR40514003, SRR40514004, SRR40514005, and SRR40514006. The Holstein_T2T genome assembly will be deposited in GenBank and/or CNGB before publication. Population WGS genotype data are not publicly available because of data-access restrictions but may be requested from the lead contact. The rumen RNA-seq dataset is available through CNGB under accession CNP0003806. Publicly available CENP-A CUT&Tag datasets (SRR28462358 and SRR28462359), CattleGTEx cis-eQTL and sQTL summary data, and FarmGTEx TWAS prediction models were also used in this study.

Analysis pipelines and scripts used for the Holstein_T2T genome analyses are available at https://github.com/liufangyuan0613/Holstein_T2T_pipeline. Scripts and workflows used for population-scale telomere-length estimation and genetic association analyses are available at https://github.com/WTao073/cattle-telomere-gwas. Additional information required to reproduce the analyses is provided in the STAR Methods and Supplementary Information.

## ACKNOWLEDGMENTS

We thank Dr. Rui Ding from the Saikexing Research Institute for assistance with sample collection. We also thank Prof. Ying Lu from the Shanghai Ocean University with helpful discussions. We acknowledge the CattleGTEx and FarmGTEx consortia for making their summary statistics and prediction models publicly available. This work was supported by the National Technology Innovation Center for Dairy (No. 2025-JSGG-011; project: “Development of Key Technologies for Functional Gene Discovery and Genetic Performance Testing in High-Yield, High-Fertility, and Long-Production-Life Dairy Cattle [China Dairy Cattle Phase II]”) and the project “Research and Development of Key Technologies for Dairy Cattle Biological Breeding and Creation of Breeding Materials” (No. 2026ZD0079).

## AUTHOR CONTRIBUTIONS

W.S., Y.J., Y.W., and X.L. conceived the study and supervised the research. P.C., L.W., Y.L., and G.Z. coordinated sample collection and provided biological materials and resources. F.L., P.C., W.W., J.L., R.W., and L.W. performed genome assembly, assembly evaluation, genome annotation, repeat annotation, and analyses of telomeric, subtelomeric, and centromeric regions. F.L., W.W., J.L., and X.B. performed telomere-length calibration, telomere variant repeat analysis, and chromosome-end sequence and methylation analyses. T.W., F., Z.Z., F.M., and Z.W. performed population-scale telomere-length estimation, phenotype correction, genome-wide association analysis, and conditional association analysis. T.W., F., Z.Z., F.M., and Z.W. performed eQTL, sQTL, TWAS, and transcriptomic integration analyses. P.L. and Z.W. contributed to data analysis and interpretation. F.L. and T.W. integrated the genomic, genetic, and transcriptomic results. F.L., T.W., P.C., W.W., F., P.L., and Z.W. prepared the figures and tables. F.L. and T.W. drafted the manuscript. Y.J., Y.W., X.L., P.C., W.W., F., P.L., and Z.W. revised the manuscript and contributed to interpretation of the results. Y.J., Y.W., and X.L. jointly supervised the project. All authors read and approved the final manuscript.

F.L., T.W., P.C., W.W., and F. contributed equally to this work. W.S., Y.J., Y.W., and X.L. are co-corresponding authors.

## STAR★METHODS

Detailed methods are provided in the online version of this paper and include the following

KEY RESOURCES TABLE

RESOURCE AVAILABILITY

Lead contact;
Materials availability;
Data and code availability

EXPERIMENTAL MODEL AND STUDY PARTICIPANT DETAILS

T2T assembly donor;
Population WGS cohort;
RNA-seq cohorts

METHOD DETAILS

Sample collection and DNA extraction;
Short-read library preparation and sequencing;
PacBio HiFi sequencing;
NT ultra-long sequencing;
Hi-C library preparation and sequencing;
Sequencing data quality control;
Draft genome assembly and version selection;
Removal of contaminant sequences;
Chromosome-scale scaffolding with Hi-C; Gap filling and genome polishing;
Telomere assembly and end calibration;
Assembly quality assessment; Previously unresolved region analysis;
Repeat annotation;
Centromere identification and comparison;
CENPA CUT&Tag profiling;
Telomere and subtelomere analysis; Gene annotation;
CpG methylation profiling;
rDNA,
SD and unique k-mer analysis;
Hi-C interaction map construction;
WGS variant processing and imputation; WGS-based telomere length estimation;
Coverage-based telomere length correction;
Telomere length GWAS;
Conditional association analysis;
eQTL and sQTL colocalization;
Multi-tissue TWAS;
RNA-seq processing and cell correction;
Weighted gene co-expression analysis;
Sliding-window connectivity analysis

QUANTIFICATION AND STATISTICAL ANALYSIS

## KEY RESOURCES TABLE

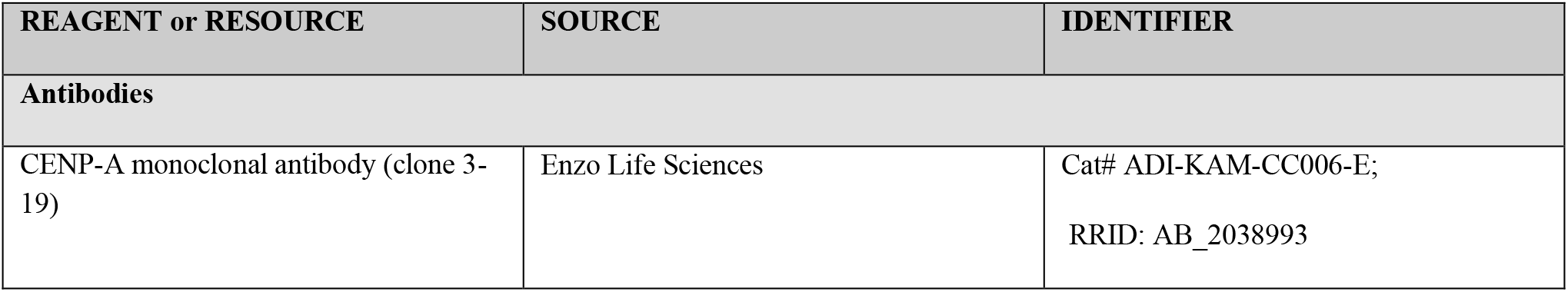
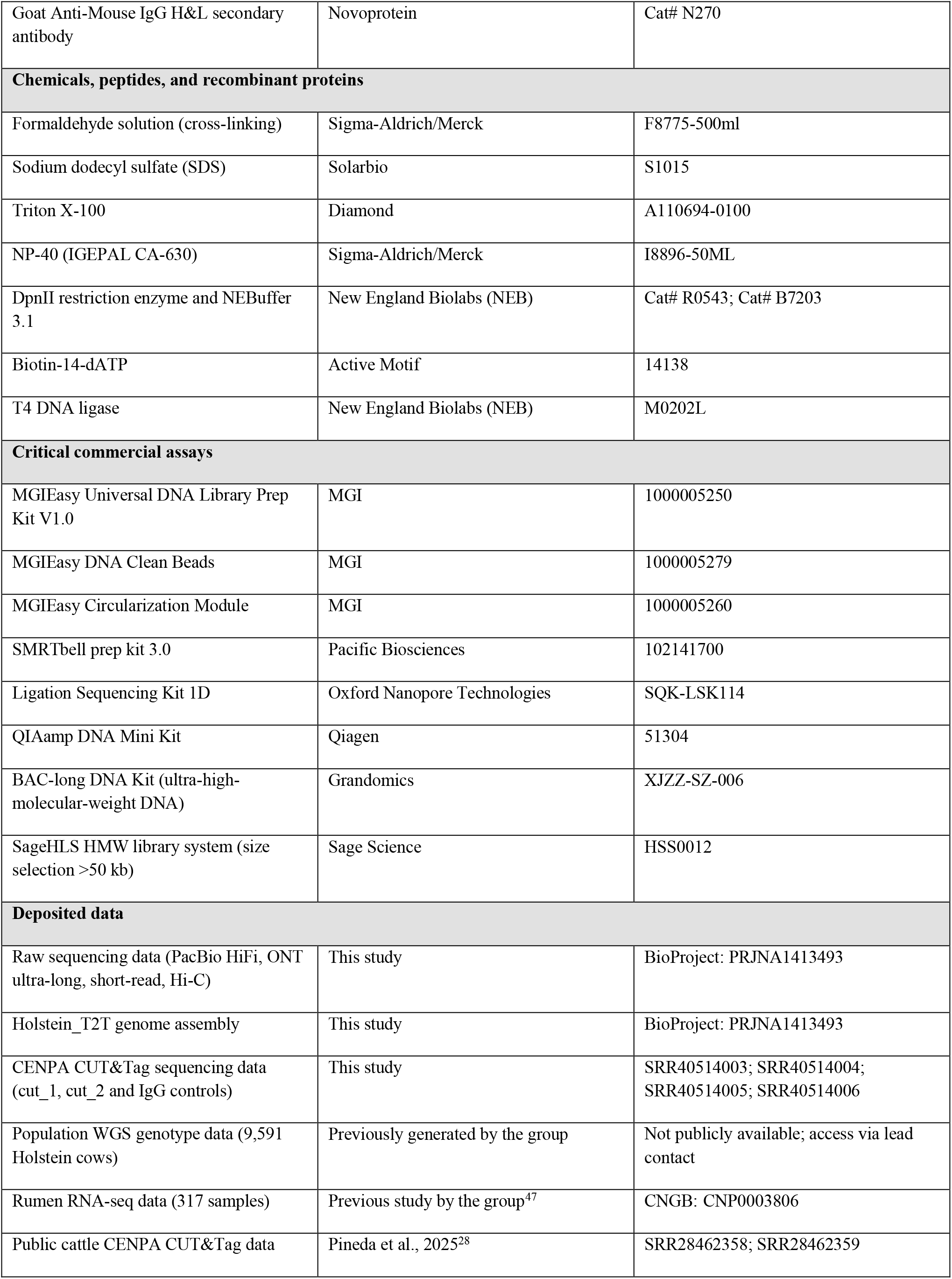
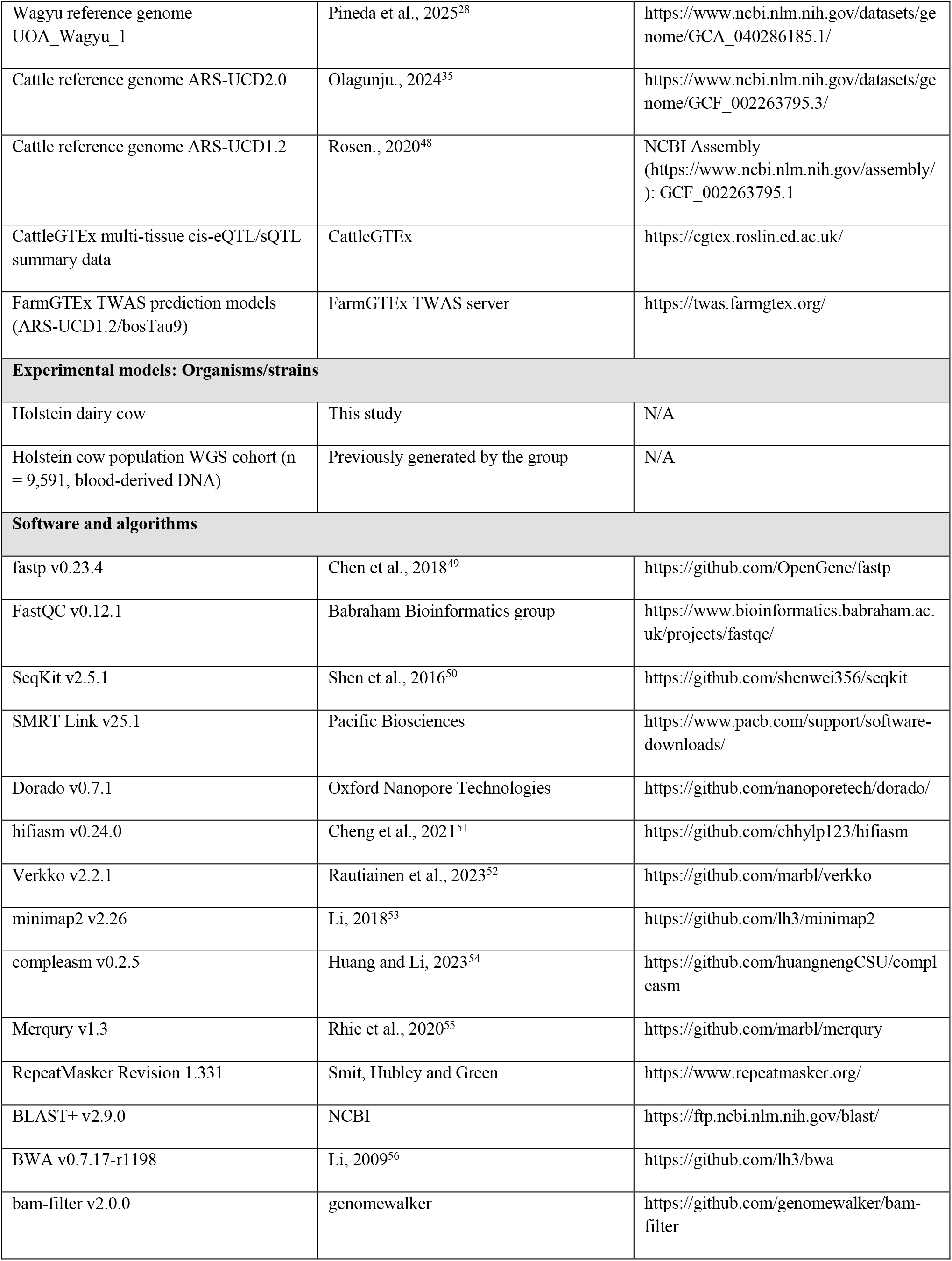
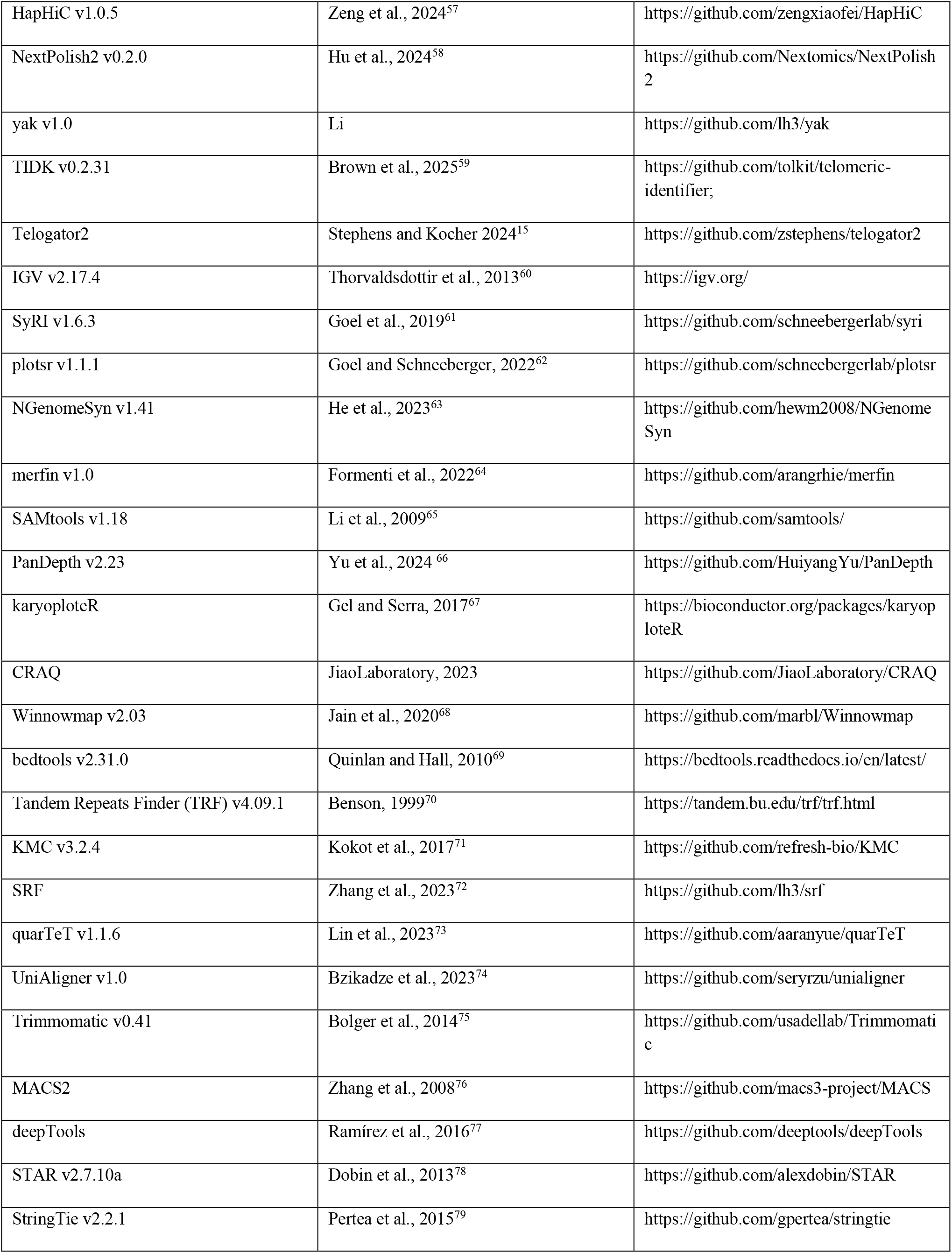
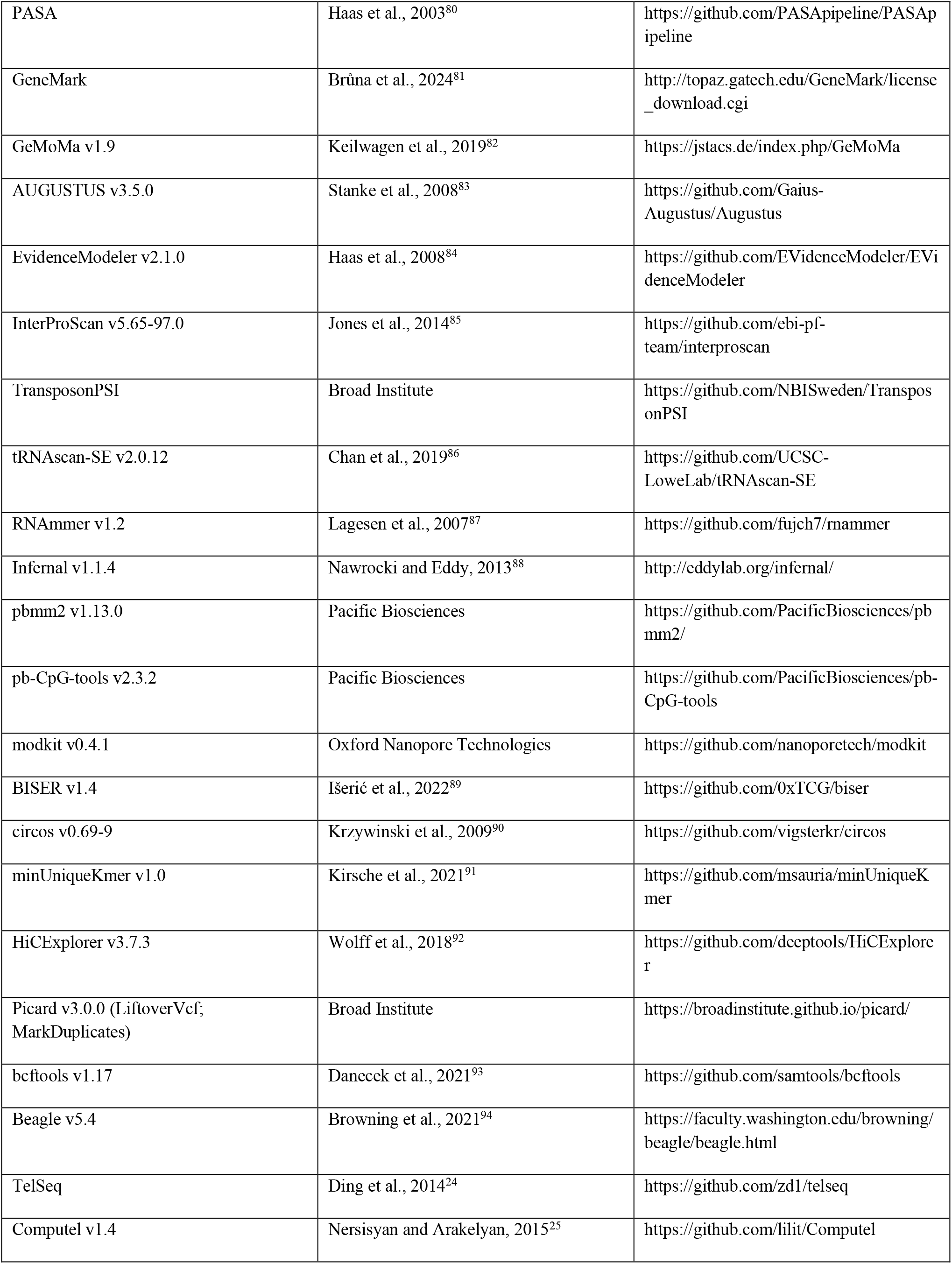
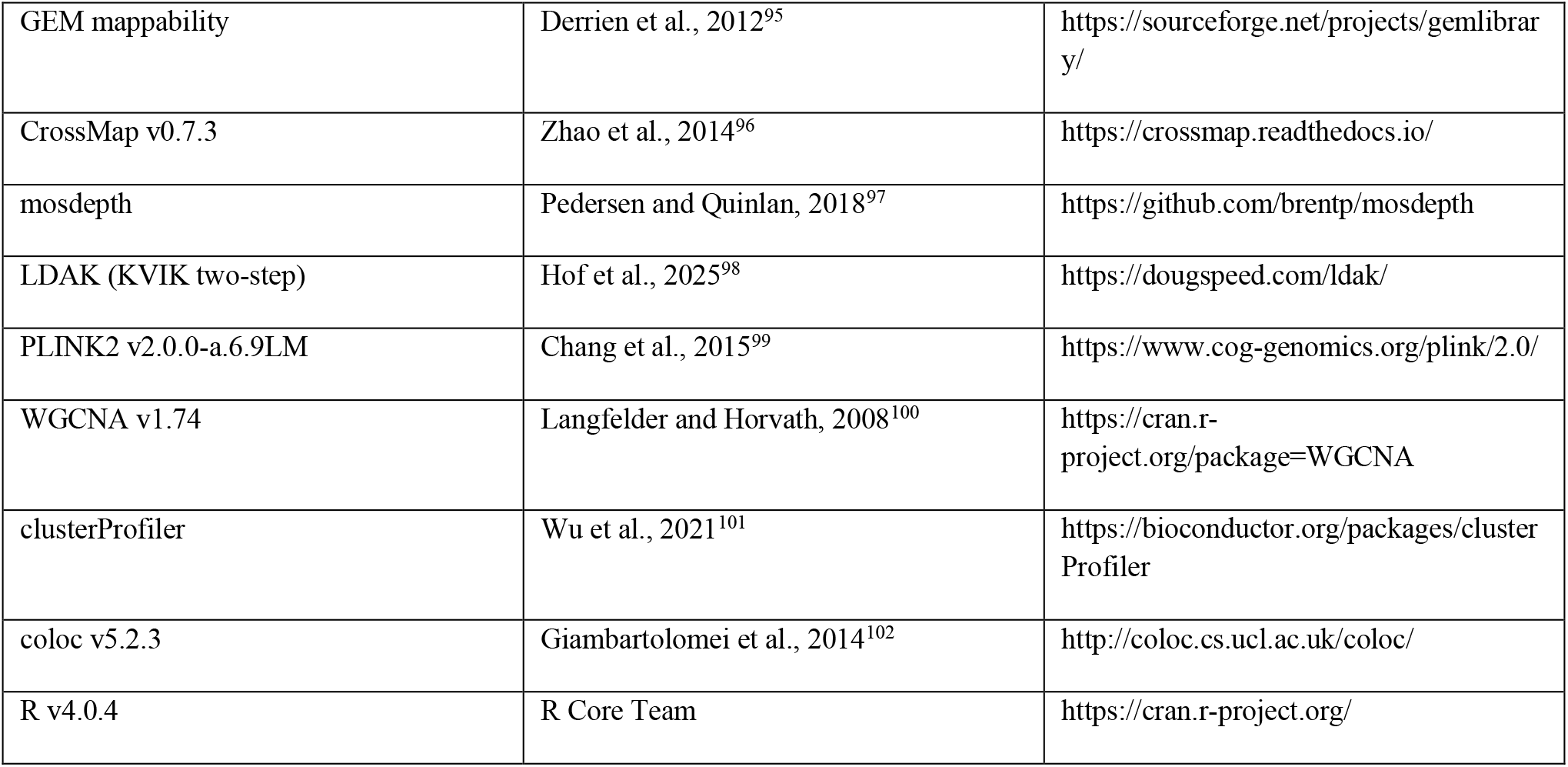

### EXPERIMENTAL MODEL AND STUDY PARTICIPANT DETAILS

#### T2T assembly donor

A single seven-year-old female Holstein dairy cow (*Bos taurus*) was used for reference genome sequencing and assembly. Peripheral blood was collected from this individual and used for all downstream genome sequencing, including PacBio HiFi, Oxford Nanopore Technologies (ONT) ultra-long, short-read, and Hi-C sequencing; all genomic data therefore originated from the same animal. The resulting reference assembly represents the genome of a single female individual and comprises the 29 bovine autosomes and the X chromosome. Because Holstein_T2T was generated from a single female animal, sex-related variation in genome structure was not assessed. All animal procedures were reviewed and approved by the **Bioethics Committee of Inner Mongolia University** (approval no. **IMU-cattle-2025-228**) and were conducted in accordance with the ethical principles approved by the committee.

#### Population WGS cohort

Whole-genome resequencing (WGS) data from Holstein cows previously generated by the group were integrated for telomere length genetic analysis. A total of 9,591 individuals with available age information were retained and stratified by sequencing depth into a high-depth group (≥4.5×; *n* = 1,498; mean depth = 7.09×) and a low-depth group (<4.5×; *n* = 8,093; mean depth = 1.23×). Genomic DNA for all individuals was extracted from peripheral blood. All animals were female Holstein cows; because the cohort was single-sex, the influence of sex on telomere length genetics was not assessed. All animal procedures for the population cohort were reviewed and approved by the Bioethics Committee of Inner Mongolia University under the same protocol (approval no. IMU-cattle-2025-228).

#### RNA-seq cohorts

RNA-seq data from 317 rumen samples and 289 whole-blood samples were used for gene expression analyses, weighted gene co-expression network analysis and telomere length-stratified connectivity analyses. The rumen RNA-seq dataset was derived from a previous study by the group^47^.

### METHOD DETAILS

#### Sample collection and DNA extraction

Peripheral blood was collected from the seven-year-old Holstein cow described above. High-molecular-weight (HMW) genomic DNA was prepared by the CTAB method and purified with the Grandomics Genomic kit following the manufacturer’s standard operating procedure. DNA degradation and contamination were monitored on 1% agarose gels. DNA purity was assessed with a NanoDrop One UV-Vis spectrophotometer (Thermo Fisher Scientific), requiring OD260/280 ratios of 1.8–2.0 and OD260/230 ratios of 2.0–2.2, and DNA concentration was measured with a Qubit 4.0 fluorometer (Invitrogen). DNA integrity was verified on the Femto Pulse system (Agilent Technologies). For ONT ultra-long sequencing, ultra-HMW DNA was additionally extracted using the BAC-long DNA Kit following the manufacturer’s guidelines, and large DNA fragments were retrieved by gel cutting on the BluePippin/SageHLS system (Sage Science).

#### Short-read library preparation and sequencing

Short-read libraries were constructed with the MGIEasy Universal DNA Library Prep Kit V1.0 (MGI, Cat# 1000005250) following the standard protocol. Briefly, 1 µg of genomic DNA was randomly fragmented with a Covaris focused-ultrasonicator, and fragments were size-selected to an average insert size of 200–400 bp with MGIEasy DNA Clean Beads (MGI, Cat# 1000005279). Size-selected fragments were end-repaired, 3′-adenylated and ligated to sequencing adapters. Adapter-ligated DNA was PCR-amplified and purified with MGIEasy DNA Clean Beads. Double-stranded PCR products were heat-denatured and circularized with the splint oligo of the MGIEasy Circularization Module (MGI, Cat# 1000005260), and the resulting single-stranded circular DNA library was quantified and sequenced in 150-bp paired-end (PE150) mode on the MGI DNBSEQ-T7 platform. A total of 309.60 Gb of raw data (2,064,006,116 reads) was generated; after quality filtering, 308.70 Gb of clean data (2,063,375,080 reads; Q30 = 95.25%; GC = 42.62%) was retained.

#### PacBio HiFi sequencing

SMRTbell libraries were prepared with the SMRTbell prep kit 3.0 (Pacific Biosciences) according to the manufacturer’s manual. Briefly, HMW genomic DNA passing Femto Pulse quality control was sheared on the Megaruptor 3 system (Diagenode), followed by DNA cleanup, DNA repair and A-tailing reactions, adapter ligation and cleanup, nuclease treatment to remove incompletely ligated molecules, and final size selection and cleanup of the library. Sequencing was performed on the PacBio Revio system (two SMRT cells) at Grandomics (Wuhan, China). Circular consensus sequencing (CCS) was performed with default parameters (min passes = 3, min predicted accuracy RQ = 0.99) to generate high-fidelity (HiFi) reads with quality above Q20, and SMRT Link v25.1 was used with default parameters to remove fail reads. In total, 213.75 Gb of HiFi data (10,668,688 reads; mean length 20,035 bp; read N50 20,657 bp; 99.9% of bases in reads >10 kb) was obtained, corresponding to approximately 71× coverage.

#### ONT ultra-long sequencing

For ultra-long Nanopore libraries, approximately 8–10 µg of HMW genomic DNA was size-selected for fragments longer than 50 kb with the SageHLS HMW library system (Sage Science) and processed with the Ligation Sequencing Kit 1D (Oxford Nanopore Technologies, Cat# SQK-LSK114) according to the manufacturer’s instructions. Approximately 400 ng of library was loaded per flow cell and sequenced on the PromethION platform (Oxford Nanopore Technologies) at Grandomics (Wuhan, China). Raw signal data (pod5) were base-called with Dorado v0.7.1 in super-accuracy (sup) mode (model dna_r10.4.1_e8.2_400bps_sup@v5.0.0), and low-quality fail reads (sup mode, QV < 7) were removed. Nine flow cells produced a total of 187.97 Gb of pass data (7,984,411 reads; read N50 90,054 bp), of which 134.2 Gb, 85.3 Gb and 27.5 Gb were contained in reads longer than 50 kb, 100 kb and 200 kb, respectively, corresponding to approximately 62× coverage.

#### Hi-C library preparation and sequencing

Hi-C libraries were prepared from 0.1 mL of peripheral blood from the same individual. Cells were cross-linked for 10 min with fresh formaldehyde at a final concentration of 1% and quenched with glycine at a final concentration of 0.2 M for 5 min. Cross-linked cells were lysed in lysis buffer (10 mM Tris-HCl pH 8.0, 10 mM NaCl, 0.2% NP-40 and cOmplete protease inhibitors [Roche]). Extracted nuclei were resuspended in 150 µL of 0.1% sodium dodecyl sulfate (SDS) and incubated at 65 °C for 10 min, after which SDS was quenched by adding 120 µL water and 30 µL 10% Triton X-100 and incubating at 37 °C for 15 min. Nuclear DNA was digested with 150 U of DpnII in 30 µL of 10× NEBuffer 3.1 (100 mM NaCl, 50 mM Tris-HCl, 10 mM MgCl2, 100 µg/mL BSA, pH 7.9) at 37 °C overnight, and the enzyme was inactivated at 65 °C for 20 min. Cohesive ends were filled in with dTTP, dCTP, dGTP and Biotin-14-dATP using 40 U of Klenow fragment at 37 °C for 2 h. Proximity ligation was performed with 20 U of T4 DNA ligase in blunt-end ligation buffer (300 mM Tris-HCl, 100 mM MgCl2, 100 mM DTT, 1 mM ATP, pH 7.8) at 16 °C for 4 h. Cross-links were reversed with 200 µg/mL proteinase K (Thermo) at 65 °C overnight. DNA was purified with the QIAamp DNA Mini Kit (Qiagen) according to the manufacturer’s instructions, sheared to approximately 400 bp, and biotin-labeled ligation junctions were pulled down with streptavidin-coated magnetic beads (Dynabeads). The library was sequenced in PE150 mode, yielding 407.22 Gb of raw data (2,714,776,684 reads); 405.85 Gb of clean data (2,713,182,440 reads; clean rate 99.94%; Q30 = 97.3%; GC = 47.0%) was retained for scaffolding, corresponding to approximately 135× coverage.

#### Sequencing data quality control

Short reads and Hi-C reads were quality-filtered with fastp v0.23.4 ^49^ to remove adapter sequences and low-quality reads (short reads: -n 0 -f 5 -F 5 -t 5 -T 5 -q 20; Hi-C reads: -z 4, discarding read pairs containing adapters, more than five ambiguous bases (N), or pairs in which more than 40% of bases in either read had quality scores below 15). Clean short-read data were further inspected with FastQC v0.12.1. PacBio HiFi reads (quality ≥ Q20) were quality-controlled with SMRT Link v25.1 using default parameters. ONT reads were base-called and filtered with Dorado in sup mode as described above, and only pass reads were used. For hybrid assembly, ONT reads were further filtered with SeqKit v2.5.1^50^ (seq --min-len 50000) to retain only reads of at least 50 kb.

#### Draft genome assembly and version selection

Three draft assembly strategies were compared. (i) HiFiasm: HiFi reads were assembled with hifiasm v0.24.0^51^ using default parameters, yielding a 3,315.78-Mb assembly of 281 contigs with a contig N50 of 90.22 Mb. (ii) HiFiasmMix: HiFi reads combined with ONT ultra-long reads longer than 50 kb were assembled with hifiasm v0.24.0 using the --ul option, yielding a 3,318.89-Mb assembly of 179 contigs with a contig N50 of 101.93 Mb. (iii) VerkkoMix: the same HiFi and >50-kb ONT read sets were assembled with Verkko v2.2.1^52^ using --hifi HiFireads --nano ONT50kreads --no-cleanup, yielding a 6,046.55-Mb assembly of 2,340 contigs with a contig N50 of 10.97 Mb. Draft assemblies were compared by contig statistics, collinearity dot plots against ARS-UCD2.0 and between assemblies (minimap2 v2.26^53^ with paf2dotplot.r), BUSCO completeness (compleasm v0.2.5^54^, mammalia_odb10) and k-mer-based consensus quality (Merqury v1.3^55^ with merfin-filtered 21-mers from short reads and HiFi reads). The HiFiasmMix assembly showed the best balance of continuity and completeness, with fewer contigs and higher contig N50 than the HiFi-only assembly, while avoiding the inflated size and fragmentation of the VerkkoMix assembly, and was therefore selected for all downstream steps.

#### Removal of contaminant sequences

Before chromosome construction, the selected assembly was screened for contaminant sequences. Repetitive sequences were soft-masked with RepeatMasker (Revision 1.331; -nolow -no_is -gff -norna -engine abblast -lib lib). Sequences longer than 1 Mb were split into non-overlapping 50-kb bins, and all sequences were searched against the NCBI nucleotide (NT) database with BLASTN (BLAST+ v2.9.0, default parameters). Sequences of 1 Mb or less were classified by their best hit, whereas longer sequences were assigned the taxonomic label supported by the majority of bins. Of 160 classified contigs (3,256.64 Mb in total), 98 contigs (99.07% of total length) were assigned to Metazoa; two contigs were of bacterial origin and one contig was of mitochondrial/plastid origin. Non-target (non-metazoan), mitochondrial and plastid-derived contigs were removed, yielding a decontaminated assembly of 3,256.47 Mb (contig N50, 101.93 Mb) with a BUSCO completeness of 99.94% (mammalia_odb10).

#### Chromosome-scale scaffolding with Hi-C

Clean Hi-C read pairs were aligned to the decontaminated contig assembly with BWA v0.7.17^56^ using mem -5SP. Alignments were filtered with bam-filter v2.0.0 (1 --nm 3), retaining read pairs with mapping quality ≥ 1 and edit distance ≤ 3. Chromosome-scale scaffolding was performed with HapHiC v1.0.5^57^ using default parameters. Briefly, HapHiC pre-processed the assembly to correct misjoined contigs, break erroneous joins, filter short contigs and remove allelic Hi-C links; contigs were then clustered into chromosome groups with the Markov clustering (MCL) algorithm, followed by a reassignment step to rescue contigs that had been filtered out or incorrectly grouped. Ordering and orientation were performed with the integrated fast 3D-DNA iterative sorting and ALLHiC optimization implemented in HapHiC, and pseudomolecules were built from the resulting chromosome assignments. The Hi-C contact map was manually inspected and curated, and the curated assembly was re-evaluated for contiguity and BUSCO completeness. A total of 3,009.12 Mb (96.02% of the input assembly) was anchored onto 30 pseudomolecules (29 autosomes and the X chromosome) with a scaffold N50 of 116.86 Mb, leaving six residual gaps.

#### Gap filling and genome polishing

Unanchored fragmented sequences were discarded before gap filling. ONT reads were aligned to the chromosome assembly with minimap2 v2.26 (--secondary=no -ax map-ont), and reads were retained as candidate gap-filling evidence if they were unmapped, poorly aligned (alignment identity < 80% or aligned read coverage < 80%), or uniquely spanned both flanking sequences of a gap. For each unresolved interval, candidate ONT reads were collected from the corresponding flanking regions and subjected to local iterative assembly. Locally assembled contigs were aligned back to the chromosome assembly with minimap2 v2.26 (-ax asm5 - -eqx), and a contig was accepted for gap replacement only when it bridged the gap and showed reliable overlap with both flanking sequences. All six gaps were closed, producing a gap-free assembly of 3,012.77 Mb in 30 chromosomes. The gap-filled assembly was then polished with NextPolish2 v0.2.0^58^ using default parameters. K-mer databases were built from quality-filtered short reads with yak v1.0 (count -b37 -k21 and count -b37 -k31), HiFi reads were aligned to the gap-filled assembly with minimap2 v2.26 (- -secondary=no -ax map-hifi), and the two k-mer databases together with the HiFi alignment BAM file were used for repeat-aware polishing. The polished assembly comprised 3.01 Gb in 30 gap-free chromosomes (contig N50, 111.23 Mb).

#### Telomere assembly and end calibration

Telomeric repeats were identified and refined using both assembled chromosome sequences and raw long-read evidence. Telomeric repeat motifs were first surveyed in the chromosome-level assembly with TIDK v0.2.31^59^ using explore --minimum 4 --maximum 12. The canonical vertebrate telomeric repeat motif TTAGGG and its reverse-complement sequence CCCTAA were used to annotate terminal telomeric arrays. For each pseudomolecule, the terminal 100-kb sequences at both ends were extracted with SeqKit v2.5.1 and searched for consecutive telomeric repeat arrays (locate -i -p TelomereSeq); a chromosome end was considered telomere-positive when at least ten consecutive copies of the canonical motif were detected. After scaffolding, 19 of the 30 chromosomes carried at least one detectable terminal telomeric array.

Chromosome ends lacking complete terminal arrays were refined by local assembly and long-read-supported end correction. HiFi reads that were unmapped to the assembly, contained telomeric repeat motifs, or mapped to the terminal 1-Mb regions of telomere-incomplete chromosomes were extracted and assembled with hifiasm v0.24.0. Candidate telomere-containing contigs were aligned back to the chromosome assembly with minimap2 v2.26 and used to extend or replace incomplete chromosome ends on the basis of sequence overlap, and the updated terminal regions were re-aligned to confirm continuity across the joins. Conversely, long-read evidence also identified overextended or misassembled terminal sequence at the 3′ ends of three chromosomes. These unsupported terminal tails, each approximately 8–13 Mb in length, were removed during end calibration, contributing substantially to the approximately 32-Mb net reduction in assembly size relative to the pre-calibration input. Thus, telomere-end refinement involved both sequence extension at incomplete termini and removal of unsupported terminal sequence where required. Raw-read-based telomere validation and coordinate calibration were then performed with Telogator2^15^. PacBio HiFi and ONT ultra-long reads were used to independently evaluate terminal repeat length, telomere orientation and chromosome-arm assignment. The p- and q-arm telomere assignments reported by Telogator2 were matched to the 5′ and 3′ ends of the assembled pseudomolecules according to chromosome orientation, and the inferred telomere coordinates were used to calibrate the final chromosome termini and to distinguish complete terminal repeat arrays from truncated or internally located telomeric repeats. After calibration, 42 chromosome ends reached base-level (±1 bp) agreement with read-supported telomere lengths, with residual deviations remaining only at the chr13 3p end (1,083 bp) and the chr19 5p end (266 bp). Finally, HiFi and ONT reads were aligned to the completed assembly with minimap2 v2.26, and read support across chromosome termini was inspected in IGV v2.17.4^60^. Only chromosome ends supported by continuous long-read alignments and terminal canonical repeat arrays were retained as completed telomeres. The final telomere coordinates were recorded separately for the 5′ and 3′ ends of each chromosome and used for downstream chromosome-arm-specific telomere and subtelomere analyses.

#### Assembly quality assessment

##### Collinearity analysis

Whole-genome alignments between the final Holstein_T2T assembly and previous cattle reference genomes were generated with minimap2 v2.26 using -ax asm5 --eqx. Structural differences were identified with SyRI v1.6.3^61^ using -F S and visualized with plotsr v1.1.1^62^. Additional chromosome-level synteny plots were produced with NGenomeSyn v1.41^63^ using default parameters.

##### Completeness assessment

Gene-space completeness was evaluated with compleasm v0.2.5 using default parameters against the mammalia_odb10 lineage dataset (n = 9,226 universal single-copy orthologs). The final assembly captured 9,218 complete BUSCO genes (99.92%; 98.8% single-copy, 1.12% duplicated), with 0.04% fragmented and 0.04% missing.

##### Consensus quality assessment

Reference-free consensus quality (QV) was estimated with Merqury v1.3 using 21-mer databases constructed from short-read and HiFi data after k-mer filtering with merfin v1.0 (k = 21)^64^. The final curated assembly reported in this study has a total length of 3.01 Gb, a contig N50 of 111.23 Mb and a Merqury QV of 72.54

##### Read mapping and coverage

Short reads, HiFi reads and ONT reads were aligned back to the final assembly with minimap2 v2.26 using --secondary=no -ax sr, --secondary=no -ax map-hifi and --secondary=no -ax map-ont, respectively, and mapping statistics were computed with SAMtools v1.18^65^ stats. Mapping rates were 96.86% for short reads (1,998,485,624 of 2,063,375,080 reads), 100% for HiFi reads (10,668,501 of 10,668,688 reads) and 93.60% for ONT reads (7,473,801 of 7,984,411 reads; 99.76% after excluding ONT reads shorter than 1 kb). Per-base coverage was averaged over 100-kb windows with PanDepth v2.23 (-w 100000)^66^ and plotted along chromosomes in R v4.0.4 with karyoplote^67^.

##### CRAQ structural accuracy assessment

Single-base- and structural-level assembly accuracy was evaluated with CRAQ using default parameters on the short-read and long-read alignments. CRAQ uses clipping information to quantify regional errors (CREs, summarized as R-AQI) and large structural errors (CSEs, summarized as S-AQI) without relying on a reference genome. The final assembly achieved a genome-wide covered rate of 99.53%, a low-confidence rate of 1.00 × 10−4, an R-AQI of 99.40 and an S-AQI of 99.22.

##### Previously unresolved region analysis

Previously unresolved regions (PURs), defined as regions newly assembled relative to ARS-UCD2.0, were identified by aligning the Holstein_T2T assembly to ARS-UCD2.0 with Winnowmap v2.03^68^ using -cx asm20 -H --MD, followed by interval summarization with bedtools v2.31.0 (intersect, window, merge)^69^. PURs longer than 50 kb were visualized along chromosomes in R.

#### Repeat annotation

Repetitive sequences were annotated by integrating tandem-repeat discovery, transposable-element annotation and satellite-repeat characterization. Tandem repeats were identified with Tandem Repeats Finder (TRF) v4.09.1^70^ using the parameters 2 7 7 80 10 50 500 -f -d -h -l 6. Interspersed repeats and transposable elements were annotated with RepeatMasker (v1.331) using a custom repeat library and the parameters -nolow -no_is -gff -norna -engine abblast -lib, and were classified into major categories including LINEs, SINEs, LTR retrotransposons, DNA transposons, satellite repeats and unclassified repetitive elements. To characterize highly repetitive satellite-rich regions, candidate repeat intervals identified by TRF and RepeatMasker were further analyzed with KMC v3.2.4^71^ to construct k-mer frequency databases, and high-copy tandem repeat units were reconstructed with SRF^72^ using - l 6, allowing the identification of abundant satellite repeat families and their consensus repeat units. Repeat density was calculated in fixed genomic windows along each chromosome to generate chromosome-scale repeat landscapes, and the distribution of major repeat classes was integrated with centromeric and subtelomeric annotations for downstream analyses of chromosome architecture.

#### Centromere identification and comparison

Centromeric regions were annotated using a repeat-based strategy. Tandem repeats were identified with TRF, and interspersed repeats were annotated with RepeatMasker as described above. High-repeat-density intervals were extracted from the genome, k-mer databases were constructed with KMC v3.2.4, and candidate satellite repeat units were identified with SRF using -l 6. Candidate centromeric repeat units were aligned back to the genome with BLASTN (BLAST+ v2.9.0, default parameters) to define repeat-enriched candidate centromeric intervals. In parallel, centromeric regions were predicted with quarTeT v1.1.6^73^ using the CentroMiner module (-t 10 -i Genome --TE TEgff) with the TRF and RepeatMasker annotations as input. The SRF/BLASTN-supported intervals and CentroMiner predictions were integrated to define the final centromere coordinates of all 30 chromosomes. The chromosomal distributions of tandem repeats, transposable elements and SRF-derived satellite units were summarized to characterize repeat composition and organization across centromeric regions. For comparative centromere analysis, homologous centromeric intervals were extracted from the Holstein_T2T and the Wagyu reference. Local alignments were generated with minimap2 v2.26 using -ax asm5 --eqx and UniAligner v1.0^74^, and alignment dotplots were used to inspect homologous sequence blocks, repeat-array continuity and local structural changes. Indels within homologous centromeric regions were classified according to alignment structure as insertion-dominant, deletion-dominant or balanced events. Higher-order repeat organization, repeat composition and local expansion or contraction patterns were then summarized across homologous chromosomes.

#### CENPA CUT&Tag profiling

CENP-A CUT&Tag (cleavage under targets and tagmentation) was performed using peripheral blood from Holstein cattle to identify and annotate centromeric regions in the Holstein_T2T. CUT&Tag libraries were generated from two biological replicates (cut_1 and cut_2), together with matched IgG control libraries (cut_1_IgG and cut_2_IgG), and sequenced on the Illumina NovaSeq platform using paired-end sequencing by Jiayin Bio (project JYKF2026020013). A CENP-A monoclonal antibody (3-19; Cat. No. ADI-KAM-CC006-E) was used as the primary antibody. Because the CENP-A primary antibody was raised in mouse, Goat Anti-Mouse IgG H&L (Cat. No. N270) was used as the secondary antibody. Spike-in DNA was included during library preparation for normalization and quality assessment. Raw sequencing reads were trimmed using Trimmomatic v0.41^75^ with default parameters, yielding 11.48 Gb and 12.56 Gb of clean data for cut_1 and cut_2, respectively, with Q30 values of 97.42% and 97.32%. Clean reads were aligned to the Holstein_T2T using BWA, and uniquely mapped reads with MAPQ ≥ 10 were retained for downstream analyses. The mapping rates for cut_1 and cut_2 were 96.69% and 92.32%, respectively. The corresponding IgG control libraries showed substantially lower mapping rates (11.67% and 8.71%), consistent with the low background signal expected for CUT&Tag control libraries containing spike-in DNA. CENP-A-enriched regions were identified using MACS2^76^ based on uniquely mapped reads with a q-value cutoff (q-value < 0.05), yielding 6,091 and 3,930 peaks for cut_1 and cut_2, respectively. Normalized genome-wide CENP-A signal tracks were generated using deepTools^77^. Signal intensity at each genomic position was calculated as R/T × 10^6^, where R represents the number of reads mapped to the locus and T represents the total number of mapped reads. IgG control tracks were processed using the same procedure and used as background controls for visualization in IGV. CENP-A enrichment patterns from the two biological replicates were integrated to identify candidate centromeric regions across the Holstein_T2T assembly. Regions showing reproducible CENP-A enrichment and overlapping or adjacent to centromeric satellite-repeat arrays were used to define sequence-resolved centromere locations. To independently validate the centromere annotations, publicly available bovine CENP-A CUT&Tag datasets (SRR28462358 and SRR28462359) were additionally analyzed and mapped to the Holstein_T2T assembly. Concordance between the independent CENP-A enrichment profiles and the centromeric regions identified in this study was used as orthogonal support for the final centromere annotations.

#### Telomere and subtelomere analysis

Telomeric sequences were annotated from the final chromosome-level assembly. Telomeric repeat motifs were surveyed with TIDK v0.2.31 (explore --minimum 4 --maximum 12), and the canonical vertebrate telomeric repeat TTAGGG was used as the primary motif for terminal repeat detection. Chromosome-terminal sequences were extracted with SeqKit v2.5.1, and telomeric repeats were located using seqkit locate -i -p TelomereSeq. Terminal repeat arrays were recorded separately for both ends of each chromosome, and telomere length was calculated for each chromosome end from the final telomere coordinates. Chromosome termini were assigned as 3p or 5p according to the orientation of the assembled pseudomolecules. Canonical telomeric repeats and telomere variant repeats (TVRs) were quantified within each terminal array: repeat motifs matching TTAGGG were classified as canonical, whereas non-canonical repeat motifs embedded within or adjacent to the terminal telomeric array were classified as TVRs. Chromosome-arm-specific telomere length and TVR composition were then compared across all chromosome ends. Subtelomeric regions were defined as the 500-kb interval immediately internal to each telomeric repeat array, excluding the telomeric tract itself; for chromosome ends with shorter available flanking sequence, the longest non-telomeric interval adjacent to the telomere was used. GC content, CpG density and repeat composition were calculated across subtelomeric regions in sliding windows, and repeat states were assigned using the genome-wide repeat annotations generated from TRF, RepeatMasker and satellite-repeat analyses. A chromosome-end feature matrix was constructed by integrating telomere length, canonical repeat content, TVR composition, GC content, CpG density and subtelomeric repeat-state information, and was used for clustering, ordination and visualization of chromosome-end-specific telomere and subtelomere states, including motif-abundance clustering separating 3p and 5p subtelomeric regions and principal component analysis of motif density with chromosome-end effects tested by PERMANOVA.

#### Gene annotation

Gene annotation was performed using an integrated strategy combining transcriptome-based, homology-based, and ab initio gene prediction. Prior to gene prediction, repetitive regions of the Holstein_T2T were masked using the genome-wide repeat annotation described above. For transcriptome-based annotation, short-read RNA-seq data from multiple tissues were quality filtered and aligned to the Holstein_T2T using STAR v2.7.10a^78^. Transcript structures were assembled from the resulting alignments using StringTie v2.2.1^79^, yielding 71,082 transcript models. These transcripts were subsequently integrated using PASA^80^, which aligned the assembled transcripts to the genome and reconstructed transcript-supported gene structures. Open reading frames were identified using GeneMark^81^, resulting in 68,338 PASA-supported transcript models corresponding to 33,898 predicted genes. PASA was also used after gene-model integration to refine untranslated regions and transcript structures. Homology-based gene prediction was performed using GeMoMa v1.9^82^ with annotated gene models from closely related bovids, including *Bison*, *Bos grunniens*, *Bos indicus*, *Bos javanicus*, and *Bos mutus*. Predictions derived from individual reference species were integrated using GeMoMa v1.9, yielding 29,740 homology-supported gene models. For ab initio prediction, 3,000 high-confidence transcript-supported genes were selected to train AUGUSTUS v3.5.0^83^. The resulting species-specific parameters were then used for genome-wide gene prediction. Transcript-supported, homology-based, and ab initio predictions were integrated using EvidenceModeler v2.1.0.^84^. The preliminary gene set was further filtered using TransposonPSI to remove models associated with transposable elements. Gene models with coding sequences not divisible by three or containing internal stop codons were also removed. The final annotation comprised 21,200 protein-coding genes. PASA was subsequently used to refine UTR annotations based on transcriptome evidence.

Functional annotation of the predicted proteins was performed against the NR, Swiss-Prot, KEGG, KOG, and GO databases. NR, Swiss-Prot, KEGG, and KOG annotations were assigned by sequence-similarity searches, whereas protein domains and associated GO terms were identified using InterProScan v5.65-97.0^85^ with Pfam domain annotations. In total, 20,358 of the 21,200 predicted protein-coding genes (96.03%) received functional annotation from at least one database. Non-coding RNAs were annotated separately. tRNA genes were identified using tRNAscan-SE v2.0.12^86^, rRNA loci using RNAmmer v1.2^87^, and other structural non-coding RNAs with Infernal v1.1.4^88^ against the Rfam database.

#### CpG methylation profiling

CpG methylation was estimated independently from PacBio HiFi and ONT reads. For HiFi-based methylation calling, HiFi reads were aligned to the final T2T assembly with pbmm2 v1.13.0 (align --preset CCS --sort), and 5-methylcytosine (5mC) levels at CpG sites were called with pb-CpG-tools v2.3.2 using default parameters. For ONT-based analysis, ONT reads with base-modification tags were converted from BAM to modification-aware FASTQ with SAMtools fastq, aligned to the assembly with minimap2 v2.26 (-ayx map-ont), and methylation calls (5mC/5hmC/6mA) were extracted with modkit v0.4.1 using default parameters. For each chromosome, 5mC-methylated CpG sites were summarized by the total number of supporting reads and the corresponding methylation level (genome-wide means of approximately 76.1% for HiFi and 80.4% for ONT data), and genome-wide 5mC CpG methylation profiles were integrated with repeat, telomere and centromere annotations for chromosome-level visualization.

#### rDNA, SD and unique k-mer analysis

Ribosomal DNA loci were annotated with RNAmmer v1.2 using the eukaryotic rRNA models (-S euk -m lsu,ssu,tsu). Predicted 5S, 18S and 28S rRNA loci were intersected with read-depth profiles generated from PacBio HiFi, ONT and short-read alignments, and read depth across rDNA intervals was compared with the genome-wide average depth to assess copy representation and local assembly support; rDNA arrays were identified on chromosomes 2, 4 and 25. Segmental duplications were identified with BISER v1.4^89^ using default parameters (soft-masked genome, --kmer-size 21 in the vendor pipeline), and duplicated intervals were summarized by chromosome, length class and genomic distribution and visualized with circos v0.69-9^90^. Minimum unique k-mer (MUK) length was calculated with minUniqueKmer v1.0^91^ using default parameters, and unique k-mer profiles were summarized across the whole genome and in non-overlapping 100-kb windows along each chromosome to evaluate sequence uniqueness across chromosome arms and repeat-rich regions, including centromeric and subtelomeric intervals.

#### Hi-C interaction map construction

Clean Hi-C read pairs were aligned to the final assembly with BWA v0.7.17 (mem -5SP) and filtered with bam-filter v2.0.0 (1 -- nm 3). The genome was divided into 100-kb bins, and the number of Hi-C read pairs spanning each pair of bins was used as the interaction intensity to build chromosome-wide contact matrices with HapHiC and HiCExplorer v3.7.3 (hicFindRestSite, hicBuildMatrix, hicCorrectMatrix, hicPlotMatrix)^92^. Genome-wide and per-chromosome interaction heatmaps were generated to confirm the structural integrity of the pseudomolecules, showing clear intra-chromosomal interaction patterns without large-scale inter-chromosomal misjoins.

#### WGS variant processing and imputation

Whole-genome resequencing data were processed using the cattle reference genome ARS-UCD2.0 for read alignment and SNP calling. Because the downstream telomere analyses were anchored to the newly assembled Holstein_T2T genome, variant coordinates were subsequently harmonized to the Holstein_T2T coordinate system before genotype integration. Chromosome nomenclature was first standardized, after which variants called against ARS-UCD2.0 were lifted over to Holstein_T2T using Picard LiftoverVcf v3.0.0 and the corresponding ARS-UCD2.0-to-Holstein_T2T chain files. Variants that could not be unambiguously mapped to the Holstein_T2T assembly were excluded. Autosomal VCF files from different sequencing batches were then merged using bcftools v1.17^93^, such that all subsequent genotype processing and association analyses were performed using SNP coordinates defined on the Holstein_T2T assembly. The merged genotype dataset was imputed with Beagle v5.4^94^ using a published Holstein cattle haplotype reference panel. After imputation, allele frequencies were recalculated with bcftools, and autosomal variants with imputation quality DR2 ≥ 0.8 and minor allele frequency (MAF) ≥ 0.05 were retained. In total, 9,925,819 SNPs from 9,591 age-annotated Holstein cows were retained for genome-wide association analyses. Thus, although initial SNP calling was performed against ARS-UCD2.0, the final genotype dataset used for teloGWAS and all downstream locus-based analyses was represented in the Holstein_T2T coordinate system.

#### WGS-based telomere length estimation

Telomere length was estimated from WGS data using TelSeq^24^. Because the default TelSeq parameters were developed for the human genome, the source-code parameters were modified for the bovine genome, with TELOMERE_ENDS, READ_LENGTH and GENOME_LENGTH_AT_TEL_GC set to 60, 150 and 321,325,950, respectively. To reduce the influence of PCR duplicates on telomeric repeat counting, duplicated reads were removed using Picard MarkDuplicates with REMOVE_DUPLICATES=true. The duplicate-filtered BAM files were then used for TelSeq analysis with -k 12, such that reads containing at least 12 copies of the TTAGGG or CCCTAA telomeric repeat motif were counted as telomere-associated reads; the read length parameter was set to 150 bp. To assess consistency across WGS-based telomere length estimation approaches, a randomly selected subset of 57 samples was additionally analyzed using Computel v1.4^25^ with the Holstein_T2T assembly generated in this study as the reference genome; TelSeq and Computel estimates were highly correlated (r = 0.965, P < 0.001). To assess the sensitivity of TelSeq estimates to sequencing depth, duplicate-removed BAM files from 10 samples were downsampled to a range of sequencing depths, and telomere length was re-estimated using the same TelSeq procedure.

#### Coverage-based telomere length correction

WGS-based telomere length estimates can be affected by systematic technical variation related to sequencing depth, sequencing batch, mappability and local copy-number differences. To account for these effects, a coverage-based NGS-PCA approach was used to derive sequencing-depth-related principal components (hereafter bPCs), which were then used to correct TelSeq-derived telomere length estimates^103^. Bovine autosomes were divided into non-overlapping 1-kb genomic bins. Low-mappability regions were identified with GEM mappability^95^ using k = 50, and bins with mappability < 1.0 were excluded; structural variant regions obtained from Ensembl were also removed. Because the original BAM files were aligned to either ARS-UCD1.2 or ARS-UCD2.0, clean bins were generated separately for each reference version, and clean bins from ARS-UCD2.0 were converted to ARS-UCD1.2 using CrossMap v0.7.3^96^. Only 1-kb clean bins with one-to-one correspondence between the two reference versions were retained. Mean sequencing depth within each clean bin was calculated for each sample using mosdepth^97^, and the resulting bin-by-sample coverage matrix was processed with NGS-PCA for normalization and principal component analysis. The number of bPCs used for correction was evaluated separately in the full cohort, the high-depth subgroup and the low-depth subgroup: for a given number of bPCs k, raw TelSeq-estimated telomere length was regressed on the first k bPCs (rawTL ∼ bPC1 + … + bPCk), and the Pearson correlation between the resulting residuals and age was calculated to evaluate the correction. Based on this procedure, 121, 5 and 131 bPCs were selected for the full cohort (n = 9,591), the high-depth subgroup (n = 1,498) and the low-depth subgroup (n = 8,093), respectively, and residuals from the corresponding linear models were used as bPC-corrected telomere length phenotypes. For the rumen WGS data, the same clean-bin and NGS-PCA procedure was applied, and eleven bPCs were selected for correction.

#### Telomere length GWAS

Telomere length GWAS was performed using the two-step KVIK approach implemented in LDAK^98^. In the first step, variants used for constructing the polygenic prediction model were filtered with PLINK2 v2.0.0-a.6.9LM^99^ using --maf 0.05 --mac 100 -- geno 0.1 --hwe 1e-15 --indep-pairwise 50 5 0.3, retaining 539,986 relatively independent SNPs. In the second step, single-variant association testing was performed for the 9,925,819 quality-controlled autosomal SNPs. The primary GWAS phenotype was the 121-bPC-corrected telomere length residual from all 9,591 individuals. Age and the first 20 genotype principal components (gPCs) were included as continuous covariates, and sequencing batch was included as a categorical fixed effect. A Bonferroni-corrected threshold of P < 5.037 × 10−9 was used to define genome-wide significance, with P = 5 × 10−8 shown as a reference threshold. For each major association region, the variant with the lowest association P value was defined as the lead SNP.

#### Conditional association analysis

Single-SNP conditional association analyses were performed to evaluate whether additional signals independent of the lead SNPs were present within the major GWAS regions. The lead variants chr13_53727328, chr19_27855458 and chr20_71539969 were used as conditioning SNPs. Allele dosages for each lead SNP were extracted from the PLINK genotype dataset and aligned according to the GWAS effect allele, and each lead SNP dosage was included as an additional continuous covariate together with age, the first 20 gPCs and sequencing batch. Genome-wide association testing was repeated using the same LDAK-KVIK framework as in the primary GWAS, and conditional association results within ± 1 Mb of each lead SNP were used to determine whether residual association signals remained after conditioning on the corresponding lead variant.

#### eQTL and sQTL colocalization

Colocalization analyses between telomere length GWAS summary statistics and multi-tissue cis-eQTL and sQTL data from CattleGTEx were performed using the coloc.abf function in the R package coloc v5.2.3^102^. For each of the three major GWAS lead SNPs, a 2-Mb colocalization window was defined by extending 1 Mb upstream and downstream of the lead SNP. Before colocalization, variants shared between the GWAS and QTL datasets were allele-harmonized and classified as directly matched, allele-swapped, strand-consistent or strand-swapped; QTL effect directions were adjusted accordingly, and variants that could not be unambiguously harmonized were removed. MAF values were compared between datasets, and variants with |ΔMAF| > 0.2 were excluded. After allele harmonization and MAF filtering, a QTL signal was retained only if it shared at least 50 SNPs with the GWAS dataset. The posterior probability PPH4 was used to quantify support for a shared causal variant, and PPH4 > 0.7 was considered evidence of colocalization. The same framework was applied to sQTL data using individual splicing events as the unit of analysis, and splicing event IDs were mapped to genes to identify candidate genes and splicing events potentially affected by regulatory variation at telomere length loci.

#### Multi-tissue TWAS

Multi-tissue transcriptome-wide association analysis (TWAS) was performed using the FarmGTEx TWAS server (https://twas.farmgtex.org/; accessed July 2026) to evaluate associations between genetically predicted gene expression and telomere length. The Imputation model and MetaXcan elastic-net method were selected, using the platform-provided ARS-UCD1.2/bosTau9 prediction models. GWAS summary statistics were used as input, and all 23 available tissues were analyzed. Gene–tissue associations with FDR < 0.05 were considered significant. To integrate TWAS and GWAS results, GWAS signals within ± 500 kb of each TWAS gene were examined, and a TWAS-significant gene was considered to have local GWAS support if at least one variant with P < 5 × 10−8 was present within this interval. For each tissue, the number of TWAS-significant genes and the number of significant TWAS genes with local GWAS support were summarized; at the gene level, the minimum FDR across tissues, the minimum GWAS P value within ± 500 kb and the number of tissues with TWAS FDR < 0.05 were calculated.

#### RNA-seq processing and cell correction

RNA-seq data from 317 rumen samples and 289 whole-blood samples were used for gene expression analysis, WGCNA and connectivity dynamics analysis. Gene expression was quantified as transcripts per million (TPM) and transformed as log2(TPM + 1), and genes with TPM ≥ 0.1 in at least 20% of samples were retained. For whole-blood samples, marker-based cell-type expression scores were constructed to account for variation in blood cell composition among individuals. Six cell categories were considered: T cells (CD3D, CD3E, CD3G, CD247, ZAP70, CD2, TRAC, LCK), B cells (MS4A1, CD79A, CD79B, CD19, BLNK, PAX5, CD22, BANK1), NK cells (NCR1, KLRD1, NKG7, GNLY, CTSW, PRF1, IL2RB), monocytes (CD14, S100A12, ADGRE1, HCK, AIF1, CSF1R, CST3, FCGR3A, LST1), platelets/megakaryocytes (PPBP, PF4, GP9, GP1BA, GP1BB, ITGA2B, ITGB3, TUBB1, TREML1) and erythrocytes (AHSP, HBB, HBM, HBA, ALAS2, SLC4A1, KLF1, EPB42). For each marker gene, expression values were Z-score standardized across the 289 whole-blood samples, and for each cell type the mean Z-score across all available marker genes was used as the cell-type expression score, requiring at least three valid marker genes per cell type. To reduce confounding from cell-composition differences, the six marker-based cell-type expression scores were included as covariates in a multiple linear regression model for each gene’s log2(TPM + 1) expression value, and residuals were used as cell-composition-corrected expression values for downstream whole-blood transcriptomic analyses. Before WGCNA, whole-blood telomere length was further regressed on age and mean sequencing depth, and the resulting residuals were used for downstream analyses.

#### Weighted gene co-expression analysis

Weighted gene co-expression network analysis (WGCNA) was performed separately for the rumen and whole-blood transcriptomes using the R package WGCNA v1.74^100^. For genes passing the initial expression filter, the median absolute deviation (MAD) was calculated across samples, and the top 75% most variable genes were retained for PCA, sample hierarchical clustering and network construction. Soft-thresholding powers were selected according to scale-free topology fit and mean connectivity across candidate powers, using signed R² = 0.85 as a reference; final powers of 5 and 3 were used for rumen and whole blood, respectively. Pearson correlations were then calculated between each module eigengene and the corresponding telomere length phenotype: for rumen, the 11-bPC-corrected telomere length residual was used; for whole blood, the telomere length residual additionally adjusted for age and mean sequencing depth was used.

#### Sliding-window connectivity analysis

To characterize continuous changes in gene co-expression connectivity across the telomere length gradient, a telomere length-stratified sliding-window analysis was performed in both tissues. Unlike the MAD-based filtering used for global WGCNA, this analysis retained the top 30% of genes with the highest expression variance across all samples. For whole blood, high-variance genes were selected from the uncorrected log2(TPM + 1) expression matrix, after which the six marker-based cell-type expression scores were regressed out and residuals were used for connectivity analysis. Samples were ordered from high to low according to the corrected telomere length residual, and a fixed sliding window covering approximately 25% of samples was applied with a step size of one sample: for the 317 rumen samples, a window size of 79 samples generated 239 windows, and for the 289 whole-blood samples, a window size of 72 samples generated 218 windows. Within each window, gene connectivity was calculated using the softConnectivity function in WGCNA with the same soft-thresholding powers as in the global network analysis (power = 5 for rumen and power = 3 for whole blood), producing a connectivity trajectory along the telomere length gradient for each candidate gene. Connectivity values for each gene were Z-score standardized across all sliding windows. The elbow method was used to evaluate clustering performance for K = 2 to 15, and K-means clustering was then performed with K = 6, nstart = 100 and a maximum of 500 iterations. GO enrichment analysis for relevant modules and high-connectivity gene clusters was performed using clusterProfiler^101^, with P < 0.05 considered statistically significant.

### QUANTIFICATION AND STATISTICAL ANALYSIS

Assembly quality was quantified with descriptive metrics, and no randomization, blinding or sample-size estimation was applicable for the reference genome, as all genomic data were generated from a single donor individual. Genome completeness is reported as the number and percentage of complete, single-copy, duplicated, fragmented and missing genes among the n = 9,226 mammalia_odb10 BUSCO markers (compleasm v0.2.5). Consensus accuracy is reported as k-mer-based QV (Merqury v1.3/merfin v1.0, k = 21). Read support is reported as the mapping rate (number of aligned reads divided by the total number of input reads; SAMtools v1.18 stats), with n representing the number of reads of each data type as listed in the Method details. Single-base- and structural-level accuracy is reported as CRAQ covered rate, low-confidence rate, R-AQI and S-AQI. Repeat content is reported as the total length and percentage of the assembly occupied by each repeat class, and CpG methylation is reported as the proportion of methylated CpG sites per chromosome and averaged genome-wide. Statistical tests used for each analysis are described in the corresponding Method details and figure legends. Unless otherwise specified, correlations were calculated using Pearson or Spearman correlation as indicated, and paired comparisons between chromosome ends (e.g., 5p versus 3p telomere lengths across the 30 chromosomes) used Wilcoxon signed-rank tests. Enrichment analyses used the statistical framework implemented in clusterProfiler, and multiple testing was controlled using FDR where indicated. GWAS significance was assessed using a Bonferroni-corrected threshold of P < 5.037 × 10−9, with P = 5 × 10−8 shown as a reference threshold; for GWAS, n represents the number of individuals (9,591 for the primary analysis) and the number of tested SNPs (9,925,819) as stated in the Method details. eQTL and sQTL colocalization was summarized by coloc PPH4, with PPH4 > 0.7 interpreted as evidence supporting a shared causal variant. TWAS gene–tissue associations with FDR < 0.05 were considered significant. WGCNA module–trait relationships were evaluated by Pearson correlation between module eigengenes and telomere length phenotypes. PERMANOVA was used to test chromosome-end effects in motif-density ordination analyses where indicated. The agreement between TelSeq- and Computel-based telomere length estimates (n = 57 samples) was evaluated by Pearson correlation. No data or subjects were excluded from the reported analyses except the explicitly described quality-control filters stated in the Method details.

### SUPPLEMENTAL FIGURES

**Figure S1.**
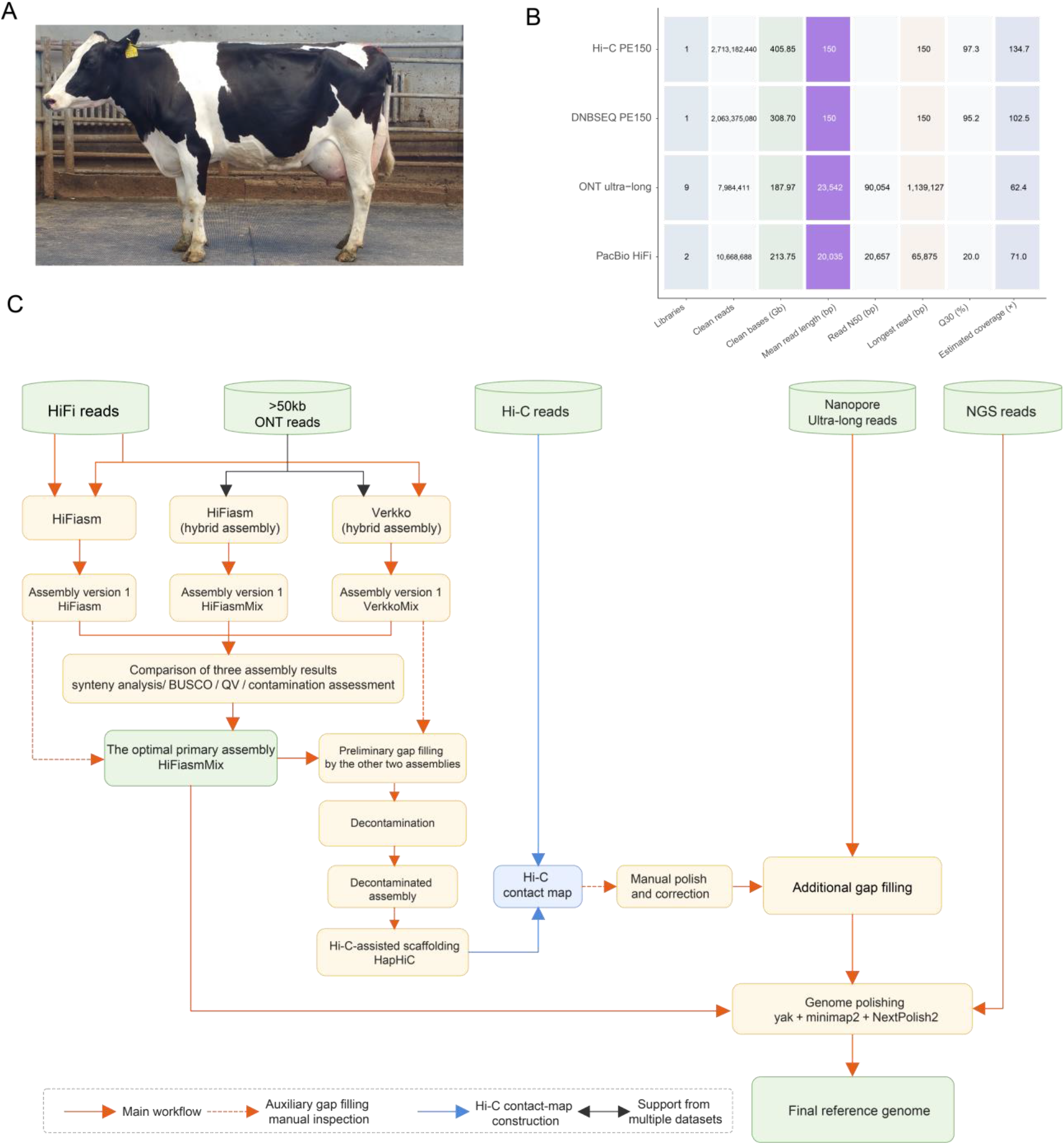
Study sample and overview of genome assembly. (A) Photograph of the donor Holstein cow, a seven-year-old female used for Holstein_T2T assembly. (B) Summary of sequencing datasets, including read number, sequencing yield, read N50, and estimated genome coverage across platforms. (C) Integrated Holstein_T2T assembly workflow using PacBio HiFi, ONT ultra-long, and Hi-C sequencing data.

**Figure S2.**
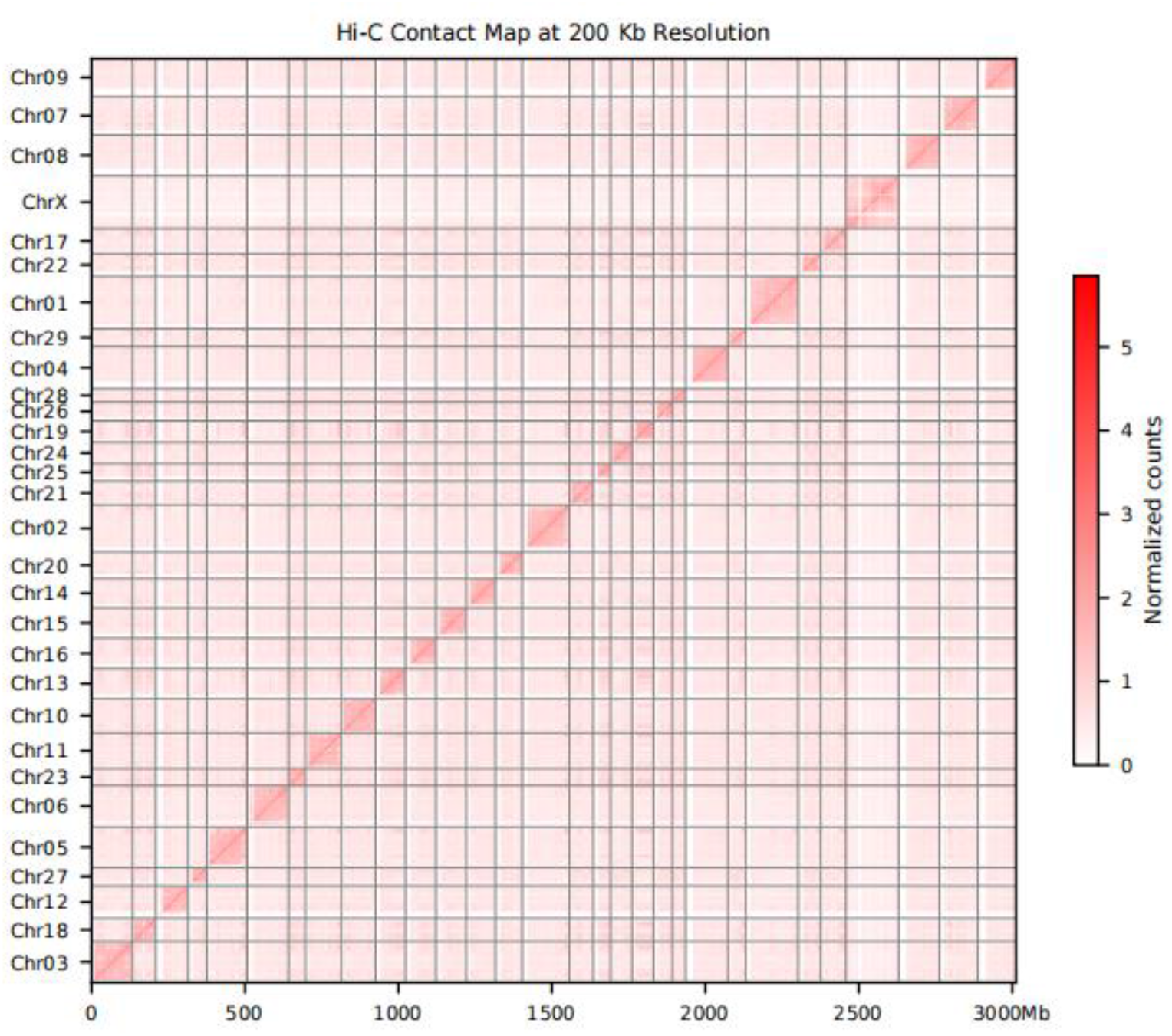
Hi-C chromatin interaction map supporting the structural accuracy of the assembly. The genome-wide Hi-C contact matrix shows clear chromosome-level intrachromosomal interaction patterns, with no evidence of large-scale interchromosomal misjoins.

**Figure S3.**
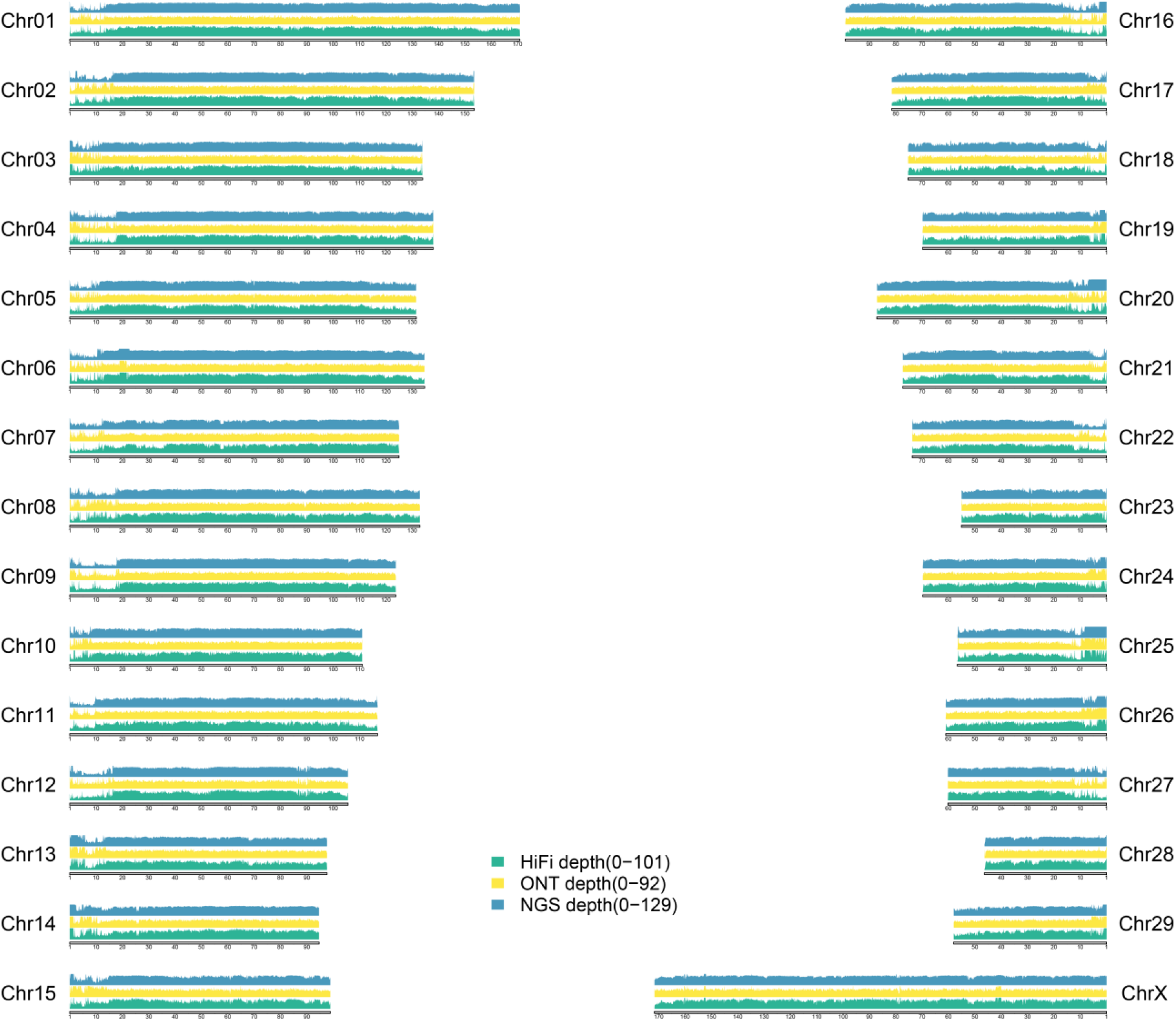
Sequencing-read coverage across the Holstein_T2T assembly. Genome-wide coverage profiles of PacBio HiFi and ONT reads across chromosomes, supporting the high continuity and sequence accuracy of the Holstein_T2T assembly.

**Figure S4.**
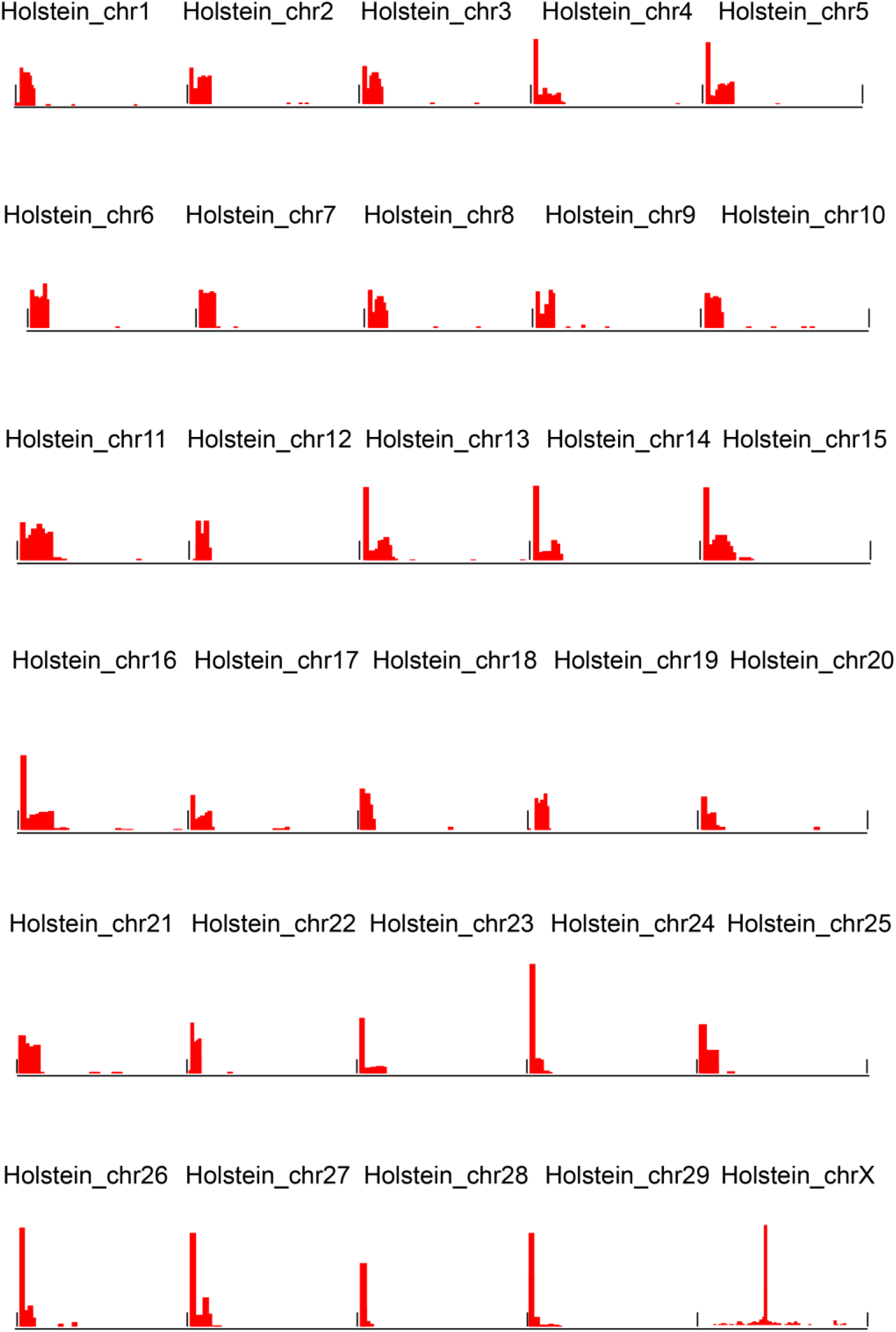
CENP-A CUT&Tag signal profiles. Genome-wide CENP-A CUT&Tag enrichment profiles used to identify and validate centromeric regions in the Holstein_T2T.

**Figure S5.**
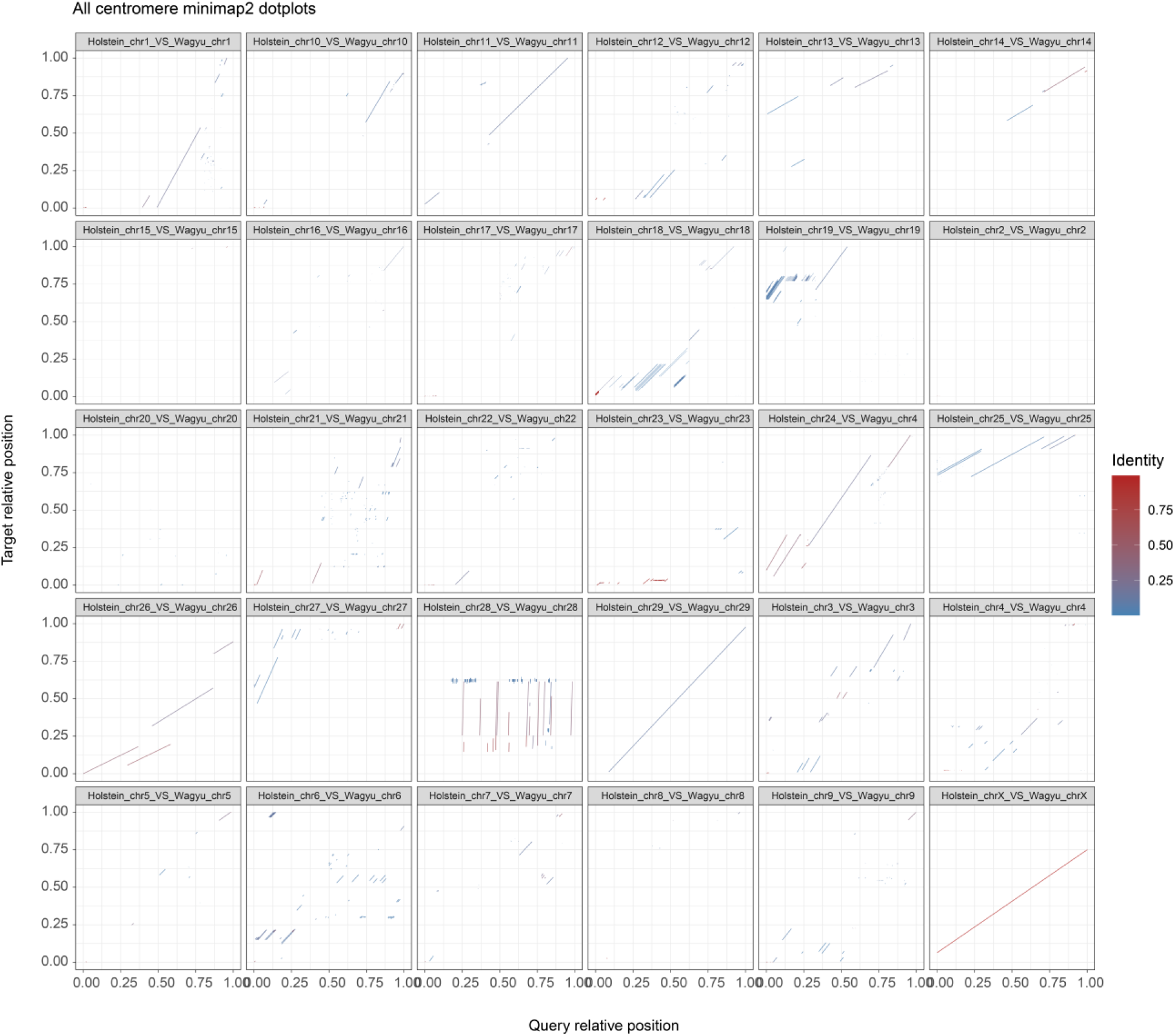
Comparative alignment of homologous centromeric regions between the Holstein_T2T and the Wagyu reference genome (UOA_Wagyu_1) by minimap2. Chromosome-by-chromosome dot plots of homologous centromeric intervals between the Holstein_T2T assembly and the Wagyu reference genome. Line color indicates local sequence identity.

**Figure S6.**
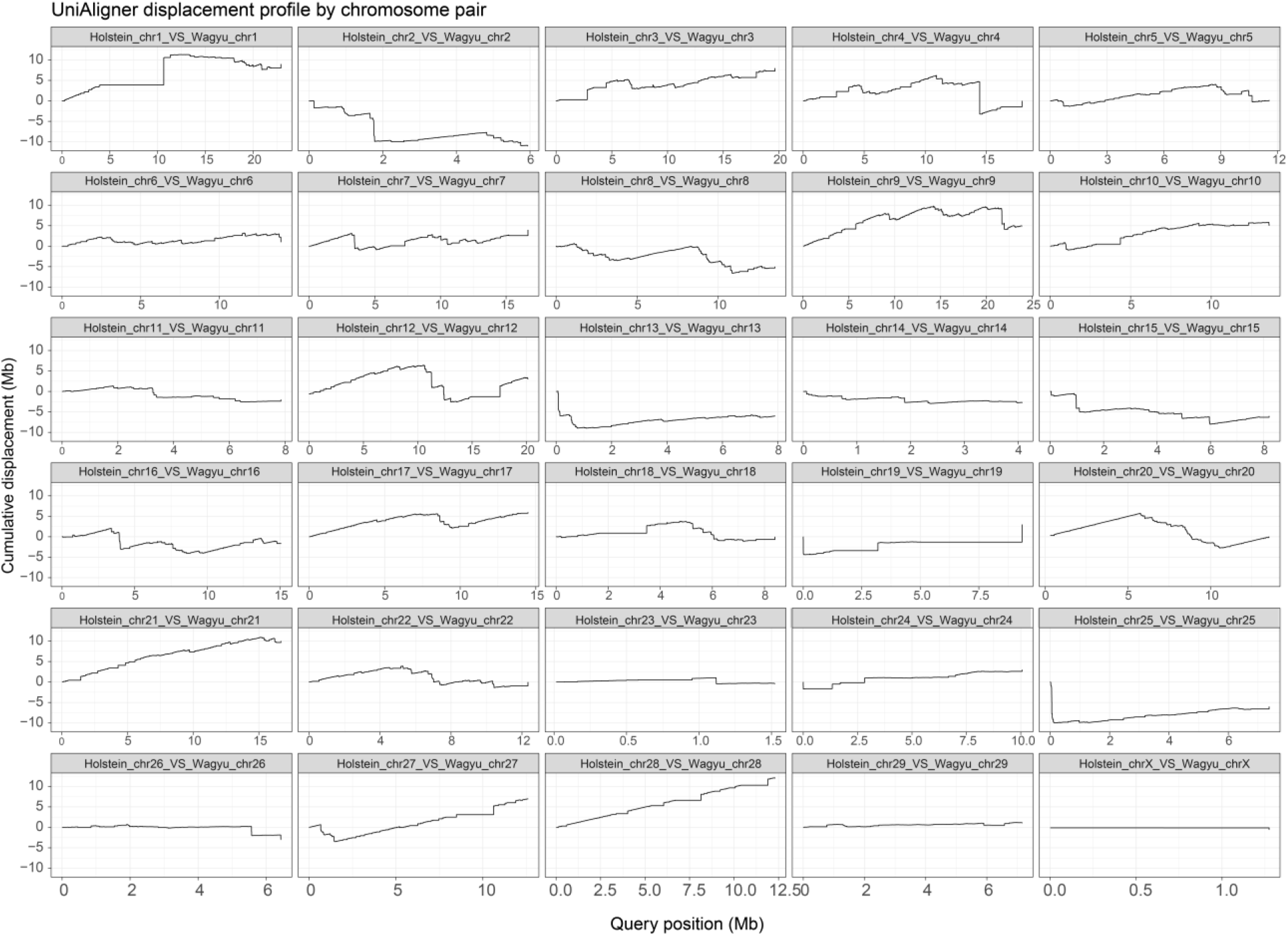
Comparative alignment of homologous centromeric regions between the Holstein_T2T and the Wagyu reference genome (UOA_Wagyu_1) by UniAligner. Sequence-divergence profiles across the corresponding homologous centromeric intervals, showing local variation in alignment structure and sequence conservation.

**Figure S7.**
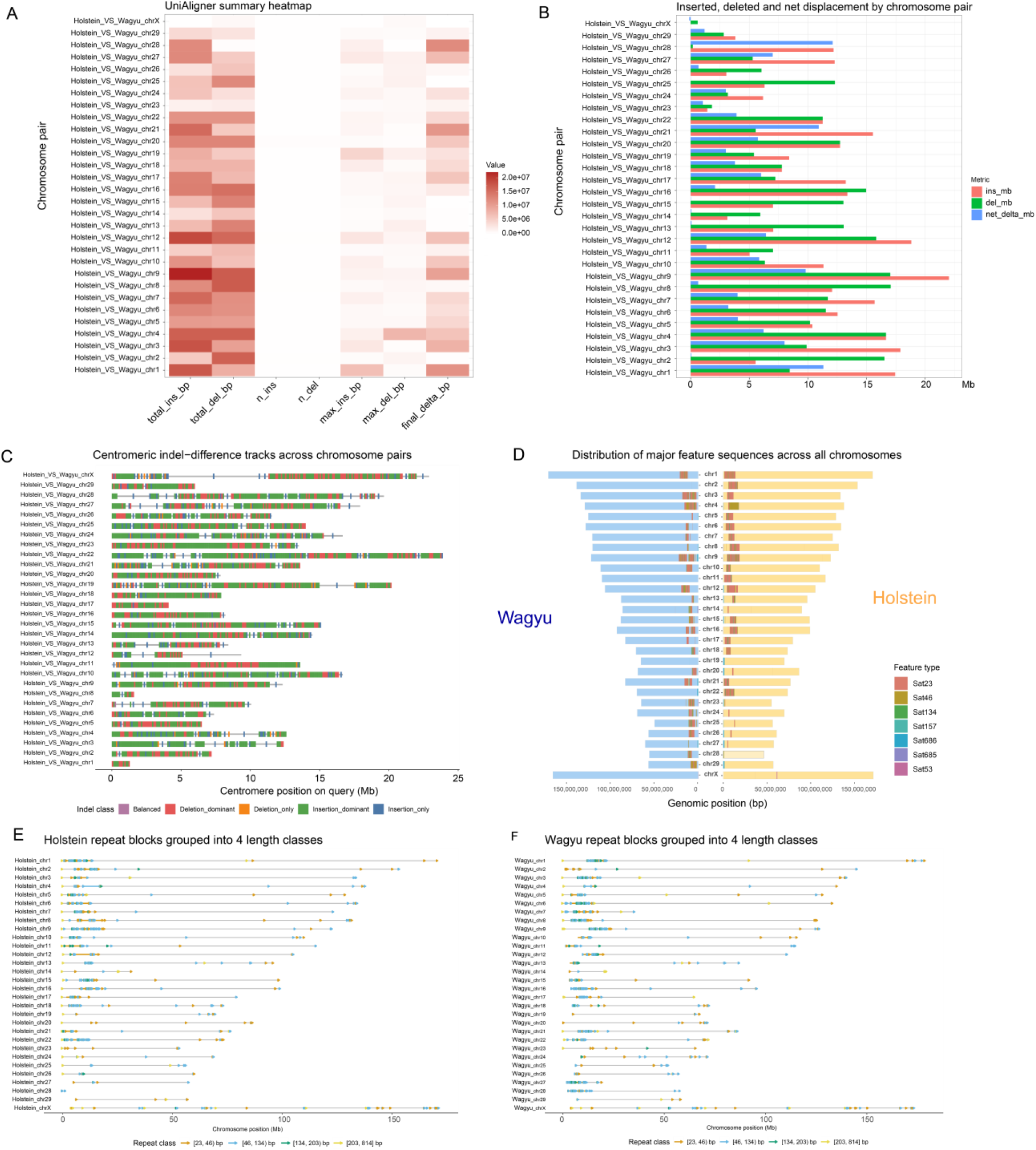
Structural variation and repeat-organization differences across homologous bovine centromeres. (A) Heatmap summarizing insertion and deletion burden across homologous centromeric intervals between the Holstein_T2T and the Wagyu reference genome. Columns represent indel-related metrics, including total insertion and deletion length, event number, maximum event size, and net sequence-length change. (B) Chromosome-level comparison of centromeric indel profiles, showing the relative contribution of insertion, deletion, and net length-change signals across homologous intervals. (C) Local classification of indel-dominated segments along each aligned centromeric interval. Colors indicate balanced, deletion-dominant, deletion-only, insertion-dominant, and insertion-only regions. (D) Comparison of centromeric interval length and repeat composition between the Wagyu reference genome and the Holstein_T2T assembly. Mirrored bars show homologous intervals in the two genomes, with colors indicating repeat categories. (E and F) Distribution of major centromeric repeat units along Holstein (E) and Wagyu (F) centromeric intervals. Colored points mark repeats units or repeat families detected within each chromosome-specific centromeric region, highlighting differences in repeat abundance and organization between the two genomes.

**Figure S8.**
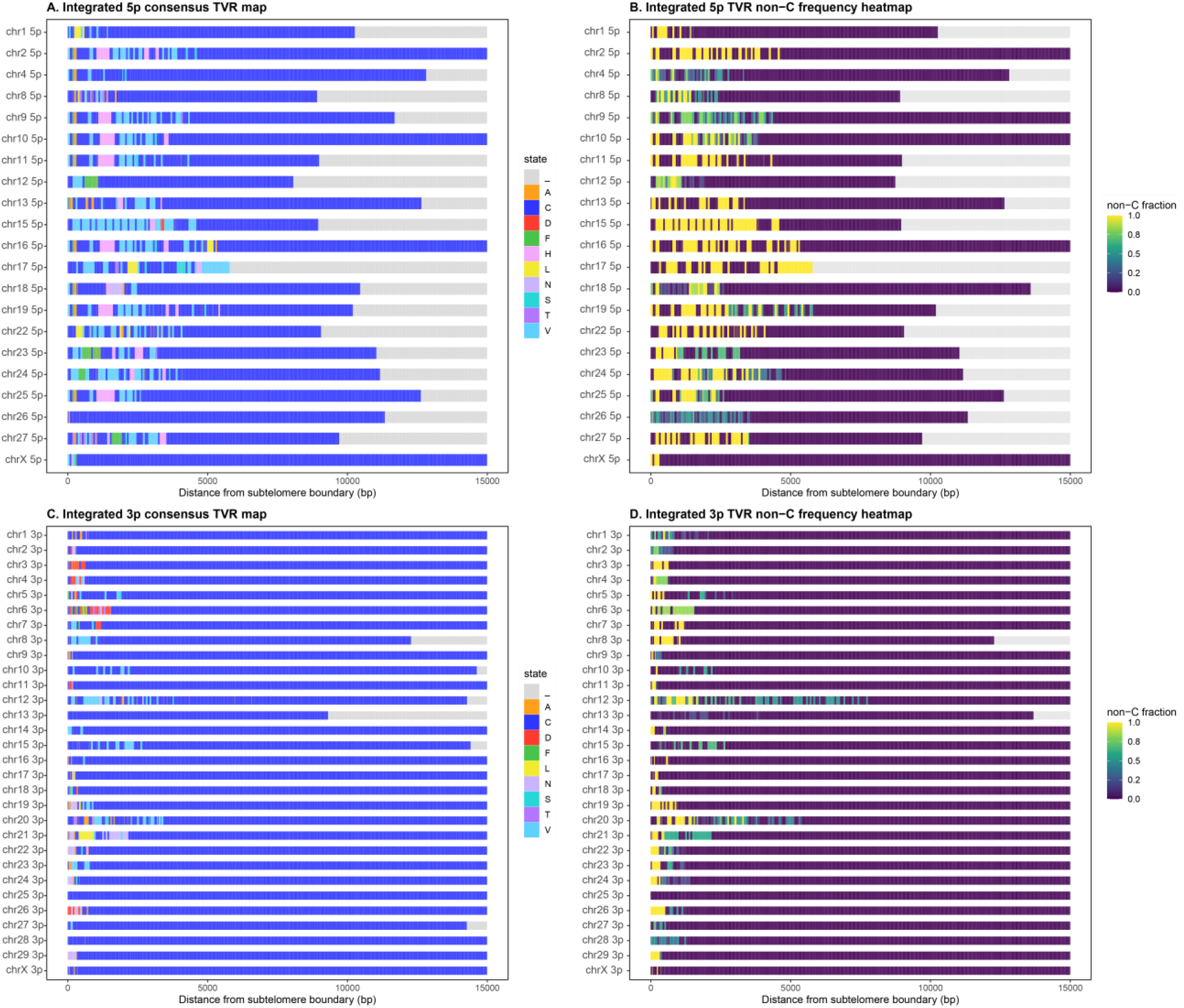
Chromosome-end-specific distribution of telomere variant repeats and non-canonical repeat content. (A) Base-state profiles of terminal telomeric repeat arrays at 5p chromosome ends. Rows represent individual chromosome ends, and columns indicate positions along the terminal repeat tract. Colors denote canonical and non-canonical repeat states, with gray indicating unassigned or gap regions. (B) Distribution of non-canonical repeat fraction across the corresponding 5p telomeric regions. Color intensity indicates the local fraction of non-canonical sequence (C) Base-state profiles of terminal telomeric repeat arrays at 3p chromosome ends, shown as in (A). (D) Distribution of non-canonical repeat fraction across the corresponding 3p telomeric regions, shown as in (B).

**Figure S9.**
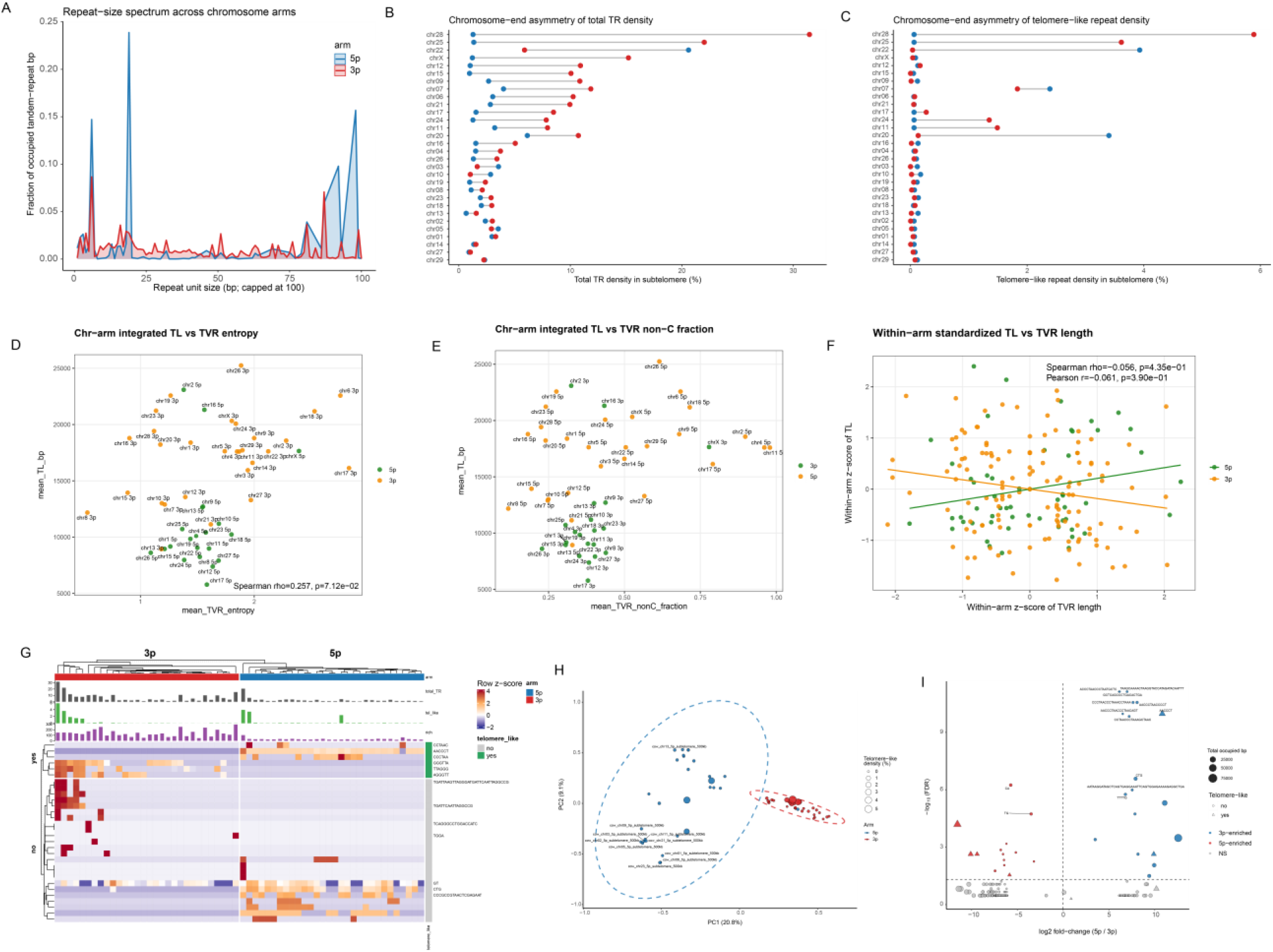
Subtelomeric tandem-repeat architecture and motif composition at chromosome ends. (A) Comparison of repeat-unit distribution patterns between 5p and 3p chromosome ends. (B) Proportion of tandem repeats at paired chromosome ends across the 30 chromosomes; 25 chromosomes show higher tandem-repeat content at 3p termini. (C) Distribution of telomere-like motif density across chromosome ends. (D–F) Correlations between telomere length and TVR abundance or non-canonical sequence composition, shown globally and by chromosome-end class. (G) Motif-abundance-based clustering of subtelomeric regions, separating 3p and 5p termini and identifying chromosome-arm-enriched motifs. (H) Principal-component analysis of motif density across subtelomeric regions (PERMANOVA, *R*² = 0.178, *P* = 0.001). (I) Differentially enriched motifs between 5p and 3p subtelomeric regions.

**Figure S10.**
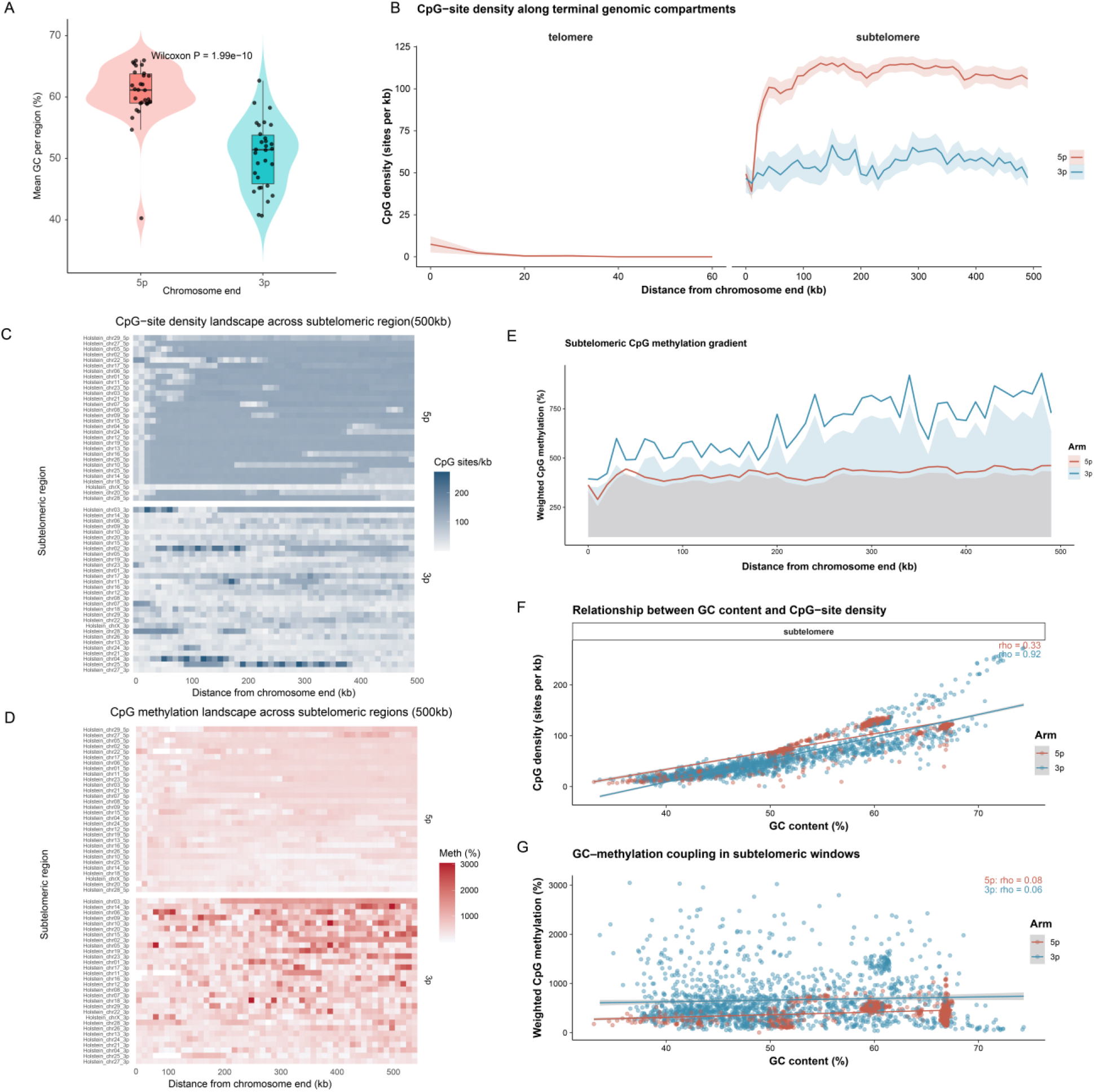
Subtelomeric GC/CpG composition and DNA methylation at chromosome ends. (A) Comparison of mean GC content between 5p and 3p chromosome ends across subtelomeric regions. Points represent individual chromosome ends, with distributions shown as violin and box plots. (B) Distance-resolved CpG-site density across terminal genomic compartments. CpG density is shown separately for telomeric repeat tracts and the adjacent 500-kb subtelomeric regions at 5p and 3p chromosome ends. Lines indicate mean profiles and shaded regions indicate variability across chromosome ends. (C) Chromosome-end-resolved heatmaps of CpG-site density across the 500-kb subtelomeric regions at 5p and 3p termini. Each row represents an individual chromosome end. (D) Chromosome-end-resolved heatmaps of CpG methylation across the 500-kb subtelomeric regions at 5p and 3p termini. Each row represents an individual chromosome end. (E) Distance-resolved profiles of weighted CpG methylation across the 500-kb subtelomeric regions at 5p and 3p chromosome ends. Lines show chromosome-arm-specific profiles, with shaded regions indicating variation across chromosome ends. (F) Relationship between GC content and CpG-site density across subtelomeric windows, shown separately for 5p and 3p chromosome ends. Lines indicate fitted trends; Spearman correlation coefficients are shown in the panel. (G) Relationship between GC content and weighted CpG methylation across subtelomeric windows at 5p and 3p chromosome ends. Lines indicate fitted trends; Spearman correlation coefficients are shown in the panel.

**Figure S11.**
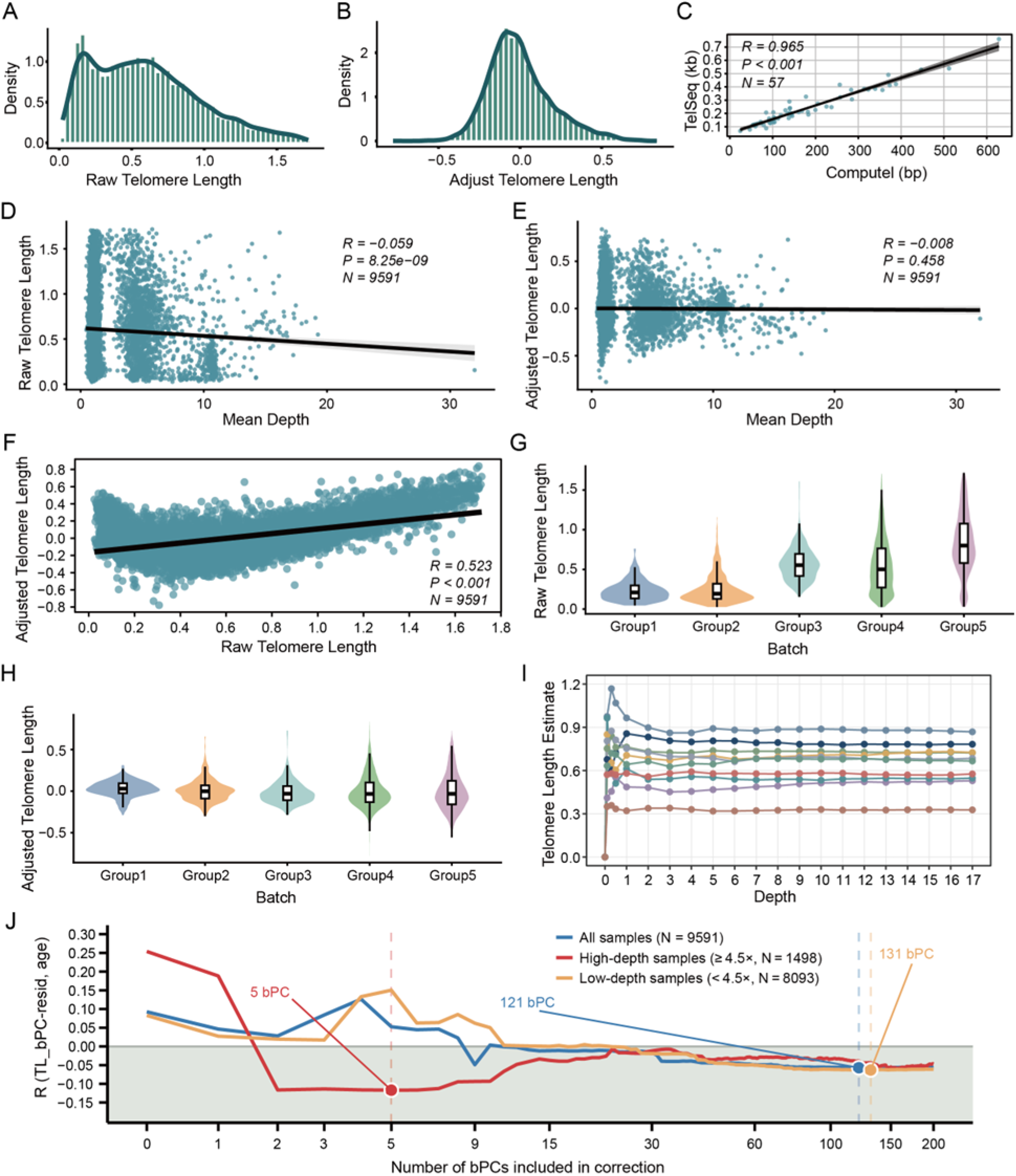
Telomere length phenotype correction and quality control. (A and B) Distribution of telomere length estimates before and after bPC correction. (B) Independent validation of sequencing-based telomere length estimates using Computel. (D and E) Correlations between telomere length and mean sequencing depth before and after correction. (F) Correlation between raw and corrected telomere length estimates (*r* = 0.523). (G–J) Evaluation of batch effects, sequencing-depth sensitivity, and correction performance of coverage-derived batch principal components (bPCs).

**Figure S12.**
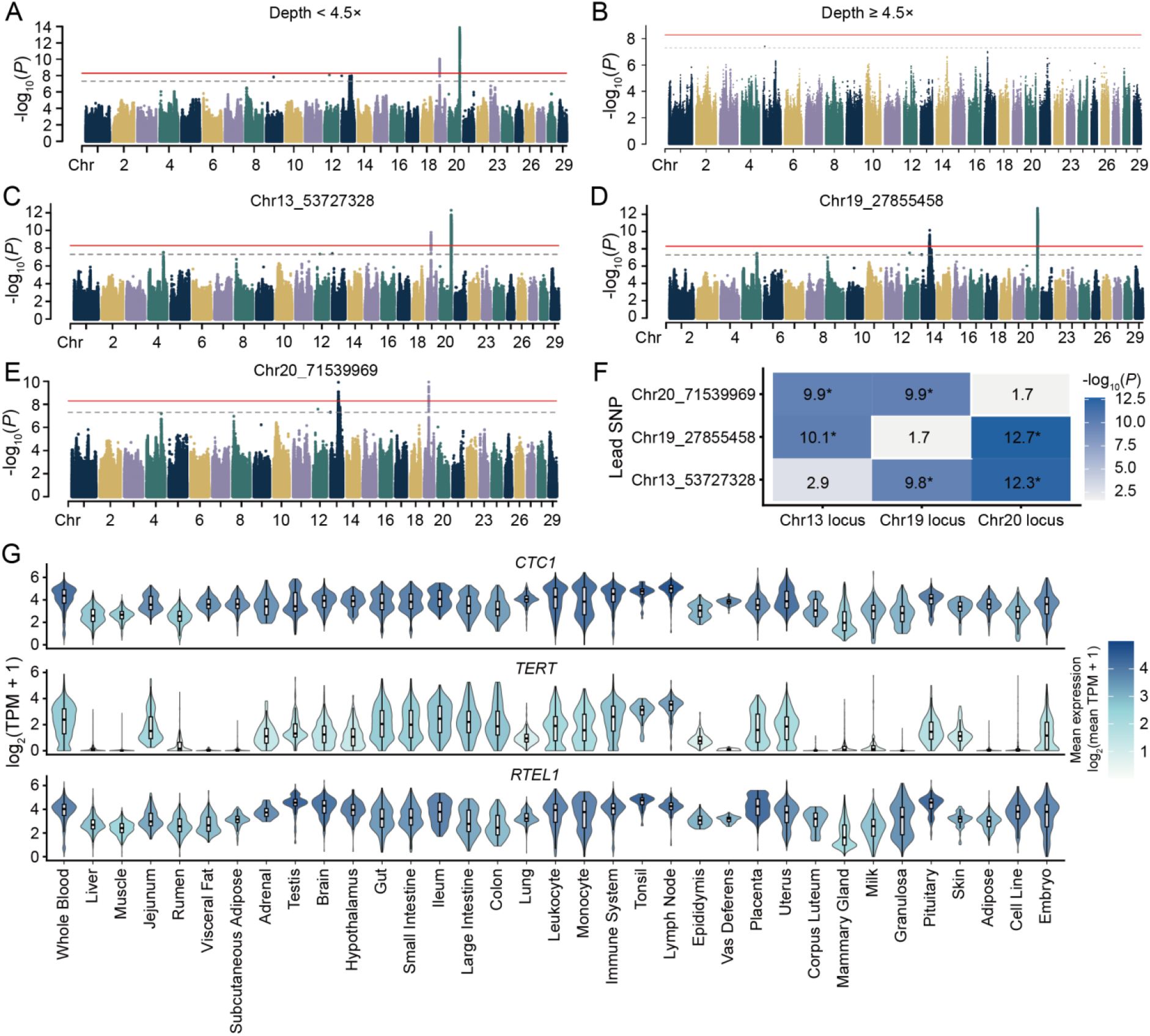
Sequencing-depth sensitivity, conditional GWAS, and tissue expression of telomere-maintenance genes. (A and B) Telomere length GWAS stratified by genome-wide mean sequencing depth in low-depth (<4.5×; A) and high-depth (≥4.5×; B) samples. The red solid line indicates the Bonferroni-corrected genome-wide significance threshold (*P* = 5.037 × 10^−9^), and the gray dashed line indicates the reference threshold (*P* = 5 × 10^−8^). (C–E) Conditional GWAS performed using chr13_53727328 (C), chr19_27855458 (D), and chr20_71539969 (E) as conditioning variants. Conditioning attenuated the corresponding local association peak while retaining signals at the other major loci. (F) Residual association strength across the three major GWAS loci after conditional analysis. Rows indicate the conditioning lead SNP, columns indicate the chromosome 13, 19, and 20 loci, and values denote the maximum residual −log10(*P*) within ±1 Mb of each locus. Asterisks indicate residual signals with *P* < 5 × 10^−8^. (G) Tissue expression profiles of *CTC1*, *TERT*, and *RTEL1* across cattle tissues. Violin plots show log2(TPM + 1) expression levels, highlighting broad expression of *CTC1* and *RTEL1* and low, tissue-restricted expression of *TERT*.

**Figure S13.**
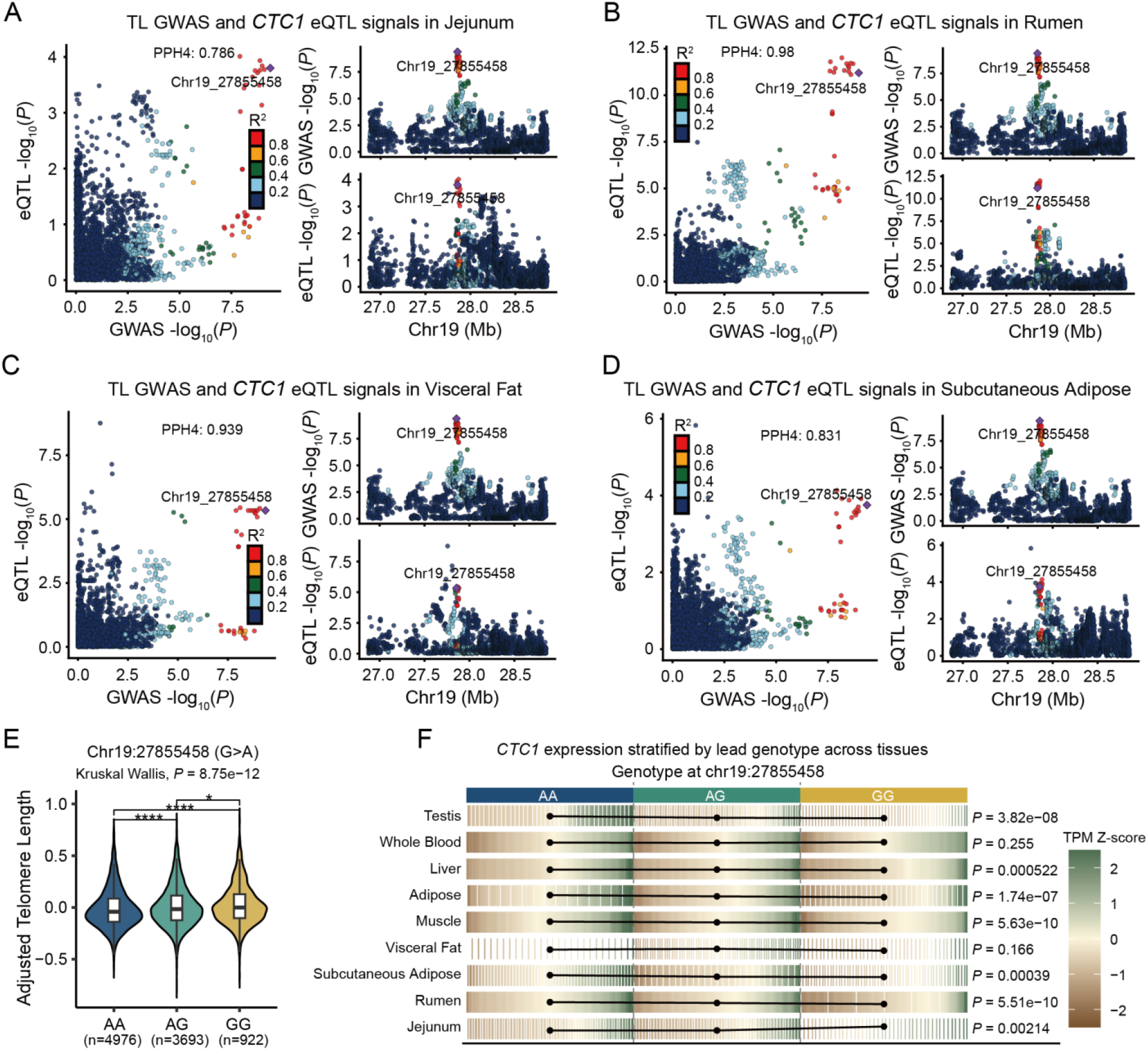
Multi-tissue eQTL colocalization at the *CTC1* locus and genotype effects of the lead variant. (A–D) Colocalization of telomere length (TL) GWAS signals with *CTC1* eQTL signals in jejunum (PPH4 = 0.786), rumen (PPH4 = 0.980), visceral fat (PPH4 = 0.939), and subcutaneous adipose tissue (PPH4 = 0.831). Left panels compare GWAS and eQTL association signals for regional variants, with point color indicating linkage disequilibrium (*R*²) with the lead variant chr19_27855458. Right panels show regional association profiles for TL GWAS and *CTC1* eQTL signals across the chromosome 19 locus. PPH4 denotes the posterior probability that the GWAS and eQTL signals share a causal variant. (E) Distribution of corrected TL across chr19_27855458 genotypes. Sample sizes for AA, AG, and GG genotypes are 4,976, 3,693, and 922, respectively. Overall genotype effects were tested using the Kruskal–Wallis test (*P* = 8.75 × 10^−12^), with asterisks indicating significant pairwise comparisons. (F) Genotype-dependent *CTC1* expression across tissues. Normalized expression levels are shown as a heatmap, with statistical significance for genotype–expression associations indicated on the right, supporting regulatory effects of chr19_27855458 on *CTC1* expression in multiple tissues.

**Figure S14.**
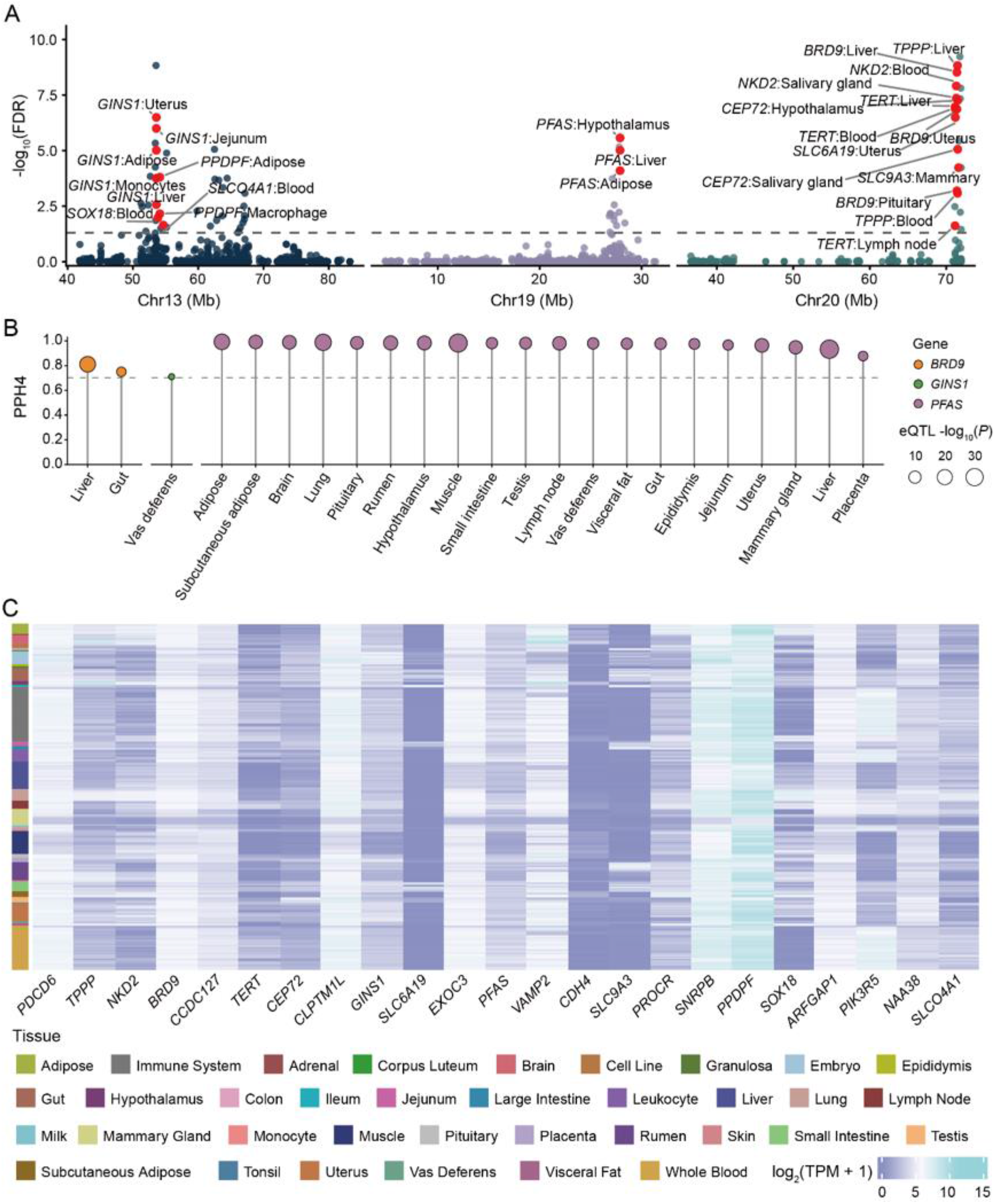
Multi-tissue colocalization, candidate-gene expression, and TWAS integration. (A) Multi-tissue TWAS results for the major telomere length-associated regions on chromosomes 13, 19, and 20. Each point represents a gene–tissue TWAS association, with genomic position on the x axis and TWAS significance on the y axis. The gray dashed line indicates the FDR = 0.05 threshold. Significant candidate gene–tissue signals are highlighted in red and labeled, including *GINS1* and neighboring genes on chromosome 13, *PFAS* on chromosome 19, and *TERT*, *BRD9*, *CEP72*, *NKD2*, and *TPPP* on chromosome 20. (B) eQTL–teloGWAS colocalization for selected candidate genes, including *BRD9*, *GINS1*, and *PFAS*, across tissues. The y axis shows PPH4, the dashed line indicates the reference threshold for strong colocalization evidence, point color denotes candidate gene, and point size reflects eQTL association strength. (C) Tissue expression profiles of teloGWAS- and TWAS-supported candidate genes. Columns represent candidate genes, rows represent transcriptome samples from different cattle tissues, the left annotation bar indicates tissue category, and heatmap color denotes expression level.

**Figure S15.**
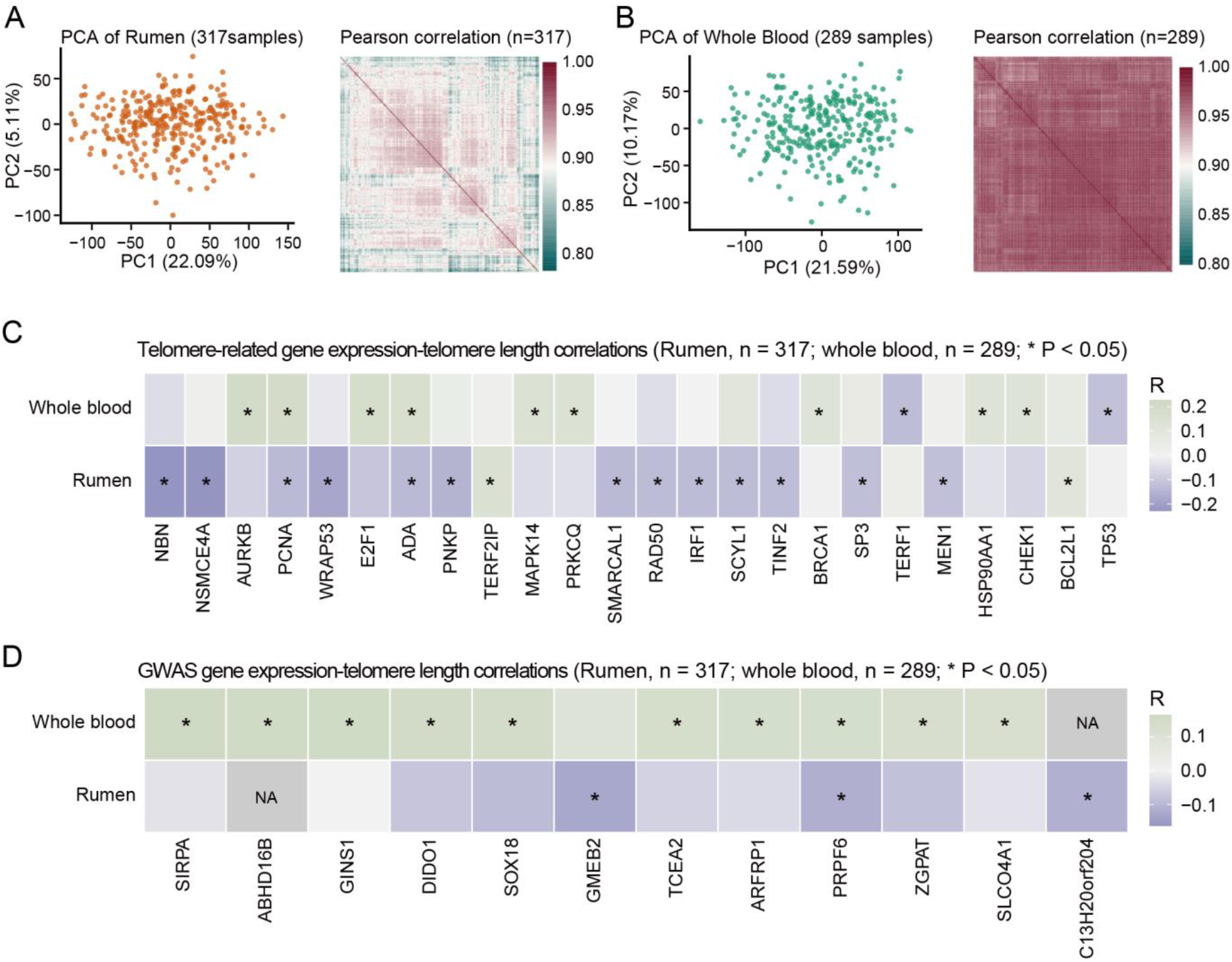
Expression–telomere length associations for telomere-related genes and GWAS candidates in rumen and whole blood. (A) Transcriptome quality assessment for 317 rumen samples. Principal-component analysis (PCA) of the gene expression matrix and inter-sample Pearson correlation heatmap were used to evaluate sample clustering and expression-data consistency. (B) Transcriptome quality assessment for 289 whole-blood samples, shown as in (A). (C) Correlations between expression levels of known telomere-maintenance genes and telomere length. Colors indicate Pearson correlation coefficients, and asterisks denote significant correlations (*P* < 0.05). Differences between rumen and whole blood indicate tissue-specific telomere-related transcriptional associations. (D) Correlations between expression levels of teloGWAS candidate genes and telomere length. Colors and asterisks are defined as in (C), highlighting tissue-dependent expression–TL associations among GWAS candidate genes.

**Figure S16.**
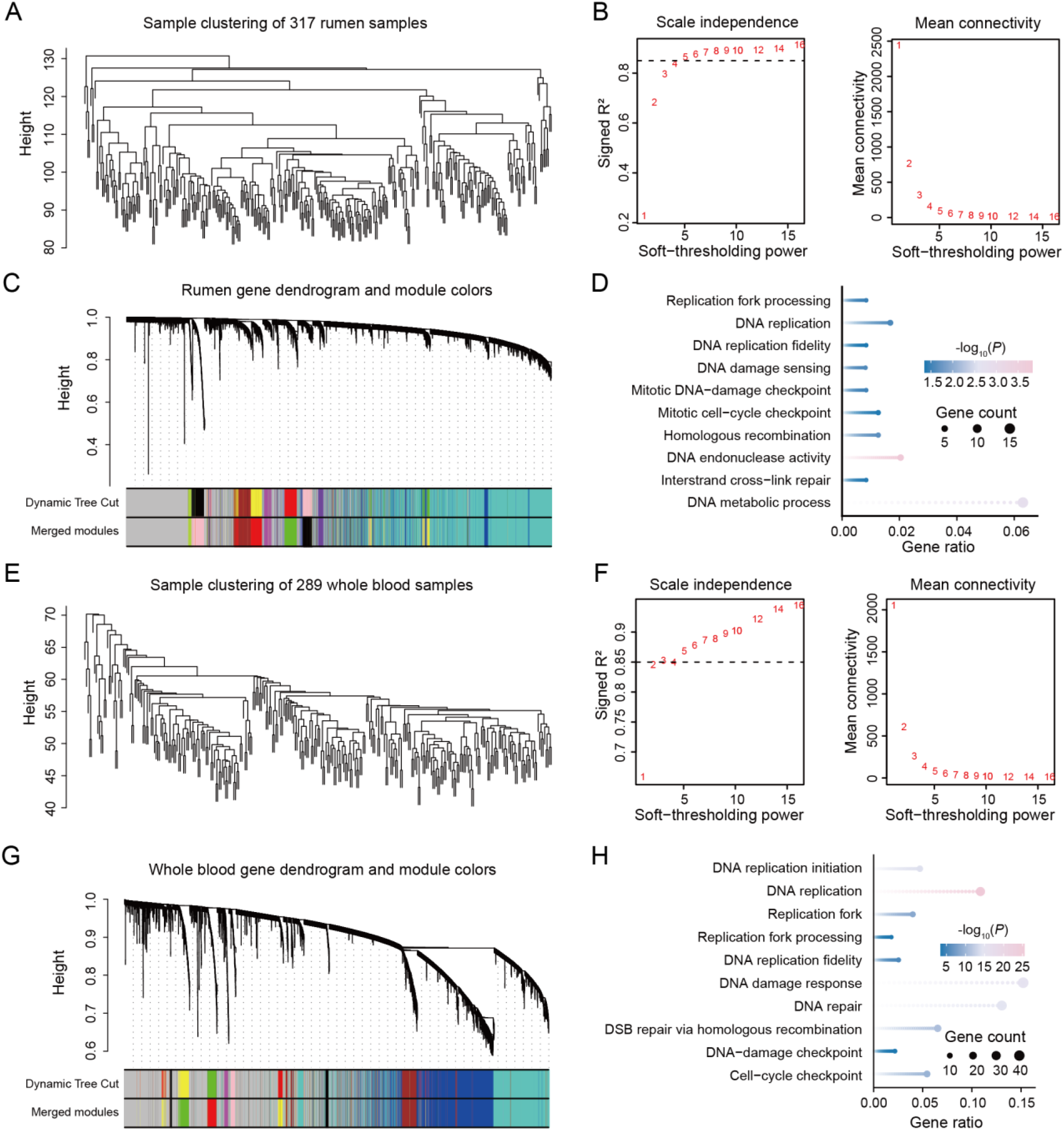
WGCNA network construction and functional enrichment in rumen and whole blood. (A) Sample clustering dendrogram for 317 rumen transcriptomes. (B) Soft-thresholding power selection for the rumen WGCNA network. The dashed line indicates signed *R*² = 0.85. (C) Gene clustering dendrogram and module assignment in rumen. (D) GO enrichment of genes in the rumen MEblack module. Abbreviated GO term labels correspond to the following full terms: DNA replication, DNA-templated DNA replication; DNA replication fidelity, DNA-templated DNA replication maintenance of fidelity; DNA damage sensing, DNA damage sensor activity; Mitotic DNA-damage checkpoint, mitotic DNA damage checkpoint signaling; and Mitotic cell-cycle checkpoint, mitotic cell cycle checkpoint signaling. (E) Sample clustering dendrogram for 289 whole-blood transcriptomes. (F) Soft-thresholding power selection for the whole-blood WGCNA network. The dashed line indicates signed *R*² = 0.85. (G) Gene clustering dendrogram and module assignment in whole blood. (H) GO enrichment of genes in the whole-blood MEgreen module. Abbreviated GO term labels correspond to the following full terms: DNA replication, DNA-templated DNA replication; DNA replication fidelity, DNA-templated DNA replication maintenance of fidelity; DSB repair via homologous recombination, double-strand break repair via homologous recombination; DNA-damage checkpoint, DNA damage checkpoint signaling; and Cell-cycle checkpoint, cell cycle checkpoint signaling.

