## Supplemental figures for "A telomere-to-telomere genome of Holstein cattle resolves chromosome-end architecture and the genetic basis of telomere length"

Document S1. Figures S1–S16 and Tables S1–S22

Supplemental tables (Supplementary Tables 1–22) are provided separately in the Excel workbook Supplementary_Tables.xlsx.

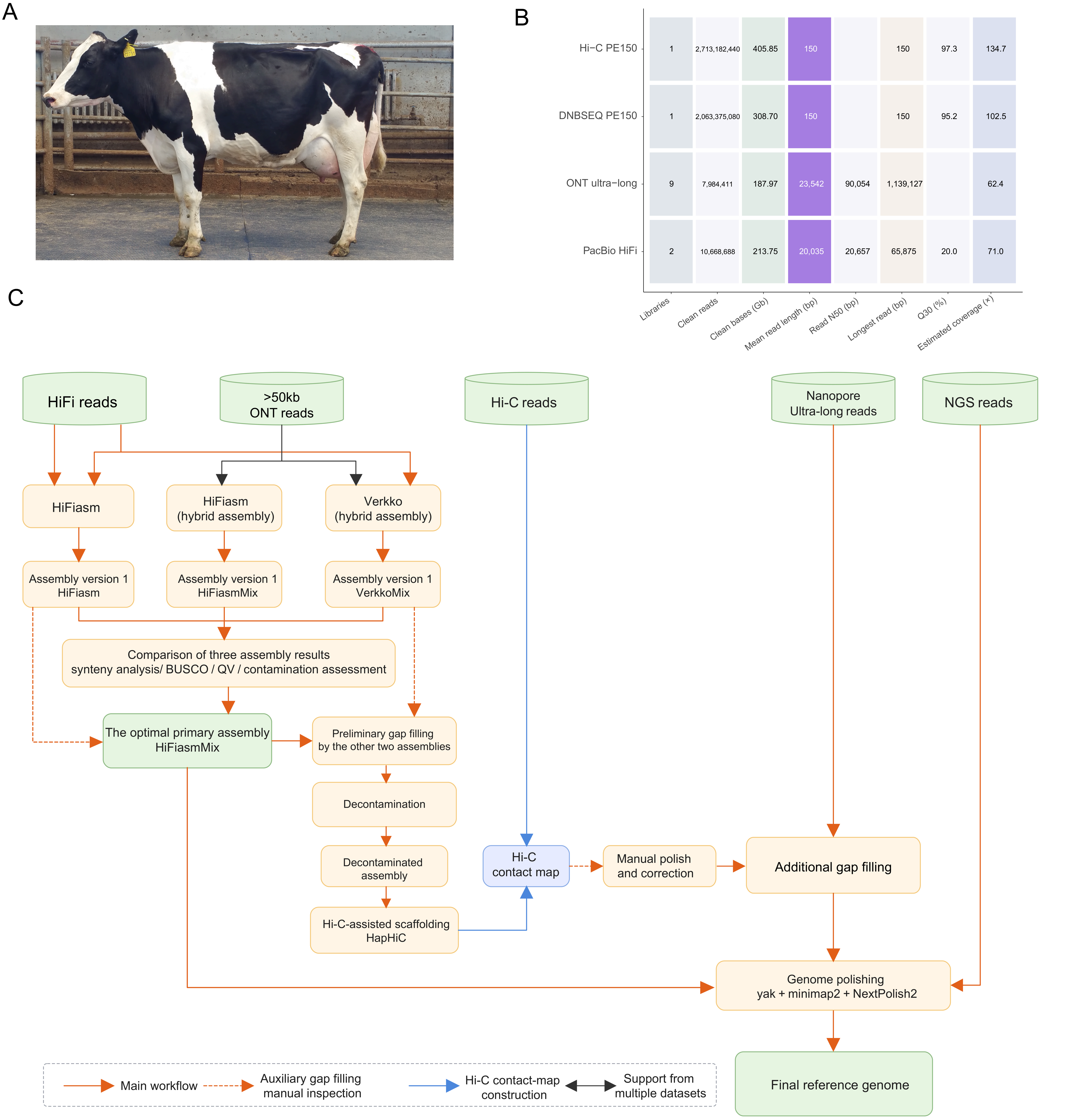


Study sample and overview of genome assembly. (A) Photograph of the donor Holstein cow, a seven-year-old female used for Holstein_T2T assembly. (B) Summary of sequencing datasets, including read number, sequencing yield, read N50, and estimated genome coverage across platforms. (C) Integrated Holstein_T2T assembly workflow using PacBio HiFi, ONT ultra-long, and Hi-C sequencing data.

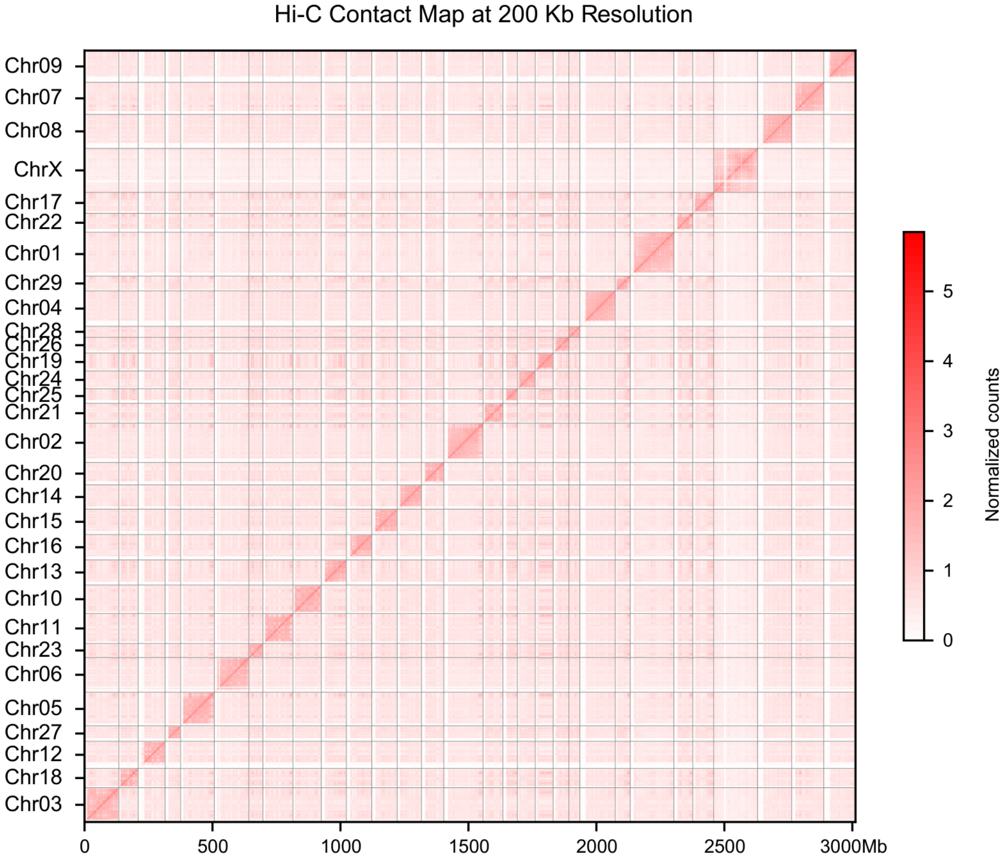


Hi-C chromatin interaction map supporting the structural accuracy of the assembly.

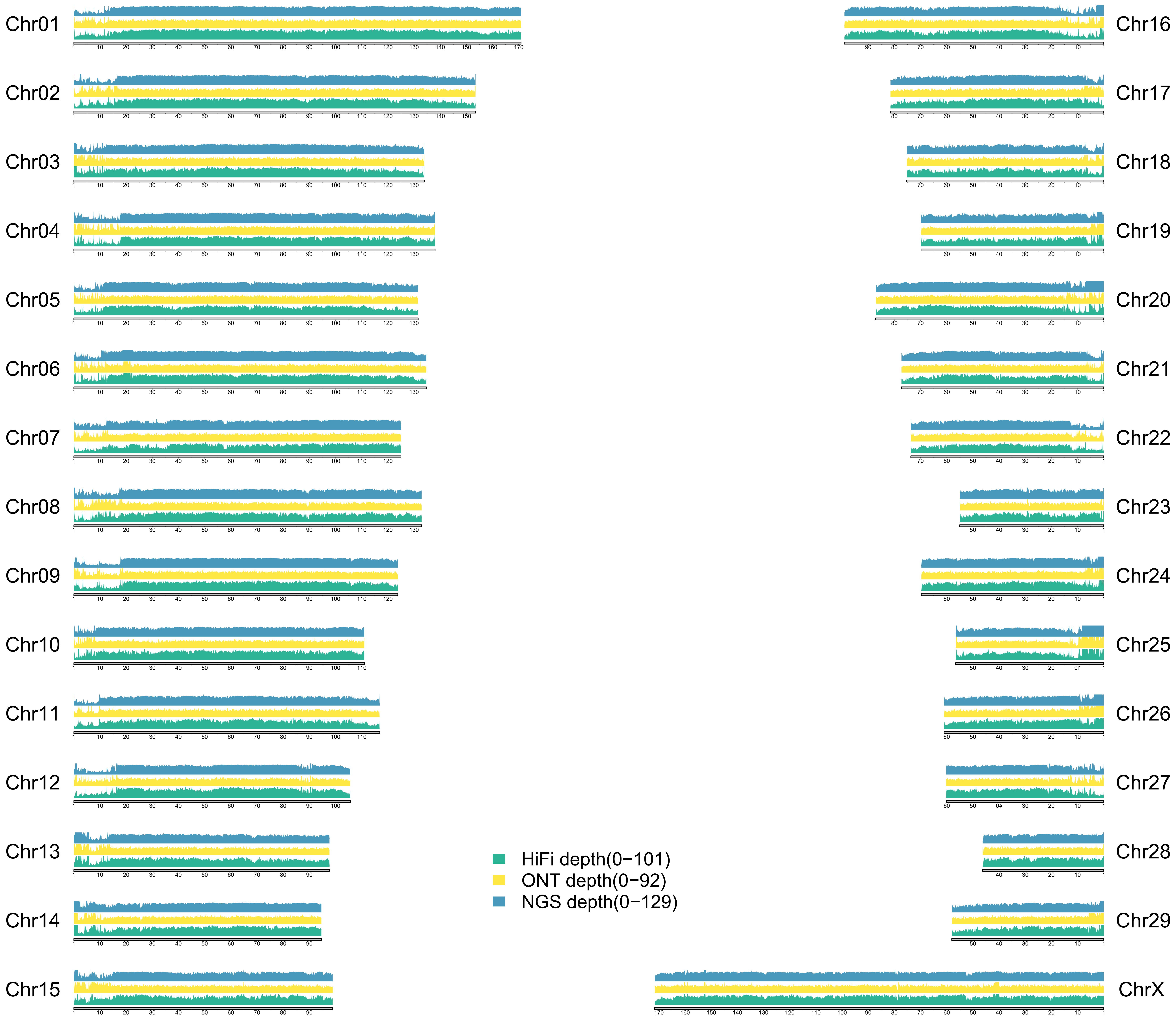


Sequencing-read coverage across the Holstein_T2T assembly.

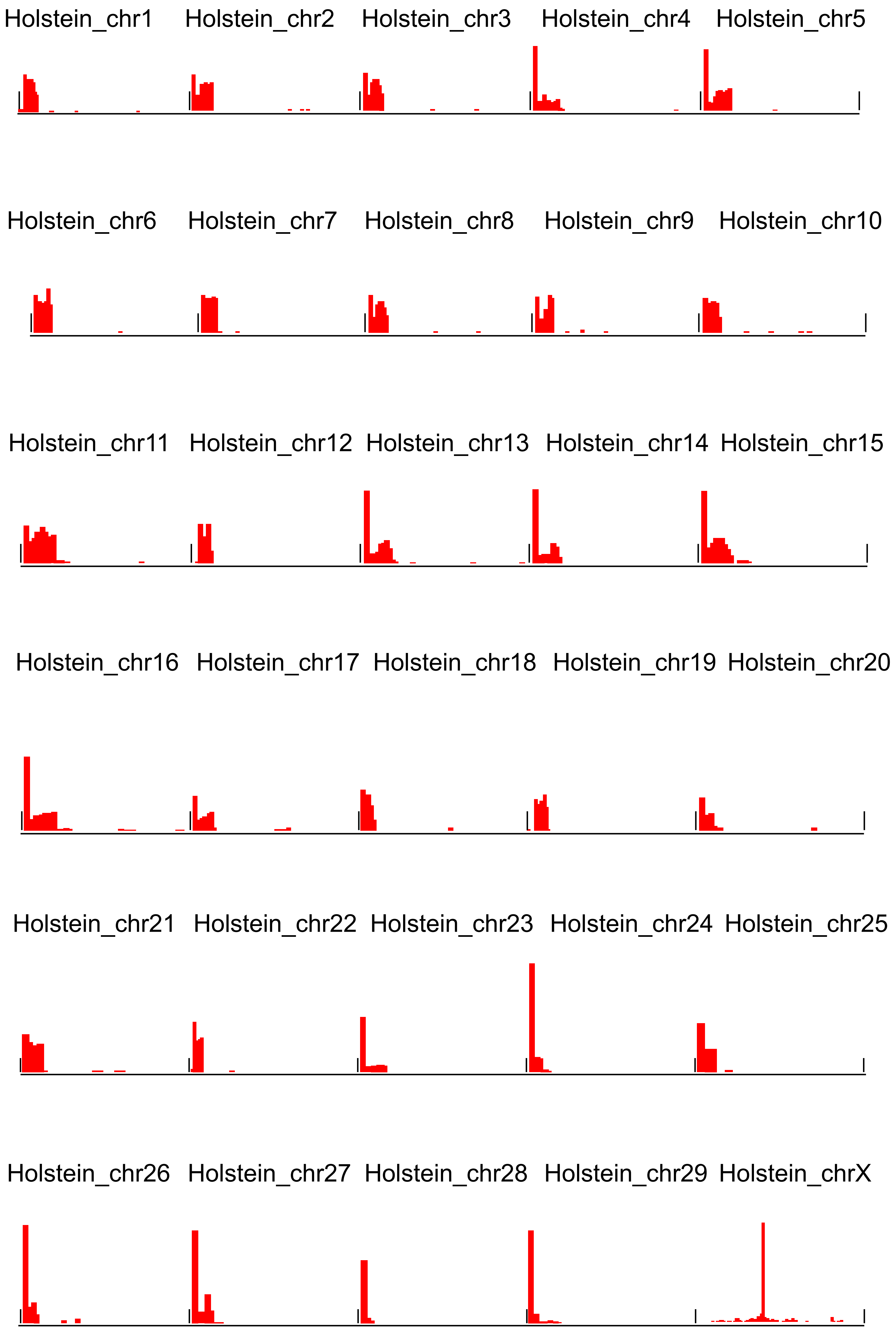


CENP-A CUT&Tag signal profiles.

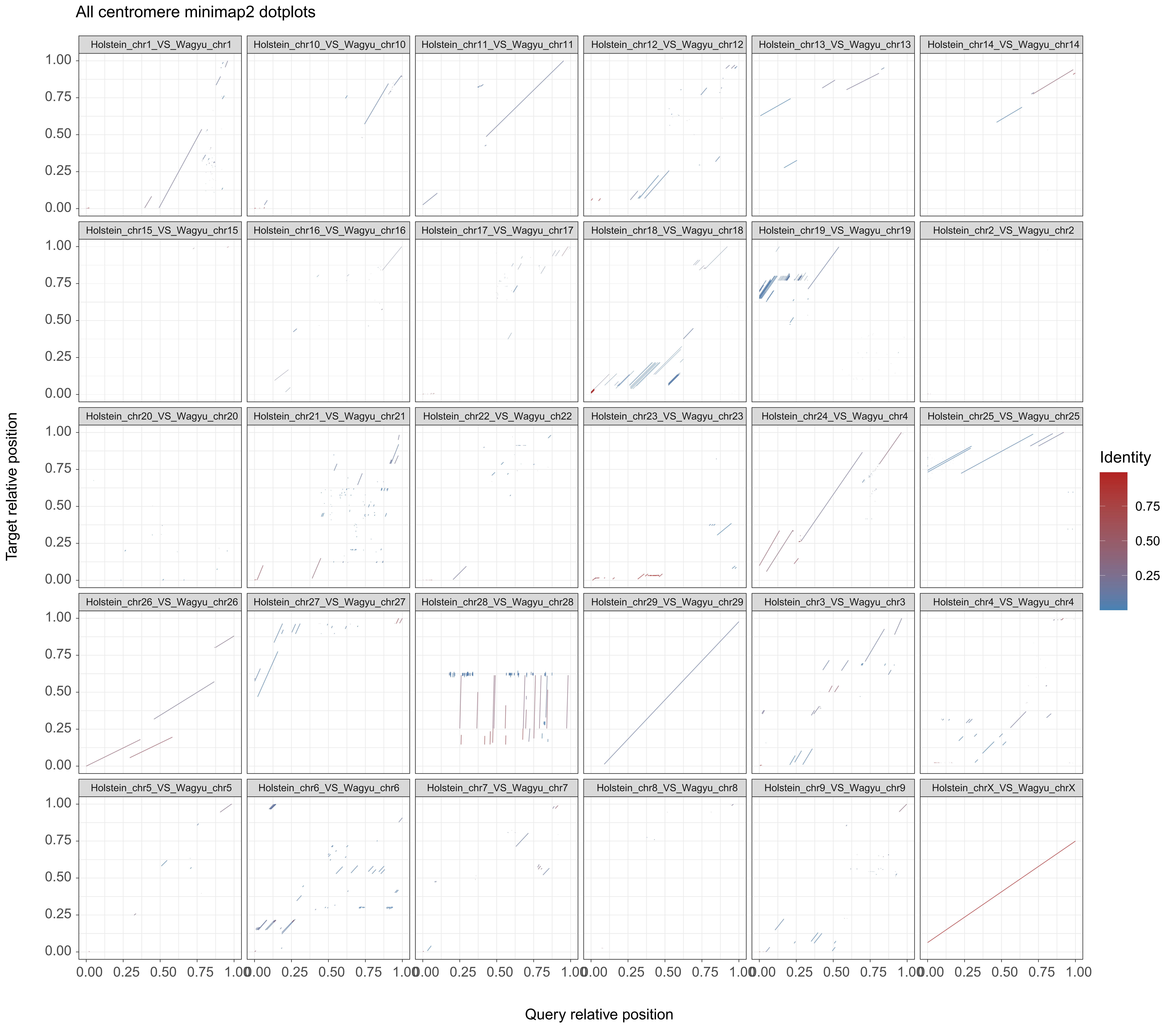


Comparative alignment of homologous centromeric regions between the Holstein_T2T and the Wagyu reference genome (UOA_Wagyu_1) by minimap2 Chromosome-by-chromosome dot plots of homologous centromeric intervals between the Holstein_T2T assembly and the Wagyu reference genome. Line color indicates local sequence identity.

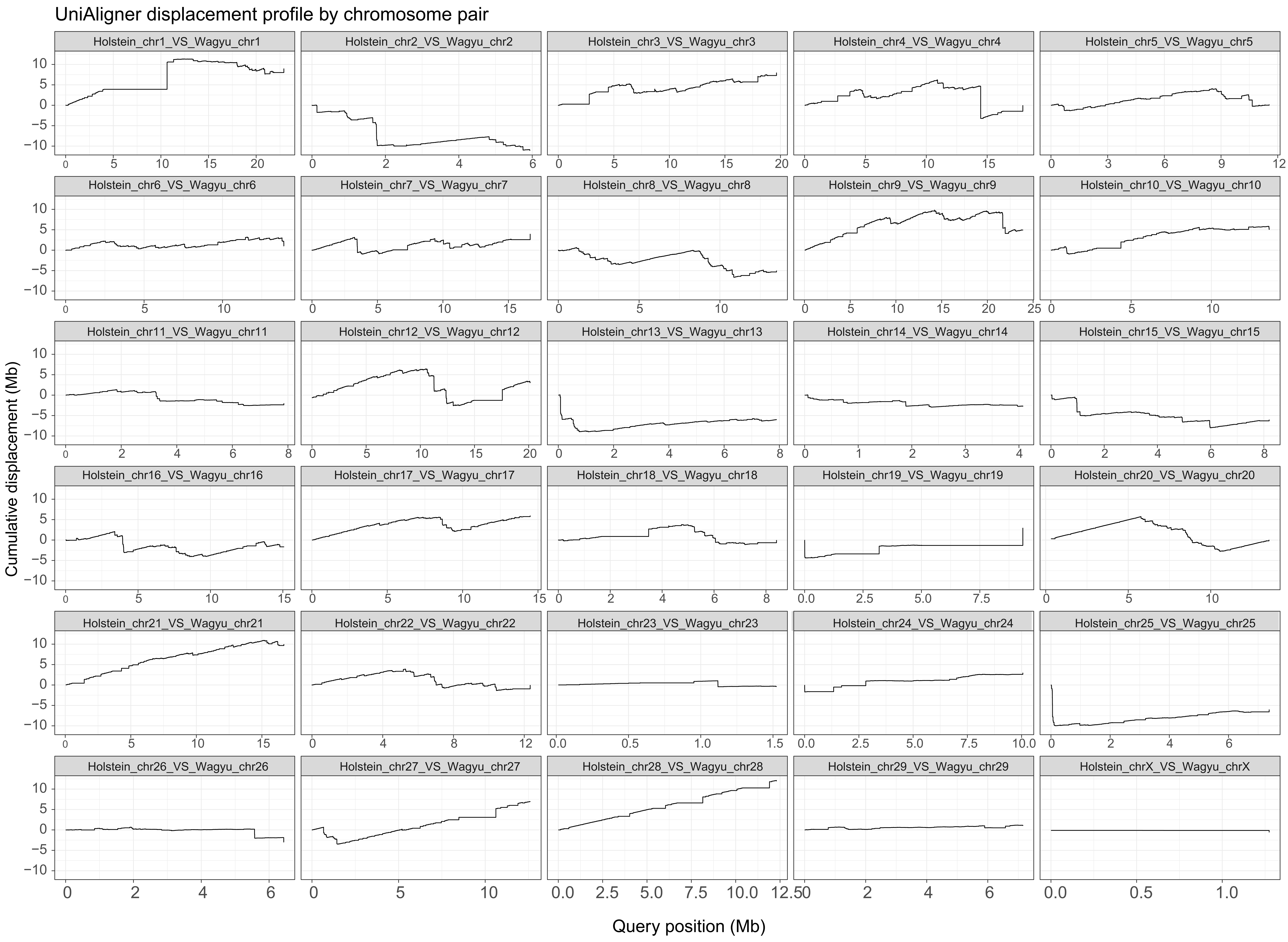


Comparative alignment of homologous centromeric regions between the Holstein_T2T and the Wagyu reference genome (UOA_Wagyu_1) by UniAligner Sequence-divergence profiles across the corresponding homologous centromeric intervals, showing local variation in alignment structure and sequence conservation.

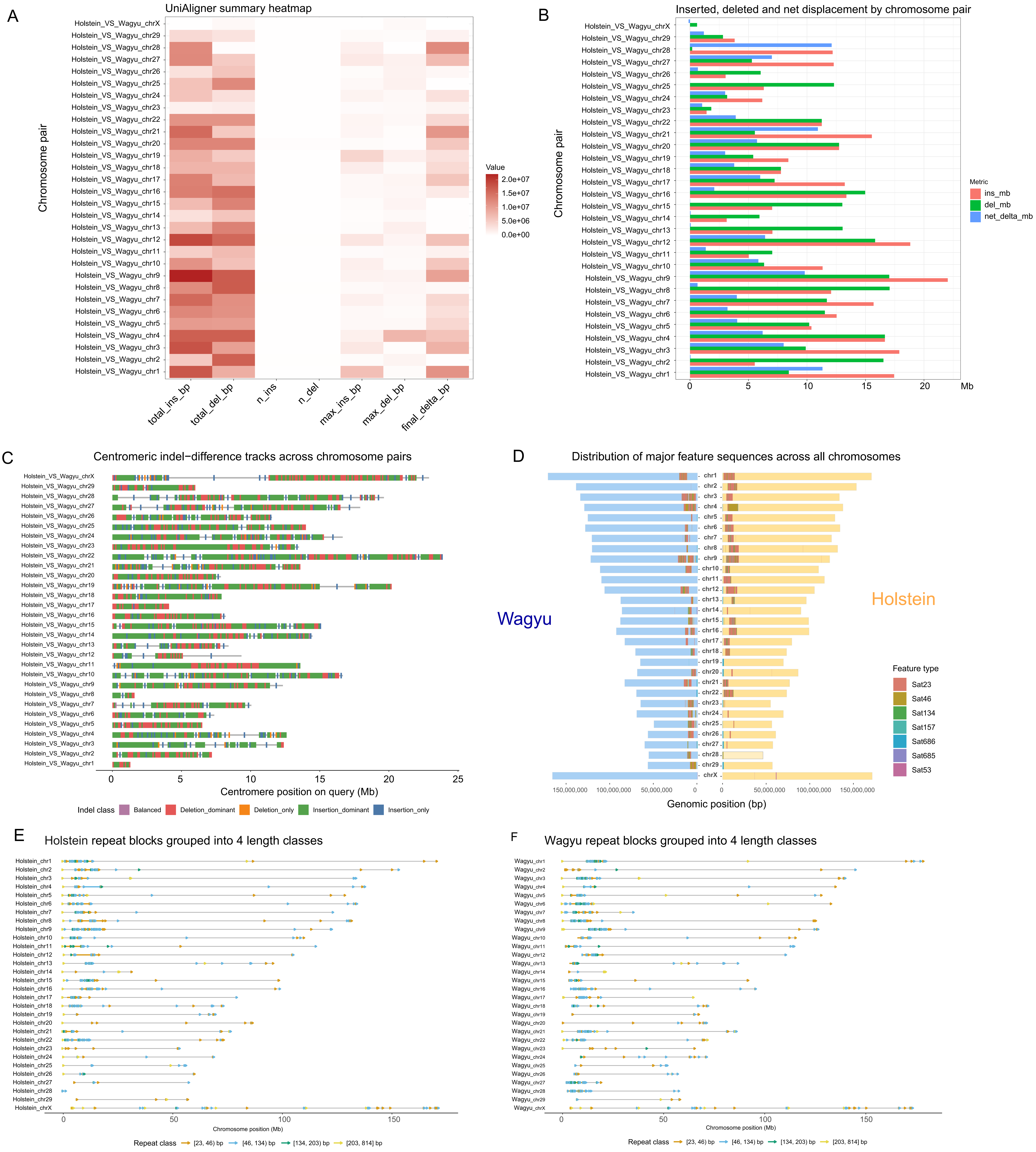


Structural variation and repeat-organization differences across homologous bovine centromeres. (A) Heatmap summarizing insertion and deletion burden across homologous centromeric intervals between the Holstein_T2T and the Wagyu reference genome. Columns represent indel-related metrics, including total insertion and deletion length, event number, maximum event size, and net sequence-length change. (B) Chromosome-level comparison of centromeric indel profiles, showing the relative contribution of insertion, deletion, and net length-change signals across homologous intervals. (C) Local classification of indel-dominated segments along each aligned centromeric interval. Colors indicate balanced, deletion-dominant, deletion-only, insertion-dominant, and insertion-only regions. (D) Comparison of centromeric interval length and repeat composition between the Wagyu reference genome and the Holstein_T2T assembly. Mirrored bars show homologous intervals in the two genomes, with colors indicating repeat categories. (E and F) Distribution of major centromeric repeat units along Holstein (E) and Wagyu (F) centromeric intervals. Colored points mark repeats units or repeat families detected within each chromosome-specific centromeric region, highlighting differences in repeat abundance and organization between the two genomes.

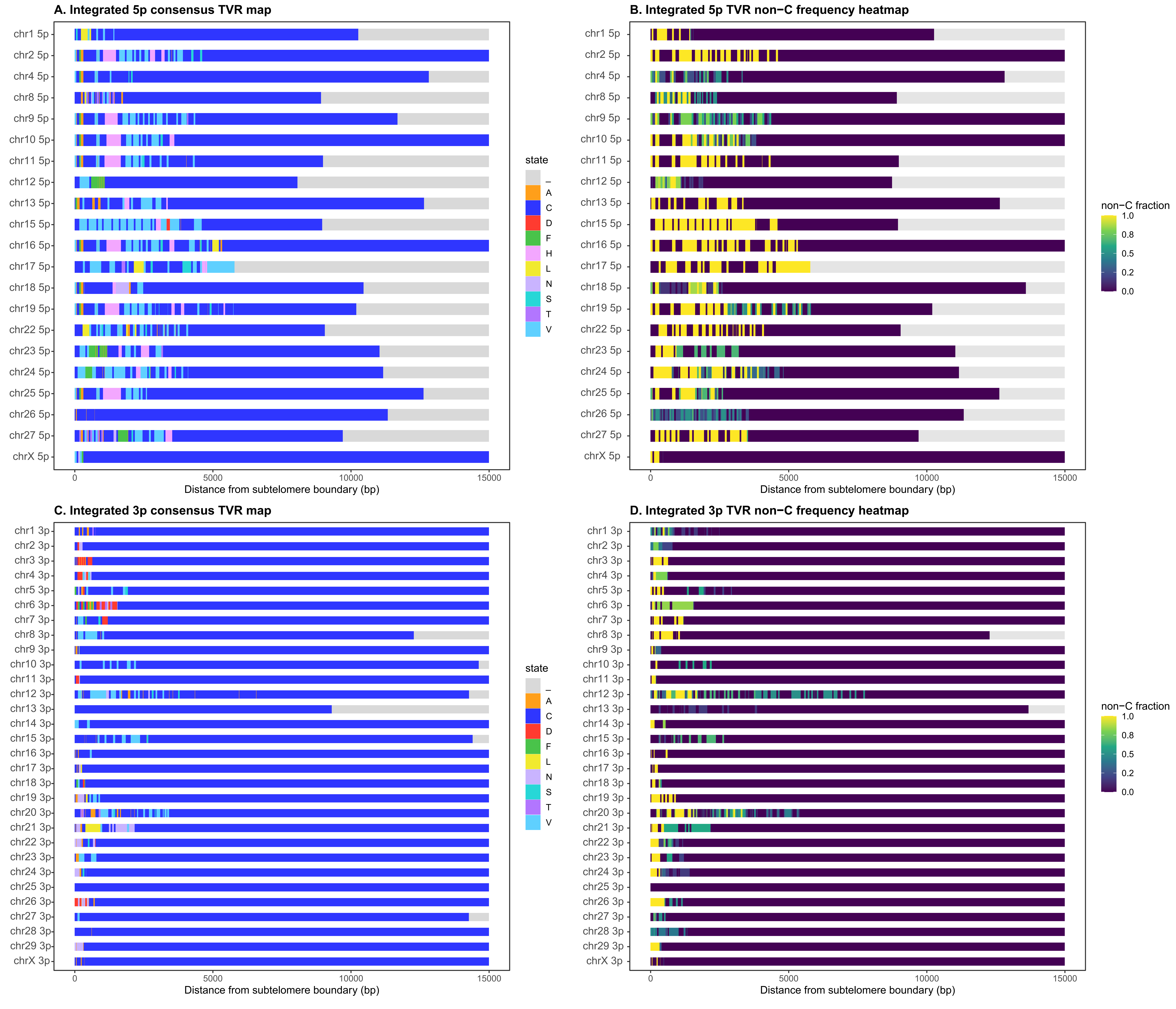


Chromosome-end-specific distribution of telomere variant repeats and non-canonical repeat content. (A) Base-state profiles of terminal telomeric repeat arrays at 5p chromosome ends. Rows represent individual chromosome ends, and columns indicate positions along the terminal repeat tract. Colors denote canonical and non-canonical repeat states, with gray indicating unassigned or gap regions. (B) Distribution of non-canonical repeat fraction across the corresponding 5p telomeric regions. Color intensity indicates the local fraction of non-canonical sequence (C) Base-state profiles of terminal telomeric repeat arrays at 3p chromosome ends, shown as in (A).
(D) Distribution of non-canonical repeat fraction across the corresponding 3p telomeric regions, shown as in (B).

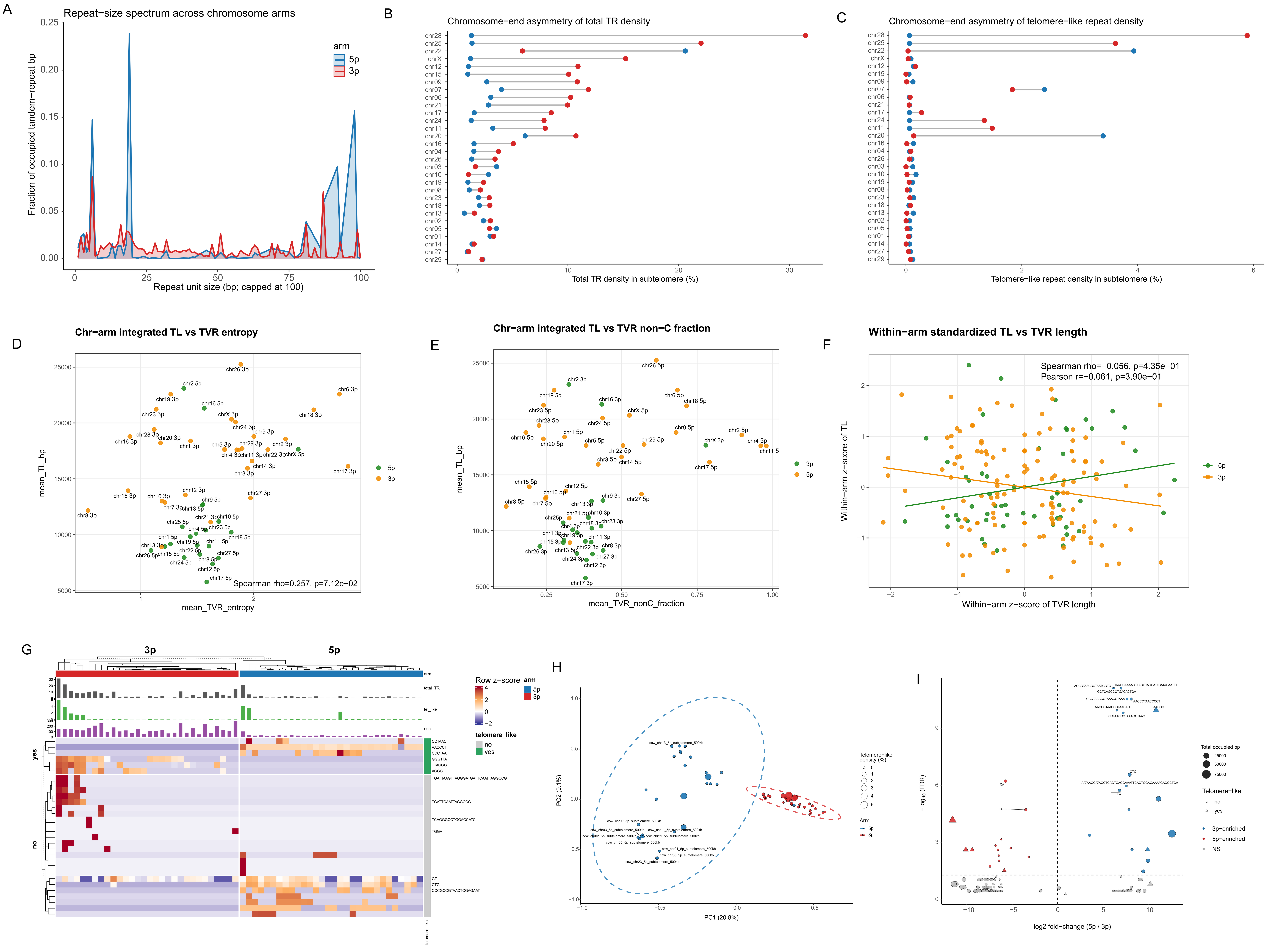


Subtelomeric tandem-repeat architecture and motif composition at chromosome ends. (A) Comparison of repeat-unit distribution patterns between 5p and 3p chromosome ends. (B) Proportion of tandem repeats at paired chromosome ends across the 30 chromosomes; 25 chromosomes show higher tandem-repeat content at 3p termini. (C) Distribution of telomere-like motif density across chromosome ends. (D–F) Correlations between telomere length and TVR abundance or non-canonical sequence composition, shown globally and by chromosome-end class. (G) Motif-abundance-based clustering of subtelomeric regions, separating 3p and 5p termini and identifying chromosome-arm-enriched motifs. (H) Principal-component analysis of motif density across subtelomeric regions (PERMANOVA, $R^2 = 0.178$, $P = 0.001$). (I) Differentially enriched motifs between 5p and 3p subtelomeric regions.

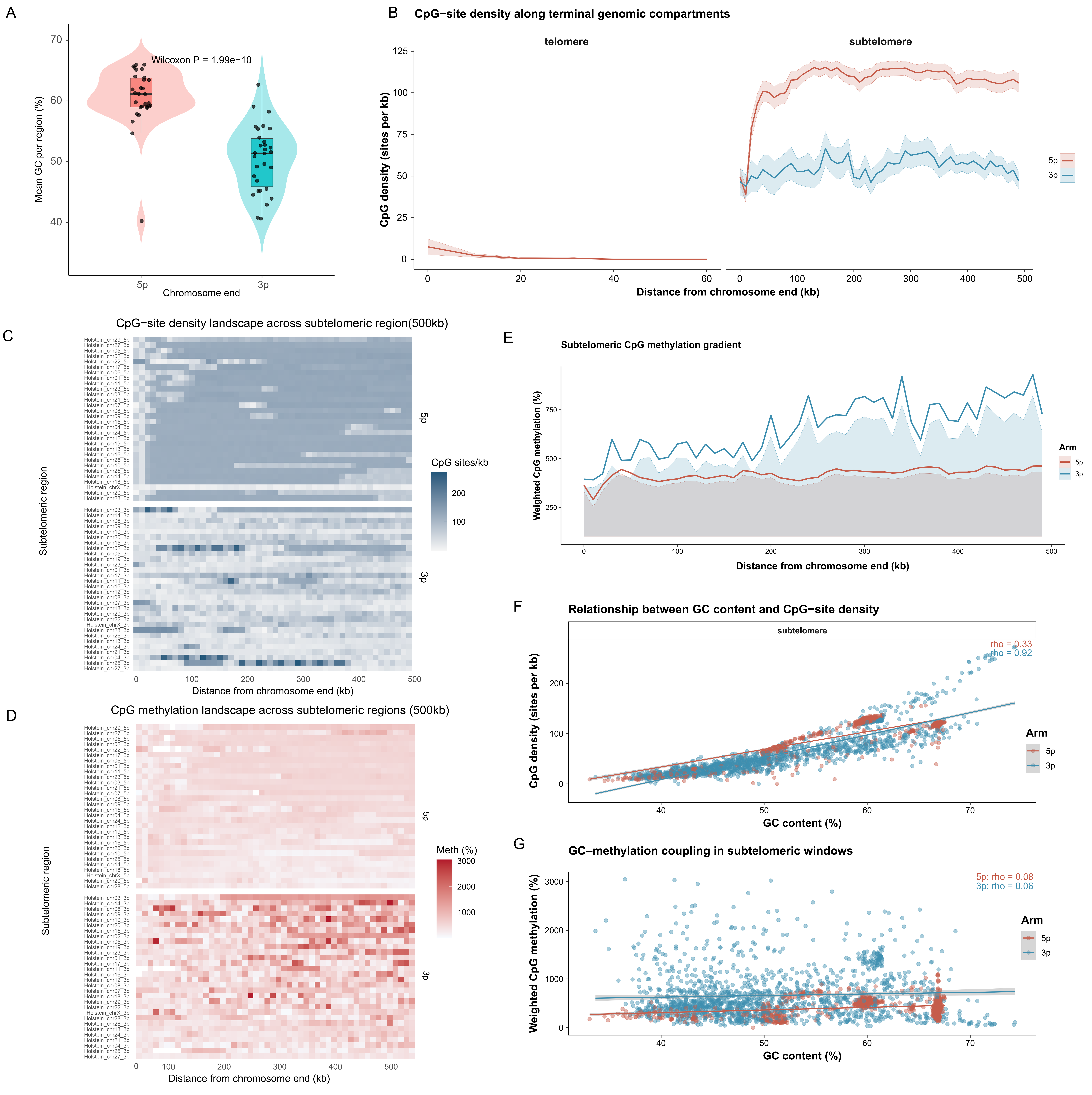


Subtelomeric GC/CpG composition and DNA methylation at chromosome ends. (B) Distance-resolved CpG-site density across terminal genomic compartments. CpG density is shown separately for telomeric repeat tracts and the adjacent 500-kb subtelomeric regions at 5p and 3p chromosome ends. Lines indicate mean profiles and shaded regions indicate variability across chromosome ends. (C) Chromosome-end-resolved heatmaps of CpG-site density across the 500-kb subtelomeric regions at 5p and 3p termini. Each row represents an individual chromosome end. (D) Chromosome-end-resolved heatmaps of CpG methylation across the 500-kb subtelomeric regions at 5p and 3p termini. Each row represents an individual chromosome end. (E) Distance-resolved profiles of weighted CpG methylation across the 500-kb subtelomeric regions at 5p and 3p chromosome ends. Lines show chromosome-arm-specific profiles, with shaded regions indicating variation across chromosome ends. (F) Relationship between GC content and CpG-site density across subtelomeric windows, shown separately for 5p and 3p chromosome ends. Lines indicate fitted trends; Spearman correlation coefficients are shown in the panel. (G) Relationship between GC content and weighted CpG methylation across subtelomeric windows at 5p and 3p chromosome ends. Lines indicate fitted trends; Spearman correlation coefficients are shown in the panel.

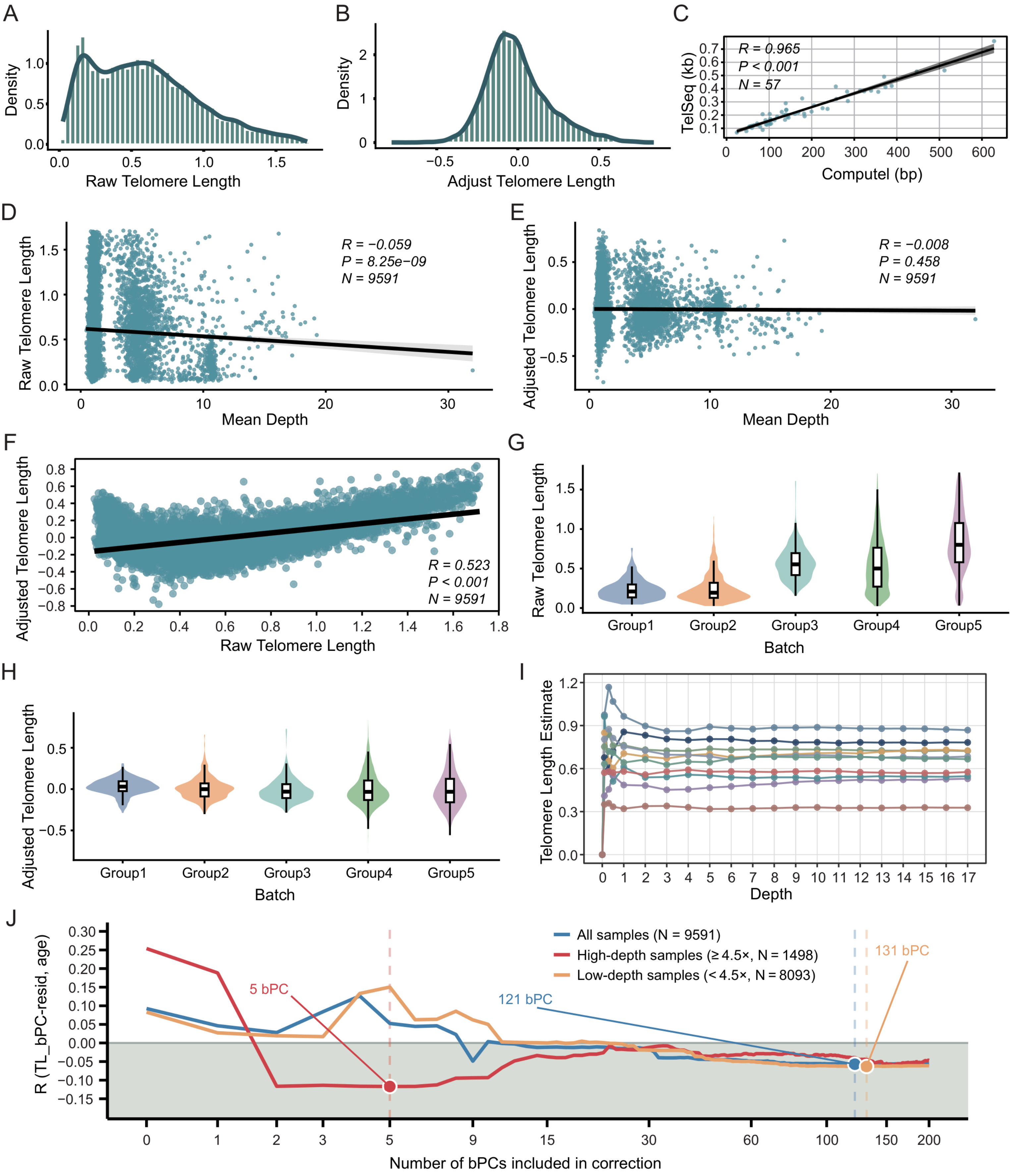


Telomere length phenotype correction and quality control. (A and B) Distribution of telomere length estimates before and after bPC correction. Independent validation of sequencing-based telomere length estimates using Computel. (D and E) Correlations between telomere length and mean sequencing depth before and after correction. (F) Correlation between raw and corrected telomere length estimates (r = 0.523). (G–J) Evaluation of batch effects, sequencing-depth sensitivity, and correction performance of coverage-derived batch principal components (bPCs).

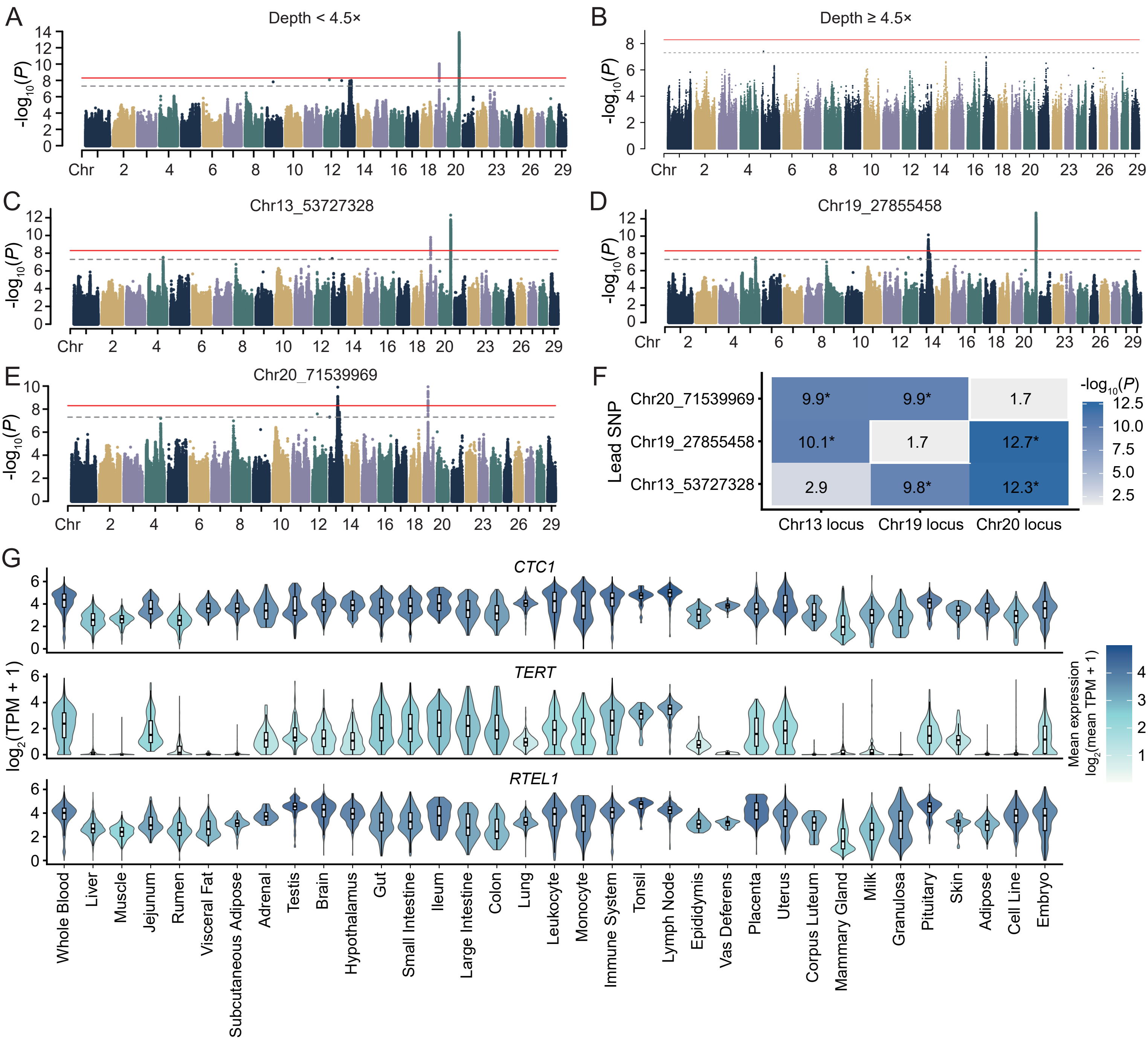


Sequencing-depth sensitivity, conditional GWAS, and tissue expression of telomere-maintenance genes. (C–E) Conditional GWAS performed using chr13_53727328 (C), chr19_27855458 (D), and chr20_71539969 (E) as conditioning variants. Conditioning attenuated the corresponding local association peak while retaining signals at the other major loci. (F) Residual association strength across the three major GWAS loci after conditional analysis. Rows indicate the conditioning lead SNP, columns indicate the chromosome 13, 19, and 20 loci, and values denote the maximum residual −log10(P) within ±1 Mb of each locus. Asterisks indicate residual signals with P < 5 × 10−8. (G) Tissue expression profiles of CTC1, TERT, and RTEL1 across cattle tissues. Violin plots show log2(TPM + 1) expression levels, highlighting broad expression of CTC1 and RTEL1 and low, tissue-restricted expression of TERT.

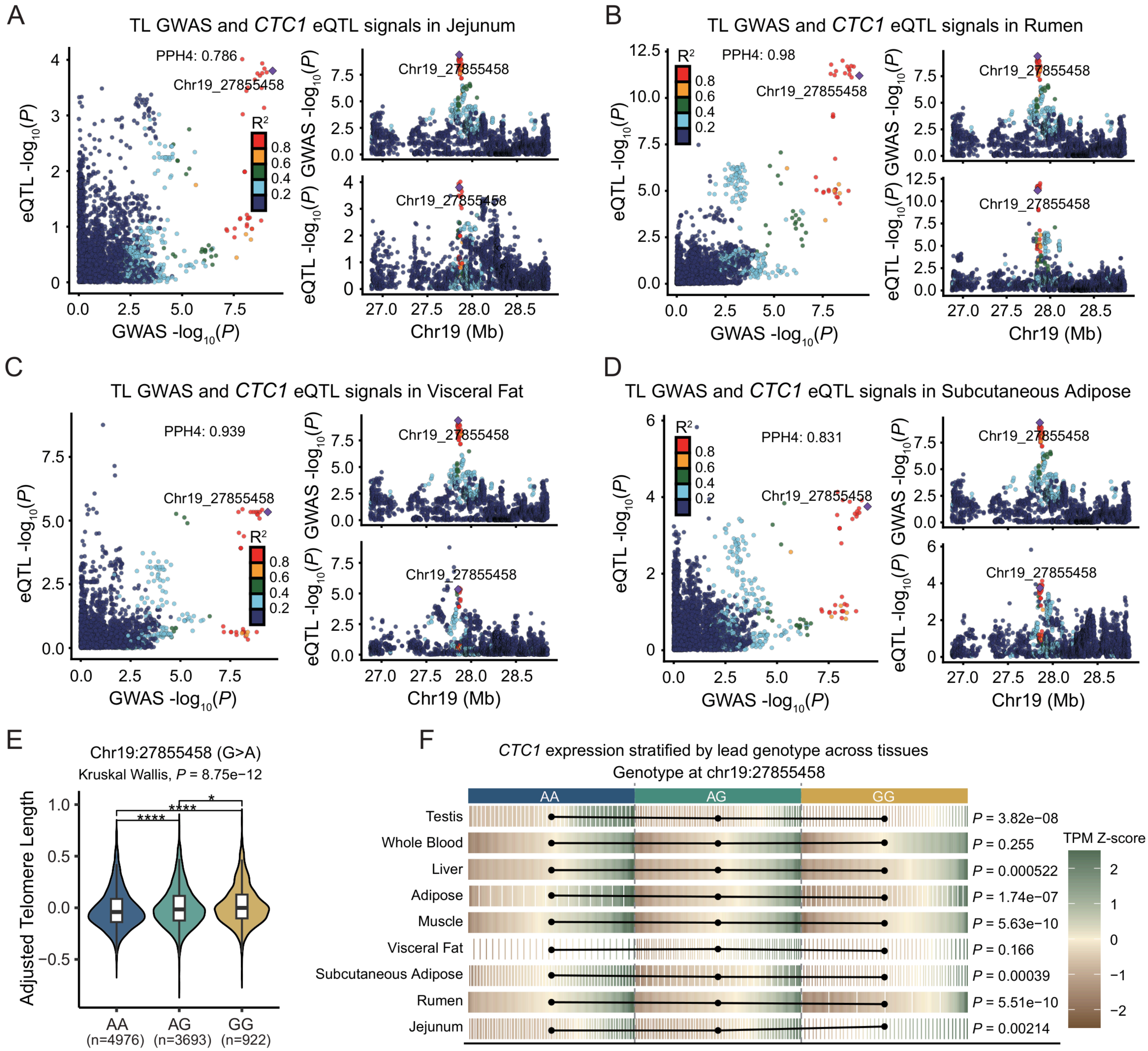


Multi-tissue eQTL colocalization at the CTC1 locus and genotype effects of the lead variant. (E) Distribution of corrected TL across chr19_27855458 genotypes. Sample sizes for AA, AG, and GG genotypes are 4,976, 3,693, and 922, respectively. Overall genotype effects were tested using the Kruskal–Wallis test ($P = 8.75 \times 10^{-12}$), with asterisks indicating significant pairwise comparisons. (F) Genotype-dependent CTC1 expression across tissues. Normalized expression levels are shown as a heatmap, with statistical significance for genotype–expression associations indicated on the right, supporting regulatory effects of chr19_27855458 on CTC1 expression in multiple tissues.

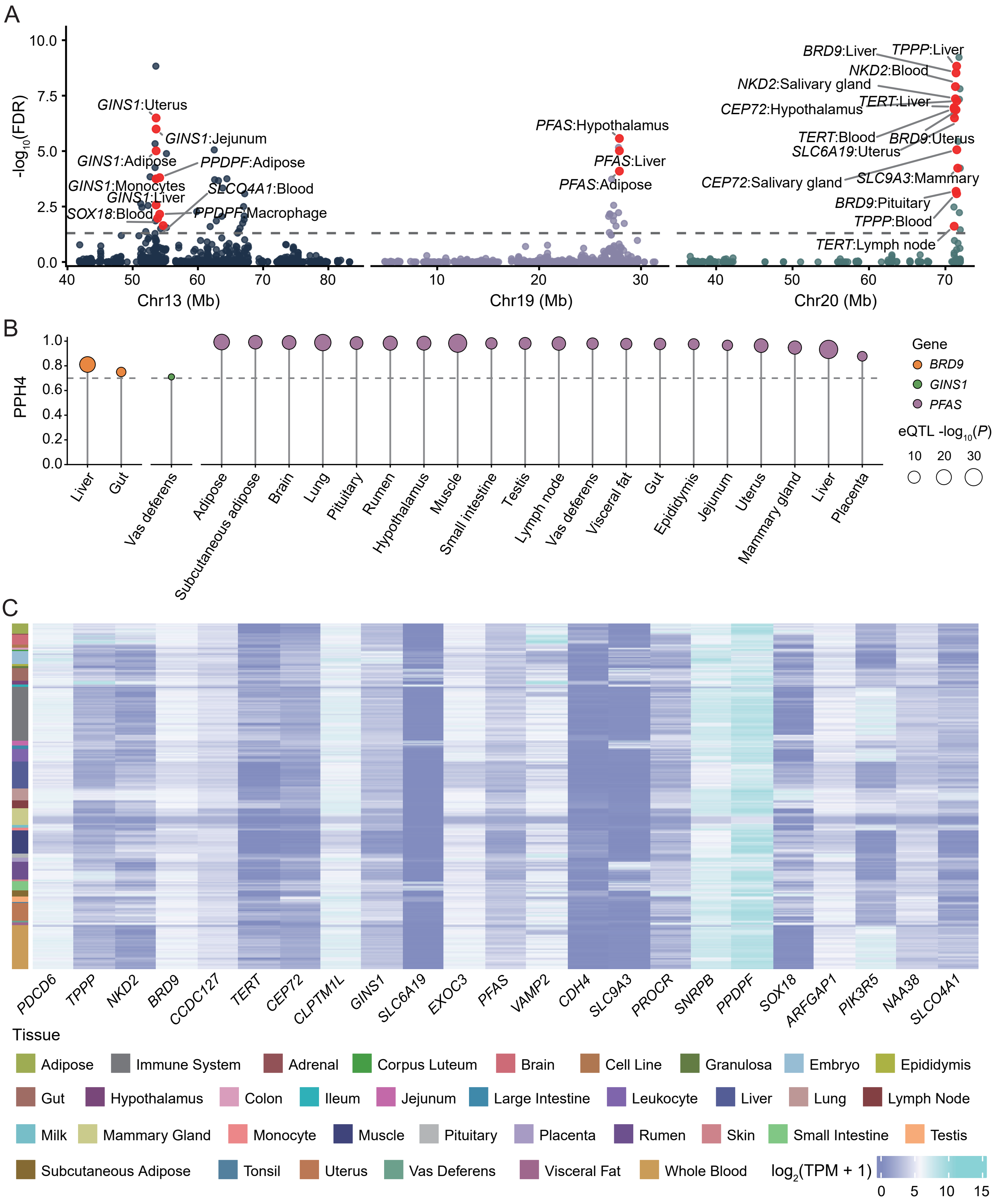


Multi-tissue colocalization, candidate-gene expression, and TWAS integration. (A) Multi-tissue TWAS results for the major telomere length-associated regions on chromosomes 13, 19, and 20. Each point represents a gene–tissue TWAS association, with genomic position on the x axis and TWAS significance on the y axis. The gray dashed line indicates the FDR = 0.05 threshold. Significant candidate gene–tissue signals are highlighted in red and labeled, including GINS1 and neighboring genes on chromosome 13, PFAS on chromosome 19, and TERT, BRD9, CEP72, NKD2, and TPPP on chromosome 20. (B) eQTL–teloGWAS colocalization for selected candidate genes, including BRD9, GINS1, and PFAS, across tissues. The y axis shows PPH4, the dashed line indicates the reference threshold for strong colocalization evidence, point color denotes candidate gene, and point size reflects eQTL association strength. (C) Tissue expression profiles of teloGWAS- and TWAS-supported candidate genes. Columns represent candidate genes, rows represent transcriptome samples from different cattle tissues, the left annotation bar indicates tissue category, and heatmap color denotes expression level.

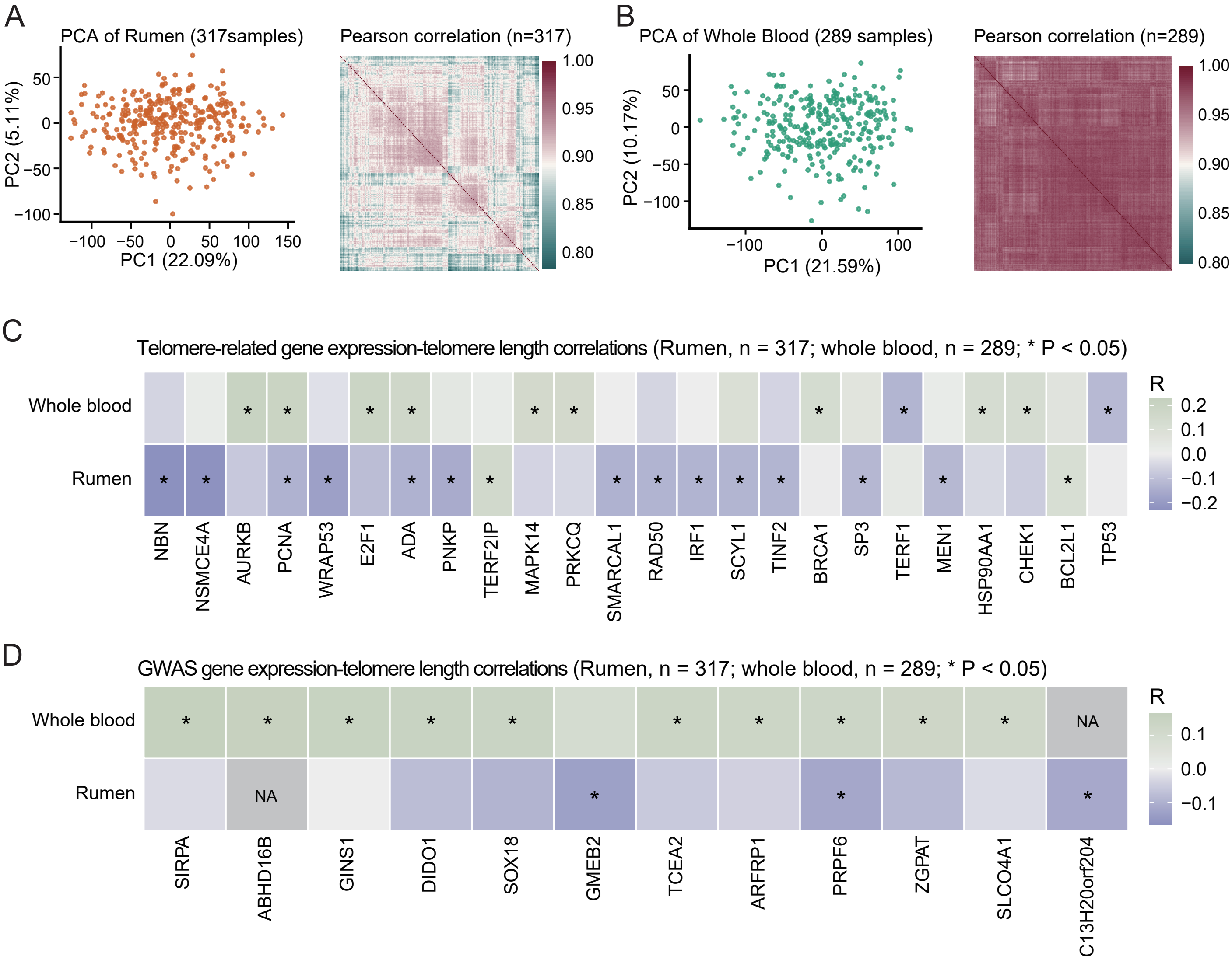


Expression–telomere length associations for telomere-related genes and GWAS candidates in rumen and whole blood. (A) Transcriptome quality assessment for 317 rumen samples. Principal-component analysis (PCA) of the gene expression matrix and inter-sample Pearson correlation heatmap were used to evaluate sample clustering and expression-data consistency. (B) Transcriptome quality assessment for 289 whole-blood samples, shown as in (A). (C) Correlations between expression levels of known telomere-maintenance genes and telomere length. Colors indicate Pearson correlation coefficients, and asterisks denote significant correlations (P < 0.05). Differences between rumen and whole blood indicate tissue-specific telomere-related transcriptional associations. (D) Correlations between expression levels of teloGWAS candidate genes and telomere length. Colors and asterisks are defined as in (C), highlighting tissue-dependent expression–TL associations among GWAS candidate genes.

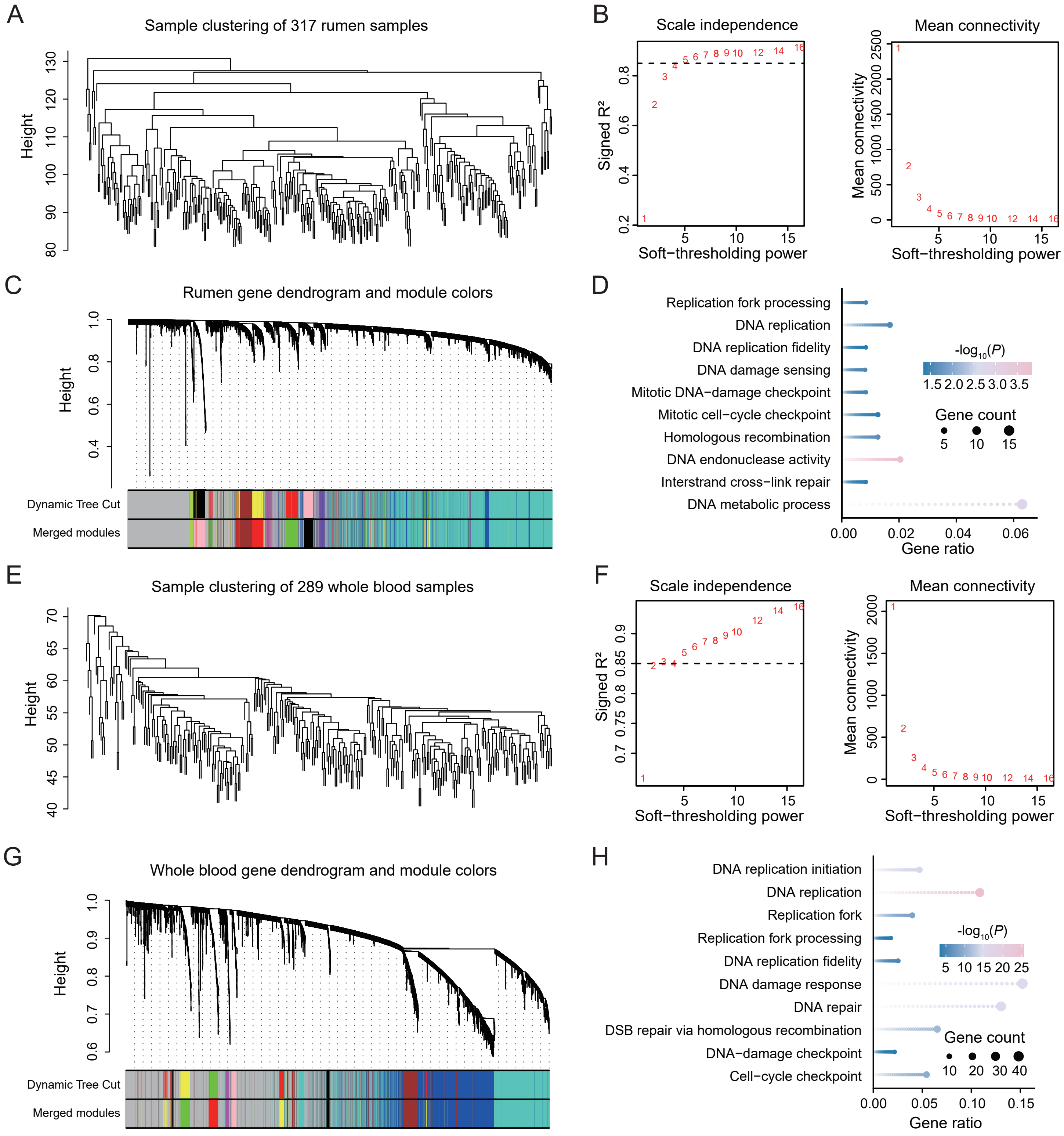


WGCNA network construction and functional enrichment in rumen and whole blood. (A) Sample clustering dendrogram for 317 rumen transcriptomes. (B) Soft-thresholding power selection for the rumen WGCNA network. The dashed line indicates signed $R^2 = 0.85$. (C) Gene clustering dendrogram and module assignment in rumen. (D) GO enrichment of genes in the rumen MEblack module. Abbreviated GO term labels correspond to the following full terms: DNA replication, DNA-templated DNA replication; DNA replication fidelity, DNA-templated DNA replication maintenance of fidelity; DNA damage sensing, DNA damage sensor activity; Mitotic DNA-damage checkpoint, mitotic DNA damage checkpoint signaling; and Mitotic cell-cycle checkpoint, mitotic cell cycle checkpoint signaling. (E) Sample clustering dendrogram for 289 whole-blood transcriptomes. (F) Soft-thresholding power selection for the whole-blood WGCNA network. The dashed line indicates signed $R^2 = 0.85$. (G) Gene clustering dendrogram and module assignment in whole blood. (H) GO enrichment of genes in the whole-blood MEgreen module. Abbreviated GO term labels correspond to the following full terms: DNA replication, DNA-templated DNA replication; DNA replication fidelity, DNA-templated DNA replication maintenance of fidelity; DSB repair via homologous recombination, double-strand break repair via homologous recombination; DNA-damage checkpoint, DNA damage checkpoint signaling; and Cell-cycle checkpoint, cell cycle checkpoint signaling.

### Supplementary Tables S1-S22

Rendered from Supplementary_Tables.xlsx (one worksheet per table)

**Supplementary Table 1. Summary of sequencing data in this study, related to Figure S1.**

| Sample ID | Breed | Sampling site | Tissue | Sequencing | Clean data (Gb) | Number of reads | Mean length or N50 (bp) | Depth (×) |
|---|---|---|---|---|---|---|---|---|
| Holstein_T2T | Holstein cattle | Hohhot, Inner Mongolia, China | Blood | DNA-seq, PacBio HiFi | 213.75 | 10668688 | 20657 | 71.13 |
| | | | | DNA-seq, ONT ultra-long | 187.97 | 7984411 | 90054 | 62.55 |
| | | | | DNA-seq, short-read (PE150) | 308.7 | 2063375080 | 2 × 150 | 102.73 |
| | | | | Hi-C (PE150) | 405.85 | 2713182440 | 2 × 150 | 135.06 |

**Supplementary Table 2. Holstein_T2T genome assembly and quality statistics, related to Figure 1.**

| Assembly feature | Value |
|---|---|
| Assembled bases (bp) | 3,005,015,558 |
| Assembly size (Gb) | 3.01 |
| Number of chromosomes | 30 |
| Gap number | 0 |
| Gap bases (bp) | 0 |
| Number of contigs | 30 |
| Contig N50 (Mb) | 111.23 Mb |
| Number of scaffolds | 30 |
| Scaffold N50 (Mb) | 111.23 Mb |
| Longest chromosome (Mb) | 171.64 |
| Merqury QV | 72.54 |
| BUSCO completeness (%) | 99.92 |
| Single-copy BUSCOs (%) | 98.8 |
| Duplicated BUSCOs (%) | 1.12 |
| Fragmented BUSCOs (%) | 0.04 |
| Missing BUSCOs (%) | 0.04 |
| Short-read mapping rate (%) | 96.86 |
| PacBio HiFi mapping rate (%) | 100 |
| ONT mapping rate (%) | 93.6 |
| CRAQ R-AQI | 99.4 |
| CRAQ S-AQI | 99.22 |
| Protein-coding genes | 21200 |
| Repeat sequences (Mb) | 1547.21 |
| Repeat sequences (%) | 51.36 |
| LINEs (Mb) | 775.64 |
| SINEs (Mb) | 141.26 |
| LTRs (Mb) | 177.15 |
| Satellite repeats (Mb) | 283.66 |

| | |
|---|---|
| DNA transposons (Mb) | 71.98 |

**Supplementary Table 3. Centromeric candidate intervals, related to Figure S4.**

| Chromosome | Chromosome length (bp) | Centromere start (bp) | Centromere end (bp) | Interval length (bp) |
|---|---|---|---|---|
| Chr01 | 170,625,057 | 21,589 | 13,912,593 | 13,891,004 |
| Chr02 | 153,687,530 | 21,589 | 16,949,247 | 16,927,658 |
| Chr03 | 133,980,150 | 21,674 | 11,649,047 | 11,627,373 |
| Chr04 | 138,061,648 | 21,624 | 17,934,424 | 17,912,800 |
| Chr05 | 131,645,373 | 21,589 | 11,352,881 | 11,331,292 |
| Chr06 | 134,763,664 | 20,402 | 12,914,308 | 12,893,906 |
| Chr07 | 125,212,660 | 21,589 | 12,668,680 | 12,647,091 |
| Chr08 | 132,963,638 | 21,589 | 18,507,991 | 18,486,402 |
| Chr09 | 111,232,015 | 21,589 | 18,881,263 | 18,859,674 |
| Chr10 | 111,247,955 | 248,461 | 8,604,860 | 8,356,399 |
| Chr11 | 105,680,519 | 21,589 | 9,867,665 | 9,846,076 |
| Chr12 | 97,706,411 | 1,209,181 | 14,174,503 | 12,965,322 |
| Chr13 | 97,727,959 | 3,766,147 | 13,914,811 | 10,148,664 |
| Chr14 | 95,023,284 | 3,671,331 | 6,177,921 | 2,506,590 |
| Chr15 | 98,952,501 | 1,346,981 | 14,875,732 | 13,528,751 |
| Chr16 | 99,349,279 | 1,751,485 | 16,675,591 | 14,924,106 |
| Chr17 | 81,655,903 | 21,589 | 8,290,433 | 8,268,844 |
| Chr18 | 75,688,686 | 323,563 | 8,414,399 | 8,090,836 |
| Chr19 | 69,880,705 | 2,659,687 | 6,347,017 | 3,687,330 |
| Chr20 | 87,425,511 | 9,281,650 | 13,555,521 | 4,273,871 |
| Chr21 | 77,391,410 | 21,589 | 6,679,489 | 6,657,900 |
| Chr22 | 73,747,813 | 196,762 | 12,516,183 | 12,319,421 |
| Chr23 | 55,160,999 | 21,589 | 1,631,926 | 1,610,337 |
| Chr24 | 69,783,461 | 1,787,603 | 7,064,780 | 5,277,177 |
| Chr25 | 56,631,963 | 7,725,712 | 13,378,837 | 5,653,125 |
| Chr26 | 61,048,865 | 3,682,167 | 9,445,711 | 5,763,544 |
| Chr27 | 60,215,355 | 38,208 | 5,418,519 | 5,380,311 |
| Chr28 | 68,732,309 | 21,589 | 206,548 | 184,959 |
| Chr29 | 58,148,320 | 1,359,097 | 6,054,088 | 4,694,991 |

| | | | | |
|---|---|---|---|---|
| ChrX | 171,644,615 | 109,800,000 | 110,300,000 | 500,000 |

*Note: chromosome lengths now refer to the final telomere-corrected assembly (30 sequences, 3,005,015,558 bp; N50 111,232,015 bp). Centromere coordinates were called on the pre-correction analysis assembly (see Methods); liftover re-validation to the final coordinate system is pending.*

**Supplementary Table 4. Telomere coordinates and telomere-length changes before and after correction in the Holstein_T2T genome, related to Figure 2.**

| Chromosome | Chromosome end | Pre-correction start | Pre-correction end | Pre-correction TL (bp) | Corrected start | Corrected end | Corrected TL (bp) | ΔTL (bp) | \|ΔTL\| (bp) | Corrected TL (kb) | TL change | Coordinate status | Telogator2 target TL | Agreement with read-based target |
|---|---|---|---|---|---|---|---|---|---|---|---|---|---|---|
| chr01 | 5p | 1195 | 14905 | 13710 | 1 | 7263 | 7262 | -6448 | 6448 | 7.262 | Shortened | Updated | 7262 | PASS |
| chr01 | 3p | 170617104 | 170619565 | 2461 | 170609461 | 170625057 | 15596 | 13135 | 13135 | 15.596 | Extended | Updated | 15596 | PASS |
| chr02 | 5p | 1 | 32827 | 32826 | 1 | 23096 | 23095 | -9731 | 9731 | 23.095 | Shortened | Updated | 23094 | PASS |
| chr02 | 3p | 153679643 | 153681460 | 1817 | 153669909 | 153687530 | 17621 | 15804 | 15804 | 17.621 | Extended | Updated | 17620 | PASS |
| chr03 | 5p | 1 | 2253 | 2252 | 1 | 2253 | 2252 | 0 | 0 | 2.252 | Unchanged | Unchanged (no read-based target) | — | — |
| chr03 | 3p | 133978454 | 133980146 | 1692 | 133978454 | 133980146 | 1692 | 0 | 0 | 1.692 | Unchanged | Unchanged (no read-based target) | — | — |
| chr04 | 5p | 1 | 2018 | 2017 | 1 | 9963 | 9962 | 7945 | 7945 | 9.962 | Extended | Updated | 9962 | PASS |
| chr04 | 3p | 138039372 | 138040906 | 1534 | 138047315 | 138061648 | 14333 | 12799 | 12799 | 14.333 | Extended | Updated | 14332 | PASS |
| chr05 | 5p | 1 | 65759 | 65758 | 1 | 65759 | 65758 | 0 | 0 | 65.758 | Unchanged | Unchanged (no read-based target) | — | — |
| chr05 | 3p | 131627891 | 131629541 | 1650 | 131627892 | 131645373 | 17481 | 15831 | 15831 | 17.481 | Extended | Updated | 17482 | PASS |
| chr06 | 5p | 5 | 1931 | 1926 | 5 | 1931 | 1926 | 0 | 0 | 1.926 | Unchanged | Unchanged (no read-based target) | — | — |
| chr06 | 3p | 134740362 | 134741889 | 1527 | 134740361 | 134763664 | 23303 | 21776 | 21776 | 23.303 | Extended | Updated | 23302 | PASS |
| chr07 | 5p | 346 | 18425 | 18079 | 1 | 9383 | 9382 | -8697 | 8697 | 9.382 | Shortened | Updated | 9383 | PASS |
| chr07 | 3p | 125212199 | 125213465 | 1266 | 125203157 | 125212660 | 9503 | 8237 | 8237 | 9.503 | Extended | Updated | 9503 | PASS |
| chr08 | 5p | 524 | 9351 | 8827 | 524 | 9351 | 8827 | 0 | 0 | 8.827 | Unchanged | Unchanged (no read-based target) | — | — |
| chr08 | 3p | 132951563 | 132952661 | 1098 | 132951563 | 132963638 | 12075 | 10977 | 10977 | 12.075 | Extended | Updated | 12075 | PASS |
| chr09 | 5p | 1102 | 45655 | 44553 | 1 | 10281 | 10280 | -34273 | 34273 | 10.28 | Shortened | Updated | 10280 | PASS |
| chr09 | 3p | 111248935 | 111252360 | 3425 | 111213561 | 111232015 | 18454 | 15029 | 15029 | 18.454 | Extended | Updated (3' terminal region replaced; 3' tail removed: chr length 124,097,578 -> 111,232,015 bp) | 18455 | PASS |
| chr10 | 5p | 932 | 13852 | 12920 | 1 | 9441 | 9440 | -3480 | 3480 | 9.44 | Shortened | Updated | 9439 | PASS |
| chr10 | 3p | 111248935 | 111252360 | 3425 | 111244522 | 111247947 | 3425 | 0 | 0 | 3.425 | Unchanged | Unchanged (source tele.bed coordinate invalid: older assembly version; arm skipped) | 14433 | — |
| chr11 | 5p | 1308 | 30033 | 28725 | 1308 | 30033 | 28725 | 0 | 0 | 28.725 | Unchanged | Unchanged (no read-based target) | — | — |
| chr11 | 3p | 105661780 | 105663422 | 1642 | 105661781 | 105680519 | 18738 | 17096 | 17096 | 18.738 | Extended | Updated (3' terminal region replaced; 3' tail removed: chr length 117,105,659 -> 105,680,519 bp) | 18739 | PASS |
| chr12 | 5p | 1 | 20996 | 20995 | 1 | 6296 | 6295 | -14700 | 14700 | 6.295 | Shortened | Updated | 6296 | PASS |
| chr12 | 3p | 97707062 | 97708051 | 989 | 97692362 | 97706411 | 14049 | 13060 | 13060 | 14.049 | Extended | Updated (3' terminal region replaced; 3' tail removed: chr length 105,663,422 -> 97,706,411 bp) | 14049 | PASS |
| chr13 | 5p | 1 | 1850 | 1849 | 1 | 12651 | 12650 | 10801 | 10801 | 12.65 | Extended | Updated | 12650 | PASS |
| chr13 | 3p | 97708145 | 97709323 | 1178 | 97717862 | 97727959 | 10097 | 8919 | 8919 | 10.097 | Extended | Updated | 9014 | FAIL (1083 bp) |
| chr14 | 5p | 499 | 11159 | 10660 | 499 | 11159 | 10660 | 0 | 0 | 10.66 | Unchanged | Unchanged (no read-based target) | — | — |
| chr14 | 3p | 95007048 | 95008543 | 1495 | 95007049 | 95023284 | 16235 | 14740 | 14740 | 16.235 | Extended | Updated | 16236 | PASS |
| chr15 | 5p | 722 | 1447 | 725 | 1 | 9356 | 9355 | 8630 | 8630 | 9.355 | Extended | Updated | 9356 | PASS |
| chr15 | 3p | 98931913 | 98932996 | 1083 | 98939822 | 98952501 | 12679 | 11596 | 11596 | 12.679 | Extended | Updated | 12679 | PASS |
| chr16 | 5p | 390 | 17191 | 16801 | 390 | 17191 | 16801 | 0 | 0 | 16.801 | Unchanged | Unchanged (no read-based target) | — | — |
| chr16 | 3p | 99348094 | 99349279 | 1185 | 99348094 | 99349279 | 1185 | 0 | 0 | 1.185 | Unchanged | Unchanged (no read-based target) | — | — |
| chr17 | 5p | 844 | 8388 | 7544 | 844 | 8388 | 7544 | 0 | 0 | 7.544 | Unchanged | Unchanged (no read-based target) | — | — |
| chr17 | 3p | 81652644 | 81653587 | 943 | 81652644 | 81653587 | 943 | 0 | 0 | 0.943 | Unchanged | Unchanged (no read-based target) | — | — |
| chr18 | 5p | 625 | 11454 | 10829 | 1 | 10089 | 10088 | -741 | 741 | 10.088 | Shortened | Updated | 10089 | PASS |
| chr18 | 3p | 75672353 | 75674794 | 2441 | 75670988 | 75688686 | 17698 | 15257 | 15257 | 17.698 | Extended | Updated | 17698 | PASS |
| chr19 | 5p | 38 | 249 | 211 | 1 | 9752 | 9751 | 9540 | 9540 | 9.751 | Extended | Updated | 9485 | FAIL (266 bp) |
| chr19 | 3p | 69836700 | 69837472 | 772 | 69845935 | 69880705 | 34770 | 33998 | 33998 | 34.77 | Extended | Updated | 34769 | PASS |
| chr20 | 5p | 1128 | 36019 | 34891 | 1 | 7220 | 7219 | -27672 | 27672 | 7.219 | Shortened | Updated | 7218 | PASS |
| chr20 | 3p | 87444206 | 87445120 | 914 | 87415406 | 87425511 | 10105 | 9191 | 9191 | 10.105 | Extended | Updated | 10106 | PASS |
| chr21 | 5p | 554 | 12667 | 12113 | 1 | 7551 | 7550 | -4563 | 4563 | 7.55 | Shortened | Updated | 7551 | PASS |
| chr21 | 3p | 77384801 | 77387364 | 2563 | 77379686 | 77391410 | 11724 | 9161 | 9161 | 11.724 | Extended | Updated | 11725 | PASS |
| chr22 | 5p | 36497 | 37572 | 1075 | 36497 | 37572 | 1075 | 0 | 0 | 1.075 | Unchanged | Unchanged (no read-based target) | — | — |
| chr22 | 3p | 73731985 | 73733004 | 1019 | 73731986 | 73747813 | 15827 | 14808 | 14808 | 15.827 | Extended | Updated | 15828 | PASS |
| chr23 | 5p | 1 | 26567 | 26566 | 1 | 8666 | 8665 | -17901 | 17901 | 8.665 | Shortened | Updated | 8666 | PASS |
| chr23 | 3p | 55161100 | 55165163 | 4063 | 55143200 | 55160999 | 17799 | 13736 | 13736 | 17.799 | Extended | Updated | 17800 | PASS |
| chr24 | 5p | 1 | 2228 | 2227 | 1 | 10353 | 10352 | 8125 | 8125 | 10.352 | Extended | Updated | 10353 | PASS |
| chr24 | 3p | 69756583 | 69759482 | 2899 | 69764708 | 69783461 | 18753 | 15854 | 15854 | 18.753 | Extended | Updated | 18753 | PASS |
| chr25 | 5p | 836 | 18108 | 17272 | 1 | 13532 | 13531 | -3741 | 3741 | 13.531 | Shortened | Updated | 13530 | PASS |
| chr25 | 3p | 56629806 | 56630881 | 1075 | 56629806 | 56630881 | 1075 | 0 | 0 | 1.075 | Unchanged | Unchanged (no read-based target) | — | — |
| chr26 | 5p | 1 | 1122 | 1121 | 1 | 9420 | 9419 | 8298 | 8298 | 9.419 | Extended | Updated | 9420 | PASS |
| chr26 | 3p | 61013773 | 61014675 | 902 | 61022072 | 61048865 | 26793 | 25891 | 25891 | 26.793 | Extended | Updated | 26794 | PASS |
| chr27 | 5p | 18 | 2532 | 2514 | 18 | 2532 | 2514 | 0 | 0 | 2.514 | Unchanged | Unchanged (no read-based target) | — | — |
| chr27 | 3p | 60201611 | 60202258 | 647 | 60201611 | 60215355 | 13744 | 13097 | 13097 | 13.744 | Extended | Updated | 13744 | PASS |
| chr28 | 5p | 1 | 15742 | 15741 | 1 | 10545 | 10544 | -5197 | 5197 | 10.544 | Shortened | Updated | 10545 | PASS |
| chr28 | 3p | 68715282 | 68718411 | 3129 | 68710086 | 68732309 | 22223 | 19094 | 19094 | 22.223 | Extended | Updated | 22224 | PASS |
| chr29 | 5p | 1 | 18606 | 18605 | 1 | 18606 | 18605 | 0 | 0 | 18.605 | Unchanged | Unchanged (no read-based target) | — | — |
| chr29 | 3p | 58129550 | 58130961 | 1411 | 58129551 | 58148320 | 18769 | 17358 | 17358 | 18.769 | Extended | Updated | 18770 | PASS |
| chrX | 5p | 1 | 739 | 738 | 1 | 15282 | 15281 | 14543 | 14543 | 15.281 | Extended | Updated | 15282 | PASS |
| chrX | 3p | 171609294 | 171612168 | 2874 | 171623837 | 171644615 | 20778 | 17904 | 17904 | 20.778 | Extended | Updated | 20778 | PASS |

**Summary of coordinate/length changes**

| Total chromosome ends | Coordinates updated | Coordinates unchanged | TL extended | TL shortened | TL unchanged |
|---|---|---|---|---|---|
| 60 | 44 | 16 | 32 | 12 | 16 |

Calibration: 42 of 44 ends with read-based targets agree at <=1 bp with Telogator2 estimates (residual deviations: chr13 3p +1,083 bp; chr19 5p +266 bp). 16 ends lacked valid read-based targets and were left unchanged.

*Note: chr09, chr11 and chr12 3p termini carried mis-scaffolded tails in the pre-correction assembly that were removed during end calibration (see Coordinate status). chr10 3p source tele.bed coordinate was invalid (older assembly version) and the arm was skipped; pre-correction coordinates for chr10 3p are shown in the pre-correction (input) coordinate system derived from the final assembly scan.*

**Supplementary Table 5. Summary statistics for chromosome-end-specific telomere and subtelomeric features in the Holstein_T2T genome, related to Figure 2**

| Analysis | Feature / comparison | 5p termini | 3p termini | Effect / statistic | P value | FDR | Source figure |
|---|---|---|---|---|---|---|---|
| Telomere length | Observed TL range (kb) | 1.08–65.76 | 0.94–34.77 | | | | Figure 2A |
| Telomere length | Chromosomes with longer 3p telomeres | | | 21 of 30 chromosomes | | | Figure 2B |
| Telomere length | Median absolute inter-end difference | | | 7.68 kb | 0.074 | | Figure 2B |
| TVR | Chromosomes with higher TVR abundance at 5p | | | 22 of 30 chromosomes | 6.0E-05 | | Figure 2C–E |
| TVR | TVR abundance vs. mean TL | | | Spearman $\rho$ = −0.405 | 0.026 | | Figure 2D |
| Motif composition | 5p vs. 3p motif-density separation | | | PERMANOVA $R^2$ = 0.178 | 0.001 | | Figure S9H |
| Subtelomere | Mean GC content | 60.65% | 50.64% | 5p > 3p | 1.2E-08 | | Figure 2G |
| Subtelomere | CpG density (sites/kb) | 106.4 | 56.2 | 5p > 3p | 4.8E-07 | | Figure 2H |
| Subtelomere | DNA methylation | Lower | Higher | 3p > 5p | 5.6E-07 | | Figure S10A |
| TL association | GC content, 100–250 kb | | | Negative association with TL | 4.6E-04 | 0.021 | Figure 2I |
| TL association | GC content, 0–500 kb | | | Negative association with TL | 0.0016 | 0.024 | Figure 2I |
| TL association | CpG density, 250–500 kb | | | Negative association with TL | 0.0021 | 0.024 | Figure 2I |
| TL association | DNA methylation at 5p boundary | | | Positive association with TL | 0.0050 | | Figure S10E |
| TL association | DNA methylation 10 kb upstream of 5p terminus | | | Negative association with TL | 0.0078 | | Figure S10E |
| TL association | Methylation vs. TL, chromosome-end level | $\rho$ = 0.08 | $\rho$ = 0.06 | Weak arm-specific trends | | | Figure S10G |

**Supplementary Table 6. Descriptive characteristics of Holstein cows, related to Figure 3.**

| Characteristic | Mean | Median (IQR) | Min-Max |
|---|---|---|---|
| Age (years) | 4.89 | 4.50 (4.00-5.50) | 2.00-13.50 |
| TelSeq estimated telomere length (kb) | 0.603 | 0.559 (0.295-0.838) | 0.024-1.716 |
| Mean sequencing depth across clean autosomal 1-kb bins (×) | 2.14 | 1.05 (0.91-1.41) | 0.42-31.98 |
| Data are from 9591 Holstein cows. IQR, interquartile range. | | | |

**Supplementary Table 7. Sensitivity of bPC adjustment across sequencing-depth strata, related to Figure S11.**

| Cohort | N | Median depth (IQR) | No. of bPCs included | Raw TL-age r | Adjusted TL-age r | Raw TL-depth r | Adjusted TL-depth r |
|---|---|---|---|---|---|---|---|
| Full cohort | 9591 | 1.05 (0.91-1.41) | 121 | 0.093 | -0.057 | -0.059 | -0.008 |
| High depth subset | 1498 | 5.73 (4.98-9.52) | 5 | 0.254 | -0.117 | -0.248 | 0.026 |
| Low depth subset | 8093 | 1.00 (0.89-1.17) | 131 | 0.083 | -0.063 | 0.1 | -0.008 |
| IQR, interquartile range; bPC, batch principal component; TL, telomere length; r, Pearson correlation coefficient. Adjusted TL denotes residuals after bPC correction performed separately within each cohort. | | | | | | | |

**Supplementary Table 8. Association statistics of genome-wide significant telomere length variants across sequencing-depth strata, related to Figures 3 and S12.**

| Chr | SNP | Position | A1 | A2 | Gene | Annotation | P value | | | Effect estimate | | | SE | | | MAF | | |
|---|---|---|---|---|---|---|---|---|---|---|---|---|---|---|---|---|---|---|
| | | | | | | | Full cohort | High depth | Low depth | Full cohort | High depth | Low depth | Full cohort | High depth | Low depth | Full cohort | High depth | Low depth |
| 13 | 13_53114773_G_A | 53114773 | A | G | STK35(dist=88975),PDYN(dist=15462 | intergenic | 6.363E-09 | 0.022679 | 2.329E-07 | -0.028196 | -0.034633 | -0.028143 | 0.0048556 | 0.015198 | 0.0054425 | 0.091805 | 0.077437 | 0.094464 |
| 13 | 13_53228852_C_A | 53228852 | A | C | SIRPA | intronic | 4.761E-08 | 0.063877 | 1.6E-07 | -0.030512 | -0.031534 | -0.03309 | 0.0055883 | 0.017018 | 0.0063141 | 0.067459 | 0.05741 | 0.069319 |
| 13 | 13_53519599_G_A | 53519599 | A | G | LOC536097(dist=1200),GINS1(dist=7 | intergenic | 1.3316E-08 | 0.05896 | 1.44E-07 | 0.027929 | 0.028981 | 0.029073 | 0.0049153 | 0.015346 | 0.005527 | 0.085497 | 0.070093 | 0.088348 |
| 13 | 13_53526947_A_G | 53526947 | G | A | LOC536097(dist=8548),GINS1(dist=6 | intergenic | 4.1718E-08 | 0.08074 | 2.593E-07 | 0.0267 | 0.026676 | 0.028166 | 0.0048692 | 0.015275 | 0.0054681 | 0.084037 | 0.067423 | 0.087112 |
| 13 | 13_53530098_A_G | 53530098 | G | A | LOC536097(dist=11699),GINS1(dist= | intergenic | 1.0442E-08 | 0.047349 | 1.137E-07 | 0.028185 | 0.031564 | 0.029204 | 0.0049245 | 0.015916 | 0.0055068 | 0.083307 | 0.065754 | 0.086556 |
| 13 | 13_53530582_G_A | 53530582 | A | G | LOC536097(dist=12183),GINS1(dist= | intergenic | 2.2716E-08 | 0.10245 | 1.2E-07 | 0.027318 | 0.026416 | 0.028795 | 0.004887 | 0.016176 | 0.0054398 | 0.084819 | 0.067089 | 0.088101 |
| 13 | 13_53530814_T_C | 53530814 | C | T | LOC536097(dist=12415),GINS1(dist= | intergenic | 1.8675E-08 | 0.18983 | 9.968E-08 | 0.028155 | 0.021775 | 0.029644 | 0.0050064 | 0.016609 | 0.0055645 | 0.07726 | 0.062417 | 0.080007 |
| 13 | 13_53530828_G_T | 53530828 | T | G | LOC536097(dist=12429),GINS1(dist= | intergenic | 2.0153E-08 | 0.16233 | 1.303E-07 | 0.028077 | 0.023205 | 0.029355 | 0.0050041 | 0.016608 | 0.0055614 | 0.077104 | 0.062417 | 0.079822 |
| 13 | 13_53530829_G_T | 53530829 | T | G | LOC536097(dist=12430),GINS1(dist= | intergenic | 2.0153E-08 | 0.16233 | 1.303E-07 | 0.028077 | 0.023205 | 0.029355 | 0.0050041 | 0.016608 | 0.0055614 | 0.077104 | 0.062417 | 0.079822 |
| 13 | 13_53530860_T_C | 53530860 | C | T | LOC536097(dist=12461),GINS1(dist= | intergenic | 4.6118E-08 | 0.17586 | 2.29E-07 | 0.027098 | 0.022169 | 0.028526 | 0.0049579 | 0.016377 | 0.0055133 | 0.077625 | 0.063084 | 0.080316 |
| 13 | 13_53530861_G_A | 53530861 | A | G | LOC536097(dist=12462),GINS1(dist= | intergenic | 1.1842E-08 | 0.13217 | 9.266E-08 | 0.026934 | 0.02408 | 0.027974 | 0.0047236 | 0.015994 | 0.0052381 | 0.086435 | 0.067423 | 0.089954 |
| 13 | 13_53531060_A_G | 53531060 | G | A | LOC536097(dist=12661),GINS1(dist= | intergenic | 1.01E-08 | 0.098998 | 5.061E-08 | 0.029909 | 0.027533 | 0.031817 | 0.0052205 | 0.01669 | 0.005839 | 0.074966 | 0.062417 | 0.077289 |
| 13 | 13_53531077_G_A | 53531077 | A | G | LOC536097(dist=12678),GINS1(dist= | intergenic | 7.0124E-09 | 0.10777 | 3.46E-08 | 0.030314 | 0.026949 | 0.0323 | 0.005235 | 0.016756 | 0.0058553 | 0.074653 | 0.061749 | 0.077042 |
| 13 | 13_53531423_C_T | 53531423 | T | C | LOC536097(dist=13024),GINS1(dist= | intergenic | 2.9735E-08 | 0.14339 | 6.718E-08 | -0.030848 | -0.026305 | -0.033585 | 0.0055652 | 0.017976 | 0.0062211 | 0.067928 | 0.054406 | 0.070431 |
| 13 | 13_53531600_T_C | 53531600 | C | T | LOC536097(dist=13201),GINS1(dist= | intergenic | 1.095E-08 | 0.14424 | 2.504E-08 | -0.031534 | -0.026153 | -0.034337 | 0.0055174 | 0.017911 | 0.0061614 | 0.068554 | 0.054072 | 0.071234 |
| 13 | 13_53531663_C_T | 53531663 | T | C | LOC536097(dist=13264),GINS1(dist= | intergenic | 1.7941E-08 | 0.10439 | 3.836E-08 | -0.031287 | -0.029398 | -0.034112 | 0.0055565 | 0.018103 | 0.0062042 | 0.068137 | 0.053405 | 0.070864 |
| 13 | 13_53531823_T_A | 53531823 | A | T | LOC536097(dist=13424),GINS1(dist= | intergenic | 1.8259E-08 | 0.13414 | 5.147E-08 | -0.030745 | -0.02646 | -0.033247 | 0.0054632 | 0.017664 | 0.0061046 | 0.069284 | 0.056075 | 0.071729 |
| 13 | 13_53531844_G_A | 53531844 | A | G | LOC536097(dist=13445),GINS1(dist= | intergenic | 9.7168E-09 | 0.11343 | 2.82E-08 | -0.031539 | -0.028319 | -0.03408 | 0.0054989 | 0.01789 | 0.0061381 | 0.068762 | 0.055073 | 0.071296 |
| 13 | 13_53531916_G_A | 53531916 | A | G | LOC536097(dist=13517),GINS1(dist= | intergenic | 1.9194E-08 | 0.11394 | 7.528E-08 | -0.030847 | -0.027883 | -0.033008 | 0.0054896 | 0.017639 | 0.0061375 | 0.068397 | 0.055407 | 0.070802 |
| 13 | 13_53531983_A_G | 53531983 | G | A | LOC536097(dist=13584),GINS1(dist= | intergenic | 1.9749E-08 | 0.074851 | 7.944E-08 | -0.03063 | -0.031055 | -0.032787 | 0.0054559 | 0.017433 | 0.0061074 | 0.068762 | 0.055407 | 0.071234 |
| 13 | 13_53532018_C_T | 53532018 | T | C | LOC536097(dist=13619),GINS1(dist= | intergenic | 1.2366E-08 | 0.077563 | 6.813E-08 | -0.031071 | -0.031016 | -0.032912 | 0.0054563 | 0.017573 | 0.0060994 | 0.068815 | 0.055073 | 0.071358 |
| 13 | 13_53532022_A_T | 53532022 | T | A | LOC536097(dist=13623),GINS1(dist= | intergenic | 1.1578E-08 | 0.080449 | 6.186E-08 | -0.031149 | -0.030899 | -0.033005 | 0.0054591 | 0.017676 | 0.006097 | 0.068867 | 0.055073 | 0.07142 |
| 13 | 13_53532061_C_T | 53532061 | T | C | LOC536097(dist=13662),GINS1(dist= | intergenic | 6.7238E-09 | 0.11602 | 4.708E-08 | -0.031556 | -0.027625 | -0.033217 | 0.0054429 | 0.017577 | 0.0060814 | 0.068971 | 0.055073 | 0.071543 |
| 13 | 13_53532073_C_T | 53532073 | T | C | LOC536097(dist=13674),GINS1(dist= | intergenic | 7.927E-09 | 0.12325 | 4.741E-08 | -0.031369 | -0.026817 | -0.033221 | 0.0054366 | 0.017399 | 0.0060836 | 0.068971 | 0.055741 | 0.07142 |
| 13 | 13_53532187_T_C | 53532187 | C | T | LOC536097(dist=13788),GINS1(dist= | intergenic | 6.1477E-09 | 0.10253 | 2.479E-08 | -0.03175 | -0.02927 | -0.033929 | 0.0054622 | 0.017927 | 0.0060862 | 0.068606 | 0.053071 | 0.071482 |
| 13 | 13_53532234_C_T | 53532234 | T | C | LOC536097(dist=13835),GINS1(dist= | intergenic | 1.0439E-08 | 0.15571 | 2.74E-08 | -0.031182 | -0.025276 | -0.033756 | 0.0054481 | 0.017805 | 0.0060743 | 0.069023 | 0.054072 | 0.07179 |
| 13 | 13_53532589_G_C | 53532589 | C | G | LOC536097(dist=14190),GINS1(dist= | intergenic | 3.1471E-08 | 0.19016 | 1.174E-07 | -0.030237 | -0.023198 | -0.032321 | 0.0054648 | 0.017707 | 0.0061011 | 0.067668 | 0.055741 | 0.069875 |
| 13 | 13_53532615_T_C | 53532615 | C | T | LOC536097(dist=14216),GINS1(dist= | intergenic | 2.8698E-08 | 0.16855 | 1.103E-07 | -0.03036 | -0.024548 | -0.032395 | 0.0054711 | 0.017829 | 0.0061019 | 0.067459 | 0.05474 | 0.069813 |
| 13 | 13_53532660_C_T | 53532660 | T | C | LOC536097(dist=14261),GINS1(dist= | intergenic | 1.5723E-08 | 0.085443 | 9.462E-08 | -0.030906 | -0.03087 | -0.032503 | 0.0054667 | 0.017948 | 0.0060903 | 0.067459 | 0.053738 | 0.069999 |
| 13 | 13_53532740_T_C | 53532740 | C | T | LOC536097(dist=14341),GINS1(dist= | intergenic | 3.3319E-08 | 0.14792 | 1.001E-07 | -0.030217 | -0.026233 | -0.032417 | 0.0054712 | 0.01813 | 0.0060859 | 0.068815 | 0.054072 | 0.071543 |
| 13 | 13_53532920_T_G | 53532920 | G | T | LOC536097(dist=14521),GINS1(dist= | intergenic | 3.6051E-08 | 0.25343 | 7.885E-08 | -0.030227 | -0.020509 | -0.032842 | 0.0054867 | 0.017958 | 0.0061162 | 0.068658 | 0.053738 | 0.07142 |
| 13 | 13_53533158_A_G | 53533158 | G | A | LOC536097(dist=14759),GINS1(dist= | intergenic | 2.7696E-08 | 0.13431 | 1.393E-07 | -0.030175 | -0.026569 | -0.031905 | 0.0054317 | 0.017744 | 0.0060583 | 0.069805 | 0.055407 | 0.07247 |
| 13 | 13_53533265_G_T | 53533265 | T | G | LOC536097(dist=14866),GINS1(dist= | intergenic | 1.8968E-08 | 0.051909 | 1.28E-07 | 0.027247 | 0.029615 | 0.028755 | 0.0048472 | 0.015235 | 0.0054442 | 0.085497 | 0.06976 | 0.08841 |
| 13 | 13_53533287_G_A | 53533287 | A | G | LOC536097(dist=14888),GINS1(dist= | intergenic | 3.7765E-08 | 0.1733 | 1.369E-07 | -0.030048 | -0.023696 | -0.03224 | 0.0054623 | 0.017402 | 0.0061184 | 0.069179 | 0.055741 | 0.071667 |
| 13 | 13_53533348_C_T | 53533348 | T | C | LOC536097(dist=14949),GINS1(dist= | intergenic | 3.4368E-08 | 0.15019 | 1.137E-07 | -0.030224 | -0.024662 | -0.032643 | 0.0054778 | 0.01714 | 0.0061551 | 0.069232 | 0.058077 | 0.071296 |
| 13 | 13_53533381_C_A | 53533381 | A | C | LOC536097(dist=14982),GINS1(dist= | intergenic | 4.6352E-08 | 0.13968 | 1.811E-07 | -0.029783 | -0.025766 | -0.031821 | 0.0054501 | 0.017445 | 0.0060986 | 0.069544 | 0.057076 | 0.071852 |
| 13 | 13_53533458_T_G | 53533458 | G | T | LOC536097(dist=15059),GINS1(dist= | intergenic | 4.8852E-08 | 0.12884 | 1.668E-07 | -0.02985 | -0.026948 | -0.031966 | 0.0054716 | 0.017744 | 0.0061086 | 0.069023 | 0.055407 | 0.071543 |
| 13 | 13_53534463_T_C | 53534463 | C | T | LOC536097(dist=16064),GINS1(dist= | intergenic | 3.9367E-08 | 0.10326 | 1.289E-07 | -0.030769 | -0.029903 | -0.032974 | 0.0056009 | 0.018354 | 0.0062447 | 0.067407 | 0.054072 | 0.069875 |
| 13 | 13_53534689_A_G | 53534689 | G | A | LOC536097(dist=16290),GINS1(dist= | intergenic | 3.4087E-08 | 0.074417 | 1.28E-07 | -0.030608 | -0.032255 | -0.032698 | 0.0055459 | 0.01808 | 0.0061907 | 0.068293 | 0.054406 | 0.070864 |
| 13 | 13_53544541_A_G | 53544541 | G | A | LOC536097(dist=26142),GINS1(dist= | intergenic | 2.639E-08 | 0.060325 | 3.729E-07 | -0.028805 | -0.032063 | -0.029317 | 0.0051771 | 0.017069 | 0.0057685 | 0.076999 | 0.060414 | 0.080069 |
| 13 | 13_53544694_G_A | 53544694 | A | G | LOC536097(dist=26295),GINS1(dist= | intergenic | 1.27E-08 | 0.047639 | 1.147E-07 | -0.029381 | -0.034785 | -0.030389 | 0.0051635 | 0.017563 | 0.0057319 | 0.076791 | 0.058411 | 0.080193 |
| 13 | 13_53544923_C_T | 53544923 | T | C | LOC536097(dist=26524),GINS1(dist= | intergenic | 1.9686E-08 | 0.072837 | 8.701E-08 | -0.028779 | -0.03075 | -0.030499 | 0.0051255 | 0.017142 | 0.0056986 | 0.077468 | 0.059746 | 0.080749 |
| 13 | 13_53544994_T_A | 53544994 | A | T | LOC536097(dist=26595),GINS1(dist= | intergenic | 2.0125E-08 | 0.054426 | 1.044E-07 | -0.028701 | -0.033363 | -0.030198 | 0.0051152 | 0.017345 | 0.0056775 | 0.077156 | 0.058745 | 0.080563 |
| 13 | 13_53544997_A_G | 53544997 | G | A | LOC536097(dist=26598),GINS1(dist= | intergenic | 2.0125E-08 | 0.054426 | 1.044E-07 | -0.028701 | -0.033363 | -0.030198 | 0.0051152 | 0.017345 | 0.0056775 | 0.077156 | 0.058745 | 0.080563 |
| 13 | 13_53545084_A_C | 53545084 | C | A | LOC536097(dist=26685),GINS1(dist= | intergenic | 1.2207E-08 | 0.073744 | 4.681E-08 | -0.029296 | -0.031022 | -0.031193 | 0.0051425 | 0.017348 | 0.0057098 | 0.076947 | 0.058745 | 0.080316 |
| 13 | 13_53545151_A_G | 53545151 | G | A | LOC536097(dist=26752),GINS1(dist= | intergenic | 1.1979E-08 | 0.042935 | 6.796E-08 | -0.029236 | -0.035449 | -0.030705 | 0.005129 | 0.017511 | 0.0056898 | 0.077156 | 0.058077 | 0.080687 |
| 13 | 13_53545179_A_G | 53545179 | G | A | LOC536097(dist=26780),GINS1(dist= | intergenic | 1.853E-08 | 0.055941 | 5.625E-08 | -0.028824 | -0.033156 | -0.030886 | 0.0051241 | 0.017346 | 0.0056877 | 0.077312 | 0.058745 | 0.080749 |
| 13 | 13_53545189_T_C | 53545189 | C | T | LOC536097(dist=26790),GINS1(dist= | intergenic | 2.0497E-08 | 0.062812 | 5.976E-08 | -0.028775 | -0.032529 | -0.030837 | 0.0051313 | 0.017484 | 0.00569 | 0.077156 | 0.058411 | 0.080625 |
| 13 | 13_53555744_G_A | 53555744 | A | G | LOC536097(dist=37345),GINS1(dist= | intergenic | 4.9571E-08 | 0.066006 | 2.165E-07 | -0.028561 | -0.032301 | -0.030186 | 0.0052379 | 0.01757 | 0.0058222 | 0.076791 | 0.059413 | 0.080007 |
| 13 | 13_53557202_A_C | 53557202 | C | A | LOC536097(dist=38803),GINS1(dist= | intergenic | 4.9423E-08 | 0.10516 | 1.521E-07 | -0.028413 | -0.026991 | -0.030644 | 0.0052102 | 0.016658 | 0.0058369 | 0.077833 | 0.06275 | 0.080625 |
| 13 | 13_53560159_A_G | 53560159 | G | A | LOC536097(dist=41760),GINS1(dist= | intergenic | 3.6792E-08 | 0.077203 | 2.617E-07 | -0.028774 | -0.030968 | -0.029918 | 0.0052263 | 0.017524 | 0.0058103 | 0.077312 | 0.058077 | 0.080872 |
| 13 | 13_53560417_C_T | 53560417 | T | C | LOC536097(dist=42018),GINS1(dist= | intergenic | 1.9292E-08 | 0.045432 | 1.4E-07 | -0.029226 | -0.034346 | -0.030557 | 0.005202 | 0.017168 | 0.0058036 | 0.077521 | 0.059413 | 0.080872 |
| 13 | 13_53567108_C_G | 53567108 | G | C | LOC536097(dist=48709),GINS1(dist= | intergenic | 4.6913E-08 | 0.12585 | 2.26E-07 | 0.027322 | 0.025049 | 0.028889 | 0.0050016 | 0.016365 | 0.0055806 | 0.083047 | 0.064085 | 0.086556 |
| 13 | 13_53570771_G_T | 53570771 | T | G | LOC536097(dist=52372),GINS1(dist= | intergenic | 1.4612E-08 | 0.1411 | 6.412E-08 | -0.029806 | -0.025237 | -0.03177 | 0.0052605 | 0.017148 | 0.0058758 | 0.076999 | 0.059746 | 0.080193 |
| 13 | 13_53571665_C_A | 53571665 | A | C | LOC536097(dist=53266),GINS1(dist= | intergenic | 4.725E-08 | 0.083869 | 1.586E-07 | -0.028549 | -0.029635 | -0.030575 | 0.0052275 | 0.017143 | 0.0058324 | 0.077833 | 0.059746 | 0.081181 |
| 13 | 13_53572928_A_G | 53572928 | G | A | LOC536097(dist=54529),GINS1(dist= | intergenic | 4.7359E-08 | 0.070417 | 2.313E-07 | -0.028564 | -0.03162 | -0.030085 | 0.0052307 | 0.017477 | 0.0058166 | 0.077364 | 0.056742 | 0.081181 |
| 13 | 13_53573372_C_G | 53573372 | G | C | LOC536097(dist=54973),GINS1(dist= | intergenic | 3.1944E-08 | 0.086347 | 1.098E-07 | -0.029626 | -0.030946 | -0.031603 | 0.0053569 | 0.018044 | 0.005952 | 0.075279 | 0.056409 | 0.078772 |
| 13 | 13_53585197_G_A | 53585197 | A | G | LOC536097(dist=66798),GINS1(dist= | intergenic | 3.9077E-08 | 0.044505 | 1.698E-07 | 0.027227 | 0.032731 | 0.028899 | 0.0049549 | 0.01629 | 0.0055259 | 0.083881 | 0.065754 | 0.087236 |
| 13 | 13_53588229_C_T | 53588229 | T | C | LOC536097(dist=69830),GINS1(dist= | intergenic | 3.6129E-08 | 0.089116 | 1.296E-07 | 0.027428 | 0.026306 | 0.029592 | 0.004979 | 0.015473 | 0.0056051 | 0.083099 | 0.068425 | 0.085815 |
| 13 | 13_53605166_A_G | 53605166 | G | A | GINS1(NM_001083701.2:c.*501A>G, | UTR3 | 2.9869E-08 | 0.07989 | 2.099E-07 | -0.02775 | -0.029978 | -0.028776 | 0.005007 | 0.017117 | 0.0055442 | 0.077208 | 0.061749 | 0.080069 |
| 13 | 13_53605826_A_G | 53605826 | G | A | GINS1(NM_001083701.2:c.*1161A>G | UTR3 | 9.7145E-09 | 0.05635 | 5.611E-08 | -0.028764 | -0.032777 | -0.030156 | 0.0050149 | 0.017176 | 0.0055528 | 0.077364 | 0.059413 | 0.080687 |
| 13 | 13_53605853_G_A | 53605853 | A | G | GINS1(NM_001083701.2:c.*1188G>A | UTR3 | 1.2106E-08 | 0.036096 | 6.603E-08 | -0.02845 | -0.035342 | -0.029952 | 0.0049928 | 0.016863 | 0.005545 | 0.077781 | 0.061415 | 0.080811 |
| 13 | 13_53605906_C_T | 53605906 | T | C | GINS1(NM_001083701.2:c.*1241C>T | UTR3 | 4.8723E-09 | 0.039122 | 2.549E-08 | -0.029204 | -0.035495 | -0.030777 | 0.0049908 | 0.017206 | 0.0055255 | 0.07799 | 0.060748 | 0.081181 |
| 13 | 13_53605914_G_A | 53605914 | A | G | GINS1(NM_001083701.2:c.*1249G>A | UTR3 | 7.1876E-09 | 0.073758 | 2.914E-08 | -0.028937 | -0.030757 | -0.030699 | 0.0050008 | 0.017201 | 0.0055348 | 0.077729 | 0.06008 | 0.080996 |
| 13 | 13_53605916_C_G | 53605916 | G | C | GINS1(NM_001083701.2:c.*1251C>G | UTR3 | 5.6615E-09 | 0.074072 | 2.271E-08 | -0.029144 | -0.030789 | -0.030935 | 0.005002 | 0.017237 | 0.0055345 | 0.077677 | 0.059746 | 0.080996 |
| 13 | 13_53605919_C_T | 53605919 | T | C | GINS1(NM_001083701.2:c.*1254C>T | UTR3 | 7.997E-09 | 0.10455 | 3.137E-08 | -0.028886 | -0.027985 | -0.030669 | 0.0050076 | 0.017241 | 0.0055424 | 0.077677 | 0.059746 | 0.080996 |
| 13 | 13_53606008_G_C | 53606008 | C | G | GINS1(NM_001083701.2:c.*1343G>C | UTR3 | 2.212E-08 | 0.082142 | 7.695E-08 | -0.028092 | -0.029517 | -0.029945 | 0.0050213 | 0.01698 | 0.0055721 | 0.07799 | 0.060414 | 0.081243 |
| 13 | 13_53606040_A_G | 53606040 | G | A | GINS1(NM_001083701.2:c.*1375A>G | UTR3 | 3.4605E-08 | 0.088797 | 1.16E-07 | -0.028049 | -0.029424 | -0.029876 | 0.0050846 | 0.01729 | 0.0056373 | 0.076947 | 0.058411 | 0.080378 |
| 13 | 13_53606054_T_C | 53606054 | C | T | GINS1(NM_001083701.2:c.*1389T>C | UTR3 | 4.1271E-08 | 0.090027 | 1.377E-07 | -0.02799 | -0.029148 | -0.029845 | 0.0051027 | 0.017194 | 0.0056652 | 0.076478 | 0.058411 | 0.079822 |

| | | | | | | | | | | | | | | | | | | |
|---|---|---|---|---|---|---|---|---|---|---|---|---|---|---|---|---|---|---|
| 13 | 13_53606055_G_C | 53606055 | C | G | GINS1(NM_001083701.2:c.*1390G>C | UTR3 | 4.1271E-08 | 0.090027 | 1.377E-07 | -0.02799 | -0.029148 | -0.029845 | 0.0051027 | 0.017194 | 0.0056652 | 0.076478 | 0.058411 | 0.079822 |
| 13 | 13_53606127_C_T | 53606127 | T | C | GINS1(NM_001083701.2:c.*1462C>T | UTR3 | 3.8542E-08 | 0.15233 | 8.523E-08 | -0.028287 | -0.025049 | -0.030556 | 0.0051455 | 0.017501 | 0.0057054 | 0.075748 | 0.055741 | 0.079451 |
| 13 | 13_53606138_T_G | 53606138 | G | T | GINS1(NM_001083701.2:c.*1473T>G | UTR3 | 3.2443E-08 | 0.11453 | 5.656E-08 | -0.028463 | -0.027492 | -0.031019 | 0.0051491 | 0.017421 | 0.0057133 | 0.075748 | 0.056409 | 0.079328 |
| 13 | 13_53606419_G_A | 53606419 | A | G | GINS1(NM_001083701.2:c.*1754G>A | UTR3 | 4.9524E-08 | 0.057044 | 2.295E-07 | -0.027926 | -0.032617 | -0.029459 | 0.0051213 | 0.01714 | 0.005694 | 0.077364 | 0.059746 | 0.080625 |
| 13 | 13_53606507_A_T | 53606507 | T | A | GINS1(NM_001083701.2:c.*1842A>T | UTR3 | 4.9672E-08 | 0.081419 | 2.074E-07 | -0.027902 | -0.029534 | -0.029599 | 0.0051172 | 0.016949 | 0.0057001 | 0.076686 | 0.058077 | 0.080131 |
| 13 | 13_53610536_A_G | 53610536 | G | A | GINS1(dist=3881),PCMTD2(dist=4027 | intergenic | 3.6333E-08 | 0.11778 | 2.107E-07 | -0.028564 | -0.026764 | -0.029988 | 0.0051862 | 0.017111 | 0.0057783 | 0.076739 | 0.06008 | 0.079822 |
| 13 | 13_53610561_T_C | 53610561 | C | T | GINS1(dist=3906),PCMTD2(dist=4024 | intergenic | 3.816E-08 | 0.062564 | 3.129E-07 | 0.027333 | 0.029825 | 0.028431 | 0.0049704 | 0.016015 | 0.0055578 | 0.082212 | 0.067089 | 0.085012 |
| 13 | 13_53620018_G_A | 53620018 | A | G | GINS1(dist=13363),PCMTD2(dist=307 | intergenic | 4.2816E-08 | 0.076173 | 4.22E-07 | -0.028623 | -0.031145 | -0.029374 | 0.0052242 | 0.017563 | 0.0058066 | 0.076791 | 0.057744 | 0.080316 |
| 13 | 13_53620558_G_A | 53620558 | A | G | GINS1(dist=13903),PCMTD2(dist=302 | intergenic | 2.6695E-08 | 0.038112 | 5.47E-07 | -0.028334 | -0.034132 | -0.028554 | 0.0050944 | 0.01646 | 0.0057005 | 0.077521 | 0.061081 | 0.080563 |
| 13 | 13_53620629_C_G | 53620629 | G | C | GINS1(dist=13974),PCMTD2(dist=301 | intergenic | 4.43E-08 | 0.084641 | 4.202E-07 | -0.028598 | -0.029982 | -0.029426 | 0.0052255 | 0.017387 | 0.005816 | 0.076113 | 0.058411 | 0.07939 |
| 13 | 13_53621066_A_C | 53621066 | C | A | GINS1(dist=14411),PCMTD2(dist=297 | intergenic | 3.4107E-08 | 0.092906 | 1.954E-07 | -0.028822 | -0.028868 | -0.030294 | 0.0052223 | 0.01718 | 0.0058217 | 0.076113 | 0.059413 | 0.079204 |
| 13 | 13_53636865_T_C | 53636865 | C | T | GINS1(dist=30210),PCMTD2(dist=139 | intergenic | 8.3725E-10 | 0.0095409 | 2.369E-08 | 0.026852 | 0.036181 | 0.027298 | 0.0043749 | 0.013958 | 0.0048898 | 0.111146 | 0.093792 | 0.114358 |
| 13 | 13_53637587_T_C | 53637587 | C | T | GINS1(dist=30932),PCMTD2(dist=132 | intergenic | 1.4645E-08 | 0.057138 | 8.821E-08 | -0.029154 | -0.032673 | -0.030618 | 0.0051458 | 0.017176 | 0.0057236 | 0.078824 | 0.059413 | 0.082417 |
| 13 | 13_53637650_G_A | 53637650 | A | G | GINS1(dist=30995),PCMTD2(dist=131 | intergenic | 3.207E-08 | 0.041618 | 1.929E-07 | 0.027198 | 0.031945 | 0.02871 | 0.0049185 | 0.01568 | 0.0055147 | 0.083933 | 0.067757 | 0.086927 |
| 13 | 13_53637726_T_C | 53637726 | C | T | GINS1(dist=31071),PCMTD2(dist=130 | intergenic | 1.8562E-08 | 0.032648 | 1.353E-07 | 0.027408 | 0.032814 | 0.028855 | 0.0048726 | 0.01536 | 0.0054737 | 0.084923 | 0.070093 | 0.087668 |
| 13 | 13_53637782_A_C | 53637782 | C | A | GINS1(dist=31127),PCMTD2(dist=130 | intergenic | 4.7126E-08 | 0.062599 | 1.144E-07 | 0.026995 | 0.029835 | 0.029285 | 0.0049424 | 0.016023 | 0.0055232 | 0.083776 | 0.066088 | 0.087051 |
| 13 | 13_53639323_C_T | 53639323 | T | C | GINS1(dist=32668),PCMTD2(dist=114 | intergenic | 7.7927E-09 | 0.0089758 | 3.51E-07 | 0.024824 | 0.037328 | 0.024308 | 0.0043001 | 0.014286 | 0.004772 | 0.111198 | 0.090454 | 0.115038 |
| 13 | 13_53639613_T_C | 53639613 | C | T | GINS1(dist=32958),PCMTD2(dist=111 | intergenic | 1.176E-08 | 0.0052181 | 3.263E-07 | 0.024273 | 0.038783 | 0.024194 | 0.004256 | 0.013885 | 0.0047369 | 0.112293 | 0.092123 | 0.116026 |
| 13 | 13_53640745_T_C | 53640745 | C | T | GINS1(dist=34090),PCMTD2(dist=100 | intergenic | 2.0526E-09 | 0.0008826 | 1.057E-07 | 0.026053 | 0.048901 | 0.025565 | 0.0043468 | 0.014705 | 0.0048085 | 0.107705 | 0.081776 | 0.112505 |
| 13 | 13_53641138_C_T | 53641138 | T | C | GINS1(dist=34483),PCMTD2(dist=967 | intergenic | 1.9319E-08 | 0.076067 | 1.606E-07 | -0.028496 | -0.029937 | -0.02958 | 0.0050723 | 0.016876 | 0.005645 | 0.078511 | 0.061415 | 0.081676 |
| 13 | 13_53641213_A_T | 53641213 | T | A | GINS1(dist=34558),PCMTD2(dist=959 | intergenic | 2.9568E-08 | 0.066323 | 1.577E-07 | -0.02817 | -0.031559 | -0.029571 | 0.0050811 | 0.017187 | 0.0056398 | 0.078615 | 0.061081 | 0.081861 |
| 13 | 13_53641235_T_G | 53641235 | G | T | GINS1(dist=34580),PCMTD2(dist=957 | intergenic | 2.9998E-08 | 0.087581 | 1.437E-07 | -0.02816 | -0.029489 | -0.029653 | 0.0050818 | 0.017262 | 0.0056371 | 0.078563 | 0.060414 | 0.081923 |
| 13 | 13_53641304_G_A | 53641304 | A | G | GINS1(dist=34649),PCMTD2(dist=950 | intergenic | 1.5074E-08 | 0.096955 | 5.929E-08 | -0.028628 | -0.028593 | -0.030403 | 0.0050572 | 0.017227 | 0.0056085 | 0.079606 | 0.060748 | 0.083097 |
| 13 | 13_53641443_G_A | 53641443 | A | G | GINS1(dist=34788),PCMTD2(dist=936 | intergenic | 9.7524E-09 | 0.074195 | 4.311E-08 | -0.029275 | -0.031192 | -0.030995 | 0.0051047 | 0.017471 | 0.0056585 | 0.078146 | 0.059413 | 0.081614 |
| 13 | 13_53641750_C_A | 53641750 | A | C | GINS1(dist=35095),PCMTD2(dist=905 | intergenic | 5.892E-09 | 0.065879 | 4.143E-08 | -0.029729 | -0.032018 | -0.031075 | 0.0051083 | 0.017408 | 0.0056658 | 0.078615 | 0.059079 | 0.082232 |
| 13 | 13_53641824_C_G | 53641824 | G | C | GINS1(dist=35169),PCMTD2(dist=898 | intergenic | 9.1043E-09 | 0.046424 | 6.242E-08 | -0.029331 | -0.034229 | -0.030719 | 0.005104 | 0.017187 | 0.0056763 | 0.07872 | 0.061081 | 0.081984 |
| 13 | 13_53642001_C_A | 53642001 | A | C | GINS1(dist=35346),PCMTD2(dist=880 | intergenic | 4.0637E-08 | 0.032716 | 4.033E-07 | -0.028026 | -0.03645 | -0.028796 | 0.0051068 | 0.017068 | 0.0056825 | 0.078511 | 0.062083 | 0.081552 |
| 13 | 13_53642005_T_C | 53642005 | C | T | GINS1(dist=35350),PCMTD2(dist=880 | intergenic | 3.929E-08 | 0.032716 | 3.854E-07 | -0.028065 | -0.03645 | -0.028854 | 0.0051083 | 0.017068 | 0.0056844 | 0.078459 | 0.062083 | 0.08149 |
| 13 | 13_53642015_T_C | 53642015 | C | T | GINS1(dist=35360),PCMTD2(dist=879 | intergenic | 3.2546E-08 | 0.032608 | 3.636E-07 | -0.028197 | -0.0362 | -0.028905 | 0.0051015 | 0.01694 | 0.005682 | 0.078563 | 0.062417 | 0.081552 |
| 13 | 13_53642052_C_T | 53642052 | T | C | GINS1(dist=35397),PCMTD2(dist=875 | intergenic | 3.4499E-08 | 0.037444 | 3.375E-07 | -0.02826 | -0.036075 | -0.029027 | 0.0051225 | 0.017336 | 0.0056903 | 0.07825 | 0.060414 | 0.081552 |
| 13 | 13_53642126_G_A | 53642126 | A | G | GINS1(dist=35471),PCMTD2(dist=868 | intergenic | 2.2938E-08 | 0.068775 | 1.331E-07 | -0.028589 | -0.031328 | -0.029983 | 0.005116 | 0.017214 | 0.0056845 | 0.078094 | 0.059079 | 0.081614 |
| 13 | 13_53642409_C_G | 53642409 | G | C | GINS1(dist=35754),PCMTD2(dist=839 | intergenic | 1.2365E-08 | 0.040627 | 8.241E-08 | -0.029016 | -0.035786 | -0.030294 | 0.0050954 | 0.017479 | 0.0056501 | 0.078563 | 0.06008 | 0.081984 |
| 13 | 13_53642465_C_T | 53642465 | T | C | GINS1(dist=35810),PCMTD2(dist=834 | intergenic | 3.5987E-08 | 0.072001 | 1.845E-07 | -0.027987 | -0.030484 | -0.029476 | 0.0050799 | 0.016944 | 0.005653 | 0.078668 | 0.060748 | 0.081984 |
| 13 | 13_53642468_G_C | 53642468 | C | G | GINS1(dist=35813),PCMTD2(dist=834 | intergenic | 3.1361E-08 | 0.070527 | 1.845E-07 | -0.028101 | -0.03058 | -0.029476 | 0.0050782 | 0.016909 | 0.005653 | 0.07872 | 0.061081 | 0.081984 |
| 13 | 13_53642479_G_C | 53642479 | C | G | GINS1(dist=35824),PCMTD2(dist=832 | intergenic | 4.2951E-08 | 0.072001 | 2.007E-07 | -0.027813 | -0.030484 | -0.029369 | 0.0050769 | 0.016944 | 0.0056493 | 0.078668 | 0.060748 | 0.081984 |
| 13 | 13_53642485_G_A | 53642485 | A | G | GINS1(dist=35830),PCMTD2(dist=832 | intergenic | 2.4889E-08 | 0.072826 | 1.132E-07 | 0.027317 | 0.028798 | 0.028997 | 0.0049007 | 0.016053 | 0.0054668 | 0.083099 | 0.065754 | 0.086309 |
| 13 | 13_53642496_G_A | 53642496 | A | G | GINS1(dist=35841),PCMTD2(dist=831 | intergenic | 4.3882E-08 | 0.05756 | 2.817E-07 | -0.027803 | -0.0328 | -0.028937 | 0.0050787 | 0.017272 | 0.005635 | 0.078615 | 0.059413 | 0.08217 |
| 13 | 13_53642498_T_C | 53642498 | C | T | GINS1(dist=35843),PCMTD2(dist=831 | intergenic | 4.8444E-08 | 0.064838 | 2.817E-07 | -0.027707 | -0.031824 | -0.028937 | 0.0050774 | 0.017236 | 0.005635 | 0.078668 | 0.059746 | 0.08217 |
| 13 | 13_53642558_C_T | 53642558 | T | C | GINS1(dist=35903),PCMTD2(dist=825 | intergenic | 2.5945E-08 | 0.043727 | 2.041E-07 | -0.028432 | -0.035212 | -0.029436 | 0.0051074 | 0.01746 | 0.0056656 | 0.078355 | 0.059413 | 0.081861 |
| 13 | 13_53642576_T_C | 53642576 | C | T | GINS1(dist=35921),PCMTD2(dist=823 | intergenic | 2.7779E-08 | 0.04251 | 2.229E-07 | -0.028401 | -0.035782 | -0.029335 | 0.0051129 | 0.01764 | 0.0056641 | 0.078303 | 0.058745 | 0.081923 |
| 13 | 13_53642829_C_T | 53642829 | T | C | GINS1(dist=36174),PCMTD2(dist=797 | intergenic | 1.473E-08 | 0.021777 | 1.959E-07 | -0.025548 | -0.035065 | -0.025973 | 0.00451 | 0.015284 | 0.0049917 | 0.101762 | 0.080107 | 0.10577 |
| 13 | 13_53643691_C_A | 53643691 | A | C | GINS1(dist=37036),PCMTD2(dist=711 | intergenic | 2.7532E-08 | 0.0039625 | 7.892E-07 | 0.024292 | 0.041427 | 0.023985 | 0.0043718 | 0.014379 | 0.0048573 | 0.107288 | 0.086782 | 0.111084 |
| 13 | 13_53645331_T_C | 53645331 | C | T | GINS1(dist=38676),PCMTD2(dist=547 | intergenic | 5.3052E-09 | 0.0049351 | 1.672E-07 | 0.025469 | 0.03932 | 0.025492 | 0.0043632 | 0.013987 | 0.0048718 | 0.108591 | 0.09012 | 0.11201 |
| 13 | 13_53646768_T_G | 53646768 | G | T | GINS1(dist=40113),PCMTD2(dist=404 | intergenic | 1.0175E-08 | 0.0088262 | 4.03E-07 | 0.024935 | 0.036682 | 0.02461 | 0.0043534 | 0.014008 | 0.0048564 | 0.108383 | 0.089786 | 0.111825 |
| 13 | 13_53649336_G_A | 53649336 | A | G | GINS1(dist=42681),PCMTD2(dist=147 | intergenic | 1.9592E-09 | 0.007313 | 1.092E-07 | 0.026506 | 0.03701 | 0.026314 | 0.0044169 | 0.013798 | 0.0049549 | 0.107497 | 0.090788 | 0.110589 |
| 13 | 13_53649923_A_C | 53649923 | C | A | PCMTD2(dist=885) | downstream | 2.0997E-08 | 0.011081 | 8.557E-07 | 0.025997 | 0.037525 | 0.025497 | 0.0046394 | 0.014773 | 0.0051799 | 0.097227 | 0.081776 | 0.100086 |
| 13 | 13_53649948_A_T | 53649948 | T | A | PCMTD2(dist=860) | downstream | 2.8892E-08 | 0.01191 | 8.78E-07 | 0.025652 | 0.037044 | 0.025382 | 0.0046237 | 0.01473 | 0.005162 | 0.097591 | 0.082443 | 0.100395 |
| 13 | 13_53649960_A_G | 53649960 | G | A | PCMTD2(dist=848) | downstream | 4.8377E-08 | 0.010187 | 1.61E-06 | 0.02528 | 0.03805 | 0.024793 | 0.0046324 | 0.014809 | 0.0051683 | 0.097227 | 0.082109 | 0.100025 |
| 13 | 13_53650207_A_T | 53650207 | T | A | PCMTD2(dist=601) | downstream | 3.2876E-08 | 0.0065126 | 1E-06 | 0.024222 | 0.039331 | 0.023822 | 0.0043837 | 0.014456 | 0.0048699 | 0.108174 | 0.086449 | 0.112196 |
| 13 | 13_53651003_T_C | 53651003 | C | T | PCMTD2(XM_025000384.1:c.*2172A> | UTR3 | 2.7674E-08 | 0.0070025 | 8.699E-07 | 0.023968 | 0.03738 | 0.023681 | 0.0043142 | 0.013861 | 0.0048141 | 0.11026 | 0.090788 | 0.113864 |
| 13 | 13_53651989_G_A | 53651989 | A | G | PCMTD2(XM_025000384.1:c.*1186C> | UTR3 | 2.6311E-08 | 0.0058866 | 8.893E-07 | 0.024292 | 0.038663 | 0.023937 | 0.0043656 | 0.014039 | 0.0048705 | 0.109426 | 0.09012 | 0.112999 |
| 13 | 13_53653035_C_G | 53653035 | G | C | PCMTD2(XM_025000384.1:c.*140G> | UTR3 | 2.4939E-09 | 0.0072271 | 5.605E-08 | 0.026521 | 0.038661 | 0.026939 | 0.0044485 | 0.014393 | 0.0049603 | 0.107027 | 0.08745 | 0.110651 |
| 13 | 13_53654392_G_T | 53654392 | T | G | PCMTD2 | intronic | 4.2092E-09 | 0.0038759 | 2.399E-07 | 0.025792 | 0.041413 | 0.025231 | 0.0043895 | 0.014339 | 0.0048845 | 0.108487 | 0.088451 | 0.112196 |
| 13 | 13_53654658_G_A | 53654658 | A | G | PCMTD2 | intronic | 1.368E-09 | 0.0033267 | 9.352E-08 | -0.027614 | -0.042531 | -0.027213 | 0.0045573 | 0.014487 | 0.0050971 | 0.102961 | 0.083778 | 0.106512 |
| 13 | 13_53655043_C_A | 53655043 | A | C | PCMTD2 | intronic | 7.7073E-09 | 0.0047824 | 8.739E-07 | 0.025083 | 0.040125 | 0.023766 | 0.0043436 | 0.014222 | 0.0048323 | 0.109321 | 0.088451 | 0.113184 |
| 13 | 13_53655940_G_A | 53655940 | A | G | PCMTD2 | intronic | 7.1056E-09 | 0.0047263 | 5.152E-07 | 0.02524 | 0.039934 | 0.024397 | 0.0043604 | 0.014135 | 0.0048594 | 0.109061 | 0.089119 | 0.112752 |
| 13 | 13_53656779_A_T | 53656779 | T | A | PCMTD2 | intronic | 6.0314E-09 | 0.0037353 | 4.07E-07 | 0.025408 | 0.040917 | 0.024672 | 0.0043688 | 0.014111 | 0.0048704 | 0.109321 | 0.089786 | 0.112937 |
| 13 | 13_53658224_C_T | 53658224 | T | C | PCMTD2 | intronic | 4.0333E-09 | 0.014242 | 1.258E-07 | -0.028264 | -0.037789 | -0.028316 | 0.0048046 | 0.015417 | 0.0053579 | 0.094151 | 0.078104 | 0.097121 |
| 13 | 13_53659897_C_T | 53659897 | C | T | PCMTD2 | intronic | 1.1227E-09 | 0.0077364 | 9.966E-08 | 0.027254 | 0.038143 | 0.026637 | 0.0044745 | 0.014321 | 0.005 | 0.10635 | 0.088785 | 0.109601 |
| 13 | 13_53664907_A_G | 53664907 | G | A | PCMTD2 | intronic | 4.6105E-09 | 0.0055051 | 1.514E-07 | 0.025692 | 0.039204 | 0.025691 | 0.0043838 | 0.014123 | 0.0048927 | 0.108696 | 0.089453 | 0.112258 |
| 13 | 13_53665008_A_G | 53665008 | G | A | PCMTD2 | intronic | 5.848E-09 | 0.0061221 | 3.046E-07 | -0.027675 | -0.042491 | -0.027079 | 0.0047543 | 0.015501 | 0.0052882 | 0.094516 | 0.075768 | 0.097986 |
| 13 | 13_53665087_A_G | 53665087 | G | A | PCMTD2 | intronic | 3.7087E-09 | 0.0079047 | 2.483E-07 | -0.028065 | -0.041268 | -0.027309 | 0.0047595 | 0.015537 | 0.0052934 | 0.094933 | 0.077103 | 0.098233 |
| 13 | 13_53666036_T_C | 53666036 | C | T | PCMTD2 | intronic | 1.0154E-09 | 0.0048405 | 8.447E-08 | 0.026753 | 0.039448 | 0.026215 | 0.0043808 | 0.014001 | 0.0048934 | 0.108331 | 0.088785 | 0.111949 |
| 13 | 13_53666133_T_C | 53666133 | C | T | PCMTD2 | intronic | 1.2727E-09 | 0.0063351 | 8.058E-08 | 0.027445 | 0.041508 | 0.02688 | 0.0045209 | 0.015205 | 0.0050095 | 0.104056 | 0.078772 | 0.108736 |
| 13 | 13_53667446_A_C | 53667446 | C | A | PCMTD2 | intronic | 3.3647E-08 | 0.010333 | 5.298E-07 | 0.02382 | 0.036353 | 0.024073 | 0.0043142 | 0.014176 | 0.0048 | 0.113961 | 0.091455 | 0.118127 |
| 13 | 13_53706565_T_C | 53706565 | C | T | MYT1 | intronic | 2.7408E-08 | 0.23792 | 1.109E-07 | -0.027438 | -0.017651 | -0.02959 | 0.0049373 | 0.014956 | 0.0055746 | 0.087426 | 0.079773 | 0.088842 |
| 13 | 13_53727328_G_A | 53727328 | A | G | MYT1 | intronic | 2.0267E-10 | 0.034508 | 1.306E-08 | -0.027516 | -0.028452 | -0.027645 | 0.004327 | 0.013458 | 0.0048626 | 0.115994 | 0.105808 | 0.11788 |
| 13 | 13_53729408_G_A | 53729408 | A | G | MYT1 | intronic | 2.5774E-08 | 0.23858 | 1.965E-07 | -0.027892 | -0.017956 | -0.029411 | 0.0050094 | 0.015236 | 0.0056531 | 0.086175 | 0.079439 | 0.087421 |
| 13 | 13_53803885_C_T | 53803885 | T | C | RGS19(XM_025000763.1:c.*229C>T, | UTR3 | 1.3058E-08 | 0.24473 | 1.214E-07 | -0.030865 | -0.018403 | -0.032689 | 0.0054289 | 0.01582 | 0.0061778 | 0.072047 | 0.067757 | 0.072841 |
| 13 | 13_53811139_C_T | 53811139 | T | C | TCEA2 | intronic | 1.2537E-09 | 0.26938 | 1.154E-08 | -0.033201 | -0.017534 | -0.035537 | 0.0054668 | 0.015876 | 0.0062277 | 0.071369 | 0.067089 | 0.072161 |
| 13 | 13_53827681_G_A | 53827681 | A | G | SOX18(dist=2394),C13H20orf204(dist | intergenic | 9.4005E-09 | 0.51832 | 3.425E-08 | -0.0319 | -0.010861 | -0.034592 | 0.0055564 | 0.016814 | 0.0062688 | 0.069961 | 0.060414 | 0.071729 |
| 13 | 13_53863355_A_T | 53863355 | T | A | PRPF6 | intronic | 2.7317E-08 | 0.38402 | 1.272E-07 | -0.030306 | -0.014059 | -0.032678 | 0.0054528 | 0.01615 | 0.0061856 | 0.07236 | 0.068425 | 0.073088 |
| 13 | 13_53964448_G_A | 53964448 | A | G | ABHD16B(dist=5687),LOC112449309 | intergenic | 2.8497E-08 | 0.092445 | 2.852E-07 | -0.030162 | -0.026165 | -0.03187 | 0.0054342 | 0.01555 | 0.0062088 | 0.071265 | 0.067423 | 0.071976 |
| 13 | 13_53987372_C_G | 53987372 | G | C | ZBTB46 | intronic | 4.0692E-08 | 0.36786 | 2.48E-07 | -0.029807 | -0.014543 | -0.03177 | 0.0054315 | 0.01615 | 0.006158 | 0.072777 | 0.068425 | 0.073582 |
| 13 | 13_54046492_C_G | 54046492 | G | C | ZGPAT | exonic | 1.5493E-08 | 0.13947 | 2.499E-07 | -0.030237 | -0.022974 | -0.031394 | 0.005346 | 0.015547 | 0.0060867 | 0.073663 | 0.068425 | 0.074632 |

| | | | | | | | | | | | | | | | | | | |
|---|---|---|---|---|---|---|---|---|---|---|---|---|---|---|---|---|---|---|
| 13 | 13_54046685_C_T | 54046685 | T | C | ZGPAT | exonic | 4.8617E-08 | 0.20894 | 5.862E-07 | -0.029265 | -0.019776 | -0.030449 | 0.0053635 | 0.015739 | 0.006095 | 0.073715 | 0.068758 | 0.074632 |
| 13 | 13_54049127_G_A | 54049127 | A | G | ARFRP1 | intronic | 2.8792E-09 | 0.11291 | 9.5E-08 | -0.031799 | -0.025374 | -0.032359 | 0.0053548 | 0.016006 | 0.0060642 | 0.074028 | 0.069092 | 0.074941 |
| 13 | 13_54049637_C_T | 54049637 | T | C | ARFRP1 | intronic | 1.0678E-08 | 0.21866 | 1.316E-07 | -0.030531 | -0.019569 | -0.031912 | 0.005338 | 0.015908 | 0.0060477 | 0.074341 | 0.069426 | 0.07525 |
| 13 | 13_54052028_C_T | 54052028 | T | C | ARFRP1 | intronic | 2.626E-08 | 0.35932 | 3.16E-07 | -0.029777 | -0.014617 | -0.031005 | 0.0053511 | 0.015946 | 0.0060632 | 0.073923 | 0.068091 | 0.075003 |
| 13 | 13_54052696_C_T | 54052696 | T | C | ARFRP1(XM_025000630.1:c.*19C>T, | UTR3 | 2.859E-08 | 0.17373 | 3.078E-07 | -0.029568 | -0.02094 | -0.0311 | 0.0053277 | 0.015393 | 0.0060759 | 0.074497 | 0.071429 | 0.075065 |
| 13 | 13_54054269_G_A | 54054269 | A | G | ARFRP1,TNFRSF6B(dist=32) | downstream | 1.9191E-08 | 0.2557 | 2.108E-07 | -0.029938 | -0.017808 | -0.031411 | 0.0053278 | 0.015668 | 0.0060526 | 0.074236 | 0.068758 | 0.07525 |
| 13 | 13_54060242_C_T | 54060242 | T | C | RTEL1 | exonic | 3.0803E-08 | 0.17534 | 9.471E-07 | -0.029391 | -0.020835 | -0.029661 | 0.0053084 | 0.015374 | 0.0060504 | 0.074653 | 0.070761 | 0.075374 |
| 13 | 13_54062728_T_C | 54062728 | C | T | RTEL1 | intronic | 3.8914E-08 | 0.22318 | 4.503E-07 | -0.029505 | -0.019472 | -0.030703 | 0.0053687 | 0.015985 | 0.0060842 | 0.073298 | 0.065754 | 0.074694 |
| 13 | 13_54074212_C_T | 54074212 | T | C | RTEL1 | intronic | 4.6435E-08 | 0.10249 | 9.381E-07 | -0.029085 | -0.026056 | -0.029546 | 0.0053226 | 0.015957 | 0.0060246 | 0.074341 | 0.068758 | 0.075374 |
| 13 | 13_54075711_T_C | 54075711 | C | T | RTEL1 | intronic | 3.5891E-08 | 0.21307 | 5.854E-07 | -0.029125 | -0.019286 | -0.030027 | 0.0052859 | 0.015488 | 0.0060103 | 0.074966 | 0.071095 | 0.075683 |
| 13 | 13_54090993_G_A | 54090993 | A | G | STMN3(dist=1764),GMEB2(dist=4140 | intergenic | 3.3371E-08 | 0.13351 | 3.085E-07 | -0.029396 | -0.02376 | -0.030869 | 0.0053227 | 0.015836 | 0.0060312 | 0.075175 | 0.068425 | 0.076424 |
| 13 | 13_54112236_T_C | 54112236 | T | C | GMEB2(XM_025000448.1:c.*1313C>T | UTR3 | 3.6518E-08 | 0.21422 | 3.564E-07 | -0.029569 | -0.019541 | -0.031093 | 0.0053694 | 0.015733 | 0.0061076 | 0.073976 | 0.06976 | 0.074756 |
| 13 | 13_54115191_T_C | 54115191 | C | T | FNDC11(dist=600) | downstream | 2.8492E-08 | 0.28646 | 3.736E-07 | -0.029651 | -0.017099 | -0.030717 | 0.005342 | 0.016042 | 0.0060443 | 0.074601 | 0.068758 | 0.075683 |
| 13 | 13_54118492_G_A | 54118492 | A | G | FNDC11 | intronic | 2.9177E-08 | 0.15548 | 4.786E-07 | -0.029773 | -0.02226 | -0.030761 | 0.0053682 | 0.015671 | 0.0061098 | 0.072881 | 0.067757 | 0.073829 |
| 13 | 13_54138151_C_A | 54138151 | A | C | PPDPF(dist=276) | upstream | 2.4995E-08 | 0.30161 | 3.804E-07 | -0.030078 | -0.016268 | -0.0312 | 0.0053967 | 0.015748 | 0.0061435 | 0.071995 | 0.067757 | 0.072779 |
| 13 | 13_54138496_G_C | 54138496 | C | G | PPDPF(dist=621) | upstream | 4.9209E-08 | 0.27265 | 7.722E-07 | -0.029644 | -0.018063 | -0.030332 | 0.0054352 | 0.016466 | 0.0061373 | 0.072255 | 0.065754 | 0.073459 |
| 13 | 13_54163666_T_G | 54163666 | G | T | EEF1A2(dist=3969),KCNQ2(dist=7739 | intergenic | 3.4044E-08 | 0.26363 | 8.024E-07 | -0.029672 | -0.01745 | -0.030233 | 0.005376 | 0.01561 | 0.0061266 | 0.072777 | 0.066756 | 0.073891 |
| 13 | 13_54166798_C_T | 54166798 | T | C | EEF1A2(dist=7101),KCNQ2(dist=4607 | intergenic | 4.2142E-08 | 0.25758 | 4.154E-07 | -0.029744 | -0.017702 | -0.031351 | 0.0054262 | 0.015636 | 0.0061937 | 0.072881 | 0.070093 | 0.073397 |
| 13 | 13_54180079_C_T | 54180079 | T | C | KCNQ2 | intronic | 3.0347E-08 | 0.042861 | 7.483E-06 | 0.024098 | 0.026666 | 0.02201 | 0.0043503 | 0.013168 | 0.0049134 | 0.110155 | 0.104473 | 0.111207 |
| 13 | 13_54186051_C_G | 54186051 | G | C | KCNQ2 | intronic | 1.119E-08 | 0.056273 | 2.338E-06 | 0.02496 | 0.024898 | 0.023325 | 0.0043701 | 0.013043 | 0.0049398 | 0.109738 | 0.104473 | 0.110713 |
| 13 | 13_54222374_T_A | 54222374 | A | T | KCNQ2(dist=2665),CHRNA4(dist=207 | intergenic | 3.0251E-08 | 0.027179 | 1.956E-06 | -0.02521 | -0.030637 | -0.024407 | 0.0045506 | 0.01387 | 0.0051297 | 0.103013 | 0.094459 | 0.104597 |
| 13 | 13_54234702_C_T | 54234702 | T | C | KCNQ2(dist=14993),CHRNA4(dist=84 | intergenic | 4.0927E-08 | 0.20388 | 3.259E-07 | -0.029864 | -0.020765 | -0.031462 | 0.0054429 | 0.016343 | 0.0061595 | 0.072464 | 0.067089 | 0.073459 |
| 13 | 13_54240830_C_T | 54240830 | T | C | KCNQ2(dist=21121),CHRNA4(dist=23 | intergenic | 2.3007E-08 | 0.1874 | 2.709E-07 | -0.030211 | -0.021144 | -0.031523 | 0.0054067 | 0.016038 | 0.0061297 | 0.072307 | 0.068758 | 0.072964 |
| 13 | 13_54344986_A_G | 54344986 | G | A | YTHDF1(XM_005214814.4:c.*729A>C | UTR3 | 1.7753E-08 | 0.27999 | 9.153E-08 | -0.030859 | -0.017602 | -0.033195 | 0.0054787 | 0.016294 | 0.006213 | 0.071682 | 0.066756 | 0.072594 |
| 13 | 13_54380263_G_A | 54380263 | A | G | LOC112449310(dist=9907),LOC1019 | intergenic | 4.7417E-08 | 0.22418 | 1.373E-06 | -0.029277 | -0.019005 | -0.029461 | 0.0053614 | 0.015635 | 0.006101 | 0.073611 | 0.070093 | 0.074262 |
| 13 | 13_54404840_G_A | 54404840 | A | G | LOC101905484(dist=6036),BHLHE23 | intergenic | 3.135E-08 | 0.082546 | 6.483E-07 | -0.030087 | -0.027369 | -0.03084 | 0.0054369 | 0.015765 | 0.0061975 | 0.072099 | 0.067423 | 0.072964 |
| 13 | 13_54504795_A_C | 54504795 | C | A | DIDO1 | intronic | 4.1338E-08 | 0.078427 | 7.652E-07 | -0.029943 | -0.028906 | -0.030512 | 0.0054591 | 0.016425 | 0.0061715 | 0.071786 | 0.066088 | 0.072841 |
| 13 | 13_54529202_G_A | 54529202 | A | G | TCFL5 | intronic | 2.9354E-08 | 0.18081 | 4.454E-07 | -0.03009 | -0.021664 | -0.031034 | 0.0054263 | 0.016188 | 0.0061473 | 0.073819 | 0.06976 | 0.074571 |
| 13 | 13_54531129_T_C | 54531129 | C | T | TCFL5 | intronic | 4.1965E-08 | 0.16753 | 5.409E-07 | -0.029907 | -0.022502 | -0.030956 | 0.0054552 | 0.016304 | 0.0061773 | 0.073246 | 0.068425 | 0.074138 |
| 13 | 13_54541047_A_G | 54541047 | G | A | TCFL5 | intronic | 2.7603E-08 | 0.10633 | 4.963E-07 | -0.029694 | -0.025415 | -0.030532 | 0.0053445 | 0.015737 | 0.0060728 | 0.074862 | 0.07243 | 0.075312 |
| 13 | 13_54546007_T_A | 54546007 | A | T | COL9A3(XM_025001125.1:c.*6A>T) | UTR3 | 2.5907E-08 | 0.14711 | 4.594E-07 | -0.029911 | -0.022516 | -0.030905 | 0.0053729 | 0.01553 | 0.0061289 | 0.074028 | 0.071429 | 0.074509 |
| 13 | 13_54589402_C_T | 54589402 | T | C | MRGBP(dist=7942),NTSR1(dist=1971 | intergenic | 4.8809E-08 | 0.23167 | 5.646E-07 | -0.029367 | -0.019111 | -0.030535 | 0.0053829 | 0.015978 | 0.0061035 | 0.074028 | 0.070427 | 0.074694 |
| 13 | 13_54622128_G_C | 54622128 | C | G | NTSR1 | intronic | 9.8119E-09 | 0.042307 | 7.334E-07 | -0.026153 | -0.028443 | -0.025405 | 0.004561 | 0.014008 | 0.00513 | 0.103535 | 0.093458 | 0.1054 |
| 13 | 13_54624261_T_C | 54624261 | C | T | NTSR1 | intronic | 2.4422E-08 | 0.24563 | 3.971E-07 | -0.029877 | -0.018471 | -0.030798 | 0.0053568 | 0.015909 | 0.0060741 | 0.074132 | 0.069426 | 0.075003 |
| 13 | 13_54635602_G_A | 54635602 | A | G | NTSR1 | intronic | 3.7916E-08 | 0.16839 | 3.278E-07 | -0.029197 | -0.020798 | -0.031032 | 0.0053083 | 0.0151 | 0.0060767 | 0.07408 | 0.072764 | 0.074323 |
| 13 | 13_54637982_G_A | 54637982 | A | G | NTSR1 | intronic | 4.2756E-08 | 0.18566 | 3.527E-07 | -0.028263 | -0.020524 | -0.0297 | 0.0051583 | 0.015507 | 0.0058316 | 0.074184 | 0.071762 | 0.074632 |
| 13 | 13_54650491_G_A | 54650491 | A | G | NTSR1 | intronic | 1.0962E-08 | 0.072967 | 8.447E-07 | 0.024491 | 0.023046 | 0.023838 | 0.0042853 | 0.012853 | 0.0048405 | 0.116411 | 0.108812 | 0.117818 |
| 13 | 13_54684548_C_T | 54684548 | T | C | SLCO4A1 | exonic | 9.5268E-09 | 0.1106 | 2.98E-07 | -0.024782 | -0.020972 | -0.024922 | 0.0043182 | 0.013144 | 0.0048632 | 0.116359 | 0.104139 | 0.118621 |
| 13 | 13_54714074_T_C | 54714074 | C | T | LOC104973829 | ncRNA_exonic | 1.4428E-08 | 0.18176 | 1.948E-07 | -0.030398 | -0.020938 | -0.031752 | 0.0053629 | 0.01568 | 0.0061011 | 0.074184 | 0.070427 | 0.07488 |
| 13 | 13_54715138_C_T | 54715138 | T | C | LOC104973829 | ncRNA_intronic | 4.6384E-08 | 0.25845 | 3.008E-07 | -0.02927 | -0.017682 | -0.031229 | 0.0053563 | 0.015646 | 0.0060958 | 0.074549 | 0.072764 | 0.07488 |
| 19 | 19_27847146_G_A | 27847146 | A | G | TRNAD-GUC_15(dist=13);TRNAG-UC | upstream;downstream | 5.3451E-09 | 0.82401 | 9.696E-11 | -0.018223 | 0.0020173 | -0.022827 | 0.0031224 | 0.0090712 | 0.0035272 | 0.263163 | 0.270694 | 0.261769 |
| 19 | 19_27847147_C_G | 27847147 | G | C | TRNAD-GUC_15(dist=14);TRNAG-UC | upstream;downstream | 6.7908E-09 | 0.82401 | 1.293E-10 | -0.018097 | 0.0020173 | -0.022673 | 0.0031224 | 0.0090712 | 0.0035272 | 0.263111 | 0.270694 | 0.261708 |
| 19 | 19_27847705_A_G | 27847705 | G | A | TRNAD-GUC_15(dist=572);TRNAP-C | upstream;downstream | 1.3261E-09 | 0.56632 | 1.28E-10 | -0.018474 | -0.005176 | -0.022031 | 0.0030463 | 0.0090264 | 0.0034264 | 0.289594 | 0.287049 | 0.290065 |
| 19 | 19_27848382_C_T | 27848382 | T | C | TRNAP-CGG_1(dist=569) | upstream | 1.0355E-09 | 0.41541 | 2.858E-10 | -0.018606 | -0.007309 | -0.02167 | 0.0030482 | 0.0089739 | 0.0034363 | 0.289021 | 0.288051 | 0.289201 |
| 19 | 19_27849059_A_G | 27849059 | G | A | LOC112442636(dist=473) | downstream | 1.4243E-09 | 0.40617 | 3.033E-10 | -0.018506 | -0.00756 | -0.021643 | 0.0030574 | 0.0091009 | 0.003437 | 0.288552 | 0.284379 | 0.289324 |
| 19 | 19_27851129_A_G | 27851129 | G | A | LOC112442636(dist=208);TRNAS-AG | upstream;downstream | 1.7164E-09 | 0.59689 | 1.895E-10 | -0.018281 | -0.004727 | -0.021797 | 0.0030354 | 0.0089368 | 0.003422 | 0.290585 | 0.288718 | 0.29093 |
| 19 | 19_27851735_C_T | 27851735 | T | C | LOC112442636,TRNAS-AGA_4,TRNA | upstream;downstream | 1.4903E-09 | 0.59531 | 9.258E-11 | -0.018889 | -0.004847 | -0.022834 | 0.0031245 | 0.0091251 | 0.0035245 | 0.263372 | 0.269693 | 0.262202 |
| 19 | 19_27852114_T_C | 27852114 | C | T | TRNAS-AGA_4,TRNAT-AGU_5(dist=4 | upstream;downstream | 7.3206E-09 | 0.6741 | 4.883E-10 | -0.018058 | -0.00385 | -0.021897 | 0.0031223 | 0.0091551 | 0.0035189 | 0.26405 | 0.27036 | 0.262882 |
| 19 | 19_27853166_C_T | 27853166 | T | C | CTC1(XM_010816067.2:c.*190G>A,X | UTR3 | 2.2562E-09 | 0.44134 | 8.533E-10 | -0.018193 | -0.006976 | -0.020983 | 0.0030432 | 0.0090601 | 0.0034203 | 0.289021 | 0.284713 | 0.289818 |
| 19 | 19_27855150_T_A | 27855150 | A | T | CTC1 | intronic | 6.734E-09 | 0.83801 | 4.187E-10 | -0.018173 | -0.001855 | -0.022123 | 0.0031346 | 0.0090742 | 0.0035414 | 0.262694 | 0.272363 | 0.260904 |
| 19 | 19_27855458_G_A | 27855458 | A | G | CTC1 | intronic | 4.1573E-10 | 0.40718 | 1.512E-10 | -0.019095 | -0.007567 | -0.021991 | 0.0030562 | 0.0091292 | 0.0034339 | 0.288656 | 0.284379 | 0.289448 |
| 19 | 19_27856693_G_A | 27856693 | A | G | CTC1 | intronic | 8.0664E-10 | 0.35728 | 3.553E-10 | -0.018687 | -0.008274 | -0.021491 | 0.0030417 | 0.0089879 | 0.0034262 | 0.290168 | 0.288385 | 0.290498 |
| 19 | 19_27860065_A_G | 27860065 | G | A | CTC1 | exonic | 6.1004E-09 | 0.62219 | 5.204E-10 | -0.018119 | -0.004444 | -0.021891 | 0.0031164 | 0.0090182 | 0.0035235 | 0.263685 | 0.272363 | 0.262078 |
| 19 | 19_27861175_A_G | 27861175 | G | A | CTC1 | exonic | 1.574E-08 | 0.66605 | 2.07E-09 | -0.017625 | -0.00392 | -0.021083 | 0.0031177 | 0.0090831 | 0.0035183 | 0.263841 | 0.269693 | 0.262758 |
| 19 | 19_27861978_A_G | 27861978 | G | A | CTC1 | intronic | 2.8345E-08 | 0.7927 | 1.99E-09 | -0.0174 | -0.002425 | -0.021152 | 0.0031344 | 0.0092269 | 0.0035261 | 0.262851 | 0.269693 | 0.261584 |
| 19 | 19_27862463_G_C | 27862463 | C | G | CTC1 | intronic | 2.9346E-08 | 0.90001 | 1.436E-09 | -0.017236 | -0.00113 | -0.021256 | 0.0031082 | 0.0089922 | 0.0035126 | 0.264623 | 0.272697 | 0.263129 |
| 19 | 19_27864225_A_C | 27864225 | C | A | CTC1 | intronic | 1.5335E-08 | 0.92805 | 4.226E-10 | -0.017692 | -0.000829 | -0.021989 | 0.003127 | 0.0091833 | 0.0035209 | 0.263372 | 0.269693 | 0.262202 |
| 19 | 19_27864563_G_T | 27864563 | T | G | CTC1 | intronic | 9.3854E-09 | 0.39747 | 2.429E-09 | -0.017237 | -0.007082 | -0.020416 | 0.0030023 | 0.0083699 | 0.003422 | 0.302419 | 0.320093 | 0.299147 |
| 19 | 19_27864564_A_C | 27864564 | C | A | CTC1 | intronic | 9.3854E-09 | 0.39747 | 2.429E-09 | -0.017237 | -0.007082 | -0.020416 | 0.0030023 | 0.0083699 | 0.003422 | 0.302419 | 0.320093 | 0.299147 |
| 19 | 19_27864568_G_A | 27864568 | A | G | CTC1 | intronic | 9.3181E-09 | 0.42756 | 2.429E-09 | -0.017236 | -0.006626 | -0.020416 | 0.0030013 | 0.0083514 | 0.003422 | 0.302523 | 0.320761 | 0.299147 |
| 19 | 19_27867730_T_A | 27867730 | A | T | CTC1 | intronic | 1.2314E-08 | 0.46138 | 1.318E-09 | -0.017202 | -0.006595 | -0.020605 | 0.0030203 | 0.0089537 | 0.0033973 | 0.300125 | 0.301068 | 0.299951 |
| 19 | 19_27868063_C_T | 27868063 | T | C | CTC1 | intronic | 2.3096E-08 | 0.80781 | 1.062E-09 | 0.018913 | -0.002404 | 0.023227 | 0.0033851 | 0.0098817 | 0.0038079 | 0.227766 | 0.216288 | 0.22989 |
| 19 | 19_27868736_G_A | 27868736 | A | G | CTC1 | intronic | 7.7207E-09 | 0.41417 | 1.574E-09 | -0.017646 | -0.007481 | -0.02069 | 0.003056 | 0.0091609 | 0.0034273 | 0.289021 | 0.285381 | 0.289695 |
| 19 | 19_27871925_G_A | 27871925 | A | G | CTC1 | intronic | 1.0783E-09 | 0.51721 | 1.484E-10 | -0.018653 | -0.005907 | -0.022008 | 0.0030592 | 0.0091208 | 0.0034349 | 0.287457 | 0.281041 | 0.288645 |
| 19 | 19_27873455_T_C | 27873455 | C | T | CTC1 | intronic | 4.6925E-09 | 0.83211 | 4.425E-10 | -0.018214 | -0.001892 | -0.021965 | 0.0031093 | 0.0089249 | 0.003521 | 0.263841 | 0.272697 | 0.262202 |
| 19 | 19_27875264_T_C | 27875264 | C | T | CTC1(dist=1569),PFAS(dist=4687) | intergenic | 1.3299E-09 | 0.38328 | 6.619E-10 | -0.018428 | -0.007866 | -0.021124 | 0.0030391 | 0.0090221 | 0.0034209 | 0.289334 | 0.286048 | 0.289942 |
| 19 | 19_27878648_A_G | 27878648 | G | A | CTC1(dist=4953),PFAS(dist=1303) | intergenic | 7.254E-09 | 0.57658 | 1.182E-09 | -0.017882 | -0.005039 | -0.021212 | 0.0030912 | 0.0090246 | 0.0034874 | 0.270566 | 0.275701 | 0.269616 |
| 19 | 19_27879943_G_A | 27879943 | A | G | PFAS(dist=8) | upstream | 6.568E-09 | 0.402 | 1.572E-09 | -0.017564 | -0.007472 | -0.020587 | 0.0030274 | 0.0089153 | 0.0034102 | 0.290481 | 0.286382 | 0.291239 |
| 19 | 19_27880163_T_C | 27880163 | C | T | PFAS | intronic | 7.3454E-09 | 0.81234 | 5.5E-10 | -0.018047 | -0.002157 | -0.021839 | 0.0031208 | 0.0090875 | 0.00352 | 0.265144 | 0.272029 | 0.26387 |

| | | | | | | | | | | | | | | | | | | |
|---|---|---|---|---|---|---|---|---|---|---|---|---|---|---|---|---|---|---|
| 19 | 19_27881447_G_A | 27881447 | A | G | PFAS | intronic | 1.1668E-08 | 0.48488 | 3.854E-09 | -0.017356 | -0.006297 | -0.020155 | 0.0030426 | 0.0090153 | 0.0034218 | 0.289125 | 0.283378 | 0.290189 |
| 19 | 19_27882688_T_G | 27882688 | G | T | PFAS | intronic | 8.5748E-09 | 0.48273 | 1.224E-09 | -0.01741 | -0.006162 | -0.020767 | 0.0030243 | 0.0087788 | 0.0034173 | 0.291367 | 0.29239 | 0.291178 |
| 19 | 19_27883894_G_A | 27883894 | A | G | PFAS | intronic | 3.0246E-09 | 0.46536 | 9.522E-10 | -0.01809 | -0.006597 | -0.020993 | 0.0030504 | 0.0090365 | 0.0034318 | 0.289647 | 0.286048 | 0.290313 |
| 19 | 19_27886686_C_T | 27886686 | T | C | PFAS | intronic | 8.1698E-09 | 0.9831 | 3.251E-10 | 0.018286 | 0.000198 | 0.022417 | 0.003172 | 0.0093476 | 0.0035661 | 0.256751 | 0.258344 | 0.256456 |
| 19 | 19_27887087_C_T | 27887087 | T | C | PFAS | intronic | 4.2066E-09 | 0.66257 | 5.141E-10 | -0.018329 | -0.003982 | -0.021861 | 0.0031193 | 0.0091243 | 0.0035176 | 0.263945 | 0.269025 | 0.263005 |
| 19 | 19_27887195_G_A | 27887195 | A | G | PFAS | intronic | 4.1296E-09 | 0.71577 | 1.869E-10 | -0.018311 | -0.003293 | -0.022413 | 0.0031146 | 0.0090437 | 0.0035175 | 0.264362 | 0.271028 | 0.263129 |
| 20 | 20_71161624_A_G | 71161624 | G | A | TERT | exonic | 2.664E-08 | 0.6468 | 8.701E-10 | 0.015703 | 0.0037725 | 0.01962 | 0.0028232 | 0.0082331 | 0.0031998 | 0.383015 | 0.414219 | 0.37724 |
| 20 | 20_71161828_T_C | 71161828 | C | T | TERT(XM_024981282.1:c.*133T>C,X | UTR3 | 3.0533E-08 | 0.5992 | 7.846E-10 | 0.015613 | 0.0043051 | 0.019663 | 0.0028191 | 0.0081915 | 0.0031983 | 0.383797 | 0.419226 | 0.37724 |
| 20 | 20_71161915_G_A | 71161915 | A | G | TERT(XM_024981282.1:c.*220G>A,X | UTR3 | 4.529E-08 | 0.56799 | 1.254E-09 | 0.0154 | 0.0046544 | 0.019417 | 0.0028159 | 0.008151 | 0.0031971 | 0.383589 | 0.421228 | 0.376622 |
| 20 | 20_71161970_A_G | 71161970 | G | A | TERT(XM_024981282.1:c.*275A>G,X | UTR3 | 1.9823E-08 | 0.47343 | 5.469E-10 | 0.015865 | 0.0059193 | 0.019868 | 0.0028262 | 0.0082566 | 0.0032019 | 0.381608 | 0.414553 | 0.37551 |
| 20 | 20_71173729_C_T | 71173729 | T | C | TERT(dist=11352),SLC6A18(dist=134 | intergenic | 1.0884E-10 | 0.21583 | 6.714E-13 | 0.018026 | 0.010258 | 0.022528 | 0.002793 | 0.0082883 | 0.0031353 | 0.489365 | 0.458611 | 0.479736 |
| 20 | 20_71173872_T_C | 71173872 | C | T | TERT(dist=11495),SLC6A18(dist=119 | intergenic | 4.2003E-08 | 0.19674 | 1.237E-09 | 0.015511 | 0.010567 | 0.01941 | 0.0028294 | 0.008186 | 0.0031948 | 0.415285 | 0.459613 | 0.40708 |
| 20 | 20_71175537_G_A | 71175537 | A | G | SLC6A18 | intronic | 1.0814E-10 | 0.10073 | 1.789E-12 | 0.018072 | 0.013383 | 0.022242 | 0.0027996 | 0.0081537 | 0.0031549 | 0.458034 | 0.498999 | 0.45008 |
| 20 | 20_71176000_G_A | 71176000 | A | G | SLC6A18 | exonic | 4.3334E-10 | 0.11589 | 7.614E-12 | 0.017544 | 0.013017 | 0.02164 | 0.0028109 | 0.0082791 | 0.0031612 | 0.456209 | 0.497664 | 0.448536 |
| 20 | 20_71177017_G_A | 71177017 | A | G | SLC6A18 | intronic | 2.4802E-10 | 0.15884 | 1.97E-12 | 0.017845 | 0.011674 | 0.022316 | 0.0028198 | 0.0082856 | 0.0031714 | 0.457147 | 0.499332 | 0.449339 |
| 20 | 20_71177331_A_G | 71177331 | G | A | SLC6A18 | intronic | 2.8996E-10 | 0.38555 | 5.731E-13 | 0.017789 | 0.0071732 | 0.022857 | 0.0028218 | 0.0082669 | 0.0031716 | 0.447138 | 0.488318 | 0.439516 |
| 20 | 20_71178266_G_A | 71178266 | A | G | SLC6A18 | intronic | 1.842E-11 | 0.079522 | 1.309E-13 | 0.018881 | 0.014343 | 0.023456 | 0.0028106 | 0.0081798 | 0.0031675 | 0.461422 | 0.492657 | 0.452922 |
| 20 | 20_71178514_A_G | 71178514 | G | A | SLC6A18 | intronic | 4.1244E-08 | 0.39516 | 6.333E-11 | 0.015395 | 0.0068706 | 0.020733 | 0.0028065 | 0.0080803 | 0.0031723 | 0.423887 | 0.476636 | 0.414123 |
| 20 | 20_71182845_G_A | 71182845 | A | G | SLC6A19(dist=93) | downstream | 1.8759E-10 | 0.10505 | 1.002E-12 | 0.017955 | 0.013419 | 0.022588 | 0.0028182 | 0.0082793 | 0.003168 | 0.456313 | 0.494993 | 0.4473 |
| 20 | 20_71187383_T_G | 71187383 | G | T | SLC6A19 | intronic | 1.7755E-09 | 0.27824 | 5.331E-12 | 0.017032 | 0.0090609 | 0.021902 | 0.0028306 | 0.0083566 | 0.0031758 | 0.44182 | 0.482977 | 0.434202 |
| 20 | 20_71187859_C_T | 71187859 | T | C | SLC6A19 | intronic | 1.3099E-09 | 0.13757 | 1.606E-11 | 0.017101 | 0.012279 | 0.021362 | 0.0028191 | 0.0082693 | 0.0031703 | 0.454124 | 0.497664 | 0.4452 |
| 20 | 20_71188377_G_A | 71188377 | A | G | SLC6A19 | intronic | 1.9889E-10 | 0.18471 | 3.36E-12 | 0.017852 | 0.010739 | 0.022034 | 0.0028059 | 0.0080963 | 0.003165 | 0.459389 | 0.486649 | 0.449401 |
| 20 | 20_71188387_T_C | 71188387 | C | T | SLC6A19 | intronic | 1.9768E-09 | 0.58711 | 5.986E-12 | 0.016884 | 0.004372 | 0.021841 | 0.0028141 | 0.0080511 | 0.0031746 | 0.445522 | 0.496328 | 0.436118 |
| 20 | 20_71188608_C_T | 71188608 | T | C | SLC6A19 | intronic | 1.9693E-10 | 0.14158 | 4.348E-12 | 0.017922 | 0.012102 | 0.02195 | 0.0028163 | 0.0082328 | 0.0031696 | 0.46038 | 0.496662 | 0.452428 |
| 20 | 20_71189372_C_T | 71189372 | T | C | SLC6A19 | intronic | 4.618E-10 | 0.13039 | 7.318E-12 | 0.017572 | 0.012436 | 0.021749 | 0.0028198 | 0.0082216 | 0.0031744 | 0.454176 | 0.499332 | 0.44557 |
| 20 | 20_71189496_A_G | 71189496 | G | A | SLC6A19 | intronic | 2.3796E-10 | 0.11049 | 6.921E-12 | 0.017895 | 0.013172 | 0.02179 | 0.002825 | 0.0082534 | 0.0031767 | 0.45647 | 0.493992 | 0.4473 |
| 20 | 20_71190263_C_T | 71190263 | T | C | SLC6A19 | intronic | 2.1959E-10 | 0.12932 | 3.913E-12 | 0.017955 | 0.012559 | 0.022069 | 0.0028289 | 0.00828 | 0.0031798 | 0.455688 | 0.495661 | 0.446682 |
| 20 | 20_71190711_G_A | 71190711 | A | G | SLC6A19 | intronic | 3.3776E-10 | 0.25997 | 3.202E-12 | 0.017725 | 0.0092768 | 0.022143 | 0.0028223 | 0.0082352 | 0.0031776 | 0.455844 | 0.496662 | 0.447053 |
| 20 | 20_71191782_C_G | 71191782 | G | C | SLC6A19 | intronic | 2.4687E-10 | 0.15272 | 2.208E-12 | 0.017876 | 0.011887 | 0.022285 | 0.0028245 | 0.0083123 | 0.0031743 | 0.453811 | 0.498999 | 0.445447 |
| 20 | 20_71192010_C_T | 71192010 | T | C | SLC6A19 | exonic | 3.0489E-10 | 0.15962 | 3.774E-12 | 0.017745 | 0.01155 | 0.022047 | 0.0028183 | 0.008213 | 0.0031743 | 0.458398 | 0.49466 | 0.44971 |
| 20 | 20_71193078_G_A | 71193078 | A | G | SLC6A19 | intronic | 5.2048E-10 | 0.10486 | 3.279E-12 | -0.017521 | -0.013507 | -0.022058 | 0.0028202 | 0.0083285 | 0.0031669 | 0.441299 | 0.492657 | 0.431793 |
| 20 | 20_71193729_C_A | 71193729 | A | C | SLC6A19 | intronic | 1.2565E-09 | 0.57052 | 1.155E-12 | -0.01716 | -0.004659 | -0.022617 | 0.0028257 | 0.0082127 | 0.0031808 | 0.42519 | 0.462951 | 0.418201 |
| 20 | 20_71194228_A_G | 71194228 | G | A | SLC6A19 | intronic | 1.5637E-09 | 0.32143 | 7.049E-12 | -0.017017 | -0.008037 | -0.021809 | 0.0028184 | 0.0081059 | 0.0031807 | 0.430195 | 0.479306 | 0.421105 |
| 20 | 20_71195675_G_A | 71195675 | A | G | SLC6A19 | intronic | 2.1148E-10 | 0.22556 | 3.73E-12 | 0.017872 | 0.010004 | 0.021985 | 0.0028132 | 0.0082553 | 0.0031646 | 0.458086 | 0.499332 | 0.450204 |
| 20 | 20_71195975_G_A | 71195975 | A | G | SLC6A19 | intronic | 1.4092E-10 | 0.043334 | 4.979E-12 | 0.017989 | 0.016978 | 0.021803 | 0.0028042 | 0.008403 | 0.0031571 | 0.456261 | 0.491322 | 0.446559 |
| 20 | 20_71196104_C_T | 71196104 | T | C | SLC6A19 | intronic | 1.148E-10 | 0.14554 | 6.856E-13 | 0.01807 | 0.011848 | 0.022695 | 0.0028032 | 0.0081404 | 0.0031599 | 0.452455 | 0.496662 | 0.444273 |
| 20 | 20_71196132_C_T | 71196132 | T | C | SLC6A19 | intronic | 1.2453E-10 | 0.13211 | 7.647E-13 | 0.018023 | 0.012265 | 0.022623 | 0.0028013 | 0.0081448 | 0.0031564 | 0.453967 | 0.498331 | 0.445138 |
| 20 | 20_71196133_T_G | 71196133 | G | T | SLC6A19 | intronic | 1.1413E-10 | 0.13006 | 7.23E-13 | 0.018068 | 0.012338 | 0.022654 | 0.0028026 | 0.0081498 | 0.0031574 | 0.454176 | 0.497997 | 0.445323 |
| 20 | 20_71197234_A_C | 71197234 | C | A | SLC6A19 | intronic | 4.7808E-11 | 0.13456 | 1.406E-12 | 0.018589 | 0.012495 | 0.022476 | 0.0028261 | 0.0083499 | 0.003173 | 0.457199 | 0.496662 | 0.449895 |
| 20 | 20_71198412_C_T | 71198412 | T | C | SLC6A19 | intronic | 1.8533E-10 | 0.14697 | 1.053E-12 | 0.017896 | 0.012207 | 0.022392 | 0.0028081 | 0.0084167 | 0.0031435 | 0.489365 | 0.456943 | 0.479427 |
| 20 | 20_71198758_G_C | 71198758 | C | G | SLC6A19 | intronic | 4.8494E-09 | 0.60925 | 9.178E-12 | 0.016486 | 0.004091 | 0.021717 | 0.002817 | 0.0080036 | 0.0031849 | 0.449901 | 0.493658 | 0.441802 |
| 20 | 20_71198831_G_A | 71198831 | A | G | SLC6A19 | intronic | 6.8552E-10 | 0.10744 | 1.205E-11 | 0.017308 | 0.012845 | 0.021533 | 0.0028054 | 0.0079791 | 0.0031761 | 0.460849 | 0.484646 | 0.45076 |
| 20 | 20_71199479_G_A | 71199479 | A | G | SLC6A19(dist=238) | upstream | 5.1152E-11 | 0.072186 | 5.858E-13 | 0.01839 | 0.014314 | 0.022802 | 0.0028001 | 0.0079612 | 0.0031653 | 0.461214 | 0.477637 | 0.449895 |
| 20 | 20_71201576_T_C | 71201576 | C | T | SLC6A19(dist=2335),LOC112443022( | intergenic | 8.766E-11 | 0.1533 | 1.008E-12 | 0.018169 | 0.011633 | 0.022487 | 0.0028009 | 0.0081463 | 0.0031541 | 0.457408 | 0.494326 | 0.448474 |
| 20 | 20_71201782_C_A | 71201782 | A | C | SLC6A19(dist=2541),LOC112443022( | intergenic | 5.7897E-11 | 0.12533 | 5.09E-13 | 0.0183 | 0.012276 | 0.022797 | 0.0027943 | 0.0080091 | 0.0031563 | 0.458972 | 0.49032 | 0.449586 |
| 20 | 20_71201844_G_A | 71201844 | A | G | SLC6A19(dist=2603),LOC112443022( | intergenic | 1.9952E-10 | 0.11412 | 1.799E-12 | 0.01782 | 0.012861 | 0.022238 | 0.0028012 | 0.0081404 | 0.0031546 | 0.455218 | 0.493324 | 0.445694 |
| 20 | 20_71201866_G_A | 71201866 | A | G | SLC6A19(dist=2625),LOC112443022( | intergenic | 2.8938E-10 | 0.12172 | 2.98E-12 | 0.017694 | 0.012758 | 0.022006 | 0.0028066 | 0.0082436 | 0.0031533 | 0.455271 | 0.493324 | 0.445756 |
| 20 | 20_71202457_G_A | 71202457 | A | G | SLC6A19(dist=3216),LOC112443022( | intergenic | 1.3949E-10 | 0.18661 | 3.977E-13 | 0.018153 | 0.011072 | 0.02303 | 0.0028291 | 0.0083839 | 0.0031738 | 0.454072 | 0.498665 | 0.445817 |
| 20 | 20_71202535_T_G | 71202535 | G | T | SLC6A19(dist=3294),LOC112443022( | intergenic | 3.5111E-10 | 0.21782 | 6.226E-13 | 0.017676 | 0.010239 | 0.022766 | 0.0028172 | 0.0083087 | 0.0031639 | 0.454019 | 0.496662 | 0.444891 |
| 20 | 20_71202705_C_T | 71202705 | T | C | SLC6A19(dist=3464),LOC112443022( | intergenic | 4.4916E-10 | 0.15594 | 3.351E-12 | 0.017552 | 0.011598 | 0.022054 | 0.0028146 | 0.0081743 | 0.0031677 | 0.457564 | 0.489319 | 0.447733 |
| 20 | 20_71202891_G_A | 71202891 | A | G | SLC6A19(dist=3650),LOC112443022( | intergenic | 9.8647E-11 | 0.22513 | 1.365E-13 | 0.018136 | 0.0099076 | 0.023351 | 0.0028036 | 0.0081678 | 0.0031557 | 0.456887 | 0.495661 | 0.448103 |
| 20 | 20_71202905_C_T | 71202905 | T | C | SLC6A19(dist=3664),LOC112443022( | intergenic | 1.421E-10 | 0.24399 | 3.715E-13 | 0.017964 | 0.0094935 | 0.022923 | 0.0028009 | 0.0081484 | 0.003155 | 0.45965 | 0.493658 | 0.451007 |
| 20 | 20_71203436_C_T | 71203436 | T | C | SLC6A19(dist=4195),LOC112443022( | intergenic | 6.7627E-11 | 0.17116 | 4.089E-13 | -0.01833 | -0.011264 | -0.022917 | 0.0028089 | 0.0082314 | 0.0031598 | 0.439162 | 0.478972 | 0.431793 |
| 20 | 20_71203454_C_T | 71203454 | T | C | SLC6A19(dist=4213),LOC112443022( | intergenic | 4.6638E-11 | 0.1715 | 2.941E-13 | -0.018488 | -0.011248 | -0.023064 | 0.0028091 | 0.0082257 | 0.0031607 | 0.439214 | 0.479306 | 0.431793 |
| 20 | 20_71203976_C_T | 71203976 | T | C | SLC6A19(dist=4735),LOC112443022( | intergenic | 2.3265E-09 | 0.33837 | 6.759E-12 | 0.016875 | 0.0078072 | 0.021825 | 0.0028251 | 0.0081546 | 0.0031803 | 0.444166 | 0.493992 | 0.434944 |
| 20 | 20_71204762_C_T | 71204762 | T | C | SLC6A19(dist=5521),LOC112443022( | intergenic | 1.2246E-09 | 0.24609 | 4.767E-12 | 0.017144 | 0.0096265 | 0.021908 | 0.0028211 | 0.0082995 | 0.0031694 | 0.457825 | 0.493324 | 0.448783 |
| 20 | 20_71205809_C_T | 71205809 | T | C | LOC112443022(dist=663) | upstream | 1.206E-10 | 0.092253 | 9.168E-13 | 0.018122 | 0.013783 | 0.022635 | 0.0028146 | 0.0081864 | 0.0031691 | 0.456574 | 0.490988 | 0.446868 |
| 20 | 20_71205893_G_A | 71205893 | A | G | LOC112443022(dist=579) | upstream | 4.6512E-11 | 0.17955 | 1.726E-13 | 0.018498 | 0.010876 | 0.023364 | 0.0028105 | 0.0081031 | 0.0031708 | 0.459598 | 0.491322 | 0.450513 |
| 20 | 20_71206009_C_T | 71206009 | T | C | LOC112443022(dist=463) | upstream | 7.7924E-11 | 0.13909 | 6.072E-13 | 0.018174 | 0.011853 | 0.022738 | 0.002794 | 0.0080128 | 0.0031585 | 0.458868 | 0.485648 | 0.448598 |
| 20 | 20_71206269_T_C | 71206269 | C | T | LOC112443022(dist=203) | upstream | 3.8065E-10 | 0.18738 | 1.039E-12 | 0.017664 | 0.010932 | 0.022593 | 0.0028209 | 0.0082918 | 0.0031708 | 0.453654 | 0.497664 | 0.444644 |
| 20 | 20_71206277_T_C | 71206277 | C | T | LOC112443022(dist=195) | upstream | 4.7571E-10 | 0.17547 | 1.739E-12 | 0.017561 | 0.011234 | 0.022359 | 0.0028202 | 0.0082917 | 0.0031697 | 0.45428 | 0.496996 | 0.445261 |
| 20 | 20_71206494_A_G | 71206494 | G | A | LOC112443022 | ncRNA_exonic | 5.7742E-10 | 0.062172 | 8.5E-12 | 0.017481 | 0.015476 | 0.021651 | 0.0028211 | 0.0082977 | 0.0031701 | 0.45501 | 0.497664 | 0.44625 |
| 20 | 20_71207015_G_C | 71207015 | C | G | LOC112443022 | ncRNA_exonic | 5.9705E-09 | 0.51985 | 4.914E-12 | 0.016497 | 0.0053567 | 0.021993 | 0.0028358 | 0.0083234 | 0.0031837 | 0.441038 | 0.485648 | 0.432781 |
| 20 | 20_71207618_A_G | 71207618 | G | A | LOC112443022 | ncRNA_exonic | 2.4639E-11 | 0.093978 | 2.211E-13 | 0.01872 | 0.013567 | 0.023198 | 0.0028042 | 0.0081007 | 0.0031625 | 0.461579 | 0.490654 | 0.452737 |
| 20 | 20_71207699_A_C | 71207699 | C | A | LOC112443022 | ncRNA_exonic | 5.0009E-10 | 0.38164 | 9.936E-13 | 0.017522 | 0.0072358 | 0.022554 | 0.0028175 | 0.0082706 | 0.0031627 | 0.446825 | 0.490654 | 0.438712 |
| 20 | 20_71210424_T_C | 71210424 | C | T | LOC112443022 | ncRNA_exonic | 2.6999E-10 | 0.21088 | 3.158E-12 | 0.017714 | 0.010458 | 0.021939 | 0.002805 | 0.0083585 | 0.0031473 | 0.481389 | 0.467623 | 0.471951 |
| 20 | 20_71211786_A_G | 71211786 | G | A | LOC112443022 | ncRNA_exonic | 4.2941E-08 | 0.36084 | 4.482E-10 | 0.015414 | 0.0073499 | 0.019857 | 0.0028136 | 0.0080435 | 0.0031841 | 0.413565 | 0.466288 | 0.403806 |
| 20 | 20_71211891_T_C | 71211891 | C | T | LOC112443022 | ncRNA_exonic | 1.0671E-10 | 0.28807 | 5.952E-13 | 0.018157 | 0.0087602 | 0.022764 | 0.0028119 | 0.0082459 | 0.003161 | 0.452768 | 0.497997 | 0.443655 |
| 20 | 20_71211934_G_A | 71211934 | A | G | LOC112443022(dist=6) | downstream | 7.6258E-11 | 0.23977 | 7.186E-13 | 0.018329 | 0.0096807 | 0.022728 | 0.0028165 | 0.0082349 | 0.0031673 | 0.453237 | 0.496662 | 0.443964 |
| 20 | 20_71212077_A_G | 71212077 | G | A | LOC112443022(dist=149) | downstream | 2.1662E-10 | 0.10406 | 3.599E-12 | 0.01801 | 0.013781 | 0.022095 | 0.0028366 | 0.0084783 | 0.0031782 | 0.451569 | 0.492991 | 0.443902 |
| 20 | 20_71212282_G_A | 71212282 | A | G | LOC112443022(dist=354) | downstream | 7.5323E-10 | 0.14586 | 1.206E-11 | 0.017463 | 0.012364 | 0.021547 | 0.0028374 | 0.0085015 | 0.0031783 | 0.454436 | 0.496328 | 0.446682 |
| 20 | 20_71212367_G_A | 71212367 | A | G | LOC112443022(dist=439) | downstream | 1.6991E-10 | 0.17837 | 1.233E-12 | 0.017947 | 0.011041 | 0.022473 | 0.0028102 | 0.0082039 | 0.0031645 | 0.455218 | 0.492657 | 0.44557 |
| 20 | 20_71212411_A_G | 71212411 | G | A | LOC112443022(dist=483) | downstream | 8.8902E-11 | 0.13882 | 9.415E-13 | 0.01825 | 0.012242 | 0.022596 | 0.0028144 | 0.0082702 | 0.0031652 | 0.455688 | 0.494326 | 0.446435 |
| 20 | 20_71212595_C_T | 71212595 | T | C | LOC112443022(dist=667) | downstream | 1.0974E-10 | 0.12799 | 1.088E-12 | 0.018313 | 0.012904 | 0.022637 | 0.002838 | 0.0084779 | 0.0031798 | 0.454072 | 0.496996 | 0.446126 |
| 20 | 20_71212923_G_A | 71212923 | A | G | LOC112443022(dist=995) | downstream | 4.4023E-11 | 0.070925 | 2.556E-13 | 0.018548 | 0.014835 | 0.022183 | 0.0028147 | 0.0082147 | 0.0031688 | 0.460275 | 0.494326 | 0.451872 |

| | | | | | | | | | | | | | | | | | | |
|---|---|---|---|---|---|---|---|---|---|---|---|---|---|---|---|---|---|---|
| 20 | 20_71213496_C_T | 71213496 | T | C | LOC112443022(dist=1568),LOC11244 | intergenic | 1.0084E-10 | 0.19305 | 1.123E-12 | 0.018135 | 0.010598 | 0.022482 | 0.0028048 | 0.0081424 | 0.0031601 | 0.462048 | 0.485648 | 0.452366 |
| 20 | 20_71213510_G_A | 71213510 | A | G | LOC112443022(dist=1582),LOC11244 | intergenic | 8.2632E-11 | 0.13086 | 8.293E-13 | 0.018325 | 0.012593 | 0.022682 | 0.0028211 | 0.0083358 | 0.0031695 | 0.464237 | 0.490988 | 0.45595 |
| 20 | 20_71213845_A_G | 71213845 | G | A | LOC112443022(dist=1917),LOC11244 | intergenic | 3.6668E-10 | 0.24974 | 4.408E-12 | 0.017634 | 0.0094551 | 0.02194 | 0.0028135 | 0.0082148 | 0.003169 | 0.460484 | 0.49466 | 0.452181 |
| 20 | 20_71214406_C_T | 71214406 | T | C | LOC112443022(dist=2478),LOC11244 | intergenic | 3.0594E-09 | 0.29792 | 1.653E-11 | 0.016733 | 0.0086618 | 0.021344 | 0.0028225 | 0.0083215 | 0.0031697 | 0.459493 | 0.493992 | 0.450883 |
| 20 | 20_71215178_G_A | 71215178 | A | G | LOC112443022(dist=3250),LOC11244 | intergenic | 2.2167E-08 | 0.053427 | 2.804E-09 | 0.015874 | 0.016188 | 0.018984 | 0.0028375 | 0.0083813 | 0.0031946 | 0.430404 | 0.475634 | 0.422031 |
| 20 | 20_71216949_C_G | 71216949 | G | C | LOC112443022(dist=5021),LOC11244 | intergenic | 2.2113E-10 | 0.22493 | 2.434E-12 | 0.017967 | 0.010263 | 0.022243 | 0.0028312 | 0.0084573 | 0.0031743 | 0.461526 | 0.499666 | 0.454343 |
| 20 | 20_71217219_C_T | 71217219 | T | C | LOC112443022(dist=5291),LOC11244 | intergenic | 1.0742E-09 | 0.3701 | 6.411E-12 | 0.017255 | 0.0074979 | 0.021817 | 0.0028296 | 0.0083655 | 0.0031756 | 0.462048 | 0.49466 | 0.454034 |
| 20 | 20_71218826_C_G | 71218826 | G | C | LOC112443022(dist=6898),LOC11244 | intergenic | 3.3013E-08 | 0.22119 | 8.93E-10 | 0.015603 | 0.010099 | 0.01952 | 0.0028243 | 0.0082549 | 0.0031857 | 0.431133 | 0.481976 | 0.421722 |
| 20 | 20_71219009_C_T | 71219009 | T | C | LOC112443022(dist=7081),LOC11244 | intergenic | 9.6942E-10 | 0.21126 | 4.722E-12 | 0.017241 | 0.010688 | 0.021778 | 0.0028197 | 0.00855 | 0.0031499 | 0.491867 | 0.45494 | 0.482022 |
| 20 | 20_71219157_G_A | 71219157 | A | G | LOC112443022(dist=7229),LOC11244 | intergenic | 2.4423E-09 | 0.32705 | 1.078E-11 | 0.016753 | 0.0080252 | 0.021467 | 0.0028084 | 0.0081884 | 0.0031588 | 0.460223 | 0.487984 | 0.450636 |
| 20 | 20_71219339_G_A | 71219339 | A | G | LOC112443022(dist=7411),LOC11244 | intergenic | 1.4203E-10 | 0.3117 | 2.953E-12 | 0.018117 | 0.0084561 | 0.022145 | 0.0028248 | 0.0083586 | 0.0031727 | 0.462256 | 0.497664 | 0.454838 |
| 20 | 20_71219816_C_T | 71219816 | T | C | LOC112443022(dist=7888),LOC11244 | intergenic | 4.3107E-10 | 0.40889 | 1.198E-12 | 0.017571 | 0.0069736 | 0.022389 | 0.0028149 | 0.0084441 | 0.003151 | 0.495621 | 0.452937 | 0.486099 |
| 20 | 20_71225610_C_A | 71225610 | A | C | LOC112443022(dist=13682),LOC1124 | intergenic | 1.6901E-09 | 0.19814 | 1.448E-11 | 0.016986 | 0.010748 | 0.021363 | 0.0028193 | 0.0083521 | 0.0031634 | 0.460119 | 0.49466 | 0.451748 |
| 20 | 20_71226172_A_G | 71226172 | G | A | LOC112443022(dist=14244),LOC1124 | intergenic | 1.5911E-10 | 0.15112 | 3.045E-12 | 0.017984 | 0.011865 | 0.022045 | 0.0028115 | 0.0082651 | 0.0031604 | 0.460536 | 0.49466 | 0.452243 |
| 20 | 20_71230082_G_A | 71230082 | A | G | LOC112443023(dist=258) | downstream | 3.6199E-08 | 0.08203 | 2.386E-09 | 0.015567 | 0.014221 | 0.019061 | 0.0028261 | 0.0081774 | 0.0031933 | 0.430091 | 0.478304 | 0.421166 |
| 20 | 20_71232631_C_T | 71232631 | T | C | LOC112443023 | ncRNA_exonic | 3.8079E-10 | 0.28192 | 2.352E-12 | 0.017652 | 0.0088995 | 0.022227 | 0.0028191 | 0.0082708 | 0.00317 | 0.460119 | 0.488652 | 0.450636 |
| 20 | 20_71235737_T_C | 71235737 | C | T | SLC12A7 | exonic | 4.9081E-08 | 0.2989 | 6.653E-10 | 0.015521 | 0.0085485 | 0.019822 | 0.0028454 | 0.0082292 | 0.0032104 | 0.408769 | 0.455607 | 0.400099 |
| 20 | 20_71281129_G_A | 71281129 | A | G | SLC12A7 | intronic | 2.3275E-08 | 0.52957 | 5.689E-11 | 0.015779 | 0.0052231 | 0.020825 | 0.0028248 | 0.0083084 | 0.0031785 | 0.499427 | 0.449266 | 0.491289 |
| 20 | 20_71290453_A_G | 71290453 | G | A | SLC12A7 | intronic | 1.9177E-09 | 0.25263 | 9.726E-12 | 0.016924 | 0.0092721 | 0.021671 | 0.0028185 | 0.0081052 | 0.003182 | 0.447346 | 0.483645 | 0.440628 |
| 20 | 20_71290462_G_A | 71290462 | A | G | SLC12A7 | intronic | 6.5058E-09 | 0.24671 | 3.165E-11 | 0.016361 | 0.0097382 | 0.020988 | 0.0028193 | 0.0084067 | 0.0031614 | 0.494265 | 0.440921 | 0.495861 |
| 20 | 20_71290589_T_C | 71290589 | C | T | SLC12A7 | intronic | 8.9643E-10 | 0.42055 | 2.357E-12 | 0.017234 | 0.0066982 | 0.022136 | 0.0028129 | 0.0083159 | 0.0031571 | 0.479721 | 0.474299 | 0.47121 |
| 20 | 20_71291864_C_T | 71291864 | T | C | SLC12A7 | intronic | 1.8852E-08 | 0.078096 | 5.3E-10 | 0.015769 | 0.014377 | 0.019638 | 0.0028048 | 0.0081601 | 0.0031623 | 0.471014 | 0.484312 | 0.462746 |
| 20 | 20_71291886_G_A | 71291886 | A | G | SLC12A7 | intronic | 9.6545E-09 | 0.072314 | 2.448E-10 | 0.0161 | 0.014653 | 0.020034 | 0.0028066 | 0.0081537 | 0.0031648 | 0.470754 | 0.485314 | 0.462622 |
| 20 | 20_71293128_C_T | 71293128 | T | C | SLC12A7 | exonic | 1.2817E-09 | 0.14587 | 5.024E-11 | 0.017059 | 0.011787 | 0.020859 | 0.0028105 | 0.0081051 | 0.0031747 | 0.462569 | 0.498331 | 0.455332 |
| 20 | 20_71293773_A_G | 71293773 | G | A | SLC12A7 | intronic | 7.0163E-10 | 0.27662 | 1.379E-11 | 0.017458 | 0.008945 | 0.021565 | 0.0028314 | 0.0082219 | 0.0031901 | 0.443541 | 0.475968 | 0.437539 |
| 20 | 20_71295284_A_G | 71295284 | G | A | SLC12A7 | intronic | 1.8481E-09 | 0.65118 | 6.285E-12 | 0.017072 | 0.0037763 | 0.021919 | 0.0028402 | 0.0083523 | 0.0031891 | 0.441925 | 0.475968 | 0.435623 |
| 20 | 20_71297262_A_G | 71297262 | G | A | SLC12A7 | intronic | 3.4113E-09 | 0.61105 | 1.376E-11 | 0.016733 | 0.0042188 | 0.021498 | 0.0028311 | 0.0082953 | 0.0031799 | 0.44229 | 0.477971 | 0.435685 |
| 20 | 20_71297709_C_T | 71297709 | T | C | SLC12A7 | intronic | 1.6555E-09 | 0.26072 | 2.526E-11 | 0.017015 | 0.0092962 | 0.021187 | 0.0028225 | 0.0082655 | 0.0031756 | 0.465332 | 0.488318 | 0.456753 |
| 20 | 20_71298057_G_T | 71298057 | T | G | SLC12A7 | intronic | 1.8282E-09 | 0.36088 | 1.956E-11 | 0.017039 | 0.0076669 | 0.021331 | 0.0028339 | 0.0083912 | 0.0031793 | 0.465124 | 0.487984 | 0.456444 |
| 20 | 20_71300352_G_A | 71300352 | A | G | SLC12A7(XM_024981553.1:c.*600G> | UTR3 | 5.9779E-09 | 0.2332 | 1.533E-10 | 0.016457 | 0.0098597 | 0.020379 | 0.0028289 | 0.0082706 | 0.0031831 | 0.466323 | 0.487984 | 0.457865 |
| 20 | 20_71300630_T_G | 71300630 | G | T | SLC12A7(XM_024981552.1:c.*878T> | UTR3 | 1.4008E-09 | 0.54929 | 4.174E-12 | 0.017173 | 0.0049201 | 0.022129 | 0.002836 | 0.0082163 | 0.0031927 | 0.440517 | 0.481976 | 0.432843 |
| 20 | 20_71301433_A_G | 71301433 | G | A | SLC12A7(dist=416) | downstream | 1.5783E-10 | 0.49898 | 1.405E-13 | 0.018193 | 0.0056619 | 0.023593 | 0.0028437 | 0.0083743 | 0.00319 | 0.434939 | 0.472296 | 0.428024 |
| 20 | 20_71301611_C_T | 71301611 | T | C | SLC12A7(dist=594) | downstream | 4.8695E-09 | 0.57128 | 8.428E-12 | 0.016524 | 0.004841 | 0.021548 | 0.0028239 | 0.0085505 | 0.0031543 | 0.495308 | 0.448932 | 0.484987 |
| 20 | 20_71301631_G_A | 71301631 | A | G | SLC12A7(dist=614) | downstream | 2.9637E-09 | 0.53894 | 5.408E-12 | 0.016765 | 0.0052563 | 0.021758 | 0.0028254 | 0.008555 | 0.0031559 | 0.495464 | 0.449266 | 0.485234 |
| 20 | 20_71301654_C_T | 71301654 | T | C | SLC12A7(dist=637) | downstream | 4.2653E-09 | 0.69389 | 4.733E-12 | 0.016583 | 0.0033646 | 0.021796 | 0.0028233 | 0.0085487 | 0.0031527 | 0.495777 | 0.447931 | 0.485358 |
| 20 | 20_71301896_G_T | 71301896 | T | G | SLC12A7(dist=879) | downstream | 4.4004E-10 | 0.60252 | 4.21E-13 | 0.017567 | 0.0043122 | 0.022898 | 0.0028156 | 0.0082801 | 0.003159 | 0.475133 | 0.473965 | 0.465711 |
| 20 | 20_71302597_G_A | 71302597 | A | G | SLC12A7(dist=1580),NKD2(dist=3964 | intergenic | 9.3556E-09 | 0.41668 | 1.904E-11 | 0.016096 | 0.0064948 | 0.021276 | 0.0028031 | 0.0079966 | 0.0031692 | 0.462778 | 0.477637 | 0.451748 |
| 20 | 20_71304744_T_C | 71304744 | C | T | SLC12A7(dist=3727),NKD2(dist=1817 | intergenic | 1.5767E-09 | 0.50593 | 3.327E-12 | 0.017086 | 0.005339 | 0.022279 | 0.0028306 | 0.0080264 | 0.0031995 | 0.443906 | 0.496662 | 0.434141 |
| 20 | 20_71306108_A_G | 71306108 | G | A | NKD2(dist=453) | downstream | 5.0354E-10 | 0.5778 | 9.934E-13 | 0.017581 | 0.0045142 | 0.022715 | 0.0028275 | 0.0081102 | 0.0031852 | 0.439214 | 0.486649 | 0.430434 |
| 20 | 20_71306715_G_A | 71306715 | A | G | NKD2(XM_024981562.1:c.*826C>T) | UTR3 | 5.2618E-09 | 0.48034 | 1.666E-11 | 0.016461 | 0.0060018 | 0.021229 | 0.0028194 | 0.0085041 | 0.0031531 | 0.493953 | 0.453939 | 0.484307 |
| 20 | 20_71309526_A_G | 71309526 | G | A | NKD2 | intronic | 1.1752E-09 | 0.42625 | 8.833E-12 | 0.017279 | 0.0066609 | 0.021774 | 0.0028403 | 0.0083718 | 0.0031907 | 0.461735 | 0.493658 | 0.453478 |
| 20 | 20_71309556_G_A | 71309556 | A | G | NKD2 | intronic | 1.3716E-09 | 0.31842 | 2.266E-11 | 0.017096 | 0.0083747 | 0.02114 | 0.0028217 | 0.008394 | 0.003161 | 0.49828 | 0.447597 | 0.488261 |
| 20 | 20_71313268_G_C | 71313268 | C | G | NKD2 | intronic | 3.5635E-10 | 0.25411 | 3.625E-12 | 0.017655 | 0.0095341 | 0.021961 | 0.0028149 | 0.0083602 | 0.0031594 | 0.495308 | 0.452603 | 0.485667 |
| 20 | 20_71313672_T_C | 71313672 | C | T | NKD2 | intronic | 5.6072E-10 | 0.3126 | 3.011E-12 | 0.017467 | 0.0082592 | 0.022157 | 0.0028168 | 0.0081793 | 0.0031757 | 0.458242 | 0.495661 | 0.44971 |
| 20 | 20_71313905_G_A | 71313905 | A | G | NKD2 | intronic | 2.6626E-09 | 0.29538 | 2.995E-11 | 0.016839 | 0.0087476 | 0.021123 | 0.0028296 | 0.0083599 | 0.0031779 | 0.462152 | 0.492991 | 0.453849 |
| 20 | 20_71313977_G_T | 71313977 | T | G | NKD2 | intronic | 8.1944E-10 | 0.22687 | 1.581E-11 | 0.017381 | 0.010049 | 0.02145 | 0.0028303 | 0.0083158 | 0.0031823 | 0.462413 | 0.492323 | 0.454034 |
| 20 | 20_71314587_G_A | 71314587 | A | G | NKD2 | intronic | 1.073E-09 | 0.63311 | 2.258E-12 | 0.01737 | 0.003995 | 0.022442 | 0.0028484 | 0.0083692 | 0.0031979 | 0.43838 | 0.475634 | 0.431484 |
| 20 | 20_71316291_T_C | 71316291 | C | T | NKD2 | intronic | 2.1116E-09 | 0.2269 | 1.752E-11 | 0.016935 | 0.009951 | 0.021417 | 0.0028277 | 0.008235 | 0.0031846 | 0.465906 | 0.486315 | 0.457062 |
| 20 | 20_71316956_T_C | 71316956 | C | T | NKD2 | intronic | 3.0877E-09 | 0.19569 | 3.552E-11 | 0.016759 | 0.010707 | 0.021068 | 0.0028277 | 0.0082749 | 0.0031818 | 0.463716 | 0.488318 | 0.454838 |
| 20 | 20_71317071_T_C | 71317071 | C | T | NKD2 | intronic | 1.1893E-09 | 0.27245 | 7.288E-12 | 0.017196 | 0.009162 | 0.021764 | 0.0028275 | 0.0083486 | 0.0031764 | 0.4621 | 0.492991 | 0.453787 |
| 20 | 20_71317607_C_G | 71317607 | G | C | NKD2 | intronic | 3.6496E-09 | 0.079213 | 4.912E-11 | 0.016581 | 0.014275 | 0.020839 | 0.0028107 | 0.0081327 | 0.0031701 | 0.461109 | 0.496996 | 0.453355 |
| 20 | 20_71318401_C_A | 71318401 | C | A | NKD2 | intronic | 1.2599E-10 | 0.24833 | 9.895E-13 | 0.018241 | 0.0097199 | 0.02269 | 0.002836 | 0.0084198 | 0.0031815 | 0.458398 | 0.493324 | 0.449462 |
| 20 | 20_71318739_C_T | 71318739 | T | C | NKD2 | intronic | 3.1447E-10 | 0.30982 | 1.405E-12 | 0.017889 | 0.0086376 | 0.022574 | 0.0028434 | 0.0085048 | 0.0031869 | 0.458138 | 0.495661 | 0.449586 |
| 20 | 20_71319648_T_C | 71319648 | C | T | NKD2 | intronic | 2.5755E-09 | 0.84894 | 2.966E-12 | 0.016917 | 0.0015604 | 0.02234 | 0.0028401 | 0.0081921 | 0.0032009 | 0.438067 | 0.477303 | 0.430804 |
| 20 | 20_71319659_G_A | 71319659 | A | G | NKD2 | intronic | 1.3455E-09 | 0.80447 | 1.658E-12 | 0.017207 | 0.0020269 | 0.02259 | 0.0028386 | 0.0081874 | 0.0031995 | 0.438432 | 0.476969 | 0.431299 |
| 20 | 20_71319678_G_A | 71319678 | A | G | NKD2 | intronic | 5.6165E-10 | 0.25535 | 4.468E-12 | 0.017615 | 0.0096017 | 0.022065 | 0.0028407 | 0.0084414 | 0.0031878 | 0.459076 | 0.495995 | 0.45076 |
| 20 | 20_71319974_C_T | 71319974 | T | C | NKD2 | intronic | 4.3887E-10 | 0.57952 | 5.733E-13 | 0.017739 | 0.0046088 | 0.023003 | 0.002843 | 0.008318 | 0.0031919 | 0.437806 | 0.473632 | 0.431175 |
| 20 | 20_71320470_A_G | 71320470 | G | A | NKD2 | intronic | 1.1932E-09 | 0.50624 | 1.794E-12 | 0.017312 | 0.0055356 | 0.022536 | 0.0028468 | 0.008328 | 0.0031967 | 0.438015 | 0.477303 | 0.430743 |
| 20 | 20_71320628_A_G | 71320628 | G | A | NKD2 | intronic | 2.9167E-10 | 0.30706 | 5.427E-13 | 0.017909 | 0.0084617 | 0.023025 | 0.0028413 | 0.0082844 | 0.0031916 | 0.436138 | 0.476636 | 0.428642 |
| 20 | 20_71320811_C_A | 71320811 | A | C | NKD2 | intronic | 3.6944E-09 | 0.36458 | 3.698E-11 | 0.016686 | 0.0075265 | 0.021055 | 0.0028294 | 0.0083013 | 0.0031826 | 0.463925 | 0.492657 | 0.455888 |
| 20 | 20_71321848_T_C | 71321848 | C | T | NKD2 | intronic | 3.5309E-10 | 0.35481 | 1.494E-12 | 0.017726 | 0.0075182 | 0.022524 | 0.0028256 | 0.0081252 | 0.0031836 | 0.43911 | 0.48498 | 0.430619 |
| 20 | 20_71322081_C_A | 71322081 | A | C | NKD2 | intronic | 7.4015E-10 | 0.28893 | 1.999E-12 | 0.017362 | 0.0087635 | 0.022287 | 0.0028198 | 0.0082637 | 0.0031682 | 0.459858 | 0.484646 | 0.449586 |
| 20 | 20_71322092_C_A | 71322092 | A | C | NKD2 | intronic | 1.3527E-09 | 0.37068 | 2.214E-12 | 0.017085 | 0.0073808 | 0.022241 | 0.0028189 | 0.0082449 | 0.0031681 | 0.460223 | 0.483311 | 0.449771 |
| 20 | 20_71322746_A_G | 71322746 | G | A | NKD2 | intronic | 5.6598E-10 | 0.47088 | 1.2E-12 | 0.01765 | 0.0060408 | 0.022698 | 0.002847 | 0.0083778 | 0.0031944 | 0.435877 | 0.473298 | 0.428951 |
| 20 | 20_71325349_A_G | 71325349 | G | A | NKD2 | intronic | 4.7119E-10 | 0.4228 | 3.563E-12 | 0.017763 | 0.0067556 | 0.022234 | 0.0028519 | 0.008428 | 0.0031975 | 0.436399 | 0.472964 | 0.429631 |
| 20 | 20_71327594_G_C | 71327594 | C | G | NKD2(dist=652) | upstream | 1.9265E-08 | 0.98405 | 7.985E-11 | 0.015967 | 0.0001636 | 0.020814 | 0.0028418 | 0.008184 | 0.0032017 | 0.460484 | 0.490988 | 0.451501 |
| 20 | 20_71328571_G_T | 71328571 | T | G | NKD2(dist=1629),LOC112443024(dist | intergenic | 4.154E-08 | 0.56519 | 4.134E-11 | 0.015484 | 0.0047566 | 0.020952 | 0.0028234 | 0.0082702 | 0.0031749 | 0.460275 | 0.493992 | 0.45181 |
| 20 | 20_71329124_C_T | 71329124 | T | C | NKD2(dist=2182),LOC112443024(dist | intergenic | 1.1931E-09 | 0.25064 | 7.269E-12 | 0.017167 | 0.0095029 | 0.021766 | 0.0028229 | 0.008272 | 0.0031765 | 0.456835 | 0.496328 | 0.448165 |
| 20 | 20_71329190_A_G | 71329190 | G | A | NKD2(dist=2248),LOC112443024(dist | intergenic | 1.3364E-09 | 0.25901 | 1.237E-11 | 0.017137 | 0.0093381 | 0.021549 | 0.0028265 | 0.008273 | 0.0031803 | 0.45892 | 0.493658 | 0.450142 |
| 20 | 20_71329260_T_C | 71329260 | C | T | NKD2(dist=2318),LOC112443024(dist | intergenic | 7.859E-10 | 0.31205 | 6.878E-12 | 0.017502 | 0.0085913 | 0.021894 | 0.0028489 | 0.0084985 | 0.0031915 | 0.46064 | 0.499332 | 0.453231 |
| 20 | 20_71329880_A_G | 71329880 | G | A | NKD2(dist=2938),LOC112443024(dist | intergenic | 9.5335E-10 | 0.58028 | 1.875E-12 | 0.017407 | 0.004617 | 0.022496 | 0.0028456 | 0.0083493 | 0.0031939 | 0.433844 | 0.474299 | 0.426356 |
| 20 | 20_71330019_A_G | 71330019 | G | A | NKD2(dist=3077),LOC112443024(dist | intergenic | 1.2063E-09 | 0.23895 | 1.103E-11 | 0.017244 | 0.010047 | 0.021568 | 0.0028365 | 0.0085321 | 0.0031753 | 0.455114 | 0.499332 | 0.446682 |
| 20 | 20_71330024_T_C | 71330024 | C | T | NKD2(dist=3082),LOC112443024(dist | intergenic | 1.321E-09 | 0.2403 | 1.103E-11 | 0.017205 | 0.010026 | 0.021568 | 0.0028369 | 0.008538 | 0.0031753 | 0.455166 | 0.498999 | 0.446682 |
| 20 | 20_71330245_G_A | 71330245 | A | G | NKD2(dist=3303),LOC112443024(dist | intergenic | 1.107E-09 | 0.39034 | 5.491E-12 | 0.017217 | 0.0071331 | 0.021897 | 0.0028255 | 0.0083039 | 0.0031771 | 0.459545 | 0.488652 | 0.449957 |
| 20 | 20_71330387_C_T | 71330387 | T | C | NKD2(dist=3445),LOC112443024(dist | intergenic | 3.7018E-09 | 0.41925 | 1.286E-11 | 0.016676 | 0.0067186 | 0.021518 | 0.0028279 | 0.008318 | 0.0031784 | 0.457929 | 0.492323 | 0.448721 |

| | | | | | | | | | | | | | | | | | | |
|---|---|---|---|---|---|---|---|---|---|---|---|---|---|---|---|---|---|---|
| 20 | 20_71330715_A_G | 71330715 | G | A | NKD2(dist=3773),LOC112443024(dist | intergenic | 4.34E-09 | 0.37743 | 2.647E-11 | 0.016671 | 0.0074319 | 0.021238 | 0.0028397 | 0.00842 | 0.0031864 | 0.458294 | 0.497997 | 0.450204 |
| 20 | 20_71330734_A_G | 71330734 | G | A | NKD2(dist=3792),LOC112443024(dist | intergenic | 3.713E-09 | 0.35712 | 2.66E-11 | 0.016744 | 0.0077484 | 0.021238 | 0.0028396 | 0.0084143 | 0.0031868 | 0.458555 | 0.497664 | 0.450451 |
| 20 | 20_71330904_T_C | 71330904 | C | T | NKD2(dist=3962),LOC112443024(dist | intergenic | 8.6019E-09 | 0.24229 | 8.258E-11 | 0.016355 | 0.0098501 | 0.02071 | 0.0028412 | 0.0084241 | 0.0031882 | 0.457981 | 0.496996 | 0.449648 |
| 20 | 20_71332478_C_T | 71332478 | T | C | NKD2(dist=5536),LOC112443024(dist | intergenic | 1.1694E-09 | 0.19828 | 9.322E-12 | 0.01718 | 0.010543 | 0.021681 | 0.0028236 | 0.0081954 | 0.0031807 | 0.458294 | 0.489987 | 0.448721 |
| 20 | 20_71332669_A_G | 71332669 | G | A | NKD2(dist=5727),LOC112443024(dist | intergenic | 1.9333E-09 | 0.26285 | 1.177E-11 | 0.016923 | 0.0091234 | 0.021574 | 0.0028189 | 0.0081483 | 0.0031806 | 0.458346 | 0.494993 | 0.44971 |
| 20 | 20_71332838_C_T | 71332838 | T | C | NKD2(dist=5896),LOC112443024(dist | intergenic | 2.6373E-10 | 0.48911 | 4.92E-13 | 0.017902 | 0.0056175 | 0.023105 | 0.0028331 | 0.008121 | 0.0031968 | 0.435669 | 0.480307 | 0.427406 |
| 20 | 20_71333741_T_C | 71333741 | C | T | NKD2(dist=6799),LOC112443024(dist | intergenic | 2.3923E-11 | 0.33706 | 1.033E-13 | 0.018995 | 0.0078884 | 0.02381 | 0.0028436 | 0.0082171 | 0.0032017 | 0.439005 | 0.479973 | 0.431422 |
| 20 | 20_71336741_G_A | 71336741 | A | G | NKD2(dist=9799),LOC112443024(dist | intergenic | 9.9494E-11 | 0.15938 | 5.222E-12 | 0.01837 | 0.011836 | 0.022015 | 0.0028402 | 0.008411 | 0.0031908 | 0.459598 | 0.498331 | 0.45181 |
| 20 | 20_71338118_T_C | 71338118 | C | T | NKD2(dist=11176),LOC112443024(di | intergenic | 1.3155E-09 | 0.25924 | 4.069E-11 | 0.017191 | 0.0094422 | 0.021013 | 0.0028342 | 0.0083693 | 0.003183 | 0.461891 | 0.495661 | 0.454034 |
| 20 | 20_71339517_A_G | 71339517 | G | A | NKD2(dist=12575),LOC112443024(di | intergenic | 1.3838E-10 | 0.54866 | 1.986E-12 | 0.01823 | 0.0049358 | 0.022499 | 0.0028407 | 0.0082294 | 0.0031979 | 0.437493 | 0.484312 | 0.428827 |
| 20 | 20_71339722_G_A | 71339722 | A | G | NKD2(dist=12780),LOC112443024(di | intergenic | 5.5737E-10 | 0.22981 | 1.499E-11 | 0.017603 | 0.0099989 | 0.021537 | 0.0028382 | 0.0083266 | 0.0031916 | 0.459337 | 0.492323 | 0.450389 |
| 20 | 20_71341393_G_A | 71341393 | A | G | LOC112443024(dist=317) | downstream | 2.2139E-10 | 0.11343 | 1.775E-11 | 0.017886 | 0.012969 | 0.021354 | 0.0028186 | 0.0081927 | 0.0031761 | 0.461891 | 0.490654 | 0.453108 |
| 20 | 20_71341407_G_A | 71341407 | A | G | LOC112443024(dist=303) | downstream | 1.9521E-10 | 0.30035 | 2.315E-12 | 0.018047 | 0.0084652 | 0.022395 | 0.0028353 | 0.0081735 | 0.0031928 | 0.439579 | 0.486983 | 0.430804 |
| 20 | 20_71342074_T_C | 71342074 | C | T | LOC112443024(XR_003031301.1:exo | ncRNA_splicing | 2.1778E-09 | 0.42183 | 2.567E-12 | 0.016976 | 0.0065744 | 0.022378 | 0.0028369 | 0.0081848 | 0.003197 | 0.434783 | 0.477637 | 0.42685 |
| 20 | 20_71342520_T_G | 71342520 | G | T | LOC112443024 | ncRNA_exonic | 5.3016E-11 | 0.28892 | 8.743E-14 | 0.018629 | 0.0087234 | 0.023808 | 0.0028389 | 0.0082258 | 0.0031921 | 0.43228 | 0.476969 | 0.424008 |
| 20 | 20_71342608_T_C | 71342608 | C | T | LOC112443024 | ncRNA_exonic | 4.6775E-10 | 0.65707 | 7.796E-13 | 0.0177 | 0.003648 | 0.02289 | 0.0028413 | 0.008217 | 0.0031949 | 0.435148 | 0.477637 | 0.427283 |
| 20 | 20_71343251_T_C | 71343251 | C | T | LOC112443024 | ncRNA_intronic | 2.4075E-10 | 0.49247 | 4.318E-13 | 0.018077 | 0.0057834 | 0.023189 | 0.0028545 | 0.008426 | 0.0032007 | 0.432541 | 0.471295 | 0.425368 |
| 20 | 20_71343298_G_A | 71343298 | A | G | LOC112443024 | ncRNA_intronic | 3.3206E-10 | 0.75714 | 5.273E-13 | 0.017943 | 0.0026141 | 0.023108 | 0.0028558 | 0.0084533 | 0.0032015 | 0.433844 | 0.472296 | 0.426727 |
| 20 | 20_71343339_T_C | 71343339 | C | T | LOC112443024 | ncRNA_intronic | 2.0046E-10 | 0.60026 | 3.612E-13 | 0.018143 | 0.0044034 | 0.023269 | 0.0028522 | 0.008403 | 0.0032009 | 0.433427 | 0.471963 | 0.426294 |
| 20 | 20_71343481_A_G | 71343481 | G | A | LOC112443024 | ncRNA_intronic | 6.8856E-11 | 0.33395 | 1.214E-13 | 0.018456 | 0.0078364 | 0.023654 | 0.0028293 | 0.0081106 | 0.0031899 | 0.43546 | 0.47964 | 0.427283 |
| 20 | 20_71343905_C_T | 71343905 | T | C | LOC112443024 | ncRNA_intronic | 5.1532E-10 | 0.16766 | 1.142E-11 | 0.017639 | 0.011613 | 0.021624 | 0.0028384 | 0.0084171 | 0.003186 | 0.461579 | 0.496662 | 0.453849 |
| 20 | 20_71344330_T_C | 71344330 | C | T | LOC112443024 | ncRNA_intronic | 1.3527E-09 | 0.79938 | 6.187E-13 | 0.017171 | 0.0021023 | 0.022894 | 0.0028329 | 0.0082719 | 0.0031814 | 0.432645 | 0.479973 | 0.423885 |
| 20 | 20_71344943_C_A | 71344943 | A | C | LOC112443024 | ncRNA_intronic | 6.1382E-10 | 0.2387 | 9.297E-12 | 0.017587 | 0.010053 | 0.0217 | 0.0028426 | 0.0085323 | 0.0031833 | 0.458816 | 0.49733 | 0.451687 |
| 20 | 20_71345082_T_C | 71345082 | C | T | LOC112443024 | ncRNA_intronic | 7.1565E-10 | 0.59337 | 1.006E-12 | 0.017586 | 0.0045004 | 0.022818 | 0.0028536 | 0.0084284 | 0.0032004 | 0.435825 | 0.476636 | 0.428271 |
| 20 | 20_71345201_T_A | 71345201 | A | T | LOC112443024 | ncRNA_intronic | 1.9116E-09 | 0.31484 | 1.782E-11 | 0.017056 | 0.0085538 | 0.02139 | 0.0028402 | 0.0085101 | 0.0031818 | 0.459337 | 0.497997 | 0.45144 |
| 20 | 20_71345389_T_C | 71345389 | C | T | LOC112443024 | ncRNA_intronic | 1.0443E-09 | 0.43094 | 1.536E-12 | 0.017418 | 0.0066246 | 0.022635 | 0.0028542 | 0.0084112 | 0.003201 | 0.435773 | 0.476302 | 0.428271 |
| 20 | 20_71346548_A_G | 71346548 | G | A | LOC112443024 | ncRNA_intronic | 9.0638E-11 | 0.39259 | 7.352E-14 | 0.018491 | 0.0071503 | 0.023959 | 0.0028527 | 0.0083638 | 0.0032024 | 0.432958 | 0.478304 | 0.424564 |
| 20 | 20_71346625_T_C | 71346625 | C | T | LOC112443024 | ncRNA_intronic | 6.6272E-11 | 0.57183 | 7.093E-14 | 0.018616 | 0.0047186 | 0.023972 | 0.0028514 | 0.0083463 | 0.0032021 | 0.43546 | 0.479973 | 0.427221 |
| 20 | 20_71347596_G_C | 71347596 | C | G | LOC112443024 | ncRNA_intronic | 2.2357E-09 | 0.38468 | 1.773E-11 | 0.016847 | 0.0073568 | 0.021221 | 0.0028173 | 0.0084629 | 0.0031562 | 0.493431 | 0.455941 | 0.48406 |
| 20 | 20_71348087_C_A | 71348087 | A | C | LOC112443024 | ncRNA_intronic | 2.4158E-10 | 0.45533 | 4.459E-13 | 0.018077 | 0.0062804 | 0.023175 | 0.0028547 | 0.0084125 | 0.0032005 | 0.435669 | 0.475634 | 0.428271 |
| 20 | 20_71348662_A_G | 71348662 | G | A | LOC112443024 | ncRNA_intronic | 1.9703E-09 | 0.82364 | 1.605E-12 | 0.017148 | 0.0018809 | 0.022624 | 0.0028579 | 0.0084397 | 0.0032022 | 0.433427 | 0.473632 | 0.425985 |
| 20 | 20_71348678_C_T | 71348678 | T | C | LOC112443024 | ncRNA_intronic | 1.1603E-08 | 0.93417 | 4.03E-12 | 0.0163 | -0.000699 | 0.0222 | 0.0028569 | 0.0084571 | 0.0032007 | 0.443906 | 0.485648 | 0.436179 |
| 20 | 20_71351659_C_G | 71351659 | G | C | LOC112443024(dist=899) | upstream | 5.7268E-12 | 0.24758 | 2.088E-14 | 0.01954 | 0.0095087 | 0.024424 | 0.0028375 | 0.0082237 | 0.0031947 | 0.431759 | 0.468625 | 0.424935 |
| 20 | 20_71352673_A_C | 71352673 | C | A | LOC112443024(dist=1913),TRIP13(di | intergenic | 3.1291E-11 | 0.21692 | 2.988E-14 | 0.018798 | 0.0099781 | 0.024278 | 0.0028308 | 0.0080811 | 0.003195 | 0.43155 | 0.474633 | 0.423576 |
| 20 | 20_71352902_A_G | 71352902 | G | A | LOC112443024(dist=2142),TRIP13(di | intergenic | 1.6015E-10 | 0.18268 | 3.39E-13 | 0.018139 | 0.01098 | 0.023211 | 0.0028363 | 0.0082396 | 0.0031892 | 0.430664 | 0.470628 | 0.423267 |
| 20 | 20_71354494_C_T | 71354494 | T | C | LOC112443024(dist=3734),TRIP13(di | intergenic | 1.5175E-10 | 0.21198 | 8.388E-13 | 0.01818 | 0.010252 | 0.022854 | 0.002839 | 0.0082135 | 0.0031942 | 0.433323 | 0.470961 | 0.426356 |
| 20 | 20_71354558_T_C | 71354558 | C | T | LOC112443024(dist=3798),TRIP13(di | intergenic | 1.7763E-10 | 0.23861 | 3.734E-13 | 0.018187 | 0.0099439 | 0.023213 | 0.0028508 | 0.008438 | 0.0031952 | 0.444219 | 0.483979 | 0.436859 |
| 20 | 20_71355096_C_T | 71355096 | T | C | LOC112443024(dist=4336),TRIP13(di | intergenic | 1.2192E-09 | 0.4175 | 2.867E-12 | 0.017335 | 0.0069498 | 0.022321 | 0.0028523 | 0.0085718 | 0.003196 | 0.463716 | 0.413885 | 0.47294 |
| 20 | 20_71355354_C_T | 71355354 | T | C | LOC112443024(dist=4594),TRIP13(di | intergenic | 4.1922E-12 | 0.23742 | 2.12E-14 | 0.019781 | 0.009936 | 0.024458 | 0.0028541 | 0.0084099 | 0.0032 | 0.432645 | 0.469626 | 0.4258 |
| 20 | 20_71355485_C_T | 71355485 | T | C | LOC112443024(dist=4725),TRIP13(di | intergenic | 8.3318E-11 | 0.25342 | 3.318E-13 | 0.018357 | 0.0096281 | 0.02302 | 0.0028265 | 0.0084303 | 0.0031618 | 0.470493 | 0.486315 | 0.462498 |
| 20 | 20_71358068_A_C | 71358068 | C | A | LOC112443024(dist=7308),TRIP13(di | intergenic | 2.3563E-11 | 0.15761 | 3.289E-13 | 0.018947 | 0.011529 | 0.023258 | 0.0028355 | 0.0081586 | 0.003194 | 0.432228 | 0.472296 | 0.424812 |
| 20 | 20_71358678_G_C | 71358678 | C | G | LOC112443024(dist=7918),TRIP13(di | intergenic | 1.5278E-08 | 0.57615 | 2.536E-11 | 0.016144 | 0.0047931 | 0.021306 | 0.0028532 | 0.0085742 | 0.0031937 | 0.460953 | 0.40988 | 0.470407 |
| 20 | 20_71359918_T_C | 71359918 | C | T | LOC112443024(dist=9158),TRIP13(di | intergenic | 2.1199E-10 | 0.39097 | 1.022E-12 | 0.017929 | 0.0070349 | 0.022604 | 0.0028223 | 0.0082005 | 0.0031714 | 0.435408 | 0.477303 | 0.427654 |
| 20 | 20_71360272_A_G | 71360272 | G | A | LOC112443024(dist=9512),TRIP13(di | intergenic | 4.358E-10 | 0.3076 | 1.502E-12 | 0.017679 | 0.0084883 | 0.022488 | 0.0028329 | 0.0083196 | 0.0031788 | 0.434835 | 0.474633 | 0.427468 |
| 20 | 20_71360601_G_C | 71360601 | C | G | LOC112443024(dist=9841),TRIP13(di | intergenic | 5.0362E-10 | 0.32972 | 1.724E-12 | 0.017596 | 0.0079634 | 0.022493 | 0.0028299 | 0.0081703 | 0.0031881 | 0.43937 | 0.480307 | 0.431793 |
| 20 | 20_71360768_G_A | 71360768 | A | G | LOC112443024(dist=10008),TRIP13( | intergenic | 3.9273E-10 | 0.31297 | 2.926E-12 | 0.017757 | 0.0084041 | 0.022231 | 0.002838 | 0.0083291 | 0.0031844 | 0.435982 | 0.472296 | 0.42926 |
| 20 | 20_71362074_T_A | 71362074 | A | T | LOC112443024(dist=11314),TRIP13( | intergenic | 8.5539E-11 | 0.31073 | 4.642E-13 | 0.018417 | 0.0084108 | 0.023053 | 0.0028375 | 0.0082971 | 0.0031861 | 0.437493 | 0.4753 | 0.430495 |
| 20 | 20_71362191_A_C | 71362191 | C | A | LOC112443024(dist=11431),TRIP13( | intergenic | 4.4676E-11 | 0.19751 | 4.377E-13 | 0.018659 | 0.01054 | 0.023091 | 0.0028324 | 0.0081789 | 0.0031879 | 0.438171 | 0.472964 | 0.431731 |
| 20 | 20_71362740_C_T | 71362740 | T | C | LOC112443024(dist=11980),TRIP13( | intergenic | 4.643E-09 | 0.5595 | 1.202E-11 | 0.016693 | 0.0049968 | 0.021627 | 0.0028489 | 0.0085623 | 0.0031899 | 0.460901 | 0.40988 | 0.470345 |
| 20 | 20_71362935_G_A | 71362935 | A | G | LOC112443024(dist=12175),TRIP13( | intergenic | 1.893E-10 | 0.35546 | 1.415E-12 | 0.018058 | 0.0076228 | 0.022562 | 0.0028349 | 0.0082494 | 0.0031856 | 0.437181 | 0.475634 | 0.430063 |
| 20 | 20_71363973_G_A | 71363973 | A | G | LOC112443024(dist=13213),TRIP13( | intergenic | 2.1444E-09 | 0.34784 | 3.121E-12 | 0.016956 | 0.0078077 | 0.02221 | 0.0028323 | 0.0083167 | 0.0031855 | 0.462621 | 0.411215 | 0.472136 |
| 20 | 20_71364008_T_G | 71364008 | G | T | LOC112443024(dist=13248),TRIP13( | intergenic | 1.1755E-10 | 0.1926 | 8.191E-13 | 0.018165 | 0.010528 | 0.022759 | 0.0028196 | 0.0080804 | 0.0031795 | 0.437598 | 0.478304 | 0.430063 |
| 20 | 20_71364398_G_A | 71364398 | A | G | LOC112443024(dist=13638),TRIP13( | intergenic | 5.5822E-11 | 0.23172 | 7.607E-13 | 0.018516 | 0.009811 | 0.022774 | 0.002825 | 0.0082036 | 0.0031772 | 0.437181 | 0.473965 | 0.430372 |
| 20 | 20_71364419_A_G | 71364419 | G | A | LOC112443024(dist=13659),TRIP13( | intergenic | 3.5579E-11 | 0.23687 | 5.425E-13 | 0.018698 | 0.0096902 | 0.022918 | 0.0028238 | 0.0081923 | 0.0031768 | 0.437546 | 0.474633 | 0.430681 |
| 20 | 20_71365376_C_T | 71365376 | T | C | LOC112443024(dist=14616),TRIP13( | intergenic | 4.9323E-11 | 0.41101 | 1.217E-13 | 0.018562 | 0.0067237 | 0.023563 | 0.002824 | 0.0081785 | 0.0031778 | 0.447711 | 0.488652 | 0.440133 |
| 20 | 20_71365864_G_A | 71365864 | A | G | LOC112443024(dist=15104),TRIP13( | intergenic | 4.7236E-12 | 0.34188 | 2.44E-14 | 0.019581 | 0.0078692 | 0.02427 | 0.0028323 | 0.0082795 | 0.0031829 | 0.437754 | 0.473298 | 0.431175 |
| 20 | 20_71366356_T_A | 71366356 | A | T | LOC112443024(dist=15596),TRIP13( | intergenic | 4.8414E-09 | 0.58239 | 8.437E-12 | 0.016698 | 0.0046791 | 0.02187 | 0.0028531 | 0.0085091 | 0.0032017 | 0.458607 | 0.408211 | 0.467935 |
| 20 | 20_71366508_T_C | 71366508 | C | T | LOC112443024(dist=15748),TRIP13( | intergenic | 2.9236E-08 | 0.59296 | 4.91E-11 | 0.015813 | 0.0045234 | 0.021044 | 0.0028513 | 0.0084619 | 0.0032012 | 0.45574 | 0.404873 | 0.465155 |
| 20 | 20_71368114_A_G | 71368114 | G | A | LOC112443024(dist=17354),TRIP13( | intergenic | 5.2791E-10 | 0.32029 | 1.94E-12 | 0.01764 | 0.0083156 | 0.022415 | 0.0028403 | 0.0083669 | 0.0031846 | 0.434366 | 0.469626 | 0.427839 |
| 20 | 20_71369075_T_C | 71369075 | C | T | LOC112443024(dist=18315),TRIP13( | intergenic | 6.0907E-10 | 0.3347 | 1.664E-12 | 0.017532 | 0.0079364 | 0.022497 | 0.0028332 | 0.0082268 | 0.0031865 | 0.434001 | 0.473298 | 0.426727 |
| 20 | 20_71369188_T_G | 71369188 | G | T | LOC112443024(dist=18428),TRIP13( | intergenic | 2.3385E-08 | 0.57371 | 2.441E-11 | 0.015847 | 0.0047293 | 0.021293 | 0.0028375 | 0.0084062 | 0.0031891 | 0.462048 | 0.410214 | 0.471642 |
| 20 | 20_71371635_A_G | 71371635 | G | A | LOC112443024(dist=20875),TRIP13( | intergenic | 3.2776E-09 | 0.73704 | 6.459E-12 | 0.01672 | 0.0027398 | 0.021876 | 0.0028258 | 0.0081595 | 0.0031847 | 0.437024 | 0.474967 | 0.430001 |
| 20 | 20_71371893_C_T | 71371893 | T | C | LOC112443024(dist=21133),TRIP13( | intergenic | 2.4133E-10 | 0.46496 | 6.484E-13 | 0.017975 | 0.0060472 | 0.022944 | 0.0028385 | 0.0082759 | 0.0031911 | 0.435148 | 0.46996 | 0.428704 |
| 20 | 20_71373981_T_C | 71373981 | C | T | LOC112443024(dist=23221),TRIP13( | intergenic | 8.429E-11 | 0.34265 | 3.148E-13 | 0.018415 | 0.0078201 | 0.023261 | 0.0028362 | 0.0082408 | 0.0031918 | 0.435982 | 0.471295 | 0.429445 |
| 20 | 20_71374588_G_A | 71374588 | A | G | TRIP13(dist=692) | downstream | 7.7748E-09 | 0.37653 | 1.847E-11 | 0.016586 | 0.0076162 | 0.021632 | 0.0028729 | 0.0086126 | 0.0032202 | 0.440621 | 0.38785 | 0.450389 |
| 20 | 20_71374595_G_A | 71374595 | A | G | TRIP13(dist=685) | downstream | 1.2251E-08 | 0.61121 | 1.326E-11 | 0.016292 | 0.004359 | 0.021688 | 0.0028602 | 0.0085749 | 0.0032055 | 0.456835 | 0.408879 | 0.465711 |
| 20 | 20_71374756_T_C | 71374756 | C | T | TRIP13(dist=524) | downstream | 1.4455E-10 | 0.45079 | 6.326E-13 | 0.018252 | 0.006353 | 0.02295 | 0.0028469 | 0.0084247 | 0.0031904 | 0.438432 | 0.474967 | 0.431669 |
| 20 | 20_71376099_T_C | 71376099 | C | T | TRIP13(XM_002696452.4:c.*32A>G,X | UTR3 | 2.4949E-10 | 0.36452 | 9.805E-13 | 0.017925 | 0.0074397 | 0.022746 | 0.002833 | 0.0082044 | 0.0031887 | 0.435252 | 0.468625 | 0.429075 |
| 20 | 20_71376761_T_C | 71376761 | C | T | TRIP13 | intronic | 2.251E-08 | 0.64393 | 2.117E-11 | 0.016035 | 0.0040373 | 0.021455 | 0.0028677 | 0.0087346 | 0.0032034 | 0.461579 | 0.419226 | 0.469418 |
| 20 | 20_71376914_G_C | 71376914 | C | G | TRIP13 | intronic | 2.5239E-08 | 0.69417 | 2.3E-11 | 0.015942 | 0.0034167 | 0.021389 | 0.0028613 | 0.0086893 | 0.0031993 | 0.464133 | 0.417557 | 0.472754 |
| 20 | 20_71377576_A_G | 71377576 | G | A | TRIP13 | intronic | 5.6661E-10 | 0.1692 | 6.476E-12 | 0.017555 | 0.011306 | 0.021875 | 0.0028317 | 0.0082241 | 0.0031848 | 0.435512 | 0.473298 | 0.428518 |
| 20 | 20_71377665_A_C | 71377665 | C | A | TRIP13 | intronic | 1.015E-09 | 0.22579 | 4.087E-12 | 0.017317 | 0.010018 | 0.022079 | 0.0028355 | 0.0082709 | 0.0031841 | 0.43374 | 0.474299 | 0.426233 |
| 20 | 20_71378023_G_A | 71378023 | A | G | TRIP13 | intronic | 1.8877E-10 | 0.29852 | 1.828E-12 | 0.018166 | 0.0088322 | 0.022484 | 0.0028517 | 0.0084957 | 0.0031905 | 0.434574 | 0.466956 | 0.42858 |
| 20 | 20_71378649_G_A | 71378649 | A | G | TRIP13 | intronic | 2.7036E-08 | 0.60561 | 1.665E-11 | 0.015872 | 0.0044317 | 0.021519 | 0.0028549 | 0.0085829 | 0.0031962 | 0.460327 | 0.408545 | 0.469912 |
| 20 | 20_71378946_A_G | 71378946 | G | A | TRIP13 | intronic | 6.7015E-11 | 0.28973 | 4.286E-13 | 0.018516 | 0.0087131 | 0.023105 | 0.0028368 | 0.0082298 | 0.0031886 | 0.436555 | 0.474299 | 0.429569 |

| | | | | | | | | | | | | | | | | | | |
|---|---|---|---|---|---|---|---|---|---|---|---|---|---|---|---|---|---|---|
| 20 | 20_71379551_A_G | 71379551 | G | A | TRIP13 | intronic | 6.5033E-10 | 0.27636 | 2.261E-12 | 0.017491 | 0.0089725 | 0.022338 | 0.0028313 | 0.0082428 | 0.0031832 | 0.437858 | 0.475968 | 0.430804 |
| 20 | 20_71379902_T_C | 71379902 | C | T | TRIP13 | intronic | 4.9627E-10 | 0.30305 | 8.352E-12 | 0.017701 | 0.008692 | 0.021777 | 0.0028458 | 0.0084395 | 0.0031873 | 0.433375 | 0.469292 | 0.426727 |
| 20 | 20_71379948_A_G | 71379948 | G | A | TRIP13 | intronic | 1.0261E-09 | 0.37459 | 7.566E-12 | 0.017348 | 0.0074668 | 0.021798 | 0.0028415 | 0.0084096 | 0.0031838 | 0.433479 | 0.467957 | 0.427097 |
| 20 | 20_71380308_C_T | 71380308 | T | C | TRIP13 | intronic | 9.9246E-11 | 0.2829 | 2.911E-12 | 0.0183 | 0.0088229 | 0.022218 | 0.0028292 | 0.0082163 | 0.0031823 | 0.437598 | 0.476302 | 0.430434 |
| 20 | 20_71380598_T_C | 71380598 | C | T | TRIP13 | intronic | 7.5217E-10 | 0.36086 | 4.276E-12 | 0.017486 | 0.0076298 | 0.022088 | 0.002841 | 0.0083501 | 0.0031884 | 0.435043 | 0.472964 | 0.428024 |
| 20 | 20_71380906_A_T | 71380906 | T | A | TRIP13 | intronic | 3.3264E-10 | 0.41645 | 2.002E-12 | 0.017788 | 0.0066654 | 0.022406 | 0.0028312 | 0.0082027 | 0.0031852 | 0.437441 | 0.47964 | 0.429631 |
| 20 | 20_71380970_A_G | 71380970 | G | A | TRIP13 | intronic | 6.4598E-10 | 0.45651 | 4.432E-12 | 0.017503 | 0.0061424 | 0.022043 | 0.0028327 | 0.0082492 | 0.0031841 | 0.436242 | 0.476302 | 0.428827 |
| 20 | 20_71381221_A_G | 71381221 | G | A | TRIP13 | intronic | 1.9942E-10 | 0.38598 | 1.331E-12 | 0.018103 | 0.0072645 | 0.022644 | 0.0028456 | 0.0083796 | 0.0031934 | 0.437546 | 0.471963 | 0.431175 |
| 20 | 20_71381258_C_G | 71381258 | G | C | TRIP13 | intronic | 9.1232E-10 | 0.71738 | 2.596E-12 | 0.017306 | 0.0030838 | 0.022134 | 0.002826 | 0.0085199 | 0.003163 | 0.491763 | 0.476636 | 0.485914 |
| 20 | 20_71382984_A_G | 71382984 | G | A | TRIP13 | intronic | 7.2708E-11 | 0.48811 | 6.36E-13 | 0.018478 | 0.0057321 | 0.02294 | 0.0028363 | 0.0082677 | 0.0031894 | 0.436868 | 0.471629 | 0.430434 |
| 20 | 20_71384455_C_T | 71384455 | T | C | TRIP13 | intronic | 8.5703E-10 | 0.32822 | 5.698E-12 | 0.017386 | 0.0081088 | 0.02192 | 0.0028344 | 0.0082936 | 0.0031828 | 0.436607 | 0.474299 | 0.429631 |
| 20 | 20_71384468_C_T | 71384468 | T | C | TRIP13 | intronic | 6.3939E-10 | 0.2812 | 4.888E-12 | 0.017518 | 0.0089741 | 0.021979 | 0.0028344 | 0.0083275 | 0.0031813 | 0.436399 | 0.472964 | 0.429631 |
| 20 | 20_71385114_G_C | 71385114 | C | G | TRIP13 | intronic | 5.1545E-10 | 0.33159 | 2.259E-12 | 0.017647 | 0.0081111 | 0.022355 | 0.0028397 | 0.008354 | 0.0031856 | 0.434209 | 0.467957 | 0.427962 |
| 20 | 20_71385601_T_A | 71385601 | A | T | TRIP13 | intronic | 2.2354E-09 | 0.31279 | 1.785E-11 | 0.016919 | 0.0083084 | 0.021382 | 0.0028295 | 0.0082312 | 0.0031807 | 0.433792 | 0.473632 | 0.426418 |
| 20 | 20_71385619_A_G | 71385619 | G | A | TRIP13 | intronic | 2.1988E-09 | 0.32206 | 1.69E-11 | 0.016942 | 0.0081453 | 0.02143 | 0.002832 | 0.0082256 | 0.0031841 | 0.434053 | 0.473965 | 0.426665 |
| 20 | 20_71385684_C_T | 71385684 | T | C | TRIP13 | intronic | 2.0399E-08 | 0.307 | 1.128E-10 | 0.01587 | 0.0086259 | 0.020477 | 0.0028295 | 0.008444 | 0.0031753 | 0.462778 | 0.413885 | 0.471828 |
| 20 | 20_71385700_A_C | 71385700 | C | A | TRIP13 | intronic | 2.7517E-10 | 0.11912 | 1.108E-11 | 0.017833 | 0.01279 | 0.021591 | 0.0028251 | 0.0082065 | 0.0031789 | 0.434887 | 0.473632 | 0.427715 |
| 20 | 20_71385750_T_C | 71385750 | C | T | TRIP13 | intronic | 2.2044E-10 | 0.12578 | 7.474E-12 | 0.017971 | 0.01264 | 0.021805 | 0.0028317 | 0.0082566 | 0.0031841 | 0.43546 | 0.471963 | 0.428704 |
| 20 | 20_71386538_C_G | 71386538 | G | C | TRIP13 | intronic | 5.8381E-10 | 0.4496 | 4.012E-12 | 0.017597 | 0.0062547 | 0.02215 | 0.0028406 | 0.0082724 | 0.0031932 | 0.436764 | 0.468959 | 0.430804 |
| 20 | 20_71386554_T_C | 71386554 | C | T | TRIP13 | intronic | 1.1072E-09 | 0.28272 | 6.736E-12 | 0.017316 | 0.0089406 | 0.021891 | 0.002842 | 0.0083227 | 0.0031897 | 0.435043 | 0.47263 | 0.428086 |
| 20 | 20_71386768_G_A | 71386768 | A | G | TRIP13 | intronic | 3.3508E-10 | 0.19059 | 3.571E-12 | 0.017779 | 0.010777 | 0.02213 | 0.0028303 | 0.0082339 | 0.0031827 | 0.434522 | 0.470961 | 0.427777 |
| 20 | 20_71386992_C_T | 71386992 | T | C | TRIP13 | intronic | 8.254E-10 | 0.32379 | 6.172E-12 | 0.017443 | 0.0082584 | 0.02191 | 0.0028408 | 0.0083697 | 0.0031866 | 0.432749 | 0.468291 | 0.426171 |
| 20 | 20_71387739_T_C | 71387739 | C | T | TRIP13 | intronic | 3.5064E-11 | 0.22679 | 5.164E-13 | 0.018756 | 0.0098534 | 0.023028 | 0.0028317 | 0.0081523 | 0.0031891 | 0.437546 | 0.473965 | 0.430804 |
| 20 | 20_71388363_A_G | 71388363 | G | A | TRIP13 | intronic | 6.5627E-10 | 0.28234 | 8.45E-12 | 0.017521 | 0.0089093 | 0.021765 | 0.0028368 | 0.0082872 | 0.0031863 | 0.436868 | 0.474633 | 0.429878 |
| 20 | 20_71392416_T_C | 71392416 | C | T | BRD9 | intronic | 1.5284E-10 | 0.23421 | 1.414E-12 | 0.018098 | 0.0097766 | 0.022505 | 0.0028268 | 0.0082185 | 0.0031774 | 0.431238 | 0.471295 | 0.423823 |
| 20 | 20_71392419_A_G | 71392419 | G | A | BRD9 | intronic | 6.6452E-11 | 0.24765 | 4.718E-13 | 0.018478 | 0.0095016 | 0.023015 | 0.0028304 | 0.0082187 | 0.0031819 | 0.431707 | 0.471295 | 0.424379 |
| 20 | 20_71392525_A_G | 71392525 | G | A | BRD9 | intronic | 4.581E-11 | 0.27557 | 1.037E-13 | 0.018675 | 0.0090186 | 0.023701 | 0.0028365 | 0.0082714 | 0.0031873 | 0.435617 | 0.477971 | 0.427777 |
| 20 | 20_71393175_C_T | 71393175 | T | C | BRD9 | intronic | 6.7342E-10 | 0.29768 | 1.36E-12 | 0.017514 | 0.0086371 | 0.022594 | 0.0028376 | 0.0082936 | 0.0031875 | 0.430143 | 0.468291 | 0.423082 |
| 20 | 20_71393225_A_G | 71393225 | G | A | BRD9 | intronic | 4.2864E-10 | 0.30903 | 1.034E-12 | 0.017626 | 0.0083383 | 0.022626 | 0.0028232 | 0.0081968 | 0.0031752 | 0.431238 | 0.469626 | 0.424132 |
| 20 | 20_71393623_A_G | 71393623 | G | A | BRD9 | intronic | 3.57E-11 | 0.17588 | 1.964E-13 | 0.018757 | 0.01127 | 0.023365 | 0.002833 | 0.0083265 | 0.0031784 | 0.433219 | 0.466956 | 0.426974 |
| 20 | 20_71393864_C_T | 71393864 | T | C | BRD9 | intronic | 9.5535E-11 | 0.17715 | 8.604E-13 | 0.018405 | 0.011332 | 0.022781 | 0.002843 | 0.008397 | 0.0031856 | 0.43228 | 0.46729 | 0.4258 |
| 20 | 20_71395443_G_A | 71395443 | A | G | BRD9 | intronic | 8.9946E-11 | 0.39231 | 2.048E-13 | 0.018355 | 0.0070825 | 0.023356 | 0.0028313 | 0.0082793 | 0.0031795 | 0.432749 | 0.467957 | 0.426233 |
| 20 | 20_71395557_A_G | 71395557 | G | A | BRD9 | intronic | 1.9642E-10 | 0.18623 | 6.91E-13 | 0.018033 | 0.010899 | 0.022863 | 0.0028335 | 0.0082455 | 0.0031837 | 0.431342 | 0.470294 | 0.424132 |
| 20 | 20_71395895_G_A | 71395895 | A | G | BRD9 | intronic | 1.7731E-10 | 0.21143 | 1.845E-12 | 0.017973 | 0.010165 | 0.022355 | 0.0028172 | 0.008134 | 0.0031728 | 0.431029 | 0.47263 | 0.423329 |
| 20 | 20_71395924_G_A | 71395924 | A | G | BRD9 | intronic | 6.8427E-11 | 0.15239 | 5.158E-13 | 0.01839 | 0.011649 | 0.022917 | 0.0028188 | 0.0081395 | 0.0031737 | 0.431915 | 0.474633 | 0.424008 |
| 20 | 20_71395933_C_T | 71395933 | T | C | BRD9 | intronic | 1.1105E-10 | 0.21489 | 5.565E-13 | 0.018166 | 0.010091 | 0.022863 | 0.0028159 | 0.0081364 | 0.0031706 | 0.432124 | 0.474299 | 0.424317 |
| 20 | 20_71395953_G_A | 71395953 | A | G | BRD9 | intronic | 1.5385E-10 | 0.26771 | 5.128E-13 | 0.01804 | 0.0090313 | 0.02291 | 0.0028181 | 0.0081484 | 0.0031723 | 0.431759 | 0.473632 | 0.424008 |
| 20 | 20_71395969_C_T | 71395969 | T | C | BRD9 | intronic | 3.5913E-10 | 0.2778 | 1.268E-12 | 0.017687 | 0.0088643 | 0.022524 | 0.0028205 | 0.0081678 | 0.0031733 | 0.432072 | 0.474299 | 0.424256 |
| 20 | 20_71395994_C_A | 71395994 | A | C | BRD9 | intronic | 2.7445E-10 | 0.37956 | 5.016E-13 | 0.017834 | 0.0071771 | 0.022959 | 0.0028251 | 0.0081679 | 0.0031778 | 0.431238 | 0.474967 | 0.423143 |
| 20 | 20_71396004_T_G | 71396004 | G | T | BRD9 | intronic | 2.3206E-08 | 0.58586 | 2.305E-11 | 0.015856 | 0.004667 | 0.021247 | 0.0028384 | 0.0085658 | 0.0031782 | 0.468147 | 0.42223 | 0.476646 |
| 20 | 20_71396007_G_T | 71396007 | T | G | BRD9 | intronic | 1.5677E-10 | 0.40884 | 2.93E-13 | 0.018076 | 0.0067549 | 0.02319 | 0.002825 | 0.0081784 | 0.0031778 | 0.431967 | 0.474967 | 0.424008 |
| 20 | 20_71396187_A_G | 71396187 | G | A | BRD9 | exonic | 5.5286E-10 | 0.33742 | 1.817E-12 | 0.017584 | 0.0079456 | 0.022434 | 0.0028347 | 0.0082829 | 0.003183 | 0.430404 | 0.471295 | 0.422835 |
| 20 | 20_71396193_C_T | 71396193 | T | C | BRD9 | exonic | 5.5286E-10 | 0.33742 | 1.817E-12 | 0.017584 | 0.0079456 | 0.022434 | 0.0028347 | 0.0082829 | 0.003183 | 0.430404 | 0.471295 | 0.422835 |
| 20 | 20_71396606_C_T | 71396606 | T | C | BRD9 | intronic | 2.0729E-10 | 0.31706 | 8.453E-13 | 0.017961 | 0.008101 | 0.022775 | 0.0028258 | 0.0080968 | 0.0031838 | 0.434157 | 0.474967 | 0.426603 |
| 20 | 20_71397314_C_T | 71397314 | T | C | BRD9 | intronic | 1.1846E-08 | 0.62634 | 1.913E-11 | 0.016117 | 0.0041087 | 0.021299 | 0.0028266 | 0.0084385 | 0.003173 | 0.46773 | 0.417223 | 0.477079 |
| 20 | 20_71397349_A_G | 71397349 | G | A | BRD9 | intronic | 1.2658E-10 | 0.3066 | 5.769E-13 | 0.018196 | 0.0083905 | 0.022934 | 0.0028293 | 0.0082068 | 0.0031827 | 0.431342 | 0.47263 | 0.423699 |
| 20 | 20_71397651_G_A | 71397651 | A | G | BRD9 | intronic | 1.35E-10 | 0.4036 | 6.926E-13 | 0.018152 | 0.0068639 | 0.022808 | 0.0028268 | 0.0082182 | 0.0031762 | 0.432176 | 0.47263 | 0.424688 |
| 20 | 20_71398468_C_A | 71398468 | A | C | BRD9 | intronic | 2.0042E-10 | 0.13489 | 1.802E-12 | 0.018003 | 0.012216 | 0.022458 | 0.0028302 | 0.0081705 | 0.003186 | 0.432332 | 0.473965 | 0.424626 |
| 20 | 20_71398529_C_T | 71398529 | T | C | BRD9 | intronic | 2.8936E-10 | 0.23995 | 2.128E-12 | 0.017889 | 0.0096604 | 0.022417 | 0.0028375 | 0.0082209 | 0.0031907 | 0.431342 | 0.472964 | 0.423638 |
| 20 | 20_71398655_A_G | 71398655 | G | A | BRD9 | intronic | 3.507E-11 | 0.30445 | 2.554E-13 | 0.018777 | 0.0084905 | 0.023295 | 0.0028349 | 0.0082678 | 0.0031842 | 0.430821 | 0.467957 | 0.423947 |
| 20 | 20_71399130_A_G | 71399130 | G | A | BRD9 | intronic | 5.3203E-11 | 0.27947 | 4.439E-13 | 0.018607 | 0.0089691 | 0.02304 | 0.0028357 | 0.008293 | 0.0031817 | 0.431498 | 0.474299 | 0.423576 |
| 20 | 20_71399255_T_C | 71399255 | C | T | BRD9 | intronic | 4.6645E-12 | 0.24692 | 1.916E-14 | 0.019557 | 0.0094491 | 0.024353 | 0.002828 | 0.0081607 | 0.0031808 | 0.431342 | 0.4753 | 0.423205 |
| 20 | 20_71399484_C_T | 71399484 | T | C | BRD9 | intronic | 3.2865E-11 | 0.2019 | 2.631E-13 | 0.018764 | 0.010458 | 0.023239 | 0.0028288 | 0.0081945 | 0.0031782 | 0.432958 | 0.479973 | 0.424256 |
| 20 | 20_71399901_T_C | 71399901 | C | T | BRD9 | intronic | 5.5449E-11 | 0.28154 | 2.734E-13 | 0.018582 | 0.0088532 | 0.023278 | 0.0028346 | 0.0082212 | 0.0031857 | 0.434053 | 0.480307 | 0.425491 |
| 20 | 20_71401122_G_C | 71401122 | C | G | BRD9 | intronic | 4.7017E-11 | 0.30305 | 1.979E-13 | 0.018607 | 0.0084624 | 0.023344 | 0.0028278 | 0.0082165 | 0.0031759 | 0.43374 | 0.476302 | 0.425862 |
| 20 | 20_71401124_C_T | 71401124 | T | C | BRD9 | intronic | 1.4531E-11 | 0.19768 | 1.121E-13 | 0.019032 | 0.010462 | 0.023557 | 0.0028184 | 0.0081215 | 0.0031723 | 0.43301 | 0.473965 | 0.425429 |
| 20 | 20_71401256_G_A | 71401256 | A | G | BRD9 | intronic | 5.985E-11 | 0.24796 | 1.929E-13 | 0.018498 | 0.0094723 | 0.023351 | 0.0028266 | 0.0081988 | 0.0031754 | 0.432697 | 0.477303 | 0.424441 |
| 20 | 20_71401617_T_C | 71401617 | C | T | BRD9 | intronic | 9.9586E-10 | 0.28724 | 3.708E-12 | 0.017173 | 0.0085868 | 0.021996 | 0.0028106 | 0.0080689 | 0.0031659 | 0.431863 | 0.480975 | 0.422773 |
| 20 | 20_71401678_T_C | 71401678 | C | T | BRD9 | intronic | 7.5576E-09 | 0.43523 | 7.497E-12 | 0.016344 | 0.0066469 | 0.021686 | 0.0028286 | 0.0085187 | 0.0031668 | 0.46747 | 0.410214 | 0.478067 |
| 20 | 20_71401763_C_A | 71401763 | A | C | BRD9 | intronic | 2.106E-11 | 0.063397 | 1.342E-12 | 0.018818 | 0.015014 | 0.022437 | 0.0028093 | 0.0080878 | 0.0031647 | 0.431185 | 0.473298 | 0.423391 |
| 20 | 20_71401861_G_A | 71401861 | A | G | BRD9 | intronic | 7.378E-09 | 0.26939 | 2.6E-11 | 0.016309 | 0.0093894 | 0.021051 | 0.0028206 | 0.0085013 | 0.0031572 | 0.465436 | 0.410881 | 0.475534 |
| 20 | 20_71401872_G_C | 71401872 | C | G | BRD9 | intronic | 6.0603E-09 | 0.25045 | 1.413E-11 | 0.016379 | 0.0097161 | 0.02132 | 0.0028166 | 0.0084544 | 0.0031555 | 0.469346 | 0.414887 | 0.479427 |
| 20 | 20_71401934_G_C | 71401934 | C | G | BRD9 | intronic | 4.1193E-09 | 0.31307 | 2.004E-11 | 0.016584 | 0.0085219 | 0.021205 | 0.0028208 | 0.0084476 | 0.0031622 | 0.469033 | 0.417223 | 0.478624 |
| 20 | 20_71401981_A_G | 71401981 | G | A | BRD9 | intronic | 1.4657E-09 | 0.20454 | 8.592E-12 | 0.017094 | 0.010764 | 0.021628 | 0.0028264 | 0.008484 | 0.0031675 | 0.469242 | 0.419559 | 0.478438 |
| 20 | 20_71402002_A_G | 71402002 | G | A | BRD9 | intronic | 1.7239E-09 | 0.20373 | 6.645E-12 | 0.017037 | 0.010783 | 0.021762 | 0.0028291 | 0.0084837 | 0.00317 | 0.468251 | 0.416555 | 0.47782 |
| 20 | 20_71402113_G_C | 71402113 | C | G | BRD9 | intronic | 1.1636E-09 | 0.31733 | 2.923E-12 | 0.017187 | 0.008478 | 0.022093 | 0.0028245 | 0.0084784 | 0.0031646 | 0.468512 | 0.412884 | 0.478809 |
| 20 | 20_71402158_A_G | 71402158 | G | A | BRD9 | intronic | 8.7817E-10 | 0.424 | 1.082E-12 | 0.017316 | 0.00675 | 0.022548 | 0.0028247 | 0.0084429 | 0.003167 | 0.466844 | 0.409546 | 0.47745 |
| 20 | 20_71402186_A_G | 71402186 | G | A | BRD9 | intronic | 3.5849E-10 | 0.53362 | 7.552E-13 | 0.017715 | 0.0053026 | 0.02268 | 0.0028249 | 0.0085183 | 0.0031635 | 0.468095 | 0.413218 | 0.478253 |
| 20 | 20_71402241_T_C | 71402241 | C | T | BRD9 | intronic | 1.4108E-09 | 0.33968 | 3.725E-12 | 0.017108 | 0.0081135 | 0.021988 | 0.0028258 | 0.0084975 | 0.0031651 | 0.468199 | 0.413551 | 0.478315 |
| 20 | 20_71402259_A_T | 71402259 | T | A | BRD9 | intronic | 1.4344E-09 | 0.32495 | 4.131E-12 | 0.017128 | 0.0083603 | 0.02199 | 0.0028303 | 0.0084934 | 0.0031719 | 0.468043 | 0.414219 | 0.478006 |
| 20 | 20_71402409_A_G | 71402409 | G | A | BRD9 | intronic | 1.0595E-10 | 0.30727 | 6.667E-13 | 0.018247 | 0.0084542 | 0.022764 | 0.0028254 | 0.0082806 | 0.0031678 | 0.429257 | 0.475634 | 0.420672 |
| 20 | 20_71402440_T_C | 71402440 | C | T | BRD9 | intronic | 3.055E-10 | 0.50408 | 1.286E-12 | 0.017731 | 0.0055041 | 0.022425 | 0.0028163 | 0.0082386 | 0.0031604 | 0.428475 | 0.473298 | 0.420178 |
| 20 | 20_71402464_T_C | 71402464 | C | T | BRD9 | intronic | 2.3747E-10 | 0.71163 | 6.812E-13 | 0.01784 | 0.0030471 | 0.022704 | 0.0028161 | 0.0082429 | 0.0031607 | 0.428266 | 0.471295 | 0.420301 |
| 20 | 20_71402557_T_C | 71402557 | C | T | BRD9 | intronic | 2.463E-11 | 0.29355 | 3.366E-13 | 0.018841 | 0.0085792 | 0.023102 | 0.0028223 | 0.0081678 | 0.0031739 | 0.432489 | 0.477971 | 0.42407 |
| 20 | 20_71402777_T_C | 71402777 | C | T | BRD9 | intronic | 1.7866E-10 | 0.33288 | 4.225E-13 | 0.017999 | 0.0079218 | 0.022991 | 0.0028218 | 0.0081809 | 0.003172 | 0.431655 | 0.472964 | 0.424008 |

| | | | | | | | | | | | | | | | | | | |
|---|---|---|---|---|---|---|---|---|---|---|---|---|---|---|---|---|---|---|
| 20 | 20_71402820_T_C | 71402820 | C | T | BRD9 | intronic | 5.7506E-11 | 0.30817 | 2.883E-13 | 0.018518 | 0.0084178 | 0.023173 | 0.0028271 | 0.0082602 | 0.0031745 | 0.430873 | 0.471963 | 0.423267 |
| 20 | 20_71402849_A_G | 71402849 | G | A | BRD9 | intronic | 1.041E-10 | 0.40738 | 2.347E-13 | 0.018242 | 0.0068602 | 0.023218 | 0.0028234 | 0.0082801 | 0.0031687 | 0.429465 | 0.470294 | 0.421908 |
| 20 | 20_71402925_G_T | 71402925 | T | G | BRD9 | intronic | 4.9942E-09 | 0.2356 | 1.658E-11 | 0.016523 | 0.0098668 | 0.021381 | 0.0028257 | 0.0083191 | 0.0031755 | 0.476384 | 0.481976 | 0.468677 |
| 20 | 20_71403320_C_A | 71403320 | A | C | BRD9 | intronic | 1.8797E-10 | 0.20931 | 4.777E-13 | 0.018033 | 0.010329 | 0.022999 | 0.0028306 | 0.0082275 | 0.0031804 | 0.428996 | 0.474967 | 0.420487 |
| 20 | 20_71403559_C_T | 71403559 | T | C | BRD9 | intronic | 2.2986E-11 | 0.16595 | 1.382E-13 | 0.018873 | 0.011331 | 0.023495 | 0.0028229 | 0.0081793 | 0.0031759 | 0.431029 | 0.472296 | 0.423391 |
| 20 | 20_71404508_T_C | 71404508 | C | T | BRD9 | intronic | 1.1883E-11 | 0.15716 | 2.281E-13 | 0.019151 | 0.011484 | 0.023307 | 0.002824 | 0.0081177 | 0.0031792 | 0.435148 | 0.47964 | 0.426912 |
| 20 | 20_71404704_C_G | 71404704 | G | C | BRD9 | intronic | 5.7832E-11 | 0.20601 | 2.921E-13 | 0.018472 | 0.010183 | 0.023219 | 0.0028205 | 0.0080524 | 0.0031815 | 0.430195 | 0.473965 | 0.422093 |
| 20 | 20_71405569_C_T | 71405569 | T | C | BRD9 | intronic | 5.7117E-10 | 0.55282 | 2.248E-12 | 0.017482 | 0.0048561 | 0.022343 | 0.0028206 | 0.0081817 | 0.0031836 | 0.465958 | 0.411215 | 0.47609 |
| 20 | 20_71405810_A_G | 71405810 | G | A | BRD9 | intronic | 1.1246E-11 | 0.21366 | 2.331E-13 | 0.019217 | 0.01012 | 0.023352 | 0.0028304 | 0.008138 | 0.0031866 | 0.43447 | 0.476636 | 0.426665 |
| 20 | 20_71406170_A_G | 71406170 | G | A | BRD9 | intronic | 2.7948E-09 | 0.27945 | 1.914E-11 | 0.016762 | 0.0088521 | 0.021331 | 0.0028203 | 0.0081847 | 0.0031779 | 0.498071 | 0.447931 | 0.488076 |
| 20 | 20_71406322_C_T | 71406322 | T | C | BRD9 | intronic | 1.1228E-09 | 0.31963 | 2.624E-12 | 0.017195 | 0.008146 | 0.022267 | 0.0028231 | 0.008185 | 0.0031826 | 0.497915 | 0.449599 | 0.4882 |
| 20 | 20_71406618_C_G | 71406618 | G | C | BRD9 | intronic | 2.7188E-09 | 0.41774 | 8.935E-12 | 0.016908 | 0.0068351 | 0.021768 | 0.0028427 | 0.0084347 | 0.0031905 | 0.464394 | 0.410881 | 0.474299 |
| 20 | 20_71407025_A_G | 71407025 | G | A | BRD9 | intronic | 7.6665E-11 | 0.24073 | 6.703E-13 | 0.018339 | 0.0093795 | 0.022885 | 0.0028183 | 0.0079951 | 0.0031849 | 0.435304 | 0.481308 | 0.426789 |
| 20 | 20_71407893_T_C | 71407893 | C | T | BRD9 | intronic | 1.7225E-09 | 0.15916 | 4.787E-11 | 0.017244 | 0.011767 | 0.021192 | 0.0028635 | 0.0083581 | 0.003222 | 0.404702 | 0.449266 | 0.396454 |
| 20 | 20_71408529_C_T | 71408529 | T | C | BRD9(XM_005221673.4:c.*170C>T,N | UTR3 | 6.8311E-10 | 0.27039 | 2.212E-11 | 0.017343 | 0.009036 | 0.021138 | 0.0028108 | 0.0081983 | 0.003159 | 0.491659 | 0.463618 | 0.496849 |
| 20 | 20_71411361_C_G | 71411361 | G | C | BRD9(dist=2446),LOC617224(dist=68 | intergenic | 1.4953E-08 | 0.35665 | 4.246E-10 | 0.015897 | 0.0076411 | 0.019659 | 0.0028077 | 0.0082896 | 0.0031481 | 0.493901 | 0.467957 | 0.498703 |
| 20 | 20_71417623_T_C | 71417623 | C | T | LOC617224(dist=636) | upstream | 7.3702E-10 | 0.36602 | 3.888E-12 | 0.017412 | 0.0074764 | 0.022045 | 0.0028276 | 0.0082709 | 0.003176 | 0.443593 | 0.483645 | 0.436179 |
| 20 | 20_71417996_C_T | 71417996 | T | C | LOC617224(dist=263) | upstream | 1.8024E-11 | 0.21779 | 1.084E-13 | 0.01909 | 0.010343 | 0.023654 | 0.0028402 | 0.0083923 | 0.0031835 | 0.432332 | 0.46996 | 0.425368 |
| 20 | 20_71418078_C_G | 71418078 | G | C | LOC617224(dist=181) | upstream | 4.2972E-11 | 0.23401 | 3.364E-13 | 0.018744 | 0.0099608 | 0.023203 | 0.0028428 | 0.0083698 | 0.0031876 | 0.431185 | 0.467623 | 0.424441 |
| 20 | 20_71418200_A_G | 71418200 | G | A | LOC617224(dist=59) | upstream | 8.7429E-11 | 0.2255 | 7.054E-13 | 0.018416 | 0.010075 | 0.02288 | 0.0028388 | 0.0083121 | 0.0031873 | 0.430456 | 0.466622 | 0.423761 |
| 20 | 20_71418771_T_C | 71418771 | C | T | LOC617224 | intronic | 1.6227E-11 | 0.14594 | 8.318E-13 | 0.019095 | 0.01203 | 0.022804 | 0.0028346 | 0.0082736 | 0.0031868 | 0.435043 | 0.466288 | 0.42926 |
| 20 | 20_71418972_G_C | 71418972 | C | G | LOC617224 | intronic | 4.7718E-09 | 0.4239 | 1.652E-11 | 0.01666 | 0.0068425 | 0.021461 | 0.0028455 | 0.0085565 | 0.0031871 | 0.465488 | 0.41255 | 0.475287 |
| 20 | 20_71419169_C_T | 71419169 | T | C | LOC617224 | intronic | 5.5776E-09 | 0.33651 | 3.433E-11 | 0.01656 | 0.0081309 | 0.021129 | 0.002841 | 0.0084601 | 0.0031885 | 0.464081 | 0.404873 | 0.47504 |
| 20 | 20_71419930_G_A | 71419930 | A | G | LOC617224 | intronic | 5.9855E-10 | 0.18475 | 9.466E-12 | 0.017545 | 0.010819 | 0.021756 | 0.0028341 | 0.0081577 | 0.0031927 | 0.433062 | 0.467623 | 0.426665 |
| 20 | 20_71420736_T_C | 71420736 | C | T | LOC617224 | intronic | 5.4245E-11 | 0.09985 | 1.287E-12 | 0.018576 | 0.013516 | 0.02262 | 0.0028323 | 0.0082138 | 0.0031878 | 0.432541 | 0.471963 | 0.425244 |
| 20 | 20_71424058_A_G | 71424058 | G | A | LOC617224 | intronic | 4.3163E-10 | 0.20551 | 1.098E-11 | 0.017651 | 0.010503 | 0.021585 | 0.0028278 | 0.0082964 | 0.0031775 | 0.437493 | 0.471629 | 0.431175 |
| 20 | 20_71426535_A_G | 71426535 | G | A | LOC617224 | exonic | 2.1358E-08 | 0.42196 | 3.073E-11 | 0.015898 | 0.0067792 | 0.021172 | 0.0028385 | 0.008442 | 0.0031871 | 0.456417 | 0.400868 | 0.4667 |
| 20 | 20_71430259_T_C | 71430259 | C | T | LOC617224 | intronic | 7.9475E-11 | 0.37362 | 8.904E-13 | 0.018439 | 0.0074594 | 0.022726 | 0.0028361 | 0.008384 | 0.0031801 | 0.435148 | 0.466956 | 0.42926 |
| 20 | 20_71430402_A_G | 71430402 | G | A | LOC617224 | intronic | 7.8859E-11 | 0.35637 | 6.596E-13 | 0.018516 | 0.0077994 | 0.022924 | 0.0028473 | 0.0084564 | 0.0031894 | 0.435043 | 0.469626 | 0.428642 |
| 20 | 20_71431631_G_T | 71431631 | T | G | LOC617224 | intronic | 1.6912E-10 | 0.37071 | 1.672E-12 | 0.018113 | 0.0073957 | 0.022497 | 0.0028359 | 0.0082619 | 0.0031868 | 0.436242 | 0.477971 | 0.428518 |
| 20 | 20_71432846_C_G | 71432846 | G | C | LOC617224 | intronic | 6.1312E-11 | 0.22866 | 7.62E-13 | 0.018538 | 0.0099539 | 0.022824 | 0.0028343 | 0.0082687 | 0.0031842 | 0.431967 | 0.469626 | 0.424997 |
| 20 | 20_71433671_C_T | 71433671 | T | C | LOC617224 | intronic | 1.8268E-10 | 0.21961 | 1.56E-12 | 0.0181 | 0.010124 | 0.022555 | 0.0028391 | 0.0082475 | 0.0031906 | 0.432593 | 0.469626 | 0.425738 |
| 20 | 20_71433810_T_C | 71433810 | C | T | LOC617224 | intronic | 1.3426E-11 | 0.17181 | 2.304E-13 | 0.019239 | 0.011327 | 0.023414 | 0.0028443 | 0.0082896 | 0.0031944 | 0.42884 | 0.466622 | 0.421846 |
| 20 | 20_71435027_T_C | 71435027 | C | T | LOC617224 | intronic | 2.5637E-11 | 0.13659 | 2.738E-13 | 0.018758 | 0.011792 | 0.023274 | 0.0028124 | 0.0079215 | 0.0031854 | 0.435148 | 0.474299 | 0.427901 |
| 20 | 20_71435752_A_G | 71435752 | G | A | LOC617224 | intronic | 7.4567E-12 | 0.17065 | 1.677E-13 | 0.019284 | 0.011085 | 0.023395 | 0.0028158 | 0.0080907 | 0.0031733 | 0.434991 | 0.477637 | 0.427097 |
| 20 | 20_71436317_A_G | 71436317 | G | A | LOC617224 | intronic | 4.4816E-09 | 0.44108 | 1.163E-11 | 0.016724 | 0.0064942 | 0.021738 | 0.0028513 | 0.00843 | 0.003204 | 0.461735 | 0.406208 | 0.472013 |
| 20 | 20_71436498_G_C | 71436498 | C | G | LOC617224 | intronic | 6.0495E-10 | 0.30889 | 2.199E-12 | 0.017342 | 0.0082253 | 0.022185 | 0.0028019 | 0.0080834 | 0.0031598 | 0.454384 | 0.496662 | 0.445323 |
| 20 | 20_71438212_C_T | 71438212 | T | C | LOC617224 | intronic | 2.7966E-10 | 0.44501 | 8.087E-13 | 0.017753 | 0.006188 | 0.022707 | 0.0028136 | 0.0081019 | 0.0031715 | 0.428787 | 0.466956 | 0.421722 |
| 20 | 20_71439982_T_A | 71439982 | A | T | LOC617224 | intronic | 1.3182E-08 | 0.55187 | 2.513E-11 | 0.015972 | 0.0050527 | 0.020992 | 0.0028101 | 0.0084925 | 0.003146 | 0.470858 | 0.422897 | 0.479736 |
| 20 | 20_71439996_T_C | 71439996 | C | T | LOC617224 | intronic | 2.3576E-08 | 0.66916 | 2.697E-11 | 0.015702 | 0.0036059 | 0.021 | 0.0028122 | 0.0084387 | 0.003152 | 0.469346 | 0.416222 | 0.47918 |
| 20 | 20_71440253_A_T | 71440253 | T | A | LOC617224 | intronic | 8.7445E-10 | 0.31577 | 2.774E-12 | 0.017155 | 0.0083676 | 0.021872 | 0.0027981 | 0.0083411 | 0.0031297 | 0.471901 | 0.482644 | 0.463487 |
| 20 | 20_71440264_G_T | 71440264 | T | G | LOC617224 | intronic | 2.0522E-08 | 0.42512 | 2.853E-11 | 0.015742 | 0.0067582 | 0.020878 | 0.0028073 | 0.0084735 | 0.0031376 | 0.483109 | 0.429907 | 0.492957 |
| 20 | 20_71440322_C_T | 71440322 | T | C | LOC617224 | intronic | 5.5091E-10 | 0.32075 | 3.503E-12 | 0.017405 | 0.0084628 | 0.021769 | 0.0028055 | 0.0085232 | 0.0031296 | 0.468773 | 0.493324 | 0.461757 |
| 20 | 20_71440714_T_C | 71440714 | C | T | LOC617224 | intronic | 2.5668E-09 | 0.40041 | 5.216E-12 | 0.016638 | 0.0072171 | 0.021476 | 0.0027929 | 0.0085826 | 0.0031127 | 0.485194 | 0.470294 | 0.476955 |
| 20 | 20_71440949_T_C | 71440949 | C | T | LOC617224 | intronic | 1.4351E-09 | 0.40577 | 1.169E-11 | 0.016838 | 0.0070277 | 0.021076 | 0.0027825 | 0.0084533 | 0.0031067 | 0.48582 | 0.468625 | 0.477388 |
| 20 | 20_71440992_C_T | 71440992 | T | C | LOC617224 | intronic | 1.3579E-09 | 0.3608 | 1.729E-11 | 0.016782 | 0.0077196 | 0.020791 | 0.0027692 | 0.0084475 | 0.0030905 | 0.486758 | 0.465955 | 0.478006 |
| 20 | 20_71441035_C_T | 71441035 | T | C | LOC617224 | intronic | 1.6342E-09 | 0.22209 | 7.854E-12 | 0.01666 | 0.010241 | 0.021071 | 0.0027627 | 0.0083877 | 0.00308 | 0.445157 | 0.479306 | 0.438836 |
| 20 | 20_71441042_G_C | 71441042 | C | G | LOC617224 | intronic | 1.8088E-09 | 0.25336 | 6.959E-12 | 0.016526 | 0.0095531 | 0.020998 | 0.0027479 | 0.0083636 | 0.0030616 | 0.447607 | 0.481308 | 0.441369 |
| 20 | 20_71441080_G_A | 71441080 | A | G | LOC617224 | intronic | 2.4046E-08 | 0.45374 | 5.337E-11 | 0.015413 | 0.0064192 | 0.020177 | 0.0027621 | 0.0085681 | 0.0030752 | 0.460484 | 0.414553 | 0.468986 |
| 20 | 20_71441084_A_G | 71441084 | G | A | LOC617224 | intronic | 9.701E-10 | 0.48107 | 2.418E-12 | 0.016683 | 0.005939 | 0.021282 | 0.0027285 | 0.0084291 | 0.0030368 | 0.495047 | 0.464619 | 0.487582 |
| 20 | 20_71441102_C_T | 71441102 | T | C | LOC617224 | intronic | 1.7793E-10 | 0.37238 | 5.63E-13 | 0.017415 | 0.00749 | 0.021898 | 0.0027299 | 0.0083966 | 0.0030376 | 0.4731 | 0.488318 | 0.465958 |
| 20 | 20_71441112_C_T | 71441112 | T | C | LOC617224 | intronic | 1.5707E-10 | 0.41498 | 3.965E-13 | 0.017479 | 0.0068539 | 0.022055 | 0.0027319 | 0.0084081 | 0.0030392 | 0.473621 | 0.488318 | 0.466576 |
| 20 | 20_71441115_A_G | 71441115 | G | A | LOC617224 | intronic | 1.1685E-08 | 0.49304 | 1.882E-11 | 0.01575 | 0.0058551 | 0.020648 | 0.0027611 | 0.0085415 | 0.003075 | 0.458503 | 0.412216 | 0.46707 |
| 20 | 20_71441122_G_A | 71441122 | A | G | LOC617224 | intronic | 1.8203E-10 | 0.39243 | 5.834E-13 | 0.017422 | 0.0071704 | 0.021922 | 0.0027326 | 0.0083843 | 0.0030429 | 0.472787 | 0.49032 | 0.465958 |
| 20 | 20_71441160_G_A | 71441160 | A | G | LOC617224 | intronic | 1.6461E-10 | 0.22732 | 1.693E-12 | 0.017681 | 0.010147 | 0.021779 | 0.0027665 | 0.0084048 | 0.0030858 | 0.469242 | 0.489653 | 0.461634 |
| 20 | 20_71441370_C_T | 71441370 | T | C | LOC617224 | intronic | 1.2834E-09 | 0.6057 | 1.205E-12 | 0.017094 | 0.0044034 | 0.022372 | 0.0028164 | 0.00853 | 0.0031488 | 0.465436 | 0.414219 | 0.474917 |
| 20 | 20_71442139_C_T | 71442139 | T | C | LOC617224 | intronic | 6.1549E-10 | 0.29361 | 2.634E-12 | 0.017333 | 0.008914 | 0.021875 | 0.0028018 | 0.0084875 | 0.0031268 | 0.465176 | 0.495327 | 0.457865 |
| 20 | 20_71442260_T_C | 71442260 | C | T | LOC617224 | intronic | 9.5438E-11 | 0.11185 | 1.686E-12 | 0.018299 | 0.013206 | 0.022383 | 0.0028265 | 0.0083063 | 0.0031713 | 0.428944 | 0.467957 | 0.421722 |
| 20 | 20_71442528_T_C | 71442528 | C | T | LOC617224 | intronic | 1.7623E-09 | 0.48586 | 5.134E-12 | 0.016879 | 0.0058852 | 0.021652 | 0.0028046 | 0.0084448 | 0.0031371 | 0.489365 | 0.470294 | 0.481898 |
| 20 | 20_71442538_C_G | 71442538 | G | C | LOC617224 | intronic | 1.1293E-09 | 0.36624 | 3.908E-12 | 0.017095 | 0.0076339 | 0.021772 | 0.0028071 | 0.0084489 | 0.003137 | 0.468147 | 0.494326 | 0.461201 |
| 20 | 20_71442561_G_C | 71442561 | C | G | LOC617224 | intronic | 1.407E-08 | 0.53789 | 2.616E-11 | 0.016056 | 0.0052782 | 0.021113 | 0.0028305 | 0.0085685 | 0.003167 | 0.465697 | 0.417557 | 0.474608 |
| 20 | 20_71442587_T_C | 71442587 | C | T | LOC617224 | intronic | 5.1679E-09 | 0.47719 | 1.184E-11 | 0.016307 | 0.0059137 | 0.021221 | 0.0027915 | 0.0083195 | 0.0031289 | 0.489417 | 0.466956 | 0.481342 |
| 20 | 20_71442692_G_A | 71442692 | A | G | LOC617224 | intronic | 3.494E-08 | 0.51403 | 8.289E-11 | 0.015654 | 0.0056121 | 0.020623 | 0.0028386 | 0.0085999 | 0.0031751 | 0.465541 | 0.417223 | 0.474484 |
| 20 | 20_71444098_T_C | 71444098 | C | T | LOC617224 | intronic | 1.0403E-09 | 0.32135 | 4.419E-12 | 0.017175 | 0.0083294 | 0.0218 | 0.0028141 | 0.0083992 | 0.0031489 | 0.46627 | 0.491989 | 0.458544 |
| 20 | 20_71446401_A_C | 71446401 | C | A | LOC617224 | intronic | 1.43E-10 | 0.28759 | 1.367E-12 | 0.01811 | 0.0087543 | 0.022519 | 0.0028242 | 0.0082322 | 0.0031773 | 0.434626 | 0.46996 | 0.428086 |
| 20 | 20_71446504_G_A | 71446504 | A | G | LOC617224 | intronic | 1.0003E-10 | 0.24572 | 6.363E-13 | 0.018195 | 0.009725 | 0.022664 | 0.0028136 | 0.0083777 | 0.003151 | 0.468877 | 0.489319 | 0.461139 |
| 20 | 20_71446746_A_G | 71446746 | G | A | LOC617224 | intronic | 6.9424E-09 | 0.28544 | 2.768E-11 | 0.01639 | 0.0089828 | 0.02115 | 0.0028297 | 0.0084094 | 0.0031765 | 0.464029 | 0.402203 | 0.475473 |
| 20 | 20_71446765_C_T | 71446765 | T | C | LOC617224 | intronic | 2.55E-10 | 0.24236 | 1.21E-12 | 0.017696 | 0.0096241 | 0.022308 | 0.0027982 | 0.0082319 | 0.0031401 | 0.470962 | 0.47964 | 0.461819 |
| 20 | 20_71446775_C_T | 71446775 | T | C | LOC617224 | intronic | 1.8195E-09 | 0.22943 | 6.622E-12 | 0.01701 | 0.010074 | 0.021818 | 0.0028288 | 0.0083821 | 0.0031779 | 0.463038 | 0.40287 | 0.474175 |
| 20 | 20_71447274_T_C | 71447274 | C | T | LOC617224 | intronic | 3.3786E-10 | 0.53448 | 3.078E-12 | 0.017744 | 0.0050604 | 0.022216 | 0.0028254 | 0.0081463 | 0.0031856 | 0.434313 | 0.470294 | 0.427654 |
| 20 | 20_71447597_G_C | 71447597 | C | G | LOC617224 | intronic | 4.0273E-11 | 0.44844 | 1.886E-13 | 0.018461 | 0.006226 | 0.023096 | 0.0027959 | 0.0082136 | 0.0031394 | 0.470754 | 0.483979 | 0.462375 |
| 20 | 20_71447601_A_G | 71447601 | G | A | LOC617224 | intronic | 3.5415E-12 | 0.3025 | 4.012E-14 | 0.019598 | 0.0083342 | 0.024032 | 0.0028181 | 0.008083 | 0.0031786 | 0.435356 | 0.477637 | 0.42753 |
| 20 | 20_71447948_T_G | 71447948 | G | T | LOC617224 | intronic | 2.3314E-10 | 0.46923 | 5.882E-13 | 0.017809 | 0.0060391 | 0.022682 | 0.0028101 | 0.0083444 | 0.0031488 | 0.464654 | 0.49466 | 0.457123 |
| 20 | 20_71448095_G_T | 71448095 | T | G | LOC617224 | intronic | 7.349E-09 | 0.23079 | 6.767E-11 | 0.016262 | 0.0097911 | 0.020675 | 0.0028122 | 0.0081707 | 0.0031682 | 0.479773 | 0.4753 | 0.471457 |
| 20 | 20_71448150_G_A | 71448150 | A | G | LOC617224 | intronic | 6.5312E-10 | 0.35179 | 2.164E-12 | 0.017376 | 0.0076536 | 0.022211 | 0.002813 | 0.0082197 | 0.0031623 | 0.445365 | 0.48231 | 0.438527 |

| | | | | | | | | | | | | | | | | | | |
|---|---|---|---|---|---|---|---|---|---|---|---|---|---|---|---|---|---|---|
| 20 | 20_71448225_G_A | 71448225 | A | G | LOC617224 | intronic | 2.3917E-11 | 0.30354 | 1.191E-13 | 0.018637 | 0.0085028 | 0.023209 | 0.0027901 | 0.0082641 | 0.0031288 | 0.48363 | 0.475968 | 0.476152 |
| 20 | 20_71448297_C_T | 71448297 | T | C | LOC617224 | intronic | 5.0546E-11 | 0.23248 | 3.998E-13 | 0.018553 | 0.0098368 | 0.023032 | 0.0028242 | 0.0082385 | 0.0031743 | 0.435617 | 0.468959 | 0.429445 |
| 20 | 20_71448849_T_C | 71448849 | C | T | LOC617224 | intronic | 3.2754E-12 | 0.10799 | 4.522E-14 | 0.019752 | 0.013308 | 0.024031 | 0.0028358 | 0.0082795 | 0.003185 | 0.433323 | 0.473632 | 0.425862 |
| 20 | 20_71449014_G_A | 71449014 | A | G | LOC617224 | intronic | 2.0312E-11 | 0.18422 | 2.119E-13 | 0.018821 | 0.011092 | 0.023075 | 0.0028075 | 0.0083535 | 0.0031433 | 0.470076 | 0.484312 | 0.461634 |
| 20 | 20_71449177_G_A | 71449177 | A | G | LOC617224 | intronic | 2.7055E-12 | 0.10734 | 4.503E-14 | 0.019793 | 0.013315 | 0.023995 | 0.0028307 | 0.0082689 | 0.00318 | 0.434783 | 0.473632 | 0.427592 |
| 20 | 20_71449819_T_C | 71449819 | C | T | LOC617224 | intronic | 3.8051E-11 | 0.26992 | 2.197E-13 | 0.018746 | 0.0091341 | 0.023373 | 0.0028353 | 0.0082793 | 0.003186 | 0.431967 | 0.46729 | 0.425429 |
| 20 | 20_71450177_A_G | 71450177 | G | A | LOC617224 | intronic | 5.5769E-11 | 0.17747 | 1.242E-12 | 0.018611 | 0.011223 | 0.022621 | 0.0028394 | 0.0083224 | 0.0031858 | 0.432124 | 0.468959 | 0.425306 |
| 20 | 20_71450498_T_C | 71450498 | C | T | LOC617224 | intronic | 1.1828E-09 | 0.16319 | 8.483E-12 | 0.01719 | 0.011343 | 0.02175 | 0.0028261 | 0.0081343 | 0.0031844 | 0.433427 | 0.474967 | 0.425738 |
| 20 | 20_71450861_A_G | 71450861 | G | A | LOC617224 | intronic | 1.1207E-09 | 0.16418 | 3.424E-11 | 0.017094 | 0.011325 | 0.020956 | 0.0028063 | 0.0081406 | 0.0031622 | 0.488739 | 0.477637 | 0.482516 |
| 20 | 20_71451005_C_T | 71451005 | T | C | LOC617224 | intronic | 1.6219E-08 | 0.32224 | 6.628E-10 | 0.015755 | 0.0081917 | 0.019317 | 0.0027894 | 0.0082757 | 0.0031284 | 0.475863 | 0.43725 | 0.48301 |
| 20 | 20_71451218_C_T | 71451218 | T | C | LOC617224 | intronic | 6.0357E-09 | 0.26819 | 2.574E-10 | 0.016216 | 0.0093371 | 0.019741 | 0.0027883 | 0.0084327 | 0.0031224 | 0.465384 | 0.432243 | 0.471519 |
| 20 | 20_71451220_A_G | 71451220 | G | A | LOC617224 | intronic | 9.0887E-09 | 0.20477 | 6.646E-10 | 0.01607 | 0.010487 | 0.019421 | 0.0027963 | 0.0082704 | 0.0031454 | 0.49927 | 0.471295 | 0.495552 |
| 20 | 20_71452023_C_T | 71452023 | T | C | LOC617224 | exonic | 1.4992E-08 | 0.32945 | 7.237E-10 | 0.015738 | 0.0080605 | 0.019216 | 0.0027797 | 0.0082653 | 0.003119 | 0.476436 | 0.43992 | 0.483195 |
| 20 | 20_71452636_C_T | 71452636 | T | C | LOC617224 | intronic | 2.7891E-11 | 0.38716 | 1.381E-13 | 0.018709 | 0.0071694 | 0.0233 | 0.0028103 | 0.0082905 | 0.0031495 | 0.466583 | 0.493324 | 0.459162 |
| 20 | 20_71454211_G_A | 71454211 | A | G | LOC617224 | intronic | 1.0615E-10 | 0.34484 | 4.252E-13 | 0.018187 | 0.0079247 | 0.02287 | 0.0028162 | 0.0083892 | 0.0031557 | 0.470754 | 0.490654 | 0.463611 |
| 20 | 20_71454235_T_C | 71454235 | C | T | LOC617224 | intronic | 1.1577E-10 | 0.36623 | 4.987E-13 | 0.018159 | 0.007591 | 0.022808 | 0.0028176 | 0.0084013 | 0.0031565 | 0.470806 | 0.489319 | 0.463425 |
| 20 | 20_71454387_T_C | 71454387 | C | T | LOC617224 | intronic | 5.486E-11 | 0.22987 | 7.187E-13 | 0.018623 | 0.0099482 | 0.022904 | 0.0028402 | 0.0082853 | 0.0031918 | 0.434678 | 0.469292 | 0.428271 |
| 20 | 20_71454622_T_C | 71454622 | C | T | LOC617224 | intronic | 3.0029E-11 | 0.3052 | 1.648E-13 | 0.018759 | 0.0085651 | 0.023339 | 0.0028224 | 0.0083534 | 0.0031647 | 0.464863 | 0.495995 | 0.457618 |
| 20 | 20_71456273_A_G | 71456273 | G | A | LOC617224 | intronic | 9.5194E-10 | 0.38404 | 6.625E-12 | 0.017299 | 0.0071426 | 0.021853 | 0.0028279 | 0.0082055 | 0.003183 | 0.431081 | 0.463284 | 0.42512 |
| 20 | 20_71456292_G_T | 71456292 | T | G | LOC617224 | intronic | 6.7758E-10 | 0.4461 | 2.106E-12 | 0.017346 | 0.0063535 | 0.022138 | 0.0028109 | 0.0083387 | 0.0031504 | 0.459598 | 0.494993 | 0.453046 |
| 20 | 20_71456341_A_G | 71456341 | G | A | LOC617224 | intronic | 1.4889E-10 | 0.28858 | 1.226E-12 | 0.018166 | 0.0087431 | 0.02265 | 0.0028356 | 0.0082386 | 0.003189 | 0.427223 | 0.464286 | 0.420363 |
| 20 | 20_71456502_T_C | 71456502 | C | T | LOC617224 | intronic | 2.9461E-10 | 0.32533 | 2.59E-12 | 0.017784 | 0.0082817 | 0.022121 | 0.0028221 | 0.0084201 | 0.0031608 | 0.461214 | 0.495327 | 0.454899 |
| 20 | 20_71458942_C_T | 71458942 | T | C | LOC617224 | intronic | 7.9333E-10 | 0.48869 | 3.459E-12 | 0.017276 | 0.0057011 | 0.021952 | 0.0028108 | 0.008234 | 0.0031551 | 0.462517 | 0.491656 | 0.454034 |
| 20 | 20_71459002_A_G | 71459002 | G | A | LOC617224 | intronic | 1.2E-10 | 0.41732 | 3.675E-13 | 0.018295 | 0.0066688 | 0.023235 | 0.0028412 | 0.0082221 | 0.0031973 | 0.425712 | 0.462951 | 0.418819 |
| 20 | 20_71459131_T_C | 71459131 | C | T | LOC617224 | intronic | 2.0838E-10 | 0.30277 | 1.584E-12 | 0.01808 | 0.008509 | 0.022621 | 0.002845 | 0.0082572 | 0.003201 | 0.425972 | 0.460948 | 0.419498 |
| 20 | 20_71459607_G_T | 71459607 | T | G | LOC617224 | intronic | 3.6452E-11 | 0.24893 | 2.439E-13 | 0.018642 | 0.0095638 | 0.023123 | 0.0028169 | 0.008295 | 0.0031579 | 0.46137 | 0.491656 | 0.452675 |
| 20 | 20_71459819_G_A | 71459819 | A | G | LOC617224 | intronic | 7.0504E-11 | 0.27244 | 8.551E-13 | 0.018391 | 0.0092088 | 0.022586 | 0.0028209 | 0.0083909 | 0.003158 | 0.461214 | 0.498999 | 0.45422 |
| 20 | 20_71461744_A_G | 71461744 | G | A | LOC617224(XM_024981564.1:c.*244/ | UTR3 | 1.5627E-10 | 0.36123 | 1.053E-12 | 0.018142 | 0.0074134 | 0.022769 | 0.0028351 | 0.0081196 | 0.0031963 | 0.427745 | 0.465287 | 0.420796 |
| 20 | 20_71462296_G_T | 71462296 | T | G | LOC617224(dist=362) | downstream | 2.8609E-11 | 0.5221 | 1.095E-13 | 0.018907 | 0.0052531 | 0.023769 | 0.0028416 | 0.0082065 | 0.0031996 | 0.42811 | 0.465621 | 0.421166 |
| 20 | 20_71462849_G_A | 71462849 | A | G | LOC617224(dist=915) | downstream | 1.8018E-09 | 0.52409 | 6.302E-12 | 0.017003 | 0.0053192 | 0.021785 | 0.0028269 | 0.0083496 | 0.0031699 | 0.460849 | 0.496996 | 0.453046 |
| 20 | 20_71463948_C_T | 71463948 | T | C | LOC104975299(XM_024981567.1:c.*8 | UTR3 | 1.1051E-08 | 0.8323 | 4.02E-11 | 0.016308 | 0.0018141 | 0.021055 | 0.0028542 | 0.0085671 | 0.0031886 | 0.475081 | 0.480307 | 0.466823 |
| 20 | 20_71464086_A_G | 71464086 | G | A | LOC104975299(XM_024981567.1:c.*7 | UTR3 | 1.2661E-10 | 0.32982 | 7.268E-13 | 0.018348 | 0.0081011 | 0.022992 | 0.002853 | 0.0083134 | 0.0032047 | 0.42837 | 0.464286 | 0.421722 |
| 20 | 20_71465001_C_G | 71465001 | G | C | LOC104975299 | exonic | 6.0855E-11 | 0.37844 | 4.038E-13 | 0.018659 | 0.0073432 | 0.023236 | 0.0028524 | 0.0083373 | 0.0032031 | 0.428266 | 0.461282 | 0.422155 |
| 20 | 20_71468477_C_A | 71468477 | A | C | TPPP | intronic | 2.496E-10 | 0.56093 | 1.525E-12 | 0.017745 | 0.0047579 | 0.022295 | 0.0028046 | 0.0081827 | 0.0031525 | 0.463768 | 0.491656 | 0.455517 |
| 20 | 20_71468859_T_G | 71468859 | G | T | TPPP | intronic | 2.2616E-10 | 0.71786 | 1.1E-12 | 0.017884 | 0.0030297 | 0.022455 | 0.0028197 | 0.0083852 | 0.0031549 | 0.464759 | 0.491656 | 0.456691 |
| 20 | 20_71468882_C_T | 71468882 | T | C | TPPP | intronic | 2.0402E-10 | 0.72557 | 8.517E-13 | 0.017932 | 0.0029453 | 0.022567 | 0.0028202 | 0.0083906 | 0.003155 | 0.465176 | 0.491989 | 0.457247 |
| 20 | 20_71472067_C_G | 71472067 | G | C | TPPP | intronic | 2.4478E-10 | 0.56421 | 1.858E-12 | 0.017824 | 0.0048185 | 0.022197 | 0.0028157 | 0.0083567 | 0.0031508 | 0.463299 | 0.492657 | 0.455146 |
| 20 | 20_71472205_C_T | 71472205 | T | C | TPPP | intronic | 1.3571E-10 | 0.60264 | 7.635E-13 | 0.018271 | 0.0043147 | 0.022919 | 0.0028457 | 0.0082878 | 0.0031975 | 0.42665 | 0.464619 | 0.419622 |
| 20 | 20_71472317_A_G | 71472317 | G | A | TPPP | intronic | 1.9148E-10 | 0.62836 | 5.257E-13 | 0.018042 | 0.0039697 | 0.023034 | 0.0028332 | 0.0082013 | 0.0031909 | 0.425555 | 0.461282 | 0.418942 |
| 20 | 20_71473111_G_C | 71473111 | C | G | TPPP | intronic | 2.3795E-10 | 0.46154 | 3.27E-12 | 0.017864 | 0.0061204 | 0.022033 | 0.0028201 | 0.0083122 | 0.0031631 | 0.464394 | 0.493992 | 0.456691 |
| 20 | 20_71475258_C_T | 71475258 | T | C | TPPP | intronic | 3.1735E-10 | 0.58167 | 2.911E-12 | 0.017773 | 0.0046398 | 0.022081 | 0.0028256 | 0.0084216 | 0.0031626 | 0.464185 | 0.496662 | 0.456938 |
| 20 | 20_71476118_A_G | 71476118 | G | A | TPPP | intronic | 6.2709E-11 | 0.41251 | 2.671E-13 | 0.018639 | 0.0068524 | 0.023418 | 0.0028512 | 0.0083618 | 0.0032035 | 0.425607 | 0.458278 | 0.41956 |
| 20 | 20_71476760_T_C | 71476760 | C | T | TPPP | intronic | 1.7674E-10 | 0.46594 | 9.76E-13 | 0.018229 | 0.0061279 | 0.022866 | 0.0028571 | 0.0084048 | 0.0032053 | 0.423261 | 0.45494 | 0.417398 |
| 20 | 20_71476853_G_C | 71476853 | C | G | TPPP | intronic | 1.6944E-10 | 0.42397 | 4.166E-13 | 0.018221 | 0.0067099 | 0.023216 | 0.002853 | 0.0083922 | 0.0032022 | 0.423157 | 0.457276 | 0.416842 |
| 20 | 20_71479838_G_A | 71479838 | A | G | TPPP | intronic | 6.5284E-10 | 0.64918 | 3.755E-12 | 0.017382 | 0.0037698 | 0.021929 | 0.002814 | 0.0082872 | 0.003157 | 0.465749 | 0.492323 | 0.457988 |
| 20 | 20_71480183_C_T | 71480183 | T | C | TPPP | exonic | 4.8494E-11 | 0.60998 | 4.062E-13 | 0.018525 | 0.0041877 | 0.022981 | 0.0028173 | 0.0082096 | 0.0031683 | 0.468199 | 0.489987 | 0.46046 |
| 20 | 20_71480448_C_G | 71480448 | G | C | TPPP | intronic | 3.8273E-11 | 0.28395 | 2.045E-13 | 0.018772 | 0.0087833 | 0.023501 | 0.0028396 | 0.0081972 | 0.0031993 | 0.426963 | 0.466622 | 0.419622 |
| 20 | 20_71480955_G_A | 71480955 | A | G | TPPP | intronic | 5.0691E-11 | 0.3179 | 4.648E-13 | 0.018741 | 0.0084061 | 0.023168 | 0.0028529 | 0.0084163 | 0.0032021 | 0.424565 | 0.452603 | 0.419375 |
| 20 | 20_71481725_A_G | 71481725 | G | A | TPPP | intronic | 4.6358E-11 | 0.29136 | 3.043E-13 | 0.018772 | 0.0088093 | 0.023374 | 0.0028519 | 0.0083488 | 0.0032053 | 0.425868 | 0.456609 | 0.420178 |
| 20 | 20_71487201_C_T | 71487201 | T | C | TPPP(XM_024981197.1:c.*1858C>T,X | UTR3 | 5.2903E-10 | 0.54411 | 9.085E-12 | 0.017479 | 0.004993 | 0.021549 | 0.0028146 | 0.008231 | 0.0031596 | 0.467209 | 0.488652 | 0.459039 |
| 20 | 20_71488622_T_C | 71488622 | C | T | TPPP(dist=129) | downstream | 1.9685E-09 | 0.58319 | 3.627E-12 | 0.017025 | 0.0044753 | 0.022246 | 0.0028373 | 0.0081557 | 0.0032003 | 0.426077 | 0.456943 | 0.420363 |
| 20 | 20_71488667_A_G | 71488667 | A | G | TPPP(dist=174) | downstream | 1.9364E-09 | 0.69252 | 7.378E-12 | 0.016804 | 0.0031959 | 0.021613 | 0.0027992 | 0.0080817 | 0.0031552 | 0.46674 | 0.493658 | 0.459409 |
| 20 | 20_71488952_G_A | 71488952 | A | G | TPPP(dist=459) | downstream | 4.2037E-10 | 0.40981 | 5.222E-12 | 0.01755 | 0.0067763 | 0.021791 | 0.0028097 | 0.0082215 | 0.0031584 | 0.46455 | 0.495661 | 0.457185 |
| 20 | 20_71490172_A_G | 71490172 | G | A | TPPP(dist=1679),CEP72(dist=1706) | intergenic | 4.0256E-10 | 0.42804 | 2.328E-12 | 0.01775 | 0.0064791 | 0.022432 | 0.0028386 | 0.008175 | 0.0031986 | 0.427223 | 0.463284 | 0.420549 |
| 20 | 20_71491300_A_G | 71491300 | G | A | CEP72(dist=578) | downstream | 3.3731E-11 | 0.43702 | 4.66E-13 | 0.018888 | 0.0064697 | 0.023155 | 0.0028491 | 0.0083241 | 0.0032004 | 0.427588 | 0.46028 | 0.421537 |
| 20 | 20_71491821_C_T | 71491821 | T | C | CEP72(dist=57) | downstream | 5.7408E-10 | 0.52575 | 3.436E-12 | 0.017467 | 0.0053019 | 0.02197 | 0.0028184 | 0.0083559 | 0.0031572 | 0.463977 | 0.493992 | 0.456197 |
| 20 | 20_71495712_G_A | 71495712 | A | G | CEP72 | intronic | 4.5789E-11 | 0.11961 | 4.038E-12 | 0.018392 | 0.012772 | 0.02176 | 0.0027935 | 0.0082061 | 0.0031373 | 0.475915 | 0.477303 | 0.467256 |
| 20 | 20_71496677_C_A | 71496677 | A | C | CEP72 | intronic | 2.1414E-11 | 0.19466 | 1.995E-13 | 0.018968 | 0.010572 | 0.023436 | 0.0028327 | 0.0081517 | 0.003189 | 0.43056 | 0.473965 | 0.422526 |
| 20 | 20_71498958_C_T | 71498958 | T | C | CEP72 | intronic | 1.5353E-10 | 0.50763 | 9.083E-13 | 0.01798 | 0.0055246 | 0.022479 | 0.0028086 | 0.0083388 | 0.0031467 | 0.464029 | 0.496328 | 0.456691 |
| 20 | 20_71499795_C_T | 71499795 | T | C | CEP72 | intronic | 2.6868E-10 | 0.33177 | 3.781E-12 | 0.017742 | 0.0080968 | 0.021855 | 0.0028092 | 0.0083425 | 0.0031469 | 0.462465 | 0.497997 | 0.455146 |
| 20 | 20_71500759_T_C | 71500759 | C | T | CEP72 | intronic | 3.3385E-11 | 0.47137 | 9.526E-13 | 0.018665 | 0.0060332 | 0.022499 | 0.0028149 | 0.0083767 | 0.0031524 | 0.465541 | 0.496662 | 0.458544 |
| 20 | 20_71500764_C_T | 71500764 | T | C | CEP72 | intronic | 3.0936E-11 | 0.44567 | 9.727E-13 | 0.018696 | 0.0063762 | 0.022494 | 0.0028147 | 0.0083605 | 0.0031529 | 0.465801 | 0.494993 | 0.458544 |
| 20 | 20_71500779_T_C | 71500779 | C | T | CEP72 | intronic | 2.5152E-11 | 0.33834 | 5.402E-13 | 0.018953 | 0.0079504 | 0.023046 | 0.0028405 | 0.0083036 | 0.0031943 | 0.426233 | 0.455607 | 0.420796 |
| 20 | 20_71501010_G_C | 71501010 | C | G | CEP72 | intronic | 1.7578E-10 | 0.36119 | 6.729E-13 | 0.018134 | 0.0075733 | 0.022945 | 0.0028418 | 0.0082941 | 0.0031935 | 0.424617 | 0.461282 | 0.41783 |
| 20 | 20_71501566_G_A | 71501566 | A | G | CEP72 | intronic | 1.1417E-10 | 0.79787 | 6.008E-13 | 0.018004 | 0.0021095 | 0.022527 | 0.0027927 | 0.0082368 | 0.0031286 | 0.462517 | 0.488985 | 0.45354 |
| 20 | 20_71501596_A_G | 71501596 | G | A | CEP72 | intronic | 7.6972E-10 | 0.60011 | 2.669E-12 | 0.017346 | 0.0043034 | 0.02217 | 0.00282 | 0.0082088 | 0.0031698 | 0.422219 | 0.46028 | 0.415174 |
| 20 | 20_71501662_C_A | 71501662 | A | C | CEP72 | intronic | 1.4309E-09 | 0.6813 | 2.837E-12 | 0.016888 | 0.0033969 | 0.021804 | 0.0027905 | 0.0082713 | 0.0031212 | 0.462882 | 0.48498 | 0.453231 |
| 20 | 20_71501663_C_G | 71501663 | G | C | CEP72 | intronic | 1.4309E-09 | 0.6813 | 2.837E-12 | 0.016888 | 0.0033969 | 0.021804 | 0.0027905 | 0.0082713 | 0.0031212 | 0.462882 | 0.48498 | 0.453231 |
| 20 | 20_71501789_G_A | 71501789 | A | G | CEP72 | intronic | 6.0584E-10 | 0.73876 | 2.902E-12 | 0.017299 | 0.0027401 | 0.021865 | 0.0027951 | 0.0082162 | 0.0031314 | 0.465384 | 0.482644 | 0.455764 |
| 20 | 20_71501848_C_T | 71501848 | T | C | CEP72 | intronic | 4.6521E-10 | 0.54895 | 6.391E-12 | 0.017373 | 0.0049567 | 0.021448 | 0.0027885 | 0.0082704 | 0.0031217 | 0.461787 | 0.493324 | 0.453478 |
| 20 | 20_71501971_T_C | 71501971 | C | T | CEP72 | intronic | 3.5424E-10 | 0.47976 | 9.523E-13 | 0.017764 | 0.0058431 | 0.022712 | 0.0028318 | 0.0082682 | 0.0031822 | 0.425659 | 0.459279 | 0.419437 |
| 20 | 20_71502108_T_C | 71502108 | C | T | CEP72 | intronic | 1.2718E-09 | 0.84166 | 1.446E-12 | 0.016983 | 0.0016385 | 0.022191 | 0.0027975 | 0.008202 | 0.0031346 | 0.465384 | 0.481642 | 0.455579 |
| 20 | 20_71502319_C_T | 71502319 | T | C | CEP72 | intronic | 1.9234E-10 | 0.79166 | 4.1E-13 | 0.017851 | 0.0021885 | 0.022753 | 0.0028035 | 0.0082849 | 0.0031374 | 0.465176 | 0.487984 | 0.456506 |
| 20 | 20_71502380_C_T | 71502380 | T | C | CEP72 | intronic | 2.4576E-10 | 0.81787 | 6.92E-13 | 0.017739 | 0.0019095 | 0.022514 | 0.0028026 | 0.0082917 | 0.0031352 | 0.464915 | 0.486315 | 0.455888 |
| 20 | 20_71502884_C_A | 71502884 | A | C | CEP72 | intronic | 1.4026E-10 | 0.46963 | 5.717E-13 | 0.018164 | 0.005938 | 0.022951 | 0.0028312 | 0.0082121 | 0.0031845 | 0.423991 | 0.462951 | 0.41678 |

| | | | | | | | | | | | | | | | | | | |
|---|---|---|---|---|---|---|---|---|---|---|---|---|---|---|---|---|---|---|
| 20 | 20_71503018_A_G | 71503018 | G | A | CEP72 | intronic | 4.1282E-10 | 0.58319 | 1.398E-12 | 0.017705 | 0.0045402 | 0.022555 | 0.0028332 | 0.0082741 | 0.0031839 | 0.424304 | 0.457276 | 0.418201 |
| 20 | 20_71503116_G_A | 71503116 | A | G | CEP72 | intronic | 1.9501E-10 | 0.69043 | 1.598E-12 | 0.017801 | 0.0032724 | 0.022156 | 0.0027966 | 0.0082167 | 0.0031357 | 0.466114 | 0.48765 | 0.457556 |
| 20 | 20_71503202_A_G | 71503202 | G | A | CEP72 | intronic | 4.4755E-10 | 0.74117 | 1.034E-12 | 0.0174 | 0.0026348 | 0.022396 | 0.0027901 | 0.0079769 | 0.003143 | 0.471588 | 0.466956 | 0.460213 |
| 20 | 20_71503463_G_A | 71503463 | A | G | CEP72 | intronic | 3.676E-10 | 0.53105 | 2.98E-12 | 0.017519 | 0.0052144 | 0.021832 | 0.0027954 | 0.0083244 | 0.0031285 | 0.473413 | 0.479973 | 0.464784 |
| 20 | 20_71503500_A_T | 71503500 | T | A | CEP72 | intronic | 2.5153E-10 | 0.46005 | 1.714E-12 | 0.017567 | 0.0059951 | 0.022021 | 0.0027769 | 0.0081151 | 0.0031209 | 0.473725 | 0.476969 | 0.464599 |
| 20 | 20_71503634_G_A | 71503634 | A | G | CEP72 | intronic | 2.8413E-09 | 0.69825 | 6.976E-12 | 0.016592 | 0.0032366 | 0.021446 | 0.0027929 | 0.0083486 | 0.0031271 | 0.474768 | 0.482977 | 0.466947 |
| 20 | 20_71503865_T_C | 71503865 | C | T | CEP72 | intronic | 1.4626E-08 | 0.51169 | 2.8E-10 | 0.015924 | 0.0054521 | 0.01992 | 0.0028106 | 0.0083086 | 0.0031571 | 0.43473 | 0.47964 | 0.426418 |
| 20 | 20_71503866_A_C | 71503866 | C | A | CEP72 | intronic | 4.8157E-09 | 0.33448 | 8.04E-11 | 0.016638 | 0.0079454 | 0.020838 | 0.0028425 | 0.0082324 | 0.0032059 | 0.398238 | 0.438919 | 0.390708 |
| 20 | 20_71504239_A_T | 71504239 | T | A | CEP72 | intronic | 9.74E-11 | 0.52092 | 1.92E-12 | 0.018197 | 0.0054203 | 0.022134 | 0.0028121 | 0.0084437 | 0.003144 | 0.461943 | 0.496662 | 0.455517 |
| 20 | 20_71504265_G_A | 71504265 | A | G | CEP72 | intronic | 6.5094E-11 | 0.35033 | 7.908E-13 | 0.018515 | 0.0077713 | 0.022788 | 0.0028347 | 0.0083208 | 0.0031815 | 0.425555 | 0.455274 | 0.420054 |
| 20 | 20_71504637_T_C | 71504637 | C | T | CEP72 | intronic | 9.5263E-10 | 0.6662 | 4.862E-12 | 0.017176 | 0.003614 | 0.021708 | 0.0028079 | 0.008378 | 0.0031418 | 0.461891 | 0.495327 | 0.453973 |
| 20 | 20_71504676_A_G | 71504676 | G | A | CEP72 | intronic | 2.9732E-10 | 0.73561 | 1.913E-12 | 0.017652 | 0.0028106 | 0.022091 | 0.0028019 | 0.0083234 | 0.0031376 | 0.461266 | 0.496662 | 0.453478 |
| 20 | 20_71504755_T_C | 71504755 | C | T | CEP72 | intronic | 5.3797E-10 | 0.36365 | 6.447E-12 | 0.017399 | 0.0076136 | 0.021549 | 0.0028029 | 0.008381 | 0.003137 | 0.460692 | 0.498331 | 0.453725 |
| 20 | 20_71504766_T_C | 71504766 | C | T | CEP72 | intronic | 7.9268E-10 | 0.33858 | 1.191E-11 | 0.017233 | 0.0080044 | 0.021286 | 0.0028038 | 0.0083642 | 0.0031389 | 0.461318 | 0.5 | 0.454158 |
| 20 | 20_71504990_T_C | 71504990 | C | T | CEP72 | intronic | 4.7537E-10 | 0.61727 | 1.075E-12 | 0.01747 | 0.0041412 | 0.022386 | 0.0028055 | 0.008287 | 0.0031438 | 0.462673 | 0.492323 | 0.454343 |
| 20 | 20_71505178_C_T | 71505178 | T | C | CEP72 | intronic | 4.3061E-10 | 0.64572 | 2.961E-12 | 0.017498 | 0.0038163 | 0.021912 | 0.002803 | 0.0083015 | 0.0031394 | 0.461318 | 0.49733 | 0.453664 |
| 20 | 20_71505236_C_G | 71505236 | G | C | CEP72 | intronic | 6.3499E-10 | 0.59921 | 3.898E-12 | 0.017293 | 0.0043298 | 0.021782 | 0.0027975 | 0.0082388 | 0.0031382 | 0.461474 | 0.493992 | 0.453231 |
| 20 | 20_71505270_T_C | 71505270 | C | T | CEP72 | intronic | 1.4665E-10 | 0.40664 | 9.601E-13 | 0.018165 | 0.0068327 | 0.022746 | 0.0028344 | 0.008234 | 0.0031875 | 0.424617 | 0.462283 | 0.417645 |
| 20 | 20_71505899_T_C | 71505899 | C | T | CEP72 | intronic | 2.1876E-09 | 0.59175 | 3.099E-11 | 0.016769 | 0.0044247 | 0.020879 | 0.0028027 | 0.0082504 | 0.0031436 | 0.464289 | 0.491322 | 0.456073 |
| 20 | 20_71505966_T_C | 71505966 | C | T | CEP72 | intronic | 1.2411E-09 | 0.45146 | 1.84E-11 | 0.017035 | 0.0062405 | 0.021106 | 0.0028041 | 0.0082877 | 0.0031416 | 0.464811 | 0.489653 | 0.456382 |
| 20 | 20_71506321_A_C | 71506321 | C | A | CEP72 | intronic | 2.674E-08 | 0.55833 | 1.347E-09 | 0.014291 | 0.0045584 | 0.017413 | 0.0025696 | 0.0077878 | 0.0028727 | 0.454645 | 0.416889 | 0.461634 |
| 20 | 20_71506322_G_A | 71506322 | A | G | CEP72 | intronic | 2.6478E-08 | 0.55833 | 1.343E-09 | 0.014298 | 0.0045584 | 0.017418 | 0.00257 | 0.0077878 | 0.0028732 | 0.454854 | 0.416889 | 0.461881 |
| 20 | 20_71506323_A_T | 71506323 | T | A | CEP72 | intronic | 4.0614E-08 | 0.55824 | 2.277E-09 | 0.014097 | 0.004556 | 0.017165 | 0.0025687 | 0.007782 | 0.002872 | 0.455166 | 0.417223 | 0.46219 |
| 20 | 20_71506411_A_G | 71506411 | G | A | CEP72 | intronic | 8.2392E-10 | 0.94507 | 9.036E-12 | 0.017052 | 0.000568 | 0.021224 | 0.0027771 | 0.0082434 | 0.0031116 | 0.479304 | 0.474299 | 0.470715 |
| 20 | 20_71506498_C_A | 71506498 | A | C | CEP72 | intronic | 1.1361E-09 | 0.58637 | 1.101E-11 | 0.017045 | 0.0044186 | 0.021367 | 0.0027994 | 0.0081208 | 0.0031456 | 0.47774 | 0.473632 | 0.468738 |
| 20 | 20_71506653_A_G | 71506653 | G | A | CEP72 | intronic | 2.9299E-10 | 0.4707 | 1.757E-12 | 0.017657 | 0.0059837 | 0.02214 | 0.0028017 | 0.0082953 | 0.0031393 | 0.46137 | 0.496996 | 0.453664 |
| 20 | 20_71506688_G_A | 71506688 | A | G | CEP72 | intronic | 1.5604E-09 | 0.49379 | 1.238E-11 | 0.01695 | 0.0056986 | 0.021297 | 0.0028072 | 0.0083277 | 0.0031432 | 0.460744 | 0.496996 | 0.452922 |
| 20 | 20_71506829_C_T | 71506829 | T | C | CEP72 | intronic | 4.4371E-10 | 0.45581 | 1.964E-12 | 0.017715 | 0.0061287 | 0.022482 | 0.0028399 | 0.0082179 | 0.0031948 | 0.423939 | 0.464953 | 0.416347 |
| 20 | 20_71506857_T_C | 71506857 | C | T | CEP72 | intronic | 3.2799E-09 | 0.97052 | 8.005E-12 | 0.016599 | 0.0003057 | 0.021505 | 0.0028053 | 0.0082711 | 0.0031447 | 0.461579 | 0.493992 | 0.453355 |
| 20 | 20_71506862_A_G | 71506862 | G | A | CEP72 | intronic | 3.5117E-09 | 0.90561 | 8.499E-12 | 0.016565 | 0.0009814 | 0.021471 | 0.002805 | 0.0082765 | 0.0031437 | 0.461526 | 0.493658 | 0.453231 |
| 20 | 20_71507030_A_C | 71507030 | C | A | CEP72 | intronic | 2.369E-10 | 0.31831 | 4.247E-12 | 0.017804 | 0.0083253 | 0.021814 | 0.0028103 | 0.0083424 | 0.0031484 | 0.461943 | 0.49733 | 0.454405 |
| 20 | 20_71507124_C_T | 71507124 | T | C | CEP72 | intronic | 4.0611E-10 | 0.59853 | 3.404E-12 | 0.017659 | 0.0044547 | 0.02198 | 0.0028247 | 0.0084608 | 0.0031581 | 0.463507 | 0.498331 | 0.456444 |
| 20 | 20_71507281_G_C | 71507281 | C | G | CEP72 | intronic | 3.8227E-10 | 0.31273 | 3.697E-12 | 0.017818 | 0.0084923 | 0.02216 | 0.0028458 | 0.0084124 | 0.0031893 | 0.426077 | 0.459613 | 0.419869 |
| 20 | 20_71507459_A_G | 71507459 | G | A | CEP72 | intronic | 6.074E-11 | 0.50655 | 4.035E-13 | 0.018431 | 0.0056498 | 0.022821 | 0.0028174 | 0.008506 | 0.0031458 | 0.463977 | 0.498331 | 0.457 |
| 20 | 20_71507601_A_G | 71507601 | G | A | CEP72 | intronic | 8.1577E-12 | 0.37366 | 2.304E-13 | 0.019248 | 0.0074416 | 0.023113 | 0.0028158 | 0.0083647 | 0.0031533 | 0.466114 | 0.497997 | 0.459471 |
| 20 | 20_71507726_G_A | 71507726 | A | G | CEP72 | intronic | 1.1661E-11 | 0.2289 | 2.611E-13 | 0.019024 | 0.0099589 | 0.022989 | 0.0028042 | 0.008277 | 0.0031435 | 0.461057 | 0.495995 | 0.453108 |
| 20 | 20_71507893_T_C | 71507893 | C | T | CEP72 | intronic | 2.7426E-10 | 0.82441 | 1.279E-12 | 0.017741 | 0.0018589 | 0.022321 | 0.0028103 | 0.008378 | 0.0031454 | 0.462621 | 0.496662 | 0.455085 |
| 20 | 20_71508020_T_C | 71508020 | C | T | CEP72 | intronic | 1.3091E-10 | 0.57219 | 1.67E-12 | 0.018077 | 0.0047286 | 0.022233 | 0.0028131 | 0.0083717 | 0.0031493 | 0.462934 | 0.496996 | 0.455517 |
| 20 | 20_71508115_G_A | 71508115 | A | G | CEP72 | intronic | 1.4051E-10 | 0.56108 | 1.776E-12 | 0.018031 | 0.0048597 | 0.022173 | 0.0028106 | 0.008361 | 0.0031446 | 0.461057 | 0.495661 | 0.453046 |
| 20 | 20_71508144_A_G | 71508144 | G | A | CEP72 | intronic | 1.1298E-10 | 0.41431 | 1.712E-12 | 0.018106 | 0.0068379 | 0.022159 | 0.0028078 | 0.0083764 | 0.0031403 | 0.461839 | 0.495995 | 0.454034 |
| 20 | 20_71508512_G_C | 71508512 | C | G | CEP72 | intronic | 1.2721E-10 | 0.72198 | 4.167E-13 | 0.018026 | 0.0029635 | 0.022749 | 0.0028032 | 0.0083288 | 0.0031378 | 0.462621 | 0.496328 | 0.455023 |
| 20 | 20_71508777_G_A | 71508777 | A | G | CEP72 | exonic | 9.672E-10 | 0.57298 | 3.937E-12 | 0.017147 | 0.0047287 | 0.021758 | 0.0028042 | 0.0083891 | 0.0031354 | 0.466427 | 0.493992 | 0.4591 |
| 20 | 20_71508826_C_T | 71508826 | T | C | CEP72 | exonic | 1.1777E-10 | 0.58527 | 4.408E-13 | 0.018026 | 0.0045838 | 0.022647 | 0.0027981 | 0.0083999 | 0.0031269 | 0.463977 | 0.49466 | 0.45632 |
| 20 | 20_71508833_C_G | 71508833 | G | C | CEP72 | exonic | 8.7383E-11 | 0.41547 | 2.727E-13 | 0.01836 | 0.0067958 | 0.023191 | 0.0028301 | 0.0083456 | 0.0031737 | 0.425659 | 0.46028 | 0.419251 |
| 20 | 20_71508910_G_T | 71508910 | T | G | CEP72 | exonic | 5.0399E-11 | 0.47372 | 6.468E-13 | 0.018426 | 0.0060091 | 0.022568 | 0.0028047 | 0.0083873 | 0.0031387 | 0.462361 | 0.5 | 0.455394 |
| 20 | 20_71509195_A_G | 71509195 | G | A | CEP72 | intronic | 1.8366E-11 | 0.56405 | 1.357E-13 | 0.018845 | 0.0048258 | 0.023225 | 0.002805 | 0.008366 | 0.0031383 | 0.461526 | 0.497997 | 0.454034 |
| 20 | 20_71509237_T_C | 71509237 | C | T | CEP72 | intronic | 1.2741E-10 | 0.56479 | 7.719E-13 | 0.017997 | 0.0047707 | 0.022466 | 0.0027988 | 0.0082862 | 0.0031351 | 0.461891 | 0.493658 | 0.453664 |
| 20 | 20_71509431_G_A | 71509431 | A | G | CEP72 | intronic | 7.3037E-11 | 0.46512 | 3.794E-13 | 0.018326 | 0.0061714 | 0.02283 | 0.0028132 | 0.0084488 | 0.0031434 | 0.463195 | 0.497664 | 0.45595 |
| 20 | 20_71509633_G_A | 71509633 | A | G | CEP72 | intronic | 8.4866E-10 | 0.48466 | 6.425E-12 | 0.017172 | 0.0057785 | 0.021562 | 0.0027988 | 0.0082688 | 0.0031386 | 0.461057 | 0.49733 | 0.453355 |
| 20 | 20_71509676_A_G | 71509676 | G | A | CEP72 | intronic | 5.9999E-10 | 0.39107 | 8.443E-12 | 0.017345 | 0.0071743 | 0.02142 | 0.0028019 | 0.0083649 | 0.0031358 | 0.462465 | 0.497997 | 0.455146 |
| 20 | 20_71509959_G_C | 71509959 | C | G | CEP72 | intronic | 4.2879E-10 | 0.45102 | 3.092E-12 | 0.017557 | 0.0063051 | 0.021956 | 0.0028123 | 0.0083653 | 0.0031484 | 0.464446 | 0.497997 | 0.457494 |
| 20 | 20_71510160_G_A | 71510160 | A | G | CEP72 | intronic | 4.3952E-10 | 0.63855 | 1.862E-12 | 0.017535 | 0.0039351 | 0.022153 | 0.0028103 | 0.0083775 | 0.0031447 | 0.461891 | 0.496662 | 0.45422 |
| 20 | 20_71510867_A_G | 71510867 | G | A | CEP72 | intronic | 7.3165E-10 | 0.54933 | 4.786E-12 | 0.017379 | 0.0050456 | 0.021821 | 0.0028218 | 0.0084267 | 0.0031571 | 0.461526 | 0.499666 | 0.454467 |
| 20 | 20_71510988_T_C | 71510988 | C | T | CEP72 | intronic | 2.1613E-09 | 0.62044 | 1.705E-11 | 0.016864 | 0.0041902 | 0.021197 | 0.0028177 | 0.0084612 | 0.00315 | 0.459285 | 0.496328 | 0.452428 |
| 20 | 20_71511009_T_C | 71511009 | C | T | CEP72 | intronic | 6.7148E-09 | 0.77686 | 2.696E-11 | 0.016321 | 0.002387 | 0.020975 | 0.002815 | 0.008422 | 0.0031483 | 0.458503 | 0.497997 | 0.451192 |
| 20 | 20_71511230_G_A | 71511230 | A | G | CEP72 | intronic | 9.0539E-11 | 0.62059 | 2.722E-13 | 0.018247 | 0.0041034 | 0.023047 | 0.002815 | 0.0082895 | 0.0031539 | 0.46382 | 0.488318 | 0.454961 |
| 20 | 20_71511306_T_C | 71511306 | C | T | CEP72 | intronic | 3.7542E-11 | 0.47725 | 1.716E-13 | 0.018788 | 0.0058337 | 0.023553 | 0.0028409 | 0.0082079 | 0.0031961 | 0.428058 | 0.46729 | 0.420796 |
| 20 | 20_71511456_A_G | 71511456 | G | A | CEP72 | intronic | 2.5968E-10 | 0.60436 | 7.799E-13 | 0.01781 | 0.0043298 | 0.022602 | 0.0028176 | 0.0083566 | 0.0031546 | 0.462256 | 0.493324 | 0.454034 |
| 20 | 20_71511580_C_T | 71511580 | T | C | CEP72 | intronic | 3.0536E-11 | 0.41376 | 1.836E-13 | 0.018925 | 0.0067528 | 0.023578 | 0.0028484 | 0.0082624 | 0.0032034 | 0.427328 | 0.463618 | 0.42061 |
| 20 | 20_71511773_G_A | 71511773 | A | G | CEP72 | intronic | 1.3724E-09 | 0.57864 | 5.184E-12 | 0.016983 | 0.0045289 | 0.021758 | 0.0028032 | 0.0081547 | 0.0031531 | 0.465697 | 0.48765 | 0.457062 |
| 20 | 20_71511780_A_G | 71511780 | G | A | CEP72 | intronic | 5.2949E-10 | 0.55278 | 1.843E-12 | 0.017427 | 0.0048501 | 0.022233 | 0.0028062 | 0.0081706 | 0.0031555 | 0.465593 | 0.486649 | 0.456753 |
| 20 | 20_71511792_C_G | 71511792 | G | C | CEP72 | intronic | 4.7231E-10 | 0.50979 | 1.748E-12 | 0.017449 | 0.0053659 | 0.022236 | 0.0028016 | 0.0081405 | 0.0031525 | 0.466166 | 0.485981 | 0.457309 |
| 20 | 20_71512430_T_G | 71512430 | G | T | CEP72 | intronic | 1.7224E-10 | 0.56034 | 3.427E-13 | 0.017956 | 0.0048685 | 0.02292 | 0.0028125 | 0.0083604 | 0.0031499 | 0.460744 | 0.499666 | 0.45354 |
| 20 | 20_71512531_C_T | 71512531 | T | C | CEP72 | intronic | 1.5154E-10 | 0.77628 | 1.814E-13 | 0.018036 | 0.0023933 | 0.023183 | 0.0028164 | 0.0084219 | 0.0031491 | 0.459389 | 0.499332 | 0.451996 |
| 20 | 20_71512676_A_G | 71512676 | G | A | CEP72 | intronic | 8.0856E-10 | 0.87809 | 1.514E-12 | 0.017291 | 0.0012902 | 0.022262 | 0.0028147 | 0.0084114 | 0.0031474 | 0.461839 | 0.495995 | 0.454034 |
| 20 | 20_71512785_C_T | 71512785 | T | C | CEP72 | intronic | 9.8053E-10 | 0.8638 | 2.178E-12 | 0.017209 | 0.0014381 | 0.022132 | 0.0028153 | 0.0083832 | 0.0031516 | 0.461474 | 0.499666 | 0.454281 |
| 20 | 20_71512982_G_A | 71512982 | A | G | CEP72 | intronic | 9.8992E-11 | 0.4537 | 1.563E-12 | 0.018333 | 0.0063431 | 0.022401 | 0.0028341 | 0.0084659 | 0.0031691 | 0.466218 | 0.496662 | 0.459348 |
| 20 | 20_71513184_A_G | 71513184 | G | A | CEP72 | intronic | 6.5839E-10 | 0.37051 | 5.334E-12 | 0.01741 | 0.0075654 | 0.021741 | 0.0028191 | 0.0084481 | 0.0031525 | 0.462152 | 0.497664 | 0.455579 |
| 20 | 20_71513195_T_C | 71513195 | C | T | CEP72 | intronic | 1.2668E-10 | 0.38704 | 3.075E-12 | 0.018143 | 0.0072835 | 0.022029 | 0.0028212 | 0.0084202 | 0.0031585 | 0.464863 | 0.49733 | 0.458853 |
| 20 | 20_71513248_A_G | 71513248 | G | A | CEP72 | intronic | 6.8661E-11 | 0.30622 | 7.2E-13 | 0.018385 | 0.0085802 | 0.022641 | 0.0028182 | 0.0083858 | 0.0031552 | 0.461839 | 0.499332 | 0.454652 |
| 20 | 20_71513273_G_A | 71513273 | A | G | CEP72 | intronic | 6.6466E-11 | 0.37745 | 4.181E-13 | 0.018596 | 0.007322 | 0.02321 | 0.0028484 | 0.0082959 | 0.0032015 | 0.425295 | 0.460614 | 0.418757 |
| 20 | 20_71513356_A_G | 71513356 | G | A | CEP72 | intronic | 6.439E-11 | 0.3425 | 4.758E-13 | 0.018566 | 0.0078254 | 0.0231 | 0.0028418 | 0.0082439 | 0.0031941 | 0.424617 | 0.462283 | 0.417645 |
| 20 | 20_71513392_C_G | 71513392 | G | C | CEP72 | intronic | 1.2519E-10 | 0.40224 | 1.086E-12 | 0.018034 | 0.0069614 | 0.022349 | 0.0028033 | 0.008311 | 0.0031393 | 0.460953 | 0.496662 | 0.453108 |
| 20 | 20_71513450_C_T | 71513450 | T | C | CEP72 | intronic | 3.1632E-10 | 0.39788 | 1.581E-12 | 0.017634 | 0.0070673 | 0.022167 | 0.0028032 | 0.0083595 | 0.0031366 | 0.460797 | 0.497664 | 0.453973 |
| 20 | 20_71513469_C_A | 71513469 | A | C | CEP72 | intronic | 5.8804E-10 | 0.45376 | 2.738E-12 | 0.017364 | 0.0062628 | 0.02193 | 0.0028035 | 0.0083598 | 0.0031371 | 0.4621 | 0.498999 | 0.454899 |

| | | | | | | | | | | | | | | | | | | |
|---|---|---|---|---|---|---|---|---|---|---|---|---|---|---|---|---|---|---|
| 20 | 20_71513498_T_C | 71513498 | C | T | CEP72 | intronic | 6.15E-10 | 0.43093 | 2.752E-12 | 0.017349 | 0.0065458 | 0.021951 | 0.0028043 | 0.0083112 | 0.0031403 | 0.46137 | 0.496662 | 0.453602 |
| 20 | 20_71513522_A_G | 71513522 | G | A | CEP72 | intronic | 5.9507E-10 | 0.60492 | 1.892E-12 | 0.017356 | 0.0042918 | 0.022116 | 0.0028031 | 0.0082959 | 0.0031404 | 0.461474 | 0.497664 | 0.453911 |
| 20 | 20_71513865_C_T | 71513865 | T | C | CEP72 | intronic | 1.3082E-10 | 0.79423 | 2.742E-13 | 0.01804 | 0.0021627 | 0.022996 | 0.0028072 | 0.0082919 | 0.0031473 | 0.463403 | 0.49466 | 0.455641 |
| 20 | 20_71513879_A_G | 71513879 | G | A | CEP72 | intronic | 3.657E-11 | 0.51974 | 1.606E-13 | 0.018781 | 0.0052801 | 0.023575 | 0.0028382 | 0.008202 | 0.0031952 | 0.426963 | 0.465287 | 0.419869 |
| 20 | 20_71514336_T_C | 71514336 | C | T | CEP72 | intronic | 1.0738E-11 | 0.41555 | 5.976E-14 | 0.019154 | 0.0064672 | 0.023949 | 0.0028183 | 0.0079434 | 0.0031896 | 0.430873 | 0.474967 | 0.422711 |
| 20 | 20_71514872_G_A | 71514872 | A | G | CEP72 | intronic | 1.5661E-10 | 0.75162 | 4.603E-13 | 0.017978 | 0.0026396 | 0.022766 | 0.0028096 | 0.0083396 | 0.003146 | 0.461162 | 0.496996 | 0.453417 |
| 20 | 20_71515508_C_A | 71515508 | A | C | CEP72 | intronic | 1.4667E-10 | 0.41938 | 1.026E-12 | 0.018112 | 0.0065476 | 0.022707 | 0.0028261 | 0.0081086 | 0.0031861 | 0.425295 | 0.463618 | 0.418201 |
| 20 | 20_71515717_C_T | 71515717 | T | C | CEP72 | intronic | 9.6319E-11 | 0.36931 | 9.594E-13 | 0.018352 | 0.0073905 | 0.022766 | 0.0028353 | 0.0082321 | 0.0031902 | 0.423783 | 0.462951 | 0.416533 |
| 20 | 20_71515894_A_G | 71515894 | G | A | CEP72 | intronic | 1.0277E-10 | 0.22666 | 4.039E-12 | 0.018263 | 0.0098917 | 0.02206 | 0.0028258 | 0.0081816 | 0.0031806 | 0.427171 | 0.466288 | 0.419931 |
| 20 | 20_71515993_A_G | 71515993 | G | A | CEP72 | intronic | 4.1386E-10 | 0.45025 | 4.964E-12 | 0.017487 | 0.006283 | 0.021666 | 0.0027984 | 0.0083218 | 0.003137 | 0.462569 | 0.498665 | 0.455394 |
| 20 | 20_71516026_G_C | 71516026 | C | G | CEP72 | intronic | 2.2333E-10 | 0.45598 | 4.333E-12 | 0.017755 | 0.0061758 | 0.02174 | 0.0027985 | 0.0082843 | 0.003139 | 0.461735 | 0.499666 | 0.45459 |
| 20 | 20_71516696_C_T | 71516696 | T | C | CEP72 | intronic | 5.9996E-11 | 0.38528 | 1.858E-13 | 0.018535 | 0.0071356 | 0.023478 | 0.0028325 | 0.0082188 | 0.0031905 | 0.425503 | 0.464286 | 0.418324 |
| 20 | 20_71517590_C_T | 71517590 | T | C | CEP72 | intronic | 6.3149E-11 | 0.45273 | 7.81E-13 | 0.018289 | 0.0061585 | 0.022509 | 0.0027982 | 0.0082017 | 0.0031418 | 0.464342 | 0.493658 | 0.456567 |
| 20 | 20_71517751_G_C | 71517751 | C | G | CEP72 | intronic | 3.6977E-11 | 0.29602 | 3.702E-13 | 0.018544 | 0.0086046 | 0.022842 | 0.002803 | 0.0082339 | 0.0031437 | 0.464654 | 0.488985 | 0.456073 |
| 20 | 20_71518014_C_A | 71518014 | A | C | CEP72 | intronic | 3.7651E-10 | 0.53231 | 1.884E-12 | 0.017583 | 0.0052107 | 0.022143 | 0.0028073 | 0.0083439 | 0.003144 | 0.462465 | 0.5 | 0.455517 |
| 20 | 20_71518465_A_C | 71518465 | C | A | CEP72 | intronic | 2.2146E-10 | 0.31389 | 1.433E-12 | 0.017998 | 0.0083941 | 0.02255 | 0.0028363 | 0.0083349 | 0.0031847 | 0.425138 | 0.461949 | 0.418324 |
| 20 | 20_71518658_A_C | 71518658 | C | A | CEP72 | intronic | 5.6385E-11 | 0.48553 | 4.269E-13 | 0.018322 | 0.0058315 | 0.02267 | 0.002796 | 0.0083613 | 0.0031282 | 0.464759 | 0.493658 | 0.457062 |
| 20 | 20_71518659_G_C | 71518659 | C | G | CEP72 | intronic | 5.6385E-11 | 0.48553 | 4.269E-13 | 0.018322 | 0.0058315 | 0.02267 | 0.002796 | 0.0083613 | 0.0031282 | 0.464759 | 0.493658 | 0.457062 |
| 20 | 20_71518731_C_T | 71518731 | C | T | CEP72 | intronic | 2.75E-10 | 0.74569 | 1.519E-12 | 0.017697 | 0.0027265 | 0.02217 | 0.0028036 | 0.0084064 | 0.0031346 | 0.465071 | 0.493658 | 0.457432 |
| 20 | 20_71519131_C_T | 71519131 | T | C | CEP72 | intronic | 2.6068E-10 | 0.55132 | 6.064E-12 | 0.017752 | 0.0050272 | 0.021605 | 0.0028086 | 0.0084381 | 0.0031412 | 0.461839 | 0.497664 | 0.455208 |
| 20 | 20_71519357_G_A | 71519357 | A | G | CEP72 | intronic | 2.2112E-10 | 0.49592 | 5.07E-12 | 0.017807 | 0.0057119 | 0.021665 | 0.002806 | 0.0083884 | 0.0031383 | 0.463351 | 0.49466 | 0.455579 |
| 20 | 20_71519390_G_A | 71519390 | A | G | CEP72 | intronic | 6.9122E-10 | 0.44729 | 1.278E-11 | 0.017317 | 0.0064291 | 0.021236 | 0.0028075 | 0.00846 | 0.0031362 | 0.462569 | 0.497664 | 0.455208 |
| 20 | 20_71519415_C_T | 71519415 | T | C | CEP72 | intronic | 7.9435E-10 | 0.58201 | 6.749E-12 | 0.017262 | 0.0046919 | 0.021496 | 0.0028087 | 0.0085237 | 0.0031323 | 0.461266 | 0.497997 | 0.453725 |
| 20 | 20_71519700_G_A | 71519700 | A | G | CEP72 | intronic | 5.9999E-10 | 0.60317 | 7.779E-12 | 0.017372 | 0.0043463 | 0.021498 | 0.0028062 | 0.0083606 | 0.0031417 | 0.463143 | 0.498999 | 0.456135 |
| 20 | 20_71519711_T_C | 71519711 | C | T | CEP72 | intronic | 8.3025E-10 | 0.438 | 4.751E-12 | 0.017362 | 0.0063875 | 0.021975 | 0.0028281 | 0.0082358 | 0.0031788 | 0.42274 | 0.461615 | 0.415544 |
| 20 | 20_71519969_G_A | 71519969 | A | G | CEP72 | intronic | 4.4348E-10 | 0.40673 | 1.08E-11 | 0.017413 | 0.0068425 | 0.021269 | 0.0027915 | 0.0082473 | 0.0031299 | 0.464029 | 0.496662 | 0.456753 |
| 20 | 20_71520317_C_T | 71520317 | T | C | CEP72 | intronic | 3.8915E-09 | 0.80527 | 3.384E-11 | 0.016427 | 0.0021015 | 0.020607 | 0.0027895 | 0.0085243 | 0.0031087 | 0.461891 | 0.499332 | 0.454961 |
| 20 | 20_71520387_C_T | 71520387 | T | C | CEP72 | intronic | 4.0412E-09 | 0.76926 | 2.282E-11 | 0.016331 | 0.0024787 | 0.020692 | 0.0027761 | 0.0084499 | 0.0030944 | 0.461162 | 0.498999 | 0.453787 |
| 20 | 20_71520414_A_G | 71520414 | G | A | CEP72 | intronic | 4.667E-09 | 0.40317 | 5.802E-11 | 0.016107 | 0.0068735 | 0.020129 | 0.0027492 | 0.0082221 | 0.0030738 | 0.463143 | 0.493658 | 0.455146 |
| 20 | 20_71520420_T_G | 71520420 | G | T | CEP72 | intronic | 4.2668E-09 | 0.40374 | 5.731E-11 | 0.016142 | 0.0068607 | 0.020131 | 0.0027483 | 0.0082168 | 0.0030731 | 0.463038 | 0.493992 | 0.455085 |
| 20 | 20_71520456_G_A | 71520456 | A | G | CEP72 | intronic | 5.8416E-09 | 0.35077 | 6.825E-11 | 0.016016 | 0.0076246 | 0.020085 | 0.0027513 | 0.0081714 | 0.0030784 | 0.463038 | 0.490988 | 0.454529 |
| 20 | 20_71520666_G_A | 71520666 | A | G | CEP72 | intronic | 7.1752E-10 | 0.61815 | 9.012E-12 | 0.017127 | 0.0042463 | 0.021126 | 0.0027794 | 0.0085186 | 0.003097 | 0.461787 | 0.495661 | 0.455517 |
| 20 | 20_71520711_G_A | 71520711 | A | G | CEP72 | intronic | 1.4762E-10 | 0.2974 | 4.4E-12 | 0.017743 | 0.008687 | 0.021422 | 0.0027689 | 0.0083366 | 0.003094 | 0.462986 | 0.498331 | 0.455826 |
| 20 | 20_71520741_G_A | 71520741 | A | G | CEP72 | intronic | 1.3541E-10 | 0.27807 | 2.936E-12 | 0.017715 | 0.0090482 | 0.021498 | 0.0027589 | 0.0083419 | 0.0030797 | 0.463247 | 0.49733 | 0.45595 |
| 20 | 20_71520769_G_A | 71520769 | A | G | CEP72 | intronic | 4.1929E-10 | 0.41245 | 6.614E-12 | 0.01721 | 0.0068203 | 0.021119 | 0.0027551 | 0.0083216 | 0.0030759 | 0.462934 | 0.497997 | 0.455702 |
| 20 | 20_71520823_T_C | 71520823 | C | T | CEP72 | intronic | 3.634E-09 | 0.65079 | 3.564E-11 | 0.016318 | 0.0038199 | 0.020422 | 0.0027657 | 0.0084386 | 0.0030843 | 0.464602 | 0.496996 | 0.457494 |
| 20 | 20_71520915_T_C | 71520915 | C | T | CEP72 | exonic | 1.9738E-09 | 0.56621 | 1.342E-11 | 0.016624 | 0.0048278 | 0.020904 | 0.0027707 | 0.008416 | 0.0030905 | 0.462308 | 0.496328 | 0.454652 |
| 20 | 20_71520972_C_T | 71520972 | T | C | CEP72 | exonic | 8.4947E-10 | 0.51394 | 6.486E-12 | 0.016939 | 0.0054969 | 0.021144 | 0.0027608 | 0.0084216 | 0.0030784 | 0.464237 | 0.495327 | 0.456753 |
| 20 | 20_71521009_C_T | 71521009 | T | C | CEP72 | exonic | 1.1725E-09 | 0.48305 | 1.169E-11 | 0.016803 | 0.0059144 | 0.020889 | 0.0027618 | 0.0084321 | 0.0030791 | 0.463403 | 0.49733 | 0.456135 |
| 20 | 20_71521098_G_C | 71521098 | C | G | CEP72 | intronic | 5.7552E-10 | 0.54816 | 3.522E-12 | 0.017271 | 0.0050154 | 0.021679 | 0.0027869 | 0.0083517 | 0.003117 | 0.42592 | 0.458278 | 0.419931 |
| 20 | 20_71521220_A_G | 71521220 | G | A | CEP72 | intronic | 4.4455E-10 | 0.45084 | 2.122E-12 | 0.017191 | 0.0063081 | 0.0216 | 0.002756 | 0.0083661 | 0.0030741 | 0.46309 | 0.49466 | 0.45527 |
| 20 | 20_71521242_A_C | 71521242 | C | A | CEP72 | intronic | 1.5754E-09 | 0.51239 | 8.286E-12 | 0.016583 | 0.0055245 | 0.020906 | 0.0027471 | 0.0084328 | 0.0030594 | 0.463768 | 0.495327 | 0.456197 |
| 20 | 20_71521288_C_G | 71521288 | G | C | CEP72 | intronic | 1.3346E-09 | 0.49757 | 8.832E-12 | 0.016655 | 0.0056977 | 0.020884 | 0.0027468 | 0.0083997 | 0.0030603 | 0.463247 | 0.493992 | 0.455332 |
| 20 | 20_71521301_T_C | 71521301 | C | T | CEP72 | intronic | 5.5078E-10 | 0.42312 | 2.922E-12 | 0.017153 | 0.0066197 | 0.021585 | 0.0027649 | 0.008264 | 0.0030918 | 0.426754 | 0.465287 | 0.419622 |
| 20 | 20_71521386_C_T | 71521386 | T | C | CEP72 | intronic | 1.7981E-09 | 0.53333 | 1.697E-11 | 0.016479 | 0.0052631 | 0.020524 | 0.0027397 | 0.0084491 | 0.0030497 | 0.462204 | 0.498999 | 0.455394 |
| 20 | 20_71521437_T_C | 71521437 | C | T | CEP72 | intronic | 1.536E-09 | 0.56716 | 7.616E-12 | 0.016529 | 0.0048063 | 0.020872 | 0.0027364 | 0.0083991 | 0.0030489 | 0.464446 | 0.497997 | 0.457494 |
| 20 | 20_71521507_T_C | 71521507 | C | T | CEP72 | intronic | 1.3558E-09 | 0.60051 | 3.091E-12 | 0.016444 | 0.0042499 | 0.021167 | 0.0027133 | 0.0081156 | 0.0030353 | 0.466792 | 0.486315 | 0.458112 |
| 20 | 20_71521510_T_C | 71521510 | C | T | CEP72 | intronic | 1.0758E-10 | 0.38108 | 7.69E-13 | 0.017817 | 0.0072972 | 0.0221 | 0.0027598 | 0.0083311 | 0.0030838 | 0.427797 | 0.461615 | 0.421537 |
| 20 | 20_71521519_C_T | 71521519 | T | C | CEP72 | intronic | 4.138E-10 | 0.54636 | 2.792E-12 | 0.017079 | 0.0050833 | 0.021268 | 0.0027333 | 0.0084268 | 0.0030436 | 0.465541 | 0.497664 | 0.45873 |
| 20 | 20_71521669_C_T | 71521669 | T | C | CEP72 | intronic | 8.9724E-10 | 0.93178 | 7.63E-13 | 0.016756 | 0.0006985 | 0.021939 | 0.002735 | 0.008159 | 0.0030608 | 0.465853 | 0.489987 | 0.457679 |
| 20 | 20_71521698_T_C | 71521698 | C | T | CEP72 | intronic | 1.709E-10 | 0.63919 | 5.405E-13 | 0.017489 | 0.0038376 | 0.022098 | 0.0027389 | 0.0081856 | 0.0030628 | 0.466479 | 0.48765 | 0.457988 |
| 20 | 20_71521711_T_C | 71521711 | C | T | CEP72 | intronic | 2.201E-10 | 0.6423 | 7.067E-13 | 0.017409 | 0.0037956 | 0.022031 | 0.002743 | 0.0081716 | 0.0030692 | 0.466375 | 0.485981 | 0.457556 |
| 20 | 20_71521773_T_C | 71521773 | C | T | CEP72 | intronic | 3.0296E-10 | 0.47081 | 2.382E-12 | 0.017289 | 0.0059077 | 0.021532 | 0.0027455 | 0.008192 | 0.0030715 | 0.471796 | 0.480307 | 0.462931 |
| 20 | 20_71522099_A_G | 71522099 | G | A | CEP72 | intronic | 1.0051E-09 | 0.37707 | 1.18E-11 | 0.016787 | 0.007323 | 0.020832 | 0.0027481 | 0.0082904 | 0.0030714 | 0.46382 | 0.493324 | 0.455888 |
| 20 | 20_71522302_G_C | 71522302 | C | G | CEP72(XM_024981570.1:c.-312C>G) | UTR5 | 2.3621E-09 | 0.62731 | 2.111E-11 | 0.016431 | 0.0040003 | 0.020618 | 0.0027518 | 0.0082391 | 0.0030782 | 0.463455 | 0.493324 | 0.455455 |
| 20 | 20_71522384_A_G | 71522384 | G | A | CEP72(XM_024981570.1:c.-394T>C) | UTR5 | 9.5248E-10 | 0.67827 | 1.427E-11 | 0.016798 | 0.0034528 | 0.020702 | 0.002746 | 0.0083236 | 0.0030645 | 0.463195 | 0.495327 | 0.455517 |
| 20 | 20_71522397_C_T | 71522397 | T | C | CEP72(XM_024981570.1:c.-407G>A) | UTR5 | 8.2673E-10 | 0.67936 | 1.268E-11 | 0.016873 | 0.0034429 | 0.020768 | 0.0027482 | 0.0083295 | 0.0030667 | 0.463768 | 0.493658 | 0.455888 |
| 20 | 20_71522403_T_C | 71522403 | C | T | CEP72(XM_024981570.1:c.-413A>G) | UTR5 | 8.211E-10 | 0.66651 | 1.133E-11 | 0.016869 | 0.0035799 | 0.02082 | 0.002747 | 0.0083073 | 0.0030669 | 0.463664 | 0.495661 | 0.456135 |
| 20 | 20_71522442_G_A | 71522442 | A | G | CEP72(XM_024981570.1:c.-452C>T) | UTR5 | 3.6229E-10 | 0.63688 | 4.137E-12 | 0.017297 | 0.0039599 | 0.021328 | 0.002759 | 0.0083885 | 0.0030765 | 0.461214 | 0.496662 | 0.453417 |
| 20 | 20_71522472_T_C | 71522472 | C | T | CEP72(XM_024981570.1:c.-482A>G) | UTR5 | 1.2536E-09 | 0.81036 | 1.438E-11 | 0.016817 | 0.0020237 | 0.020856 | 0.002769 | 0.0084331 | 0.0030879 | 0.46137 | 0.499332 | 0.454096 |
| 20 | 20_71522592_A_C | 71522592 | C | A | CEP72(XM_024981570.1:c.-602T>G) | UTR5 | 1.3191E-09 | 0.75921 | 9.944E-12 | 0.016831 | 0.0025952 | 0.021056 | 0.0027752 | 0.0084669 | 0.0030931 | 0.460327 | 0.499332 | 0.453108 |
| 20 | 20_71522628_A_C | 71522628 | C | A | CEP72(XM_024981570.1:c.-638T>G) | UTR5 | 1.3254E-09 | 0.71903 | 8.673E-12 | 0.016788 | 0.0030119 | 0.021096 | 0.0027684 | 0.0083719 | 0.00309 | 0.461318 | 0.498999 | 0.453973 |
| 20 | 20_71522678_A_G | 71522678 | G | A | CEP72(XM_024981570.1:c.-688T>C) | UTR5 | 4.283E-10 | 0.62856 | 2.416E-12 | 0.017453 | 0.0039765 | 0.021975 | 0.0027954 | 0.0082203 | 0.0031357 | 0.425399 | 0.464286 | 0.418201 |
| 20 | 20_71522781_C_T | 71522781 | T | C | CEP72(XM_024981570.1:c.-791G>A) | UTR5 | 1.6832E-10 | 0.42955 | 1.378E-12 | 0.017974 | 0.0066862 | 0.022276 | 0.0028138 | 0.008464 | 0.0031435 | 0.424304 | 0.45761 | 0.418139 |
| 20 | 20_71522953_G_C | 71522953 | C | G | CEP72(XM_024981570.1:c.-963C>G) | UTR5 | 1.0525E-09 | 0.6188 | 4.425E-12 | 0.017005 | 0.0041962 | 0.021538 | 0.0027872 | 0.0084338 | 0.0031112 | 0.463403 | 0.493992 | 0.455517 |
| 20 | 20_71523392_A_G | 71523392 | G | A | CEP72 | intronic | 7.4326E-10 | 0.69984 | 1.352E-11 | 0.01761 | 0.0032885 | 0.021149 | 0.0028036 | 0.0085299 | 0.0031272 | 0.465906 | 0.498999 | 0.45978 |
| 20 | 20_71523426_A_C | 71523426 | C | A | CEP72 | intronic | 2.3193E-10 | 0.56807 | 4.055E-12 | 0.017635 | 0.0048269 | 0.021526 | 0.0027823 | 0.0084549 | 0.0031038 | 0.467991 | 0.498665 | 0.461819 |
| 20 | 20_71523471_A_C | 71523471 | C | A | CEP72 | intronic | 3.89E-10 | 0.42612 | 9.305E-12 | 0.01744 | 0.0067326 | 0.021199 | 0.0027866 | 0.0084597 | 0.0031098 | 0.468251 | 0.499666 | 0.462313 |
| 20 | 20_71523553_T_C | 71523553 | C | T | CEP72 | intronic | 5.7543E-10 | 0.49448 | 1.182E-11 | 0.017252 | 0.0057838 | 0.021062 | 0.002784 | 0.0084658 | 0.0031054 | 0.467365 | 0.499332 | 0.461448 |
| 20 | 20_71523584_G_C | 71523584 | C | G | CEP72 | intronic | 1.4523E-09 | 0.67194 | 2.052E-11 | 0.016856 | 0.0035807 | 0.020836 | 0.0027864 | 0.0084553 | 0.0031088 | 0.468564 | 0.499332 | 0.462622 |
| 20 | 20_71523961_A_C | 71523961 | C | A | CEP72 | intronic | 1.0995E-09 | 0.40865 | 2.888E-11 | 0.016908 | 0.0069574 | 0.020596 | 0.0027745 | 0.0084202 | 0.0030961 | 0.461005 | 0.5 | 0.453787 |
| 20 | 20_71523977_A_C | 71523977 | C | A | CEP72 | intronic | 1.1915E-09 | 0.41947 | 2.716E-11 | 0.016885 | 0.0068025 | 0.020642 | 0.0027765 | 0.0084258 | 0.0030988 | 0.460536 | 0.499666 | 0.453169 |
| 20 | 20_71523986_T_C | 71523986 | C | T | CEP72 | intronic | 1.3717E-09 | 0.41324 | 3.28E-11 | 0.016827 | 0.0068895 | 0.020564 | 0.0027773 | 0.0084203 | 0.0031001 | 0.460953 | 0.498665 | 0.453478 |
| 20 | 20_71524031_A_G | 71524031 | G | A | CEP72 | intronic | 3.5727E-10 | 0.39908 | 1.484E-11 | 0.017424 | 0.0070636 | 0.02095 | 0.0027782 | 0.0083765 | 0.003104 | 0.461787 | 0.495327 | 0.453849 |
| 20 | 20_71524330_C_T | 71524330 | T | C | CEP72 | intronic | 3.8921E-10 | 0.30853 | 8.153E-12 | 0.017554 | 0.0083579 | 0.021531 | 0.0028049 | 0.0082077 | 0.0031497 | 0.423939 | 0.464286 | 0.416471 |

| | | | | | | | | | | | | | | | | | | |
|---|---|---|---|---|---|---|---|---|---|---|---|---|---|---|---|---|---|---|
| 20 | 20_71524447_C_T | 71524447 | T | C | CEP72 | intronic | 2.9013E-10 | 0.5198 | 2.52E-12 | 0.017518 | 0.0053257 | 0.021795 | 0.0027789 | 0.0082743 | 0.0031126 | 0.459598 | 0.497664 | 0.451687 |
| 20 | 20_71524529_T_C | 71524529 | C | T | CEP72 | intronic | 3.7929E-10 | 0.50393 | 7.616E-12 | 0.017373 | 0.0055335 | 0.021254 | 0.0027742 | 0.0082796 | 0.0031047 | 0.462725 | 0.496662 | 0.455208 |
| 20 | 20_71524534_A_G | 71524534 | G | A | CEP72 | intronic | 5.5938E-10 | 0.5431 | 1.026E-11 | 0.017195 | 0.0050289 | 0.021113 | 0.0027727 | 0.0082693 | 0.0031036 | 0.462621 | 0.496662 | 0.455085 |
| 20 | 20_71524890_T_A | 71524890 | A | T | CEP72 | intronic | 1.2612E-09 | 0.70986 | 2.143E-11 | 0.016899 | 0.0031131 | 0.020821 | 0.0027829 | 0.0083676 | 0.0031095 | 0.462725 | 0.493992 | 0.454714 |
| 20 | 20_71524905_A_C | 71524905 | C | A | CEP72 | intronic | 4.9911E-10 | 0.49813 | 9.559E-12 | 0.017503 | 0.0056183 | 0.0215 | 0.0028143 | 0.0082935 | 0.0031557 | 0.425555 | 0.466622 | 0.417954 |
| 20 | 20_71524931_C_T | 71524931 | T | C | CEP72 | intronic | 4.4174E-10 | 0.46267 | 7.48E-12 | 0.017531 | 0.0060373 | 0.021593 | 0.0028101 | 0.0082201 | 0.0031532 | 0.425816 | 0.468959 | 0.41783 |
| 20 | 20_71525465_T_C | 71525465 | C | T | CEP72 | intronic | 1.3001E-09 | 0.58059 | 2.509E-11 | 0.016817 | 0.004576 | 0.020684 | 0.0027718 | 0.0082821 | 0.0030997 | 0.462361 | 0.491322 | 0.453787 |
| 20 | 20_71525502_C_T | 71525502 | T | C | CEP72 | intronic | 1.9545E-09 | 0.37771 | 4.737E-11 | 0.01664 | 0.0072113 | 0.020441 | 0.0027726 | 0.0081749 | 0.0031071 | 0.462621 | 0.486315 | 0.453169 |
| 20 | 20_71526031_T_A | 71526031 | A | T | CEP72 | intronic | 3.5388E-10 | 0.61618 | 3.984E-12 | 0.017276 | 0.0041264 | 0.021375 | 0.002754 | 0.0082318 | 0.003081 | 0.467052 | 0.487984 | 0.45873 |
| 20 | 20_71526047_C_G | 71526047 | G | C | CEP72 | intronic | 1.5116E-10 | 0.60849 | 1.358E-12 | 0.017629 | 0.0042006 | 0.021843 | 0.0027528 | 0.0082006 | 0.0030816 | 0.467 | 0.487984 | 0.458668 |
| 20 | 20_71526076_G_A | 71526076 | A | G | CEP72 | intronic | 2.0551E-10 | 0.53431 | 3.488E-12 | 0.017546 | 0.0051881 | 0.021426 | 0.0027601 | 0.0083486 | 0.0030799 | 0.467574 | 0.487984 | 0.459348 |
| 20 | 20_71526081_C_A | 71526081 | A | C | CEP72 | intronic | 4.9866E-11 | 0.34435 | 1.14E-12 | 0.018298 | 0.007774 | 0.022196 | 0.0027845 | 0.0082212 | 0.0031207 | 0.430716 | 0.470628 | 0.423329 |
| 20 | 20_71526107_T_C | 71526107 | C | T | CEP72 | intronic | 1.8303E-10 | 0.26748 | 5.267E-12 | 0.0177 | 0.0090383 | 0.021485 | 0.0027765 | 0.0081509 | 0.0031146 | 0.432541 | 0.4753 | 0.424626 |
| 20 | 20_71526148_C_T | 71526148 | T | C | CEP72 | intronic | 1.1693E-09 | 0.43148 | 2.985E-11 | 0.016821 | 0.0066119 | 0.020501 | 0.0027647 | 0.008405 | 0.003084 | 0.46309 | 0.493658 | 0.455085 |
| 20 | 20_71526195_C_T | 71526195 | T | C | CEP72 | intronic | 1.5682E-09 | 0.40937 | 3.641E-11 | 0.016677 | 0.0068904 | 0.020399 | 0.0027624 | 0.008352 | 0.0030824 | 0.465541 | 0.489653 | 0.457247 |
| 20 | 20_71526196_A_G | 71526196 | G | A | CEP72 | intronic | 1.5682E-09 | 0.40937 | 3.641E-11 | 0.016677 | 0.0068904 | 0.020399 | 0.0027624 | 0.008352 | 0.0030824 | 0.465541 | 0.489653 | 0.457247 |
| 20 | 20_71526331_G_C | 71526331 | C | G | CEP72 | intronic | 1.8203E-09 | 0.34055 | 1.952E-11 | 0.016849 | 0.0078118 | 0.021118 | 0.0028021 | 0.0081963 | 0.0031475 | 0.423835 | 0.462617 | 0.416656 |
| 20 | 20_71526355_G_A | 71526355 | A | G | CEP72 | intronic | 3.1649E-09 | 0.56059 | 2.204E-11 | 0.016406 | 0.0048159 | 0.020733 | 0.00277 | 0.0082753 | 0.0030982 | 0.46137 | 0.494326 | 0.453169 |
| 20 | 20_71527024_C_T | 71527024 | T | C | CEP72 | intronic | 1.3063E-10 | 0.39572 | 2.046E-12 | 0.017898 | 0.0069528 | 0.021972 | 0.0027851 | 0.0081866 | 0.0031249 | 0.465384 | 0.491989 | 0.457494 |
| 20 | 20_71527382_C_A | 71527382 | A | C | CEP72 | intronic | 2.3815E-10 | 0.46495 | 2.033E-12 | 0.017691 | 0.0060156 | 0.022026 | 0.0027928 | 0.0082325 | 0.0031322 | 0.463507 | 0.494993 | 0.455826 |
| 20 | 20_71527455_T_G | 71527455 | G | T | CEP72 | intronic | 1.0894E-10 | 0.54901 | 8.487E-13 | 0.018115 | 0.0050564 | 0.022453 | 0.0028068 | 0.0084381 | 0.0031389 | 0.463403 | 0.497664 | 0.456197 |
| 20 | 20_71527778_C_T | 71527778 | T | C | CEP72 | intronic | 5.9973E-10 | 0.35788 | 6.436E-13 | -0.017672 | -0.00762 | -0.023093 | 0.0028547 | 0.0082879 | 0.0032114 | 0.404702 | 0.448598 | 0.396577 |
| 20 | 20_71527779_A_G | 71527779 | G | A | CEP72 | intronic | 5.475E-10 | 0.34826 | 6.496E-13 | -0.017707 | -0.007766 | -0.023083 | 0.0028538 | 0.0082794 | 0.0032106 | 0.404754 | 0.447931 | 0.396763 |
| 20 | 20_71530143_C_A | 71530143 | A | C | CEP72(dist=1008),SLC9A3(dist=7719 | intergenic | 2.6327E-09 | 0.72771 | 4.54E-12 | 0.016742 | 0.0028058 | 0.021925 | 0.0028123 | 0.0080589 | 0.0031687 | 0.473413 | 0.479306 | 0.464661 |
| 20 | 20_71531961_T_C | 71531961 | C | T | CEP72(dist=2826),SLC9A3(dist=7537 | intergenic | 9.9588E-12 | 0.24104 | 4.797E-13 | 0.019215 | 0.009614 | 0.022963 | 0.0028227 | 0.0082004 | 0.0031756 | 0.428318 | 0.46996 | 0.42061 |
| 20 | 20_71532095_G_T | 71532095 | T | G | CEP72(dist=2960),SLC9A3(dist=7524 | intergenic | 1.3236E-10 | 0.32639 | 1.696E-12 | 0.018132 | 0.0080687 | 0.022383 | 0.0028224 | 0.0082216 | 0.0031716 | 0.426389 | 0.467623 | 0.418757 |
| 20 | 20_71532189_A_G | 71532189 | G | A | CEP72(dist=3054),SLC9A3(dist=7514 | intergenic | 6.1527E-11 | 0.48396 | 1.783E-13 | 0.018447 | 0.0057009 | 0.023375 | 0.0028207 | 0.0081448 | 0.0031742 | 0.427953 | 0.470961 | 0.419993 |
| 20 | 20_71532552_C_T | 71532552 | T | C | CEP72(dist=3417),SLC9A3(dist=7478 | intergenic | 7.4216E-11 | 0.57175 | 7.679E-13 | 0.018235 | 0.0047153 | 0.022469 | 0.0028003 | 0.0083387 | 0.0031351 | 0.462048 | 0.498999 | 0.455208 |
| 20 | 20_71532825_T_C | 71532825 | C | T | CEP72(dist=3690),SLC9A3(dist=7451 | intergenic | 7.5268E-10 | 0.63524 | 1.189E-11 | 0.01726 | 0.0039948 | 0.021267 | 0.0028043 | 0.0084215 | 0.003136 | 0.461891 | 0.499332 | 0.454961 |
| 20 | 20_71532950_C_A | 71532950 | A | C | CEP72(dist=3815),SLC9A3(dist=7438 | intergenic | 5.0057E-09 | 0.79052 | 2.981E-11 | 0.016393 | 0.0022254 | 0.020856 | 0.0028037 | 0.0083776 | 0.0031375 | 0.461787 | 0.499332 | 0.45459 |
| 20 | 20_71533058_C_T | 71533058 | T | C | CEP72(dist=3923),SLC9A3(dist=7427 | intergenic | 1.3403E-10 | 0.26249 | 4.787E-12 | 0.018197 | 0.0092212 | 0.022031 | 0.0028333 | 0.0082293 | 0.0031875 | 0.425399 | 0.463618 | 0.418324 |
| 20 | 20_71533193_A_G | 71533193 | G | A | CEP72(dist=4058),SLC9A3(dist=7414 | intergenic | 7.8356E-11 | 0.30971 | 4.539E-12 | 0.018285 | 0.0085527 | 0.021768 | 0.0028115 | 0.0084194 | 0.003146 | 0.462621 | 0.49733 | 0.456197 |
| 20 | 20_71533606_C_T | 71533606 | T | C | CEP72(dist=4471),SLC9A3(dist=7373 | intergenic | 2.8891E-10 | 0.62059 | 3.02E-12 | 0.017686 | 0.0041172 | 0.021924 | 0.0028052 | 0.0083173 | 0.0031425 | 0.462725 | 0.498331 | 0.455517 |
| 20 | 20_71534423_G_A | 71534423 | A | G | CEP72(dist=5288),SLC9A3(dist=7291 | intergenic | 1.3693E-10 | 0.43332 | 1.273E-12 | 0.018002 | 0.0065543 | 0.022288 | 0.0028043 | 0.0083652 | 0.0031404 | 0.460953 | 0.497997 | 0.454096 |
| 20 | 20_71534440_T_C | 71534440 | C | T | CEP72(dist=5305),SLC9A3(dist=7289 | intergenic | 2.267E-10 | 0.48418 | 2.098E-12 | 0.017769 | 0.0058486 | 0.022051 | 0.0028018 | 0.00836 | 0.0031377 | 0.460901 | 0.498999 | 0.453849 |
| 20 | 20_71534877_A_G | 71534877 | G | A | CEP72(dist=5742),SLC9A3(dist=7246 | intergenic | 1.1758E-10 | 0.40155 | 4.198E-13 | 0.018256 | 0.0069225 | 0.02309 | 0.0028337 | 0.0082523 | 0.0031853 | 0.429934 | 0.468291 | 0.422835 |
| 20 | 20_71534917_G_C | 71534917 | C | G | CEP72(dist=5782),SLC9A3(dist=7242 | intergenic | 7.9257E-10 | 0.66177 | 7.038E-13 | 0.017194 | 0.003594 | 0.022528 | 0.0027974 | 0.0082156 | 0.0031382 | 0.467313 | 0.488985 | 0.459224 |
| 20 | 20_71535521_G_A | 71535521 | A | G | CEP72(dist=6386),SLC9A3(dist=7181 | intergenic | 2.1779E-10 | 0.40394 | 1.5E-12 | 0.017627 | 0.0069515 | 0.021967 | 0.0027767 | 0.008329 | 0.0031051 | 0.469138 | 0.491656 | 0.461881 |
| 20 | 20_71535527_A_G | 71535527 | G | A | CEP72(dist=6392),SLC9A3(dist=7181 | intergenic | 1.9865E-10 | 0.45816 | 1.215E-12 | 0.017663 | 0.0061837 | 0.02205 | 0.0027761 | 0.008335 | 0.003104 | 0.469033 | 0.491322 | 0.461695 |
| 20 | 20_71535532_G_A | 71535532 | A | G | CEP72(dist=6397),SLC9A3(dist=7180 | intergenic | 2.9336E-10 | 0.45921 | 1.782E-12 | 0.017498 | 0.006173 | 0.021886 | 0.0027765 | 0.0083403 | 0.0031041 | 0.469294 | 0.491656 | 0.462066 |
| 20 | 20_71535561_A_G | 71535561 | G | A | CEP72(dist=6426),SLC9A3(dist=7177 | intergenic | 5.1287E-10 | 0.2817 | 4.334E-12 | 0.017333 | 0.0089616 | 0.021614 | 0.0027889 | 0.0083247 | 0.0031207 | 0.466323 | 0.487984 | 0.457865 |
| 20 | 20_71535753_C_T | 71535753 | T | C | CEP72(dist=6618),SLC9A3(dist=7158 | intergenic | 1.2903E-11 | 0.39188 | 5.375E-14 | 0.019024 | 0.0071628 | 0.02366 | 0.0028102 | 0.0083657 | 0.0031452 | 0.462986 | 0.49466 | 0.455146 |
| 20 | 20_71536051_T_G | 71536051 | G | T | CEP72(dist=6916),SLC9A3(dist=7128 | intergenic | 3.5183E-11 | 0.29899 | 3.183E-13 | 0.018841 | 0.0086953 | 0.023253 | 0.0028447 | 0.0083721 | 0.0031913 | 0.425868 | 0.462283 | 0.419128 |
| 20 | 20_71536513_T_C | 71536513 | C | T | CEP72(dist=7378),SLC9A3(dist=7082 | intergenic | 8.0391E-12 | 0.27464 | 4.92E-14 | 0.019317 | 0.0089324 | 0.023944 | 0.002825 | 0.0081765 | 0.0031782 | 0.425242 | 0.466622 | 0.417583 |
| 20 | 20_71536579_A_G | 71536579 | G | A | CEP72(dist=7444),SLC9A3(dist=7075 | intergenic | 1.0068E-11 | 0.33184 | 1.034E-13 | 0.019223 | 0.0079623 | 0.023605 | 0.0028246 | 0.008205 | 0.0031742 | 0.426441 | 0.465621 | 0.419189 |
| 20 | 20_71536716_A_T | 71536716 | T | A | CEP72(dist=7581),SLC9A3(dist=7062 | intergenic | 7.1765E-11 | 0.55987 | 2.632E-13 | 0.018292 | 0.0048713 | 0.022968 | 0.0028069 | 0.008355 | 0.0031412 | 0.461683 | 0.498665 | 0.454343 |
| 20 | 20_71536874_T_C | 71536874 | C | T | CEP72(dist=7739),SLC9A3(dist=7046 | intergenic | 1.8408E-10 | 0.31168 | 2.184E-12 | 0.017916 | 0.0082966 | 0.022192 | 0.0028108 | 0.0082007 | 0.0031602 | 0.431707 | 0.467623 | 0.425059 |
| 20 | 20_71536996_C_T | 71536996 | T | C | CEP72(dist=7861),SLC9A3(dist=7034 | intergenic | 3.1086E-12 | 0.23851 | 1.023E-13 | 0.019474 | 0.0098015 | 0.023257 | 0.0027929 | 0.0083154 | 0.0031268 | 0.461526 | 0.493658 | 0.453231 |
| 20 | 20_71537174_C_T | 71537174 | T | C | CEP72(dist=8039),SLC9A3(dist=7016 | intergenic | 1.2597E-10 | 0.32412 | 8.843E-13 | 0.01805 | 0.0082963 | 0.022432 | 0.0028063 | 0.0084139 | 0.0031385 | 0.462465 | 0.498331 | 0.455208 |
| 20 | 20_71537303_C_T | 71537303 | T | C | CEP72(dist=8168),SLC9A3(dist=7003 | intergenic | 2.7486E-09 | 0.6621 | 1.535E-11 | 0.016655 | 0.0036283 | 0.021168 | 0.002801 | 0.0083025 | 0.0031386 | 0.461631 | 0.493992 | 0.453417 |
| 20 | 20_71537401_T_G | 71537401 | G | T | CEP72(dist=8266),SLC9A3(dist=6993 | intergenic | 1.4775E-10 | 0.3806 | 9.946E-13 | 0.018151 | 0.0072574 | 0.022687 | 0.0028328 | 0.0082773 | 0.0031813 | 0.424721 | 0.463952 | 0.41746 |
| 20 | 20_71538042_C_T | 71538042 | T | C | CEP72(dist=8907),SLC9A3(dist=6929 | intergenic | 7.3772E-11 | 0.48991 | 1.121E-13 | 0.01838 | 0.005687 | 0.023553 | 0.0028221 | 0.0082367 | 0.0031717 | 0.423105 | 0.459279 | 0.416409 |
| 20 | 20_71538102_C_T | 71538102 | T | C | CEP72(dist=8967),SLC9A3(dist=6923 | intergenic | 2.9472E-11 | 0.36742 | 1.049E-13 | 0.01879 | 0.0074909 | 0.023575 | 0.002826 | 0.008311 | 0.003171 | 0.424043 | 0.460948 | 0.417212 |
| 20 | 20_71538164_C_T | 71538164 | T | C | CEP72(dist=9029),SLC9A3(dist=6917 | intergenic | 2.9462E-11 | 0.37125 | 1.581E-13 | 0.018829 | 0.0074665 | 0.023447 | 0.0028318 | 0.0083505 | 0.003177 | 0.424252 | 0.458278 | 0.417954 |
| 20 | 20_71538204_G_A | 71538204 | A | G | CEP72(dist=9069),SLC9A3(dist=6913 | intergenic | 4.8758E-12 | 0.19704 | 7.638E-14 | 0.019561 | 0.010799 | 0.023734 | 0.0028312 | 0.0083711 | 0.0031745 | 0.424408 | 0.455941 | 0.418572 |
| 20 | 20_71538219_C_T | 71538219 | T | C | CEP72(dist=9084),SLC9A3(dist=6911 | intergenic | 6.0975E-12 | 0.37554 | 9.813E-14 | 0.019281 | 0.0075066 | 0.023305 | 0.0028035 | 0.0084712 | 0.0031309 | 0.460953 | 0.495661 | 0.454529 |
| 20 | 20_71538537_G_A | 71538537 | A | G | CEP72(dist=9402),SLC9A3(dist=6880 | intergenic | 1.6229E-10 | 0.69085 | 8.192E-13 | 0.017891 | 0.0033102 | 0.02242 | 0.0027984 | 0.0083232 | 0.0031322 | 0.462673 | 0.496662 | 0.455146 |
| 20 | 20_71538924_C_G | 71538924 | G | C | CEP72(dist=9789),SLC9A3(dist=6841 | intergenic | 2.8117E-11 | 0.32026 | 7.292E-14 | 0.018791 | 0.0081729 | 0.023739 | 0.0028232 | 0.0082227 | 0.0031726 | 0.423783 | 0.462283 | 0.416656 |
| 20 | 20_71538970_A_T | 71538970 | T | A | CEP72(dist=9835),SLC9A3(dist=6836 | intergenic | 1.1886E-11 | 0.37788 | 2.798E-13 | 0.019126 | 0.0072685 | 0.024085 | 0.0028203 | 0.0082426 | 0.0031661 | 0.423626 | 0.462951 | 0.416347 |
| 20 | 20_71539055_G_A | 71539055 | A | G | CEP72(dist=9920),SLC9A3(dist=6828 | intergenic | 4.634E-11 | 0.46642 | 1.377E-13 | 0.018524 | 0.0060588 | 0.023336 | 0.0028142 | 0.0083188 | 0.0031542 | 0.423939 | 0.458278 | 0.417583 |
| 20 | 20_71539093_C_T | 71539093 | T | C | CEP72(dist=9958),SLC9A3(dist=6824 | intergenic | 4.9266E-11 | 0.46628 | 2.049E-13 | 0.018455 | 0.0060424 | 0.023123 | 0.0028077 | 0.0082938 | 0.0031479 | 0.423783 | 0.459613 | 0.417151 |
| 20 | 20_71539135_T_C | 71539135 | C | T | CEP72(dist=10000),SLC9A3(dist=682 | intergenic | 6.1215E-11 | 0.4865 | 1.918E-13 | 0.018364 | 0.0057199 | 0.023182 | 0.0028076 | 0.0082196 | 0.0031521 | 0.425242 | 0.464286 | 0.418016 |
| 20 | 20_71539195_G_A | 71539195 | A | G | CEP72(dist=10060),SLC9A3(dist=681 | intergenic | 4.1417E-11 | 0.67568 | 1.461E-13 | 0.018364 | 0.0034488 | 0.023042 | 0.0027829 | 0.0082435 | 0.0031177 | 0.46356 | 0.486649 | 0.454343 |
| 20 | 20_71539969_C_A | 71539969 | A | C | CEP72(dist=10834),SLC9A3(dist=673 | intergenic | 1.0526E-12 | 0.45767 | 1.348E-14 | 0.019854 | 0.006221 | 0.023999 | 0.0027871 | 0.0083762 | 0.0031163 | 0.462621 | 0.5 | 0.455702 |
| 20 | 20_71540077_C_T | 71540077 | T | C | CEP72(dist=10942),SLC9A3(dist=672 | intergenic | 5.7504E-12 | 0.45261 | 1.01E-13 | 0.018893 | 0.0062153 | 0.022803 | 0.0027438 | 0.0082752 | 0.003065 | 0.467887 | 0.492991 | 0.460645 |
| 20 | 20_71540108_T_C | 71540108 | C | T | CEP72(dist=10973),SLC9A3(dist=672 | intergenic | 1.5259E-11 | 0.57306 | 3.98E-13 | 0.01846 | 0.004697 | 0.022146 | 0.0027367 | 0.0083347 | 0.003052 | 0.470128 | 0.493324 | 0.463363 |
| 20 | 20_71540117_T_C | 71540117 | C | T | CEP72(dist=10982),SLC9A3(dist=672 | intergenic | 1.3773E-11 | 0.52393 | 3.926E-13 | 0.018479 | 0.0053046 | 0.022126 | 0.0027334 | 0.0083236 | 0.0030485 | 0.470337 | 0.493324 | 0.463611 |
| 20 | 20_71540127_T_C | 71540127 | C | T | CEP72(dist=10992),SLC9A3(dist=672 | intergenic | 1.8681E-11 | 0.55849 | 3.474E-13 | 0.018334 | 0.0048705 | 0.022138 | 0.0027299 | 0.0083244 | 0.0030432 | 0.471484 | 0.491989 | 0.464723 |
| 20 | 20_71540138_C_T | 71540138 | T | C | CEP72(dist=11003),SLC9A3(dist=671 | intergenic | 2.2201E-11 | 0.57665 | 4.573E-13 | 0.018266 | 0.0046472 | 0.022027 | 0.00273 | 0.0083242 | 0.0030435 | 0.470806 | 0.492657 | 0.464043 |
| 20 | 20_71540171_T_C | 71540171 | C | T | CEP72(dist=11036),SLC9A3(dist=671 | intergenic | 5.2355E-11 | 0.43898 | 1.686E-12 | 0.01796 | 0.0064606 | 0.021525 | 0.0027361 | 0.008348 | 0.0030496 | 0.474664 | 0.487984 | 0.46775 |
| 20 | 20_71540174_C_A | 71540174 | A | C | CEP72(dist=11039),SLC9A3(dist=671 | intergenic | 4.7637E-11 | 0.43937 | 1.544E-12 | 0.017997 | 0.0064512 | 0.021564 | 0.0027359 | 0.0083428 | 0.0030499 | 0.474768 | 0.48765 | 0.467812 |
| 20 | 20_71540744_C_T | 71540744 | T | C | CEP72(dist=11609),SLC9A3(dist=665 | intergenic | 7.0261E-11 | 0.59093 | 1.848E-13 | 0.018235 | 0.0044801 | 0.023024 | 0.0027967 | 0.0083354 | 0.0031285 | 0.462517 | 0.491989 | 0.454096 |
| 20 | 20_71540829_A_G | 71540829 | G | A | CEP72(dist=11694),SLC9A3(dist=665 | intergenic | 1.7659E-10 | 0.59253 | 8.528E-13 | 0.017806 | 0.004492 | 0.022297 | 0.0027907 | 0.0083935 | 0.0031174 | 0.460171 | 0.499666 | 0.452737 |
| 20 | 20_71540855_C_T | 71540855 | T | C | CEP72(dist=11720),SLC9A3(dist=664 | intergenic | 7.6523E-10 | 0.32407 | 1.283E-11 | 0.017051 | 0.0080516 | 0.021044 | 0.0027716 | 0.0081649 | 0.0031082 | 0.463664 | 0.493324 | 0.455702 |
| 20 | 20_71540861_C_T | 71540861 | T | C | CEP72(dist=11726),SLC9A3(dist=664 | intergenic | 6.4371E-10 | 0.29791 | 1.011E-11 | 0.017135 | 0.0085097 | 0.02116 | 0.0027729 | 0.008175 | 0.0031094 | 0.464029 | 0.492657 | 0.456011 |

| | | | | | | | | | | | | | | | | |
|---|---|---|---|---|---|---|---|---|---|---|---|---|---|---|---|---|
| 20 | 20_71540911_C_T | 71540911 | T | C | CEP72(dist=11776),SLC9A3(dist=664 | intergenic | 1.6922E-10 | 0.30936 | 1.401E-12 | 0.017798 | 0.0083788 | 0.022102 | 0.0027866 | 0.0082423 | 0.00312 | 0.46309 | 0.492991 | 0.454961 |
| 20 | 20_71541032_T_C | 71541032 | C | T | CEP72(dist=11897),SLC9A3(dist=663 | intergenic | 5.6511E-10 | 0.64242 | 2.837E-12 | 0.017409 | 0.0039077 | 0.021923 | 0.0028079 | 0.008416 | 0.0031383 | 0.462048 | 0.498331 | 0.454714 |
| 20 | 20_71541249_T_C | 71541249 | C | T | CEP72(dist=12114),SLC9A3(dist=660 | intergenic | 4.0766E-10 | 0.38249 | 2.399E-12 | 0.017529 | 0.0073194 | 0.021996 | 0.0028042 | 0.0083811 | 0.0031381 | 0.462152 | 0.499666 | 0.455208 |
| 20 | 20_71541467_C_T | 71541467 | T | C | CEP72(dist=12332),SLC9A3(dist=658 | intergenic | 6.3971E-11 | 0.29334 | 7.061E-13 | 0.018522 | 0.0087571 | 0.022828 | 0.0028346 | 0.0083335 | 0.0031802 | 0.423053 | 0.46028 | 0.416162 |
| 20 | 20_71541493_G_A | 71541493 | A | G | CEP72(dist=12358),SLC9A3(dist=658 | intergenic | 1.805E-10 | 0.48972 | 1.041E-12 | 0.017805 | 0.0057304 | 0.022272 | 0.002792 | 0.0082958 | 0.0031258 | 0.46038 | 0.495661 | 0.452243 |
| 20 | 20_71541548_C_T | 71541548 | T | C | CEP72(dist=12413),SLC9A3(dist=657 | intergenic | 6.0808E-10 | 0.54315 | 2.749E-12 | 0.017395 | 0.0051548 | 0.021937 | 0.0028109 | 0.0084774 | 0.0031384 | 0.459285 | 0.5 | 0.451748 |
| 20 | 20_71541794_C_G | 71541794 | G | C | CEP72(dist=12659),SLC9A3(dist=655 | intergenic | 1.6207E-10 | 0.54509 | 5.18E-13 | 0.017827 | 0.0050073 | 0.022564 | 0.0027883 | 0.0082746 | 0.003125 | 0.457356 | 0.496328 | 0.450142 |
| 20 | 20_71541883_T_G | 71541883 | G | T | CEP72(dist=12748),SLC9A3(dist=654 | intergenic | 3.0733E-10 | 0.3793 | 2.917E-12 | 0.017562 | 0.0072829 | 0.021828 | 0.0027898 | 0.0082837 | 0.0031264 | 0.457564 | 0.499666 | 0.449771 |
| 20 | 20_71541911_G_A | 71541911 | A | G | CEP72(dist=12776),SLC9A3(dist=654 | intergenic | 4.7382E-10 | 0.40631 | 4.111E-12 | 0.017569 | 0.0068017 | 0.022006 | 0.0028212 | 0.0081907 | 0.003174 | 0.420029 | 0.45494 | 0.413567 |
| 20 | 20_71542157_T_C | 71542157 | C | T | CEP72(dist=13022),SLC9A3(dist=651 | intergenic | 9.8308E-11 | 0.25543 | 9.292E-13 | 0.01836 | 0.0094096 | 0.022774 | 0.0028379 | 0.0082739 | 0.0031894 | 0.422896 | 0.458278 | 0.416347 |
| 20 | 20_71544917_C_A | 71544917 | A | C | CEP72(dist=15782),SLC9A3(dist=624 | intergenic | 3.7095E-11 | 0.39763 | 4.37E-13 | 0.018629 | 0.00709 | 0.022836 | 0.0028161 | 0.0083821 | 0.0031526 | 0.46627 | 0.494993 | 0.4591 |
| 20 | 20_71547230_A_G | 71547230 | G | A | CEP72(dist=18095),SLC9A3(dist=601 | intergenic | 8.8118E-11 | 0.36694 | 9.28E-13 | 0.018425 | 0.007442 | 0.022831 | 0.0028407 | 0.0082486 | 0.0031973 | 0.427067 | 0.462617 | 0.420487 |
| 20 | 20_71548980_T_C | 71548980 | C | T | CEP72(dist=19845),SLC9A3(dist=583 | intergenic | 3.7486E-10 | 0.23435 | 1.722E-11 | 0.017613 | 0.009869 | 0.021236 | 0.0028116 | 0.0082986 | 0.0031564 | 0.464654 | 0.495327 | 0.457247 |
| 20 | 20_71549790_A_G | 71549790 | G | A | CEP72(dist=20655),SLC9A3(dist=575 | intergenic | 4.1892E-11 | 0.25461 | 6.699E-13 | 0.01879 | 0.0094699 | 0.022997 | 0.0028481 | 0.0083126 | 0.0032004 | 0.426129 | 0.461949 | 0.419498 |
| 20 | 20_71549881_T_C | 71549881 | C | T | CEP72(dist=20746),SLC9A3(dist=574 | intergenic | 4.4785E-10 | 0.22658 | 3.936E-12 | 0.017661 | 0.0099092 | 0.02213 | 0.0028319 | 0.0081948 | 0.0031891 | 0.426598 | 0.462617 | 0.419931 |
| 20 | 20_71550464_G_A | 71550464 | A | G | CEP72(dist=21329),SLC9A3(dist=568 | intergenic | 2.0365E-10 | 0.284 | 2.912E-12 | 0.017916 | 0.0089917 | 0.022015 | 0.0028175 | 0.0083927 | 0.0031531 | 0.465749 | 0.492991 | 0.458112 |
| 20 | 20_71555394_G_A | 71555394 | A | G | CEP72(dist=26259),SLC9A3(dist=519 | intergenic | 4.9626E-10 | 0.59886 | 7.107E-12 | 0.017553 | 0.0044126 | 0.021665 | 0.0028219 | 0.0083884 | 0.0031603 | 0.465488 | 0.496662 | 0.458483 |
| 20 | 20_71557560_T_G | 71557560 | G | T | CEP72(dist=28425),SLC9A3(dist=497 | intergenic | 6.6517E-11 | 0.33796 | 5.75E-13 | 0.018541 | 0.0079657 | 0.022994 | 0.0028401 | 0.0083132 | 0.0031908 | 0.426077 | 0.457944 | 0.420178 |
| 20 | 20_71559096_A_G | 71559096 | G | A | CEP72(dist=29961),SLC9A3(dist=482 | intergenic | 5.3969E-10 | 0.3673 | 5.054E-12 | 0.017674 | 0.0075262 | 0.022068 | 0.0028473 | 0.008348 | 0.0031964 | 0.426129 | 0.461282 | 0.419622 |
| 20 | 20_71559463_C_T | 71559463 | T | C | CEP72(dist=30328),SLC9A3(dist=478 | intergenic | 2.8507E-10 | 0.34997 | 1.819E-12 | 0.017715 | 0.0077772 | 0.022193 | 0.0028088 | 0.008321 | 0.003149 | 0.463143 | 0.497997 | 0.45595 |
| 20 | 20_71566079_G_A | 71566079 | A | G | CEP72(dist=36944),SLC9A3(dist=412 | intergenic | 1.6628E-08 | 0.92694 | 5.273E-11 | 0.015847 | -0.000758 | 0.020687 | 0.0028078 | 0.0082625 | 0.0031521 | 0.472996 | 0.489987 | 0.466144 |
| 20 | 20_71569980_C_T | 71569980 | T | C | CEP72(dist=40845),SLC9A3(dist=373 | intergenic | 2.3578E-10 | 0.45984 | 2.593E-12 | 0.017867 | 0.0062571 | 0.022063 | 0.00282 | 0.0084657 | 0.0031527 | 0.465228 | 0.497997 | 0.459162 |
| 20 | 20_71572364_A_G | 71572364 | G | A | CEP72(dist=43229),SLC9A3(dist=349 | intergenic | 4.0942E-10 | 0.81788 | 3.125E-12 | 0.017627 | 0.0019445 | 0.021995 | 0.0028202 | 0.0084445 | 0.0031548 | 0.467626 | 0.497997 | 0.462004 |
| 20 | 20_71572812_G_A | 71572812 | A | G | CEP72(dist=43677),SLC9A3(dist=345 | intergenic | 1.8434E-09 | 0.73514 | 1.619E-11 | 0.016937 | 0.0028491 | 0.021258 | 0.0028176 | 0.0084219 | 0.0031555 | 0.470076 | 0.486983 | 0.462128 |
| 20 | 20_71579814_T_C | 71579814 | C | T | CEP72(dist=50679),SLC9A3(dist=275 | intergenic | 8.6704E-10 | 0.44348 | 6.656E-12 | 0.017419 | 0.0063428 | 0.021931 | 0.0028405 | 0.0082768 | 0.0031947 | 0.426702 | 0.459947 | 0.420549 |
| 20 | 20_71580815_A_C | 71580815 | C | A | CEP72(dist=51680),SLC9A3(dist=265 | intergenic | 3.1718E-10 | 0.24282 | 6.613E-12 | 0.017824 | 0.0094967 | 0.021939 | 0.0028337 | 0.008131 | 0.0031954 | 0.428892 | 0.463952 | 0.422402 |
| 20 | 20_71581802_C_T | 71581802 | T | C | CEP72(dist=52667),SLC9A3(dist=255 | intergenic | 2.412E-09 | 0.74954 | 4.987E-11 | 0.016795 | 0.0026624 | 0.020725 | 0.0028146 | 0.0083396 | 0.0031539 | 0.46382 | 0.496996 | 0.456567 |
| 20 | 20_71584662_C_T | 71584662 | T | C | CEP72(dist=55527),SLC9A3(dist=226 | intergenic | 1.0403E-10 | 0.32466 | 1.702E-12 | 0.018376 | 0.0081358 | 0.022588 | 0.0028442 | 0.0082602 | 0.0032008 | 0.428735 | 0.461949 | 0.422587 |
| 20 | 20_71587457_G_A | 71587457 | A | G | CEP72(dist=58322),SLC9A3(dist=198 | intergenic | 1.0647E-10 | 0.544 | 4.147E-13 | 0.018422 | 0.0051055 | 0.023202 | 0.0028529 | 0.0084142 | 0.0031999 | 0.426859 | 0.459613 | 0.420796 |
| 20 | 20_71590417_G_A | 71590417 | A | G | CEP72(dist=61282),SLC9A3(dist=169 | intergenic | 2.7126E-10 | 0.46161 | 7.733E-12 | 0.017798 | 0.0061035 | 0.021645 | 0.0028187 | 0.0082906 | 0.0031628 | 0.468356 | 0.49466 | 0.46151 |
| 20 | 20_71592997_G_C | 71592997 | C | G | CEP72(dist=63862),SLC9A3(dist=143 | intergenic | 2.6889E-10 | 0.36385 | 2.517E-12 | 0.017878 | 0.0073388 | 0.022377 | 0.0028307 | 0.0080819 | 0.0031956 | 0.428683 | 0.469292 | 0.421166 |
| 20 | 20_71593779_G_A | 71593779 | A | G | CEP72(dist=64644),SLC9A3(dist=135 | intergenic | 2.7798E-09 | 0.44912 | 2.561E-11 | 0.016656 | 0.006119 | 0.021043 | 0.002802 | 0.0080844 | 0.0031549 | 0.46674 | 0.486983 | 0.458174 |
| 20 | 20_71595604_G_A | 71595604 | A | G | CEP72(dist=66469),SLC9A3(dist=117 | intergenic | 6.3619E-10 | 0.35647 | 2.251E-11 | 0.017387 | 0.007544 | 0.021137 | 0.0028129 | 0.0081813 | 0.0031601 | 0.468825 | 0.484646 | 0.460213 |
| 20 | 20_71596825_T_C | 71596825 | C | T | CEP72(dist=67690),SLC9A3(dist=105 | intergenic | 1.2241E-10 | 0.27827 | 2.557E-12 | 0.018266 | 0.0088904 | 0.022358 | 0.002838 | 0.0081999 | 0.003194 | 0.428318 | 0.465287 | 0.421475 |
| 20 | 20_71597174_C_G | 71597174 | G | C | CEP72(dist=68039),SLC9A3(dist=101 | intergenic | 7.0475E-11 | 0.23938 | 1.084E-12 | 0.018528 | 0.0096199 | 0.022812 | 0.0028419 | 0.0081765 | 0.0032042 | 0.431342 | 0.468959 | 0.424379 |
| 20 | 20_71597678_T_C | 71597678 | C | T | CEP72(dist=68543),SLC9A3(dist=965 | intergenic | 3.4584E-10 | 0.40369 | 5.238E-12 | 0.017743 | 0.0069963 | 0.021845 | 0.0028269 | 0.0083783 | 0.0031664 | 0.464915 | 0.491322 | 0.456815 |
| 20 | 20_71598070_C_T | 71598070 | T | C | CEP72(dist=68935),SLC9A3(dist=926 | intergenic | 6.6053E-12 | 0.25745 | 5.083E-13 | 0.019563 | 0.0094225 | 0.023133 | 0.0028493 | 0.0083205 | 0.0032026 | 0.427901 | 0.460948 | 0.421784 |
| 20 | 20_71599866_G_T | 71599866 | T | G | CEP72(dist=70731),SLC9A3(dist=747 | intergenic | 2.6871E-10 | 0.62384 | 2.826E-12 | 0.017849 | 0.0041277 | 0.022111 | 0.0028261 | 0.0084167 | 0.003165 | 0.465436 | 0.494993 | 0.458112 |
| 20 | 20_71600702_G_A | 71600702 | A | G | CEP72(dist=71567),SLC9A3(dist=663 | intergenic | 1.1579E-11 | 0.16183 | 3.923E-13 | 0.019439 | 0.011536 | 0.023377 | 0.0028648 | 0.0082458 | 0.0032208 | 0.437129 | 0.491322 | 0.427097 |
| 20 | 20_71604615_T_C | 71604615 | C | T | CEP72(dist=75480),SLC9A3(dist=272 | intergenic | 1.8639E-10 | 0.27625 | 1.331E-12 | 0.018094 | 0.0090044 | 0.022645 | 0.0028395 | 0.0082702 | 0.0031934 | 0.427693 | 0.461949 | 0.421352 |
| 20 | 20_71604672_G_T | 71604672 | T | G | CEP72(dist=75537),SLC9A3(dist=266 | intergenic | 1.913E-09 | 0.51558 | 9.797E-12 | 0.016813 | 0.0053389 | 0.021408 | 0.0027997 | 0.0082115 | 0.0031439 | 0.46747 | 0.496996 | 0.460892 |
| 20 | 20_71607420_G_A | 71607420 | A | G | SLC9A3(XM_005221659.4:c.-32G>A) | UTR5 | 5.4821E-10 | 0.33448 | 5.722E-12 | 0.017463 | 0.0079285 | 0.02177 | 0.0028145 | 0.008215 | 0.0031612 | 0.462778 | 0.499332 | 0.455764 |
| 20 | 20_71608437_T_C | 71608437 | C | T | SLC9A3 | intronic | 6.0488E-11 | 0.10964 | 2.572E-12 | 0.018538 | 0.013042 | 0.02235 | 0.0028335 | 0.0081524 | 0.0031932 | 0.428944 | 0.461949 | 0.422835 |
| 20 | 20_71608962_G_C | 71608962 | C | G | SLC9A3 | intronic | 4.1752E-10 | 0.3524 | 3.473E-12 | 0.017589 | 0.0076643 | 0.021995 | 0.0028155 | 0.0082417 | 0.0031615 | 0.462725 | 0.496328 | 0.456506 |
| 20 | 20_71611019_A_C | 71611019 | C | A | SLC9A3 | intronic | 5.2486E-11 | 0.21465 | 1E-12 | 0.018599 | 0.010195 | 0.022739 | 0.0028337 | 0.0082157 | 0.003189 | 0.4291 | 0.46028 | 0.423329 |
| 20 | 20_71611774_C_G | 71611774 | G | C | SLC9A3 | intronic | 1.2132E-09 | 0.32714 | 1.451E-11 | 0.017214 | 0.0080537 | 0.02151 | 0.0028319 | 0.0082189 | 0.0031855 | 0.427015 | 0.459613 | 0.420981 |
| 20 | 20_71611783_C_G | 71611783 | G | C | SLC9A3 | intronic | 3.0578E-09 | 0.54064 | 3.249E-11 | 0.016635 | 0.0050662 | 0.02088 | 0.002806 | 0.0082802 | 0.0031471 | 0.462204 | 0.495327 | 0.456073 |
| 20 | 20_71612096_C_T | 71612096 | T | C | SLC9A3 | intronic | 1.56E-09 | 0.41256 | 1.926E-11 | 0.016949 | 0.0067662 | 0.021139 | 0.0028071 | 0.0082576 | 0.0031496 | 0.461787 | 0.498665 | 0.454961 |
| 20 | 20_71612335_T_C | 71612335 | C | T | SLC9A3 | intronic | 3.6785E-09 | 0.29126 | 3.306E-11 | 0.016772 | 0.008774 | 0.021193 | 0.0028436 | 0.0083138 | 0.0031955 | 0.422167 | 0.455607 | 0.415977 |
| 20 | 20_71612470_T_C | 71612470 | C | T | SLC9A3 | intronic | 1.2698E-09 | 0.37429 | 1.798E-11 | 0.017067 | 0.0074949 | 0.021128 | 0.0028111 | 0.0084359 | 0.0031433 | 0.458398 | 0.487984 | 0.452922 |
| 20 | 20_71613508_T_C | 71613508 | C | T | SLC9A3 | intronic | 1.0107E-09 | 0.3957 | 1.325E-11 | 0.017124 | 0.0069548 | 0.021291 | 0.0028036 | 0.0081886 | 0.0031468 | 0.467939 | 0.488652 | 0.459904 |
| 20 | 20_71614585_A_C | 71614585 | C | A | SLC9A3 | intronic | 5.6949E-10 | 0.35484 | 2.348E-12 | 0.017563 | 0.0075314 | 0.022387 | 0.0028333 | 0.0081399 | 0.0031927 | 0.427849 | 0.467623 | 0.420487 |
| 20 | 20_71615040_T_C | 71615040 | C | T | SLC9A3 | intronic | 7.4153E-10 | 0.37316 | 3.28E-12 | 0.017474 | 0.0073102 | 0.02225 | 0.002838 | 0.0082085 | 0.0031945 | 0.424773 | 0.461949 | 0.417892 |
| 20 | 20_71616076_C_T | 71616076 | T | C | SLC9A3 | intronic | 8.6001E-10 | 0.52926 | 4.704E-12 | 0.01739 | 0.0051097 | 0.022111 | 0.0028353 | 0.0081217 | 0.0031979 | 0.427536 | 0.46996 | 0.419684 |
| 20 | 20_71620561_T_C | 71620561 | C | T | SLC9A3 | intronic | 1.874E-11 | 0.51664 | 1.593E-13 | 0.019073 | 0.0053318 | 0.023588 | 0.0028402 | 0.0082213 | 0.0031966 | 0.427119 | 0.465955 | 0.419931 |
| 20 | 20_71622142_C_T | 71622142 | T | C | SLC9A3 | intronic | 6.8893E-11 | 0.39902 | 7.356E-13 | 0.018535 | 0.0069644 | 0.022916 | 0.0028415 | 0.0082578 | 0.0031949 | 0.42665 | 0.463284 | 0.419869 |

**Supplementary Table 9. Conditional association analysis of genome-wide significant telomere length loci on chromosomes 13, 19, and 20, related to Figure S12.**

| Locus | Conditioning SNP | Marginal P value | No. Significant variants before conditioning | No. Significant variants after conditioning | Top residual SNP | Top residual P value |
|---|---|---|---|---|---|---|
| Chr13 | 13_53727328_G_A | 2.0267E-10 | 185 | 0 | 13_54655792_G_T | 0.0012864 |
| Chr19 | 19_27855458_G_A | 4.1573E-10 | 35 | 0 | 19_28491238_A_G | 0.021219 |
| Chr20 | 20_71539969_C_A | 1.0526E-12 | 725 | 0 | 20_71862779_G_A | 0.022121 |
| Conditional association analyses were performed by including the genotype dosage of the lead SNP at each locus as an additional covariate. Significant variants were defined as variants with P < 5 × 10-8 within ±1 Mb of the conditioning SNP. The top residual SNP denotes the most significant variant remaining within the same ±1 Mb region after conditioning. The marginal P value refers to the association P value of the conditioning SNP in the original GWAS. | | | | | | |

**Supplementary Table 10. Colocalization analysis between telomere length GWAS loci and cattle eQTL signals across tissues, related to Figures 4 and S13.**

| Tissue | Chr | GWAS lead SNP position | Gene ID | Gene symbol | No. of SNPs used | PP.H0 | PP.H1 | PP.H2 | PP.H3 | PP.H4 |
|---|---|---|---|---|---|---|---|---|---|---|
| Brain | chr20 | chr20:71539969 | ENSBTAG00000052982 | ENSBTAG00000052982 | 1437 | 4.22E-12 | 0.0000373 | 3.04E-08 | 0.26753 | 0.732433 |
| Whole blood | chr19 | chr19:27855458 | ENSBTAG00000007117 | CTC1 | 3526 | 1.16072E-10 | 1.64099E-06 | 5.47623E-06 | 0.076497644 | 0.923495239 |
| Whole blood | chr19 | chr19:27855458 | ENSBTAG00000017895 | CLEC10A | 710 | 0.031960417 | 0.062055264 | 0.005829846 | 0.010429673 | 0.8897248 |
| Whole blood | chr19 | chr19:27855458 | ENSBTAG00000013392 | PLD2 | 482 | 0.001277 | 0.00078 | 0.163504 | 0.099094 | 0.735345 |
| Leukocyte | chr13 | chr13:53727328 | ENSBTAG00000011638 | ENSBTAG00000011638 | 3539 | 1.78684E-10 | 6.01224E-06 | 7.20887E-06 | 0.241800953 | 0.758185826 |
| Immune system | chr13 | chr13:53727328 | ENSBTAG00000007497 | ARFRP1 | 3736 | 2.28694E-06 | 0.081248648 | 3.02342E-06 | 0.106601568 | 0.812144474 |
| Immune system | chr19 | chr19:27855458 | ENSBTAG00000003699 | ZFP3 | 358 | 0.08130789 | 0.044747466 | 0.046254754 | 0.024653079 | 0.803036811 |
| Immune system | chr19 | chr19:27855458 | ENSBTAG00000017895 | CLEC10A | 713 | 0.074865515 | 0.146316027 | 0.007836354 | 0.014558828 | 0.756423276 |
| Immune system | chr19 | chr19:27855458 | ENSBTAG00000013392 | PLD2 | 485 | 0.00086 | 0.000536 | 0.163872 | 0.101424 | 0.733309 |
| Immune system | chr19 | chr19:27855458 | ENSBTAG00000004910 | SLC25A11 | 447 | 0.036253 | 0.021369 | 0.144624 | 0.084535 | 0.713219 |
| Testis | chr19 | chr19:27855458 | ENSBTAG00000031509 | PFAS | 3366 | 1.23519E-11 | 1.74594E-07 | 1.35627E-06 | 0.018189087 | 0.981809382 |
| Testis | chr19 | chr19:27855458 | ENSBTAG00000007117 | CTC1 | 3373 | 2.96127E-12 | 4.1859E-08 | 3.46526E-06 | 0.048031223 | 0.95196527 |
| Testis | chr19 | chr19:27855458 | ENSBTAG00000031725 | TNFSF12 | 1736 | 1.04704E-05 | 0.147354599 | 7.2278E-06 | 0.10096829 | 0.751659413 |
| Lymph node | chr19 | chr19:27855458 | ENSBTAG00000031509 | PFAS | 3535 | 5.69775E-15 | 8.05524E-11 | 1.44392E-06 | 0.019432993 | 0.980565563 |
| Mammary gland | chr19 | chr19:27855458 | ENSBTAG00000031509 | PFAS | 3517 | 6.35384E-16 | 8.9828E-12 | 3.74448E-06 | 0.051989849 | 0.948006407 |
| Placenta | chr19 | chr19:27855458 | ENSBTAG00000031509 | PFAS | 2911 | 2.48385E-06 | 0.033885988 | 6.56702E-06 | 0.088713394 | 0.877391567 |
| Uterus | chr19 | chr19:27855458 | ENSBTAG00000031509 | PFAS | 3507 | 3.00898E-17 | 4.25401E-13 | 2.62445E-06 | 0.03613988 | 0.963857495 |
| Uterus | chr19 | chr19:27855458 | ENSBTAG00000003958 | TMEM220 | 426 | 2.07988E-10 | 3.13696E-11 | 0.207236622 | 0.030494059 | 0.762269319 |
| Adipose | chr19 | chr19:27855458 | ENSBTAG00000031509 | PFAS | 3308 | 1.75132E-23 | 2.47549E-19 | 5.9041E-07 | 0.007352814 | 0.992646596 |
| Adipose | chr19 | chr19:27855458 | ENSBTAG00000007117 | CTC1 | 3316 | 4.85496E-10 | 6.86262E-06 | 1.42142E-06 | 0.019111324 | 0.980880392 |
| Adipose | chr19 | chr19:27855458 | ENSBTAG00000052025 | EFNB3 | 2307 | 8.48296E-06 | 0.119588378 | 3.58148E-06 | 0.049659062 | 0.830740496 |
| Brain | chr19 | chr19:27855458 | ENSBTAG00000031509 | PFAS | 3241 | 3.31812E-16 | 4.68111E-12 | 8.35519E-07 | 0.010798081 | 0.989201083 |
| Epididymis | chr19 | chr19:27855458 | ENSBTAG00000031509 | PFAS | 2249 | 4.50185E-08 | 0.000636016 | 1.75061E-06 | 0.023756833 | 0.975605355 |
| Gut | chr19 | chr19:27855458 | ENSBTAG00000031509 | PFAS | 3538 | 2.31691E-10 | 3.27556E-06 | 1.68107E-06 | 0.022789123 | 0.97720592 |
| Gut | chr20 | chr20:71539969 | ENSBTAG00000053495 | ENSBTAG00000053495 | 1552 | 1.03411E-11 | 9.12975E-05 | 2.16834E-08 | 0.190625016 | 0.809283665 |
| Gut | chr20 | chr20:71539969 | ENSBTAG00000006971 | BRD9 | 1552 | 2.29933E-11 | 0.000202999 | 2.8357E-08 | 0.24960207 | 0.750194902 |
| Hypothalamus | chr19 | chr19:27855458 | ENSBTAG00000031509 | PFAS | 3318 | 1.40677E-16 | 1.98831E-12 | 1.22904E-06 | 0.016387454 | 0.983611317 |
| Jejunum | chr19 | chr19:27855458 | ENSBTAG00000031509 | PFAS | 3504 | 6.03199E-07 | 0.008527821 | 1.84194E-06 | 0.025074365 | 0.966395368 |
| Jejunum | chr19 | chr19:27855458 | ENSBTAG00000007117 | CTC1 | 3512 | 9.01571E-06 | 0.127463198 | 6.19937E-06 | 0.086860319 | 0.785661268 |
| Jejunum | chr19 | chr19:27855458 | ENSBTAG00000003958 | TMEM220 | 428 | 0.000485659 | 7.45357E-05 | 0.191889559 | 0.028671053 | 0.778879193 |
| Large intestine | chr19 | chr19:27855458 | ENSBTAG00000003958 | TMEM220 | 428 | 0.003370354 | 0.000517275 | 0.187307955 | 0.027966817 | 0.7808376 |
| Liver | chr13 | chr13:53727328 | ENSBTAG00000054447 | ENSBTAG00000054447 | 3584 | 9.44101E-48 | 3.19051E-43 | 2.65993E-06 | 0.088978918 | 0.911018422 |
| Liver | chr13 | chr13:53727328 | ENSBTAG00000031715 | PCMTD2 | 3952 | 6.09504E-12 | 2.07775E-07 | 3.96986E-06 | 0.134463934 | 0.865531888 |
| Liver | chr19 | chr19:27855458 | ENSBTAG00000007117 | CTC1 | 3510 | 2.52143E-22 | 3.5648E-18 | 1.15345E-06 | 0.01532275 | 0.984676097 |
| Liver | chr19 | chr19:27855458 | ENSBTAG00000031509 | PFAS | 3502 | 2.18783E-39 | 3.09311E-35 | 4.80122E-06 | 0.066945537 | 0.933049662 |
| Liver | chr19 | chr19:27855458 | ENSBTAG00000003958 | TMEM220 | 425 | 1.93155E-10 | 2.91397E-11 | 0.156211579 | 0.022745346 | 0.821043075 |
| Liver | chr20 | chr20:71539969 | ENSBTAG00000006971 | BRD9 | 1556 | 9.80548E-46 | 8.65694E-39 | 2.15115E-08 | 0.189106973 | 0.810893006 |
| Liver | chr20 | chr20:71539969 | ENSBTAG00000049710 | ENSBTAG00000049710 | 1556 | 5.93E-18 | 5.23E-11 | 2.86E-08 | 0.251779 | 0.748221 |
| Liver | chr20 | chr20:71539969 | ENSBTAG00000052982 | ENSBTAG00000052982 | 1556 | 2.12E-32 | 1.87E-25 | 3.24E-08 | 0.285444 | 0.714556 |
| Lung | chr19 | chr19:27855458 | ENSBTAG00000031509 | PFAS | 3535 | 2.71195E-27 | 3.83405E-23 | 8.82245E-07 | 0.011484331 | 0.988514787 |
| Lung | chr19 | chr19:27855458 | ENSBTAG00000003958 | TMEM220 | 433 | 3.91145E-22 | 5.99193E-23 | 0.211565004 | 0.031652752 | 0.756782244 |
| Muscle | chr19 | chr19:27855458 | ENSBTAG00000031509 | PFAS | 3027 | 9.18009E-39 | 1.29457E-34 | 1.27947E-06 | 0.017060117 | 0.982938604 |
| Muscle | chr19 | chr19:27855458 | ENSBTAG00000007117 | CTC1 | 3034 | 4.52046E-25 | 6.37498E-21 | 2.55524E-06 | 0.035070298 | 0.964927147 |
| Pituitary | chr19 | chr19:27855458 | ENSBTAG00000031509 | PFAS | 3245 | 1.76693E-12 | 2.49254E-08 | 1.15219E-06 | 0.015268755 | 0.984730068 |
| Pituitary | chr19 | chr19:27855458 | ENSBTAG00000025101 | ASGR2 | 737 | 1.63685E-05 | 0.200423115 | 3.105E-06 | 0.037256832 | 0.76230058 |
| Rumen | chr19 | chr19:27855458 | ENSBTAG00000031509 | PFAS | 3528 | 1.55372E-18 | 2.19659E-14 | 1.21432E-06 | 0.01618382 | 0.983814966 |
| Rumen | chr19 | chr19:27855458 | ENSBTAG00000007117 | CTC1 | 3536 | 4.74566E-13 | 6.70936E-09 | 1.46604E-06 | 0.019746415 | 0.980252112 |
| Small intestine | chr19 | chr19:27855458 | ENSBTAG00000031509 | PFAS | 3527 | 2.06858E-09 | 2.92446E-05 | 1.34348E-06 | 0.01801157 | 0.98195784 |

| | | | | | | | | | | |
|---|---|---|---|---|---|---|---|---|---|---|
| Subcutaneous adipose | chr19 | chr19:27855458 | ENSBTAG00000031509 | PFAS | 3307 | 5.64513E-15 | 7.97859E-11 | 6.6763E-07 | 0.008444454 | 0.991554878 |
| Subcutaneous adipose | chr19 | chr19:27855458 | ENSBTAG00000007117 | CTC1 | 3315 | 6.81843E-06 | 0.096370505 | 5.21946E-06 | 0.072940303 | 0.830677155 |
| Vas deferens | chr19 | chr19:27855458 | ENSBTAG00000031509 | PFAS | 2007 | 1.49626E-08 | 0.00021133 | 1.46862E-06 | 0.019762458 | 0.980024728 |
| Vas deferens | chr20 | chr20:71539969 | ENSBTAG00000012567 | TERT | 791 | 9.32324E-11 | 0.000275112 | 6.21719E-08 | 0.182640733 | 0.817084093 |
| Vas deferens | chr13 | chr13:53727328 | ENSBTAG00000019984 | GINS1 | 2444 | 0.0000003 | 0.007503 | 0.0000113 | 0.282219 | 0.710267 |
| Visceral fat | chr19 | chr19:27855458 | ENSBTAG00000031509 | PFAS | 3137 | 1.37185E-08 | 0.000192641 | 1.65317E-06 | 0.022236903 | 0.977568789 |
| Visceral fat | chr19 | chr19:27855458 | ENSBTAG00000007117 | CTC1 | 3145 | 1.25833E-06 | 0.017670334 | 3.18423E-06 | 0.043776487 | 0.938548737 |

**Supplementary Table 11. Colocalization analysis between telomere length GWAS loci and cattle sQTL signals across tissues, related to Figure 4.**

| Tissue | Chr | GWAS lead SNP position | Splicing event | Gene ID | Gene symbol | No. of SNPs used | PP.H0 | PP.H1 | PP.H2 | PP.H3 | PP.H4 |
|---|---|---|---|---|---|---|---|---|---|---|---|
| Whole blood | chr19 | chr19:27855458 | 19:27859828:27859948:clu_203187_NA | ENSBTAG00000007117 | CTC1 | 3526 | 2.53605E-16 | 3.58538E-12 | 1.42061E-06 | 0.019103186 | 0.980895393 |
| Whole blood | chr19 | chr19:27855458 | 19:27859443:27859651:clu_203186_NA | ENSBTAG00000007117 | CTC1 | 3526 | 3.31886E-06 | 0.046920953 | 9.52479E-06 | 0.133839164 | 0.819227039 |
| Whole blood | chr19 | chr19:27855458 | 19:27859555:27859651:clu_203186_NA | ENSBTAG00000007117 | CTC1 | 3526 | 8.26659E-06 | 0.116870361 | 8.59446E-06 | 0.120743335 | 0.762369443 |
| Whole blood | chr19 | chr19:27855458 | 19:26028211:26030550:clu_202278_NA | ENSBTAG00000020433 | NLRP1 | 268 | 0.069709199 | 0.036924179 | 0.10178876 | 0.053177961 | 0.738399901 |
| Whole blood | chr19 | chr19:27855458 | 19:27859196:27840746:clu_203104_NA | ENSBTAG00000001717 | AURKB | 3537 | 1.29845E-09 | 1.8359E-05 | 1.97101E-05 | 0.277961482 | 0.722000447 |
| Whole blood | chr19 | chr19:27855458 | 19:27866051:27873582:clu_203199_NA | ENSBTAG00000007117 | CTC1 | 3526 | 1.40976E-05 | 0.199306846 | 6.36536E-06 | 0.089279947 | 0.711392744 |
| Leukocyte | chr19 | chr19:27855458 | 19:27859828:27860076:clu_203187_NA | ENSBTAG00000007117 | CTC1 | 3533 | 8.93696E-06 | 0.126349556 | 7.71868E-06 | 0.108360502 | 0.765273287 |
| Immune system | chr13 | chr13:53727328 | 13:52522383:52522800:clu_146256_NA | ENSBTAG00000018812 | NOP56 | 1690 | 7.21285E-05 | 0.054524294 | 0.000131399 | 0.098482146 | 0.846790032 |
| Immune system | chr19 | chr19:27855458 | 19:27859828:27859948:clu_203187_NA | ENSBTAG00000007117 | CTC1 | 3537 | 6.36787E-22 | 9.00281E-18 | 1.32115E-06 | 0.017695967 | 0.982302711 |
| Immune system | chr19 | chr19:27855458 | 19:27866051:27873582:clu_203199_NA | ENSBTAG00000007117 | CTC1 | 3537 | 6.2298E-06 | 0.088076086 | 4.91986E-06 | 0.068713174 | 0.84319959 |
| Testis | chr13 | chr13:53727328 | 13:53604142:53659483:clu_146507_NA | ENSBTAG00000031715 | PCMTD2 | 3806 | 3.18187E-09 | 0.000105862 | 3.06479E-06 | 0.1010682 | 0.898822869 |
| Testis | chr19 | chr19:27855458 | 19:27870143:27873582:clu_203199_NA | ENSBTAG00000007117 | CTC1 | 3373 | 8.03026E-08 | 0.001135116 | 2.26927E-06 | 0.031109413 | 0.967753121 |
| Lymph node | chr19 | chr19:27855458 | 19:27859828:27860076:clu_203187_NA | ENSBTAG00000007117 | CTC1 | 3543 | 9.60576E-09 | 0.000135805 | 1.43253E-06 | 0.019272195 | 0.980590559 |
| Lymph node | chr19 | chr19:27855458 | 19:27859828:27859948:clu_203187_NA | ENSBTAG00000007117 | CTC1 | 3543 | 2.41011E-07 | 0.003407368 | 1.65272E-06 | 0.022391679 | 0.974199059 |
| Lymph node | chr19 | chr19:27855458 | 19:26097078:26099143:clu_202310_NA | ENSBTAG00000032083 | DHX33 | 297 | 0.043893735 | 0.02368765 | 0.092682244 | 0.049226299 | 0.790510072 |
| Mammary gland | chr19 | chr19:27855458 | 19:26097078:26099143:clu_202310_NA | ENSBTAG00000032083 | DHX33 | 298 | 6.87393E-05 | 3.71312E-05 | 0.084180865 | 0.044601208 | 0.871112057 |
| Uterus | chr19 | chr19:27855458 | 19:27859828:27859948:clu_203187_NA | ENSBTAG00000007117 | CTC1 | 3515 | 5.81617E-07 | 0.008222863 | 4.91572E-06 | 0.06857498 | 0.92319666 |
| Adipose | chr19 | chr19:27855458 | 19:27859828:27859948:clu_203187_NA | ENSBTAG00000007117 | CTC1 | 3316 | 4.90966E-07 | 0.006939943 | 1.41539E-06 | 0.019032942 | 0.974025208 |
| Brain | chr19 | chr19:27855458 | 19:27859828:27859948:clu_203187_NA | ENSBTAG00000007117 | CTC1 | 3248 | 1.7438E-06 | 0.024601893 | 2.58396E-06 | 0.035515293 | 0.939878486 |
| Gut | chr13 | chr13:53727328 | 13:54468251:54470730:clu_146853_NA | ENSBTAG00000001827 | GID8 | 2649 | 4.82514E-06 | 0.169386931 | 1.80337E-06 | 0.062539216 | 0.768067225 |
| Gut | chr20 | chr20:71539969 | 20:71828147:71829066:clu_323439_NA | ENSBTAG00000047780 | CCDC127 | 1546 | 7.13651E-10 | 0.006300506 | 1.9259E-08 | 0.169204481 | 0.824494993 |
| Gut | chr20 | chr20:71539969 | 20:71294782:71295331:clu_323148_NA | ENSBTAG00000006775 | SLC12A7 | 1552 | 1.33436E-15 | 1.17806E-08 | 3.01759E-08 | 0.265676682 | 0.734323276 |
| Liver | chr19 | chr19:27855458 | 19:27859828:27859948:clu_203187_NA | ENSBTAG00000007117 | CTC1 | 3510 | 1.32218E-06 | 0.018693021 | 2.54151E-06 | 0.034985547 | 0.946317568 |
| Lung | chr19 | chr19:27855458 | 19:27859828:27859948:clu_203187_NA | ENSBTAG00000007117 | CTC1 | 3543 | 2.30724E-08 | 0.000326193 | 1.64855E-06 | 0.022329672 | 0.977342463 |
| Milk | chr20 | chr20:71539969 | 20:71294782:71299240:clu_323148_NA | ENSBTAG00000006775 | SLC12A7 | 1553 | 2.70197E-16 | 2.38548E-09 | 1.31405E-08 | 0.115127745 | 0.88487224 |
| Milk | chr20 | chr20:71539969 | 20:71294782:71295331:clu_323148_NA | ENSBTAG00000006775 | SLC12A7 | 1553 | 1.51323E-14 | 1.33598E-07 | 1.48401E-08 | 0.130148192 | 0.86985166 |
| Muscle | chr13 | chr13:53727328 | 13:55064963:55065468:clu_146890_NA | ENSBTAG00000021558 | LSM14B | 1145 | 1.04292E-07 | 0.000246246 | 7.66919E-06 | 0.017125334 | 0.982620646 |
| Muscle | chr13 | chr13:53727328 | 13:53943480:53946243:clu_146525_NA | ENSBTAG00000021232 | TPD52L2 | 2989 | 1.21944E-05 | 0.216901763 | 3.85624E-06 | 0.067875486 | 0.7152067 |
| Muscle | chr13 | chr13:53727328 | 13:53955223:53955302:clu_146527_NA | ENSBTAG00000021232 | TPD52L2 | 2989 | 2.19655E-07 | 0.003906995 | 1.66265E-05 | 0.295033977 | 0.701042182 |
| Muscle | chr19 | chr19:27855458 | 19:27859828:27859948:clu_203187_NA | ENSBTAG00000007117 | CTC1 | 3034 | 8.10653E-06 | 0.114322368 | 2.79849E-06 | 0.03861862 | 0.847048107 |
| Rumen | chr13 | chr13:53727328 | 13:55064963:55065468:clu_146890_NA | ENSBTAG00000021558 | LSM14B | 1379 | 1.11567E-07 | 0.000379363 | 8.52239E-06 | 0.028007294 | 0.971604709 |
| Subcutaneous adipose | chr19 | chr19:27855458 | 19:26835125:26835429:clu_202366_NA | ENSBTAG00000017895 | CLEC10A | 661 | 0.009519017 | 0.016865738 | 0.045091246 | 0.079042918 | 0.849481081 |

**Supplementary Table 12. Significant gene-telomere length associations identified by transcriptome-wide association analysis across tissues, related to Figures 4 and S14.**

| Tissue | Gene ID | Gene symbol | Chr | Start | End | TWAS Z | TWAS P value | FDR |
|---|---|---|---|---|---|---|---|---|
| Adipose | ENSBTAG00000019984 | GINS1 | 13 | 53589805 | 53606654 | 5.787832777 | 7.13003E-09 | 9.7325E-06 |
| Adipose | ENSBTAG00000031509 | PFAS | 19 | 27879985 | 27895903 | -5.297918598 | 1.1713E-07 | 7.99413E-05 |
| Adipose | ENSBTAG00000031800 | PPDPF | 13 | 54136171 | 54137886 | 5.096788253 | 3.45464E-07 | 0.000157186 |
| Blood | ENSBTAG00000034435 | NKD2 | 20 | 71307541 | 71311025 | -6.997900895 | 2.59826E-12 | 1.23417E-08 |
| Blood | ENSBTAG00000012567 | TERT | 20 | 71145309 | 71162695 | -6.560335128 | 5.3687E-11 | 1.27507E-07 |
| Blood | ENSBTAG00000008606 | EXOC3 | 20 | 71649687 | 71674393 | -5.974046647 | 2.3144E-09 | 3.66446E-06 |
| Blood | ENSBTAG00000011638 | ENSBTAG00000011638 | 13 | 53538138 | 53538619 | 5.200556419 | 1.98693E-07 | 0.00017669 |
| Blood | ENSBTAG00000008291 | PROCR | 13 | 64444427 | 64449124 | 5.195052872 | 2.04662E-07 | 0.00017669 |
| Blood | ENSBTAG00000039520 | ENSBTAG00000039520 | 13 | 53431673 | 53469664 | 5.17890947 | 2.23187E-07 | 0.00017669 |
| Blood | ENSBTAG00000051215 | AGBL2 | 15 | 77451766 | 77481415 | 5.142345754 | 2.71329E-07 | 0.000184116 |
| Blood | ENSBTAG00000014046 | BPI | 13 | 67172370 | 67201998 | 4.811028439 | 1.50156E-06 | 0.000835378 |
| Blood | ENSBTAG00000052247 | TPPP | 20 | 71467623 | 71488177 | -4.800484576 | 1.58282E-06 | 0.000835378 |
| Blood | ENSBTAG00000001329 | CLPTM1L | 20 | 71117460 | 71129901 | 4.493784613 | 6.99684E-06 | 0.003323499 |
| Blood | ENSBTAG00000006143 | CREB3L1 | 15 | 76282608 | 76319986 | 4.413096307 | 1.01903E-05 | 0.004400341 |
| Blood | ENSBTAG00000047780 | CCDC127 | 20 | 71828109 | 71843714 | 4.320840538 | 1.55436E-05 | 0.005945955 |
| Blood | ENSBTAG00000054934 | VAMP2 | 19 | 27795436 | 27799326 | -4.310709745 | 1.62731E-05 | 0.005945955 |
| Blood | ENSBTAG00000014881 | PHF23 | 19 | 26948820 | 26953274 | 4.253628825 | 2.10334E-05 | 0.007136327 |
| Blood | ENSBTAG00000034077 | ASIP | 13 | 63662796 | 63668123 | 4.19037011 | 2.785E-05 | 0.008727465 |
| Blood | ENSBTAG00000050745 | ENSBTAG00000050745 | 4 | 105653052 | 105654500 | -4.178081926 | 2.93978E-05 | 0.008727465 |
| Blood | ENSBTAG00000011662 | SOX18 | 13 | 53823431 | 53825284 | 4.119047424 | 3.80442E-05 | 0.010629993 |
| Blood | ENSBTAG00000002191 | RRP12 | 26 | 18603839 | 18632724 | 4.06717293 | 4.75869E-05 | 0.012557662 |
| Blood | ENSBTAG00000013401 | ARHGEF40 | 10 | 26001808 | 26023736 | 4.011429228 | 6.03523E-05 | 0.014232249 |
| Blood | ENSBTAG00000004367 | COQ10A | 5 | 57065720 | 57069211 | 4.004927891 | 6.20364E-05 | 0.014232249 |
| Blood | ENSBTAG00000032055 | VPS39 | 10 | 37550085 | 37596971 | -4.001577716 | 6.29215E-05 | 0.014232249 |
| Blood | ENSBTAG00000052281 | ENSBTAG00000052281 | 10 | 25040667 | 25042311 | -3.966389585 | 7.29696E-05 | 0.015280065 |
| Blood | ENSBTAG00000012774 | RAB7B | 16 | 3728988 | 3757499 | -3.963084162 | 7.39877E-05 | 0.015280065 |
| Blood | ENSBTAG00000010778 | NDUFAF1 | 10 | 36820290 | 36837237 | 3.860304416 | 0.000113246 | 0.022413242 |
| Blood | ENSBTAG00000006448 | ERI2 | 25 | 18541026 | 18565241 | 3.848590918 | 0.000118799 | 0.022571849 |
| Blood | ENSBTAG00000016388 | SLCO4A1 | 13 | 54671725 | 54695783 | -3.836721644 | 0.000124688 | 0.022779477 |
| Blood | ENSBTAG00000013105 | SYT3 | 18 | 56713476 | 56727569 | 3.721373965 | 0.000198142 | 0.033521818 |
| Blood | ENSBTAG00000011553 | FAHD2A | 11 | 2153786 | 2164529 | 3.720973367 | 0.000198456 | 0.033521818 |
| Blood | ENSBTAG00000007397 | FOLH1 | 29 | 5885258 | 5958145 | 3.713194449 | 0.00020466 | 0.033521818 |
| Blood | ENSBTAG00000000528 | MANBAL | 13 | 66243119 | 66271549 | 3.638151534 | 0.000274602 | 0.042599477 |
| Blood | ENSBTAG00000051794 | ENSBTAG00000051794 | 7 | 18669526 | 18683341 | 3.634965687 | 0.000278018 | 0.042599477 |
| Blood | ENSBTAG00000003394 | RNF215 | 17 | 69378978 | 69382634 | 3.588898508 | 0.000332078 | 0.047872 |
| Blood | ENSBTAG00000050781 | ENSBTAG00000050781 | 10 | 22893872 | 22897961 | 3.588501233 | 0.000332584 | 0.047872 |
| Embryo | ENSBTAG00000004916 | ARFGAP1 | 13 | 54287348 | 54297677 | 3.964024433 | 7.36967E-05 | 0.031173706 |
| Hypothalamus | ENSBTAG00000011405 | CEP72 | 20 | 71491866 | 71529187 | -6.538083707 | 6.2312E-11 | 5.53954E-08 |
| Hypothalamus | ENSBTAG00000031509 | PFAS | 19 | 27879985 | 27895903 | -5.81542912 | 6.04785E-09 | 2.68827E-06 |
| Hypothalamus | ENSBTAG00000010236 | CIDEB | 10 | 20862517 | 20865361 | 3.842683153 | 0.000121697 | 0.03606274 |
| Hypothalamus | ENSBTAG00000004411 | CRISPLD1 | 14 | 38295064 | 38345541 | -3.717082827 | 0.000201536 | 0.042625636 |
| Hypothalamus | ENSBTAG00000013753 | DDRGK1 | 13 | 52004036 | 52014649 | 3.672978953 | 0.000239739 | 0.042625636 |
| Jejunum | ENSBTAG00000019984 | GINS1 | 13 | 53589805 | 53606654 | 6.115167547 | 9.64556E-10 | 1.00989E-06 |
| Jejunum | ENSBTAG00000039520 | ENSBTAG00000039520 | 13 | 53431673 | 53469664 | 5.750851302 | 8.87952E-09 | 4.64843E-06 |
| Jejunum | ENSBTAG00000004409 | IVD | 10 | 36022720 | 36034673 | 4.47782249 | 7.54083E-06 | 0.002631748 |
| Jejunum | ENSBTAG00000000132 | EIF4A1 | 19 | 27305005 | 27311552 | 4.04325769 | 5.27136E-05 | 0.013797793 |
| Jejunum | ENSBTAG00000003209 | BLCAP | 13 | 66461377 | 66471861 | 3.783025958 | 0.000154933 | 0.029520546 |
| Jejunum | ENSBTAG00000018742 | MTCH2 | 15 | 77409614 | 77429460 | -3.761093488 | 0.000169172 | 0.029520546 |
| Leukocyte | ENSBTAG00000000132 | EIF4A1 | 19 | 27305005 | 27311552 | 4.326384211 | 1.51577E-05 | 0.008260943 |
| Liver | ENSBTAG00000054447 | ENSBTAG00000054447 | 13 | 53548961 | 53553855 | -7.155094939 | 8.3615E-13 | 1.47814E-09 |
| Liver | ENSBTAG00000052247 | TPPP | 20 | 71467623 | 71488177 | -7.137568849 | 9.4996E-13 | 1.47814E-09 |

| | | | | | | | | |
|---|---|---|---|---|---|---|---|---|
| Liver | ENSBTAG00000006971 | BRD9 | 20 | 71390369 | 71408924 | -6.983492283 | 2.87931E-12 | 2.98681E-09 |
| Liver | ENSBTAG00000012567 | TERT | 20 | 71145309 | 71162695 | -6.416481699 | 1.3946E-10 | 1.085E-07 |
| Liver | ENSBTAG00000031509 | PFAS | 19 | 27879985 | 27895903 | -5.655054477 | 1.55797E-08 | 9.6968E-06 |
| Liver | ENSBTAG00000012608 | GDAP1 | 14 | 37626644 | 37645604 | -5.514072406 | 3.50624E-08 | 1.81857E-05 |
| Liver | ENSBTAG00000046623 | ENSBTAG00000046623 | 13 | 63799750 | 63800272 | -4.883341085 | 1.04303E-06 | 0.000463702 |
| Liver | ENSBTAG00000019984 | GINS1 | 13 | 53589805 | 53606654 | 4.496184604 | 6.91837E-06 | 0.002691245 |
| Liver | ENSBTAG00000000134 | MPDU1 | 19 | 27315837 | 27319239 | -4.46457369 | 8.02283E-06 | 0.002774117 |
| Liver | ENSBTAG00000021874 | KIF14 | 16 | 79237466 | 79278728 | 4.441689957 | 8.92551E-06 | 0.002777619 |
| Liver | ENSBTAG00000005634 | LSM2 | 23 | 27528402 | 27533357 | 4.395185986 | 1.10678E-05 | 0.00313118 |
| Liver | ENSBTAG00000045784 | CENPB | 13 | 51417094 | 51419830 | -4.351074872 | 1.35472E-05 | 0.003442494 |
| Liver | ENSBTAG00000004490 | TRIM31 | 23 | 28868629 | 28879147 | 4.33796816 | 1.43806E-05 | 0.003442494 |
| Liver | ENSBTAG00000054962 | OR4B1GP | 15 | 78022381 | 78023309 | 4.159540998 | 3.18888E-05 | 0.00708842 |
| Liver | ENSBTAG00000046668 | ANG2 | 10 | 26401174 | 26413011 | 4.074977034 | 4.60189E-05 | 0.009547385 |
| Liver | ENSBTAG00000019378 | KIAA1755 | 13 | 67088081 | 67126785 | 4.001087973 | 6.30519E-05 | 0.012263596 |
| Liver | ENSBTAG00000019967 | RASA3 | 12 | 86908880 | 86988568 | -3.976532669 | 6.99274E-05 | 0.012380489 |
| Liver | ENSBTAG00000045985 | ENSBTAG00000045985 | 18 | 59968692 | 59970062 | 3.969163721 | 7.21253E-05 | 0.012380489 |
| Liver | ENSBTAG00000018742 | MTCH2 | 15 | 77409614 | 77429460 | -3.957975154 | 7.55878E-05 | 0.012380489 |
| Liver | ENSBTAG00000004780 | CDKL1 | 10 | 43233495 | 43284821 | 3.928432754 | 8.55013E-05 | 0.013303995 |
| Liver | ENSBTAG00000031354 | ENSBTAG00000031354 | 13 | 62824630 | 62843810 | -3.903926657 | 9.46445E-05 | 0.01402541 |
| Liver | ENSBTAG00000047613 | HTRA3 | 6 | 114593970 | 114613187 | 3.709624118 | 0.000207567 | 0.029361321 |
| Liver | ENSBTAG00000009949 | DARS1 | 2 | 61390060 | 61455629 | -3.654059741 | 0.000258126 | 0.034618971 |
| Liver | ENSBTAG00000020489 | SEMA6A | 7 | 37316829 | 37446102 | 3.645392163 | 0.000266984 | 0.034618971 |
| Liver | ENSBTAG00000000621 | CATSPERG | 18 | 48153486 | 48177891 | -3.62009695 | 0.000294493 | 0.036658445 |
| Liver | ENSBTAG00000013956 | BCL2L13 | 5 | 109082265 | 109133396 | -3.57944046 | 0.000344331 | 0.040856567 |
| Liver | ENSBTAG00000008842 | JPH1 | 14 | 37502881 | 37595533 | -3.571844882 | 0.000354475 | 0.040856567 |
| Liver | ENSBTAG00000017836 | ZFP57 | 23 | 28938566 | 28943319 | 3.535790051 | 0.000406558 | 0.045181045 |
| Liver | ENSBTAG00000003490 | ELMO1 | 4 | 59867412 | 60450510 | -3.526537465 | 0.000421032 | 0.045181045 |
| Lung | ENSBTAG00000047632 | ENSBTAG00000047632 | 20 | 71944313 | 71950871 | 4.237265137 | 2.26259E-05 | 0.035273762 |
| Lymph node | ENSBTAG00000003072 | ACADVL | 19 | 26934465 | 26940043 | 4.621917227 | 3.8021E-06 | 0.006767733 |
| Lymph node | ENSBTAG00000012567 | TERT | 20 | 71145309 | 71162695 | -4.194416067 | 2.73575E-05 | 0.024348213 |
| Macrophage | ENSBTAG00000031800 | PPDPF | 13 | 54136171 | 54137886 | -4.651144319 | 3.30098E-06 | 0.006974975 |
| Mammary | ENSBTAG00000004629 | SLC9A3 | 20 | 71607452 | 71645487 | 5.494908761 | 3.90912E-08 | 5.86559E-05 |
| Mammary | ENSBTAG00000047700 | ENSBTAG00000047700 | 20 | 71943352 | 71965884 | 5.413678155 | 6.17431E-08 | 5.86559E-05 |
| Milk cell | ENSBTAG00000046668 | ANG2 | 10 | 26401174 | 26413011 | 4.093121611 | 4.25604E-05 | 0.025663943 |
| Milk cell | ENSBTAG00000004371 | CS | 5 | 57042171 | 57065001 | -3.690045225 | 0.000224214 | 0.046257135 |
| Milk cell | ENSBTAG00000013919 | ENSBTAG00000013919 | 23 | 25723715 | 25735337 | 3.683409513 | 0.000230135 | 0.046257135 |
| Monocytes | ENSBTAG00000039520 | ENSBTAG00000039520 | 13 | 53431673 | 53469664 | 5.337236381 | 9.4374E-08 | 5.52088E-05 |
| Monocytes | ENSBTAG00000019984 | GINS1 | 13 | 53589805 | 53606654 | 4.988089204 | 6.09794E-07 | 0.000178365 |
| Monocytes | ENSBTAG00000003183 | NTN4 | 5 | 60031676 | 60161682 | -3.749787277 | 0.000176985 | 0.034512005 |
| Monocytes | ENSBTAG00000047281 | C10H15orf62 | 10 | 36319056 | 36320976 | -3.641289519 | 0.000271276 | 0.039674095 |
| Muscle | ENSBTAG00000047780 | CCDC127 | 20 | 71828109 | 71843714 | 6.724447998 | 1.76259E-11 | 4.61976E-08 |
| Muscle | ENSBTAG00000001329 | CLPTM1L | 20 | 71117460 | 71129901 | 6.38583742 | 1.70462E-10 | 2.2339E-07 |
| Muscle | ENSBTAG00000015058 | CDH4 | 13 | 55175303 | 55306529 | -5.6636372 | 1.48198E-08 | 1.29475E-05 |
| Muscle | ENSBTAG00000001038 | SNRPB | 13 | 52666467 | 52675560 | -5.181733147 | 2.19834E-07 | 0.000144046 |
| Muscle | ENSBTAG00000013032 | CCNDBP1 | 10 | 38393529 | 38403862 | -4.959929166 | 7.05189E-07 | 0.00036966 |
| Muscle | ENSBTAG00000046189 | RNASE13 | 10 | 26069296 | 26071780 | -4.702115856 | 2.57479E-06 | 0.001124756 |
| Muscle | ENSBTAG00000000328 | TPPP2 | 10 | 26072695 | 26074864 | -4.535770185 | 5.73937E-06 | 0.002148982 |
| Muscle | ENSBTAG00000007922 | RNF24 | 13 | 51228908 | 51267554 | -4.489552713 | 7.13729E-06 | 0.002338354 |
| Muscle | ENSBTAG00000016208 | TGM2 | 13 | 67002552 | 67037111 | 4.398755639 | 1.08873E-05 | 0.003170633 |
| Muscle | ENSBTAG00000048366 | SNPH | 13 | 59816076 | 59859854 | 4.259457103 | 2.04924E-05 | 0.005371059 |
| Muscle | ENSBTAG00000019378 | KIAA1755 | 13 | 67088081 | 67126785 | -4.137396981 | 3.51268E-05 | 0.00836976 |
| Muscle | ENSBTAG00000046990 | GON7 | 21 | 57823069 | 57830906 | 3.929818819 | 8.50099E-05 | 0.018567577 |
| Muscle | ENSBTAG00000014695 | NAA38 | 19 | 27546606 | 27548927 | 3.848910141 | 0.000118645 | 0.02392056 |
| Muscle | ENSBTAG00000005436 | SHQ1 | 22 | 28835679 | 28939485 | -3.773953901 | 0.00016068 | 0.030081678 |

| | | | | | | | | |
|---|---|---|---|---|---|---|---|---|
| Muscle | ENSBTAG00000020999 | DDB2 | 15 | 77097152 | 77118221 | 3.63453049 | 0.000278487 | 0.048661018 |
| Oviduct | ENSBTAG00000000552 | DISP2 | 10 | 35975164 | 35985917 | 5.625245539 | 1.85244E-08 | 1.27263E-05 |
| Oviduct | ENSBTAG00000004421 | CHST14 | 10 | 36077444 | 36079398 | -4.295941356 | 1.73954E-05 | 0.005975305 |
| Oviduct | ENSBTAG00000025593 | CCDC32 | 10 | 36139789 | 36160950 | -3.73984653 | 0.000184133 | 0.042166375 |
| Pituitary | ENSBTAG00000006971 | BRD9 | 20 | 71390369 | 71408924 | -4.989038698 | 6.06805E-07 | 0.000648068 |
| Pituitary | ENSBTAG00000054594 | ENSBTAG00000054594 | 13 | 53505053 | 53517410 | -4.204658078 | 2.61477E-05 | 0.013962879 |
| Rumen | ENSBTAG00000046493 | PDCD6 | 20 | 71780646 | 71796533 | -7.248198291 | 4.22352E-13 | 5.94672E-10 |
| Rumen | ENSBTAG00000009144 | BPIFA2A | 13 | 62503682 | 62514220 | 5.69258064 | 1.25133E-08 | 8.8094E-06 |
| Rumen | ENSBTAG00000031376 | ENSBTAG00000031376 | 13 | 62565844 | 62575200 | 5.06153623 | 4.15892E-07 | 0.000195192 |
| Rumen | ENSBTAG00000049093 | ENSBTAG00000049093 | 10 | 35969942 | 35973540 | -4.116167812 | 3.85224E-05 | 0.01355988 |
| Rumen | ENSBTAG00000004100 | TTI1 | 13 | 66868177 | 66910251 | 4.002499957 | 6.26767E-05 | 0.017649753 |
| Rumen | ENSBTAG00000005978 | PIK3R5 | 19 | 28354288 | 28428847 | -3.951044404 | 7.78109E-05 | 0.018259621 |
| Salivary gland | ENSBTAG00000055240 | ENSBTAG00000055240 | 20 | 71902799 | 71910531 | -6.662769769 | 2.68714E-11 | 1.55586E-08 |
| Salivary gland | ENSBTAG00000034435 | NKD2 | 20 | 71307541 | 71311025 | 6.403737772 | 1.51618E-10 | 4.38935E-08 |
| Salivary gland | ENSBTAG00000011405 | CEP72 | 20 | 71491866 | 71529187 | -5.469563942 | 4.51144E-08 | 8.70708E-06 |
| Salivary gland | ENSBTAG00000008348 | NOL4L | 13 | 61874277 | 61955381 | -3.713810998 | 0.000204161 | 0.02521047 |
| Salivary gland | ENSBTAG00000000552 | DISP2 | 10 | 35975164 | 35985917 | 3.697530585 | 0.000217707 | 0.02521047 |
| Salivary gland | ENSBTAG00000032055 | VPS39 | 10 | 37550085 | 37596971 | -3.63097981 | 0.000282347 | 0.027246512 |
| Testis | ENSBTAG00000019242 | CHRNB1 | 19 | 27116487 | 27131137 | 5.0394373 | 4.66903E-07 | 0.000184427 |
| Testis | ENSBTAG00000038738 | PXMP4 | 13 | 63150881 | 63167809 | -4.874894813 | 1.08866E-06 | 0.000215011 |
| Testis | ENSBTAG00000020852 | SLC16A11 | 19 | 26825722 | 26827663 | 3.988263845 | 6.65586E-05 | 0.008763551 |
| Testis | ENSBTAG00000011567 | VPS18 | 10 | 36433611 | 36442758 | -3.796845461 | 0.000146549 | 0.014471722 |
| Testis | ENSBTAG00000003961 | SNX20 | 18 | 19065512 | 19074769 | 3.508241892 | 0.000451079 | 0.035635218 |
| Testis | ENSBTAG00000012071 | ASGR1 | 19 | 26893934 | 26899708 | 3.383893882 | 0.000714656 | 0.047048199 |
| Testis | ENSBTAG00000013737 | MROH8 | 13 | 66086809 | 66143601 | 3.325950823 | 0.000881175 | 0.049723432 |
| Uterus | ENSBTAG00000006971 | BRD9 | 20 | 71390369 | 71408924 | -6.555409167 | 5.54895E-11 | 1.36449E-07 |
| Uterus | ENSBTAG00000019984 | GINS1 | 13 | 53589805 | 53606654 | 6.277932582 | 3.43105E-10 | 3.21088E-07 |
| Uterus | ENSBTAG00000045695 | SLC6A19 | 20 | 71183646 | 71199637 | 6.257287815 | 3.9173E-10 | 3.21088E-07 |
| Uterus | ENSBTAG00000054934 | VAMP2 | 19 | 27795436 | 27799326 | -5.713138012 | 1.10912E-08 | 6.81829E-06 |
| Uterus | ENSBTAG00000053461 | OR11G2 | 10 | 26911104 | 26912039 | 4.904039441 | 9.38856E-07 | 0.00046173 |
| Uterus | ENSBTAG00000032055 | VPS39 | 10 | 37550085 | 37596971 | -4.512311203 | 6.4125E-06 | 0.002628057 |
| Uterus | ENSBTAG00000003863 | PCED1A | 13 | 52347705 | 52352814 | -4.46268971 | 8.09372E-06 | 0.002843209 |
| Uterus | ENSBTAG00000032801 | OR11H7 | 10 | 26879265 | 26880236 | 4.433366769 | 9.27728E-06 | 0.002851605 |
| Uterus | ENSBTAG00000013032 | CCNDBP1 | 10 | 38393529 | 38403862 | -4.051135766 | 5.09696E-05 | 0.013926033 |
| Uterus | ENSBTAG00000008842 | JPH1 | 14 | 37502881 | 37595533 | 3.979534064 | 6.90505E-05 | 0.01697951 |
| Uterus | ENSBTAG00000048533 | ENSBTAG00000048533 | 14 | 37324498 | 37325736 | -3.896778398 | 9.74807E-05 | 0.021791376 |
| Uterus | ENSBTAG00000015058 | CDH4 | 13 | 55175303 | 55306529 | -3.812319987 | 0.000137668 | 0.028210568 |
| Uterus | ENSBTAG00000007397 | FOLH1 | 29 | 5885258 | 5958145 | 3.784319426 | 0.00015413 | 0.02911278 |
| Uterus | ENSBTAG00000020596 | TMEM126B | 29 | 9925891 | 9932507 | -3.76620176 | 0.00016575 | 0.02911278 |
| Uterus | ENSBTAG00000055135 | ENSBTAG00000055135 | 5 | 74358375 | 74359379 | -3.696737656 | 0.000218388 | 0.035801043 |

**Supplementary Table 13. Genes with significant TWAS associations and nearby genome-wide significant GWAS signals, related to Figure 4.**

| Gene ID | Gene symbol | Chr | Minimum TWAS FDR | No. of supporting tissues | Supporting tissues | Top GWAS SNP | GWAS P value |
|---|---|---|---|---|---|---|---|
| ENSBTAG00000046493 | PDCD6 | 20 | 5.94672E-10 | 1 | Rumen | 20_71539969_C_A | 1.0526E-12 |
| ENSBTAG00000052247 | TPPP | 20 | 1.47814E-09 | 2 | Blood;Liver | 20_71539969_C_A | 1.0526E-12 |
| ENSBTAG00000054447 | ENSBTAG00000054447 | 13 | 1.47814E-09 | 1 | Liver | 13_53727328_G_A | 2.0267E-10 |
| ENSBTAG00000006971 | BRD9 | 20 | 2.98681E-09 | 3 | Liver; Pituitary; Uterus | 20_71539969_C_A | 1.0526E-12 |
| ENSBTAG00000034435 | NKD2 | 20 | 1.23417E-08 | 2 | Blood; Salivary gland | 20_71539969_C_A | 1.0526E-12 |
| ENSBTAG00000055240 | ENSBTAG00000055240 | 20 | 1.55586E-08 | 1 | Salivary gland | 20_71539969_C_A | 1.0526E-12 |
| ENSBTAG00000047780 | CCDC127 | 20 | 4.61976E-08 | 2 | Blood; Muscle | 20_71539969_C_A | 1.0526E-12 |
| ENSBTAG00000011405 | CEP72 | 20 | 5.53954E-08 | 2 | Hypothalamus; Salivary gland | 20_71539969_C_A | 1.0526E-12 |
| ENSBTAG00000012567 | TERT | 20 | 1.085E-07 | 3 | Blood; Liver; Lymph node | 20_71539969_C_A | 1.0526E-12 |
| ENSBTAG00000001329 | CLPTM1L | 20 | 2.2339E-07 | 2 | Blood; Muscle | 20_71539969_C_A | 1.0526E-12 |
| ENSBTAG00000045695 | SLC6A19 | 20 | 3.21088E-07 | 1 | Uterus | 20_71539969_C_A | 1.0526E-12 |
| ENSBTAG00000019984 | GINS1 | 13 | 3.21088E-07 | 5 | Adipose; Jejunum; Liver; Monocytes; Uterus | 13_53727328_G_A | 2.0267E-10 |
| ENSBTAG00000031509 | PFAS | 19 | 2.68827E-06 | 3 | Adipose; Hypothalamus; Liver | 19_27855458_G_A | 4.1573E-10 |
| ENSBTAG00000008606 | EXOC3 | 20 | 3.66446E-06 | 1 | Blood | 20_71539969_C_A | 1.0526E-12 |
| ENSBTAG00000039520 | ENSBTAG00000039520 | 13 | 4.64843E-06 | 3 | Blood; Jejunum; Monocytes | 13_53727328_G_A | 2.0267E-10 |
| ENSBTAG00000054934 | VAMP2 | 19 | 6.81829E-06 | 2 | Blood; Uterus | 19_27855458_G_A | 4.1573E-10 |
| ENSBTAG00000015058 | CDH4 | 13 | 1.29475E-05 | 2 | Muscle; Uterus | 13_54737771_C_T | 2.213E-09 |
| ENSBTAG00000004629 | SLC9A3 | 20 | 5.86559E-05 | 1 | Mammary | 20_71539969_C_A | 1.0526E-12 |
| ENSBTAG00000047700 | ENSBTAG00000047700 | 20 | 5.86559E-05 | 1 | Mammary | 20_71539969_C_A | 1.0526E-12 |
| ENSBTAG00000001038 | SNRPB | 13 | 0.000144046 | 1 | Muscle | 13_53114773_G_A | 6.363E-09 |
| ENSBTAG00000031800 | PPDPF | 13 | 0.000157186 | 2 | Adipose; Macrophage | 13_53727328_G_A | 2.0267E-10 |
| ENSBTAG00000011638 | ENSBTAG00000011638 | 13 | 0.00017669 | 1 | Blood | 13_53727328_G_A | 2.0267E-10 |
| ENSBTAG00000008291 | PROCR | 13 | 0.00017669 | 1 | Blood | 13_64409704_T_C | 3.5059E-08 |
| ENSBTAG00000011662 | SOX18 | 13 | 0.010629993 | 1 | Blood | 13_53727328_G_A | 2.0267E-10 |
| ENSBTAG00000054594 | ENSBTAG00000054594 | 13 | 0.013962879 | 1 | Pituitary | 13_53727328_G_A | 2.0267E-10 |
| ENSBTAG00000005978 | PIK3R5 | 19 | 0.018259621 | 1 | Rumen | 19_27855458_G_A | 4.1573E-10 |
| ENSBTAG00000016388 | SLCO4A1 | 13 | 0.022779477 | 1 | Blood | 13_54737771_C_T | 2.213E-09 |
| ENSBTAG00000014695 | NAA38 | 19 | 0.02392056 | 1 | Muscle | 19_27855458_G_A | 4.1573E-10 |
| ENSBTAG00000004916 | ARFGAP1 | 13 | 0.031173706 | 1 | Embryo | 13_53811139_C_T | 1.2537E-09 |
| ENSBTAG00000047632 | ENSBTAG00000047632 | 20 | 0.035273762 | 1 | Lung | 20_71539969_C_A | 1.0526E-12 |

**Supplementary Table 14. GWAS-annotated telomere length candidate genes supported by both TWAS and eQTL colocalization, related to Figure S14.**

| Gene | Chr | GWAS lead SNP position | TWAS min FDR | TWAS tissues | eQTL max PP.H4 | eQTL tissues |
|---|---|---|---|---|---|---|
| GINS1 | chr13 | chr13:53727328 | 0.000000321 | Adipose;Jejunum;Liver;Mo | 0.710267 | Vas deferens |
| PFAS | chr19 | chr19:27855458 | 0.00000269 | Adipose;Hypothalamus;Liv | 0.992646596 | Adipose; Brain; Epididymis; Gut; Hypothalamus; Jejunum; Liver; Lung; Lymph node; Mammary gland; Muscle; Pituitary; Placenta; Rumen; Small intestine; Subcutaneous adipose; Testis; Uterus; Vas |
| BRD9 | chr20 | chr20:71539969 | 2.99E-09 | Liver;Pituitary;Uterus | 0.810893006 | Gut; Liver |
| TERT | chr20 | chr20:71539969 | 0.000000108 | Blood;Liver;Lymph node | 0.817084093 | Vas deferens |

**Supplementary Table 15. Pearson correlations between telomere-related gene expression and telomere length in rumen and whole blood, related to Figure S15.**

| Gene | Rumen | | Whole Blood | |
|---|---|---|---|---|
| | Pearson r | P value | Pearson r | P value |
| NBN | -0.228502164 | 4.01257E-05 | -0.050277307 | 0.39446464 |
| NSMCE4A | -0.224914687 | 5.33234E-05 | 0.017623139 | 0.765460365 |
| AURKB | -0.079157564 | 0.159724148 | 0.205468976 | 0.000439066 |
| PCNA | -0.136042156 | 0.015356005 | 0.196475879 | 0.00078393 |
| WRAP53 | -0.182757291 | 0.001080741 | -0.024462475 | 0.67878494 |
| E2F1 | -0.104957063 | 0.061973748 | 0.187113614 | 0.001396304 |
| ADA | -0.135993959 | 0.015392694 | 0.164641025 | 0.005016998 |
| PNKP | -0.152214529 | 0.006622856 | 0.046391754 | 0.43206379 |
| TERF2IP | 0.152090964 | 0.006667594 | 0.019048896 | 0.747106478 |
| MAPK14 | -0.051125041 | 0.364272991 | 0.147071068 | 0.012314632 |
| PRKCQ | -0.045501385 | 0.419466567 | 0.144742229 | 0.013780407 |
| SMARCAL1 | -0.135988523 | 0.015396836 | 0.001309875 | 0.982311242 |
| RAD50 | -0.131429326 | 0.019234646 | -0.047765422 | 0.418541112 |
| IRF1 | -0.130251778 | 0.020351881 | 0.002015654 | 0.972783335 |
| SCYL1 | -0.129468755 | 0.021125612 | 0.104204821 | 0.076956902 |
| TINF2 | -0.129238074 | 0.021358366 | -0.051338023 | 0.38455318 |
| BRCA1 | -0.005639615 | 0.920332714 | 0.130997797 | 0.025952896 |
| SP3 | -0.124124805 | 0.027118945 | 0.063047667 | 0.285417532 |
| TERF1 | 0.037234544 | 0.508899388 | -0.129036185 | 0.028286824 |
| MEN1 | -0.121030515 | 0.031216675 | 0.021069722 | 0.72133697 |
| HSP90AA1 | -0.043032435 | 0.445166579 | 0.126351694 | 0.031770787 |
| CHEK1 | -0.070487696 | 0.21071663 | 0.124122979 | 0.034935225 |
| BCL2L1 | 0.115625469 | 0.039645281 | 0.06606061 | 0.26297531 |
| TP53 | -0.003965895 | 0.94392934 | -0.117262967 | 0.046403171 |

**Supplementary Table 16. Pearson correlations between the expression of GWAS-annotated candidate genes and telomere length in rumen and whole blood, related to Figures 5 and S15.**

| Gene | Rumen | | Whole blood | |
|---|---|---|---|---|
| | Pearson r | P value | Pearson r | P value |
| SIRPA | -0.027552262 | 0.625047741 | 0.16409207 | 0.005166652 |
| ABHD16B | NA | NA | 0.156487988 | 0.007694119 |
| GINS1 | 0.000317078 | 0.995513426 | 0.152256277 | 0.009534497 |
| DIDO1 | -0.076499654 | 0.174257926 | 0.148033778 | 0.011750014 |
| SOX18 | -0.094525557 | 0.092938826 | 0.141781203 | 0.015863909 |
| GMEB2 | -0.130497945 | 0.020113774 | 0.092487963 | 0.116683926 |
| TCEA2 | -0.05619142 | 0.318620978 | 0.133977845 | 0.022724664 |
| ARFRP1 | -0.040766445 | 0.469520567 | 0.130323693 | 0.026735546 |
| PRPF6 | -0.117131468 | 0.037123052 | 0.128486091 | 0.028972777 |
| ZGPAT | -0.084528733 | 0.133165604 | 0.126656243 | 0.031357968 |
| SLCO4A1 | -0.025909456 | 0.645832382 | 0.11990544 | 0.041658791 |
| C13H20orf204 | -0.11429423 | 0.041993903 | NA | NA |

**Supplementary Table 17. Associations between WGCNA module eigengenes and telomere length in rumen and whole blood, related to Figures 5 and S16.**

| Tissue | Module | Pearson r | P value |
|---|---|---|---|
| Rumen | MEbrown | -0.143829979 | 0.010345899 |
| Rumen | MEblack | -0.132415888 | 0.018340073 |
| Rumen | MEpurple | -0.116592832 | 0.038009074 |
| Rumen | MEgreenyellow | -0.108784966 | 0.052995703 |
| Rumen | MEblue | -0.103422062 | 0.065909643 |
| Rumen | MEgreen | 0.079916536 | 0.155746336 |
| Rumen | MEmagenta | -0.076552392 | 0.173960296 |
| Rumen | MEred | -0.066585852 | 0.237143622 |
| Rumen | MEturquoise | -0.047327405 | 0.401030399 |
| Rumen | MEyellow | -0.046938737 | 0.404913425 |
| Rumen | MEpink | -0.032819455 | 0.5604439 |
| Rumen | MEgrey | 0.014511999 | 0.796891878 |
| Whole blood | MEgreen | 0.173542831 | 0.003077051 |
| Whole blood | MEbrown | 0.150326353 | 0.010496454 |
| Whole blood | MEred | 0.133285396 | 0.023442084 |
| Whole blood | MEyellow | 0.092441312 | 0.116869265 |
| Whole blood | MEblack | 0.073301917 | 0.214087538 |
| Whole blood | MEblue | 0.061950109 | 0.293903869 |
| Whole blood | MEpink | -0.059227129 | 0.315678205 |
| Whole blood | MEgrey | 0.042584575 | 0.470831074 |
| Whole blood | MEmagenta | -0.038255932 | 0.517134065 |
| Whole blood | MEturquoise | 0.019369758 | 0.74299517 |

**Supplementary Table 18. Gene Ontology enrichment of the telomere length-associated MEblack module in rumen, related to Figure S16.**

| Module | ONTOLOGY | ID | Description | GeneRatio | BgRatio | RichFactor | FoldEnrichment | zScore | pvalue | p.adjust | qvalue | geneID | Count |
|---|---|---|---|---|---|---|---|---|---|---|---|---|---|
| MEblack | BP | GO:006151 | protein localization to cilium | 5/238 | 34/16092 | 0.147058824 | 9.943153732 | 6.395801802 | 0.000133016 | 0.157490649 | 0.053341426 | SPATA7/ASB3/CC2D2B/E | 5 |
| MEblack | MF | GO:004452 | DNA endonuclease activity | 5/245 | 34/16185 | 0.147058824 | 9.714885954 | 6.306425602 | 0.000148315 | 0.053245176 | 0.016185217 | DNASE1/EME2/DNASE2/I | 5 |
| MEblack | MF | GO:004453 | DNA nuclease activity | 5/245 | 46/16185 | 0.108695652 | 7.180567879 | 5.204161962 | 0.000630382 | 0.104813042 | 0.031860574 | DNASE1/EME2/DNASE2/I | 5 |
| MEblack | BP | GO:000734 | regulation of mitotic cell cycle | 8/238 | 130/16092 | 0.061538462 | 4.160827408 | 4.433425087 | 0.000702566 | 0.232216015 | 0.078650596 | KNTC1/EME2/PIM2/EME1 | 8 |
| MEblack | BP | GO:000838 | RNA splicing | 12/238 | 275/16092 | 0.043636364 | 2.950404889 | 3.997050502 | 0.00083235 | 0.232216015 | 0.078650596 | DDX39B/RBM41/CLK4/CL | 12 |
| MEblack | MF | GO:000998 | pseudouridine synthase activity | 3/245 | 13/16185 | 0.230769231 | 15.24489796 | 6.369875894 | 0.000875875 | 0.104813042 | 0.031860574 | PUS1/RPUSD3/PUS10 | 3 |
| MEblack | CC | GO:000581 | microtubule organizing center | 11/230 | 244/15882 | 0.045081967 | 3.11300784 | 4.032026136 | 0.000880632 | 0.106371374 | 0.005908906 | CCDC77/CROCC/SPATA7 | 11 |
| MEblack | CC | GO:001563 | microtubule cytoskeleton | 17/230 | 497/15882 | 0.034205231 | 2.361945587 | 3.739444759 | 0.000945523 | 0.106371374 | 0.005908906 | KNTC1/ATAT1/CCDC77/C | 17 |
| MEblack | BP | GO:000945 | RNA modification | 7/238 | 112/16092 | 0.0625 | 4.225840336 | 4.197339138 | 0.001377297 | 0.232216015 | 0.078650596 | TRMT44/PUS1/NSUN6/G | 7 |
| MEblack | BP | GO:000625 | DNA metabolic process | 15/238 | 418/16092 | 0.035885167 | 2.42631981 | 3.620142807 | 0.001444736 | 0.232216015 | 0.078650596 | DNASE1/DBF4B/EME2/FA | 15 |
| MEblack | BP | GO:000630 | DNA catabolic process | 3/238 | 16/16092 | 0.1875 | 12.67752101 | 5.725747852 | 0.001551486 | 0.232216015 | 0.078650596 | DNASE1/DNASE2/DFFB | 3 |
| MEblack | BP | GO:005172 | regulation of cell cycle | 11/238 | 259/16092 | 0.042471042 | 2.87161351 | 3.720446097 | 0.001686916 | 0.232216015 | 0.078650596 | DBF4B/KNTC1/EME2/PIM | 11 |
| MEblack | BP | GO:000152 | pseudouridine synthesis | 3/238 | 17/16092 | 0.176470588 | 11.93178448 | 5.525233036 | 0.001863497 | 0.232216015 | 0.078650596 | PUS1/RPUSD3/PUS10 | 3 |
| MEblack | BP | GO:190198 | regulation of cell cycle phase transition | 7/238 | 119/16092 | 0.058823529 | 3.977261493 | 3.993997459 | 0.001952534 | 0.232216015 | 0.078650596 | DBF4B/KNTC1/EME2/EME | 7 |
| MEblack | BP | GO:000640 | tRNA modification | 5/238 | 61/16092 | 0.081967213 | 5.542085687 | 4.354622402 | 0.002057913 | 0.232216015 | 0.078650596 | TRMT44/PUS1/GTPBP3/F | 5 |
| MEblack | CC | GO:007100 | U2-type prespliceosome | 3/230 | 18/15882 | 0.166666667 | 11.50869565 | 5.407501363 | 0.0020829 | 0.117163101 | 0.006508384 | LUC7L3/LUC7L/PRPF40B | 3 |
| MEblack | CC | GO:007101 | prespliceosome | 3/230 | 18/15882 | 0.166666667 | 11.50869565 | 5.407501363 | 0.0020829 | 0.117163101 | 0.006508384 | LUC7L3/LUC7L/PRPF40B | 3 |
| MEblack | MF | GO:014009 | catalytic activity, acting on DNA | 9/245 | 186/16185 | 0.048387097 | 3.196510862 | 3.735297755 | 0.002138669 | 0.191945506 | 0.058346689 | DNASE1/EME2/MUTYH/D | 9 |
| MEblack | BP | GO:000027 | mitotic cell cycle | 13/238 | 352/16092 | 0.036931818 | 2.497087471 | 3.479579092 | 0.002306864 | 0.232216015 | 0.078650596 | KNTC1/EME2/PIM2/EME1 | 13 |
| MEblack | BP | GO:004477 | cell cycle phase transition | 8/238 | 157/16092 | 0.050955414 | 3.445271102 | 3.772344761 | 0.002353541 | 0.232216015 | 0.078650596 | DBF4B/KNTC1/EME2/EME | 8 |
| MEblack | BP | GO:001607 | mRNA metabolic process | 13/238 | 374/16092 | 0.034759358 | 2.350199973 | 3.237023281 | 0.003870302 | 0.352495154 | 0.119388639 | DDX39B/UPF3B/PUS1/RE | 13 |
| MEblack | BP | GO:004478 | cilium organization | 9/238 | 220/16092 | 0.040909091 | 2.766004584 | 3.231453482 | 0.005552873 | 0.395244725 | 0.13386774 | CCDC66/CROCC/CCDC1 | 9 |
| MEblack | BP | GO:001056 | regulation of cell cycle process | 8/238 | 183/16092 | 0.043715847 | 2.955779033 | 3.260126552 | 0.005947007 | 0.395244725 | 0.13386774 | DBF4B/KNTC1/EME2/EME | 8 |
| MEblack | BP | GO:003048 | tRNA methylation | 3/238 | 26/16092 | 0.115384615 | 7.80155139 | 4.25256877 | 0.006460842 | 0.395244725 | 0.13386774 | TRMT44/GTPBP3/TRMT1 | 3 |
| MEblack | BP | GO:004341 | macromolecule methylation | 5/238 | 80/16092 | 0.0625 | 4.225840336 | 3.543852518 | 0.006635195 | 0.395244725 | 0.13386774 | TRMT44/NSUN6/GTPBP3 | 5 |
| MEblack | BP | GO:190304 | mitotic cell cycle process | 10/238 | 268/16092 | 0.037313433 | 2.522889753 | 3.080266505 | 0.006697487 | 0.395244725 | 0.13386774 | KNTC1/EME2/EME1/RTKI | 10 |
| MEblack | BP | GO:000151 | RNA methylation | 4/238 | 51/16092 | 0.078431373 | 5.303015324 | 3.770967855 | 0.006767507 | 0.395244725 | 0.13386774 | TRMT44/NSUN6/GTPBP3 | 4 |
| MEblack | CC | GO:000587 | spindle microtubule | 3/230 | 27/15882 | 0.111111111 | 7.672463768 | 4.206325767 | 0.006784621 | 0.255189628 | 0.014175728 | KNTC1/CCDC57/AURKC | 3 |
| MEblack | MF | GO:001686 | intramolecular transferase activity | 3/245 | 27/16185 | 0.111111111 | 7.340136054 | 4.087600808 | 0.007663814 | 0.508884201 | 0.154688219 | PUS1/RPUSD3/PUS10 | 3 |
| MEblack | BP | GO:000037 | RNA splicing, via transesterification reactions | 9/238 | 232/16092 | 0.038793103 | 2.622935381 | 3.050732988 | 0.007767782 | 0.395244725 | 0.13386774 | DDX39B/RBM41/RBMX2/L | 9 |
| MEblack | BP | GO:000037 | RNA splicing, via transesterification reactions with bulged adenosine as nucleophile | 9/238 | 232/16092 | 0.038793103 | 2.622935381 | 3.050732988 | 0.007767782 | 0.395244725 | 0.13386774 | DDX39B/RBM41/RBMX2/L | 9 |
| MEblack | BP | GO:000039 | mRNA splicing, via spliceosome | 9/238 | 232/16092 | 0.038793103 | 2.622935381 | 3.050732988 | 0.007767782 | 0.395244725 | 0.13386774 | DDX39B/RBM41/RBMX2/L | 9 |
| MEblack | BP | GO:000738 | pattern specification process | 5/238 | 84/16092 | 0.05952381 | 4.024609844 | 3.405265948 | 0.008127003 | 0.395244725 | 0.13386774 | CCDC40/HOXD4/HOXB6/ | 5 |
| MEblack | BP | GO:003225 | methylation | 5/238 | 84/16092 | 0.05952381 | 4.024609844 | 3.405265948 | 0.008127003 | 0.395244725 | 0.13386774 | TRMT44/NSUN6/GTPBP3 | 5 |
| MEblack | CC | GO:000568 | U1 snRNP | 3/230 | 29/15882 | 0.103448276 | 7.143328336 | 4.013884927 | 0.008297847 | 0.255189628 | 0.014175728 | LUC7L3/LUC7L/PRPF40B | 3 |
| MEblack | BP | GO:006027 | cilium assembly | 8/238 | 194/16092 | 0.041237113 | 2.788183315 | 3.070097909 | 0.008345539 | 0.395244725 | 0.13386774 | CCDC66/CROCC/CCDC1 | 8 |
| MEblack | MF | GO:000451 | endonuclease activity | 5/245 | 83/16185 | 0.060240964 | 3.979591837 | 3.373942907 | 0.008505028 | 0.508884201 | 0.154688219 | DNASE1/EME2/DNASE2/I | 5 |
| MEblack | CC | GO:000592 | cilium | 10/230 | 285/15882 | 0.035087719 | 2.422883295 | 2.938252375 | 0.008773062 | 0.255189628 | 0.014175728 | CCDC66/SPATA7/CATSP | 10 |
| MEblack | BP | GO:004558 | regulation of T cell differentiation | 2/238 | 10/16092 | 0.2 | 13.52268908 | 4.853311179 | 0.009064575 | 0.41278681 | 0.139809171 | IL23A/FANCA | 2 |
| MEblack | CC | GO:000581 | centrosome | 7/230 | 161/15882 | 0.043478261 | 3.002268431 | 3.095373609 | 0.009073409 | 0.255189628 | 0.014175728 | CCDC77/CROCC/CCDC1 | 7 |
| MEblack | BP | GO:000639 | mRNA processing | 10/238 | 284/16092 | 0.035211268 | 2.380755119 | 2.876391107 | 0.00986181 | 0.432458634 | 0.146471936 | DDX39B/RBM41/RBMX2/L | 10 |
| MEblack | BP | GO:003019 | extracellular matrix organization | 6/238 | 126/16092 | 0.047619048 | 3.219687875 | 3.064712141 | 0.01111444 | 0.442060533 | 0.149724059 | ADAMTS10/COL27A1/LAM | 6 |
| MEblack | BP | GO:004522 | external encapsulating structure organization | 6/238 | 126/16092 | 0.047619048 | 3.219687875 | 3.064712141 | 0.01111444 | 0.442060533 | 0.149724059 | ADAMTS10/COL27A1/LAM | 6 |
| MEblack | MF | GO:003098 | mismatched DNA binding | 2/245 | 11/16185 | 0.181818182 | 12.01113173 | 4.528990397 | 0.011470672 | 0.550774431 | 0.167421813 | MUTYH/MLH1 | 2 |
| MEblack | BP | GO:000008 | G2/M transition of mitotic cell cycle | 3/238 | 32/16092 | 0.09375 | 6.338760504 | 3.703848606 | 0.011550999 | 0.442060533 | 0.149724059 | MIIP/CDK3/SKP2 | 3 |
| MEblack | BP | GO:000803 | tRNA processing | 5/238 | 92/16092 | 0.054347826 | 3.674643771 | 3.152175861 | 0.01177925 | 0.442060533 | 0.149724059 | TRMT44/PUS1/GTPBP3/F | 5 |
| MEblack | BP | GO:004306 | extracellular structure organization | 6/238 | 128/16092 | 0.046875 | 3.169380252 | 3.019120011 | 0.011947582 | 0.442060533 | 0.149724059 | ADAMTS10/COL27A1/LAM | 6 |
| MEblack | CC | GO:004324 | Fanconi anaemia nuclear complex | 2/230 | 12/15882 | 0.166666667 | 11.50869565 | 4.414371663 | 0.012524916 | 0.287586542 | 0.015975369 | FANCG/FANCA | 2 |
| MEblack | CC | GO:009752 | spliceosomal snRNP complex | 4/230 | 65/15882 | 0.061538462 | 4.249364548 | 3.182079166 | 0.014569541 | 0.287586542 | 0.015975369 | RBMX2/LUC7L3/LUC7L/P | 4 |
| MEblack | CC | GO:009879 | plasma membrane protein complex | 11/230 | 357/15882 | 0.030812325 | 2.12765802 | 2.612243383 | 0.015185017 | 0.287586542 | 0.015975369 | GNAT1/KCNIP2/SCN11A/ | 11 |
| MEblack | BP | GO:000071 | resolution of meiotic recombination intermediates | 2/238 | 13/16092 | 0.153846154 | 10.40206852 | 4.155048902 | 0.01526039 | 0.461473588 | 0.15629918 | EME2/EME1 | 2 |
| MEblack | BP | GO:003019 | collagen fibril organization | 2/238 | 13/16092 | 0.153846154 | 10.40206852 | 4.155048902 | 0.01526039 | 0.461473588 | 0.15629918 | OPTC/LOXL3 | 2 |
| MEblack | BP | GO:004255 | superoxide anion generation | 2/238 | 13/16092 | 0.153846154 | 10.40206852 | 4.155048902 | 0.01526039 | 0.461473588 | 0.15629918 | NOXO1/NOXA1 | 2 |
| MEblack | CC | GO:003053 | small nuclear ribonucleoprotein complex | 4/230 | 66/15882 | 0.060606061 | 4.184980237 | 3.143028382 | 0.015337949 | 0.287586542 | 0.015975369 | RBMX2/LUC7L3/LUC7L/P | 4 |
| MEblack | BP | GO:004483 | cell cycle G2/M phase transition | 3/238 | 37/16092 | 0.081081081 | 5.482171247 | 3.344219088 | 0.017146509 | 0.461473588 | 0.15629918 | MIIP/CDK3/SKP2 | 3 |
| MEblack | BP | GO:000637 | mRNA splice site recognition | 2/238 | 14/16092 | 0.142857143 | 9.659063625 | 3.971270827 | 0.017631997 | 0.461473588 | 0.15629918 | LUC7L3/LUC7L | 2 |

| | | | | | | | | | | | | | |
|---|---|---|---|---|---|---|---|---|---|---|---|---|---|
| MEblack | BP | GO:000677 | porphyrin-containing compound biosynthetic process | 2/238 | 14/16092 | 0.142857143 | 9.659063625 | 3.971270827 | 0.017631997 | 0.461473588 | 0.15629918 | PPOX/RSAD1 | 2 |
| MEblack | BP | GO:000721 | gamma-aminobutyric acid signaling pathway | 2/238 | 14/16092 | 0.142857143 | 9.659063625 | 3.971270827 | 0.017631997 | 0.461473588 | 0.15629918 | GABBR1/GABRE | 2 |
| MEblack | BP | GO:003301 | tetrapyrrole biosynthetic process | 2/238 | 14/16092 | 0.142857143 | 9.659063625 | 3.971270827 | 0.017631997 | 0.461473588 | 0.15629918 | PPOX/RSAD1 | 2 |
| MEblack | BP | GO:000300 | regionalization | 4/238 | 68/16092 | 0.058823529 | 3.977261493 | 3.014369852 | 0.018170202 | 0.461473588 | 0.15629918 | CCDC40/HOXD4/HOXB6/ | 4 |
| MEblack | BP | GO:004001 | negative regulation of locomotion | 3/238 | 38/16092 | 0.078947368 | 5.337903582 | 3.280126802 | 0.018417279 | 0.461473588 | 0.15629918 | SEMA3B/MIIP/CCDC125 | 3 |
| MEblack | CC | GO:000568 | spliceosomal complex | 7/230 | 186/15882 | 0.037634409 | 2.598737728 | 2.658625232 | 0.018805999 | 0.297382778 | 0.016519548 | RBM41/RBMX2/LUC7L3/R | 7 |
| MEblack | CC | GO:003445 | centriolar satellite | 2/230 | 15/15882 | 0.133333333 | 9.206956522 | 3.854768244 | 0.019366528 | 0.297382778 | 0.016519548 | CCDC13/CCDC57 | 2 |
| MEblack | MF | GO:001990 | syntaxin binding | 3/245 | 38/16185 | 0.078947368 | 5.215359828 | 3.225242297 | 0.019576178 | 0.550774431 | 0.167421813 | SYT2/NAPB/VAMP1 | 3 |
| MEblack | BP | GO:004593 | positive regulation of mitotic cell cycle | 2/238 | 15/16092 | 0.133333333 | 9.01512605 | 3.805082255 | 0.020148546 | 0.461473588 | 0.15629918 | ANAPC5/CCDC57 | 2 |
| MEblack | CC | GO:000587 | microtubule | 6/230 | 147/15882 | 0.040816327 | 2.818456078 | 2.685010762 | 0.020211952 | 0.297382778 | 0.016519548 | KNTC1/ATAT1/CCDC66/K | 6 |
| MEblack | BP | GO:000695 | inflammatory response | 6/238 | 144/16092 | 0.041666667 | 2.817226891 | 2.683785835 | 0.02025059 | 0.461473588 | 0.15629918 | CST7/TLR6/LOXL3/GGT5/ | 6 |
| MEblack | BP | GO:000600 | glucose metabolic process | 3/238 | 40/16092 | 0.075 | 5.071008403 | 3.158478958 | 0.021111797 | 0.461473588 | 0.15629918 | PCK1/GAPDHS/GNMT | 3 |
| MEblack | BP | GO:000602 | aminoglycan metabolic process | 3/238 | 40/16092 | 0.075 | 5.071008403 | 3.158478958 | 0.021111797 | 0.461473588 | 0.15629918 | OVGP1/ITIH4/HEXB | 3 |
| MEblack | CC | GO:000581 | spindle | 5/230 | 109/15882 | 0.04587156 | 3.167530913 | 2.752575999 | 0.02114722 | 0.297382778 | 0.016519548 | KNTC1/CCDC57/PLK1/CE | 5 |
| MEblack | BP | GO:004593 | negative regulation of mitotic cell cycle | 4/238 | 72/16092 | 0.055555556 | 3.756302521 | 2.871920698 | 0.021936784 | 0.461473588 | 0.15629918 | KNTC1/EME2/EME1/MIIP | 4 |
| MEblack | BP | GO:000674 | glutathione metabolic process | 3/238 | 41/16092 | 0.073170732 | 4.947325272 | 3.100661559 | 0.022535677 | 0.461473588 | 0.15629918 | TXNRD3/GGT5/OPLAH | 3 |
| MEblack | BP | GO:000713 | reciprocal meiotic recombination | 3/238 | 41/16092 | 0.073170732 | 4.947325272 | 3.100661559 | 0.022535677 | 0.461473588 | 0.15629918 | EME2/EME1/CCNB1IP1 | 3 |
| MEblack | BP | GO:003582 | homologous recombination | 3/238 | 41/16092 | 0.073170732 | 4.947325272 | 3.100661559 | 0.022535677 | 0.461473588 | 0.15629918 | EME2/EME1/CCNB1IP1 | 3 |
| MEblack | BP | GO:014052 | reciprocal homologous recombination | 3/238 | 41/16092 | 0.073170732 | 4.947325272 | 3.100661559 | 0.022535677 | 0.461473588 | 0.15629918 | EME2/EME1/CCNB1IP1 | 3 |
| MEblack | CC | GO:000568 | U2-type spliceosomal complex | 3/230 | 42/15882 | 0.071428571 | 4.932298137 | 3.093224315 | 0.022732868 | 0.300876192 | 0.016713606 | LUC7L3/LUC7L/PRPF40B | 3 |
| MEblack | BP | GO:000609 | gluconeogenesis | 2/238 | 16/16092 | 0.125 | 8.451680672 | 3.6537245 | 0.022805269 | 0.461473588 | 0.15629918 | PCK1/GNMT | 2 |
| MEblack | BP | GO:000629 | mismatch repair | 2/238 | 16/16092 | 0.125 | 8.451680672 | 3.6537245 | 0.022805269 | 0.461473588 | 0.15629918 | MUTYH/MLH1 | 2 |
| MEblack | BP | GO:004561 | regulation of lymphocyte differentiation | 2/238 | 16/16092 | 0.125 | 8.451680672 | 3.6537245 | 0.022805269 | 0.461473588 | 0.15629918 | IL23A/FANCA | 2 |
| MEblack | BP | GO:004749 | vesicle transport along microtubule | 2/238 | 16/16092 | 0.125 | 8.451680672 | 3.6537245 | 0.022805269 | 0.461473588 | 0.15629918 | BICDL1/HAP1 | 2 |
| MEblack | BP | GO:003465 | nucleobase-containing compound catabolic process | 7/238 | 190/16092 | 0.036842105 | 2.491021672 | 2.533059848 | 0.023094758 | 0.461473588 | 0.15629918 | DNASE1/UPF3B/DNASE2 | 7 |
| MEblack | MF | GO:001674 | transferase activity, transferring one-carbon groups | 7/245 | 186/16185 | 0.037634409 | 2.486175115 | 2.527329441 | 0.02331074 | 0.550774431 | 0.167421813 | TRMT44/NSUN6/AMT/GN | 7 |
| MEblack | MF | GO:003600 | pre-mRNA binding | 2/245 | 16/16185 | 0.125 | 8.257653061 | 3.600778718 | 0.0238154 | 0.550774431 | 0.167421813 | RBM41/RNPC3 | 2 |
| MEblack | MF | GO:001990 | phosphatase binding | 3/245 | 41/16185 | 0.073170732 | 4.833748133 | 3.047135382 | 0.023935809 | 0.550774431 | 0.167421813 | PPP1R36/PPP6R1/SFI1 | 3 |
| MEblack | MF | GO:000467 | protein serine/threonine kinase activity | 10/245 | 321/16185 | 0.031152648 | 2.057982071 | 2.373593217 | 0.024694888 | 0.550774431 | 0.167421813 | ULK3/CLK4/CLK1/BRSK1/ | 10 |
| MEblack | MF | GO:000808 | phosphoric diester hydrolase activity | 4/245 | 73/16185 | 0.054794521 | 3.619793123 | 2.781221127 | 0.02473043 | 0.550774431 | 0.167421813 | GDPD3/SMPD2/MPPE1/P | 4 |
| MEblack | CC | GO:012011 | Sm-like protein family complex | 4/230 | 77/15882 | 0.051948052 | 3.587125918 | 2.758567406 | 0.025492817 | 0.318660212 | 0.017701505 | RBMX2/LUC7L3/LUC7L/P | 4 |
| MEblack | BP | GO:012003 | plasma membrane bounded cell projection assembly | 8/238 | 238/16092 | 0.033613445 | 2.272720853 | 2.42361025 | 0.025514418 | 0.461473588 | 0.15629918 | CCDC66/CROCC/CCDC1 | 8 |
| MEblack | BP | GO:000673 | one-carbon metabolic process | 2/238 | 17/16092 | 0.117647059 | 7.954522986 | 3.515012597 | 0.025597505 | 0.461473588 | 0.15629918 | GNMT/CA9 | 2 |
| MEblack | BP | GO:000677 | porphyrin-containing compound metabolic process | 2/238 | 17/16092 | 0.117647059 | 7.954522986 | 3.515012597 | 0.025597505 | 0.461473588 | 0.15629918 | PPOX/RSAD1 | 2 |
| MEblack | BP | GO:000907 | aromatic amino acid family catabolic process | 2/238 | 17/16092 | 0.117647059 | 7.954522986 | 3.515012597 | 0.025597505 | 0.461473588 | 0.15629918 | AFMID/AMDHD1 | 2 |
| MEblack | BP | GO:001931 | hexose biosynthetic process | 2/238 | 17/16092 | 0.117647059 | 7.954522986 | 3.515012597 | 0.025597505 | 0.461473588 | 0.15629918 | PCK1/GNMT | 2 |
| MEblack | BP | GO:001569 | inorganic anion transport | 4/238 | 76/16092 | 0.052631579 | 3.558602388 | 2.739322319 | 0.026147153 | 0.461473588 | 0.15629918 | CLCN2/SLC12A9/GABRE/ | 4 |
| MEblack | MF | GO:001664 | oxidoreductase activity, acting on the CH-NH2 group of donors, oxygen as acceptor | 2/245 | 17/16185 | 0.117647059 | 7.771908764 | 3.463292938 | 0.02672527 | 0.550774431 | 0.167421813 | LOXL3/AOC2 | 2 |
| MEblack | MF | GO:014061 | DNA damage sensor activity | 2/245 | 17/16185 | 0.117647059 | 7.771908764 | 3.463292938 | 0.02672527 | 0.550774431 | 0.167421813 | MLH1/XRCC3 | 2 |
| MEblack | MF | GO:014066 | ATP-dependent DNA damage sensor activity | 2/245 | 17/16185 | 0.117647059 | 7.771908764 | 3.463292938 | 0.02672527 | 0.550774431 | 0.167421813 | MLH1/XRCC3 | 2 |
| MEblack | BP | GO:000836 | regulation of cell size | 3/238 | 44/16092 | 0.068181818 | 4.610007639 | 2.937883502 | 0.027113782 | 0.461473588 | 0.15629918 | SEMA3B/SLC12A9/ZFYVE | 3 |
| MEblack | BP | GO:001810 | peptidyl-tyrosine phosphorylation | 3/238 | 44/16092 | 0.068181818 | 4.610007639 | 2.937883502 | 0.027113782 | 0.461473588 | 0.15629918 | CLK4/CLK1/CLCF1 | 3 |
| MEblack | BP | GO:003253 | regulation of cellular component size | 5/238 | 115/16092 | 0.043478261 | 2.939715016 | 2.557701155 | 0.028069002 | 0.461473588 | 0.15629918 | SEMA3B/IQANK1/SLC12A | 5 |
| MEblack | BP | GO:000626 | DNA-templated DNA replication | 4/238 | 78/16092 | 0.051282051 | 3.467356173 | 2.676330877 | 0.028421808 | 0.461473588 | 0.15629918 | DBF4B/EME2/POLG2/EMI | 4 |
| MEblack | BP | GO:190199 | regulation of mitotic cell cycle phase transition | 4/238 | 78/16092 | 0.051282051 | 3.467356173 | 2.676330877 | 0.028421808 | 0.461473588 | 0.15629918 | KNTC1/MIIP/ANAPC5/CDI | 4 |
| MEblack | BP | GO:001967 | NAD+ metabolic process | 2/238 | 18/16092 | 0.111111111 | 7.512605042 | 3.387190173 | 0.028520701 | 0.461473588 | 0.15629918 | NAPRT/NUDT17 | 2 |
| MEblack | BP | GO:004636 | monosaccharide biosynthetic process | 2/238 | 18/16092 | 0.111111111 | 7.512605042 | 3.387190173 | 0.028520701 | 0.461473588 | 0.15629918 | PCK1/GNMT | 2 |
| MEblack | BP | GO:009719 | execution phase of apoptosis | 2/238 | 18/16092 | 0.111111111 | 7.512605042 | 3.387190173 | 0.028520701 | 0.461473588 | 0.15629918 | DNASE2/DFFB | 2 |
| MEblack | BP | GO:200014 | negative regulation of cell motility | 2/238 | 18/16092 | 0.111111111 | 7.512605042 | 3.387190173 | 0.028520701 | 0.461473588 | 0.15629918 | MIIP/CCDC125 | 2 |
| MEblack | BP | GO:004578 | positive regulation of cell cycle | 3/238 | 45/16092 | 0.066666667 | 4.507563025 | 2.886857739 | 0.028741751 | 0.461473588 | 0.15629918 | DBF4B/ANAPC5/CCDC57 | 3 |
| MEblack | BP | GO:004477 | mitotic cell cycle phase transition | 5/238 | 116/16092 | 0.043103448 | 2.914372646 | 2.535315536 | 0.028993011 | 0.461473588 | 0.15629918 | KNTC1/MIIP/ANAPC5/CDI | 5 |
| MEblack | BP | GO:004501 | glycerolipid biosynthetic process | 5/238 | 116/16092 | 0.043103448 | 2.914372646 | 2.535315536 | 0.028993011 | 0.461473588 | 0.15629918 | DGAT1/PCK1/MPPE1/PIG | 5 |
| MEblack | MF | GO:003015 | signaling receptor complex adaptor activity | 2/245 | 18/16185 | 0.111111111 | 7.340136054 | 3.336582976 | 0.029770539 | 0.550774431 | 0.167421813 | FRS3/GNB3 | 2 |
| MEblack | CC | GO:009880 | plasma membrane signaling receptor complex | 4/230 | 81/15882 | 0.049382716 | 3.409983897 | 2.635920362 | 0.029987089 | 0.341656679 | 0.018978953 | TLR6/GABBR1/ITGA10/IT | 4 |

| | | | | | | | | | | | | | |
|---|---|---|---|---|---|---|---|---|---|---|---|---|---|
| MEblack | CC | GO:000566 | RNA polymerase III complex | 2/230 | 19/15882 | 0.105263158 | 7.268649886 | 3.314179862 | 0.030369483 | 0.341656679 | 0.018978953 | POLR3F/POLR3A | 2 |
| MEblack | BP | GO:001821 | peptidyl-tyrosine modification | 3/238 | 46/16092 | 0.065217391 | 4.409572525 | 2.837305002 | 0.030420489 | 0.461473588 | 0.15629918 | CLK4/CLK1/CLCF1 | 3 |
| MEblack | MF | GO:001662 | oxidoreductase activity, acting on the CH-CH group of donors | 3/245 | 45/16185 | 0.066666667 | 4.404081633 | 2.834890421 | 0.030497216 | 0.550774431 | 0.167421813 | ACAD9/PPOX/RSAD1 | 3 |
| MEblack | BP | GO:009911 | microtubule-based transport | 4/238 | 80/16092 | 0.05 | 3.380672269 | 2.615365466 | 0.030810701 | 0.461473588 | 0.15629918 | BICDL1/CCDC40/HAP1/T | 4 |
| MEblack | BP | GO:003003 | cell projection assembly | 8/238 | 248/16092 | 0.032258065 | 2.181078883 | 2.296586974 | 0.031506681 | 0.461473588 | 0.15629918 | CCDC66/CROCC/CCDC1 | 8 |
| MEblack | BP | GO:001943 | triglyceride biosynthetic process | 2/238 | 19/16092 | 0.105263158 | 7.117204777 | 3.268826847 | 0.031570406 | 0.461473588 | 0.15629918 | DGAT1/PCK1 | 2 |
| MEblack | BP | GO:004477 | mitotic DNA damage checkpoint signaling | 2/238 | 19/16092 | 0.105263158 | 7.117204777 | 3.268826847 | 0.031570406 | 0.461473588 | 0.15629918 | EME2/EME1 | 2 |
| MEblack | BP | GO:004646 | neutral lipid biosynthetic process | 2/238 | 19/16092 | 0.105263158 | 7.117204777 | 3.268826847 | 0.031570406 | 0.461473588 | 0.15629918 | DGAT1/PCK1 | 2 |
| MEblack | BP | GO:004646 | acylglycerol biosynthetic process | 2/238 | 19/16092 | 0.105263158 | 7.117204777 | 3.268826847 | 0.031570406 | 0.461473588 | 0.15629918 | DGAT1/PCK1 | 2 |
| MEblack | BP | GO:190198 | negative regulation of cell cycle phase transition | 4/238 | 81/16092 | 0.049382716 | 3.338935574 | 2.585604584 | 0.032048218 | 0.462744995 | 0.1567298 | KNTC1/EME2/EME1/MIIP | 4 |
| MEblack | MF | GO:014029 | molecular sensor activity | 2/245 | 19/16185 | 0.105263158 | 6.953813104 | 3.219233878 | 0.032946477 | 0.550774431 | 0.167421813 | MLH1/XRCC3 | 2 |
| MEblack | BP | GO:002240 | cell cycle process | 13/238 | 495/16092 | 0.026262626 | 1.775706646 | 2.147781736 | 0.033008646 | 0.464822807 | 0.157433546 | DBF4B/KNTC1/EME2/EM | 13 |
| MEblack | BP | GO:000995 | anterior/posterior pattern specification | 3/238 | 48/16092 | 0.0625 | 4.225840336 | 2.742317462 | 0.033929576 | 0.464822807 | 0.157433546 | HOXD4/HOXB6/HOXA5 | 3 |
| MEblack | BP | GO:000680 | superoxide metabolic process | 2/238 | 20/16092 | 0.1 | 6.761344538 | 3.158744164 | 0.034742272 | 0.464822807 | 0.157433546 | NOXO1/NOXA1 | 2 |
| MEblack | BP | GO:003129 | replication fork processing | 2/238 | 20/16092 | 0.1 | 6.761344538 | 3.158744164 | 0.034742272 | 0.464822807 | 0.157433546 | EME2/EME1 | 2 |
| MEblack | BP | GO:003301 | tetrapyrrole metabolic process | 2/238 | 20/16092 | 0.1 | 6.761344538 | 3.158744164 | 0.034742272 | 0.464822807 | 0.157433546 | PPOX/RSAD1 | 2 |
| MEblack | BP | GO:001605 | organic acid catabolic process | 5/238 | 122/16092 | 0.040983607 | 2.771042843 | 2.405841952 | 0.034940228 | 0.464822807 | 0.157433546 | AFMID/PCK1/AMT/FAAH/ | 5 |
| MEblack | BP | GO:004639 | carboxylic acid catabolic process | 5/238 | 122/16092 | 0.040983607 | 2.771042843 | 2.405841952 | 0.034940228 | 0.464822807 | 0.157433546 | AFMID/PCK1/AMT/FAAH/ | 5 |
| MEblack | BP | GO:003336 | protein localization to organelle | 8/238 | 254/16092 | 0.031496063 | 2.129557335 | 2.223235856 | 0.035531235 | 0.467433131 | 0.158317652 | KNTC1/SPATA7/ASB3/PE | 8 |
| MEblack | BP | GO:000639 | tRNA metabolic process | 6/238 | 165/16092 | 0.036363636 | 2.458670741 | 2.307505914 | 0.036144908 | 0.470281003 | 0.159282214 | TRMT44/PUS1/EXOSC7/ | 6 |
| MEblack | MF | GO:000817 | tRNA methyltransferase activity | 2/245 | 20/16185 | 0.1 | 6.606122449 | 3.110079863 | 0.036248464 | 0.550774431 | 0.167421813 | TRMT44/TRMT13 | 2 |
| MEblack | MF | GO:001663 | oxidoreductase activity, acting on the CH-NH2 group of donors | 2/245 | 20/16185 | 0.1 | 6.606122449 | 3.110079863 | 0.036248464 | 0.550774431 | 0.167421813 | LOXL3/AOC2 | 2 |
| MEblack | MF | GO:005153 | 4 iron, 4 sulfur cluster binding | 2/245 | 20/16185 | 0.1 | 6.606122449 | 3.110079863 | 0.036248464 | 0.550774431 | 0.167421813 | MUTYH/RSAD1 | 2 |
| MEblack | CC | GO:000092 | spindle pole | 3/230 | 51/15882 | 0.058823529 | 4.061892583 | 2.654842648 | 0.037537184 | 0.373905238 | 0.020770354 | PLK1/CEP95/AURKC | 3 |
| MEblack | BP | GO:014118 | nucleic acid catabolic process | 5/238 | 126/16092 | 0.03968254 | 2.683073229 | 2.323810937 | 0.039295061 | 0.505710344 | 0.171281984 | DNASE1/UPF3B/DNASE2 | 5 |
| MEblack | CC | GO:000581 | centriole | 3/230 | 52/15882 | 0.057692308 | 3.983779264 | 2.612437007 | 0.039422982 | 0.373905238 | 0.020770354 | CROCC/WDR90/CCDC57 | 3 |
| MEblack | CC | GO:000560 | basement membrane | 2/230 | 22/15882 | 0.090909091 | 6.277470356 | 3.00264143 | 0.039883225 | 0.373905238 | 0.020770354 | COL27A1/LAMB3 | 2 |
| MEblack | CC | GO:000568 | U12-type spliceosomal complex | 2/230 | 22/15882 | 0.090909091 | 6.277470356 | 3.00264143 | 0.039883225 | 0.373905238 | 0.020770354 | RBM41/RNPC3 | 2 |
| MEblack | MF | GO:014064 | catalytic activity, acting on a nucleic acid | 13/245 | 498/16185 | 0.026104418 | 1.724489796 | 2.035911121 | 0.040274702 | 0.550774431 | 0.167421813 | DNASE1/DDX39B/EME2/T | 13 |
| MEblack | BP | GO:003629 | interstrand cross-link repair | 2/238 | 22/16092 | 0.090909091 | 6.146676853 | 2.959653518 | 0.041435596 | 0.515915559 | 0.174738448 | FANCG/FANCA | 2 |
| MEblack | BP | GO:007238 | organelle transport along microtubule | 2/238 | 22/16092 | 0.090909091 | 6.146676853 | 2.959653518 | 0.041435596 | 0.515915559 | 0.174738448 | BICDL1/HAP1 | 2 |
| MEblack | BP | GO:001094 | negative regulation of cell cycle process | 4/238 | 88/16092 | 0.045454545 | 3.073338426 | 2.389504378 | 0.041520393 | 0.515915559 | 0.174738448 | KNTC1/EME2/EME1/MIIP | 4 |
| MEblack | MF | GO:001679 | hydrolase activity, acting on glycosyl bonds | 5/245 | 126/16185 | 0.03968254 | 2.621477162 | 2.265263244 | 0.042711704 | 0.550774431 | 0.167421813 | MUTYH/OVGP1/AMY2B/H | 5 |
| MEblack | BP | GO:004648 | glycerolipid metabolic process | 7/238 | 217/16092 | 0.032258065 | 2.181078883 | 2.146161865 | 0.042779284 | 0.515915559 | 0.174738448 | GDPD3/DGAT1/PCK1/MP | 7 |
| MEblack | MF | GO:001510 | chloride transmembrane transporter activity | 4/245 | 87/16185 | 0.045977011 | 3.037297678 | 2.362163603 | 0.043038514 | 0.550774431 | 0.167421813 | CLCN2/SLC12A9/GABRE/ | 4 |
| MEblack | MF | GO:001691 | GABA receptor activity | 2/245 | 22/16185 | 0.090909091 | 6.005565863 | 2.912630229 | 0.043212653 | 0.550774431 | 0.167421813 | GABBR1/GABRE | 2 |
| MEblack | BP | GO:000709 | mitotic cell cycle checkpoint signaling | 3/238 | 53/16092 | 0.056603774 | 3.827176153 | 2.525880087 | 0.04357549 | 0.515915559 | 0.174738448 | KNTC1/EME2/EME1 | 3 |
| MEblack | BP | GO:017004 | proteinogenic amino acid catabolic process | 3/238 | 53/16092 | 0.056603774 | 3.827176153 | 2.525880087 | 0.04357549 | 0.515915559 | 0.174738448 | AFMID/AMT/AMDHD1 | 3 |
| MEblack | MF | GO:001681 | hydrolase activity, acting on carbon-nitrogen (but not peptide) bonds | 4/245 | 88/16185 | 0.045454545 | 3.002782931 | 2.335525235 | 0.044573563 | 0.550774431 | 0.167421813 | AFMID/FAAH/OPLAH/AM | 4 |
| MEblack | BP | GO:000152 | retinoid metabolic process | 2/238 | 23/16092 | 0.086956522 | 5.879430033 | 2.869122932 | 0.04494885 | 0.515915559 | 0.174738448 | RDH13/RDH5 | 2 |
| MEblack | BP | GO:000703 | peroxisome organization | 2/238 | 23/16092 | 0.086956522 | 5.879430033 | 2.869122932 | 0.04494885 | 0.515915559 | 0.174738448 | PEX16/PEX10 | 2 |
| MEblack | BP | GO:003447 | snRNA 3'-end processing | 2/238 | 23/16092 | 0.086956522 | 5.879430033 | 2.869122932 | 0.04494885 | 0.515915559 | 0.174738448 | EXOSC7/INTS6L | 2 |
| MEblack | BP | GO:004500 | DNA-templated DNA replication maintenance of fidelity | 2/238 | 23/16092 | 0.086956522 | 5.879430033 | 2.869122932 | 0.04494885 | 0.515915559 | 0.174738448 | EME2/EME1 | 2 |
| MEblack | BP | GO:004677 | protein autophosphorylation | 2/238 | 23/16092 | 0.086956522 | 5.879430033 | 2.869122932 | 0.04494885 | 0.515915559 | 0.174738448 | ULK3/PIM2 | 2 |
| MEblack | BP | GO:004643 | organophosphate catabolic process | 4/238 | 91/16092 | 0.043956044 | 2.972019577 | 2.311367086 | 0.046014982 | 0.515915559 | 0.174738448 | GDPD3/GAPDHS/NUDT1 | 4 |
| MEblack | MF | GO:000848 | transaminase activity | 2/245 | 23/16185 | 0.086956522 | 5.744454303 | 2.822828239 | 0.046866152 | 0.550774431 | 0.167421813 | LOC104968656/PHYKPL | 2 |
| MEblack | MF | GO:001706 | snRNA binding | 2/245 | 23/16185 | 0.086956522 | 5.744454303 | 2.822828239 | 0.046866152 | 0.550774431 | 0.167421813 | RBM41/RNPC3 | 2 |
| MEblack | CC | GO:003606 | ciliary basal body | 3/230 | 56/15882 | 0.053571429 | 3.699223602 | 2.452817392 | 0.047436055 | 0.41972948 | 0.02331588 | SPATA7/DZIP1L/CFAP10 | 3 |
| MEblack | BP | GO:003009 | hemopoiesis | 4/238 | 92/16092 | 0.043478261 | 2.939715016 | 2.2860326 | 0.047571151 | 0.515915559 | 0.174738448 | EFNA2/IL23A/FANCA/THI | 4 |
| MEblack | MF | GO:000817 | RNA methyltransferase activity | 3/245 | 54/16185 | 0.055555556 | 3.670068027 | 2.436522236 | 0.048333624 | 0.550774431 | 0.167421813 | TRMT44/NSUN6/TRMT13 | 3 |
| MEblack | BP | GO:000024 | spliceosomal complex assembly | 2/238 | 24/16092 | 0.083333333 | 5.634453782 | 2.783773099 | 0.048567853 | 0.515915559 | 0.174738448 | LUC7L3/LUC7L | 2 |
| MEblack | BP | GO:000907 | aromatic amino acid metabolic process | 2/238 | 24/16092 | 0.083333333 | 5.634453782 | 2.783773099 | 0.048567853 | 0.515915559 | 0.174738448 | AFMID/AMDHD1 | 2 |
| MEblack | BP | GO:001610 | diterpenoid metabolic process | 2/238 | 24/16092 | 0.083333333 | 5.634453782 | 2.783773099 | 0.048567853 | 0.515915559 | 0.174738448 | RDH13/RDH5 | 2 |
| MEblack | BP | GO:001715 | regulation of calcium ion-dependent exocytosis | 2/238 | 24/16092 | 0.083333333 | 5.634453782 | 2.783773099 | 0.048567853 | 0.515915559 | 0.174738448 | SYT2/DOC2G | 2 |
| MEblack | BP | GO:003021 | T cell differentiation | 2/238 | 24/16092 | 0.083333333 | 5.634453782 | 2.783773099 | 0.048567853 | 0.515915559 | 0.174738448 | IL23A/FANCA | 2 |
| MEblack | BP | GO:004477 | mitotic DNA integrity checkpoint signaling | 2/238 | 24/16092 | 0.083333333 | 5.634453782 | 2.783773099 | 0.048567853 | 0.515915559 | 0.174738448 | EME2/EME1 | 2 |
| MEblack | BP | GO:190160 | alpha-amino acid catabolic process | 3/238 | 56/16092 | 0.053571429 | 3.62214886 | 2.408318389 | 0.049949312 | 0.515915559 | 0.174738448 | AFMID/AMT/AMDHD1 | 3 |

**Supplementary Table 19. Gene Ontology enrichment of the telomere length-associated MEgreen module in whole blood, related to Figure S16.**

| Module | ONTOLOGY | ID | Description | GeneRatio | BgRatio | RichFactor | FoldEnrichment | zScore | pvalue | p.adjust | qvalue | geneID | Count |
|---|---|---|---|---|---|---|---|---|---|---|---|---|---|
| MEgreen | BP | GO:000704 | cell cycle | 106/276 | 424/7644 | 0.25 | 6.923913043 | 24.29040001 | 1.91606E-64 | 1.61907E-61 | 1.61907E-61 | INCENP/CCNB1/MCM4/R | 106 |
| MEgreen | BP | GO:002240 | cell cycle process | 93/276 | 351/7644 | 0.264957265 | 7.338164251 | 23.52749031 | 3.24168E-58 | 1.36961E-55 | 1.36961E-55 | INCENP/CCNB1/MCM4/R | 93 |
| MEgreen | BP | GO:000027 | mitotic cell cycle | 79/276 | 253/7644 | 0.312252964 | 8.648049493 | 23.94258354 | 6.33519E-55 | 1.78441E-52 | 1.78441E-52 | INCENP/CCNB1/MCM4/R | 79 |
| MEgreen | BP | GO:190304 | mitotic cell cycle process | 64/276 | 197/7644 | 0.324873096 | 8.99757228 | 22.0096062 | 2.82243E-45 | 5.96238E-43 | 5.96238E-43 | INCENP/CCNB1/MCM4/R | 64 |
| MEgreen | BP | GO:000705 | chromosome segregation | 46/276 | 121/7644 | 0.380165289 | 10.52892562 | 20.44809026 | 5.90005E-36 | 9.97108E-34 | 9.97108E-34 | INCENP/CCNB1/BUB1B/S | 46 |
| MEgreen | BP | GO:000028 | nuclear division | 38/276 | 107/7644 | 0.355140187 | 9.83583909 | 17.81361623 | 1.87901E-28 | 2.64628E-26 | 2.64628E-26 | INCENP/CCNB1/RAD51/B | 38 |
| MEgreen | BP | GO:004828 | organelle fission | 39/276 | 120/7644 | 0.325 | 9.001086957 | 17.09727441 | 1.68332E-27 | 2.03201E-25 | 2.03201E-25 | INCENP/CCNB1/RAD51/B | 39 |
| MEgreen | CC | GO:000569 | chromosome | 67/276 | 398/7449 | 0.168341709 | 4.543396329 | 14.25147677 | 2.27415E-27 | 3.41122E-25 | 3.41122E-25 | INCENP/RAD51/SKA1/MA | 67 |
| MEgreen | BP | GO:000626 | DNA replication | 34/276 | 88/7644 | 0.386363636 | 10.70059289 | 17.71353754 | 5.83506E-27 | 6.16328E-25 | 6.16328E-25 | MCM4/RAD51/RFC3/MGM | 34 |
| MEgreen | BP | GO:009881 | nuclear chromosome segregation | 33/276 | 84/7644 | 0.392857143 | 10.88043478 | 17.62245716 | 1.8073E-26 | 1.69685E-24 | 1.69685E-24 | INCENP/CCNB1/BUB1B/M | 33 |
| MEgreen | BP | GO:000626 | DNA-templated DNA replication | 30/276 | 67/7644 | 0.447761194 | 12.40103829 | 18.140315 | 3.39137E-26 | 2.86571E-24 | 2.86571E-24 | MCM4/RAD51/RFC3/MGM | 30 |
| MEgreen | CC | GO:000077 | chromosome, centromeric region | 28/276 | 65/7449 | 0.430769231 | 11.62608696 | 16.87750166 | 1.25527E-23 | 9.41452E-22 | 9.41452E-22 | INCENP/SKA1/MAD2L1/C | 28 |
| MEgreen | BP | GO:000625 | DNA metabolic process | 54/276 | 305/7644 | 0.17704918 | 4.903492516 | 13.46469403 | 1.49139E-23 | 1.14566E-21 | 1.14566E-21 | MCM4/RAD51/RFC3/ANK | 54 |
| MEgreen | BP | GO:005127 | chromosome organization | 39/276 | 151/7644 | 0.258278146 | 7.153181687 | 14.77991745 | 2.42444E-23 | 1.70721E-21 | 1.70721E-21 | CCNB1/RAD51/BUB1B/M | 39 |
| MEgreen | CC | GO:009868 | chromosomal region | 28/276 | 73/7449 | 0.383561644 | 10.35199524 | 15.7499512 | 6.01563E-22 | 3.00781E-20 | 3.00781E-20 | INCENP/SKA1/MAD2L1/C | 28 |
| MEgreen | BP | GO:014001 | mitotic nuclear division | 26/276 | 70/7644 | 0.371428571 | 10.28695652 | 15.10676398 | 2.6385E-20 | 1.71503E-18 | 1.71503E-18 | CCNB1/BUB1B/MAD2L1/ | 26 |
| MEgreen | BP | GO:004477 | cell cycle phase transition | 31/276 | 113/7644 | 0.274336283 | 7.597922278 | 13.67513919 | 1.49844E-19 | 9.04418E-18 | 9.04418E-18 | CCNB1/BUB1B/MAD2L1/ | 31 |
| MEgreen | BP | GO:004477 | mitotic cell cycle phase transition | 27/276 | 82/7644 | 0.329268293 | 9.119300106 | 14.3060206 | 1.87716E-19 | 1.05747E-17 | 1.05747E-17 | CCNB1/BUB1B/MAD2L1/ | 27 |
| MEgreen | BP | GO:000081 | sister chromatid segregation | 24/276 | 63/7644 | 0.380952381 | 10.55072464 | 14.7317728 | 3.96626E-19 | 2.09468E-17 | 2.09468E-17 | CCNB1/BUB1B/MAD2L1/ | 24 |
| MEgreen | BP | GO:005172 | regulation of cell cycle | 38/276 | 184/7644 | 0.206521739 | 5.719754253 | 12.5421083 | 4.45772E-19 | 2.21575E-17 | 2.21575E-17 | BUB1B/MAD2L1/MKI67/C | 38 |
| MEgreen | BP | GO:001056 | regulation of cell cycle process | 32/276 | 134/7644 | 0.23880597 | 6.613887086 | 12.68842644 | 3.54624E-18 | 1.66476E-16 | 1.66476E-16 | BUB1B/MAD2L1/MKI67/C | 32 |
| MEgreen | BP | GO:000007 | mitotic sister chromatid segregation | 22/276 | 56/7644 | 0.392857143 | 10.88043478 | 14.36210411 | 5.73953E-18 | 2.55258E-16 | 2.55258E-16 | CCNB1/BUB1B/MAD2L1/ | 22 |
| MEgreen | CC | GO:000079 | condensed chromosome | 23/276 | 65/7449 | 0.353846154 | 9.55 | 13.58003561 | 2.74061E-17 | 1.02773E-15 | 1.02773E-15 | INCENP/RAD51/SKA1/MA | 23 |
| MEgreen | CC | GO:000077 | kinetochore | 20/276 | 48/7449 | 0.416666667 | 11.24547101 | 13.96790605 | 7.91827E-17 | 2.37548E-15 | 2.37548E-15 | INCENP/SKA1/MAD2L1/C | 20 |
| MEgreen | CC | GO:000077 | condensed chromosome, centromeric region | 20/276 | 50/7449 | 0.4 | 10.79565217 | 13.63188155 | 2.0857E-16 | 5.21426E-15 | 5.21426E-15 | INCENP/SKA1/MAD2L1/C | 20 |
| MEgreen | BP | GO:000697 | DNA damage response | 42/276 | 277/7644 | 0.151624549 | 4.199340763 | 10.49703659 | 7.87584E-16 | 3.32754E-14 | 3.32754E-14 | MCM4/RAD51/CLSPN/RF | 42 |
| MEgreen | BP | GO:000628 | DNA repair | 36/276 | 212/7644 | 0.169811321 | 4.703035275 | 10.58241019 | 2.99089E-15 | 1.20348E-13 | 1.20348E-13 | MCM4/RAD51/RFC3/ANK | 36 |
| MEgreen | BP | GO:000734 | regulation of mitotic cell cycle | 24/276 | 90/7644 | 0.266666667 | 7.385507246 | 11.79342172 | 4.64602E-15 | 1.78449E-13 | 1.78449E-13 | BUB1B/MAD2L1/MKI67/C | 24 |
| MEgreen | BP | GO:000627 | DNA replication initiation | 13/276 | 20/7644 | 0.65 | 18.00217391 | 14.73464478 | 8.29848E-15 | 3.04879E-13 | 3.04879E-13 | MCM4/MCM6/MCM10/PR | 13 |
| MEgreen | BP | GO:190304 | meiotic cell cycle process | 19/276 | 54/7644 | 0.351851852 | 9.744766506 | 12.48061172 | 1.25197E-14 | 4.40799E-13 | 4.40799E-13 | INCENP/RAD51/ANKLE1/ | 19 |
| MEgreen | BP | GO:000631 | DNA recombination | 25/276 | 106/7644 | 0.235849057 | 6.531993437 | 11.09987747 | 2.75256E-14 | 9.30366E-13 | 9.30366E-13 | MCM4/RAD51/ANKLE1/M | 25 |
| MEgreen | BP | GO:003355 | cellular response to stress | 48/276 | 392/7644 | 0.12244898 | 3.391304348 | 9.407225938 | 2.97106E-14 | 9.65594E-13 | 9.65594E-13 | MCM4/RAD51/CLSPN/P4 | 48 |
| MEgreen | MF | GO:014009 | catalytic activity, acting on DNA | 28/251 | 156/7669 | 0.179487179 | 5.484012667 | 10.40775197 | 9.23266E-14 | 1.52341E-11 | 1.52341E-11 | MCM4/RAD51/RFC3/ANK | 28 |
| MEgreen | MF | GO:014065 | ATP-dependent activity | 40/251 | 320/7669 | 0.125 | 3.819223108 | 9.475994724 | 1.06161E-13 | 1.52341E-11 | 1.52341E-11 | KIFC1/MCM4/RAD51/RFC | 40 |
| MEgreen | BP | GO:005132 | meiotic cell cycle | 19/276 | 61/7644 | 0.31147541 | 8.626514612 | 11.57396483 | 1.61162E-13 | 5.04377E-12 | 5.04377E-12 | INCENP/RAD51/ANKLE1/ | 19 |
| MEgreen | BP | GO:190198 | regulation of cell cycle phase transition | 22/276 | 89/7644 | 0.247191011 | 6.846116268 | 10.73635541 | 3.69201E-13 | 1.1142E-11 | 1.1142E-11 | BUB1B/MAD2L1/CLSPN/S | 22 |
| MEgreen | BP | GO:190199 | regulation of mitotic cell cycle phase transition | 18/276 | 58/7644 | 0.310344828 | 8.595202399 | 11.23721244 | 7.69576E-13 | 2.24239E-11 | 2.24239E-11 | BUB1B/MAD2L1/CLSPN/S | 18 |
| MEgreen | BP | GO:003326 | nuclear DNA replication | 10/276 | 13/7644 | 0.769230769 | 21.30434783 | 14.18017714 | 8.34143E-13 | 2.27371E-11 | 2.27371E-11 | MCM4/RAD51/MCM6/LIG | 10 |
| MEgreen | BP | GO:004478 | cell cycle DNA replication | 10/276 | 13/7644 | 0.769230769 | 21.30434783 | 14.18017714 | 8.34143E-13 | 2.27371E-11 | 2.27371E-11 | MCM4/RAD51/MCM6/LIG | 10 |
| MEgreen | BP | GO:190296 | mitotic DNA replication | 9/276 | 10/7644 | 0.9 | 24.92608696 | 14.65234923 | 8.89643E-13 | 2.34921E-11 | 2.34921E-11 | MCM4/RAD51/MCM6/LIG | 9 |
| MEgreen | BP | GO:000072 | recombinational repair | 19/276 | 69/7644 | 0.275362319 | 7.626339004 | 10.70085649 | 1.93382E-12 | 4.95174E-11 | 4.95174E-11 | MCM4/RAD51/ANKLE1/M | 19 |
| MEgreen | BP | GO:000705 | spindle organization | 19/276 | 71/7644 | 0.267605634 | 7.411512554 | 10.50430577 | 3.39532E-12 | 8.43837E-11 | 8.43837E-11 | INCENP/HAUS4/TPX2/KIF | 19 |
| MEgreen | CC | GO:000581 | spindle | 21/276 | 87/7449 | 0.24137931 | 6.514617691 | 10.14848208 | 3.47734E-12 | 7.45144E-11 | 7.45144E-11 | INCENP/SKA1/TPX2/KNS | 21 |
| MEgreen | BP | GO:014001 | meiotic nuclear division | 15/276 | 41/7644 | 0.365853659 | 10.13255567 | 11.34758446 | 4.32383E-12 | 1.0439E-10 | 1.0439E-10 | INCENP/RAD51/ANKLE1/ | 15 |
| MEgreen | MF | GO:000809 | ATP-dependent activity, acting on DNA | 19/251 | 83/7669 | 0.228915663 | 6.994239908 | 10.09953192 | 1.21313E-11 | 1.16056E-09 | 1.16056E-09 | MCM4/RAD51/RFC3/MCM | 19 |
| MEgreen | BP | GO:000072 | double-strand break repair via homologous recombination | 18/276 | 68/7644 | 0.264705882 | 7.331202046 | 10.14921164 | 1.57207E-11 | 3.69E-10 | 3.69E-10 | MCM4/RAD51/ANKLE1/M | 18 |
| MEgreen | BP | GO:005131 | metaphase chromosome alignment | 10/276 | 18/7644 | 0.555555556 | 15.38647343 | 11.82643536 | 1.08665E-10 | 2.48167E-09 | 2.48167E-09 | INCENP/CCNB1/NDC80/K | 10 |
| MEgreen | CC | GO:009908 | supramolecular complex | 35/276 | 282/7449 | 0.124113475 | 3.34971477 | 7.890326206 | 1.83869E-10 | 3.44755E-09 | 3.44755E-09 | INCENP/SKA1/MAD2L1/C | 35 |
| MEgreen | BP | GO:000007 | cell cycle checkpoint signaling | 15/276 | 52/7644 | 0.288461538 | 7.989130435 | 9.787198124 | 2.13457E-10 | 4.74661E-09 | 4.74661E-09 | BUB1B/MAD2L1/CLSPN/S | 15 |
| MEgreen | BP | GO:005000 | chromosome localization | 10/276 | 19/7644 | 0.526315789 | 14.57665904 | 11.46730728 | 2.22146E-10 | 4.81317E-09 | 4.81317E-09 | INCENP/CCNB1/NDC80/K | 10 |
| MEgreen | MF | GO:012054 | macromolecular conformation isomerase activity | 22/251 | 134/7669 | 0.164179104 | 5.016293037 | 8.627215799 | 2.9148E-10 | 1.93108E-08 | 1.93108E-08 | KIFC1/MCM4/CENPE/KIF | 22 |
| MEgreen | BP | GO:000008 | G2/M transition of mitotic cell cycle | 11/276 | 25/7644 | 0.44 | 12.18608696 | 10.84203665 | 3.17338E-10 | 6.70377E-09 | 6.70377E-09 | CLSPN/CDC25A/FOXM1/ | 11 |
| MEgreen | MF | GO:000377 | microtubule motor activity | 12/251 | 34/7669 | 0.352941176 | 10.78368877 | 10.51652032 | 3.36426E-10 | 1.93108E-08 | 1.93108E-08 | KIFC1/CENPE/KIF4A/KIF | 12 |
| MEgreen | CC | GO:000094 | outer kinetochore | 8/276 | 11/7449 | 0.727272727 | 19.6284585 | 12.12743997 | 4.81683E-10 | 8.02805E-09 | 8.02805E-09 | SKA1/NDC80/SPDL1/NU | 8 |
| MEgreen | BP | GO:000701 | microtubule-based process | 35/276 | 300/7644 | 0.116666667 | 3.23115942 | 7.63021118 | 5.15568E-10 | 1.06257E-08 | 1.06257E-08 | INCENP/KIFC1/SKA1/HA | 35 |
| MEgreen | BP | GO:000705 | mitotic spindle organization | 14/276 | 47/7644 | 0.29787234 | 8.249768733 | 9.648586192 | 5.40509E-10 | 1.08745E-08 | 1.08745E-08 | TPX2/KIF4A/RACGAP1/P | 14 |
| MEgreen | BP | GO:190199 | negative regulation of mitotic cell cycle phase transition | 12/276 | 33/7644 | 0.363636364 | 10.07114625 | 10.10671316 | 6.97868E-10 | 1.37056E-08 | 1.37056E-08 | BUB1B/MAD2L1/CLSPN/S | 12 |
| MEgreen | BP | GO:001094 | negative regulation of cell cycle process | 16/276 | 65/7644 | 0.246153846 | 6.817391304 | 9.115713442 | 7.13663E-10 | 1.37056E-08 | 1.37056E-08 | BUB1B/MAD2L1/CLSPN/S | 16 |

| | | | | | | | | | | | | | |
|---|---|---|---|---|---|---|---|---|---|---|---|---|---|
| MEgreen | MF | GO:001685 | isomerase activity | 27/251 | 209/7669 | 0.129186603 | 3.947139671 | 7.94582409 | 7.14643E-10 | 3.41838E-08 | 3.41838E-08 | KIFC1/MCM4/P4HB/CENF | 27 |
| MEgreen | BP | GO:005178 | regulation of nuclear division | 11/276 | 27/7644 | 0.407407407 | 11.28341385 | 10.35950205 | 8.70171E-10 | 1.63399E-08 | 1.63399E-08 | BUB1B/MAD2L1/MKI67/SI | 11 |
| MEgreen | CC | GO:001563 | microtubule cytoskeleton | 37/276 | 329/7449 | 0.112462006 | 3.035251751 | 7.406278669 | 9.17503E-10 | 1.37625E-08 | 1.37625E-08 | INCENP/SKA1/HAUS4/TP | 37 |
| MEgreen | BP | GO:004578 | negative regulation of cell cycle | 17/276 | 76/7644 | 0.223684211 | 6.195080092 | 8.808868849 | 1.03532E-09 | 1.90184E-08 | 1.90184E-08 | BUB1B/MAD2L1/CLSPN/S | 17 |
| MEgreen | BP | GO:190285 | microtubule cytoskeleton organization involved in mitosis | 15/276 | 58/7644 | 0.25862069 | 7.162668666 | 9.117757944 | 1.16557E-09 | 2.09554E-08 | 2.09554E-08 | TPX2/KIF4A/SPDL1/RAC( | 15 |
| MEgreen | BP | GO:001995 | sexual reproduction | 20/276 | 108/7644 | 0.185185185 | 5.128824477 | 8.363335435 | 1.20082E-09 | 2.11395E-08 | 2.11395E-08 | INCENP/RAD51/ANKLE1/( | 20 |
| MEgreen | BP | GO:001038 | regulation of G2/M transition of mitotic cell cycle | 10/276 | 22/7644 | 0.454545455 | 12.58893281 | 10.53493779 | 1.4121E-09 | 2.43516E-08 | 2.43516E-08 | CLSPN/CDC25A/CENPF/T | 10 |
| MEgreen | BP | GO:004483 | cell cycle G2/M phase transition | 11/276 | 29/7644 | 0.379310345 | 10.50524738 | 9.925196902 | 2.16473E-09 | 3.65839E-08 | 3.65839E-08 | CLSPN/CDC25A/FOXM1/( | 11 |
| MEgreen | BP | GO:190198 | negative regulation of cell cycle phase transition | 15/276 | 61/7644 | 0.245901639 | 6.810406272 | 8.817846851 | 2.50617E-09 | 4.15238E-08 | 4.15238E-08 | BUB1B/MAD2L1/CLSPN/S | 15 |
| MEgreen | CC | GO:000565 | replication fork | 11/276 | 29/7449 | 0.379310345 | 10.23725637 | 9.776063259 | 2.83276E-09 | 3.86285E-08 | 3.86285E-08 | RFC3/MMS22L/XRCC2/W | 11 |
| MEgreen | BP | GO:000709 | mitotic cell cycle checkpoint signaling | 12/276 | 37/7644 | 0.324324324 | 8.982373678 | 9.419717332 | 3.2008E-09 | 5.2013E-08 | 5.2013E-08 | BUB1B/MAD2L1/CLSPN/S | 12 |
| MEgreen | CC | GO:000022 | nuclear chromosome | 16/276 | 70/7449 | 0.228571429 | 6.168944099 | 8.522677596 | 3.34898E-09 | 4.18622E-08 | 4.18622E-08 | RAD51/SMARCB1/MMS22 | 16 |
| MEgreen | BP | GO:000630 | double-strand break repair | 19/276 | 103/7644 | 0.184466019 | 5.108906712 | 8.125350151 | 3.38835E-09 | 5.40217E-08 | 5.40217E-08 | MCM4/RAD51/ANKLE1/M( | 19 |
| MEgreen | BP | GO:002241 | reproductive process | 20/276 | 115/7644 | 0.173913043 | 4.816635161 | 7.98128062 | 3.78939E-09 | 5.9297E-08 | 5.9297E-08 | INCENP/RAD51/ANKLE1/( | 20 |
| MEgreen | BP | GO:000708 | regulation of mitotic nuclear division | 10/276 | 24/7644 | 0.416666667 | 11.53985507 | 10.00862544 | 4.01636E-09 | 6.17058E-08 | 6.17058E-08 | BUB1B/MAD2L1/MKI67/SI | 10 |
| MEgreen | BP | GO:005198 | regulation of chromosome segregation | 10/276 | 25/7644 | 0.4 | 11.07826087 | 9.768283949 | 6.48237E-09 | 9.78143E-08 | 9.78143E-08 | BUB1B/MAD2L1/MKI67/SI | 10 |
| MEgreen | BP | GO:005138 | kinetochore organization | 7/276 | 10/7644 | 0.7 | 19.38695652 | 11.26018261 | 8.11738E-09 | 1.20337E-07 | 1.20337E-07 | CENPT/CENPH/NUF2/DL( | 7 |
| MEgreen | MF | GO:000801 | microtubule binding | 20/251 | 133/7669 | 0.15037594 | 4.594554114 | 7.691922503 | 9.40478E-09 | 3.40925E-07 | 3.40925E-07 | KIFC1/SKA1/HAUS4/CEN | 20 |
| MEgreen | MF | GO:012054 | polypeptide conformation or assembly isomerase activity | 13/251 | 53/7669 | 0.245283019 | 7.494324588 | 8.726557537 | 9.50314E-09 | 3.40925E-07 | 3.40925E-07 | KIFC1/CENPE/KIF4A/KIF2 | 13 |
| MEgreen | BP | GO:190274 | regulation of cell cycle G2/M phase transition | 10/276 | 26/7644 | 0.384615385 | 10.65217391 | 9.541199802 | 1.0201E-08 | 1.48618E-07 | 1.48618E-07 | CLSPN/CDC25A/CENPF/T | 10 |
| MEgreen | MF | GO:001711 | single-stranded DNA helicase activity | 7/251 | 11/7669 | 0.636363636 | 19.44331764 | 11.25932113 | 1.09277E-08 | 3.48471E-07 | 3.48471E-07 | MCM4/MCM6/MCM2/CHT | 7 |
| MEgreen | BP | GO:005178 | negative regulation of nuclear division | 8/276 | 15/7644 | 0.533333333 | 14.77101449 | 10.33211948 | 1.34932E-08 | 1.93251E-07 | 1.93251E-07 | BUB1B/MAD2L1/SPDL1/T | 8 |
| MEgreen | BP | GO:006500 | protein-DNA complex assembly | 14/276 | 59/7644 | 0.237288136 | 6.571849668 | 8.314942798 | 1.42404E-08 | 2.00553E-07 | 2.00553E-07 | RAD51/CENPT/CENPN/H. | 14 |
| MEgreen | MF | GO:001563 | tubulin binding | 21/251 | 154/7669 | 0.136363636 | 4.166425208 | 7.301290036 | 2.35723E-08 | 6.76526E-07 | 6.76526E-07 | KIFC1/SKA1/HAUS4/CEN | 21 |
| MEgreen | BP | GO:006198 | meiosis I cell cycle process | 11/276 | 36/7644 | 0.305555556 | 8.462560386 | 8.68591388 | 2.99935E-08 | 4.15483E-07 | 4.15483E-07 | RAD51/ANKLE1/CDC25A/ | 11 |
| MEgreen | BP | GO:000022 | microtubule cytoskeleton organization | 26/276 | 212/7644 | 0.122641509 | 3.396636587 | 6.849027772 | 3.63715E-08 | 4.95709E-07 | 4.95709E-07 | INCENP/SKA1/HAUS4/TP | 26 |
| MEgreen | BP | GO:007182 | protein-DNA complex organization | 14/276 | 64/7644 | 0.21875 | 6.058423913 | 7.864701942 | 4.35569E-08 | 5.84216E-07 | 5.84216E-07 | RAD51/CENPT/CENPN/H. | 14 |
| MEgreen | MF | GO:000377 | cytoskeletal motor activity | 12/251 | 50/7669 | 0.24 | 7.332908367 | 8.263684309 | 4.66985E-08 | 1.21841E-06 | 1.21841E-06 | KIFC1/CENPE/KIF4A/KIF2 | 12 |
| MEgreen | CC | GO:000030 | cyclin-dependent protein kinase holoenzyme complex | 10/276 | 30/7449 | 0.333333333 | 8.996376812 | 8.608057365 | 6.46088E-08 | 7.45486E-07 | 7.45486E-07 | CCNB1/CCNF/CCNB3/CK | 10 |
| MEgreen | BP | GO:004593 | negative regulation of mitotic cell cycle | 12/276 | 47/7644 | 0.255319149 | 7.071230342 | 8.080090746 | 6.53195E-08 | 8.62421E-07 | 8.62421E-07 | BUB1B/MAD2L1/CLSPN/S | 12 |
| MEgreen | BP | GO:005130 | chromosome separation | 8/276 | 21/7644 | 0.380952381 | 10.55072464 | 8.481929797 | 3.52987E-07 | 4.58883E-06 | 4.58883E-06 | BUB1B/MAD2L1/SPDL1/C | 8 |
| MEgreen | MF | GO:001653 | cyclin-dependent protein serine/threonine kinase regulator activity | 9/251 | 31/7669 | 0.290322581 | 8.870453669 | 8.076538822 | 4.00763E-07 | 8.84762E-06 | 8.84762E-06 | CCNB1/CCNF/CCNB3/CK | 9 |
| MEgreen | MF | GO:001991 | cyclin-dependent protein kinase regulator activity | 9/251 | 31/7669 | 0.290322581 | 8.870453669 | 8.076538822 | 4.00763E-07 | 8.84762E-06 | 8.84762E-06 | CCNB1/CCNF/CCNB3/CK | 9 |
| MEgreen | BP | GO:005130 | cell division | 13/276 | 66/7644 | 0.196969697 | 5.455204216 | 7.035159456 | 4.91645E-07 | 6.29454E-06 | 6.29454E-06 | INCENP/SKA1/ECT2/RAC | 13 |
| MEgreen | BP | GO:003304 | regulation of sister chromatid segregation | 8/276 | 22/7644 | 0.363636364 | 10.07114625 | 8.246139911 | 5.37456E-07 | 6.77836E-06 | 6.77836E-06 | BUB1B/MAD2L1/SPDL1/C | 8 |
| MEgreen | CC | GO:000587 | microtubule | 16/276 | 99/7449 | 0.161616162 | 4.361879666 | 6.605109956 | 5.78587E-07 | 6.19915E-06 | 6.19915E-06 | SKA1/TPX2/TUBA1B/TUB | 16 |
| MEgreen | BP | GO:000712 | meiosis I | 9/276 | 30/7644 | 0.3 | 8.308695652 | 7.762573597 | 6.72441E-07 | 8.35607E-06 | 8.35607E-06 | RAD51/ANKLE1/CENPS/M | 9 |
| MEgreen | MF | GO:001688 | ATP hydrolysis activity | 18/251 | 142/7669 | 0.126760563 | 3.873014982 | 6.356317808 | 7.47032E-07 | 1.53141E-05 | 1.53141E-05 | MCM4/RAD51/RFC3/KIF2 | 18 |
| MEgreen | CC | GO:005123 | spindle midzone | 6/276 | 11/7449 | 0.545454545 | 14.72134387 | 8.932825774 | 9.68521E-07 | 9.68521E-06 | 9.68521E-06 | INCENP/RACGAP1/CDCA | 6 |
| MEgreen | BP | GO:001096 | regulation of mitotic sister chromatid separation | 7/276 | 17/7644 | 0.411764706 | 11.40409207 | 8.311194322 | 1.05723E-06 | 1.29473E-05 | 1.29473E-05 | BUB1B/MAD2L1/SPDL1/C | 7 |
| MEgreen | BP | GO:004481 | mitotic G2/M transition checkpoint | 6/276 | 12/7644 | 0.5 | 13.84782609 | 8.620108924 | 1.61521E-06 | 1.94979E-05 | 1.94979E-05 | CLSPN/TICRR/CDK1/CDC | 6 |
| MEgreen | BP | GO:005130 | mitotic sister chromatid separation | 7/276 | 18/7644 | 0.388888889 | 10.7705314 | 8.031889022 | 1.67691E-06 | 1.96804E-05 | 1.96804E-05 | BUB1B/MAD2L1/SPDL1/C | 7 |
| MEgreen | BP | GO:190581 | regulation of chromosome separation | 7/276 | 18/7644 | 0.388888889 | 10.7705314 | 8.031889022 | 1.67691E-06 | 1.96804E-05 | 1.96804E-05 | BUB1B/MAD2L1/SPDL1/C | 7 |
| MEgreen | MF | GO:000453 | DNA nuclease activity | 9/251 | 37/7669 | 0.243243243 | 7.432001723 | 7.213770668 | 2.08175E-06 | 3.98308E-05 | 3.98308E-05 | ANKLE1/MGME1/POLE/FI | 9 |
| MEgreen | BP | GO:003007 | regulation of mitotic metaphase/anaphase transition | 7/276 | 19/7644 | 0.368421053 | 10.20366133 | 7.773724908 | 2.57364E-06 | 2.79338E-05 | 2.79338E-05 | BUB1B/MAD2L1/SPDL1/C | 7 |
| MEgreen | BP | GO:190209 | regulation of metaphase/anaphase transition of cell cycle | 7/276 | 19/7644 | 0.368421053 | 10.20366133 | 7.773724908 | 2.57364E-06 | 2.79338E-05 | 2.79338E-05 | BUB1B/MAD2L1/SPDL1/C | 7 |
| MEgreen | BP | GO:000709 | mitotic spindle assembly checkpoint signaling | 6/276 | 13/7644 | 0.461538462 | 12.7826087 | 8.228753758 | 2.90909E-06 | 2.79338E-05 | 2.79338E-05 | BUB1B/MAD2L1/SPDL1/B | 6 |
| MEgreen | BP | GO:003157 | spindle checkpoint signaling | 6/276 | 13/7644 | 0.461538462 | 12.7826087 | 8.228753758 | 2.90909E-06 | 2.79338E-05 | 2.79338E-05 | BUB1B/MAD2L1/SPDL1/B | 6 |
| MEgreen | BP | GO:003304 | negative regulation of sister chromatid segregation | 6/276 | 13/7644 | 0.461538462 | 12.7826087 | 8.228753758 | 2.90909E-06 | 2.79338E-05 | 2.79338E-05 | BUB1B/MAD2L1/SPDL1/B | 6 |
| MEgreen | BP | GO:003304 | regulation of mitotic sister chromatid segregation | 6/276 | 13/7644 | 0.461538462 | 12.7826087 | 8.228753758 | 2.90909E-06 | 2.79338E-05 | 2.79338E-05 | BUB1B/MAD2L1/SPDL1/B | 6 |
| MEgreen | BP | GO:003304 | negative regulation of mitotic sister chromatid segregation | 6/276 | 13/7644 | 0.461538462 | 12.7826087 | 8.228753758 | 2.90909E-06 | 2.79338E-05 | 2.79338E-05 | BUB1B/MAD2L1/SPDL1/B | 6 |
| MEgreen | BP | GO:004513 | meiotic chromosome segregation | 6/276 | 13/7644 | 0.461538462 | 12.7826087 | 8.228753758 | 2.90909E-06 | 2.79338E-05 | 2.79338E-05 | NUF2/PTTG1/SGO1/ESPL | 6 |

| | | | | | | | | | | | | | |
|---|---|---|---|---|---|---|---|---|---|---|---|---|---|
| MEgreen | BP | GO:004583 | negative regulation of mitotic nuclear division | 6/276 | 13/7644 | 0.461538462 | 12.7826087 | 8.228753758 | 2.90909E-06 | 2.79338E-05 | 2.79338E-05 | BUB1B/MAD2L1/SPDL1/B | 6 |
| MEgreen | BP | GO:004584 | negative regulation of mitotic metaphase/anaphase transition | 6/276 | 13/7644 | 0.461538462 | 12.7826087 | 8.228753758 | 2.90909E-06 | 2.79338E-05 | 2.79338E-05 | BUB1B/MAD2L1/SPDL1/B | 6 |
| MEgreen | BP | GO:005198 | negative regulation of chromosome segregation | 6/276 | 13/7644 | 0.461538462 | 12.7826087 | 8.228753758 | 2.90909E-06 | 2.79338E-05 | 2.79338E-05 | BUB1B/MAD2L1/SPDL1/B | 6 |
| MEgreen | BP | GO:007117 | spindle assembly checkpoint signaling | 6/276 | 13/7644 | 0.461538462 | 12.7826087 | 8.228753758 | 2.90909E-06 | 2.79338E-05 | 2.79338E-05 | BUB1B/MAD2L1/SPDL1/B | 6 |
| MEgreen | BP | GO:007117 | mitotic spindle checkpoint signaling | 6/276 | 13/7644 | 0.461538462 | 12.7826087 | 8.228753758 | 2.90909E-06 | 2.79338E-05 | 2.79338E-05 | BUB1B/MAD2L1/SPDL1/B | 6 |
| MEgreen | BP | GO:190210 | negative regulation of metaphase/anaphase transition of cell | 6/276 | 13/7644 | 0.461538462 | 12.7826087 | 8.228753758 | 2.90909E-06 | 2.79338E-05 | 2.79338E-05 | BUB1B/MAD2L1/SPDL1/B | 6 |
| MEgreen | BP | GO:190581 | negative regulation of chromosome separation | 6/276 | 13/7644 | 0.461538462 | 12.7826087 | 8.228753758 | 2.90909E-06 | 2.79338E-05 | 2.79338E-05 | BUB1B/MAD2L1/SPDL1/B | 6 |
| MEgreen | BP | GO:200081 | negative regulation of mitotic sister chromatid separation | 6/276 | 13/7644 | 0.461538462 | 12.7826087 | 8.228753758 | 2.90909E-06 | 2.79338E-05 | 2.79338E-05 | BUB1B/MAD2L1/SPDL1/B | 6 |
| MEgreen | BP | GO:000632 | chromatin organization | 18/276 | 142/7644 | 0.126760563 | 3.510716473 | 5.844737038 | 3.08646E-06 | 2.9304E-05 | 2.9304E-05 | SMARCB1/EZH2/CENPN/ | 18 |
| MEgreen | MF | GO:000367 | DNA helicase activity | 8/251 | 31/7669 | 0.258064516 | 7.884847706 | 7.065125102 | 4.83375E-06 | 8.67054E-05 | 8.67054E-05 | MCM4/MCM6/MCM2/CHT | 8 |
| MEgreen | CC | GO:003198 | nuclear lumen | 37/276 | 459/7449 | 0.080610022 | 2.175594392 | 5.099753918 | 5.15629E-06 | 4.83402E-05 | 4.83402E-05 | RAD51/MAD2L1/SMARCE | 37 |
| MEgreen | BP | GO:000709 | metaphase/anaphase transition of mitotic cell cycle | 7/276 | 21/7644 | 0.333333333 | 9.231884058 | 7.310676972 | 5.58052E-06 | 4.96373E-05 | 4.96373E-05 | BUB1B/MAD2L1/SPDL1/C | 7 |
| MEgreen | BP | GO:000713 | reciprocal meiotic recombination | 7/276 | 21/7644 | 0.333333333 | 9.231884058 | 7.310676972 | 5.58052E-06 | 4.96373E-05 | 4.96373E-05 | RAD51/ANKLE1/CENPS/M | 7 |
| MEgreen | BP | GO:003582 | homologous recombination | 7/276 | 21/7644 | 0.333333333 | 9.231884058 | 7.310676972 | 5.58052E-06 | 4.96373E-05 | 4.96373E-05 | RAD51/ANKLE1/CENPS/M | 7 |
| MEgreen | BP | GO:004478 | metaphase/anaphase transition of cell cycle | 7/276 | 21/7644 | 0.333333333 | 9.231884058 | 7.310676972 | 5.58052E-06 | 4.96373E-05 | 4.96373E-05 | BUB1B/MAD2L1/SPDL1/C | 7 |
| MEgreen | BP | GO:004500 | DNA-templated DNA replication maintenance of fidelity | 7/276 | 21/7644 | 0.333333333 | 9.231884058 | 7.310676972 | 5.58052E-06 | 4.96373E-05 | 4.96373E-05 | RAD51/MMS22L/CENPS/F | 7 |
| MEgreen | BP | GO:014052 | reciprocal homologous recombination | 7/276 | 21/7644 | 0.333333333 | 9.231884058 | 7.310676972 | 5.58052E-06 | 4.96373E-05 | 4.96373E-05 | RAD51/ANKLE1/CENPS/M | 7 |
| MEgreen | MF | GO:014064 | catalytic activity, acting on a nucleic acid | 30/251 | 382/7669 | 0.078534031 | 2.3995119 | 5.161410377 | 6.44358E-06 | 0.000108783 | 0.000108783 | MCM4/RAD51/RFC3/ANK | 30 |
| MEgreen | MF | GO:000369 | single-stranded DNA binding | 9/251 | 43/7669 | 0.209302326 | 6.394978227 | 6.525443323 | 7.94539E-06 | 0.000120107 | 0.000120107 | MCM4/RAD51/MCM6/MCI | 9 |
| MEgreen | MF | GO:000368 | DNA replication origin binding | 5/251 | 10/7669 | 0.5 | 15.27689243 | 8.309638885 | 7.95136E-06 | 0.000120107 | 0.000120107 | MCM10/CDC45/CDC6/MC | 5 |
| MEgreen | BP | GO:001097 | negative regulation of G2/M transition of mitotic cell cycle | 6/276 | 15/7644 | 0.4 | 11.07826087 | 7.56151956 | 7.98051E-06 | 7.02451E-05 | 7.02451E-05 | CLSPN/TICRR/CDK1/CDC | 6 |
| MEgreen | CC | GO:009951 | polymeric cytoskeletal fiber | 17/276 | 135/7449 | 0.125925926 | 3.39863124 | 5.51666804 | 8.95104E-06 | 7.89798E-05 | 7.89798E-05 | SKA1/TPX2/TUBA1B/TUB | 17 |
| MEgreen | CC | GO:003299 | protein-DNA complex | 11/276 | 61/7449 | 0.180327869 | 4.866892373 | 5.948226785 | 1.18878E-05 | 9.50848E-05 | 9.50848E-05 | H2BC11/PRIM2/CENPA/G | 11 |
| MEgreen | CC | GO:007268 | mitotic spindle | 8/276 | 31/7449 | 0.258064516 | 6.964936886 | 6.527806086 | 1.20441E-05 | 9.50848E-05 | 9.50848E-05 | SKA1/KNSTRN/RACGAP1 | 8 |
| MEgreen | BP | GO:200125 | negative regulation of chromosome organization | 6/276 | 16/7644 | 0.375 | 10.38586957 | 7.273456167 | 1.23839E-05 | 0.00010788 | 0.00010788 | BUB1B/MAD2L1/SPDL1/B | 6 |
| MEgreen | BP | GO:000627 | DNA strand elongation involved in DNA replication | 5/276 | 10/7644 | 0.5 | 13.84782609 | 7.868015986 | 1.28537E-05 | 0.000107927 | 0.000107927 | MCM4/LIG1/POLE/MCM3/ | 5 |
| MEgreen | BP | GO:002261 | DNA strand elongation | 5/276 | 10/7644 | 0.5 | 13.84782609 | 7.868015986 | 1.28537E-05 | 0.000107927 | 0.000107927 | MCM4/LIG1/POLE/MCM3/ | 5 |
| MEgreen | BP | GO:005144 | regulation of meiotic cell cycle | 5/276 | 10/7644 | 0.5 | 13.84782609 | 7.868015986 | 1.28537E-05 | 0.000107927 | 0.000107927 | CDC25A/TRIP13/CDC25C | 5 |
| MEgreen | BP | GO:003304 | regulation of chromosome organization | 8/276 | 32/7644 | 0.25 | 6.923913043 | 6.498991015 | 1.29002E-05 | 0.000107927 | 0.000107927 | BUB1B/MAD2L1/SPDL1/C | 8 |
| MEgreen | BP | GO:003157 | DNA integrity checkpoint signaling | 8/276 | 33/7644 | 0.242424242 | 6.714097497 | 6.366421925 | 1.6501E-05 | 0.0001367 | 0.0001367 | CLSPN/TICRR/CHEK1/CD | 8 |
| MEgreen | CC | GO:190255 | serine/threonine protein kinase complex | 10/276 | 52/7449 | 0.192307692 | 5.190217391 | 5.947498992 | 1.65103E-05 | 0.000123827 | 0.000123827 | CCNB1/CCNF/CCNB3/CK | 10 |
| MEgreen | CC | GO:000092 | spindle pole | 9/276 | 42/7449 | 0.214285714 | 5.783385093 | 6.09765401 | 1.75656E-05 | 0.000125468 | 0.000125468 | SPDL1/AURKB/CENPF/PL | 9 |
| MEgreen | BP | GO:000633 | chromatin remodeling | 15/276 | 118/7644 | 0.127118644 | 3.520633751 | 5.34048016 | 2.01269E-05 | 0.000165119 | 0.000165119 | SMARCB1/EZH2/CENPN/ | 15 |
| MEgreen | BP | GO:000007 | regulation of cyclin-dependent protein serine/threonine kinase activity | 7/276 | 25/7644 | 0.28 | 7.754782609 | 6.54702586 | 2.03711E-05 | 0.000165515 | 0.000165515 | CCNB1/CCNF/CCNB3/CC | 7 |
| MEgreen | CC | GO:004359 | nuclear replication fork | 6/276 | 17/7449 | 0.352941176 | 9.525575448 | 6.90270434 | 2.1483E-05 | 0.000146475 | 0.000146475 | MMS22L/WDHD1/MCM10 | 6 |
| MEgreen | BP | GO:000860 | attachment of spindle microtubules to kinetochore | 5/276 | 11/7644 | 0.454545455 | 12.58893281 | 7.44395636 | 2.28711E-05 | 0.000184058 | 0.000184058 | NDC80/KNSTRN/NUF2/KI | 5 |
| MEgreen | BP | GO:190275 | negative regulation of cell cycle G2/M phase transition | 6/276 | 18/7644 | 0.333333333 | 9.231884058 | 6.767040244 | 2.70046E-05 | 0.000215272 | 0.000215272 | CLSPN/TICRR/CDK1/CDC | 6 |
| MEgreen | MF | GO:004239 | histone binding | 10/251 | 62/7669 | 0.161290323 | 4.928029816 | 5.712134312 | 2.8286E-05 | 0.000405904 | 0.000405904 | COPRS/HJURP/HPF1/CK | 10 |
| MEgreen | MF | GO:000452 | DNA exonuclease activity | 5/251 | 13/7669 | 0.384615385 | 11.75145572 | 7.136289872 | 3.74492E-05 | 0.000488542 | 0.000488542 | MGME1/POLE/POLD1/EX | 5 |
| MEgreen | MF | GO:001689 | DNA exonuclease activity, producing 5'-phosphomonoesters | 5/251 | 13/7669 | 0.384615385 | 11.75145572 | 7.136289872 | 3.74492E-05 | 0.000488542 | 0.000488542 | MGME1/POLE/POLD1/EX | 5 |
| MEgreen | BP | GO:190198 | positive regulation of cell cycle phase transition | 5/276 | 12/7644 | 0.416666667 | 11.53985507 | 7.071600919 | 3.80538E-05 | 0.000299509 | 0.000299509 | CDC25A/CDCA5/CDC16/F | 5 |
| MEgreen | CC | GO:190291 | protein kinase complex | 10/276 | 57/7449 | 0.175438596 | 4.734935164 | 5.552179209 | 3.82573E-05 | 0.000249504 | 0.000249504 | CCNB1/CCNF/CCNB3/CK | 10 |
| MEgreen | BP | GO:004477 | mitotic DNA integrity checkpoint signaling | 6/276 | 19/7644 | 0.315789474 | 8.745995423 | 6.542530784 | 3.82804E-05 | 0.000299509 | 0.000299509 | CLSPN/TICRR/CDK1/CDC | 6 |
| MEgreen | BP | GO:000926 | deoxyribonucleotide metabolic process | 5/276 | 13/7644 | 0.384615385 | 10.65217391 | 6.740897913 | 6.00193E-05 | 0.000456903 | 0.000456903 | DUT/RRM1/DTYMK/RRM2 | 5 |
| MEgreen | BP | GO:005123 | spindle elongation | 5/276 | 13/7644 | 0.384615385 | 10.65217391 | 6.740897913 | 6.00193E-05 | 0.000456903 | 0.000456903 | KIF4A/RACGAP1/PRC1/K | 5 |
| MEgreen | BP | GO:200024 | regulation of reproductive process | 5/276 | 13/7644 | 0.384615385 | 10.65217391 | 6.740897913 | 6.00193E-05 | 0.000456903 | 0.000456903 | CDC25A/TRIP13/CDC25C | 5 |
| MEgreen | CC | GO:000587 | microtubule associated complex | 9/276 | 50/7449 | 0.18 | 4.858043478 | 5.36895255 | 7.60869E-05 | 0.000475543 | 0.000475543 | INCENP/HAUS4/KIF4A/KI | 9 |
| MEgreen | CC | GO:009908 | supramolecular polymer | 17/276 | 163/7449 | 0.104294479 | 2.814817285 | 4.595217639 | 0.000104287 | 0.000601654 | 0.000601654 | SKA1/TPX2/TUBA1B/TUB | 17 |
| MEgreen | CC | GO:009951 | supramolecular fiber | 17/276 | 163/7449 | 0.104294479 | 2.814817285 | 4.595217639 | 0.000104287 | 0.000601654 | 0.000601654 | SKA1/TPX2/TUBA1B/TUB | 17 |
| MEgreen | CC | GO:000587 | spindle microtubule | 6/276 | 22/7449 | 0.272727273 | 7.360671937 | 5.860458362 | 0.000110729 | 0.000615162 | 0.000615162 | SKA1/AURKB/KIF11/AURI | 6 |
| MEgreen | BP | GO:014069 | membraneless organelle assembly | 13/276 | 108/7644 | 0.12037037 | 3.33373591 | 4.727209001 | 0.000127326 | 0.000960627 | 0.000960627 | INCENP/HAUS4/CENPT/F | 13 |
| MEgreen | CC | GO:004257 | DNA polymerase complex | 5/276 | 15/7449 | 0.333333333 | 8.996376812 | 6.08067178 | 0.000148958 | 0.00079799 | 0.00079799 | POLE2/PRIM2/POLE/POL | 5 |

| | | | | | | | | | | | | | |
|---|---|---|---|---|---|---|---|---|---|---|---|---|---|
| MEgreen | MF | GO:001646 | pyrophosphatase activity | 25/251 | 351/7669 | 0.071225071 | 2.176195503 | 4.149266872 | 0.000193184 | 0.002410602 | 0.002410602 | GPN2/MCM4/RAD51/RFC | 25 |
| MEgreen | MF | GO:001988 | protein kinase regulator activity | 9/251 | 64/7669 | 0.140625 | 4.296625996 | 4.871296395 | 0.000212951 | 0.002546534 | 0.002546534 | CCNB1/CCNF/CCNB3/CK | 9 |
| MEgreen | MF | GO:000452 | DNA endonuclease activity | 6/251 | 28/7669 | 0.214285714 | 6.547239613 | 5.408979554 | 0.000237832 | 0.002730307 | 0.002730307 | ANKLE1/FEN1/NEIL3/DN | 6 |
| MEgreen | BP | GO:004581 | negative regulation of gene expression, epigenetic | 7/276 | 36/7644 | 0.194444444 | 5.3852657 | 5.104151691 | 0.000251556 | 0.001881108 | 0.001881108 | EZH2/LMNB2/HELLS/UHF | 7 |
| MEgreen | MF | GO:001711 | ribonucleoside triphosphate phosphatase activity | 23/251 | 316/7669 | 0.07278481 | 2.22385143 | 4.086715982 | 0.000256577 | 0.002786848 | 0.002786848 | GPN2/MCM4/RAD51/RFC | 23 |
| MEgreen | MF | GO:001681 | hydrolase activity, acting on acid anhydrides, in phosphorus-containing | 25/251 | 358/7669 | 0.069832402 | 2.133644194 | 4.040772389 | 0.000262177 | 0.002786848 | 0.002786848 | GPN2/MCM4/RAD51/RFC | 25 |
| MEgreen | MF | GO:001681 | hydrolase activity, acting on acid anhydrides | 25/251 | 359/7669 | 0.069637883 | 2.127700896 | 4.025473384 | 0.000273636 | 0.002804767 | 0.002804767 | GPN2/MCM4/RAD51/RFC | 25 |
| MEgreen | BP | GO:190199 | positive regulation of mitotic cell cycle phase transition | 4/276 | 10/7644 | 0.4 | 11.07826087 | 6.171932675 | 0.000294004 | 0.00217924 | 0.00217924 | CDC25A/CDCA5/CDC16/ | 4 |
| MEgreen | BP | GO:004002 | epigenetic regulation of gene expression | 7/276 | 37/7644 | 0.189189189 | 5.239717979 | 5.003141489 | 0.000300811 | 0.002210308 | 0.002210308 | EZH2/LMNB2/HELLS/UHF | 7 |
| MEgreen | BP | GO:003129 | replication fork processing | 5/276 | 18/7644 | 0.277777778 | 7.693236715 | 5.502191466 | 0.000344308 | 0.002508109 | 0.002508109 | RAD51/MMS22L/CENPS/ | 5 |
| MEgreen | BP | GO:000007 | DNA damage checkpoint signaling | 6/276 | 28/7644 | 0.214285714 | 5.934782609 | 5.062852043 | 0.000404074 | 0.002918315 | 0.002918315 | CLSPN/TICRR/CHEK1/CD | 6 |
| MEgreen | CC | GO:006169 | transferase complex, transferring phosphorus-containing groups | 15/276 | 150/7449 | 0.1 | 2.698913043 | 4.122959398 | 0.000419951 | 0.002172158 | 0.002172158 | CCNB1/CCNF/CCNB3/PO | 15 |
| MEgreen | MF | GO:004313 | 3'-5' DNA helicase activity | 4/251 | 12/7669 | 0.333333333 | 10.18459495 | 5.856738133 | 0.000450648 | 0.00445986 | 0.00445986 | MCM6/MCM2/RECQL4/M | 4 |
| MEgreen | MF | GO:000809 | cytoskeletal protein binding | 23/251 | 330/7669 | 0.06969697 | 2.129506218 | 3.85799491 | 0.000477474 | 0.004567836 | 0.004567836 | KIFC1/SKA1/HAUS4/CEN | 23 |
| MEgreen | BP | GO:000701 | microtubule-based movement | 12/276 | 108/7644 | 0.111111111 | 3.077294686 | 4.207762368 | 0.000488553 | 0.003480312 | 0.003480312 | KIFC1/CENPE/KIF4A/KIF | 12 |
| MEgreen | BP | GO:001063 | negative regulation of organelle organization | 8/276 | 52/7644 | 0.153846154 | 4.260869565 | 4.566344381 | 0.000493109 | 0.003480312 | 0.003480312 | BUB1B/MAD2L1/SPDL1/T | 8 |
| MEgreen | BP | GO:003150 | heterochromatin formation | 6/276 | 29/7644 | 0.206896552 | 5.730134933 | 4.939116124 | 0.000494246 | 0.003480312 | 0.003480312 | EZH2/LMNB2/HELLS/H2A | 6 |
| MEgreen | BP | GO:000007 | DNA replication checkpoint signaling | 4/276 | 12/7644 | 0.333333333 | 9.231884058 | 5.523092914 | 0.000654379 | 0.004495528 | 0.004495528 | CLSPN/TICRR/CDC6/OR | 4 |
| MEgreen | BP | GO:000622 | pyrimidine nucleotide biosynthetic process | 4/276 | 12/7644 | 0.333333333 | 9.231884058 | 5.523092914 | 0.000654379 | 0.004495528 | 0.004495528 | DHODH/DUT/DTYMK/TY | 4 |
| MEgreen | BP | GO:004593 | positive regulation of mitotic cell cycle | 4/276 | 12/7644 | 0.333333333 | 9.231884058 | 5.523092914 | 0.000654379 | 0.004495528 | 0.004495528 | CDC25A/CDCA5/CDC16/ | 4 |
| MEgreen | BP | GO:004277 | signal transduction in response to DNA damage | 6/276 | 31/7644 | 0.193548387 | 5.360448808 | 4.708101168 | 0.000720979 | 0.004913126 | 0.004913126 | CLSPN/TICRR/CHEK1/CD | 6 |
| MEgreen | CC | GO:000581 | centrosome | 13/276 | 126/7449 | 0.103174603 | 2.784592823 | 3.962815537 | 0.000751117 | 0.003755587 | 0.003755587 | LRRC45/CCNF/XRCC2/K | 13 |
| MEgreen | BP | GO:003304 | regulation of organelle organization | 16/276 | 182/7644 | 0.087912088 | 2.434782609 | 3.791454406 | 0.000860785 | 0.005818908 | 0.005818908 | BUB1B/SKA1/MAD2L1/M | 16 |
| MEgreen | BP | GO:007190 | regulation of protein serine/threonine kinase activity | 7/276 | 44/7644 | 0.159090909 | 4.406126482 | 4.385225948 | 0.000902227 | 0.006050647 | 0.006050647 | CCNB1/CCNF/CCNB3/CC | 7 |
| MEgreen | BP | GO:003450 | protein localization to chromosome | 4/276 | 13/7644 | 0.307692308 | 8.52173913 | 5.253042067 | 0.000918538 | 0.006063789 | 0.006063789 | SPDL1/KNL1/KNTC1/TTK | 4 |
| MEgreen | BP | GO:007252 | pyrimidine-containing compound biosynthetic process | 4/276 | 13/7644 | 0.307692308 | 8.52173913 | 5.253042067 | 0.000918538 | 0.006063789 | 0.006063789 | DHODH/DUT/DTYMK/TY | 4 |
| MEgreen | MF | GO:000368 | chromatin binding | 13/251 | 146/7669 | 0.089041096 | 2.720542488 | 3.860831047 | 0.000954557 | 0.008837349 | 0.008837349 | TOX2/EZH2/HMGN2/WD | 13 |
| MEgreen | CC | GO:000578 | endoplasmic reticulum lumen | 4/276 | 13/7449 | 0.307692308 | 8.304347826 | 5.170200566 | 0.001011634 | 0.004895002 | 0.004895002 | HSPA5/PDIA6/TOR2A/PD | 4 |
| MEgreen | MF | GO:000520 | structural constituent of cytoskeleton | 5/251 | 25/7669 | 0.2 | 6.110756972 | 4.707923456 | 0.001120135 | 0.010000497 | 0.010000497 | TUBA1B/TUBB4A/TUBB3/ | 5 |
| MEgreen | MF | GO:001664 | oxidoreductase activity, acting on the CH-NH group of donors | 4/251 | 15/7669 | 0.266666667 | 8.147675963 | 5.096837115 | 0.001149883 | 0.010000497 | 0.010000497 | MTHFD1L/ETFDH/PYCR1 | 4 |
| MEgreen | BP | GO:005133 | regulation of transferase activity | 11/276 | 103/7644 | 0.106796117 | 2.957788096 | 3.871519827 | 0.001162385 | 0.007614073 | 0.007614073 | CCNB1/CLSPN/TPX2/CC | 11 |
| MEgreen | BP | GO:000701 | cytoskeleton organization | 29/276 | 441/7644 | 0.065759637 | 1.821256039 | 3.438371715 | 0.001186226 | 0.007710469 | 0.007710469 | INCENP/SKA1/HAUS4/TP | 29 |
| MEgreen | MF | GO:001920 | kinase regulator activity | 9/251 | 81/7669 | 0.111111111 | 3.394864985 | 3.985606155 | 0.001244215 | 0.010202565 | 0.010202565 | CCNB1/CCNF/CCNB3/CK | 9 |
| MEgreen | MF | GO:012054 | nucleic acid conformation isomerase activity | 9/251 | 81/7669 | 0.111111111 | 3.394864985 | 3.985606155 | 0.001244215 | 0.010202565 | 0.010202565 | MCM4/MCM6/TOP2A/MC | 9 |
| MEgreen | BP | GO:190129 | nucleoside phosphate biosynthetic process | 10/276 | 89/7644 | 0.112359551 | 3.111871031 | 3.878437739 | 0.001309661 | 0.00844781 | 0.00844781 | DHODH/DUT/IMPDH1/IM | 10 |
| MEgreen | BP | GO:003336 | protein localization to organelle | 15/276 | 174/7644 | 0.086206897 | 2.387556222 | 3.583245587 | 0.001512959 | 0.009685228 | 0.009685228 | TIMM44/NUP85/SPDL1/H | 15 |
| MEgreen | BP | GO:005122 | spindle assembly | 6/276 | 36/7644 | 0.166666667 | 4.615942029 | 4.208711144 | 0.001639444 | 0.010416016 | 0.010416016 | INCENP/HAUS4/RACGAP | 6 |
| MEgreen | BP | GO:004484 | cell cycle G1/S phase transition | 5/276 | 25/7644 | 0.2 | 5.539130435 | 4.399520467 | 0.001735281 | 0.010781707 | 0.010781707 | E2F1/CCNE2/FAM83D/C | 5 |
| MEgreen | BP | GO:007252 | pyrimidine-containing compound metabolic process | 5/276 | 25/7644 | 0.2 | 5.539130435 | 4.399520467 | 0.001735281 | 0.010781707 | 0.010781707 | DHODH/DUT/TK1/DTYMK | 5 |
| MEgreen | BP | GO:009006 | positive regulation of cell cycle process | 5/276 | 25/7644 | 0.2 | 5.539130435 | 4.399520467 | 0.001735281 | 0.010781707 | 0.010781707 | CDC25A/CDCA5/CDC16/ | 5 |
| MEgreen | BP | GO:000916 | nucleotide biosynthetic process | 8/276 | 64/7644 | 0.125 | 3.461956522 | 3.827784023 | 0.001995934 | 0.012310687 | 0.012310687 | DHODH/DUT/IMPDH1/IM | 8 |
| MEgreen | BP | GO:005164 | organelle localization | 13/276 | 144/7644 | 0.090277778 | 2.500301932 | 3.517555393 | 0.002044006 | 0.012515833 | 0.012515833 | INCENP/CCNB1/NDC80/ | 13 |
| MEgreen | BP | GO:004477 | mitotic DNA damage checkpoint | 4/276 | 16/7644 | 0.25 | 6.923913043 | 4.590658496 | 0.002146126 | 0.013046595 | 0.013046595 | CLSPN/TICRR/CDK1/DTL | 4 |
| MEgreen | CC | GO:003204 | small-subunit processome | 5/276 | 26/7449 | 0.192307692 | 5.190217391 | 4.198145229 | 0.002334955 | 0.010945103 | 0.010945103 | FBL/NOP56/WDR46/UTP | 5 |
| MEgreen | BP | GO:004585 | regulation of protein kinase activity | 9/276 | 84/7644 | 0.107142857 | 2.967391304 | 3.508982118 | 0.003144595 | 0.018979874 | 0.018979874 | CCNB1/CLSPN/TPX2/CC | 9 |
| MEgreen | MF | GO:000438 | helicase activity | 8/251 | 76/7669 | 0.105263158 | 3.21618788 | 3.571422004 | 0.003245724 | 0.025875634 | 0.025875634 | MCM4/MCM6/MCM2/CHT | 8 |
| MEgreen | BP | GO:000622 | pyrimidine nucleotide metabolic process | 4/276 | 18/7644 | 0.222222222 | 6.154589372 | 4.237342688 | 0.003408607 | 0.020141768 | 0.020141768 | DHODH/DUT/DTYMK/TY | 4 |
| MEgreen | BP | GO:000628 | base-excision repair | 4/276 | 18/7644 | 0.222222222 | 6.154589372 | 4.237342688 | 0.003408607 | 0.020141768 | 0.020141768 | POLE/NEIL3/POLD1/APE | 4 |
| MEgreen | BP | GO:009030 | mitotic spindle assembly | 4/276 | 18/7644 | 0.222222222 | 6.154589372 | 4.237342688 | 0.003408607 | 0.020141768 | 0.020141768 | RACGAP1/PRC1/KIF11/K | 4 |
| MEgreen | BP | GO:004578 | positive regulation of cell cycle | 5/276 | 29/7644 | 0.172413793 | 4.775112444 | 3.941899968 | 0.003447337 | 0.020229165 | 0.020229165 | CDC25A/CDCA5/CDC16/ | 5 |
| MEgreen | MF | GO:001664 | oxidoreductase activity, acting on the CH-NH group of donors, NAD or NADP as | 3/251 | 10/7669 | 0.3 | 9.166135458 | 4.752969673 | 0.003504562 | 0.026405378 | 0.026405378 | MTHFD1L/PYCR1/MTHFD | 3 |

| | | | | | | | | | | | | | |
|---|---|---|---|---|---|---|---|---|---|---|---|---|---|
| MEgreen | MF | GO:001688 | DNA endonuclease activity, producing 5'-phosphomonoesters | 3/251 | 10/7669 | 0.3 | 9.166135458 | 4.752969673 | 0.003504562 | 0.026405378 | 0.026405378 | FEN1/DNA2/EXO1 | 3 |
| MEgreen | MF | GO:001686 | intramolecular oxidoreductase activity | 4/251 | 20/7669 | 0.2 | 6.110756972 | 4.209518238 | 0.003588187 | 0.026405378 | 0.026405378 | P4HB/PDIA6/TXNDC5/PD | 4 |
| MEgreen | BP | GO:190113 | carbohydrate derivative biosynthetic process | 14/276 | 174/7644 | 0.08045977 | 2.228385807 | 3.172201567 | 0.00405838 | 0.023650561 | 0.023650561 | DHODH/ADK/DUT/IMPDH | 14 |
| MEgreen | BP | GO:005112 | negative regulation of cellular component organization | 8/276 | 73/7644 | 0.109589041 | 3.035139964 | 3.381354869 | 0.004585996 | 0.026414559 | 0.026414559 | BUB1B/MAD2L1/SPDL1/T | 8 |
| MEgreen | BP | GO:000673 | one-carbon metabolic process | 3/276 | 10/7644 | 0.3 | 8.308695652 | 4.475849365 | 0.004626455 | 0.026414559 | 0.026414559 | MTHFD1L/SHMT2/MTHFD | 3 |
| MEgreen | BP | GO:004665 | tetrahydrofolate metabolic process | 3/276 | 10/7644 | 0.3 | 8.308695652 | 4.475849365 | 0.004626455 | 0.026414559 | 0.026414559 | MTHFD1L/SHMT2/MTHFD | 3 |
| MEgreen | CC | GO:000587 | kinesin complex | 3/276 | 10/7449 | 0.3 | 8.09673913 | 4.404793179 | 0.004974532 | 0.021319422 | 0.021319422 | KIF20A/KIF22/KIF23 | 3 |
| MEgreen | CC | GO:003089 | replisome | 3/276 | 10/7449 | 0.3 | 8.09673913 | 4.404793179 | 0.004974532 | 0.021319422 | 0.021319422 | PRIM2/POLD1/POLA2 | 3 |
| MEgreen | CC | GO:004360 | nuclear replisome | 3/276 | 10/7449 | 0.3 | 8.09673913 | 4.404793179 | 0.004974532 | 0.021319422 | 0.021319422 | PRIM2/POLD1/POLA2 | 3 |
| MEgreen | MF | GO:000452 | exonuclease activity | 6/251 | 50/7669 | 0.12 | 3.666454183 | 3.479402246 | 0.005502193 | 0.039478238 | 0.039478238 | MGME1/POLE/FEN1/POL | 6 |
| MEgreen | CC | GO:014016 | nuclear ribonucleoprotein granule | 4/276 | 20/7449 | 0.2 | 5.397826087 | 3.862885446 | 0.005586545 | 0.023277271 | 0.023277271 | FBL/SRSF12/WRAP53/RN | 4 |
| MEgreen | BP | GO:000706 | sister chromatid cohesion | 4/276 | 21/7644 | 0.19047619 | 5.275362319 | 3.796918498 | 0.006122716 | 0.034428327 | 0.034428327 | BUB1B/CDCA5/ESCO2/BU | 4 |
| MEgreen | BP | GO:000071 | resolution of meiotic recombination intermediates | 3/276 | 11/7644 | 0.272727273 | 7.553359684 | 4.209440534 | 0.006193025 | 0.034428327 | 0.034428327 | ANKLE1/CENPS/TOP2A | 3 |
| MEgreen | BP | GO:000630 | DNA damage tolerance | 3/276 | 11/7644 | 0.272727273 | 7.553359684 | 4.209440534 | 0.006193025 | 0.034428327 | 0.034428327 | POLE2/PCLAF/RAD18 | 3 |
| MEgreen | BP | GO:000911 | nucleoside metabolic process | 3/276 | 11/7644 | 0.272727273 | 7.553359684 | 4.209440534 | 0.006193025 | 0.034428327 | 0.034428327 | ADK/TK1/ADA | 3 |
| MEgreen | CC | GO:000581 | microtubule organizing center | 14/276 | 180/7449 | 0.077777778 | 2.099154589 | 2.928071351 | 0.006831857 | 0.027416177 | 0.027416177 | LRRC45/CCNF/XRCC2/KN | 14 |
| MEgreen | CC | GO:000565 | nucleoplasm | 19/276 | 277/7449 | 0.068592058 | 1.851239994 | 2.832016136 | 0.006945432 | 0.027416177 | 0.027416177 | MAD2L1/EZH2/ANKLE1/F | 19 |
| MEgreen | BP | GO:000609 | tricarboxylic acid cycle | 3/276 | 12/7644 | 0.25 | 6.923913043 | 3.97458491 | 0.00803924 | 0.043545882 | 0.043545882 | MDH1/IDH3A/ACO2 | 3 |
| MEgreen | BP | GO:000676 | folic acid-containing compound metabolic process | 3/276 | 12/7644 | 0.25 | 6.923913043 | 3.97458491 | 0.00803924 | 0.043545882 | 0.043545882 | MTHFD1L/SHMT2/MTHFD | 3 |
| MEgreen | BP | GO:004438 | protein localization to cytoskeleton | 3/276 | 12/7644 | 0.25 | 6.923913043 | 3.97458491 | 0.00803924 | 0.043545882 | 0.043545882 | CEP72/STIL/FAM83D | 3 |
| MEgreen | BP | GO:007269 | protein localization to microtubule cytoskeleton | 3/276 | 12/7644 | 0.25 | 6.923913043 | 3.97458491 | 0.00803924 | 0.043545882 | 0.043545882 | CEP72/STIL/FAM83D | 3 |
| MEgreen | BP | GO:000028 | mitotic cytokinesis | 4/276 | 23/7644 | 0.173913043 | 4.816635161 | 3.547727434 | 0.008560467 | 0.046073851 | 0.046073851 | INCENP/ECT2/RACGAP1/ | 4 |
| MEgreen | BP | GO:000091 | cytokinesis | 6/276 | 50/7644 | 0.12 | 3.323478261 | 3.1900683 | 0.008780025 | 0.04695646 | 0.04695646 | INCENP/ECT2/RACGAP1/ | 6 |
| MEgreen | MF | GO:014061 | DNA damage sensor activity | 3/251 | 14/7669 | 0.214285714 | 6.547239613 | 3.821227058 | 0.009648714 | 0.065932879 | 0.065932879 | RAD51/XRCC2/XRCC3 | 3 |
| MEgreen | MF | GO:014066 | ATP-dependent DNA damage sensor activity | 3/251 | 14/7669 | 0.214285714 | 6.547239613 | 3.821227058 | 0.009648714 | 0.065932879 | 0.065932879 | RAD51/XRCC2/XRCC3 | 3 |
| MEgreen | BP | GO:004354 | regulation of kinase activity | 9/276 | 100/7644 | 0.09 | 2.492608696 | 2.907748161 | 0.009864978 | 0.052427083 | 0.052427083 | CCNB1/CLSPN/TPX2/CCN | 9 |
| MEgreen | CC | GO:014053 | endoplasmic reticulum protein-containing complex | 5/276 | 36/7449 | 0.138888889 | 3.748490338 | 3.242443555 | 0.009901356 | 0.038082139 | 0.038082139 | HSPA5/SEC61A1/SEC11 | 5 |
| MEgreen | BP | GO:000008 | G1/S transition of mitotic cell cycle | 4/276 | 24/7644 | 0.166666667 | 4.615942029 | 3.433691368 | 0.009986592 | 0.052741688 | 0.052741688 | E2F1/CCNE2/CCNE1/CDK | 4 |
| MEgreen | MF | GO:001649 | oxidoreductase activity | 19/251 | 327/7669 | 0.058103976 | 1.775296368 | 2.635535886 | 0.010674316 | 0.070431021 | 0.070431021 | DHODH/MTHFD1L/ACAD | 19 |
| MEgreen | MF | GO:000451 | nuclease activity | 9/251 | 112/7669 | 0.080357143 | 2.455214855 | 2.85361854 | 0.010930463 | 0.070431021 | 0.070431021 | ANKLE1/MGME1/POLE/FI | 9 |
| MEgreen | MF | GO:001661 | oxidoreductase activity, acting on the CH-OH group of donors, NAD or NADP as | 6/251 | 58/7669 | 0.103448276 | 3.160736365 | 3.038291336 | 0.011288596 | 0.070431021 | 0.070431021 | MDH1/IDH2/IDH3A/IMPDH | 6 |
| MEgreen | MF | GO:004698 | protein heterodimerization activity | 6/251 | 58/7669 | 0.103448276 | 3.160736365 | 3.038291336 | 0.011288596 | 0.070431021 | 0.070431021 | CENPT/H2BC11/CENPS/ | 6 |
| MEgreen | BP | GO:005112 | regulation of cellular component organization | 18/276 | 278/7644 | 0.064748201 | 1.793243666 | 2.607502995 | 0.011635284 | 0.061067174 | 0.061067174 | BUB1B/SKA1/MAD2L1/SM | 18 |
| MEgreen | MF | GO:000028 | magnesium ion binding | 6/251 | 59/7669 | 0.101694915 | 3.107164562 | 2.988592 | 0.012232337 | 0.074695334 | 0.074695334 | IDH2/DUT/IDH3A/PSPH/F | 6 |
| MEgreen | BP | GO:003026 | chromosome condensation | 3/276 | 14/7644 | 0.214285714 | 5.934782609 | 3.576691121 | 0.012609744 | 0.065773049 | 0.065773049 | NCAPG/TOP2A/NCAPH | 3 |
| MEgreen | CC | GO:199023 | transferase complex | 24/276 | 400/7449 | 0.06 | 1.619347826 | 2.49761111 | 0.012988155 | 0.048705581 | 0.048705581 | CCNB1/EZH2/CCNF/CCN | 24 |
| MEgreen | MF | GO:001661 | oxidoreductase activity, acting on CH-OH group of donors | 6/251 | 60/7669 | 0.1 | 3.055378486 | 2.939937771 | 0.013230265 | 0.079105959 | 0.079105959 | MDH1/IDH2/IDH3A/IMPDH | 6 |
| MEgreen | BP | GO:005508 | nucleobase-containing small molecule metabolic process | 13/276 | 181/7644 | 0.071823204 | 1.989190488 | 2.606598616 | 0.013779168 | 0.071431884 | 0.071431884 | DHODH/IDH2/ADK/DUT/IM | 13 |
| MEgreen | MF | GO:014029 | molecular sensor activity | 3/251 | 16/7669 | 0.1875 | 5.728834661 | 3.482833897 | 0.014145573 | 0.08198936 | 0.08198936 | RAD51/XRCC2/XRCC3 | 3 |
| MEgreen | MF | GO:000451 | endonuclease activity | 6/251 | 61/7669 | 0.098360656 | 3.005290314 | 2.892287229 | 0.014283861 | 0.08198936 | 0.08198936 | ANKLE1/FEN1/NEIL3/DN | 6 |
| MEgreen | BP | GO:000912 | nucleoside monophosphate biosynthetic process | 3/276 | 15/7644 | 0.2 | 5.539130435 | 3.405619681 | 0.015348139 | 0.078601075 | 0.078601075 | DUT/ADA/TYMS | 3 |
| MEgreen | BP | GO:004255 | pteridine-containing compound metabolic process | 3/276 | 15/7644 | 0.2 | 5.539130435 | 3.405619681 | 0.015348139 | 0.078601075 | 0.078601075 | MTHFD1L/SHMT2/MTHFD | 3 |
| MEgreen | MF | GO:014099 | histone modifying activity | 7/251 | 80/7669 | 0.0875 | 2.673456175 | 2.767588765 | 0.015428845 | 0.086825069 | 0.086825069 | EZH2/HDAC8/NSD2/FBL/ | 7 |
| MEgreen | BP | GO:000912 | nucleoside monophosphate metabolic process | 3/276 | 16/7644 | 0.1875 | 5.192934783 | 3.24925966 | 0.018394757 | 0.093635963 | 0.093635963 | DUT/ADA/TYMS | 3 |
| MEgreen | BP | GO:017003 | proteinogenic amino acid biosynthetic process | 4/276 | 29/7644 | 0.137931034 | 3.820089955 | 2.944683813 | 0.019394031 | 0.098131473 | 0.098131473 | SHMT2/PSPH/PYCR1/PS | 4 |
| MEgreen | MF | GO:004205 | histone methyltransferase activity | 4/251 | 32/7669 | 0.125 | 3.819223108 | 2.939527575 | 0.0196089 | 0.106381328 | 0.106381328 | EZH2/NSD2/FBL/SUV39H | 4 |
| MEgreen | MF | GO:001920 | nucleobase-containing compound kinase activity | 3/251 | 18/7669 | 0.166666667 | 5.092297477 | 3.197266563 | 0.019645332 | 0.106381328 | 0.106381328 | ADK/TK1/DTYMK | 3 |
| MEgreen | CC | GO:003049 | midbody | 3/276 | 16/7449 | 0.1875 | 5.060461957 | 3.189167505 | 0.019696348 | 0.07205981 | 0.07205981 | INCENP/RACGAP1/CEP5 | 3 |
| MEgreen | BP | GO:000911 | nucleobase metabolic process | 3/276 | 17/7644 | 0.176470588 | 4.887468031 | 3.105461005 | 0.021751994 | 0.109407351 | 0.109407351 | DHODH/ADK/ADA | 3 |
| MEgreen | BP | GO:000646 | protein phosphorylation | 25/276 | 457/7644 | 0.054704595 | 1.515079441 | 2.197708841 | 0.024124283 | 0.120621414 | 0.120621414 | CCNB1/BUB1B/CLSPN/TI | 25 |
| MEgreen | BP | GO:000645 | protein folding | 6/276 | 63/7644 | 0.095238095 | 2.637681159 | 2.526085473 | 0.025498828 | 0.126744175 | 0.126744175 | P4HB/HSPD1/DNAJB11/T | 6 |
| MEgreen | MF | GO:014066 | ATP-dependent protein folding | 3/251 | 20/7669 | 0.15 | 4.583067729 | 2.951224063 | 0.026162779 | 0.139050325 | 0.139050325 | HSPA5/HSPD1/HSP90B1 | 3 |

| | | | | | | | | | | | | | |
|---|---|---|---|---|---|---|---|---|---|---|---|---|---|
| MEgreen | BP | GO:003139 | regulation of protein modification process | 10/276 | 137/7644 | 0.072992701 | 2.021580451 | 2.335131334 | 0.026188928 | 0.129413123 | 0.129413123 | CCNB1/CLSPN/TPX2/CCI | 10 |
| MEgreen | BP | GO:000911 | nucleotide metabolic process | 9/276 | 119/7644 | 0.075630252 | 2.094629156 | 2.329157744 | 0.027860213 | 0.136871395 | 0.136871395 | DHODH/IDH2/DUT/IMPDH | 9 |
| MEgreen | MF | GO:004698 | protein dimerization activity | 10/251 | 153/7669 | 0.065359477 | 1.996979403 | 2.291252828 | 0.028320288 | 0.147780415 | 0.147780415 | CENPT/E2F1/H2BC11/CE | 10 |
| MEgreen | BP | GO:005105 | negative regulation of DNA metabolic process | 3/276 | 19/7644 | 0.157894737 | 4.372997712 | 2.848948412 | 0.029400184 | 0.142776753 | 0.142776753 | PARPBP/GMNN/RMI2 | 3 |
| MEgreen | BP | GO:007058 | protein localization to mitochondrion | 3/276 | 19/7644 | 0.157894737 | 4.372997712 | 2.848948412 | 0.029400184 | 0.142776753 | 0.142776753 | TIMM44/HSPD1/BCS1L | 3 |
| MEgreen | BP | GO:000193 | regulation of protein phosphorylation | 9/276 | 121/7644 | 0.074380165 | 2.060007186 | 2.274665947 | 0.030595952 | 0.147734739 | 0.147734739 | CCNB1/CLSPN/TPX2/CCI | 9 |
| MEgreen | CC | GO:001660 | nuclear body | 4/276 | 33/7449 | 0.121212121 | 3.27140975 | 2.565024718 | 0.032430034 | 0.115821552 | 0.115821552 | FBL/SRSF12/WRAP53/RI | 4 |
| MEgreen | BP | GO:000914 | nucleoside triphosphate biosynthetic process | 3/276 | 20/7644 | 0.15 | 4.154347826 | 2.733661882 | 0.033688631 | 0.160829907 | 0.160829907 | IMPDH1/IMPDH2/DTYMK | 3 |
| MEgreen | BP | GO:190265 | secondary alcohol metabolic process | 3/276 | 20/7644 | 0.15 | 4.154347826 | 2.733661882 | 0.033688631 | 0.160829907 | 0.160829907 | IDH2/IDH3A/DHCR7 | 3 |
| MEgreen | BP | GO:006164 | cytoskeleton-dependent cytokinesis | 4/276 | 35/7644 | 0.114285714 | 3.165217391 | 2.484754004 | 0.036163269 | 0.170714874 | 0.170714874 | INCENP/ECT2/RACGAP1 | 4 |
| MEgreen | BP | GO:190160 | alpha-amino acid biosynthetic process | 4/276 | 35/7644 | 0.114285714 | 3.165217391 | 2.484754004 | 0.036163269 | 0.170714874 | 0.170714874 | SHMT2/PSPH/PYCR1/PS | 4 |
| MEgreen | BP | GO:000675 | nucleoside phosphate metabolic process | 11/276 | 167/7644 | 0.065868263 | 1.824264514 | 2.084360387 | 0.039050736 | 0.183321509 | 0.183321509 | DHODH/IDH2/DUT/IMPDH | 11 |
| MEgreen | BP | GO:001631 | phosphorylation | 25/276 | 479/7644 | 0.052192067 | 1.445493328 | 1.949003083 | 0.039422606 | 0.184044762 | 0.184044762 | CCNB1/BUB1B/CLSPN/TI | 25 |
| MEgreen | MF | GO:004418 | protein folding chaperone | 3/251 | 24/7669 | 0.125 | 3.819223108 | 2.544373248 | 0.042231043 | 0.216434093 | 0.216434093 | HSPA5/HSPD1/HSP90B1 | 3 |
| MEgreen | CC | GO:003068 | preribosome | 5/276 | 52/7449 | 0.096153846 | 2.595108696 | 2.2640609 | 0.042427158 | 0.148001714 | 0.148001714 | FBL/NOP56/WDR46/UTP | 5 |
| MEgreen | BP | GO:004428 | small molecule biosynthetic process | 10/276 | 149/7644 | 0.067114094 | 1.858768602 | 2.048779102 | 0.042973176 | 0.199256649 | 0.199256649 | ADK/SHMT2/PDSS1/DHC | 10 |
| MEgreen | BP | GO:000865 | amino acid biosynthetic process | 4/276 | 37/7644 | 0.108108108 | 2.994124559 | 2.353195983 | 0.043152623 | 0.199256649 | 0.199256649 | SHMT2/PSPH/PYCR1/PS | 4 |
| MEgreen | MF | GO:000472 | protein tyrosine phosphatase activity | 4/251 | 41/7669 | 0.097560976 | 2.98085706 | 2.339236986 | 0.044023337 | 0.221661366 | 0.221661366 | CDC25A/CDKN3/CDC25C | 4 |
| MEgreen | BP | GO:006500 | protein-containing complex assembly | 18/276 | 324/7644 | 0.055555556 | 1.538647343 | 1.917490181 | 0.045186181 | 0.207512625 | 0.207512625 | RAD51/CENPT/CENPN/D | 18 |
| MEgreen | MF | GO:000504 | signal sequence binding | 3/251 | 25/7669 | 0.12 | 3.666454183 | 2.45628236 | 0.046865873 | 0.231905266 | 0.231905266 | SEC61A1/KDELR3/KPNA | 3 |
| MEgreen | BP | GO:003497 | response to endoplasmic reticulum stress | 4/276 | 38/7644 | 0.105263158 | 2.915331808 | 2.290705829 | 0.046912436 | 0.21400095 | 0.21400095 | P4HB/HSPA5/PDIA6/PDIA | 4 |
| MEgreen | BP | GO:000627 | regulation of DNA replication | 2/276 | 10/7644 | 0.2 | 5.539130435 | 2.779766054 | 0.048273135 | 0.21400095 | 0.21400095 | TICRR/GMNN | 2 |
| MEgreen | BP | GO:000657 | biogenic amine metabolic process | 2/276 | 10/7644 | 0.2 | 5.539130435 | 2.779766054 | 0.048273135 | 0.21400095 | 0.21400095 | SRM/ODC1 | 2 |
| MEgreen | BP | GO:000706 | mitotic sister chromatid cohesion | 2/276 | 10/7644 | 0.2 | 5.539130435 | 2.779766054 | 0.048273135 | 0.21400095 | 0.21400095 | CDCA5/ESCO2 | 2 |
| MEgreen | BP | GO:000914 | pyrimidine nucleoside triphosphate metabolic process | 2/276 | 10/7644 | 0.2 | 5.539130435 | 2.779766054 | 0.048273135 | 0.21400095 | 0.21400095 | DUT/DTYMK | 2 |
| MEgreen | BP | GO:001021 | response to ionizing radiation | 2/276 | 10/7644 | 0.2 | 5.539130435 | 2.779766054 | 0.048273135 | 0.21400095 | 0.21400095 | TICRR/RHNO1 | 2 |
| MEgreen | BP | GO:007189 | DNA biosynthetic process | 3/276 | 23/7644 | 0.130434783 | 3.612476371 | 2.428409764 | 0.048371812 | 0.21400095 | 0.21400095 | POLE2/LIG1/PCLAF | 3 |

**Supplementary Table 20. Summary of gene connectivity clusters from telomere length-ordered sliding-window analysis in rumen and whole blood, related to Figure 5.**

| Tissue | Cluster | No. of genes | Pearson r between mean Z-connectivity and mean TL residual |
|---|---|---|---|
| Rumen | C1 | 878 | -0.462031731 |
| Rumen | C2 | 730 | -0.707532815 |
| Rumen | C3 | 788 | 0.037573309 |
| Rumen | C4 | 638 | 0.322803359 |
| Rumen | C5 | 1645 | -0.155367705 |
| Rumen | C6 | 644 | 0.836462329 |
| Whole blood | C1 | 1342 | 0.350563472 |
| Whole blood | C2 | 687 | 0.085905411 |
| Whole blood | C3 | 622 | 0.697492422 |
| Whole blood | C4 | 1398 | 0.826160929 |
| Whole blood | C5 | 883 | -0.705800454 |
| Whole blood | C6 | 591 | -0.498889843 |

**Supplementary Table 21. Gene Ontology enrichment of rumen connectivity clusters C2 and C6 identified from telomere length-ordered sliding-window analysis, related to Figure 5.**

| Cluster | ONTOLOGY | ID | Description | GeneRatio | BgRatio | RichFactor | FoldEnrichment | zScore | pvalue | p.adjust | qvalue | geneID | Count |
|---|---|---|---|---|---|---|---|---|---|---|---|---|---|
| Cluster_2 | CC | GO:000573 | mitochondrion | 50/435 | 239/3473 | 0.209205021 | 1.67027365 | 4.062764894 | 0.000102301 | 0.021892508 | 0.021892508 | ADO/AK3/COX14/CRLS1/ | 50 |
| Cluster_2 | MF | GO:004830 | calcium-dependent protein binding | 7/419 | 13/3517 | 0.538461538 | 4.519735634 | 4.675094678 | 0.000292642 | 0.087500019 | 0.087500019 | S100A9/S100A4/S100A8/ | 7 |
| Cluster_2 | CC | GO:009879 | mitochondrial protein-containing complex | 11/435 | 31/3473 | 0.35483871 | 2.832999629 | 3.878630364 | 0.000847405 | 0.065489601 | 0.065489601 | MRPL16/CHCHD3/TIMM2 | 11 |
| Cluster_2 | CC | GO:003197 | membrane-enclosed lumen | 39/435 | 195/3473 | 0.2 | 1.596781609 | 3.245404726 | 0.001530131 | 0.065489601 | 0.065489601 | LSM6/RRS1/MAD2L1/ZNF | 39 |
| Cluster_2 | CC | GO:004323 | organelle lumen | 39/435 | 195/3473 | 0.2 | 1.596781609 | 3.245404726 | 0.001530131 | 0.065489601 | 0.065489601 | LSM6/RRS1/MAD2L1/ZNF | 39 |
| Cluster_2 | CC | GO:007001 | intracellular organelle lumen | 39/435 | 195/3473 | 0.2 | 1.596781609 | 3.245404726 | 0.001530131 | 0.065489601 | 0.065489601 | LSM6/RRS1/MAD2L1/ZNF | 39 |
| Cluster_2 | MF | GO:001682 | lyase activity | 12/419 | 41/3517 | 0.292682927 | 2.456720414 | 3.449999157 | 0.002153989 | 0.322021307 | 0.322021307 | LOC100847874/GADL1/P | 12 |
| Cluster_2 | BP | GO:000690 | vesicle fusion | 8/442 | 21/3586 | 0.380952381 | 3.090713208 | 3.602373915 | 0.002356676 | 0.839437462 | 0.839437462 | STXBP6/VAMP3/VAMP8/ | 8 |
| Cluster_2 | BP | GO:009017 | organelle membrane fusion | 8/442 | 22/3586 | 0.363636364 | 2.950226244 | 3.439869249 | 0.003311641 | 0.839437462 | 0.839437462 | STXBP6/VAMP3/VAMP8/ | 8 |
| Cluster_2 | CC | GO:000031 | organellar ribosome | 7/435 | 18/3473 | 0.388888889 | 3.104853129 | 3.387463513 | 0.004243839 | 0.129740232 | 0.129740232 | MRPL16/MRPL57/MRPL5 | 7 |
| Cluster_2 | CC | GO:000576 | mitochondrial ribosome | 7/435 | 18/3473 | 0.388888889 | 3.104853129 | 3.387463513 | 0.004243839 | 0.129740232 | 0.129740232 | MRPL16/MRPL57/MRPL5 | 7 |
| Cluster_2 | BP | GO:001598 | energy derivation by oxidation of organic compounds | 12/442 | 43/3586 | 0.279069767 | 2.264127118 | 3.126466974 | 0.004455466 | 0.839437462 | 0.839437462 | PPP1R3C/SDHD/PPP1R3 | 12 |
| Cluster_2 | BP | GO:000609 | generation of precursor metabolites and energy | 13/442 | 49/3586 | 0.265306122 | 2.152460984 | 3.04523763 | 0.005030246 | 0.839437462 | 0.839437462 | PPP1R3C/SDHD/PPP1R3 | 13 |
| Cluster_2 | MF | GO:001683 | carbon-oxygen lyase activity | 6/419 | 16/3517 | 0.375 | 3.147673031 | 3.166086949 | 0.007750561 | 0.727919186 | 0.727919186 | LOC100847874/PCBD1/C | 6 |
| Cluster_2 | BP | GO:000674 | glutathione metabolic process | 6/442 | 16/3586 | 0.375 | 3.042420814 | 3.069626652 | 0.009160677 | 0.839437462 | 0.839437462 | GSTT1/GSR/GSTM2/GST | 6 |
| Cluster_2 | BP | GO:007162 | granulocyte chemotaxis | 6/442 | 16/3586 | 0.375 | 3.042420814 | 3.069626652 | 0.009160677 | 0.839437462 | 0.839437462 | S100A9/LOC508666/S100 | 6 |
| Cluster_2 | BP | GO:006102 | membrane fusion | 8/442 | 26/3586 | 0.307692308 | 2.496345284 | 2.870833068 | 0.010367559 | 0.839437462 | 0.839437462 | STXBP6/VAMP3/VAMP8/ | 8 |
| Cluster_2 | BP | GO:000854 | fibroblast growth factor receptor signaling pathway | 5/442 | 12/3586 | 0.416666667 | 3.380467572 | 3.096635331 | 0.010503499 | 0.839437462 | 0.839437462 | FGF5/SPRY1/FGF2/FGFR | 5 |
| Cluster_2 | BP | GO:004434 | cellular response to fibroblast growth factor stimulus | 5/442 | 12/3586 | 0.416666667 | 3.380467572 | 3.096635331 | 0.010503499 | 0.839437462 | 0.839437462 | FGF5/SPRY1/FGF2/FGFR | 5 |
| Cluster_2 | BP | GO:007177 | response to fibroblast growth factor | 5/442 | 12/3586 | 0.416666667 | 3.380467572 | 3.096635331 | 0.010503499 | 0.839437462 | 0.839437462 | FGF5/SPRY1/FGF2/FGFR | 5 |
| Cluster_2 | MF | GO:003509 | phosphatidylinositol binding | 9/419 | 33/3517 | 0.272727273 | 2.28921675 | 2.736116431 | 0.012436056 | 0.727919186 | 0.727919186 | STXBP6/LOC101902366/ | 9 |
| Cluster_2 | BP | GO:004828 | organelle fusion | 8/442 | 27/3586 | 0.296296296 | 2.403888051 | 2.745141982 | 0.013188802 | 0.839437462 | 0.839437462 | STXBP6/VAMP3/VAMP8/ | 8 |
| Cluster_2 | BP | GO:001062 | negative regulation of gene expression | 12/442 | 49/3586 | 0.244897959 | 1.986887063 | 2.60772873 | 0.013470261 | 0.839437462 | 0.839437462 | NANOS1/TYROBP/DDX6/ | 12 |
| Cluster_2 | BP | GO:006102 | membrane organization | 17/442 | 79/3586 | 0.215189873 | 1.745861733 | 2.513150139 | 0.013886986 | 0.839437462 | 0.839437462 | S100A9/STXBP6/VAMP3/ | 17 |
| Cluster_2 | BP | GO:004212 | regulation of cell population proliferation | 14/442 | 61/3586 | 0.229508197 | 1.862028039 | 2.545781905 | 0.014153169 | 0.839437462 | 0.839437462 | IL1B/FGF5/TYROBP/FGF | 14 |
| Cluster_2 | MF | GO:005108 | protein-folding chaperone binding | 7/419 | 23/3517 | 0.304347826 | 2.554633185 | 2.750560104 | 0.014449615 | 0.727919186 | 0.727919186 | LOC614226/BAG5/LOC61 | 7 |
| Cluster_2 | MF | GO:004698 | protein heterodimerization activity | 6/419 | 18/3517 | 0.333333333 | 2.797931583 | 2.812083297 | 0.014607074 | 0.727919186 | 0.727919186 | POLE3/BCL2L1/TAF11/D | 6 |
| Cluster_2 | BP | GO:004533 | cellular respiration | 9/442 | 33/3586 | 0.272727273 | 2.212669683 | 2.623713033 | 0.015390139 | 0.839437462 | 0.839437462 | SDHD/NDUFC2/SDHC/C | 9 |
| Cluster_2 | CC | GO:000574 | mitochondrial envelope | 14/435 | 61/3473 | 0.229508197 | 1.832372338 | 2.481524958 | 0.01613358 | 0.423235633 | 0.423235633 | SDHD/CHCHD3/COX19/T | 14 |
| Cluster_2 | BP | GO:001055 | negative regulation of macromolecule biosynthetic process | 27/442 | 145/3586 | 0.186206897 | 1.510719301 | 2.353631033 | 0.016596099 | 0.839437462 | 0.839437462 | NANOS1/ZNF397/TYROB | 27 |
| Cluster_2 | CC | GO:003196 | organelle envelope | 18/435 | 86/3473 | 0.209302326 | 1.671050521 | 2.384171254 | 0.017799629 | 0.423235633 | 0.423235633 | SDHD/CHCHD3/COX19/T | 18 |
| Cluster_2 | MF | GO:000813 | transcription factor binding | 9/419 | 35/3517 | 0.257142857 | 2.158404364 | 2.532623088 | 0.01835954 | 0.761591815 | 0.761591815 | NR4A1/NFAT5/LRIF1/PE | 9 |
| Cluster_2 | BP | GO:000828 | cell population proliferation | 14/442 | 63/3586 | 0.222222222 | 1.802919038 | 2.410452247 | 0.01866717 | 0.839437462 | 0.839437462 | IL1B/FGF5/TYROBP/FGF | 14 |
| Cluster_2 | BP | GO:000625 | DNA metabolic process | 13/442 | 57/3586 | 0.228070175 | 1.850361197 | 2.426216094 | 0.018681201 | 0.839437462 | 0.839437462 | LOC112445063/SWSAP1/ | 13 |
| Cluster_2 | BP | GO:000989 | negative regulation of biosynthetic process | 27/442 | 147/3586 | 0.183673469 | 1.490165297 | 2.275095262 | 0.019678593 | 0.839437462 | 0.839437462 | NANOS1/ZNF397/TYROB | 27 |
| Cluster_2 | BP | GO:001590 | fatty acid transport | 5/442 | 14/3586 | 0.357142857 | 2.897543633 | 2.666947766 | 0.02154356 | 0.839437462 | 0.839437462 | RBP1/CRABP2/FABP5/P | 5 |
| Cluster_2 | BP | GO:001050 | regulation of autophagy | 6/442 | 19/3586 | 0.315789474 | 2.562038581 | 2.559360412 | 0.022510055 | 0.839437462 | 0.839437462 | RRAGD/PLEKHF1/TBC1D | 6 |
| Cluster_2 | BP | GO:001964 | aerobic electron transport chain | 6/442 | 19/3586 | 0.315789474 | 2.562038581 | 2.559360412 | 0.022510055 | 0.839437462 | 0.839437462 | SDHD/NDUFC2/SDHC/C | 6 |
| Cluster_2 | BP | GO:003059 | leukocyte chemotaxis | 6/442 | 19/3586 | 0.315789474 | 2.562038581 | 2.559360412 | 0.022510055 | 0.839437462 | 0.839437462 | S100A9/LOC508666/S100 | 6 |
| Cluster_2 | BP | GO:009753 | granulocyte migration | 6/442 | 19/3586 | 0.315789474 | 2.562038581 | 2.559360412 | 0.022510055 | 0.839437462 | 0.839437462 | S100A9/LOC508666/S100 | 6 |
| Cluster_2 | MF | GO:000408 | carbonate dehydratase activity | 4/419 | 10/3517 | 0.4 | 3.3575179 | 2.745224423 | 0.02317882 | 0.761591815 | 0.761591815 | LOC100847874/CA2/CA1 | 4 |
| Cluster_2 | MF | GO:000550 | fatty acid binding | 4/419 | 10/3517 | 0.4 | 3.3575179 | 2.745224423 | 0.02317882 | 0.761591815 | 0.761591815 | RBP1/CRABP2/FABP5/C | 4 |
| Cluster_2 | CC | GO:000574 | mitochondrial outer membrane | 6/435 | 19/3473 | 0.315789474 | 2.52123412 | 2.515662022 | 0.024200651 | 0.423731976 | 0.423731976 | MIEF2/BCL2L1/SNN/CYB | 6 |
| Cluster_2 | CC | GO:001986 | outer membrane | 6/435 | 19/3473 | 0.315789474 | 2.52123412 | 2.515662022 | 0.024200651 | 0.423731976 | 0.423731976 | MIEF2/BCL2L1/SNN/CYB | 6 |
| Cluster_2 | CC | GO:003196 | organelle outer membrane | 6/435 | 19/3473 | 0.315789474 | 2.52123412 | 2.515662022 | 0.024200651 | 0.423731976 | 0.423731976 | MIEF2/BCL2L1/SNN/CYB | 6 |
| Cluster_2 | BP | GO:000906 | aerobic respiration | 8/442 | 30/3586 | 0.266666667 | 2.163499246 | 2.399166029 | 0.024979205 | 0.894255534 | 0.894255534 | SDHD/NDUFC2/SDHC/C | 8 |
| Cluster_2 | MF | GO:001683 | hydro-lyase activity | 5/419 | 15/3517 | 0.333333333 | 2.797931583 | 2.565969312 | 0.025471298 | 0.761591815 | 0.761591815 | LOC100847874/PCBD1/C | 5 |
| Cluster_2 | CC | GO:003198 | nuclear lumen | 27/435 | 148/3473 | 0.182432432 | 1.456523765 | 2.147528941 | 0.025740728 | 0.423731976 | 0.423731976 | LSM6/RRS1/MAD2L1/ZNF | 27 |
| Cluster_2 | BP | GO:000611 | oxidative phosphorylation | 6/442 | 20/3586 | 0.3 | 2.433936652 | 2.41084195 | 0.028911125 | 0.924123463 | 0.924123463 | SDHD/NDUFC2/SDHC/C | 6 |
| Cluster_2 | BP | GO:004277 | ATP synthesis coupled electron transport | 6/442 | 20/3586 | 0.3 | 2.433936652 | 2.41084195 | 0.028911125 | 0.924123463 | 0.924123463 | SDHD/NDUFC2/SDHC/C | 6 |
| Cluster_2 | BP | GO:004277 | mitochondrial ATP synthesis coupled electron transport | 6/442 | 20/3586 | 0.3 | 2.433936652 | 2.41084195 | 0.028911125 | 0.924123463 | 0.924123463 | SDHD/NDUFC2/SDHC/C | 6 |
| Cluster_2 | MF | GO:000548 | SNAP receptor activity | 4/419 | 11/3517 | 0.363636364 | 3.052289 | 2.506799746 | 0.033071803 | 0.824039081 | 0.824039081 | VAMP3/VAMP8/STX7/SE | 4 |
| Cluster_2 | MF | GO:000837 | galactosyltransferase activity | 4/419 | 11/3517 | 0.363636364 | 3.052289 | 2.506799746 | 0.033071803 | 0.824039081 | 0.824039081 | A4GALT/B3GALT2/C1GA | 4 |
| Cluster_2 | BP | GO:001060 | negative regulation of macromolecule metabolic process | 33/442 | 196/3586 | 0.168367347 | 1.365984855 | 1.975630613 | 0.035007776 | 0.998839738 | 0.998839738 | NANOS1/ZNF397/TYROB | 33 |

| | | | | | | | | | | | | | |
|---|---|---|---|---|---|---|---|---|---|---|---|---|---|
| Cluster_2 | BP | GO:000828 | negative regulation of cell population proliferation | 6/442 | 21/3586 | 0.285714286 | 2.318034906 | 2.271021461 | 0.036402376 | 0.998839738 | 0.998839738 | TYROBP/NUPR1/CDKN1 | 6 |
| Cluster_2 | BP | GO:002290 | respiratory electron transport chain | 6/442 | 21/3586 | 0.285714286 | 2.318034906 | 2.271021461 | 0.036402376 | 0.998839738 | 0.998839738 | SDHD/NDUFC2/SDHC/CC | 6 |
| Cluster_2 | BP | GO:009752 | myeloid leukocyte migration | 6/442 | 21/3586 | 0.285714286 | 2.318034906 | 2.271021461 | 0.036402376 | 0.998839738 | 0.998839738 | S100A9/LOC508666/S100 | 6 |
| Cluster_2 | BP | GO:001571 | monocarboxylic acid transport | 5/442 | 16/3586 | 0.3125 | 2.535350679 | 2.307532968 | 0.03820266 | 0.998839738 | 0.998839738 | RBP1/CRABP2/FABP5/PF | 5 |
| Cluster_2 | CC | GO:003120 | SNARE complex | 4/435 | 11/3473 | 0.363636364 | 2.903239289 | 2.39203102 | 0.038968403 | 0.595659878 | 0.595659878 | VAMP3/VAMP8/STX7/SE | 4 |
| Cluster_2 | BP | GO:000989 | negative regulation of metabolic process | 34/442 | 206/3586 | 0.165048544 | 1.339058999 | 1.879165289 | 0.042161162 | 0.998839738 | 0.998839738 | NANOS1/ZNF397/RRAGD | 34 |
| Cluster_2 | BP | GO:000657 | modified amino acid metabolic process | 8/442 | 33/3586 | 0.242424242 | 1.966817496 | 2.091791112 | 0.042663819 | 0.998839738 | 0.998839738 | GSTT1/VNN1/GSR/GSTM | 8 |
| Cluster_2 | BP | GO:190253 | positive regulation of intracellular signal transduction | 14/442 | 70/3586 | 0.2 | 1.622624434 | 1.972264661 | 0.043243424 | 0.998839738 | 0.998839738 | LAMTOR3/PLEKHF1/S10 | 14 |
| Cluster_2 | BP | GO:001605 | vesicle organization | 10/442 | 45/3586 | 0.222222222 | 1.802916038 | 2.032019506 | 0.043349271 | 0.998839738 | 0.998839738 | STXBP6/VAMP3/VAMP8/ | 10 |
| Cluster_2 | BP | GO:000628 | DNA repair | 9/442 | 39/3586 | 0.230769231 | 1.872258963 | 2.053343444 | 0.043446768 | 0.998839738 | 0.998839738 | LOC112445063/SWSAP1/ | 9 |
| Cluster_2 | BP | GO:002290 | electron transport chain | 6/442 | 22/3586 | 0.272727273 | 2.212669683 | 2.13894422 | 0.045032155 | 0.998839738 | 0.998839738 | SDHD/NDUFC2/SDHC/CC | 6 |
| Cluster_2 | CC | GO:000575 | mitochondrial matrix | 8/435 | 33/3473 | 0.242424242 | 1.93549286 | 2.042955127 | 0.04628532 | 0.606330899 | 0.606330899 | MRPL16/MRPL57/MRPL5 | 8 |
| Cluster_2 | CC | GO:003196 | mitochondrial membrane | 11/435 | 51/3473 | 0.215686275 | 1.722019382 | 1.965325747 | 0.04711689 | 0.606330899 | 0.606330899 | CHCHD3/TIMM23/MIEF2/ | 11 |
| Cluster_2 | BP | GO:190253 | regulation of intracellular signal transduction | 23/442 | 131/3586 | 0.175572519 | 1.424441297 | 1.85542627 | 0.047500622 | 0.998839738 | 0.998839738 | LAMTOR3/PLEKHF1/S10 | 23 |
| Cluster_2 | CC | GO:001660 | nuclear body | 6/435 | 22/3473 | 0.272727273 | 2.177429467 | 2.096111416 | 0.048166473 | 0.606330899 | 0.606330899 | TRIM52/SNURF/PPP1R8/ | 6 |
| Cluster_6 | BP | GO:000762 | circadian rhythm | 8/394 | 18/3586 | 0.444444444 | 4.045121263 | 4.549772393 | 0.000319916 | 0.218347569 | 0.218347569 | CIART/CIPC/ID1/PASD1/I | 8 |
| Cluster_6 | MF | GO:000370 | DNA-binding transcription factor activity | 58/369 | 360/3517 | 0.161111111 | 1.535576634 | 3.671604005 | 0.000337823 | 0.098982179 | 0.098982179 | ZNF180/ZNF275/ATMIN/T | 58 |
| Cluster_6 | BP | GO:004851 | rhythmic process | 8/394 | 19/3586 | 0.421052632 | 3.832220144 | 4.348239959 | 0.000500224 | 0.218347569 | 0.218347569 | CIART/CIPC/ID1/PASD1/I | 8 |
| Cluster_6 | BP | GO:003292 | circadian regulation of gene expression | 6/394 | 13/3586 | 0.461538462 | 4.20070285 | 4.061269169 | 0.001480709 | 0.430886229 | 0.430886229 | CIART/ID1/PASD1/ID4/BH | 6 |
| Cluster_6 | MF | GO:001620 | antioxidant activity | 7/369 | 18/3517 | 0.388888889 | 3.706564288 | 3.940967768 | 0.001506592 | 0.145927418 | 0.145927418 | GPX1/GPX8/GPX7/PRDX | 7 |
| Cluster_6 | MF | GO:001668 | oxidoreductase activity, acting on peroxide as acceptor | 6/369 | 14/3517 | 0.428571429 | 4.084785134 | 3.95903301 | 0.001847503 | 0.145927418 | 0.145927418 | GPX1/GPX8/GPX7/PRDX | 6 |
| Cluster_6 | MF | GO:000098 | DNA-binding transcription factor activity, RNA polymerase II-specific | 52/369 | 340/3517 | 0.152941176 | 1.457707636 | 3.039759772 | 0.002394824 | 0.145927418 | 0.145927418 | ZNF180/ZNF275/ATMIN/T | 52 |
| Cluster_6 | MF | GO:014011 | transcription regulator activity | 64/369 | 439/3517 | 0.145785877 | 1.389509294 | 2.986317857 | 0.002556935 | 0.145927418 | 0.145927418 | NAB1/ZNF180/ZNF275/AT | 64 |
| Cluster_6 | CC | GO:000989 | external side of plasma membrane | 14/359 | 63/3473 | 0.222222222 | 2.149798824 | 3.12674687 | 0.004076441 | 0.693648781 | 0.693648781 | TNFRSF4/CCRL2/IL4R/IT | 14 |
| Cluster_6 | MF | GO:000369 | double-stranded DNA binding | 48/369 | 319/3517 | 0.150470219 | 1.434156536 | 2.783701796 | 0.0049298 | 0.145927418 | 0.145927418 | CIART/ZNF180/ZNF275/A | 48 |
| Cluster_6 | MF | GO:000460 | peroxidase activity | 5/369 | 12/3517 | 0.416666667 | 3.97131888 | 3.529523914 | 0.005248103 | 0.145927418 | 0.145927418 | GPX1/GPX8/GPX7/PRDX | 5 |
| Cluster_6 | MF | GO:004356 | sequence-specific DNA binding | 51/369 | 345/3517 | 0.147826087 | 1.408954872 | 2.738032503 | 0.005435464 | 0.145927418 | 0.145927418 | CIART/ZNF180/ZNF275/A | 51 |
| Cluster_6 | MF | GO:000367 | DNA binding | 62/369 | 436/3517 | 0.142201835 | 1.355349196 | 2.713679685 | 0.005438418 | 0.145927418 | 0.145927418 | CIART/ZNF180/ZNF275/Z | 62 |
| Cluster_6 | MF | GO:000097 | transcription cis-regulatory region binding | 47/369 | 314/3517 | 0.149681529 | 1.426639394 | 2.711861857 | 0.00597655 | 0.145927418 | 0.145927418 | CIART/ZNF180/ZNF275/A | 47 |
| Cluster_6 | MF | GO:000106 | transcription regulatory region nucleic acid binding | 47/369 | 314/3517 | 0.149681529 | 1.426639394 | 2.711861857 | 0.00597655 | 0.145927418 | 0.145927418 | CIART/ZNF180/ZNF275/A | 47 |
| Cluster_6 | MF | GO:199083 | sequence-specific double-stranded DNA binding | 47/369 | 314/3517 | 0.149681529 | 1.426639394 | 2.711861857 | 0.00597655 | 0.145927418 | 0.145927418 | CIART/ZNF180/ZNF275/A | 47 |
| Cluster_6 | CC | GO:000998 | cell surface | 19/359 | 102/3473 | 0.18627451 | 1.802037249 | 2.791213307 | 0.007225508 | 0.693648781 | 0.693648781 | TNFRSF4/CCRL2/IL4R/IT | 19 |
| Cluster_6 | MF | GO:000097 | RNA polymerase II transcription regulatory region sequence-specific DNA | 45/369 | 301/3517 | 0.149501661 | 1.424925047 | 2.639111948 | 0.007284919 | 0.164190872 | 0.164190872 | CIART/ZNF180/ZNF275/T | 45 |
| Cluster_6 | BP | GO:000636 | transcription by RNA polymerase II | 68/394 | 477/3586 | 0.142557652 | 1.297491726 | 2.451235115 | 0.01038553 | 0.988129247 | 0.988129247 | ZNF180/ZNF275/CHD2/A | 68 |
| Cluster_6 | BP | GO:000635 | regulation of transcription by RNA polymerase II | 66/394 | 465/3586 | 0.141935484 | 1.291829049 | 2.369569008 | 0.012760561 | 0.988129247 | 0.988129247 | ZNF180/ZNF275/CHD2/A | 66 |
| Cluster_6 | MF | GO:000097 | RNA polymerase II cis-regulatory region sequence-specific DNA binding | 39/369 | 263/3517 | 0.148288973 | 1.413366719 | 2.385749615 | 0.013948053 | 0.291912831 | 0.291912831 | CIART/ZNF180/ZNF275/R | 39 |
| Cluster_6 | BP | GO:000697 | response to oxidative stress | 7/394 | 25/3586 | 0.28 | 2.548426396 | 2.729202957 | 0.015184615 | 0.988129247 | 0.988129247 | GPX1/GPX8/GPX7/PRDX | 7 |
| Cluster_6 | BP | GO:001995 | sexual reproduction | 7/394 | 25/3586 | 0.28 | 2.548426396 | 2.729202957 | 0.015184615 | 0.988129247 | 0.988129247 | CFAP97/ERCC4/CCDC13 | 7 |
| Cluster_6 | MF | GO:000098 | cis-regulatory region sequence-specific DNA binding | 39/369 | 265/3517 | 0.147169811 | 1.402699801 | 2.333723167 | 0.015706702 | 0.306804246 | 0.306804246 | CIART/ZNF180/ZNF275/R | 39 |
| Cluster_6 | BP | GO:006027 | cilium assembly | 6/394 | 20/3586 | 0.3 | 2.730456853 | 2.726128688 | 0.017158217 | 0.988129247 | 0.988129247 | TMEM216/LOC101902172 | 6 |
| Cluster_6 | BP | GO:000190 | leukocyte mediated cytotoxicity | 4/394 | 10/3586 | 0.4 | 3.640609137 | 2.937423846 | 0.01757063 | 0.988129247 | 0.988129247 | JSP.1/LOC616942/BoLA/ | 4 |
| Cluster_6 | BP | GO:000191 | regulation of leukocyte mediated cytotoxicity | 4/394 | 10/3586 | 0.4 | 3.640609137 | 2.937423846 | 0.01757063 | 0.988129247 | 0.988129247 | JSP.1/LOC616942/BoLA/ | 4 |
| Cluster_6 | BP | GO:000191 | positive regulation of leukocyte mediated cytotoxicity | 4/394 | 10/3586 | 0.4 | 3.640609137 | 2.937423846 | 0.01757063 | 0.988129247 | 0.988129247 | JSP.1/LOC616942/BoLA/ | 4 |
| Cluster_6 | BP | GO:000717 | epidermal growth factor receptor signaling pathway | 4/394 | 10/3586 | 0.4 | 3.640609137 | 2.937423846 | 0.01757063 | 0.988129247 | 0.988129247 | SOCS5/SOCS4/BTC/ERR | 4 |
| Cluster_6 | BP | GO:003134 | regulation of cell killing | 4/394 | 10/3586 | 0.4 | 3.640609137 | 2.937423846 | 0.01757063 | 0.988129247 | 0.988129247 | JSP.1/LOC616942/BoLA/ | 4 |
| Cluster_6 | BP | GO:003134 | positive regulation of cell killing | 4/394 | 10/3586 | 0.4 | 3.640609137 | 2.937423846 | 0.01757063 | 0.988129247 | 0.988129247 | JSP.1/LOC616942/BoLA/ | 4 |
| Cluster_6 | MF | GO:000486 | protein kinase inhibitor activity | 4/369 | 11/3517 | 0.363636364 | 3.465878295 | 2.804027568 | 0.021617685 | 0.372587151 | 0.372587151 | PKIB/CDKN2B/ITPRIP/SH | 4 |
| Cluster_6 | MF | GO:001921 | kinase inhibitor activity | 4/369 | 11/3517 | 0.363636364 | 3.465878295 | 2.804027568 | 0.021617685 | 0.372587151 | 0.372587151 | PKIB/CDKN2B/ITPRIP/SH | 4 |
| Cluster_6 | BP | GO:004478 | cilium organization | 6/394 | 21/3586 | 0.285714286 | 2.600435098 | 2.583920794 | 0.021858354 | 0.988129247 | 0.988129247 | TMEM216/LOC101902172 | 6 |
| Cluster_6 | BP | GO:000246 | adaptive immune response based on somatic recombination of immune receptors built from immunoglobulin | 5/394 | 16/3586 | 0.3125 | 2.844225888 | 2.597175199 | 0.024445032 | 0.988129247 | 0.988129247 | IL18BP/JSP.1/LOC616942 | 5 |
| Cluster_6 | BP | GO:007092 | organelle assembly | 13/394 | 66/3586 | 0.196969697 | 1.792724196 | 2.283410532 | 0.024872306 | 0.988129247 | 0.988129247 | TCAP/TMOD4/CCDC136/ | 13 |

| | | | | | | | | | | | | | |
|---|---|---|---|---|---|---|---|---|---|---|---|---|---|
| Cluster_6 | BP | GO:000072 | double-strand break repair via homologous recombination | 4/394 | 11/3586 | 0.363636364 | 3.30964467 | 2.695037047 | 0.025263142 | 0.988129247 | 0.988129247 | ERCC4/RMI1/LOC101907 | 4 |
| Cluster_6 | BP | GO:000072 | recombinational repair | 4/394 | 11/3586 | 0.363636364 | 3.30964467 | 2.695037047 | 0.025263142 | 0.988129247 | 0.988129247 | ERCC4/RMI1/LOC101907 | 4 |
| Cluster_6 | BP | GO:000190 | cell killing | 4/394 | 11/3586 | 0.363636364 | 3.30964467 | 2.695037047 | 0.025263142 | 0.988129247 | 0.988129247 | JSP.1/LOC616942/BoLA/E | 4 |
| Cluster_6 | BP | GO:000245 | T cell mediated immunity | 4/394 | 11/3586 | 0.363636364 | 3.30964467 | 2.695037047 | 0.025263142 | 0.988129247 | 0.988129247 | JSP.1/LOC616942/BoLA/E | 4 |
| Cluster_6 | BP | GO:003812 | ERBB signaling pathway | 4/394 | 11/3586 | 0.363636364 | 3.30964467 | 2.695037047 | 0.025263142 | 0.988129247 | 0.988129247 | SOCS5/SOCS4/BTC/ERR | 4 |
| Cluster_6 | MF | GO:004227 | peptide binding | 6/369 | 23/3517 | 0.260869565 | 2.486390951 | 2.448250066 | 0.02751741 | 0.44792229 | 0.44792229 | RANBP6/JSP.1/APOE/LO | 6 |
| Cluster_6 | BP | GO:004589 | negative regulation of DNA-templated transcription | 17/394 | 95/3586 | 0.178947368 | 1.628693561 | 2.181660358 | 0.027523588 | 0.988129247 | 0.988129247 | CIART/NAB1/CIPC/ID1/PA | 17 |
| Cluster_6 | CC | GO:001632 | basolateral plasma membrane | 4/359 | 12/3473 | 0.333333333 | 3.224698236 | 2.62082916 | 0.028366436 | 0.962272701 | 0.962272701 | AQP7/PDPN/TAX1BP3/DL | 4 |
| Cluster_6 | BP | GO:190267 | negative regulation of RNA biosynthetic process | 17/394 | 96/3586 | 0.177083333 | 1.611728003 | 2.13423636 | 0.03019145 | 0.988129247 | 0.988129247 | CIART/NAB1/CIPC/ID1/PA | 17 |
| Cluster_6 | CC | GO:009855 | side of membrane | 15/359 | 87/3473 | 0.172413793 | 1.667947363 | 2.142082374 | 0.030999623 | 0.962272701 | 0.962272701 | TNFRSF4/CCRL2/IL4R/IT | 15 |
| Cluster_6 | BP | GO:000270 | positive regulation of leukocyte mediated immunity | 4/394 | 12/3586 | 0.333333333 | 3.033840948 | 2.479085957 | 0.03468647 | 0.988129247 | 0.988129247 | JSP.1/LOC616942/BoLA/E | 4 |
| Cluster_6 | BP | GO:000270 | regulation of lymphocyte mediated immunity | 4/394 | 12/3586 | 0.333333333 | 3.033840948 | 2.479085957 | 0.03468647 | 0.988129247 | 0.988129247 | JSP.1/LOC616942/BoLA/E | 4 |
| Cluster_6 | BP | GO:012003 | plasma membrane bounded cell projection assembly | 8/394 | 36/3586 | 0.222222222 | 2.022560632 | 2.166144338 | 0.037775335 | 0.988129247 | 0.988129247 | TMEM216/LOC101902172 | 8 |
| Cluster_6 | BP | GO:000225 | adaptive immune response | 5/394 | 18/3586 | 0.277777778 | 2.52820079 | 2.283313261 | 0.039928317 | 0.988129247 | 0.988129247 | IL18BP/JSP.1/LOC616942 | 5 |
| Cluster_6 | BP | GO:000905 | catabolic process | 39/394 | 271/3586 | 0.143911439 | 1.30981325 | 1.863391529 | 0.042526193 | 0.988129247 | 0.988129247 | SOCS5/SEL1L/ENDOG/G | 39 |
| Cluster_6 | BP | GO:005125 | negative regulation of RNA metabolic process | 17/394 | 100/3586 | 0.17 | 1.547258883 | 1.949801401 | 0.042807335 | 0.988129247 | 0.988129247 | CIART/NAB1/CIPC/ID1/PA | 17 |
| Cluster_6 | BP | GO:000281 | regulation of adaptive immune response | 4/394 | 13/3586 | 0.307692308 | 2.800468567 | 2.284556806 | 0.045879858 | 0.988129247 | 0.988129247 | JSP.1/LOC616942/BoLA/E | 4 |
| Cluster_6 | BP | GO:000282 | regulation of adaptive immune response based on somatic recombination of immune receptors built from immunoglobulin superfamily domains | 4/394 | 13/3586 | 0.307692308 | 2.800468567 | 2.284556806 | 0.045879858 | 0.988129247 | 0.988129247 | JSP.1/LOC616942/BoLA/E | 4 |
| Cluster_6 | BP | GO:003070 | cytoskeleton-dependent intracellular transport | 4/394 | 13/3586 | 0.307692308 | 2.800468567 | 2.284556806 | 0.045879858 | 0.988129247 | 0.988129247 | CCDC88C/LOC101902172 | 4 |
| Cluster_6 | BP | GO:000695 | immune response | 24/394 | 154/3586 | 0.155844156 | 1.418419144 | 1.864487677 | 0.04634815 | 0.988129247 | 0.988129247 | IL18BP/CCRL2/TRIM59/LT | 24 |
| Cluster_6 | MF | GO:000510 | signaling receptor binding | 30/369 | 211/3517 | 0.142180095 | 1.355141987 | 1.821428292 | 0.04820409 | 0.711158475 | 0.711158475 | EFNA4/ITGA5/TSHB/HCS | 30 |
| Cluster_6 | CC | GO:000992 | basal plasma membrane | 4/359 | 14/3473 | 0.285714286 | 2.764027059 | 2.245286463 | 0.048631422 | 0.962272701 | 0.962272701 | AQP7/PDPN/TAX1BP3/DL | 4 |
| Cluster_6 | CC | GO:007101 | precatalytic spliceosome | 4/359 | 14/3473 | 0.285714286 | 2.764027059 | 2.245286463 | 0.048631422 | 0.962272701 | 0.962272701 | PRPF38B/LOC107131494 | 4 |
| Cluster_6 | BP | GO:000004 | transition metal ion transport | 5/394 | 19/3586 | 0.263157895 | 2.39513759 | 2.141921429 | 0.049473167 | 0.988129247 | 0.988129247 | ATP2C2/SLC30A1/LOC50 | 5 |

**Supplementary Table 22. Gene Ontology enrichment of whole-blood connectivity clusters C4 and C5 identified from telomere length-ordered sliding-window analysis, related to Figure 5.**

| Cluster | ONTOLOGY | ID | Description | GeneRatio | BgRatio | RichFactor | FoldEnrichment | zScore | pvalue | p.adjust | qvalue | geneID | Count |
|---|---|---|---|---|---|---|---|---|---|---|---|---|---|
| Cluster_4 | BP | GO:000695 | response to stress | 89/524 | 263/2312 | 0.338403042 | 1.493106551 | 4.597582449 | 6.80113E-06 | 0.004142534 | 0.004142534 | C5AR1/AGER/RAD1/PPP | 89 |
| Cluster_4 | BP | GO:000695 | defense response | 56/524 | 149/2312 | 0.375838926 | 1.658281674 | 4.496333319 | 1.48468E-05 | 0.004142534 | 0.004142534 | C5AR1/AGER/OAS1Z/IL1 | 56 |
| Cluster_4 | BP | GO:000704 | cell cycle | 51/524 | 132/2312 | 0.386363636 | 1.704718945 | 4.512908464 | 1.50638E-05 | 0.004142534 | 0.004142534 | CDCA8/PSMC3IP/NCAPG | 51 |
| Cluster_4 | BP | GO:000960 | response to external stimulus | 61/524 | 174/2312 | 0.350574713 | 1.546810564 | 4.059658752 | 7.27502E-05 | 0.015004733 | 0.015004733 | C5AR1/JAML/AGER/SEM | 61 |
| Cluster_4 | BP | GO:001995 | sexual reproduction | 19/524 | 38/2312 | 0.5 | 2.20610687 | 4.05755316 | 0.000181434 | 0.026026931 | 0.026026931 | PSMC3IP/SYCE2/NUF2/R | 19 |
| Cluster_4 | BP | GO:002241 | reproductive process | 20/524 | 41/2312 | 0.487804878 | 2.152299386 | 4.029308784 | 0.000189287 | 0.026026931 | 0.026026931 | PSMC3IP/SYCE2/NUF2/R | 20 |
| Cluster_4 | BP | GO:000225 | activation of immune response | 22/524 | 49/2312 | 0.448979592 | 1.980993924 | 3.756677724 | 0.000409325 | 0.048241906 | 0.048241906 | C5AR1/PTGDR2/RIGI/IFIH | 22 |
| Cluster_4 | BP | GO:005160 | defense response to virus | 12/524 | 21/2312 | 0.571428571 | 2.521264995 | 3.790389442 | 0.000624786 | 0.064431012 | 0.064431012 | OAS1Z/RIGI/IFIH1/OASL/I | 12 |
| Cluster_4 | MF | GO:003802 | signaling receptor activity | 71/540 | 219/2316 | 0.324200913 | 1.390461695 | 3.347743487 | 0.000768774 | 0.077121259 | 0.077121259 | GPR141/C5AR1/TNFRSF | 71 |
| Cluster_4 | MF | GO:006008 | molecular transducer activity | 71/540 | 219/2316 | 0.324200913 | 1.390461695 | 3.347743487 | 0.000768774 | 0.077121259 | 0.077121259 | GPR141/C5AR1/TNFRSF | 71 |
| Cluster_4 | BP | GO:002241 | cellular process involved in reproduction in multicellular organism | 9/524 | 14/2312 | 0.642857143 | 2.836423119 | 3.730298762 | 0.000979101 | 0.082903011 | 0.082903011 | RSPH1/CCDC136/FOLR3 | 9 |
| Cluster_4 | BP | GO:000960 | response to biotic stimulus | 38/524 | 106/2312 | 0.358490566 | 1.581737001 | 3.318629164 | 0.001105373 | 0.082903011 | 0.082903011 | OAS1Z/STAT2/PGLYRP1/ | 38 |
| Cluster_4 | BP | GO:002240 | cell cycle process | 38/524 | 106/2312 | 0.358490566 | 1.581737001 | 3.318629164 | 0.001105373 | 0.082903011 | 0.082903011 | CDCA8/PSMC3IP/NCAPG | 38 |
| Cluster_4 | CC | GO:000022 | nuclear chromosome | 9/519 | 14/2248 | 0.642857143 | 2.78447564 | 3.668772446 | 0.001129526 | 0.176206078 | 0.176206078 | PSMC3IP/RAD1/SYCE2/P | 9 |
| Cluster_4 | MF | GO:001653 | cyclin-dependent protein serine/threonine kinase regulator activity | 9/540 | 14/2316 | 0.642857143 | 2.757142857 | 3.635543341 | 0.001220794 | 0.077121259 | 0.077121259 | CCNB1/CCNF/CDK5R1/C | 9 |
| Cluster_4 | MF | GO:001991 | cyclin-dependent protein kinase regulator activity | 9/540 | 14/2316 | 0.642857143 | 2.757142857 | 3.635543341 | 0.001220794 | 0.077121259 | 0.077121259 | CCNB1/CCNF/CDK5R1/C | 9 |
| Cluster_4 | BP | GO:000275 | immune response-activating signaling pathway | 18/524 | 40/2312 | 0.45 | 1.985496183 | 3.403005396 | 0.00133083 | 0.084093009 | 0.084093009 | C5AR1/PTGDR2/RIGI/IFIH | 18 |
| Cluster_4 | MF | GO:001988 | protein kinase regulator activity | 12/540 | 22/2316 | 0.545454545 | 2.339393939 | 3.479956858 | 0.001433481 | 0.077121259 | 0.077121259 | CCNB1/CCNF/CDK5R1/C | 12 |
| Cluster_4 | BP | GO:004320 | response to external biotic stimulus | 37/524 | 104/2312 | 0.355769231 | 1.569729888 | 3.217866315 | 0.001507592 | 0.084093009 | 0.084093009 | OAS1Z/STAT2/PGLYRP1/ | 37 |
| Cluster_4 | BP | GO:005170 | response to other organism | 37/524 | 104/2312 | 0.355769231 | 1.569729888 | 3.217866315 | 0.001507592 | 0.084093009 | 0.084093009 | OAS1Z/STAT2/PGLYRP1/ | 37 |
| Cluster_4 | BP | GO:004860 | multicellular organismal reproductive process | 11/524 | 20/2312 | 0.55 | 2.426717557 | 3.468387049 | 0.001625946 | 0.084093009 | 0.084093009 | RSPH1/SPATA6L/CCDC1 | 11 |
| Cluster_4 | BP | GO:000027 | mitotic cell cycle | 30/524 | 80/2312 | 0.375 | 1.654580153 | 3.225095583 | 0.001630895 | 0.084093009 | 0.084093009 | CDCA8/NCAPG/CCNB1/S | 30 |
| Cluster_4 | BP | GO:000961 | response to virus | 12/524 | 23/2312 | 0.52173913 | 2.30202456 | 3.396581635 | 0.001826603 | 0.088643952 | 0.088643952 | OAS1Z/RIGI/IFIH1/OASL/I | 12 |
| Cluster_4 | BP | GO:000695 | inflammatory response | 23/524 | 58/2312 | 0.396551724 | 1.749670966 | 3.129605464 | 0.002436416 | 0.111284179 | 0.111284179 | C5AR1/AGER/IL1RN/HRH | 23 |
| Cluster_4 | MF | GO:001920 | kinase regulator activity | 13/540 | 26/2316 | 0.5 | 2.144444444 | 3.235299905 | 0.002570664 | 0.115251415 | 0.115251415 | CCNB1/CCNF/SOCS6/CD | 13 |
| Cluster_4 | BP | GO:000276 | immune response-regulating signaling pathway | 18/524 | 42/2312 | 0.428571429 | 1.890948746 | 3.153888216 | 0.002615847 | 0.111284179 | 0.111284179 | C5AR1/PTGDR2/RIGI/IFIH | 18 |
| Cluster_4 | BP | GO:004441 | biological process involved in interspecies interaction between organisms | 37/524 | 107/2312 | 0.345794393 | 1.52571877 | 3.013859025 | 0.002697798 | 0.111284179 | 0.111284179 | OAS1Z/STAT2/PGLYRP1/ | 37 |
| Cluster_4 | BP | GO:190304 | mitotic cell cycle process | 24/524 | 62/2312 | 0.387096774 | 1.707953706 | 3.058379009 | 0.002889934 | 0.112887923 | 0.112887923 | CDCA8/NCAPG/CCNB1/N | 24 |
| Cluster_4 | BP | GO:000720 | positive regulation of cytosolic calcium ion concentration | 14/524 | 30/2312 | 0.466666667 | 2.059033079 | 3.160051728 | 0.003010345 | 0.112887923 | 0.112887923 | C5AR1/HRH4/PTGDR2/X | 14 |
| Cluster_4 | BP | GO:005077 | positive regulation of immune response | 25/524 | 66/2312 | 0.378787879 | 1.671293084 | 2.994753791 | 0.003359549 | 0.12050557 | 0.12050557 | C5AR1/PTGDR2/RIGI/IFIH | 25 |
| Cluster_4 | CC | GO:000079 | condensed chromosome | 13/519 | 27/2248 | 0.481481481 | 2.085492043 | 3.108293109 | 0.003569159 | 0.278394391 | 0.278394391 | PSMC3IP/NCAPG/RAD1/ | 13 |
| Cluster_4 | BP | GO:000300 | developmental process involved in reproduction | 9/524 | 16/2312 | 0.5625 | 2.481870229 | 3.219334336 | 0.00359318 | 0.123515575 | 0.123515575 | RSPH1/SPATA6L/CCDC1 | 9 |
| Cluster_4 | BP | GO:000007 | regulation of cyclin-dependent protein serine/threonine kinase activity | 7/524 | 11/2312 | 0.636363636 | 2.80777238 | 3.25284702 | 0.004122423 | 0.125962922 | 0.125962922 | CCNB1/CCNF/CCNJL/CC | 7 |
| Cluster_4 | BP | GO:000728 | spermatid development | 7/524 | 11/2312 | 0.636363636 | 2.80777238 | 3.25284702 | 0.004122423 | 0.125962922 | 0.125962922 | RSPH1/CCDC136/NECTI | 7 |
| Cluster_4 | BP | GO:004851 | spermatid differentiation | 7/524 | 11/2312 | 0.636363636 | 2.80777238 | 3.25284702 | 0.004122423 | 0.125962922 | 0.125962922 | RSPH1/CCDC136/NECTI | 7 |
| Cluster_4 | BP | GO:000695 | immune response | 60/524 | 197/2312 | 0.304568528 | 1.343821444 | 2.730817932 | 0.005022653 | 0.147988894 | 0.147988894 | C5AR1/IL1RN/STAT2/PGL | 60 |
| Cluster_4 | BP | GO:000705 | chromosome segregation | 18/524 | 45/2312 | 0.4 | 1.764885496 | 2.804521551 | 0.006359162 | 0.179759074 | 0.179759074 | CDCA8/PSMC3IP/NCAPG | 18 |
| Cluster_4 | BP | GO:005132 | meiotic cell cycle | 10/524 | 20/2312 | 0.5 | 2.20610687 | 2.932076799 | 0.006536694 | 0.179759074 | 0.179759074 | PSMC3IP/SYCE2/NUF2/M | 10 |
| Cluster_4 | MF | GO:014037 | immune receptor activity | 21/540 | 54/2316 | 0.388888889 | 1.667901235 | 2.737875672 | 0.006946608 | 0.266948227 | 0.266948227 | C5AR1/IL18R1/PTGDR2/I | 21 |
| Cluster_4 | BP | GO:000028 | nuclear division | 16/524 | 39/2312 | 0.41025641 | 1.81013897 | 2.761683902 | 0.007457748 | 0.19847233 | 0.19847233 | CDCA8/PSMC3IP/NCAPG | 16 |
| Cluster_4 | BP | GO:000728 | germ cell development | 7/524 | 12/2312 | 0.583333333 | 2.573791349 | 2.958391988 | 0.007981243 | 0.205766427 | 0.205766427 | RSPH1/CCDC136/NECTI | 7 |
| Cluster_4 | BP | GO:000242 | immune response-activating cell surface receptor signaling pathway | 14/524 | 33/2312 | 0.424242424 | 1.871848254 | 2.730282392 | 0.008573693 | 0.21434232 | 0.21434232 | C5AR1/PTGDR2/TMIGD2/ | 14 |
| Cluster_4 | CC | GO:000030 | cyclin-dependent protein kinase holoenzyme complex | 7/519 | 12/2248 | 0.583333333 | 2.526653821 | 2.904577783 | 0.00888181 | 0.404108808 | 0.404108808 | CCNB1/CCNF/CCNJL/CC | 7 |
| Cluster_4 | BP | GO:003355 | cellular response to stress | 26/524 | 74/2312 | 0.351351351 | 1.55023726 | 2.603860262 | 0.008937092 | 0.21685591 | 0.21685591 | RAD1/PPP1R15A/HSPB8/ | 26 |
| Cluster_4 | BP | GO:000728 | spermatogenesis | 8/524 | 15/2312 | 0.533333333 | 2.353180662 | 2.845789506 | 0.009226921 | 0.217491698 | 0.217491698 | RSPH1/SPATA6L/CCDC1 | 8 |
| Cluster_4 | BP | GO:000625 | DNA metabolic process | 19/524 | 50/2312 | 0.38 | 1.676641221 | 2.618056487 | 0.009710004 | 0.220668794 | 0.220668794 | PSMC3IP/RAD1/NUDT1/M | 19 |
| Cluster_4 | BP | GO:000237 | immune system process | 70/524 | 242/2312 | 0.289256198 | 1.276260173 | 2.458230759 | 0.009896661 | 0.220668794 | 0.220668794 | C5AR1/JAML/IL1RN/ORM | 70 |
| Cluster_4 | CC | GO:190255 | serine/threonine protein kinase complex | 8/519 | 15/2248 | 0.533333333 | 2.310083494 | 2.788611162 | 0.010361764 | 0.404108808 | 0.404108808 | CCNB1/CCNF/CDK5R1/C | 8 |
| Cluster_4 | MF | GO:000488 | transmembrane signaling receptor activity | 55/540 | 179/2316 | 0.30726257 | 1.317815022 | 2.440335692 | 0.010853639 | 0.364953626 | 0.364953626 | GPR141/C5AR1/PDGFRA | 55 |
| Cluster_4 | BP | GO:000701 | microtubule-based movement | 14/524 | 34/2312 | 0.411764706 | 1.816793893 | 2.596910245 | 0.011622849 | 0.233703722 | 0.233703722 | TRAF3IP1/TMEM201/IFT2 | 14 |
| Cluster_4 | BP | GO:005079 | regulation of viral process | 6/524 | 10/2312 | 0.6 | 2.647328244 | 2.825591296 | 0.011913881 | 0.233703722 | 0.233703722 | OAS1Z/OASL/ISG20/NEC | 6 |

| | | | | | | | | | | | | | |
|---|---|---|---|---|---|---|---|---|---|---|---|---|---|
| Cluster_4 | BP | GO:190390 | regulation of viral life cycle | 6/524 | 10/2312 | 0.6 | 2.647328244 | 2.825591296 | 0.011913881 | 0.233703722 | 0.233703722 | OAS1Z/OASL/ISG20/NEC | 6 |
| Cluster_4 | BP | GO:004828 | organelle fission | 16/524 | 41/2312 | 0.390243902 | 1.721839508 | 2.524096002 | 0.012918411 | 0.233703722 | 0.233703722 | CDCA8/PSMC3IP/NCAPG | 16 |
| Cluster_4 | BP | GO:004477 | mitotic cell cycle phase transition | 12/524 | 28/2312 | 0.428571429 | 1.890948746 | 2.567234531 | 0.01334445 | 0.233703722 | 0.233703722 | CCNB1/CDK1/CCNF/SPD | 12 |
| Cluster_4 | BP | GO:005172 | regulation of cell cycle | 22/524 | 62/2312 | 0.35483871 | 1.56562423 | 2.443511855 | 0.013791449 | 0.233703722 | 0.233703722 | RAD1/CDK1/GADD45A/FE | 22 |
| Cluster_4 | BP | GO:009854 | defense response to other organism | 20/524 | 55/2312 | 0.363636364 | 1.60444136 | 2.455565926 | 0.013859534 | 0.233703722 | 0.233703722 | STAT2/PGLYRP1/RIGI/IFI | 20 |
| Cluster_4 | BP | GO:014054 | defense response to symbiont | 20/524 | 55/2312 | 0.363636364 | 1.60444136 | 2.455565926 | 0.013859534 | 0.233703722 | 0.233703722 | STAT2/PGLYRP1/RIGI/IFI | 20 |
| Cluster_4 | BP | GO:000626 | DNA replication | 7/524 | 13/2312 | 0.538461538 | 2.375807399 | 2.69241307 | 0.013964093 | 0.233703722 | 0.233703722 | POLE2/MCM2/MCM10/GIN | 7 |
| Cluster_4 | BP | GO:000697 | response to oxidative stress | 7/524 | 13/2312 | 0.538461538 | 2.375807399 | 2.69241307 | 0.013964093 | 0.233703722 | 0.233703722 | NFE2L2/GPX6/PTGS2/HM | 7 |
| Cluster_4 | BP | GO:014001 | mitotic nuclear division | 11/524 | 25/2312 | 0.44 | 1.941374046 | 2.56142121 | 0.014104264 | 0.233703722 | 0.233703722 | CDCA8/NCAPG/CCNB1/N | 11 |
| Cluster_4 | BP | GO:000727 | gamete generation | 8/524 | 16/2312 | 0.5 | 2.20610687 | 2.620243786 | 0.014860661 | 0.233703722 | 0.233703722 | RSPH1/SPATA6L/CCDC1 | 8 |
| Cluster_4 | BP | GO:004823 | male gamete generation | 8/524 | 16/2312 | 0.5 | 2.20610687 | 2.620243786 | 0.014860661 | 0.233703722 | 0.233703722 | RSPH1/SPATA6L/CCDC1 | 8 |
| Cluster_4 | BP | GO:007190 | regulation of protein serine/threonine kinase activity | 8/524 | 16/2312 | 0.5 | 2.20610687 | 2.620243786 | 0.014860661 | 0.233703722 | 0.233703722 | TRIB1/CCNB1/CCNF/CCN | 8 |
| Cluster_4 | BP | GO:000007 | mitotic sister chromatid segregation | 9/524 | 19/2312 | 0.473684211 | 2.089995982 | 2.582150368 | 0.015013694 | 0.233703722 | 0.233703722 | CDCA8/NCAPG/CCNB1/N | 9 |
| Cluster_4 | BP | GO:190304 | meiotic cell cycle process | 9/524 | 19/2312 | 0.473684211 | 2.089995982 | 2.582150368 | 0.015013694 | 0.233703722 | 0.233703722 | PSMC3IP/SYCE2/NUF2/M | 9 |
| Cluster_4 | BP | GO:000276 | immune response-regulating cell surface receptor signaling pathway | 14/524 | 35/2312 | 0.4 | 1.764885496 | 2.467918376 | 0.015454341 | 0.23610799 | 0.23610799 | C5AR1/PTGDR2/TMIGD2/ | 14 |
| Cluster_4 | BP | GO:000268 | positive regulation of immune system process | 30/524 | 92/2312 | 0.326086957 | 1.43876535 | 2.324511017 | 0.016520835 | 0.243381348 | 0.243381348 | C5AR1/CSF1/PTGDR2/RI | 30 |
| Cluster_4 | BP | GO:004477 | cell cycle phase transition | 13/524 | 32/2312 | 0.40625 | 1.792461832 | 2.443249276 | 0.016815439 | 0.243381348 | 0.243381348 | RAD1/CCNB1/CDK1/CCN | 13 |
| Cluster_4 | BP | GO:009881 | nuclear chromosome segregation | 13/524 | 32/2312 | 0.40625 | 1.792461832 | 2.443249276 | 0.016815439 | 0.243381348 | 0.243381348 | CDCA8/PSMC3IP/NCAPG | 13 |
| Cluster_4 | BP | GO:004508 | innate immune response | 18/524 | 49/2312 | 0.367346939 | 1.620813211 | 2.377379821 | 0.01711305 | 0.243418377 | 0.243418377 | STAT2/PGLYRP1/RIGI/IFI | 18 |
| Cluster_4 | MF | GO:001995 | cytokine binding | 15/540 | 38/2316 | 0.394736842 | 1.692982456 | 2.374586942 | 0.018316044 | 0.547446197 | 0.547446197 | IL1RN/IL18R1/IL23R/XCR | 15 |
| Cluster_4 | BP | GO:003134 | positive regulation of defense response | 10/524 | 23/2312 | 0.434782609 | 1.9183538 | 2.395702622 | 0.020870291 | 0.291830334 | 0.291830334 | RIGI/IFIH1/TREM2/MYD8 | 10 |
| Cluster_4 | BP | GO:000222 | pattern recognition receptor signaling pathway | 6/524 | 11/2312 | 0.545454545 | 2.40666204 | 2.531102077 | 0.021263631 | 0.292374922 | 0.292374922 | RIGI/IFIH1/MYD88/TLR8/S | 6 |
| Cluster_4 | BP | GO:001605 | organic acid catabolic process | 8/524 | 17/2312 | 0.470588235 | 2.076335878 | 2.410808931 | 0.02262532 | 0.301062727 | 0.301062727 | PRODH/HAL/ALDH5A1/ID | 8 |
| Cluster_4 | BP | GO:004639 | carboxylic acid catabolic process | 8/524 | 17/2312 | 0.470588235 | 2.076335878 | 2.410808931 | 0.02262532 | 0.301062727 | 0.301062727 | PRODH/HAL/ALDH5A1/ID | 8 |
| Cluster_4 | CC | GO:003014 | secretory granule | 6/519 | 11/2248 | 0.545454545 | 2.362585391 | 2.48150296 | 0.023245455 | 0.654900462 | 0.654900462 | SRGN/CCDC136/ENKUR/ | 6 |
| Cluster_4 | CC | GO:190291 | protein kinase complex | 8/519 | 17/2248 | 0.470588235 | 2.038308965 | 2.353905361 | 0.025188479 | 0.654900462 | 0.654900462 | CCNB1/CCNF/CDK5R1/C | 8 |
| Cluster_4 | BP | GO:000961 | response to bacterium | 18/524 | 51/2312 | 0.352941176 | 1.557251908 | 2.178051931 | 0.026175287 | 0.342771613 | 0.342771613 | PGLYRP1/IL1A/TNIP3/GN | 18 |
| Cluster_4 | MF | GO:000489 | cytokine receptor activity | 17/540 | 47/2316 | 0.361702128 | 1.551300236 | 2.105090429 | 0.03083268 | 0.829399079 | 0.829399079 | IL18R1/IL23R/XCR1/CXCF | 17 |
| Cluster_4 | BP | GO:000081 | sister chromatid segregation | 9/524 | 21/2312 | 0.428571429 | 1.890948746 | 2.219891164 | 0.030911599 | 0.386394989 | 0.386394989 | CDCA8/NCAPG/CCNB1/N | 9 |
| Cluster_4 | BP | GO:004428 | small molecule catabolic process | 9/524 | 21/2312 | 0.428571429 | 1.890948746 | 2.219891164 | 0.030911599 | 0.386394989 | 0.386394989 | PRODH/ADA/HAL/ALDH5 | 9 |
| Cluster_4 | BP | GO:004508 | positive regulation of innate immune response | 9/524 | 21/2312 | 0.428571429 | 1.890948746 | 2.219891164 | 0.030911599 | 0.386394989 | 0.386394989 | RIGI/IFIH1/MYD88/TLR8/K | 9 |
| Cluster_4 | BP | GO:005127 | chromosome organization | 16/524 | 45/2312 | 0.355555556 | 1.568787108 | 2.085509151 | 0.032699029 | 0.390417328 | 0.390417328 | CDCA8/PSMC3IP/NCAPG | 16 |
| Cluster_4 | BP | GO:005077 | regulation of immune response | 25/524 | 78/2312 | 0.320512821 | 1.414171071 | 2.014036499 | 0.033717905 | 0.390417328 | 0.390417328 | C5AR1/PTGDR2/RIGI/IFIH | 25 |
| Cluster_4 | BP | GO:000221 | activation of innate immune response | 7/524 | 15/2312 | 0.466666667 | 2.059033079 | 2.227186146 | 0.03425729 | 0.390417328 | 0.390417328 | RIGI/IFIH1/MYD88/TLR8/K | 7 |
| Cluster_4 | BP | GO:000275 | innate immune response-activating signaling pathway | 7/524 | 15/2312 | 0.466666667 | 2.059033079 | 2.227186146 | 0.03425729 | 0.390417328 | 0.390417328 | RIGI/IFIH1/MYD88/TLR8/K | 7 |
| Cluster_4 | BP | GO:007121 | cellular response to biotic stimulus | 7/524 | 15/2312 | 0.466666667 | 2.059033079 | 2.227186146 | 0.03425729 | 0.390417328 | 0.390417328 | IL1A/TNIP3/CLEC7A/DDIT | 7 |
| Cluster_4 | BP | GO:001603 | viral process | 6/524 | 12/2312 | 0.5 | 2.20610687 | 2.267223609 | 0.034546018 | 0.390417328 | 0.390417328 | OAS1Z/OASL/ISG20/NEC | 6 |
| Cluster_4 | BP | GO:001905 | viral life cycle | 6/524 | 12/2312 | 0.5 | 2.20610687 | 2.267223609 | 0.034546018 | 0.390417328 | 0.390417328 | OAS1Z/OASL/ISG20/NEC | 6 |
| Cluster_4 | BP | GO:000701 | microtubule-based process | 26/524 | 82/2312 | 0.317073171 | 1.398994601 | 1.991147223 | 0.035027673 | 0.390511217 | 0.390511217 | TRAF3IP1/TMEM201/SKA | 26 |
| Cluster_4 | BP | GO:003134 | regulation of defense response | 15/524 | 42/2312 | 0.357142857 | 1.575790622 | 2.038253339 | 0.036564935 | 0.40221428 | 0.40221428 | AGER/RIGI/IFIH1/NR1H3/ | 15 |
| Cluster_4 | CC | GO:000587 | microtubule associated complex | 7/519 | 15/2248 | 0.466666667 | 2.021323057 | 2.173962874 | 0.037637378 | 0.744770532 | 0.744770532 | CDCA8/KIF23/AURKB/DY | 7 |
| Cluster_4 | BP | GO:000695 | humoral immune response | 10/524 | 25/2312 | 0.4 | 1.764885496 | 2.08120666 | 0.038480647 | 0.417717548 | 0.417717548 | GNLY/CATHL2/CATHL5/C | 10 |
| Cluster_4 | CC | GO:003299 | protein-containing complex | 107/519 | 403/2248 | 0.265508685 | 1.150026057 | 1.820994992 | 0.040770579 | 0.744770532 | 0.744770532 | TRAF3IP1/CDCA8/PSMC | 107 |
| Cluster_4 | BP | GO:000283 | positive regulation of response to biotic stimulus | 9/524 | 22/2312 | 0.409090909 | 1.80499653 | 2.053380657 | 0.042134558 | 0.445653984 | 0.445653984 | RIGI/IFIH1/MYD88/TLR8/K | 9 |
| Cluster_4 | BP | GO:004508 | regulation of innate immune response | 9/524 | 22/2312 | 0.409090909 | 1.80499653 | 2.053380657 | 0.042134558 | 0.445653984 | 0.445653984 | RIGI/IFIH1/MYD88/TLR8/K | 9 |
| Cluster_4 | BP | GO:000693 | chemotaxis | 20/524 | 61/2312 | 0.327868852 | 1.446627456 | 1.91339417 | 0.043270967 | 0.446231845 | 0.446231845 | C5AR1/JAML/SEMA4A/XC | 20 |
| Cluster_4 | BP | GO:004233 | taxis | 20/524 | 61/2312 | 0.327868852 | 1.446627456 | 1.91339417 | 0.043270967 | 0.446231845 | 0.446231845 | C5AR1/JAML/SEMA4A/XC | 20 |
| Cluster_4 | BP | GO:004222 | response to chemical | 52/524 | 186/2312 | 0.279569892 | 1.233522121 | 1.797565514 | 0.046178624 | 0.470337833 | 0.470337833 | C5AR1/JAML/SEMA4A/IL | 52 |
| Cluster_4 | BP | GO:001975 | carboxylic acid metabolic process | 23/524 | 73/2312 | 0.315068493 | 1.390149535 | 1.833358243 | 0.048990073 | 0.489355665 | 0.489355665 | PRODH/CPT1C/EPHX1/L | 23 |
| Cluster_5 | CC | GO:009859 | plasma membrane region | 15/356 | 39/2248 | 0.384615385 | 2.428694901 | 3.903371702 | 0.000480905 | 0.070212146 | 0.070212146 | CD36/AMN/SHANK3/SHR | 15 |
| Cluster_5 | MF | GO:005101 | actin filament binding | 14/376 | 37/2316 | 0.378378378 | 2.330649799 | 3.59137725 | 0.001165846 | 0.310114958 | 0.310114958 | TPM2/MYO9A/SHROOM4 | 14 |
| Cluster_5 | BP | GO:001571 | organic anion transport | 13/366 | 34/2312 | 0.382352941 | 2.415300546 | 3.604793327 | 0.00122429 | 0.551022384 | 0.551022384 | CD36/FABP4/SLC1A4/RB | 13 |
| Cluster_5 | BP | GO:000689 | receptor-mediated endocytosis | 7/366 | 14/2312 | 0.5 | 3.158469945 | 3.512401137 | 0.00294257 | 0.551022384 | 0.551022384 | CD36/LDLRAD3/AMN/LDL | 7 |
| Cluster_5 | BP | GO:004694 | carboxylic acid transport | 10/366 | 25/2312 | 0.4 | 2.526775956 | 3.327984781 | 0.003057283 | 0.551022384 | 0.551022384 | CD36/FABP4/SLC1A4/RB | 10 |
| Cluster_5 | BP | GO:005117 | localization | 85/366 | 417/2312 | 0.20383693 | 1.287625637 | 2.812937687 | 0.003700379 | 0.551022384 | 0.551022384 | CD36/SLC12A7/LDLRAD3 | 85 |
| Cluster_5 | BP | GO:000681 | transport | 77/366 | 372/2312 | 0.206989247 | 1.307538633 | 2.807626821 | 0.003864962 | 0.551022384 | 0.551022384 | CD36/SLC12A7/LDLRAD3 | 77 |
| Cluster_5 | BP | GO:001584 | organic acid transport | 10/366 | 26/2312 | 0.384615385 | 2.429592266 | 3.178555861 | 0.004282557 | 0.551022384 | 0.551022384 | CD36/FABP4/SLC1A4/RB | 10 |
| Cluster_5 | MF | GO:000377 | actin binding | 17/376 | 55/2316 | 0.309090909 | 1.903868472 | 2.986103501 | 0.004560584 | 0.606557727 | 0.606557727 | TPM2/MYO9A/SHROOM4 | 17 |
| Cluster_5 | CC | GO:004300 | neuron projection | 19/356 | 67/2248 | 0.28358209 | 1.790709374 | 2.849655926 | 0.005968345 | 0.248787209 | 0.248787209 | SHANK3/MYO9A/RAC3/C | 19 |

| | | | | | | | | | | | | | |
|---|---|---|---|---|---|---|---|---|---|---|---|---|---|
| Cluster_5 | BP | GO:005123 | establishment of localization | 78/366 | 384/2312 | 0.203125 | 1.283128415 | 2.634296489 | 0.006177693 | 0.658212775 | 0.658212775 | CD36/SLC12A7/LDLRAD3 | 78 |
| Cluster_5 | CC | GO:004299 | cell projection | 28/356 | 112/2248 | 0.25 | 1.578651685 | 2.724529902 | 0.006769732 | 0.248787209 | 0.248787209 | DZIP1/MORN4/SHANK3/M | 28 |
| Cluster_5 | CC | GO:012002 | plasma membrane bounded cell projection | 27/356 | 107/2248 | 0.252336449 | 1.593405439 | 2.727727909 | 0.006816088 | 0.248787209 | 0.248787209 | DZIP1/SHANK3/MYO9A/T | 27 |
| Cluster_5 | BP | GO:005508 | transmembrane transport | 35/366 | 149/2312 | 0.234899329 | 1.483844941 | 2.647525649 | 0.007541145 | 0.658212775 | 0.658212775 | SLC12A7/SLC9A9/SLC1A | 35 |
| Cluster_5 | BP | GO:009743 | supramolecular fiber organization | 17/366 | 59/2312 | 0.288135593 | 1.820135223 | 2.766941464 | 0.007673465 | 0.658212775 | 0.658212775 | CCDC57/TPM2/SHROOM | 17 |
| Cluster_5 | BP | GO:009865 | import into cell | 18/366 | 66/2312 | 0.272727273 | 1.722801788 | 2.583184391 | 0.011325479 | 0.874326989 | 0.874326989 | CD36/SLC12A7/LDLRAD3 | 18 |
| Cluster_5 | MF | GO:001529 | secondary active transmembrane transporter activity | 11/376 | 34/2316 | 0.323529412 | 1.992803504 | 2.566926864 | 0.014794451 | 0.85173711 | 0.85173711 | SLC12A7/SLC9A9/SLC1A | 11 |
| Cluster_5 | BP | GO:003476 | regulation of transmembrane transport | 8/366 | 22/2312 | 0.363636364 | 2.297069051 | 2.650487703 | 0.015156432 | 0.975927853 | 0.975927853 | RANGRF/SCN1B/SHISA8 | 8 |
| Cluster_5 | BP | GO:000689 | endocytosis | 13/366 | 44/2312 | 0.295454545 | 1.866368604 | 2.515802917 | 0.015169863 | 0.975927853 | 0.975927853 | CD36/LDLRAD3/AMN/LDL | 13 |
| Cluster_5 | MF | GO:004487 | protein-containing complex binding | 22/376 | 87/2316 | 0.252873563 | 1.557593544 | 2.333409081 | 0.018152368 | 0.85173711 | 0.85173711 | CD36/TPM2/GNAQ/ADAM | 22 |
| Cluster_5 | CC | GO:003042 | axon | 9/356 | 27/2248 | 0.333333333 | 2.104868914 | 2.504865012 | 0.018712943 | 0.478602139 | 0.478602139 | MYO9A/NRP1/FSCN1/NA | 9 |
| Cluster_5 | BP | GO:003303 | macromolecule localization | 31/366 | 137/2312 | 0.226277372 | 1.429380559 | 2.2466847 | 0.019899995 | 0.993661976 | 0.993661976 | CD36/FABP4/AMN/TFRC/ | 31 |
| Cluster_5 | BP | GO:001571 | monocarboxylic acid transport | 6/366 | 15/2312 | 0.4 | 2.526775956 | 2.572228474 | 0.021243822 | 0.993661976 | 0.993661976 | CD36/FABP4/RBP1/FABP | 6 |
| Cluster_5 | BP | GO:004239 | regulation of membrane potential | 6/366 | 15/2312 | 0.4 | 2.526775956 | 2.572228474 | 0.021243822 | 0.993661976 | 0.993661976 | GNAQ/RANGRF/KCNK13/ | 6 |
| Cluster_5 | CC | GO:000013 | Golgi membrane | 6/356 | 15/2248 | 0.4 | 2.525842697 | 2.571443583 | 0.021269668 | 0.478602139 | 0.478602139 | TVP23A/ACER2/IER3IP1/ | 6 |
| Cluster_5 | MF | GO:002285 | transmembrane transporter activity | 32/376 | 140/2316 | 0.228571429 | 1.407902736 | 2.191592196 | 0.022244599 | 0.85173711 | 0.85173711 | SLC12A7/SLC9A9/SLC1A | 32 |
| Cluster_5 | CC | GO:001562 | actin cytoskeleton | 14/356 | 51/2248 | 0.274509804 | 1.733421458 | 2.297680514 | 0.022946678 | 0.478602139 | 0.478602139 | TPM2/MYO9A/SHROOM4 | 14 |
| Cluster_5 | MF | GO:000823 | peptidase activity | 24/376 | 99/2316 | 0.242424242 | 1.493230174 | 2.207765504 | 0.023131935 | 0.85173711 | 0.85173711 | DPEP2/HP/ADAM15/CTS | 24 |
| Cluster_5 | MF | GO:002280 | active transmembrane transporter activity | 14/376 | 50/2316 | 0.28 | 1.724680851 | 2.280186606 | 0.023727335 | 0.85173711 | 0.85173711 | SLC12A7/SLC9A9/SLC1A | 14 |
| Cluster_5 | BP | GO:004326 | regulation of monoatomic ion transport | 8/366 | 24/2312 | 0.333333333 | 2.10564663 | 2.360820974 | 0.026140392 | 0.993661976 | 0.993661976 | RANGRF/SCN1B/HFE/KC | 8 |
| Cluster_5 | BP | GO:001095 | regulation of metal ion transport | 7/366 | 20/2312 | 0.35 | 2.210928962 | 2.358280473 | 0.028212446 | 0.993661976 | 0.993661976 | RANGRF/SCN1B/HFE/KC | 7 |
| Cluster_5 | BP | GO:003476 | regulation of monoatomic ion transmembrane transport | 7/366 | 20/2312 | 0.35 | 2.210928962 | 2.358280473 | 0.028212446 | 0.993661976 | 0.993661976 | RANGRF/SCN1B/KCNIP3 | 7 |
| Cluster_5 | BP | GO:001590 | fatty acid transport | 5/366 | 12/2312 | 0.416666667 | 2.632058288 | 2.457714704 | 0.029312819 | 0.993661976 | 0.993661976 | CD36/FABP4/RBP1/FABP | 5 |
| Cluster_5 | BP | GO:003019 | extracellular matrix organization | 5/366 | 12/2312 | 0.416666667 | 2.632058288 | 2.457714704 | 0.029312819 | 0.993661976 | 0.993661976 | MIA/MMP28/MMP19/ADA | 5 |
| Cluster_5 | BP | GO:004306 | extracellular structure organization | 5/366 | 12/2312 | 0.416666667 | 2.632058288 | 2.457714704 | 0.029312819 | 0.993661976 | 0.993661976 | MIA/MMP28/MMP19/ADA | 5 |
| Cluster_5 | BP | GO:004522 | external encapsulating structure organization | 5/366 | 12/2312 | 0.416666667 | 2.632058288 | 2.457714704 | 0.029312819 | 0.993661976 | 0.993661976 | MIA/MMP28/MMP19/ADA | 5 |
| Cluster_5 | BP | GO:009953 | synaptic signaling | 9/366 | 29/2312 | 0.310344828 | 1.960429621 | 2.256735635 | 0.029866588 | 0.993661976 | 0.993661976 | SLC12A7/LY6G6E/CHRM | 9 |
| Cluster_5 | MF | GO:000521 | transporter activity | 34/376 | 154/2316 | 0.220779221 | 1.359906051 | 2.034652375 | 0.030626404 | 0.85173711 | 0.85173711 | SLC12A7/SLC9A9/SLC1A | 34 |
| Cluster_5 | MF | GO:000823 | metallopeptidase activity | 10/376 | 33/2316 | 0.303030303 | 1.866537718 | 2.206785086 | 0.031226405 | 0.85173711 | 0.85173711 | DPEP2/ADAM15/TRABD2 | 10 |
| Cluster_5 | MF | GO:004694 | carboxylic acid transmembrane transporter activity | 7/376 | 20/2316 | 0.35 | 2.155851064 | 2.285069893 | 0.032020192 | 0.85173711 | 0.85173711 | SLC1A4/SLC19A1/SLC7A | 7 |
| Cluster_5 | BP | GO:000701 | actin filament organization | 12/366 | 44/2312 | 0.272727273 | 1.722801788 | 2.098906677 | 0.035650827 | 0.993661976 | 0.993661976 | TPM2/SHROOM4/ARHGE | 12 |
| Cluster_5 | BP | GO:007084 | response to growth factor | 9/366 | 30/2312 | 0.3 | 1.895081967 | 2.139610461 | 0.036919148 | 0.993661976 | 0.993661976 | TSPAN16/SKI/TGFB2/FS | 9 |
| Cluster_5 | BP | GO:007136 | cellular response to growth factor stimulus | 9/366 | 30/2312 | 0.3 | 1.895081967 | 2.139610461 | 0.036919148 | 0.993661976 | 0.993661976 | TSPAN16/SKI/TGFB2/FS | 9 |
| Cluster_5 | BP | GO:000301 | muscle system process | 6/366 | 17/2312 | 0.352941176 | 2.229508197 | 2.206142909 | 0.039793131 | 0.993661976 | 0.993661976 | TPM2/P2RY1/CHRM3/SC | 6 |
| Cluster_5 | BP | GO:190406 | regulation of monoatomic cation transmembrane transport | 6/366 | 17/2312 | 0.352941176 | 2.229508197 | 2.206142909 | 0.039793131 | 0.993661976 | 0.993661976 | RANGRF/SCN1B/KCNIP3 | 6 |
| Cluster_5 | BP | GO:009028 | regulation of cellular response to growth factor stimulus | 5/366 | 13/2312 | 0.384615385 | 2.429592266 | 2.24121478 | 0.041610353 | 0.993661976 | 0.993661976 | TSPAN16/SKI/FSTL1/NR | 5 |
| Cluster_5 | CC | GO:009706 | synaptic membrane | 5/356 | 13/2248 | 0.384615385 | 2.428694901 | 2.240466095 | 0.041656441 | 0.682687278 | 0.682687278 | SHANK3/SHISA8/NRP2/S | 5 |
| Cluster_5 | BP | GO:001619 | vesicle-mediated transport | 23/366 | 102/2312 | 0.225490196 | 1.424408015 | 1.900888412 | 0.043378621 | 0.993661976 | 0.993661976 | CD36/LDLRAD3/AMN/TV | 23 |
| Cluster_5 | MF | GO:000422 | metalloendopeptidase activity | 6/376 | 17/2316 | 0.352941176 | 2.173967459 | 2.138353931 | 0.044420127 | 0.870999319 | 0.870999319 | ADAM15/TRABD2B/MMP | 6 |
| Cluster_5 | MF | GO:000808 | growth factor activity | 6/376 | 17/2316 | 0.352941176 | 2.173967459 | 2.138353931 | 0.044420127 | 0.870999319 | 0.870999319 | TGFB2/AREG/GDF11/IL2/ | 6 |
| Cluster_5 | BP | GO:005166 | localization within membrane | 9/366 | 31/2312 | 0.290322581 | 1.833950291 | 2.026877335 | 0.045031754 | 0.993661976 | 0.993661976 | RAB11FIP4/RTP4/LIN7B/ | 9 |
| Cluster_5 | MF | GO:000850 | monoatomic anion transmembrane transporter activity | 5/376 | 13/2316 | 0.384615385 | 2.369067103 | 2.178803165 | 0.045842069 | 0.870999319 | 0.870999319 | SLC12A7/SLC19A1/TTYH | 5 |
| Cluster_5 | MF | GO:003802 | cargo receptor activity | 5/376 | 13/2316 | 0.384615385 | 2.369067103 | 2.178803165 | 0.045842069 | 0.870999319 | 0.870999319 | CD36/TFRC/LDLR/OLR1/ | 5 |
| Cluster_5 | BP | GO:003002 | actin filament-based process | 17/366 | 71/2312 | 0.23943662 | 1.512506734 | 1.901852886 | 0.046406998 | 0.993661976 | 0.993661976 | TPM2/SHROOM4/ARHGE | 17 |
| Cluster_5 | BP | GO:000681 | monoatomic cation transport | 21/366 | 92/2312 | 0.22826087 | 1.441910192 | 1.875515002 | 0.04641843 | 0.993661976 | 0.993661976 | SLC12A7/SLC9A9/TFRC/ | 21 |
| Cluster_5 | BP | GO:000810 | intracellular protein localization | 21/366 | 92/2312 | 0.22826087 | 1.441910192 | 1.875515002 | 0.04641843 | 0.993661976 | 0.993661976 | CD36/AMN/TFRC/TVP23 | 21 |
| Cluster_5 | BP | GO:000657 | modified amino acid metabolic process | 7/366 | 22/2312 | 0.318181818 | 2.00993542 | 2.063746266 | 0.046821644 | 0.993661976 | 0.993661976 | GGT1/GSTA4/GSTT1/GG | 7 |
| Cluster_5 | BP | GO:000300 | system process | 13/366 | 51/2312 | 0.254901961 | 1.610200364 | 1.910629007 | 0.049029978 | 0.993661976 | 0.993661976 | IMPG2/TPM2/P2RY1/CRY | 13 |
| Cluster_5 | BP | GO:000681 | monoatomic ion transport | 22/366 | 98/2312 | 0.224489796 | 1.418088547 | 1.83384066 | 0.049700787 | 0.993661976 | 0.993661976 | SLC12A7/SLC9A9/TFRC/ | 22 |
